# ScanNet: An interpretable geometric deep learning model for structure-based protein binding site prediction

**DOI:** 10.1101/2021.09.05.459013

**Authors:** Jérôme Tubiana, Dina Schneidman-Duhovny, Haim J. Wolfson

## Abstract

Predicting the functional sites of a protein from its structure, such as the binding sites of small molecules, other proteins or antibodies sheds light on its function *in vivo*. Currently, two classes of methods prevail: Machine Learning (ML) models built on top of handcrafted features and comparative modeling. They are respectively limited by the expressivity of the handcrafted features and the availability of similar proteins. Here, we introduce ScanNet, an end-to-end, interpretable geometric deep learning model that learns features directly from 3D structures. ScanNet builds representations of atoms and amino acids based on the spatio-chemical arrangement of their neighbors. We train ScanNet for detecting protein-protein and protein-antibody binding sites, demonstrate its accuracy - including for unseen protein folds - and interpret the filters learned. Finally, we predict epitopes of the SARS-CoV-2 spike protein, validating known antigenic regions and predicting previously uncharacterized ones. Overall, ScanNet is a versatile, powerful, and interpretable model suitable for functional site prediction tasks. A webserver for ScanNet is available from http://bioinfo3d.cs.tau.ac.il/ScanNet/

## INTRODUCTION

Despite recent progresses in experimental [1] and AI-based [2, 3] protein structure determination, there remains a gap between structure and function [4]. The most accurate functional site prediction method is comparative modelling [5–13]: given a query protein, similar proteins with known functional sites are searched for and their sites are mapped onto the query structure. Comparative modelling has several shortcomings. First and foremost, its coverage is limited, as the pool of experimentally characterized protein folds or structural motifs is small. Second, functional sites are variably preserved throughout evolution. On the one hand, the B-cell epitopes of viral proteins frequently undergo antigenic drift, *i.e*. the abolition of recognition by antibodies after only one or few mutations. On the other hand, some protein-protein interactions are mainly driven by few “hotspot” residues; mutations and/or conformational changes of the other interface residues preserve the interaction. Put differently, the *invariances* in both sequence and conformation spaces of such function-determining structural motifs are in general motif-dependent and therefore unknown. This hampers our ability to both define and recognize such motifs using conventional comparative approaches.

An alternative to comparative modelling is feature-based Machine Learning (ML) [12–18]. For each amino acid of a query protein, various features of geometrical (e.g. secondary structure, solvent accessibility, molecular surface curvature), physico-chemical (e.g. hydrophobicity, polarity, electrostatic potential) and evolutionary (e.g. conservation, position-weight matrices, coevolution) nature are calculated. Then, the target property is predicted using a ML model for tabular data such as Random Forest or Gradient Boosting. Reasoning on mathematically defined features offers three advantages: i) ability to generalize to proteins with no similarity to any of the train set proteins, ii) high sequence sensitivity, *i.e*. ability to output distinct predictions for highly similar protein sequences and iii) fast inference speed. ML models are however limited by the expressiveness of the features employed, as these cannot capture the spatio-chemical arrangements of atoms or amino acids characterizing function-bearing motifs. Examples of such function-bearing motifs include Zinc fingers that are signature of DNA/RNA binding sites [19], or protein-protein interaction hotspots ”O-rings” [20], namely exposed hydrophobic/aromatic amino acids surrounded by polar/charged ones. Despite over 50 years of experimental structural determination, novel function-determining motifs are still being discovered [21].

End-to-end differentiable models, i.e. Deep Learning (DL) can potentially overcome the limitations of both approaches. Indeed, DL models can learn the data features and their invariances directly by backpropagation, and generalize well despite large number of parameters. Adapting the DL approach to protein structures requires defining an appropriate representation for proteins. Proteins can indeed be represented in multiple, complementary ways, e.g. as sequences [22, 23], residue graphs [24–27], atomic density maps [28–34], atomic point clouds [35] or molecular surfaces [36, 37], each capturing different functionally relevant features. Voxelated atomic density maps can be readily processed using classical 3D Convolutional Neural Networks, but the approach is computationally intensive and the predictions are not invariant upon rotation of the input structure. Point clouds, graphs and surfaces can be analyzed via Geometric Deep Learning [38, 39], i.e. end-to-end differentiable models tailored for data with no natural grid-like topology or shared global coordinate system. Graphs can be derived from 3D structures by taking residues as nodes and the distances and angles between them as edges and processed using Graph Neural Networks (GNN) such as Message Passing Neural Networks [40] or Graph Attention Networks [41]. By design, GNNs are invariant upon euclidean transformation and expressive, but can be challenging to regularize and interpret. In particular, it is unclear whether - and if yes, which - structural motifs are captured by GNNs.

Here, we introduce ScanNet (Spatio-Chemical Arrangement of Neighbors Neural Network), a novel geometric deep learning architecture tailored for protein structures. ScanNet builds representations of atoms and amino acids based on the spatio-chemical arrangement of their neighbors and exploits them to predict labels for each amino acid. By construction, ScanNet is end-to-end differentiable with minimal structure preprocessing, yielding fast training and inference. ScanNet predictions are local, invariant upon euclidean transformations and integrate information from multiple scales (atom, amino acid) and modalities (structure, MSA) in a synergistic fashion. Its corresponding parametric function is expressive, meaning that it can efficiently approximate known handcrafted features. Crucially, through appropriate parameterization and regularization, the filters learnt by ScanNet can be readily visualized and interpreted. We showcase the capabilities of ScanNet on two related tasks: prediction of protein-protein binding sites and B-cell epitopes (i.e. antibody binding sites). ScanNet outperforms baseline methods based on ML, structural homology and surface-based geometric deep learning. We further visualize and interpret the representations learnt by the network. We find that they encompass known handcrafted features, and find filters detecting simple, generic structural motifs such as hydrogen bonds as well as filters recognizing complex, task-specific motifs such as O-rings and transmembrane helical domains. Applied to the SARS-CoV-2 spike protein, ScanNet predictions validate known antigenic regions and predict a previously uncharacterized one.

## RESULTS

### A. Spatio-chemical Arrangement of Neighbors Network (ScanNet)

ScanNet takes as input a protein structure file, and optionally, a position-weight matrix derived from a multiple sequence alignment and outputs a residue-wise label probability. Its four main stages, shown in Fig. 1 and detailed in Materials and Methods, are: atomic neighborhood embedding, atom to amino acid pooling, amino acid neighborhood embedding and neighborhood attention.

**FIG. 1.**
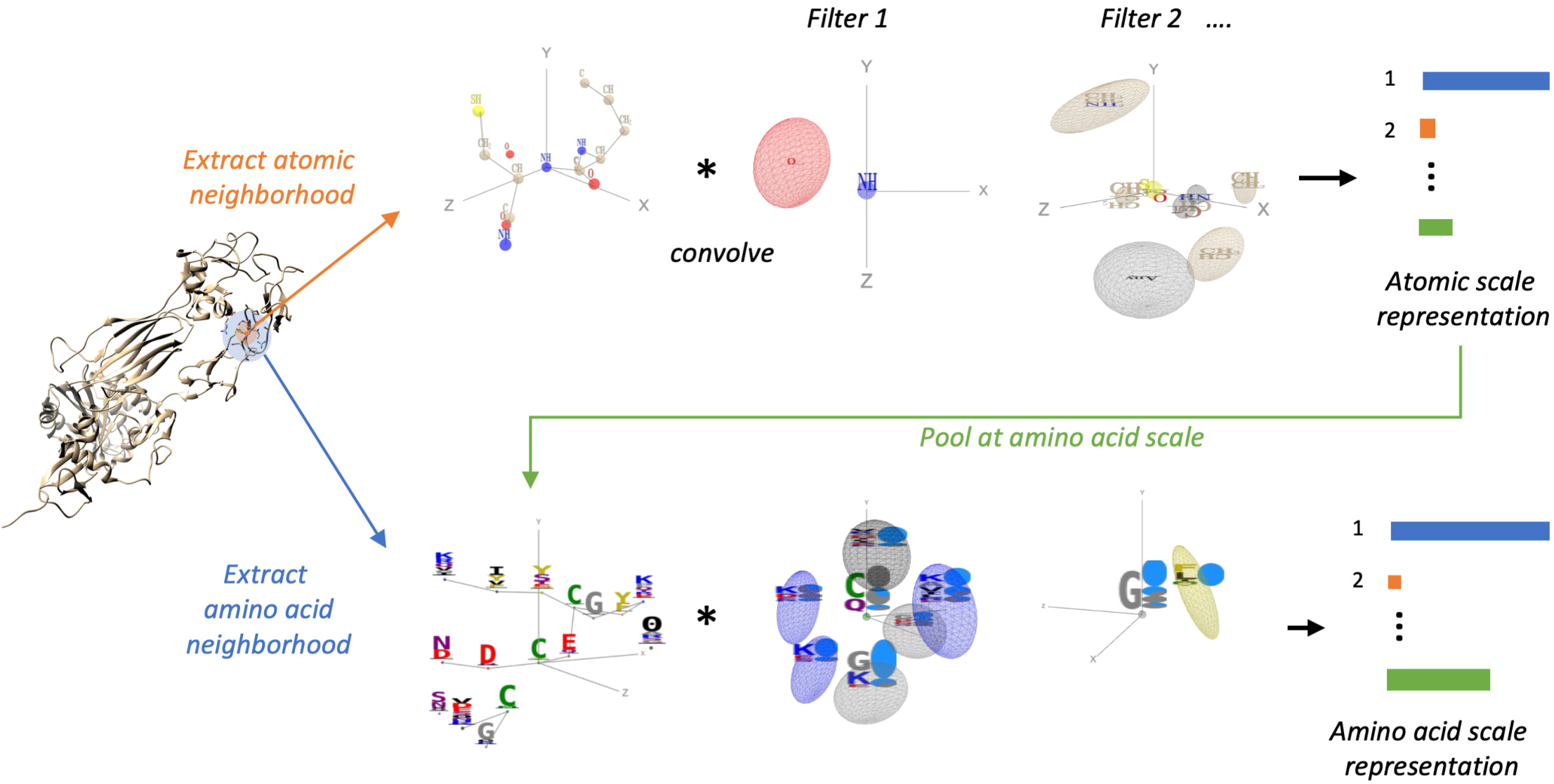
Overview of the ScanNet architecture. ScanNet inputs are the primary sequence, tertiary structure, and, optionally, position-weight matrix (PWM) computed from a multiple sequence alignment (MSA) of evolutionary-related proteins. Firstly, for each atom, neighboring atoms are extracted from the structure and positioned in a local coordinate frame (top left). The resulting point cloud is passed through a set of trainable, linear filters detecting specific spatio-chemical arrangements (top middle), yielding an atomic scale representation (top right). After aggregation of the atomic representation at amino acid (AA) level and concatenation with AA attributes, the process is reiterated with AA to obtain a representation of AA (bottom). The later is projected and locally averaged for residue-wise classification.

ScanNet first builds, for each heavy atom, a local coordinate frame centered on its position and oriented according to its covalent bonds. Next, it identifies its closest neighboring atoms. The resulting neighborhood, formally a point cloud with coordinates and attributes (atom group type) is passed through a set of spatio-chemical linear filters to yield an atom-wise representation. Each filter outputs a matching score between its (trainable) spatio-chemical pattern and the neighborhood. The patterns, which are parameterized using Gaussian kernels and sparse bilinear products, are localized in both physical and attribute space. Localization facilitates interpretation and is biologically motivated since motif functionality is often born by a few key atomic groups / amino acids in a specific arrangement, whereas other neighbors are irrelevant and interchangeable. Trainable, localized spatio-chemical patterns generalize to proteins the well-known concept of pharmacophores for small molecules.

Towards calculation of amino acid-wise output, the atom-wise representation is pooled at the amino acid scale and concatenated with embedded amino acid-level information (either amino acid type or position-weight matrix). Importantly, the constituting atoms of an amino acid have various types and may play different functional roles. In particular, some handcrafted features such as accessible surface area average information over all the atoms, whereas others, such as secondary structure consider only subsets (the backbone atoms). Therefore, a trainable, multi-headed attention pooling operation capable of learning which atoms are relevant for each feature is employed rather than a conventional symmetric pooling operation like average or maximum.

The neighborhood embedding procedure is then repeated at the amino acid scale: a local coordinate frame is constructed for each amino acid from its *C_α_* atom, side-chain orientation and local backbone orientation and its nearest neighbors are identified. The resulting neighborhood with *learnt* attributes is passed through a set of trainable filters to yield an amino-acid wise representation.

Finally, spatially-consistent output probabilities are obtained by projecting the amino-acid representations to scalar values, smoothing them across a local neighborhood and converting to probabilities with a logistic function. The smoothing scheme integrates two specifics of protein binding sites. First, protein-protein interactions are frequently driven by key ”hotspot” residues that contribute most of the binding energy, whereas other ”passenger” nearby residues have a small contribution to the binding energy [20, 42]. Such passenger residues are harder to detect directly as they do not necessarily have the salient features of protein-protein binding sites [43]. Second, some amino acid pairs consistently have *opposite* binding site labels - in particular, consecutive amino acids along the sequence because their side chains typically point in opposite directions. Altogether, this motivates the introduction of trainable, attention-based weighted averages, with *algebraic* weights.

### B. ScanNet for prediction of protein-protein binding sites

The protein-protein Binding Sites (PPBS) of a protein are defined as the residues directly involved in one or more native, high affinity protein-protein interaction (PPI). Not every surface residue is a PPBS, as (i) binding propensity competes with structural stability and (ii) PPI are highly partner and conformation-specific. Knowledge of the PPBS of a protein provides insight about its *in-vivo* behavior, particularly when its partners are unknown and can guide docking algorithms. Prediction of PPBS with conventional approaches is challenging as PPBS structural motifs are more diverse, less conserved and more extended than small molecule binding sites. Additionally, only incomplete and noisy labels can be derived from structural data, as (i) most PPIs of a given protein are not structurally characterized, and (ii) a substantial fraction (~ 15% [44]) of the structurally characterized protein-protein interfaces are not physiological but crystal-induced.

We constructed a non-redundant data set of 20K representative protein chains with annotated binding sites derived from the Dockground database of protein complexes [45]. The PPBS data set covers a wide range of complex sizes, types, organism taxonomies, protein lengths (Fig.S4 (a)-(d)) and contains around 5M amino acids, of which 22.7% are PPBS. To address the uneven sampling of the protein space, we introduced sample weights for each chain that are inversely proportional to the number of similar chains found in the data set (Materials and Methods and Sup. Fig. S4(h)). To investigate the relationship between homology and generalization error, we divided the validation/test sets into four splits based on the degree of homology with respect to their closest train set example (see Fig. 2 and Sup. Fig. S4(g)).

**FIG. 2.**
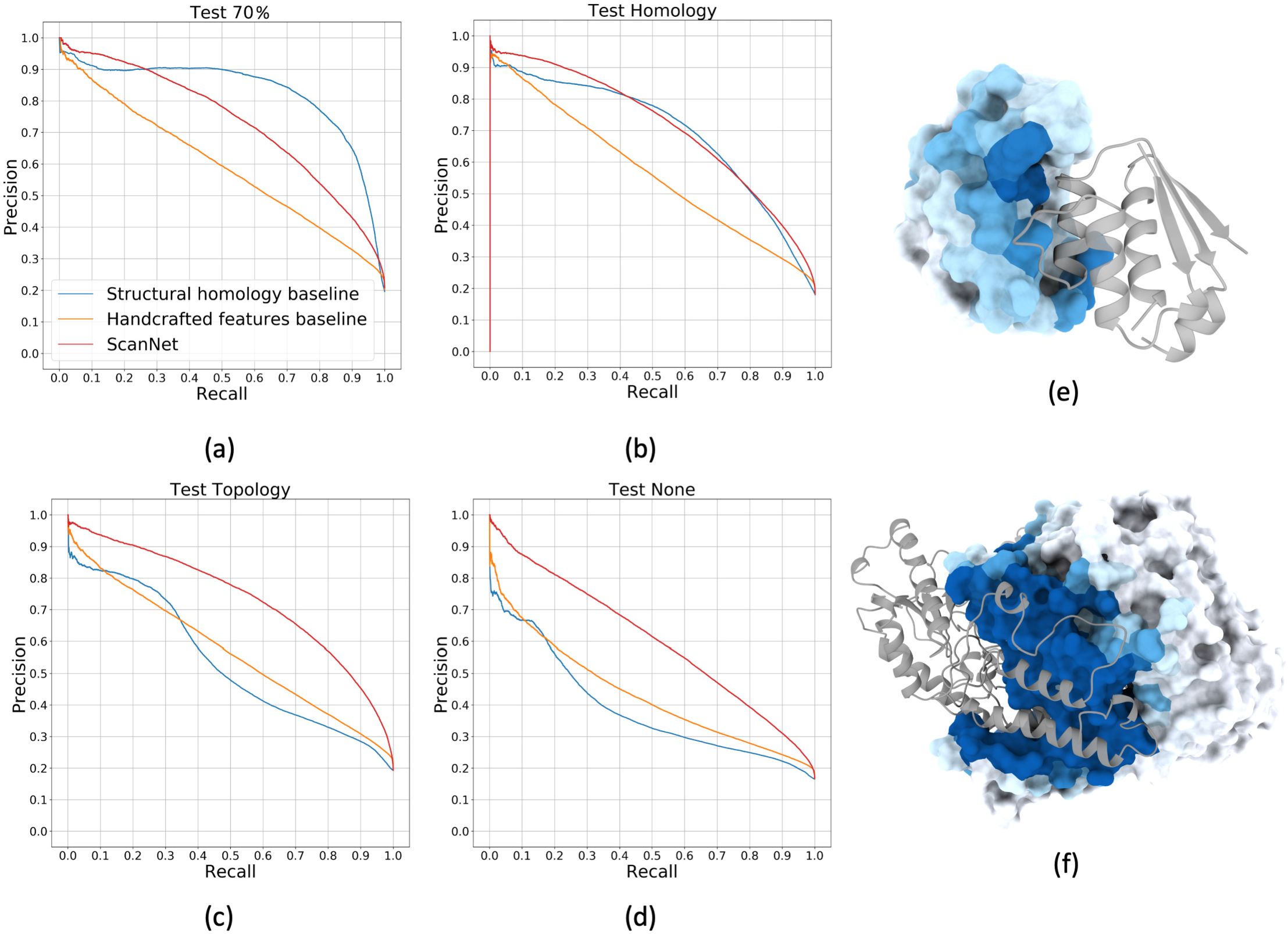
Prediction of Protein-Protein Binding Sites (PPBS) with ScanNet. (a)-(d) Precision-Recall curves of PPBS prediction for ScanNet, Structural homology, and Handcrafted features baseline methods (see main text). Train and test sets constructed from the redundant Dockground template database [45]. Proteins of the test set are subdivided into four nonoverlapping groups (a) *Test 70*%: At least 70% sequence identity with at least one train set example. (b) *Test Homology*: At most 70% sequence identity with any train set example, at least one train set example belonging to same protein superfamily (H level of CATH classification [48]). (c) *Test Topology*: At least one train set example with similar protein topology (T level of CATH classification [48]), none with similar protein superfamily. (d) *Test None*: None of the above. (e),(f) Illustration of predicted PPBS for (e) an enzyme (barnase, PDB ID: 1brs:A [57], Val. Homology dataset) with its inhibitor overlaid and (f) an homodimer (glutamic acid decarboxylase GAD67, PDB ID: 2okj:A [58], Test Topology dataset). The molecular surface of the query protein is shown with coloring based on predicted probability, ranging from low (white) to high (dark blue). The partner protein is shown in cartoon representation (gray transparent). Visualization software: ChimeraX [59]

We evaluated three models on the PPBS data set: (i) ScanNet, (ii) a ML pipeline based on handcrafted features and (iii) a structural homology pipeline (see Materials and Methods for technical details). For the handcrafted features baseline, we computed for each amino acid various geometric, chemical and evolutionary features, and used xgboost, a state-of-the-art tree-based classification algorithm [46]. For the structural homology pipeline, pairwise local structural alignments between the train set chains and the query chain were first constructed using MultiProt [47]. Then, alignments were weighted and aggregated to produce binding site probabilities for each amino acid. For all three models, the validation set was used for hyperparameters selection and early stopping, and performance is reported on the test set. Training and evaluation of a single model took one to two hours for ScanNet (excluding preprocessing time, ~ 10*ms* per step using a single Nvidia V100 GPU), few minutes for the ML baseline (excluding feature calculation time, using Intel Xeon Phi processor with 28 cores) and one month for the structural homology baseline (Intel Xeon Phi processor with 28 cores). We also evaluated Masif-site [36], a surface-based geometric deep learning model. Since Masif-site was not trained on the same data set, we only report its global test set performance.

We found that for the full test set, ScanNet achieved an AUCPR of 0.694 (Table I), accuracy of 87.7% (Sup. Table S3) and 73.5% precision at 50% recall (Sup. Fig. S17), the best performance by a substantial margin. The next best model was the structural homology baseline, whereas Masif-site and the handcrafted features model performed similarly. The model ranks differed when considering only subsets (Fig. 2 (a)-(d)). Unsurprinsingly, the structural homology baseline performed best in the high homology setting, but its performance degraded rapidly with the degree of relatedness; when the test protein had no similar fold in the train set, it was the worst algorithm. Conversely, the performance of the handcrafted features baseline increased slowly with the degree of homology, meaning that it could not faithfully recognize previously seen folds. In contrast, ScanNet could both recognize previously seen folds and generalize to unseen ones.

**TABLE I:** Performance evaluation for prediction of Protein-protein binding sites. Area under Precision Recall Curve (AUCPR) is shown. Proteins of the test set are subdivided into four non-overlapping groups. *Test 70*%: At least 70% sequence identity with at least one train set example. *Test Homology*: At most 70% sequence identity with any train set example, at least one train set example belonging to same protein superfamily (H level of CATH classification [48]). *Test Topology*: At least one train set example with similar protein topology (T level of CATH classification [48]), none with similar protein superfamily. *Test None*: None of the above. For Masif-site, only the aggregated performance is shown since its training set differs from ours. See Sup. Tables S2,S3 for additional evaluation metrics

| Algorithm | Test (70%) | Test (Homology) | Test (Topology) | Test (None) | Test (All) |
| --- | --- | --- | --- | --- | --- |
| Structural homology baseline | **0.828** | 0.696 | 0.535 | 0.387 | 0.613 |
| Handcrafted features baseline | 0.596 | 0.567 | 0.568 | 0.432 | 0.537 |
| Masif-site [36] | NA | NA | NA | NA | 0.533 |
| ScanNet | 0.732 | **0.712** | **0.735** | **0.605** | **0.694** |
| ScanNet (no evolutionary information) | 0.672 | 0.648 | 0.685 | 0.565 | 0.639 |
| ScanNet (no atomic information) | 0.697 | 0.672 | 0.689 | 0.547 | 0.648 |
| ScanNet (no regularization) | 0.756 | 0.702 | 0.701 | 0.572 | 0.678 |
| ScanNet (no reweighting) | 0.702 | 0.668 | 0.683 | 0.553 | 0.648 |

Visualizations of ScanNet predictions for representative examples (Fig. 2 (e),(f) and Sup. Fig. S9,S10,S11,S12) illustrate that predictions are spatially coherent and that in most cases, the binding sites are correctly identified. Overall, the network performed uniformly well across complex types and sizes, protein lengths and organisms (Sup. Fig.S8). PPBS identification was slightly harder when no or few homologs were found in the MSA (Sup. Fig.S8 b) and slightly easier for enzymes (Sup. Fig.S8 d). We next identified and visualized train and test examples on which ScanNet performed poorly (Sup. Fig. S13). We found *bona fide* false negative (undetected interacting patches) and false positives (predicted interacting patches), although for the later we could not rule out involvement in another PPI for which no structural data was available. Another source of mistake was confusion between types of binding sites: we found at least one instance where the incorrectly predicted PPBS were actually RNA binding sites. However, only a minority of RNA binding domain were confused as protein binding (Sup. Fig. S15) Finally, confusion between crystal and native interfaces was a substantial source of apparent mistakes. We found several train set examples in which the network ”refused” to learn the train label and instead predicted another binding interface with high confidence (Sup. Fig. S14). The predicted binding sites matched well the interface found in another biological assembly file. We found a posteriori that the biological assembly files used in the train set were annotated as probably incorrect by QSbio [44]. Overall, this demonstrated the robustness of predictions with respect to noise in training labels.

We next performed ablation experiments to investigate the importance of the network components (Table I, Sup. Fig. S16, Sup. Fig. S17). ScanNet performance decreased but remained above the other methods when discarding the evolutionary information (by replacing the position-weight matrix by the one-hot encoded sequence) or all the atomic-scale information (by removing the first two modules). Removing the sparse regularization on the spatio-chemical patterns and the early stopping yielded an homology-like performance profile, with better performance in the high homology setting but poorer otherwise. Lastly, training the model on all chains without redundancy reduction nor using sample weights yielded worse performance, highlighting the importance of sample weights.

Finally, we investigated the impact of conformational changes upon binding (*i.e*. induced fit) on ScanNet predictions using the Dockground unbound X-ray and simulated data sets [45]. Overall, predictions based on bound and unbound structures were highly consistent, and accuracy decreased only mildly from bound to unbound (Materials and Methods Sec. D, Sup. Fig. S5, Sup. Table S1).

### C. Visualization and interpretation of the learnt representations

What did ScanNet learn? Does the network reason solely by comparison with training instances or does it learn the underlying chemical principles of binding? How will it behave in out-of-sample settings such as disordered regions? To better understand the learnt representations, we visualized the spatio-chemical patterns and low-dimensional projections of the representations at the atomic (Fig. 3) and amino acid (Fig. 4) levels.

**FIG. 3.**
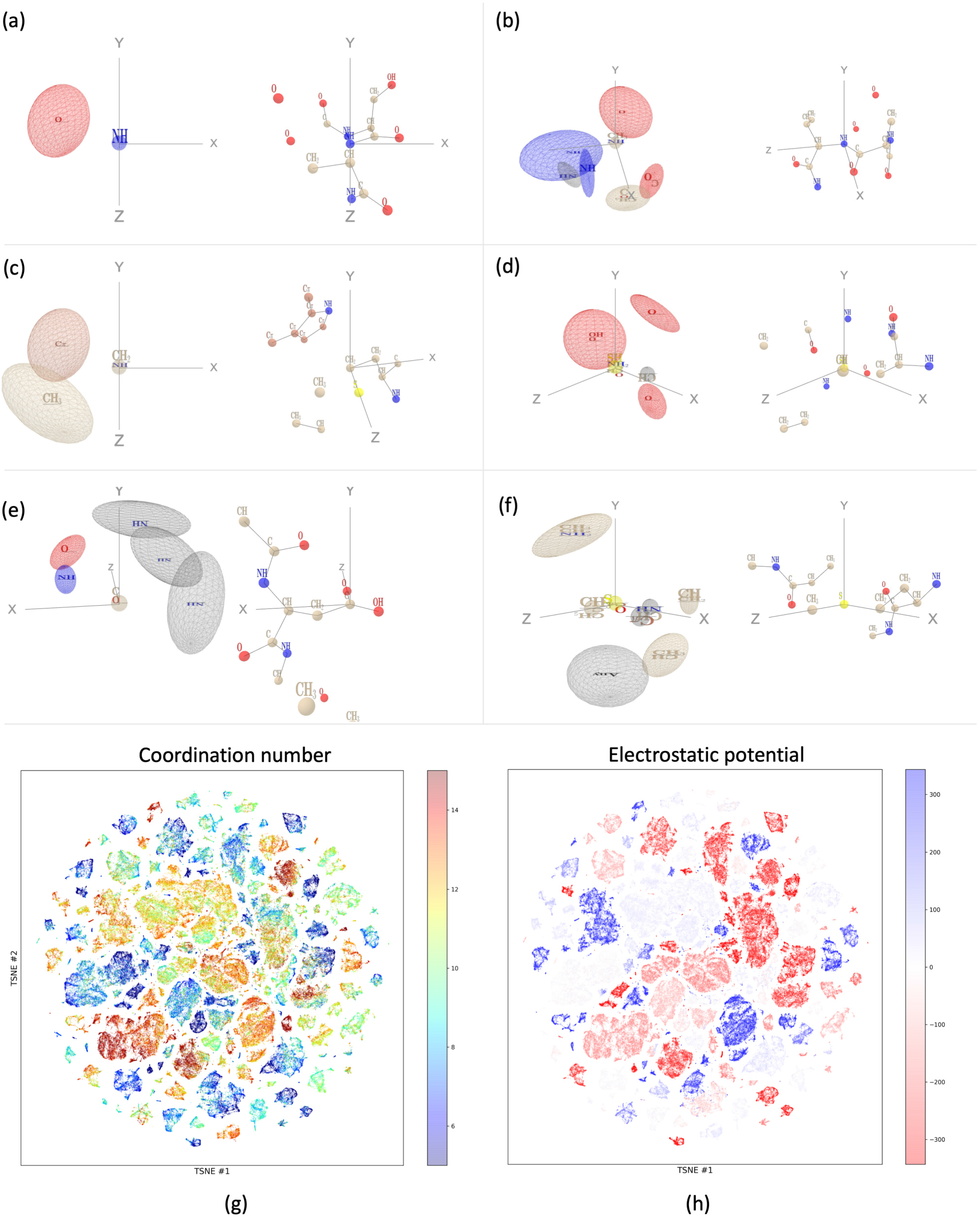
Visualisation of the learnt atomic representation. The panels (a)-(f) show each one spatio-chemical pattern on the left and one corresponding top-activating neighborhood on the right. Each pattern is depicted as follows: only the gaussian kernels relevant to the pattern are shown; they are represented by their unit ellipsoid. The corresponding locationwise attribute specificity is depicted as a weight logo inside the ellipsoid, similar to a position weight matrix: attributes with *non-zero* weights are stacked on top of one another with letter height proportional to their *algebraic* weight value, sorted from strongest positive (top) to strongest negative (bottom, reversed letters). The unit ellipsoid is colored based on the maximally activating attribute type if it is positive, or gray otherwise. Color code: carbon (beige), oxygen (red), nitrogen (blue), sulfur (yellow). The frame is overlaid in gray, with axes extending over 3.75 Å. Each filter/neighborhood pair is oriented independently for clarity. Visualizations created with pythreejs. (g), (h) Two-dimensional projection of the learnt atomic scale representation using T-SNE [60]. Each point corresponds to one atom of a representative set of proteins. Coloring based on atom coordination index (g) or electrostatic potential at the atom location, computed using APBS [49] (h).

**FIG. 4.**
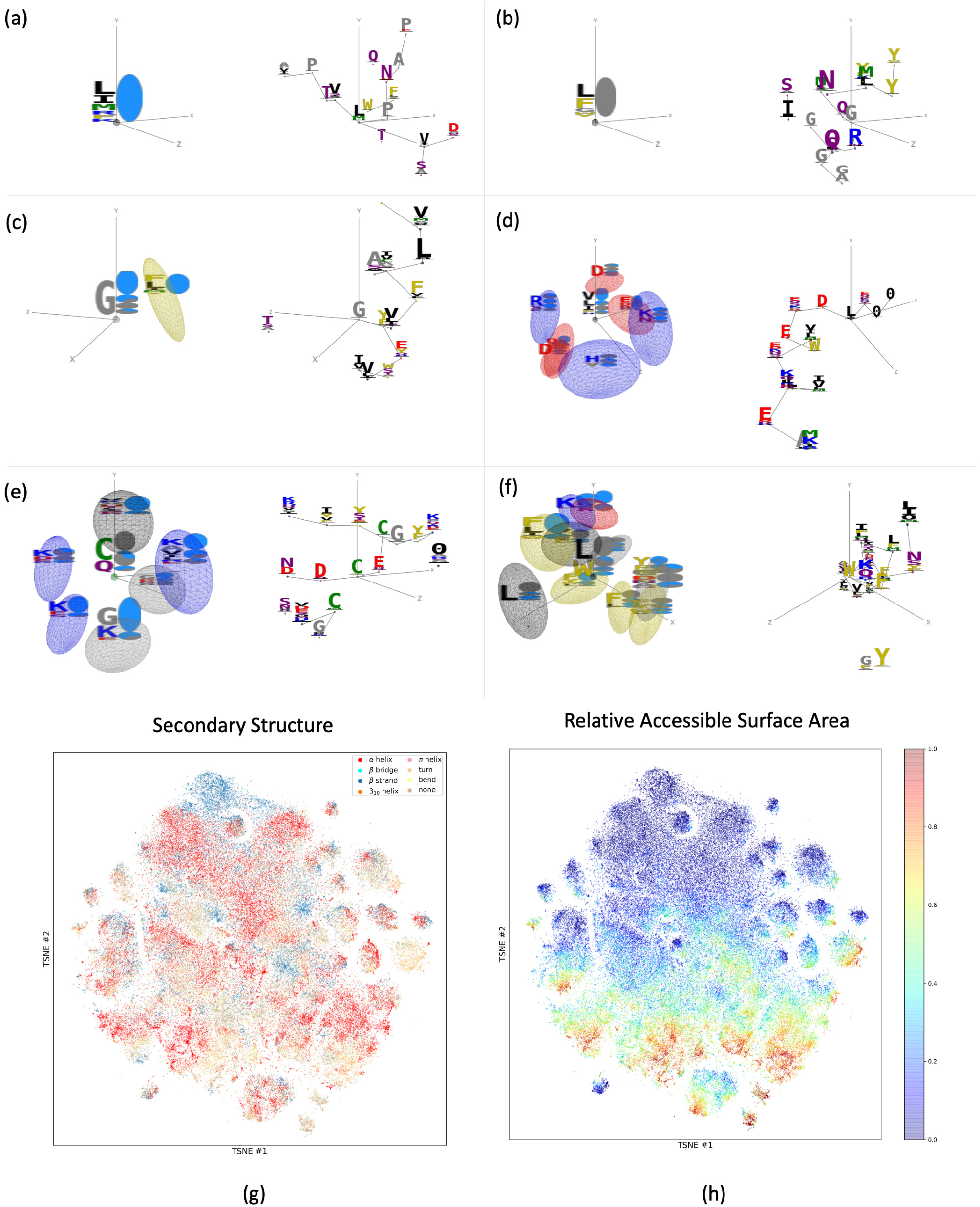
Visualisation of the learnt amino acid representation. The panels (a)-(f) show each one spatio-chemical pattern on the left and one corresponding top-activating neighborhood on the right. Gaussian kernels are depicted similarly as in Fig. 3. Since the input attributes are learnt, each component of a pattern is characterized by a complex specificity in attribute space. We represent it by the distributions of amino acid types and accessible surface areas of its top 1% maximally activating residues. The distributions are shown as a logo (each letter or symbol is proportional to the probability), with a total height proportional to the mean activation of the set. Accessible surface area values are discretized into four quartiles and represented as pie charts (from full gray = buried to full blue = accessible). Amino acids are colored by chemical properties: negatively charged (red), positively charged (blue), polar (purple), hydrophobic (black), sulfur-containing (green), aromatic (gold), tiny/proline (gray). The frame is overlaid in gray, with axes extending over 9 Å. Each filter/neighborhood pair is oriented independently for clarity. Visualizations created with pythreejs. (g), (h) Two-dimensional projection of the learnt amino acid scale representation using T-SNE [60]. Each point corresponds to one amino acid of a representative set of proteins. Coloring based on secondary structure (g) or accessible surface area (h) calculated with DSSP [61]. Additional T-SNE plots available in Sup. Fig. S19.

Recall that each pattern is composed by a set of gaussian kernels characterized by their location in the local coordinate system and specificity in attribute space. At the atomic scale, the origin corresponds to the central atom and the z-axis and xz-plane are oriented according to its covalent bonds. Panels (a)-(f) of Fig. 3,4 each show one pattern (left), together with a maximally activating neighborhood (right) taken from the validation set and the remaining patterns are provided as Supplementary Data. The atomic pattern shown in Fig. 3 (a) has two main components: a *NH* group located at the center and an oxygen located few Å away, in the (*x* < 0, *y* < 0, *z* < 0) quadrant, i.e. opposite from the two covalent bonds. It is the well known signature of a *N* – *H* – *O* hydrogen bond, ubiquitous in protein backbones. The corresponding maximally activating atom is indeed a backbone nitrogen within a beta sheet. Patterns may have more than two components, and several possible groups per location. The atomic pattern shown in panel (b) features two oxygen atoms and three NH groups in a specific arrangement; the corresponding maximally activating neighborhoods are backbone nitrogens located at contact zones between two helical fragments (right of panel (b) and Sup Fig.S18). Patterns shown in panels (c),(d) focus on side chains. Pattern (c) is defined as a carbon in the vicinity of a methyl group and an aromatic ring. Pattern (d) consists of *SH* or *NH*_2_ groups - two side chains-located hydrogen donors - surrounded by oxygen atoms. Lastly, patterns may include prescribed absence of atoms in specific regions. Pattern (e) is defined by a backbone carbon or oxygen without any *NH* groups in its vicinity, meaning that it identifies backbones available for hydrogen bonding. Pattern (f) identifies a methionine side chain with one solvent-exposed side, and is associated with high PPBS probability. Together, the filters collectively define a rich representation capturing various properties of a neighborhood, as seen from the 2D T-SNE projections colored by properties (Fig. 3 (g),(h)). In the space of filter activities, atoms cluster by coordination number (number of other atoms in range of Van Der Waals interaction) and electrostatic potential (calculated with the Adaptive Poisson-Boltzmann Solver [49]).

The amino acid scale patterns can be similarly analyzed: the origin, *z* axis and *xz* plane are respectively defined by the *C_α_*, side-chain and backbone orientation of the central amino acid. Neighborhoods are shown as backbone segments, with position weight matrices as attributes; the learnt attributes pooled from the atomic scale are not shown. Each gaussian component of a pattern is characterized by a complex specificity in attribute space. We represent it by the distributions of amino acid types and accessible surface areas of its top 1% maximally activating residues. Patterns (a) and (b) focus only on the central amino acid, i.e. they recombine and propagate features from the previous layers. Pattern a) consists of solvent exposed residues of type frequently encountered in protein-protein interfaces such as Leucine or Arginine. It is positively correlated with the output probability (*r* = 0.31). Conversely, pattern (b), which consists of buried hydrophobic amino acids, is activated by residues within the protein cores and is negatively correlated with the output (*r* = −0.32).

Multi-component patterns are also found: pattern (c) consists of an exposed glycine together with and exposed aromatic or leucine amino acid and is correlated with binding (*r* = 0.18). Pattern (d) is constituted by an exposed hydrophobic amino acid surrounded by exposed, charged amino acids and is strongly correlated with binding (*r* = 0.29). It is remarkably similar to the hotspot O-ring architecture previously described by Bogan and Thorn [20]. Conversely, pattern (e), which consists of a central cysteine (possibly involved in a disulfide bond) surrounded by exposed lysines is negatively correlated with binding (*r* = −0.13).

Distributed patterns such as pattern (f) are found and hypothetically contribute to prediction by identifying domain-level context. Pattern (f), which consists of multiple aromatic and hydrophobic components, is strongly activated by transmembrane helical domains. Identification of transmembrane domain is indeed required for accurate prediction as the hydrophobic core / hydrophilic rim rule is reversed within membranes. Inversely, we expect that for disordered regions, only the filters with patterns focusing on a single amino acid such as (a) and (b) or a linear stretch such as (c) will contribute to the prediction, whereas the others will be silent. ScanNet will thus effectively behave as a convolutional sequence model with a short kernel width.

Finally, the two dimensional T-SNE projections of the representation (Fig. 4 (f), (g) and Sup. Fig. S19) show that the filter activities encompass various amino-acid level handcrafted features, including amino acid type, secondary structure, accessible surface area, surface convexity and evolutionary conservation.

Overall, these findings support the hypothesis that ScanNet learns some of the underlying physico-chemical principles of protein-protein interactions. To consolidate these findings, we compared ScanNet predictions to experimental alanine scans and residue contributions to the binding energy using Rosetta (Materials and Methods, Sup. Fig.S7). We found that among the binding residues, the ones with higher binding probability and larger attention coefficients tend to contribute more to the binding free energy. Additionally, the amino acid filter activities reflected the type of interaction (van der Waals, electrostatic, etc.) involved in binding.

### D. ScanNet for prediction of B-cell Epitopes

B-cell epitopes (BCE) are defined as residues directly involved in a antibody-antigen complex. Although *a priori* every surface residue is potentially immunogenic, some are preferred in the sense that it is easier to mature antibodies targeting them with high affinity and specificity. Exhaustive, high-throughput experimental determination of BCEs is challenging, because they can span across multiple non-contiguous protein fragments. Prediction is challenging owing to their instability throughout evolution, and the lack of exhaustive epitope mappings for a given antigen. *In-silico* prediction of BCE can be leveraged for constructing epitope-based vaccines and for designing non-immunogenic therapeutic proteins.

We derived from the SabDab database [50] a data set of 3756 protein chains (796 95% sequence identity clusters) with annotated BCE. 8.9% of the residues were labeled as BCE, likely an underestimation of the true fraction. The data set was split into five subsets for cross-validation training, with no more than 70% sequence identity between pairs of sequences from different subsets. We evaluated ScanNet in three settings: trained from scratch, trained for PPBS prediction without finetuning, and trained via transfer learning using the PPBS network as starting point. We compared it with the handcrafted features baseline, structural homology baseline and Discotope, a popular tool based on geometric features and propensity scores [51]. We also report the performance of ScanNet without evolutionary data, of the null predictor and of a predictor based on solvent accessibility only. ScanNet trained via transfer learning outperformed the other models, with an AUCPR of 0.178 and a Positive Predicted Value at L/10 of 27.5% (Fig. 5 (a), Sup. Table S4). This represents an enrichment of respectively 143%, 153% and 309% over Discotope, solvent accessibility-based and null prediction. ScanNet perfomed equally well with or without evolutionary information unlike for PPBS. Visualization of representative spatio-chemical patterns associated with high BCE probability sheds light on the similarities and differences between PPBS and BCE (Fig. 5 (b)-(e), the remaining filters are provided as Supplementary Data). We find Asparagine and Arginine-containing patterns (b,c) as well as linear epitopes ((c), shared with PPBS). Pattern (d) consists of exposed residues with alternate charges, and putatively indicates availability for salt-bridge formation (d). Finally, pattern (e) is constituted by an exposed, charged amino acid in the vicinity of two cysteines forming a disulfide bond. A possible explanation is that disulfide bond-rich regions are more structurally stable, hence easier to recognize with high affinity and specificity.

**FIG. 5.**
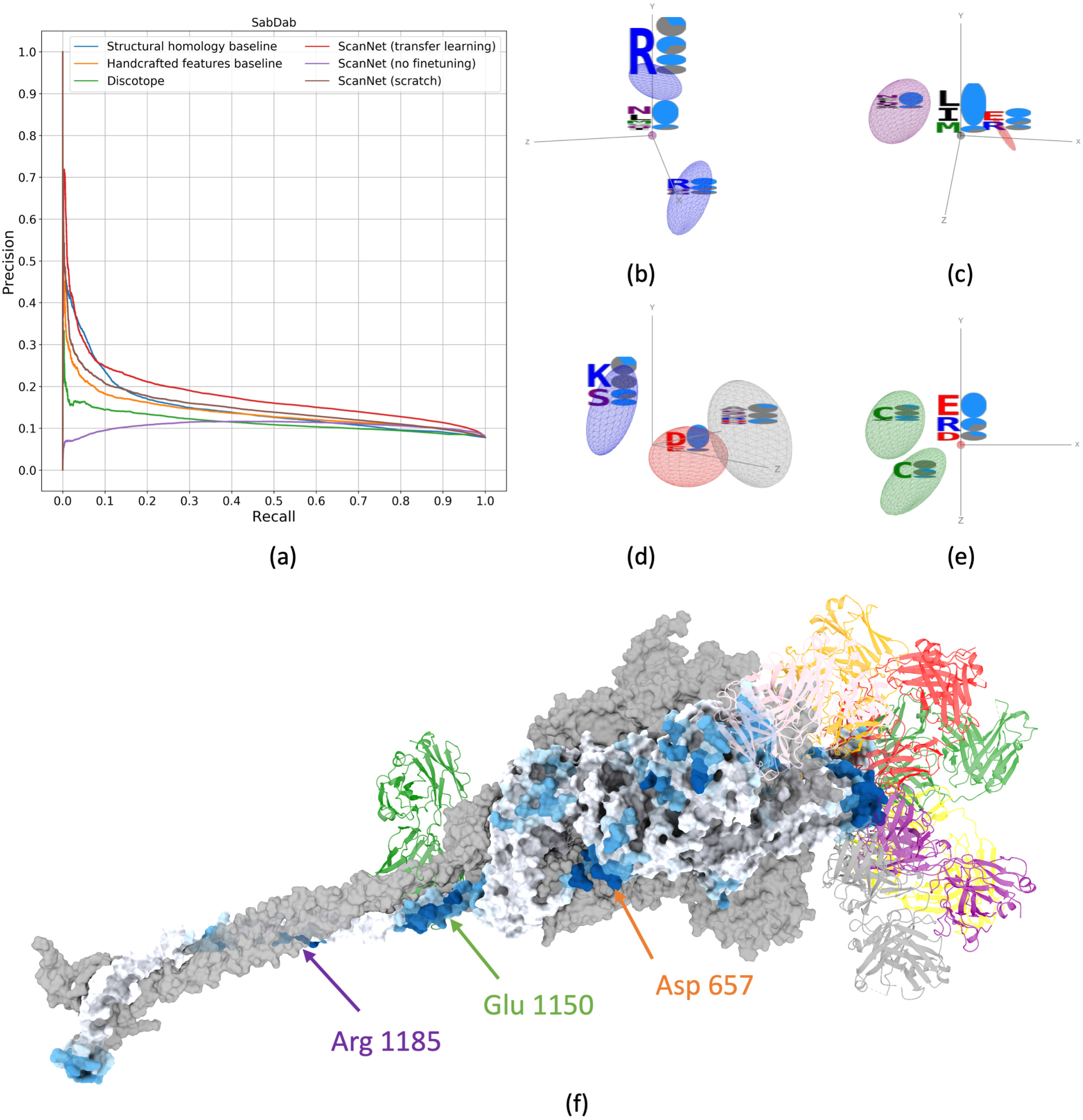
Prediction of B-cell Epitopes with ScanNet. (a) Precision-Recall curve of B-cell epitope prediction for baseline methods, Discotope [51] and ScanNet. Epitope database constructed from SabDab (timestamp: 04/19/2021) [50]; 5-fold crossvalidation performance is shown. (b)-(e) Selected learnt amino acid neighborhood filters whose activity is positively correlated with epitope probability. Same visualization as Fig. 4. (f) Application to Spike protein of SARS-CoV-2. Predictions performed on a MD snapshot of the spike trimer with one RBD open [62]. The monomer with open conformation is represented as molecular surface with color corresponding to BCE probability, from white (low) to dark blue (high). Representative antibodies binding the main epitopes are superimposed in color, cartoon representation, see full list in Sup. Table S6.

We next predicted and visualized BCE of the SARS-CoV-2 spike protein. Predictions are shown with representative antibodies superimposed for the trimer with one open Receptor Binding Domain (RBD) (Fig. 5 (e)) and for the isolated RBD and N-terminal domain (NTD) (Sup. Fig. S20). For the spike protein, the RBD was correctly identified as a major antigenic site. The six main epitopes previously described [52] all had high probabilities, including the cryptic epitope CR3022 (exposed in the open conformation). The tip of the N-terminal Domain (NTD) was also correctly identified as a highly antigenic site. Two linear epitopes located in the S2 fusion machinery are also predicted around Glu 1150 and Arg 1185 respectively. Previously, Shrock et al. [53] reported that both regions were targeted by antibodies from recovered Covid-19 patients. For the first one, a broadly neutralizing mAB targeting this epitope was recently isolated [54] and shown to neutralize several beta-coronaviruses but not SARS-CoV-2. Finally, the network predicted with high confidence one previously unreported conformational epitope constituted by three fragments in the vicinity of the glycosylated [55] Asn 657. Provided that the network is correct, and since the presence of the glycosyl group is unknown at run-time but can be imputed by ScanNet from the Asn-X-Ser/Thr linear motif, two interpretations are possible: either the glycosyl group shields an otherwise highly immunogenic region from antibodies or it directly induces immune response via glycosyl-binding antibodies. We similarly found two additional cryptic epitopes of the NTD which are centered on glycosylated asparagine when performing prediction on the NTD domain alone (Sup. Fig. S20 (b)).

Overall, ScanNet predictions are in excellent agreement with the known antigenic profile of the spike protein and predict a novel epitope that could not be detected via high-throughput linear epitope scanning. We additionally predicted BCE for three other viral protein: HIV envelope protein, influenza HA-1 and influenza HA-3 Hemagglutinin (Sup. Fig. S21). We notably found that the Hemagglutinin epitope predictions differed between the HA-1 and HA-3 strand despite the similar fold, suggesting that ScanNet could be suitable for studying antigenic drift.

## DISCUSSION

Protein function is born by a diverse set of structural motifs. These motifs, characterized by their complex spatio-chemical arrangements of atoms and amino acids, cannot be fully encompassed by handcrafted features. Conversely, detection via comparative modeling is challenging because their invariants, i.e. the set of function-preserving sequence/conformational perturbations are unknown. ScanNet is an end-to-end geometric deep learning model capable of learning such motifs together with their invariants directly from raw structural data by backpropagation. We demonstrated, through a detailed comparison on newly compiled datasets of annotated protein-protein binding sites and B-cell epitopes that it efficiently leverages these motifs to outperform feature-based methods, comparative modeling, and surface-based geometric deep learning. ScanNet reaches an accuracy of 87.7% for PPBS prediction and a positive prediction value at L/10 of 27.5 % for BCE prediction. Through appropriate parameterization and regularization, the spatio-chemical patterns learned by the model can be explicitly visualized and interpreted as previously known motifs and as novel ones. A breakthrough was recently achieved in protein structure prediction using DL [2], leading to the release of a vast set of accurate protein structure models [3]. We anticipate that ScanNet will prove insightful for analyzing these proteins, of which little is known regarding their function. A webserver is made available at http://bioinfo3d.cs.tau.ac.il/ScanNet/ and linked to both the PDB and AlphaFoldDB for ease of use. Very recently, Evans et al. [56] introduced AlphaFold-multimer, a novel approach for prediction of protein complexes from paired multiple sequence alignments and demonstrated impressive performance. We further compared ScanNet to AlphaFold-mutimer for prediction of partner-specific protein-protein binding sites, partner-agnostic protein-protein binding sites, and B-cell epitopes (Materials and Methods). We found that AlphaFold outperformed ScanNet for partner-specific PPBS, whereas both performed comparably for partner-agnostic PPBS. For B-cell epitopes, ScanNet could identify all the main epitopes of the Receptor Binding Domain of the SARS-CoV-2 spike protein, whereas AlphaFold-multimer could only identify one. This showcases the complementarity between MSA-based, partner-specific and structure-based, partner-agnostic approaches. Owing to its generality, it is straightforward to extend ScanNet to other classes of binding sites provided that sufficient training data is available. Extension to partner-specific binding prediction for prediction of interactions and guiding molecular docking is a promising future direction, as the amino acid filter activities are correlated between interacting binding sites (Sup. Fig. S22). Meanwhile, the learnt atom-wise and amino acid-wise representations can be readily used as drop-in replacement for handcrafted features in any structure-based machine learning pipeline. A second class of applications is protein design: ScanNet, which is differentiable with respect to its inputs and does not require evolutionary information, could be used in conjunction with structure prediction tools to guide design of proteins with prescribed binding or non-binding properties (e.g. non-immunogenic therapeutic proteins).

Finally, interpretable, end-to-end learning, combined with self-supervised learning techniques could pave the way towards a complete dictionary of function-bearing structural motifs found in nature, deepening our understanding of the core principles underlying protein function.

## Supporting information

3D visualisation and activation statistics of all the atomic filters of the protein-protein binding site network

3D visualisation and activation statistics of all the amino acid filters of the protein-protein binding site network

3D visualisation and activation statistics of all the amino acid filters of the B-cell epitope network

## ACKNOWLEDGEMENTS

J.T. acknowledges financial support from the Edmond J. Safra Center for Bioinformatics at Tel Aviv University and from the Human Frontier Science Program (cross-disciplinary postdoctoral fellowship LT001058/2019-C). D.S. was supported by ISF 1466/18, Israel ministry of Science and Technology and HUJI-CIDR. This work was supported by Len Blavatnik and the Blavatnik Family Foundation. We are grateful to Sonia Lichtenzveig Sela and the CS system team for their technical support. We thank Raphaël Groscot for his help on pythreejs visualizations. We thank Léa Naccache, Michael Nissan, Yoav Lotem, Mark Rozanov, Lirane Bitton, Matan Halfon and Shon Cohen for helpful discussions.

## AVAILABILITY

Data sets and source codes for training and evaluating ScanNet will be released upon publication.

## MATERIALS AND METHODS

The Materials and Methods is organized as follows. Section A provides all mathematical and implementation details for ScanNet. Section B is dedicated to the baselines methods. Section C covers data set construction, partition and sample weights. In Section D, we evaluate the impact of induced fit on changes on ScanNet predictions. We compare ScanNet to AlphaFold-multimer in section E. Section F covers the link between ScanNet prediction and binding site predictions. Section G contains additional results for the Protein-protein binding site and B-cell epitope prediction tasks.

### A. ScanNet network

#### 1. Preprocessing

##### PDB parsing

PDB files are parsed using Biopython [63]. We gather, for each chain, the amino acid sequence and the point cloud of heavy atoms, formally a list of triplets {(coordinates_*l*_, residue id_*l*_, atom id_*l*_) *l* ∈ [[1, *N*_atoms_]]}, e.g. ([10.1,101.3,-12.6], 97, CA). Only atoms belonging to classical residues are considered; exotic residues, additional molecules bound to the chain (e.g. heme, ATP, glycosyl groups, ions…) are excluded.

Towards definition of a local reference frame for each atom, we reconstruct the molecular graph (i.e. atom as nodes and covalent bonds as edges) using the residue and atom ids. Each heavy atom has one, two or three neighbors on the molecular graph; if it has only one (e.g. for methyl group *CH_3_*), a virtual hydrogen atom is appended to the graph. Two neighbors are selected to define a triplet of points 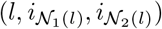 from which a frame can be derived. The coordinates of atoms 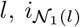 and 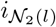 respectively define the center, *xz* plane and *z* direction, see paragraph on Frame Computation Module and Equation 3. The first (“previous”) neighbor is chosen as the closest from the N-terminal nitrogen. For the second (“next”) neighbor, if the atom has three neighbors, the furthest from the C-terminal carbon among the remaining two is used [64]. For instance, the two neighbors of the *C* atom of residue l are the *C_α_* of the residue *l* and the *N* of the residue *l* + 1. The two neighbors of the *C_β_* atom are the *C_α_* atom and the *C_γ_* atom of the side chain.

Also based on the molecular graph, an attribute is assigned to each heavy atom based on its type and the number of bound hydrogens. Twelve categories are defined: *C,CH,CH*_2_,*CH*_3_,*C_π_* (aromatic carbon), *O,OH,N,NH,NH*_2_,*S,SH*. Overall, four atomic arrays are constructed:

- The point cloud of atoms and virtual atoms (float, size [*N*_atoms_ + *N*_virtualatoms_, 3]).
- The triplets of indices for constructing atomic local frames (integer, size [*N*_atoms_, 3]).
- The atom groups (integer, size [*N*_atoms_,]).
- The residue index of each atom (integer, size [*N*_atoms_,]).

For the amino acid level, four similar arrays are constructed. The point cloud consists of the *C_α_* and the side chain centers of mass (SCoM) of each amino acid. For Glycines - which do not have a sidechain - a virtual SCoM is defined as 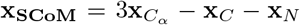, where **x***_C_α__*, **x***_C_*, **x***_N_* denote the coordinate of respectively the *C_α_,N* and *C* atom of the residue. The reference frame of each amino acid is defined by the *C_α_* (center), previous *C_α_* along the backbone (xz-plane) and SCoM (z-axis). Previous works [24, 30] considered other amino acid frames constructed from the backbone atoms only. Here, our rationale was that neighboring amino acids located in the opposite direction from the side chain (i.e. the interior of the protein) should not matter for functionality. It also facilitates filter interpretation, as for exposed residues the side-chain points towards the exterior of the protein. We also experimented with frames constructed from consecutive *C_α_* and found no difference performance-wise, but have not visualized the corresponding filters.

The per-residue attribute is given by the position-weight matrix (21-dimensional probability distribution, see below) or the one-hot-encoded sequence for the models without evolutionary information.

##### Derivation of the Position Weight Matrix

Given the sequence, we first construct a Multiple Sequence Alignment (MSA) by homology search using HHblits 2 (4 iterations, default values of other parameters) [65] on the Uni-Clust30_2018_06 database [66] (except for the SARS-Cov-2 Spike Protein for which we used the UniRef30_2020_06). Next, a sequence dependent weight *w*(*S*) was computed so as to i) address sampling redundancy [67] and ii) focus the alignment around the wild type [68]:

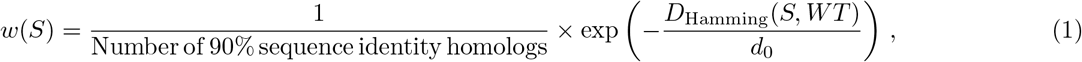

where *d*_0_ is adjusted such that the effective number of samples is 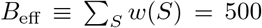. If the alignment is initially too small, *d*_0_ = ∞ is used. Focusing the alignments allows to detect local evolutionary conservation patterns as opposed to family-level conservation patterns; this is relevant as protein-protein interfaces are not always conserved at the superfamily level.

#### 2. ScanNet modules

##### Notations

The following notations are used throughout presentation of the modules; *x*: Global coordinates; *f*: Frames; *x^ℓ^*: Local coordinates; *a*: Attributes; *a^ℓ^*: Local attributes; *L:* Size of point set; *K*: Number of points in a neighborhood; *D:* Dimension of coordinates; *N/M*: Dimension of attributes. *G:* Number of Gaussian kernels. All upper case letters are integer dimension numbers. The corresponding lower case letter denote running indices e.g. *a_ln_* denotes the *n*’th 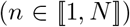 attribute of the *l*’th 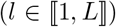 point of the point cloud and 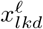 is the *d*’th local coordinate of the *k*’th neigbhor of point *i*. Bold letters denote vectors or matrices.

##### Attribute Embedding Module (AEM)

applies an element-wise non-linear transformation to the attributes *a_ln_* of each point. Here, we used a element-wise dense layer, *i.e*. a matrix product followed by ReLU non-linearity for all AEM except for the initial atomic attribute embedding module - for which the input is a categorical variable and a one-hot encoding layer is applied. The equation for the AEM writes:

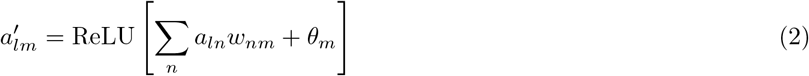

##### Frame Computation Module

(FCM) takes as input a point cloud *x_ld_* and a set of triplets of indices (*i*_*l*1_, *i*_*l*2_, *i*_*l*3_) and calculates, for every triplet, a frame *f_ldd′_* of size [*L*, 4, 3]), constituted by the center and the three unit vectors. The equation writes:

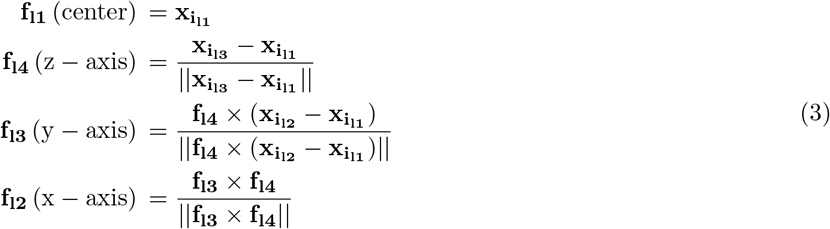

Where × denotes the cross-product. Examples of frames overlaid on a protein structure are shown in Sup. Fig. S1 (a,b). The FCM has no trainable parameters.

##### Neighborhood Computation Module

(NCM) determines, for each point, its *K* closest neighbors in space (including itself), computes their local coordinates and duplicates their attributes. Its inputs are a set of frames *f_lid_* and attributes *a_ln_*, and outputs are the neighborhoods 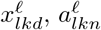. The nearest neighbor search is implemented naively by computing distances between all pairs of frame centers. For the atomic and amino acid neighborhoods, we use as local coordinates the three euclidean coordinates of the second frame center in the first frame and take *K* = 16. For the neighborhood attention module, we take *K* = 32 and use five coordinates: the distance between both frames centers 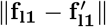, the dot product between the side-chain directions 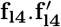, the dot product between the side-chain directions and the center to center vectors 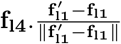 (and symmetric) and the distance between amino acids along the sequence (clipped at *d*_max_ = 8). They are shown respectively as *d,ω,θ,θ′,d*_sequence_ in Sup. Fig. S1 (c); The NCM has no trainable parameters.

##### Neighborhood Embedding Module

(NEM) is the core module of ScanNet. NEM convolves each neighborhood with a set of trainable spatio-chemical filters, akin to convolutional filters in image CNNs (Fig. 1). Its inputs are a set of *K* points with local coordinates 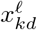 and attributes 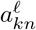, where *k* ∈ [1, *K*], *d* ∈ [1,*D*], *n* ∈ [1, *N*] respectively denote neighbor, coordinate and attribute indices. NEM outputs a set of *M* filter activities *y_m_*. It is parameterized using *G* = 32 gaussian kernels (as in [69]) and a bilinear product as follows:

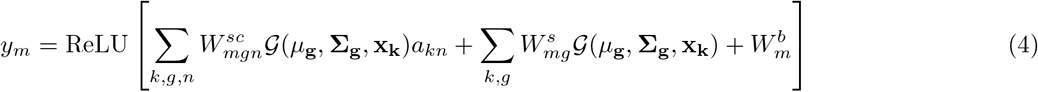

Where 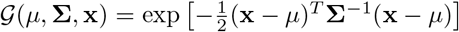 is a Gaussian kernel of center *μ* and (full) covariance matrix Σ, and **W^sc^, W^s^, W^b^** are trainable tensors of sizes [*M, G, N*], [*M, G*], [*M*,]. See a graphical sketch in Sup. Fig. S1 (d). The gaussian kernels are trainable and shared between all filters of a given layer, see implementation in Sup. Fig. S1 (e).

The above parameterization offers several advantages over other choices such as Multilayer perceptrons [25, 70, 71] or spherical harmonics [35, 72]. First, it is straightforward to interpret: a filter *m* with large entries of the tensor **W^sc^** for some *g,n* is positively activated by points having attribute *n* and located near the center of the Gaussian *g*. Similarly, the matrix *W^s^* encodes attribute-independent spatial sensitivity and *W^b^* is a bias vector. Second, localized filters, *i.e*. filters detecting only one or few combinations of point/attributes can be obtained by simply enforcing sparsity of the weights *W^sc^* and *W^s^* via a regularization penalty. Third, the filters are guaranteed to have an almost compact support, as the gaussian functions decay rapidly as ||*x*|| → ∞). This ensures that the diameter of the neighborhood is effectively capped irrespective of the local point density - in particular for unpacked or disordered regions. Last but not least, the gaussian kernels can be initialized using unsupervised learning, thereby improving performance and limiting run-to-run performance variance (initialization protocol detailed below).

For the sparsity regularization, we use the following combination of cost function and norm constraint:

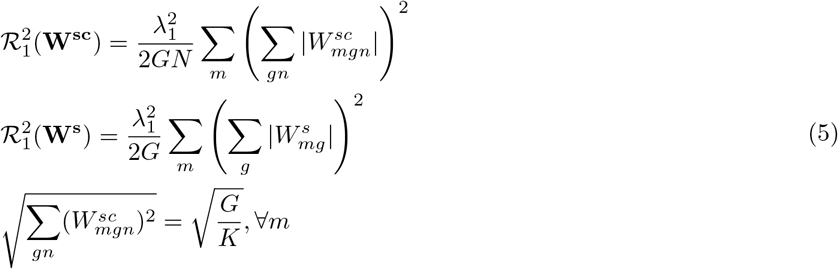

The so-called 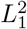 regularization (as previously described in [73]) is a variant of the *L*_1_ regularization 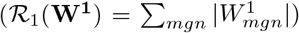 that promotes homogeneity of the filter sparsity values. This can be seen from the expression of the gradients, which write:

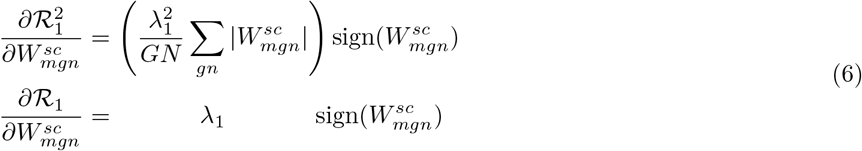

The 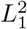 regularization is effectively a *L*_2_ regularization with a filter-dependent regularization strength: filters that are sparse (resp. not sparse) have a small (resp. large) *L*_1_ norm, hence a small (resp. large) effective *L*^1^ regularization strength; which in turn further relaxes or tightens the sparsity constraint. The *L*_2_ filter norm constraint is necessary to ensure a well-defined optimization problem because of the downstream batch norm layers. Indeed, the operation 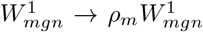 leaves the final output invariant, as it is exactly compensated by the covariation of the slope of the subsequent batch norm layer through 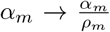 (using notations from [74]). Therefore, without constraint the optimum would be the asymptote 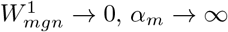 with 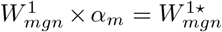, the optimum weight value without any regularization. The norm value is chosen such that the filter output *y_m_* (Eqn. 4) has roughly variance 1 when the attributes have variance 1.

To determine the value of the regularization penalty 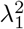, we searched for a satisfying compromise between interpretability (localized filters) and classification performance. We first determined the order of magnitude of 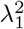 as follows: assuming filters weights *W*^1^ with sparse entries (a fraction *p* of non-zero weight, with typical weight value *W*), the *L*_2_ norm writes 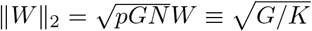, *i.e*. 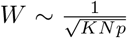 and 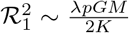. Further assuming that the regularization penalties and cross-entropy variations (about 10^−2^ per site in our experiments) should approximately balance each other, and with *G/K* = 2,*M* = 128 for both atomic and amino acid filters, we find that λ ~ 10^−2^ /*pM*. With a target *p* ~ 10^−2^, we conclude that 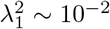. After experimentation, we chose 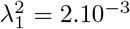 for both atomic and amino acid filters, as this value yielded the most satisfactory filter visualizations and prediction performances.

##### Atomic to Amino Acid Pooling

Towards calculation of residue-wise outputs, the learnt atomic scale representation must be aggregated at the amino acid scale. We recall that the constituting atoms of an amino acid may play different functional roles, hence symmetric pooling operations may not be sufficiently expressive. ScanNet instead employs a trainable multi-headed attention pooling. It writes:

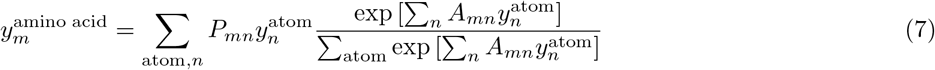

Where **P**, **A** are trainable projection and attention weighting) matrices. Eqn. 7 generalizes the average pooling (*A_m_*,. = 0) and maximum pooling (*A_m_*,. = *αP_m_*,. with large α) operations. A sparsity regularization is also employed for both **P**, **A** to simplify correspondence between atomic and amino acid filters.

##### Neighborhood Attention Module

(NAM) computes spatially coherent, residue-wise output probabilities from amino acid frames and spatio-chemical filter activities. The computation is done in four stages, see Sup. Fig. S2. First, local amino acid scale neighborhoods of size *K* = 32 are constructed, with graph-type local coordinates: distances, angles and sequence distances (see Sup. Fig.S1(c)). Second, the five-dimensional edges are projected element-wise into a single algebraic value using trainable Gaussian kernels followed by a dense layer with linear activation function. No bias is used for the dense layer, such that the edge value decays to zero as the distance increases. Third, the filter activities are projected to scalar values and locally averaged using attention-based weights. Our expression of the weighting coefficients slightly differs from the graph attention network formulation of [41] as follows: each node is characterized by a trainable output feature (unnormalized binding site probability), self-attention (”passenger” residues should have weak self-attention), cross-attention (hotspots should have strong cross-attention) and contrast coefficients (residues can follow either the majority or the hotspot residue). The weights may also take negative values depending on the edge values. Finally, a logistic function is applied to obtain normalized probabilities. Intuitively, the purpose of the NAM is to smooth out the probabilities such that if a residue has high binding propensity, its solvent-exposed neighbors should too. To this end, the NAM learns i) a diffusion kernel on the residue-residue graph (the algebraic edges) and ii) importance coefficients for each node.

#### 3. Full architecture

A diagram showing the architecture of the network is drawn in Sup. Fig. S3, and a table listing each module with its input(s) and output(s) sizes and comments is provided as Supplementary Data. In total, the network contains 475K parameters, of which about 200K are non-zero.

#### 4. Training

##### Initialization

For the neighborhood embedding modules, the gaussian kernels were initialized by unsupervised learning; using a subset of the training set, we computed atomic and amino acid neighborhoods, and fitted the spatial point density using a Gaussian Mixture Model (as implemented in Scikit-learn [75], best of 10 runs with Kmeans++ initialization, full covariance matrix and 10^−1^ covariance matrix regularization). For the trainable graph edges of the Neighborhood Attention module computed from distances and angles, we initialized them as a least square parametric fit of the label autocorrelation function (normalized):

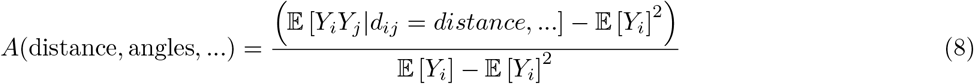

Intuitively, this initialization choice corresponds to a diffusion kernel over the residue-residue graph. All remaining weights are initialized using symmetric random distributions, see details in supporting table.

##### Padding and protein serialization trick

In our implementation, ScanNet takes as input an entire protein and computes neighborhoods on-the-fly, akin to a fully convolutional segmentation network [76]. Training on GPUs requires fixed size inputs but the lengths of proteins varied by almost two orders of magnitude in our dataset (see Sup. Fig. S4 (e)). To avoid truncating large proteins or wasting most of the computational power, we used the following protein serialization trick. We choose a relatively large maximal protein length (*L*_max_ = 1024, 2120 for the PPBS and BCE datasets), concatenate several proteins into a single example and translate each protein far away from the others, such that no two proteins overlap in space. Since ScanNet exploits only local neighborhoods, the predictions for each protein are fully independent from one another. Before training or prediction, we group proteins in a greedy fashion that minimizes the unused placeholders. Proteins are first sorted by length and the largest ones are first picked; then, we pick among the remaining proteins the largest that fits into the placeholder (if any), concatenate it, and continue until the placeholder is full. For the PPBS dataset, we found that about 96% of the amino acids placeholders were used, as opposed to less than 25% with naive padding. This results in a speed-up of about 4-fold. Finally, we used masking layers across the network to prevent backpropagating errors for the remaining placeholders that do not contain any residue.

##### Optimization

The network is trained by minimizing the binary cross-entropy loss function by backpropagation using the ADAM optimizer [77]. We set the maximum number of epochs to 100, the batch size to 1, the learning rate to 10^−3^ (10^−4^ for the transfer learning) and perform learning rate annealing and early stopping based on the validation cross-entropy; the optimal model was usually reached before 10 epochs. We used batch normalization layers before each ReLU non-linearity throughout the network to avoid vanishing gradients. Finally, regarding sample weighting, a complication of the protein serialization trick is that residues of a single example may have different sample weight as they come from different proteins. To account for this, we formally replaced the binary cross-entropy loss function and logistic non-linearity with a categorical cross-entropy and softmax function with two output classes; training labels are multiplied by their weight so as to replicate the weighted loss function.

##### Software and runtime

The network was implemented using Tensorflow v1.14.0 [78] and Keras v2.2.5 [79]. Training was completed in about one to two hours using a single Nvidia V100 GPU. The inference time is dominated by the construction of the MSA and the calculation of the Position Weight Matrix - it is of the order of one to few minutes depending on sequence length and MSA depth.

### B. Baseline Methods

#### 1. Handcrafted features baseline

For the handcrafted features baseline, we computed for each amino acid geometric, chemical and evolutionary features as described in recent works on prediction of protein-protein / protein-antibody binding sites [12–18]. The following features were computed:

- Amino acid type (one-hot encoded, 20 dim.).
- Secondary structure type (one-hot encoded, 8 dim.); computed with DSSP [61].
- Relative Accessible Surface Area (1 dim.); computed with DSSP [61].
- Coordination Number (1 dim.), defined as the number of *C_α_* atoms in a ball of radius 13 center around the *C_α_* atom of the amino acid.
- Half Sphere Exposure Index [80] (1 dim.), defined as follows: let *N*_1_ be the coordination number, and *N*_2_ the number of *C_α_* atoms in the intersection of a ball of radius 13 center and above the plane defined by the *C_α_* – *C_β_* vector. The half sphere exposure index is 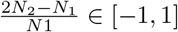.
- Backbone and Sidechain Depth [81] (2 dim.). The molecular surface was computed using MSMS (probe radius 1.5Å) [82], and the distance to the surface was computed and averaged for all backbone (resp. sidechain) atoms.
- Surface convexity index (3 dim.) [83]. For each atom, we construct a ball of radius 5/8/11 Åcentered on it, and compute the *f* fraction of its volume located on the inside of molecular surface; the index is given by 2*f* – 1 ∈ [–1, –1]. The surface convexity index is averaged at amino acid level.
- Position Weight Matrix (PWM, 21 dim.)
- Conservation score 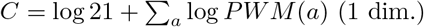.

In total, 58 features are used. For classification, we used the xgboost algorithm (boosted trees) [46]. The classifier was trained by cross-entropy minimization, using the same training and validation sets. We used 100 boosting rounds (with early stopping on validation loss), and the following four parameters were determined by grid search: tree depth (5,10,20), minimum child weight (5,10,50,100), *γ* (0.01,0.1,1.0,5.), *η* (0.5, 1.0).

#### 2. Structural homology baseline

Several approaches leveraging sequence and structure homology were previously developed [5–11], but were not readily available for large scale benchmarking, which prompted us to develop an in-house structural homology baseline method. It features three key components:

1. A non-redundant database of template protein chains with known binding sites. We used here as template the training set of ScanNet for a fair comparison. The template database was further clustered at the 90% (resp. 95%) sequence identity for the PPBS and BCE datasets, for speed gain purposes and to simplify alignment weighting, see below.
2. A local pairwise structure comparison engine. Compared to sequence homology or global structural homology, local structural homology were shown to outperform other methods in terms of coverage [7, 10]. Here, we used MultiProt [47], an algorithm we previously developed which, given two proteins, outputs a set of local structural alignments.
3. An alignment weighting scheme. Typically, MultiProt always finds at least few local alignments even when there is no homology between a query and a template protein, albeit with low coverage and low sequence identity. The alignments hence must be weighted so as to give higher importance to the most significant alignments [11]. Formally, for a given query protein with length *L*, MultiProt produces a set of *R* local alignments 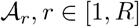. Each alignment is characterized by:

- The list of query residues included in the alignment, encoded as a binary vector:

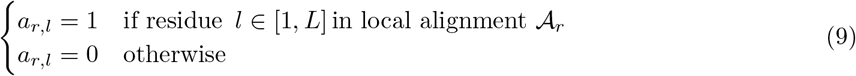
- The coverage of the local alignment: 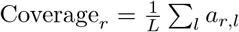
- The average root mean square deviation between matching pairs of *C_α_* atoms RMSD_*r*_
- The average sequence identity between query and template residues of the local alignment SeqID_*r*_

Combining the alignment and the corresponding binding site labels of the templates, we define the following label alignment matrix:

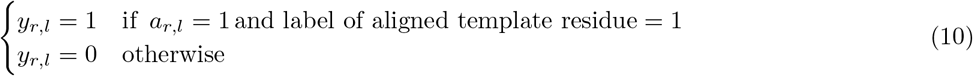

and write predicted binding site probability as:

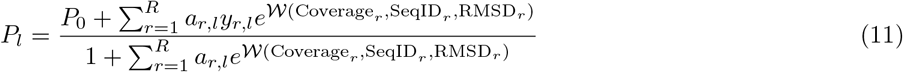

Where 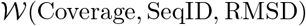 is a trainable log-weight function and *P*_0_ is a pseudo-count regularization term, such that *P_l_* = *P*_0_ if no alignment is found for a given residue. The log-weight function 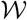 is parameterized by a two-layer perceptron with 20 hidden nodes and hyperbolic tangent activation function and was trained by crossentropy minimization on a subset of the validation set; after training, we found that 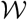 is a increasing function of both alignment coverage and sequence identity, in agreement with our intuition that high coverage/sequence identity alignments should be favored. For *P*_0_, we use the fraction of interface residues in the train set (resp. 0.22 and 0.09 for the PPBS and BCE train sets). Note that since the labels are already defined using multiple pdb files and redundancy reduction on templates was employed, there is no need to further reweight alignments by ligand diversity as described in [11].

As expected, the baseline performed very well when high quality homologs were available, and underperformed otherwise.

#### 3. Masif-site

We used the Docker image of Masif-site as made available at https://github.com/LPDI-EPFL/masif. Masif-site predicts binding site propensity at the surface vertex level. To aggregate at the amino acid level, we followed the aggregation scheme provided for the Masif vs Sppider comparison (https://github.com/LPDI-EPFL/masif/blob/master/comparison/masif_site/masif_vs_sppider/masif_sppider_Intpred_comp.ipynb): each surface vertex is first assigned to its closest atom and corresponding amino acid and the binding site probability of an amino acid is taken as the maximum binding site probability over all its corresponding vertices. We stress that the comparison with Masif-site should be interpreted with caution, as:(i) Masif-site predicts at surface vertex level rather than amino acid level. Its residue-wise probabilities are therefore not calibrated, resulting in bad likelihood scores (see Sup. Table S2). (ii) We did not retrain Masif-site owing to limited computational resources and ts training set used was smaller than ours (iii) Our test set overlaps with Masif-site training set, hence Masif-site should overperform on a fraction of our test set.

#### 4. Discotope

We used the Discotope version 1.1 as made available at https://services.healthtech.dtu.dk/software.php. To emulate the behavior of Discotope version 2.0, which processes entire protein assemblies rather than individual protein chains [51], we fused each multi-chain antigens into a single chain, and verified on a few examples that the outputs were consistent with the ones from the Discotope 2.0 webserver.

### C. Data preparation

#### Initial database and filtering

We use the Dockground database of protein-protein interfaces [45] (Jan. 2020 full redundant version) as a starting point for our Protein-Protein Binding Sites (PPBS) database. Each unique PDB chain involved in one interface or more was considered as a single example; we excluded chains with sequence length less than 10, chains involved in a protein-antibody complex (as classified in the SabDab database [50]), or designed proteins (identified as having two or more of the following red flags: no Uniprot ID, no known CATH class, no sequence homologs found, and engineered/synthetic/designed/de novo appearing in chain name). We obtained 70583 unique chains (grouped in 20025 clusters at 95% sequence identity) from 41466 distinct PDB files, involved in 240506 PPIs.

The dataset covers a wide range of complex sizes, types, organism taxonomies, protein lengths (Fig.S4 (a)-(d)). For the B-cell epitopes database, we used the SabDab database (timestamp: 04/19/2021 [50]) and included all antigens with length 10 or more forming an interface with an antibody with both heavy and light chain appearing in the PDB files. We obtained 3756 chains (grouped in 796 clusters at 95% sequence identity).

#### Data partition

For the PPBS database, we investigated the impact of homology between train and test set examples on generalization of ScanNet and our baseline models. We enforced a maximum sequence identity (90%) between a val/test example and any train set example and grouped validation and test examples into four subgroups based on their degrees of homology, see Sup. Fig. S4(g):

1. *Val/Test 70*%: At least 70% sequence identity with at least one train set example.
2. *Val/Test Homology*: At most 70% sequence identity with any train set example, at least one train set example belonging to same protein superfamily (H level of CATH classification [48]).
3. *Val/Test Topology*: At least one train set example with similar protein topology (T level of CATH classification [48]), none with similar protein superfamily.
4. *Val/Test None*: None of the above.

Subgroups are ordered by decreasing degree of homology; generalization is expected to be increasingly difficult. To ensure that the four subsets have approximately equal sizes, the following partitioning algorithm was employed. The chains are first iteratively clustered by sequence identity at several levels (100%, 95%, 90% seqID, 70% seqID) using CD-HIT [84] followed by clustering at homology and topology identifiers. If a 70% (resp. homology) cluster contains several distinct homology (resp. topology) categories, these categories are merged into a single one. Next, we constructed the *Val/Test None* by randomly drawing topology clusters and assigning all its members to either validation and test; this is repeated until *Val/Test None* are full. The *Val/Test topology* sets were constructed by randomly drawing from the remaining topology clusters with more than one homology cluster, and assigning half of the homology clusters to train and half to val/test. Similarly, the *Val/Test homology* and *Val/Test 70* are constructed similarly by drawing homology (resp. 70%) clusters with more than one 70% (resp. 90%) sequence identity cluster, and allocating each 70% (resp. 90%) cluster to either train or val/test. Finally, the remaining 90% clusters are randomly allocated to fill the training, validation and test sets (64% - 16% - 20% split).

For the BCE, the dataset was subdivided into 5 folds for cross-validation. Antigens were clustered at 70% sequence identity, and each cluster was assigned to one fold at random (except for SARS-CoV-2 antigens, which were all assigned to fold 1).

#### Label computation

An amino acid of a protein chain is labeled as a binding site if at least one of its heavy atoms is within 4^Å^ of another heavy atom from another chain within the biological assembly [12]. Next, since the same protein may appear in multiple assemblies, we take the union of all its binding sites found across pdb files. This is done by clustering sequences at 95% sequence identity using CD-HIT [84, 85], aligning the sequences and labels of each cluster using MAFFT [86] and propagating the labels along each column. We found that for the PPBS dataset, 91.2% of the binding sites were identified from the original pdb complex file and 8.8% were propagated from other pdb files.

For SabDab, we found that several epitopes appeared as accessible in one conformation of the protein and buried in another conformation; labels were propagated from one structure to another only if the residues had similar relative accessible surface area and coordination number (number of amino acids within 13Å). The propagation criterion writes:

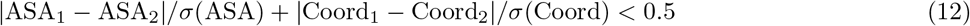

For the PPBS, we obtained 22.7% positive labels - 30% when only considering the surface residues, with relative accessible surface ≥ 25% (distributions shown in Sup. Fig.S4 (e,f)). For the BCE, we found 8.9% positive labels.

#### Sample weighting and subsampling

PDB covers unevenly the protein sequence space: many protein families do not have any representative structure, whereas others such as immunoglobulins have tens of thousands. The sampling is also biased within one family, as some genes and/or organisms are more frequently studied than others. To correct for the biases occurring at multiple scales, we apply the following hierarchical reweighting scheme:

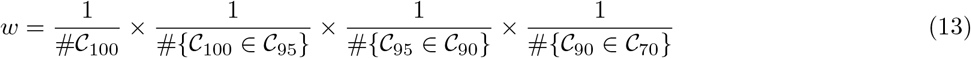

Where 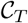 denotes the clusters at sequence identity cut-off *T*. This choice is such that each cluster at 70% sequence identity contributes a total weight 1; within each 70% cluster, each of the *K* 90% clusters contributes a total weight 1/*K*, and so on. An example of set of weights is illustrated in Fig. S4 (h).

In addition, this hierarchical choice ensures that the total weight of a cluster is invariant upon subsampling at some higher cluster identity level (e.g. the total weight of a 90% sequence identity cluster is invariant upon subsampling at 100%, 95% or 90% sequence identity). For the PPBS dataset, when hierarchical reweighting was used, we found no significant change of performance when training on the full set of chains or on 95% sequence identity representatives and therefore used the 95% sequence identity subset for speed gain purposes. When no reweighting or subsampling was used, performance significantly degraded, see Table I and Sup. Fig. S16. For the BCE database, the same approach was followed, without any subsampling - in order to include as many conformations as possible - and using a 90% sequence identity cut-off for the reweighting scheme, as similar proteins may have different epitopes.

### D. Impact of induced fit on ScanNet predictions

Protein structures undergo induced fit (i.e. conformational changes) upon binding. The magnitude of conformational changes varies, ranging from minimal rearrangement of side chain rotamers to extensive allosteric motion. ScanNet is mostly [87] trained on bound chains but applied to unbound ones. Owing to its high expressivity, it is a priori capable of picking up signature of bound conformations such as over-stretched side chains or unpacked helixes (see e.g. 4wwx:B of Sup. Fig. S14).

We evaluated the predictive performance of ScanNet on unbound chains for two datasets: the Dockground simulated and Dockground X-Ray [45]. The Dockground simulated data set consists of chains extracted from complex pdb files and relaxed using Langevin Dynamics simulations [88]. Simulating separately the bound protein structures, without the interacting partner for a short time period (1 ns) relaxes the side-chain conformations of the interface residues and reliably approximates the unbound form of the protein if conformational changes are small (< 2Å RMSD). We considered only the proteins that appeared in our data set and excluded four tetramers, obtaining 6012 chains. We used as ground truth the binding site labels of the PPBS data set (18.5% positive labels).

The Dockground X-Ray consists of chains that are both crystallized alone and in complex with their partner. It features chains undergoing larger conformational changes than the simulated data set one. We selected *N* = 709 (bound, unbound) pairs with at least 95% sequence identity between chains. As some complex component were multi-chains, there was no direct correspondence with our data set labels (which included inter-domain, intra-protein binding sites); instead, we used as ground truth labels the interface residues of the complex (6.6% positive labels). The reduction in the fraction of positive labels also stems from the longer length of proteins on average (respectively 331 and 221 for X-ray and simulated).

For both data set, we computed ScanNet predictions separately for the bound and unbound structures, excluded residues that did not match between the bound and unbound structure and compared both predictions residue-wise. Results are reported in Supplementary Table S1 and Supplementary Fig. S5. We find a good agreement between bound and unbound predictions (Pearson correlations of *r* = 0.86, *r* = 0.78 for simulated and X-ray data sets respectively). A slight drop in accuracy between bound and unbound structures is found: from 88.3% to 86.6% for the simulated set and from 91.9% to 91.3% for the X-ray set.

To further quantify the impact of global and local conformational changes on prediction, we calculated for each chain the root mean square deviation (RMSD) between the bound and unbound atomic coordinates and the RMSD between the bound and unbound solvent accessible surface area. Supplementary Fig. S5 (e),(f) show the per-chain Pearson correlation between bound and unbound predictions against the coordinate (resp. solvent accessibility) RMSD. As expected, structures with larger global/local conformational changes tend to exhibit significant changes in binding site predictions. Overall, we conclude that ScanNet predictions are overall robust to conformational changes, although improvements could be obtained by training on unbound structures.

**TABLE S1.** Performance evaluation for prediction of Protein-protein binding sites in the unbound setting. The gap of AUCPR and precision values between simulated and X-ray structures stems from the difference in the proportion of positive labels (respectively 18.5% and 6.6%).

| Data set | Accuracy | Likelihood | AUCPR | Precision at 50% Recall |
| --- | --- | --- | --- | --- |
| Dockground simulated (bound) | 0.884 | -0.280 | 0.736 | 0.793 |
| Dockground simulated (unbound) | 0.866 | -0.318 | 0.662 | 0.687 |
| Dockground X-ray (bound) | 0.919 | -0.216 | 0.294 | 0.276 |
| Dockground X-ray (unbound) | 0.912 | -0.234 | 0.238 | 0.225 |

### E. Comparison between ScanNet and AlphaFold2 binding site predictions

AlphaFold-multimer (AF2) is a recently released model for predicting the structure of protein complexes from paired multiple sequence alignments [56]. It is difficult to compare fairly AF2 and ScanNet, as the first one assumes knowledge of the partner and predicts partner-specific binding sites, whereas the second one does not assume knowledge of the partner and predicts partner-agnostic binding sites. We nonetheless benchmarked both approaches as follows. We considered Benchmark2, a set of 17 recently released dimers that do not appear in the training sets of AF2 and ScanNet [89]. For each of the 34 chains, we determined the ground truth partner-specific binding sites (i.e. involved in the complex) and partner-agnostic binding site (i.e. the union of all binding sites involved in any complex found among PDB structures with 95% or more sequence identity to the chain). Next, we ran AF2 (ColabFold version, no relaxation [90]) to predict the structure of the complex given the pair of sequences, obtaining five models. For each residue, the binding site probability was defined as the fraction of models in which it belongs to the interface (taking fractional values ∈ {0, 0.2,0.4, 0.6,0.8,1.0}; we also tested continuous values using the predicted alignment error at contacts, but found no improvement). We also experimented defining continuous binding site probability by using the predicted alignment error at contacts, but found no improvement. ScanNet was run on each chain to produce binding site probabilities (averaging of 11 models). The Area under the precision recall curve (AUCPR) was computed for each chain separately, and for both the partner-specific and partner-agnostic binding sites. Results are reported in Supplementary Fig.S6 (a),(b),(c). We found that for partner-specific binding site, AF2 outperformed ScanNet in 27/34 chains (Supplementary Fig.S6 (a)) whereas for partner-agnostic binding, the performances were comparable (19/34 better for AF2 and 15/34 better for ScanNet, Supplementary Fig.S6 (b)). Generically, ScanNet outperformed AF2 when a protein had multiple binding sites, whereas AF2 outperformed ScanNet when only a single binding site was known. Other examples where ScanNet outperformed AF2 were a mammal cell surface protein (6pnq, Supplementary Fig.S6 (c)), and a rice host-pathogen interaction (5zng) - for which no paired MSA can be constructed.

For B-cell epitope prediction, we tested AF2 on the Receptor Binding Domain (RBD) of the SARS-CoV-2 spike protein as follows. We first selected six representative antibody-antigen complexes spanning all the known epitopes of the RBD, following [52] (Sup. Table S6). We then predicted their structure with AF2, obtaining 6 × 5 = 30 models. The B-cell epitope propensity was defined residue-wise as the fraction of all models in which the residue is bound by antibodies. We found that AF2 systematically predicted a single binding mode roughly corresponding to the RBD-C epitope (Sup. Fig. S6 (d),(e)), whereas ScanNet correctly predicted multiple epitopes. We compared AF2 and ScanNet epitope propensity predictions with the empirical antibody hit rate calculated from 290 experimental structures of antibody-spike protein found in the PDB (Sup. Fig. S6 (d)), and found that the ScanNet profile better correlated (Spearman coefficient of respectively 0.74 and 0.6). Allegedly, AF2 failure stems from i) unavailability of a paired MSA ii) low sensitivity with respect to the antibody sequence and iii) unimodal rather than multimodal prediction.

### F. Link between ScanNet predictions and residue contribution to binding energy

Presumably, residues with high binding probability correspond to hotspots residues, i.e. residues with high contribution to the binding free energy of the complexes [43]. To test this hypothesis, we first compared ScanNet predictions to changes in binding affinity ΔΔ*G* measured after mutation of binding residues to alanine. Positive ΔΔ*G* indicate important residues, and hotspots are typically defined as ΔΔ*G* > 2*kcal.mol*^−1^. We found in the SKEMPI v2.0 database [91] 2122 mutations of binding residues to alanine, spread across 130 complexes. We calculated for each residue its binding site probability *p* and aggregated attention coefficient *a* (defined as 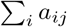 where *a* is computed as in Sup. Fig. S2). The later score quantifies the importance of the residue within the neighborhood; residues with high aggregated attention drive prediction of their neighborhood. Next, we estimated the conditional average 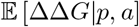 using a one-layer perceptron with 20 hidden units, hyperbolic tangent activation and non-negative kernel weights to enforce monotonicity (Sup. Fig. S7 (a),(b)). We indeed find that residues with high binding probability and large attention coefficient tend to be more important for binding.

We next performed a similar analysis using the Benchmark 5.5 dataset [92] (271 dimers, 10444 binding sites) and the Rosetta REF15 all-atom energy function [93]. For each dimer, the binding energy was estimated as the difference between the energy of the complex and the sum of the energies of the unbound structures. The FastRelax protocol of PyRosetta [94] was used to remove steric clashes prior to computation of the energies. We similarly find that residues with high binding probability and large attention coefficient tend to contribute a lower energy (Sup. Fig. S7 (c),(d)).

In addition, Rosetta allows to calculate the contribution of individual energy terms to the residue-wise binding energy. This raises the question of whether the types of interaction involved in binding can be predicted from the intermediate layer activities of ScanNet. We grouped the 19 energy terms into eight groups: Solvation (fa_sol+lk_ball_wtd+fa_intra_sol_xover4), van der Waals (fa_atr+fa_rep), Coulomb (fa_elec), Backbone-side chain hydrogen bonds (hbond_bb_sc), Side chain - side chain hydrogen bonds (hbond_sc), Side chain internal energy (fa_intra_rep+fa_dun+yhh_planarity), Backbone internal energy (omega+p_aa_pp+rama_prepro+hbond_sr_bb+hbond_lr_bb) and Others (pro_close+dslf_fa13+ref).

Next, we computed for each binding residue the vector of activities of the amino-acid spatio-chemical filters. We then performed a LASSO regression to predict residue-wise the value of each energy term from the filter activities (optimal regularization determined by cross-validation with scikit-learn [75]). The regression and correlation coefficients are shown in Sup. Fig. S7 (e). We find several ”hotspot” filters associated with negative binding energies, such as filters 81, 17, 57, 41, 2 and 22 (the O-ring filter represented in the main text). As expected, they are also strongly correlated with binding (*r* =0.47,0.19, 0.12, 0.31, 0.09,0.29 see filter depiction in Supplementary Data).

Interestingly, each filter displays a distinct energetic profile. For instance, filter 81 is associated with strong van der Waals binding without any cost in solvation energy; consistently, it is activated by hydrophobic residues already fully exposed in the unbound state (see filter depiction in Supplementary Data). The O-ring filter 22 is associated with both strong van der Waals and electrostatic energy, but at the expense of a higher solvation cost. Backbone-mediated interactions are also captured; for instance, filter 54, which corresponds to an exposed glycine/lysine tandem, is associated with strong backbone - sidechain hydrogen bonding.

Altogether, the comparative analysis with mutagenesis assays and Rosetta energy supports the claim that ScanNet learns some of the underlying physical principles of binding.

### G. Protein-protein binding site and B-cell epitope prediction: additional material

**TABLE S2.** Performance evaluation for prediction of Protein-protein binding sites. Average likelihood per residue (higher is better) is shown for the four subsets of test and the entire test.

| Algorithm | Test (70%) | Test (Homology) | Test (Topology) | Test (None) | Test (All) |
| --- | --- | --- | --- | --- | --- |
| Structural homology baseline | -0.241 | -0.312 | -0.407 | -0.427 | -0.358 |
| Masif-site | NA | NA | NA | NA | -0.571 |
| Handcrafted features baseline | -0.360 | -0.350 | -0.371 | -0.377 | -0.364 |
| ScanNet | -0.293 | -0.284 | -0.284 | -0.315 | -0.294 |

**TABLE S3.** Performance evaluation for prediction of Protein-protein binding sites. Accuracy is shown for the four subsets of test and the entire test.

| Algorithm | Test (70%) | Test (Homology) | Test (Topology) | Test (None) | Test (All) |
| --- | --- | --- | --- | --- | --- |
| Structural homology baseline | 0.916 | 0.886 | 0.844 | 0.844 | 0.868 |
| Masif-site | NA | NA | NA | NA | 0.712 |
| Handcrafted features baseline | 0.843 | 0.852 | 0.841 | 0.841 | 0.845 |
| ScanNet | 0.876 | 0.882 | 0.879 | 0.868 | 0.877 |

**TABLE S4.** Predictive performance for B-cell conformational epitopes. Area under Precision Recall Curve (AUCPR) and Positive Predicted Value at L/10 are shown. Evaluation by 5-fold cross-validation on the SabDab database.

| Algorithm | AUCPR | PPV L/10 |
| --- | --- | --- |
| Structural homology baseline | 0.147 | 0.216 |
| Handcrafted features baseline | 0.139 | 0.191 |
| Discotope [51] | 0.114 | 0.191 |
| ScanNet (transfer learning) | **0.178** | 0.273 |
| ScanNet (no finetuning) | 0.106 | 0.211 |
| ScanNet (scratch) | 0.150 | 0.221 |
| ScanNet (transfer learning, no evolutionary information) | 0.173 | **0.275** |
| Null prediction | 0.089 | 0.089 |
| Solvent Accessibility baseline | 0.121 | 0.178 |

**TABLE S5.**
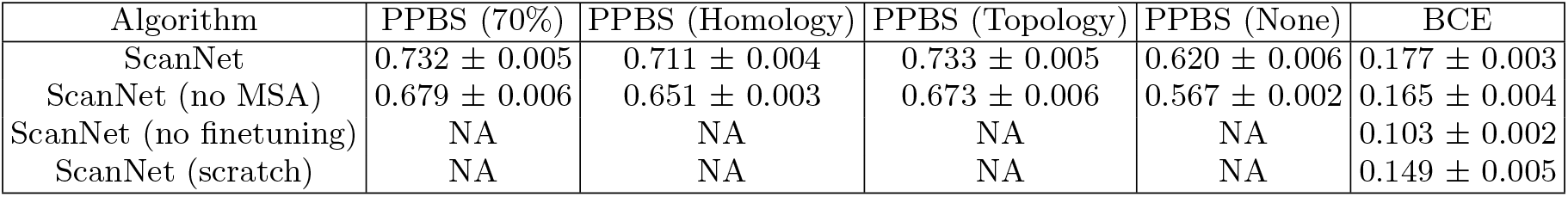
Performance evaluation for prediction of Protein-protein binding sites and Protein-antibody binding sites: impact of random seed. Each ScanNet network was retrained 10 times using different random seeds. The average and standard deviation of the Area Under the Precision-Recall Curve (AUCPR) are shown for the four test sets. The standard deviation is about 10-20 times smaller than the difference between ScanNet and the baselines.

**TABLE S6.** List of SARS-CoV2 representative antibodies shown in Fig. 5. Epitopes are classified following [52]

| Epitope | Antibody | PDB ID | Reference |
| --- | --- | --- | --- |
| RBD-A | CC12.1 | 6xc2:HL | [95] |
| RBD-B | COVA2-39 | 7jmp:HL | [96] |
| RBD-C | CV07-270 | 6xkp:HL | [97] |
| RBD-D | REGN10987 | 6xdg:AC | [98] |
| S309 | CV38-142 | 7lm8:MN | [99] |
| CR3022 (cryptic) | COVA1-16 | 7jmw:HL | [100] |
| NTD | 2-51 | 7l2c:CD | [101] |
| Stem | B6 | 7m53:HL | [54] |

**FIG. S1.**
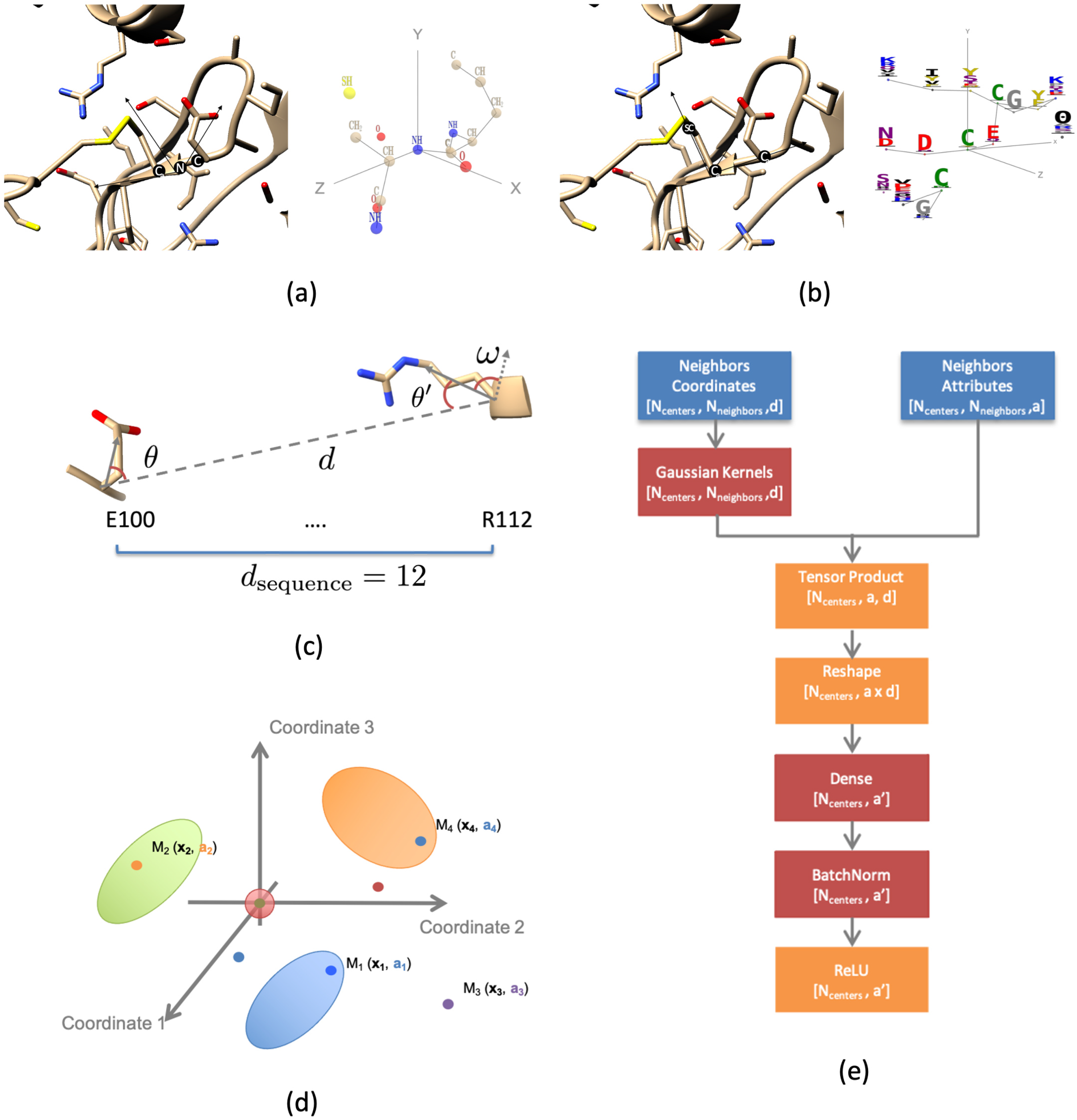
Overview of the frame computation, neighborhood computation and neighborhood embedding modules. (a) Construction of an atomic neighborhood from structure. For each atom, the *K* =16 closest atoms (including itself) are identified. Next, a frame is constructed from its position and the directions of its covalent bonds. The neighboring atoms are characterized by their coordinates in the local frame and group type (12 subclasses: C,CH,CH2,CH3,CII (aromatic ring), O, OH, N, NH, NH2, S,SH. (b) Construction of an amino acid neighborhood from structure. For each amino acid, the *K* =16 closest amino acid (including itself) are identified. Next, a frame is constructed from its *C_α_* atom, side-chain center of mass and the previous *C_α_* atom along the backbone.The neighboring amino acid are represented by their coordinates in the local frame and their attributes learnt from the position weight matrix and pooled atomic filters. (c) Local coordinate system used for the neighborhood attention module (d) Principle of neighborhood embedding module: a generic neighborhood consists of a set of *K* points *M_k_* characterized by their local coordinates x_k_ and attributes a_k_; (e) Implementation of the neighborhood embedding module.

**FIG. S2.**
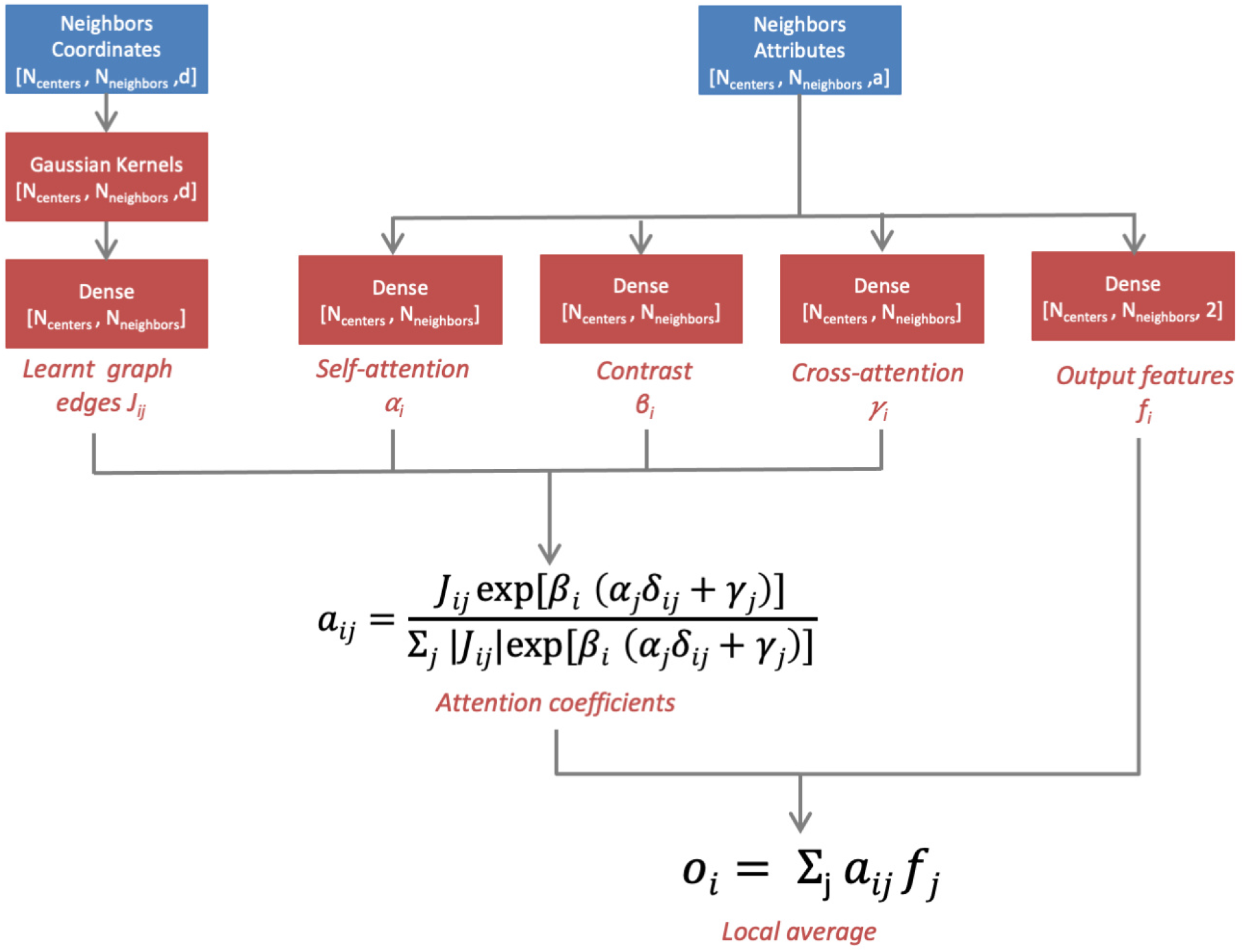
Overview of the neighborhood attention module. The neighborhood attention module is the final module of ScanNet; its purpose is to locally average predictions to produce spatially consistent predictions. An attention mechanism is included to account for driver/passenger binding sites. *δ_i,j_* = 1 if *i* = *j*; 0 otherwise is the Kronecker symbol.

**FIG. S3.**
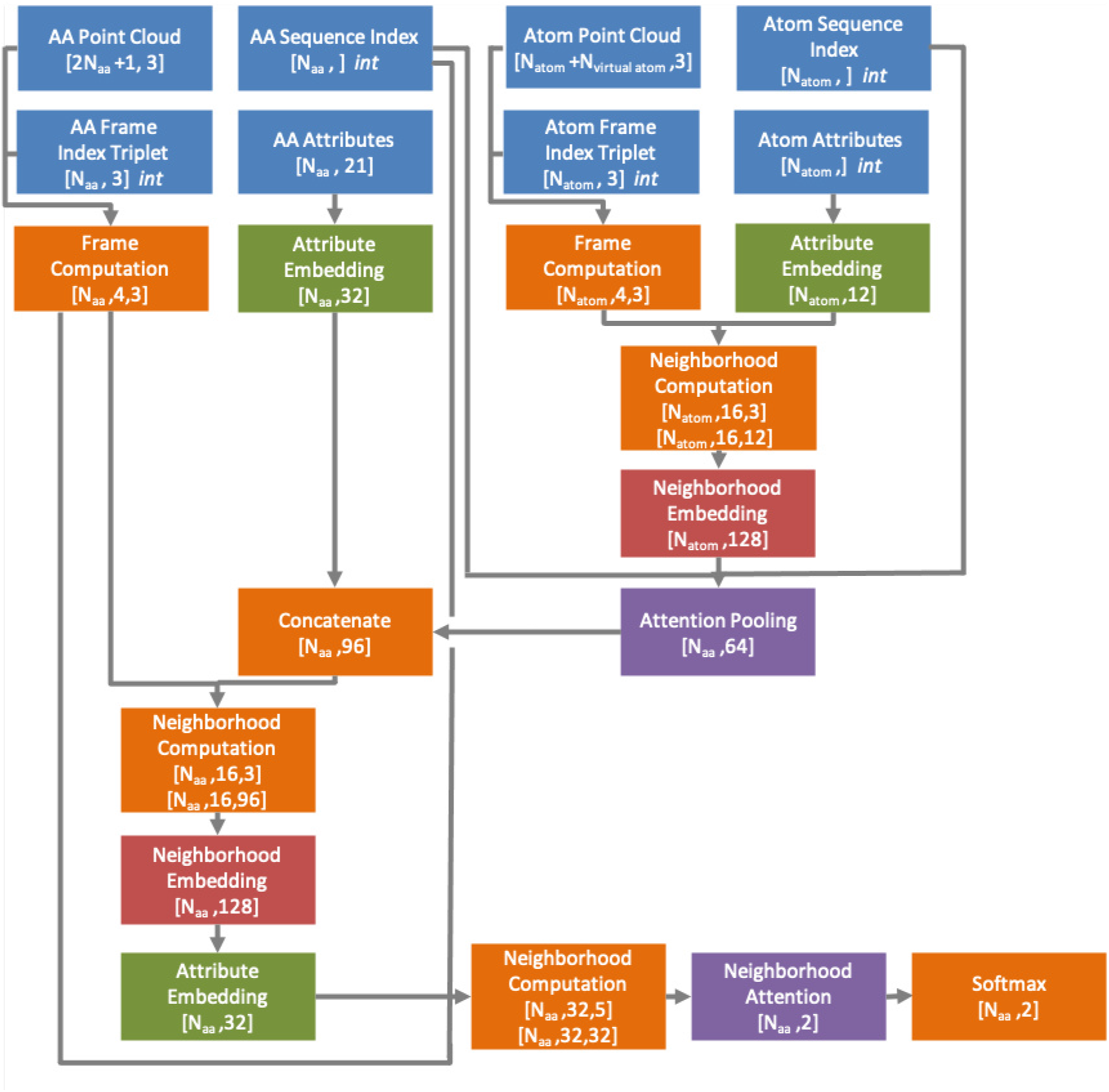
Complete architecture of ScanNet. Orange modules are not trainable.

**FIG. S4.**
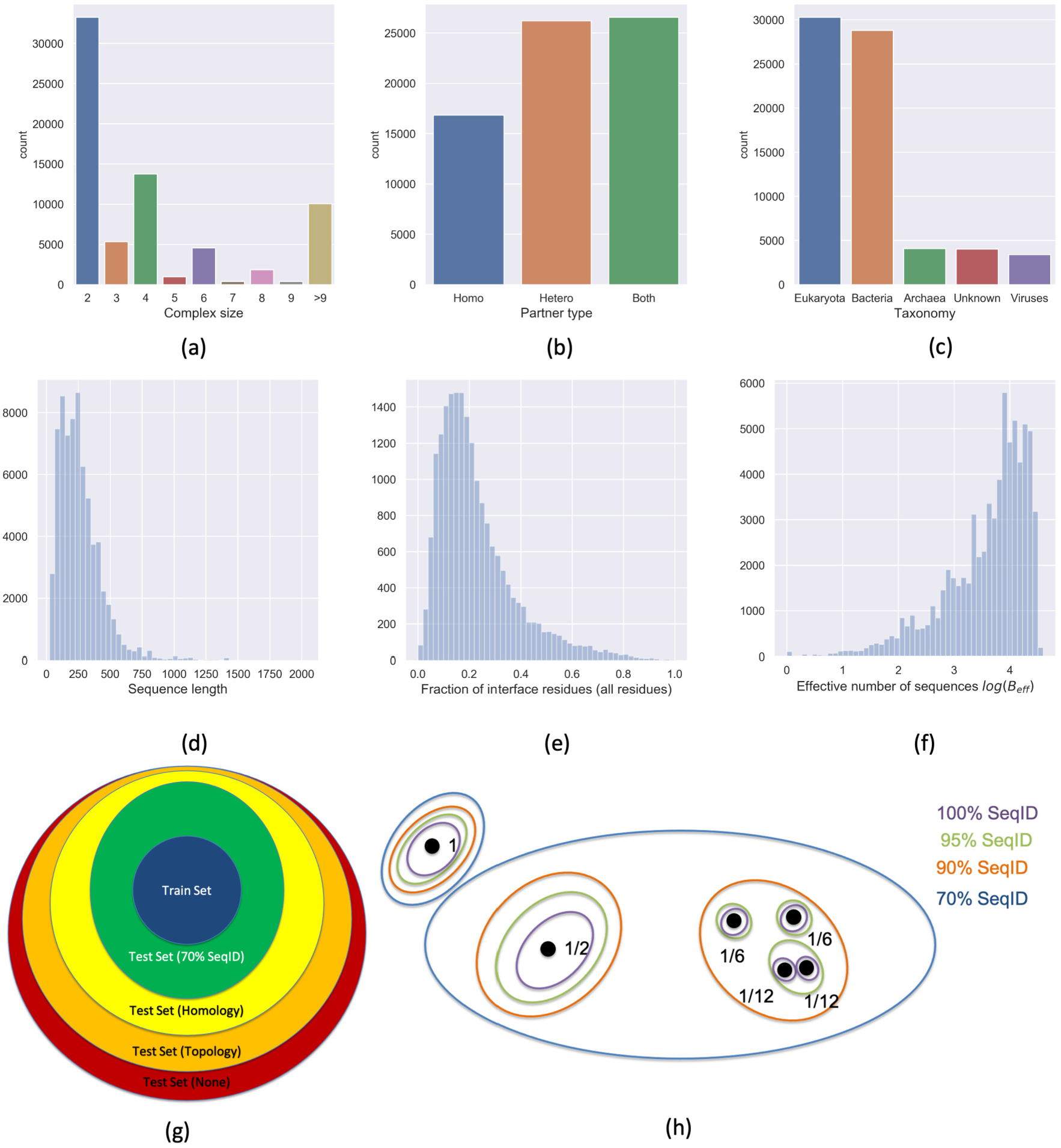
Overview of the Protein-protein binding sites database. (a-) Distribution of (a) complex sizes (b) complex types (c) source organism taxonomy (d) protein length (e) fraction of interface residues (f) effective number of sequences in corresponding the multiple sequence alignment. (g) Data partition. Proteins of the validation/test set are subdivided into four non-overlapping groups, depending on the degree of similarity with the closest protein found in the train set: (i) ≥ 70% Sequence identity (ii) Same CATH superfamily (iii) Same fold topology CAT. (iv) None of the above. Generalization is increasingly difficult. (h) Illustration of the hierarchical sample reweighting used to counterbalance heterogeneity in the sampling of the protein space at multiple levels. Sequences are first clustered at four sequence identity thresholds (100%, 95%,90%,70%). Each cluster at 70% sequence identity (blue ellipses) contributes an identical total weight of 1 irrespective of its size. Within each 70% cluster, each of the 90% clusters (orange ellipses) contributes an identical total weight 1/Ncluster90, etc. The weight of a sample is: Num(sequences in cluster 100) × Num(cluster 100 in cluster 95) × Num(cluster 95 in cluster 90) × Num(cluster 90 in cluster 70). The weight of each cluster 70% is invariant upon subsampling of the dataset.

**FIG. S5.**
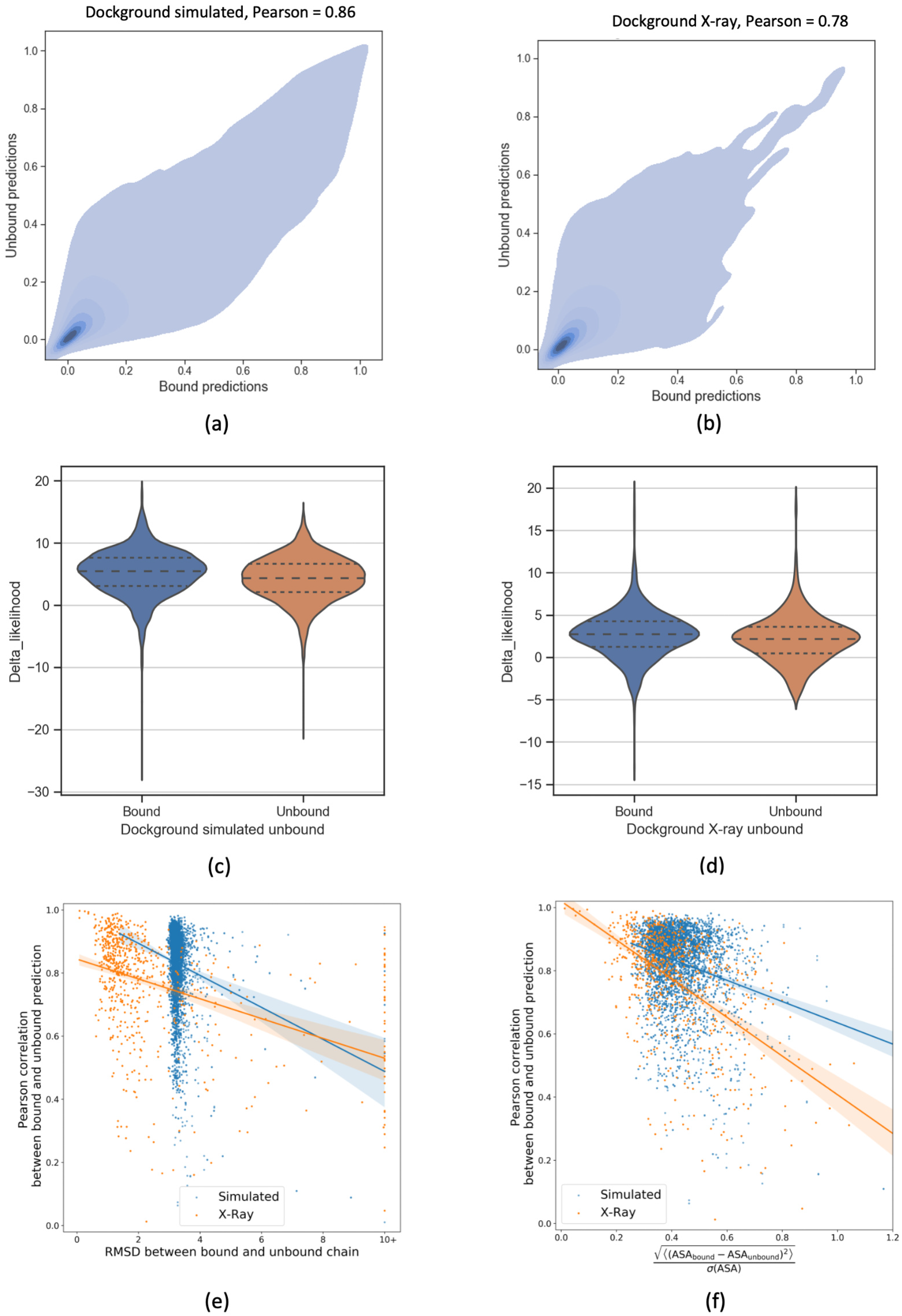
Comparison between predictions performed on bound and unbound structures. Two data sets of (bound,unbound) pairs of protein structures are considered: the Dockground simulated data set and Dockground X-ray data set. Panels (a),(b) display 2D-density plots of the distribution of ScanNet predictions on bound and unbound structures for each data set. Panels (c),(d) show for each data set the distributions of protein-wise prediction performance, measured as the difference 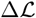 between the likelihood of ScanNet prediction and null prediction (uniform probability *p* ~ 0.2), divided by the standard deviation of the null model likelihood 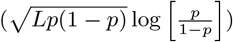. Higher is better. By construction, for a null predictor, 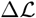 has zero variance across the data set whereas other metrics such as likelihood or accuracy have substantial variance owing to the variability of fraction interface residues across proteins, see Sup. Figure S4 (g); using 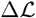 therefore facilitates detection of trends. A statistically significant but overall limited drop in performance is observed from bound to unbound. (e),(f) Impact of the degree of global (e) and local (f) conformational changes on the consistency between bound and unbound prediction. The correlation between bound and unbound predictions is represented against the RMSD between bound and unbound atomic coordinates (e), and between bound and unbound relative solvent accessibility values (f)

**FIG. S6.**
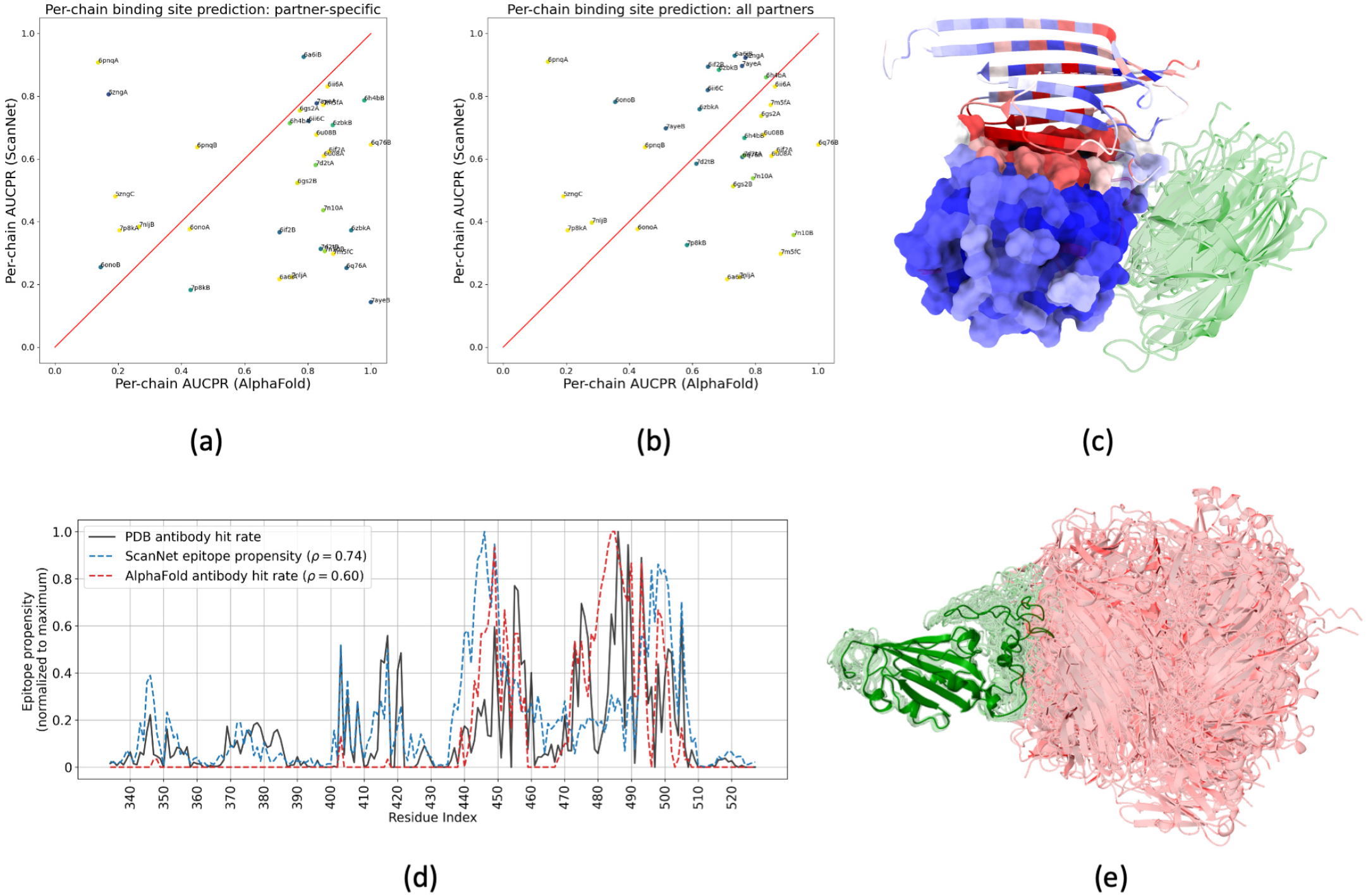
Comparison of AlphaFold-Multimer and ScanNet for binding site prediction. (a) Scatter plot depiction of the per-chain AUCPR metric for prediction of *partner-specific* binding sites on the Benchmark2 data set. (b) Same for *partner-agnostic* binding sites, determined by taking the union of all binding sites found in related complexes. In both panels, each chain is colored by the ratio of the number of partner-specific binding sites divided by the number of partner-agnostic binding sites (from 0=Blue to 1=Yellow). By definition, the ratio equals one for proteins with only one known partner and is low for multivalent proteins. (c) Depiction of complex 6pnq (chains A and B respectively in surface and cartoon representations), colored by ScanNet binding site probability (from blue=low to red=high). The five models produced by AF2 for chain B are superimposed in green. ScanNet correctly predicts the binding sites of both chains, but not AF2. (d) B-cell epitope propensity profile for the Receptor Binding Domain of the Spike Protein, as estimated from i) available structures in the Protein Data Bank ii) ScanNet B-cell epitope network and iii) AF2-based docking with representative antibodies. AF2 fails to identify all the main epitopes. (e) Depiction of the 30 RBD-antibody complex models predicted by AF2, featuring only a single binding mode.

**FIG. S7.**
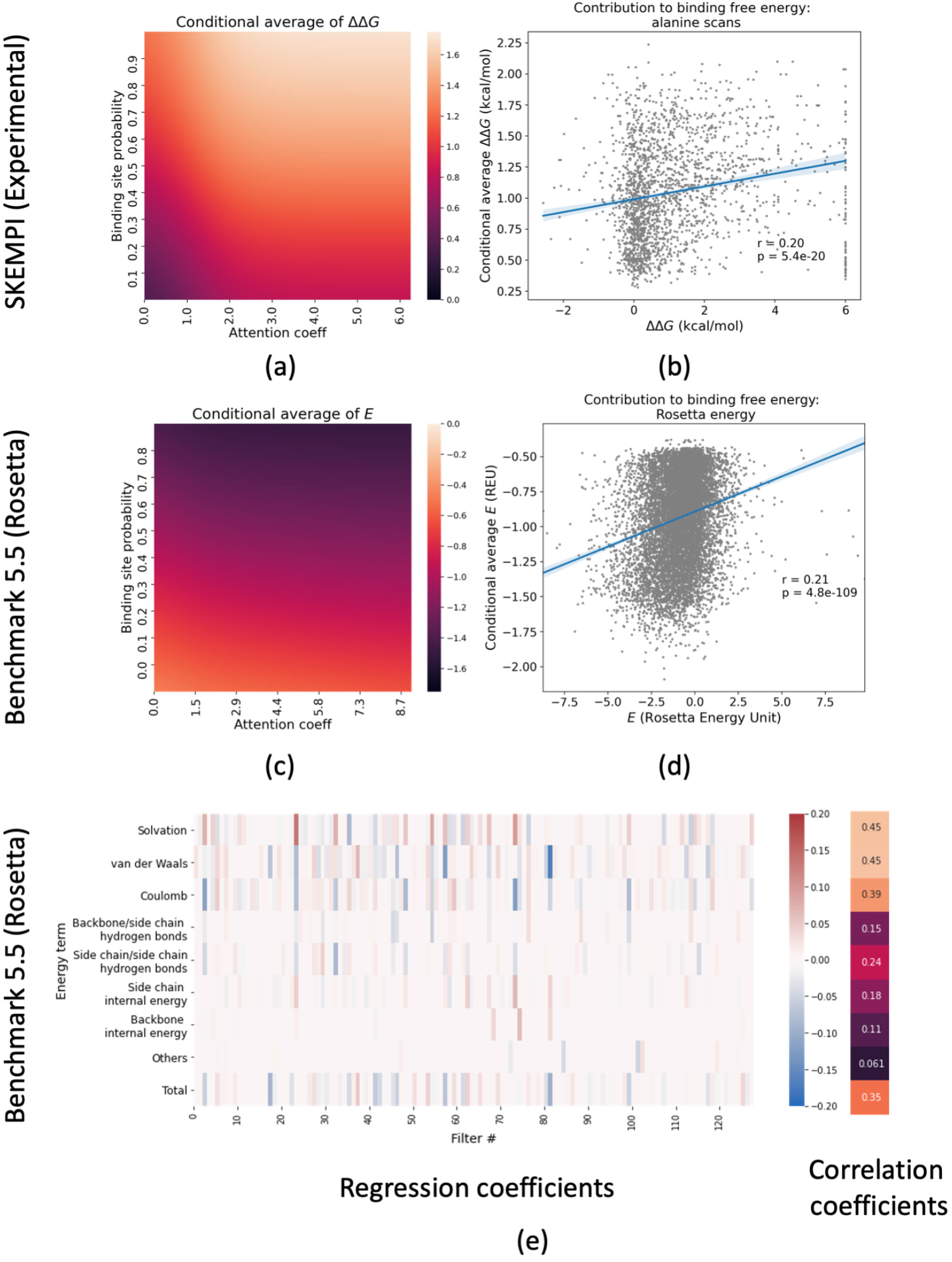
Link between ScanNet predictions and residue contribution to the binding free energy. (a) Parametric fit of the conditional average of experimentally determined changes of binding affinity upon mutation to alanine ΔΔ*G* (obtained from the SKEMPI v2 database), as function of the predicted binding site propensity p and aggregated attention coefficient *a* of the residue. (b) Scatter plot of the predicted ΔΔ*G* given *p*, *a* to the experimental one (cross-validation predictions) (c) Parametric fit of the conditional average of Rosetta binding energy *E* (computed from the Benchmark v5.5 database), as function of the predicted binding site propensity *p* and aggregated attention coefficient *a* of the residue. (d) Scatter plot of the predicted *E* given *p, a* to the experimental one (cross-validation predictions). (e) Sparse regression and correlation coefficients of Rosetta energy terms from ScanNet amino acid filter activities. Displayed values are 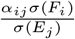, where *Y_j_* is the j’th energy term, and *F_i_* is the i’th filter activity and *α_ij_* is the regression coefficient determined by LASSO regression

**FIG. S8.**
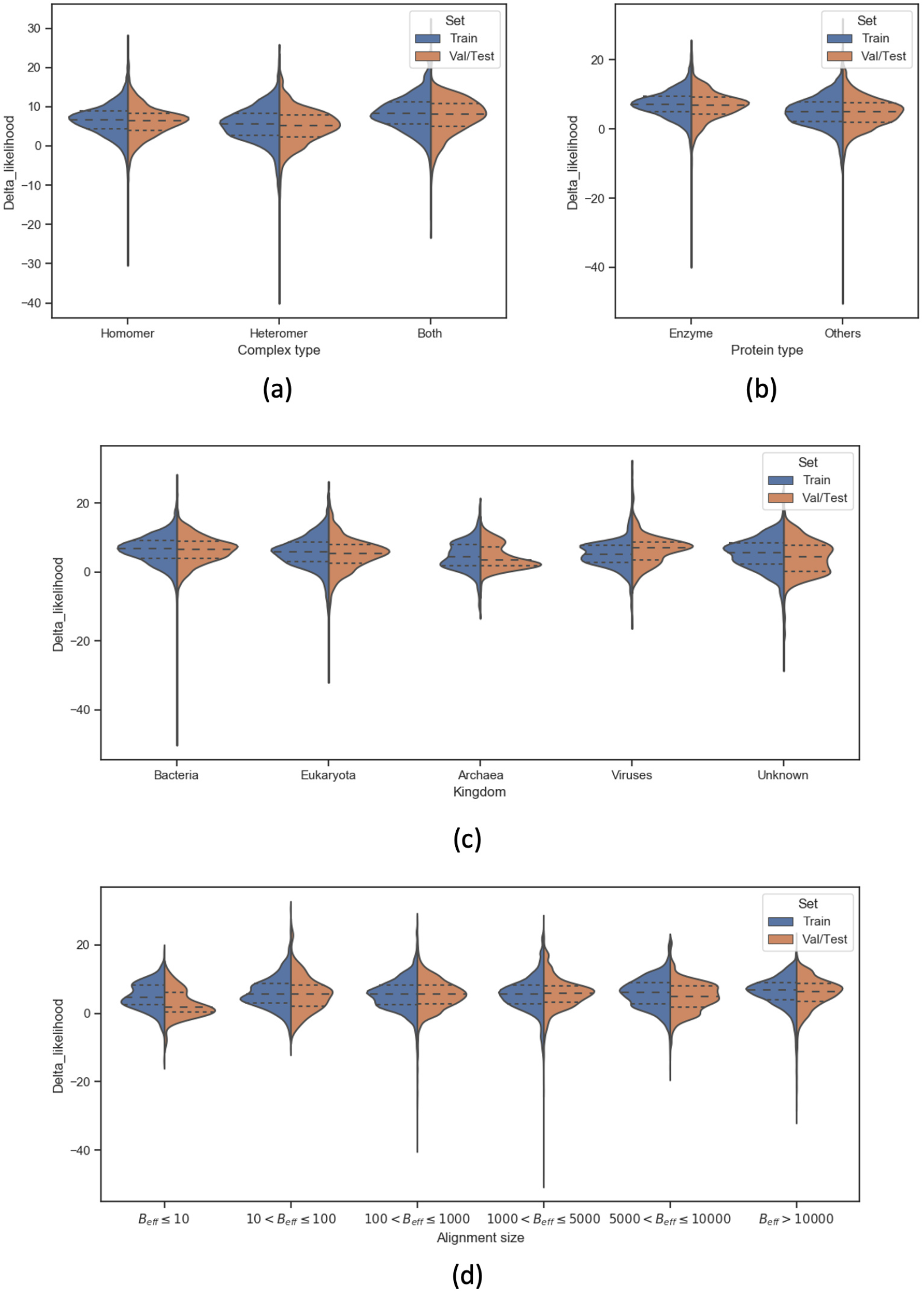
ScanNet performance by sample type. The metric shown is the difference between the likelihood of the ScanNet and the likelihood of the null predictor (constant probability ~ 0.2); higher is better. Prediction performance is shown against complex type, protein type, source organism and effective alignment size.

**FIG. S9.**
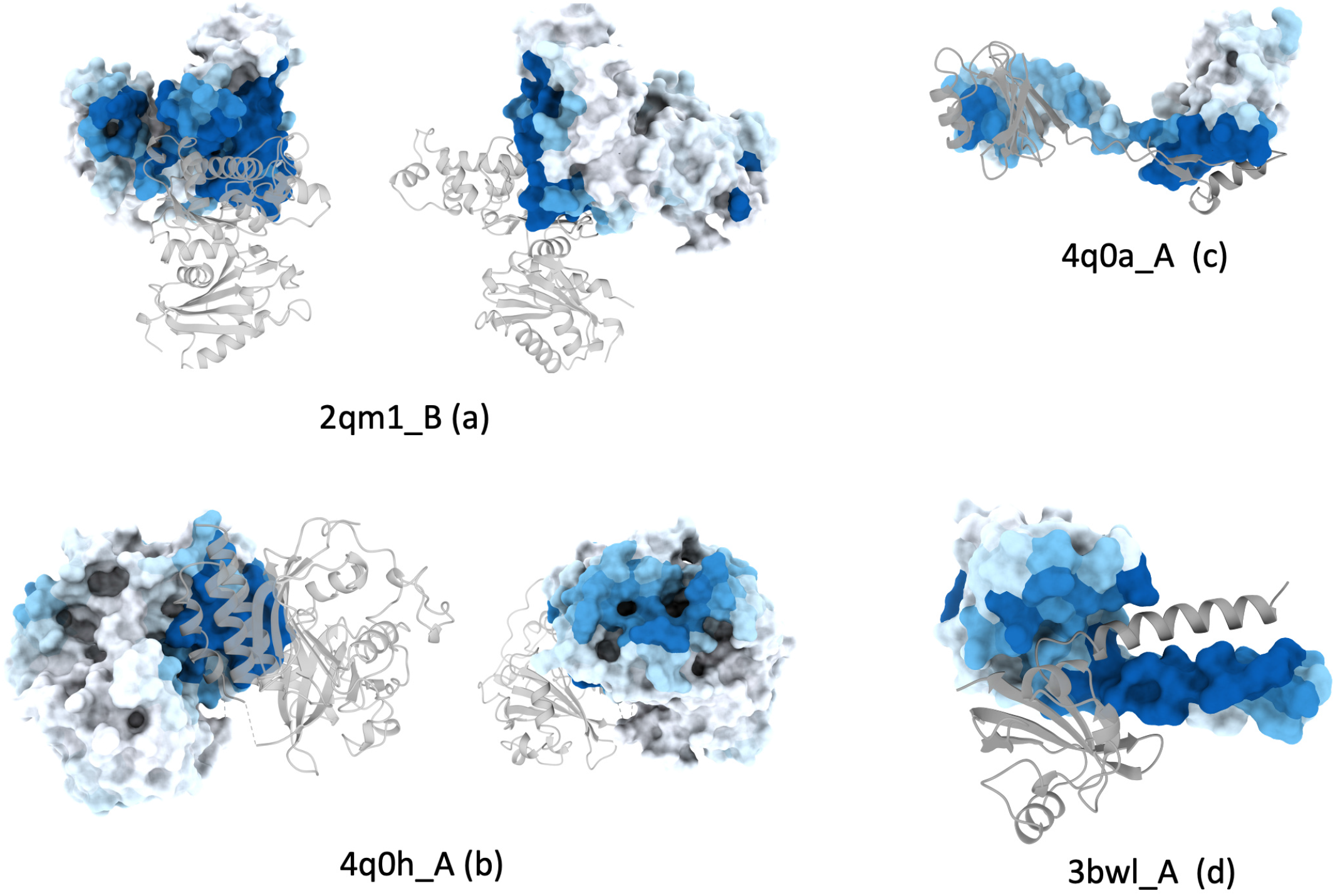
Visualization of ScanNet PPBS prediction for homodimers. Typical test sets examples are shown, with delta likelihoood values close to the median performance of the test set.

**FIG. S10.**
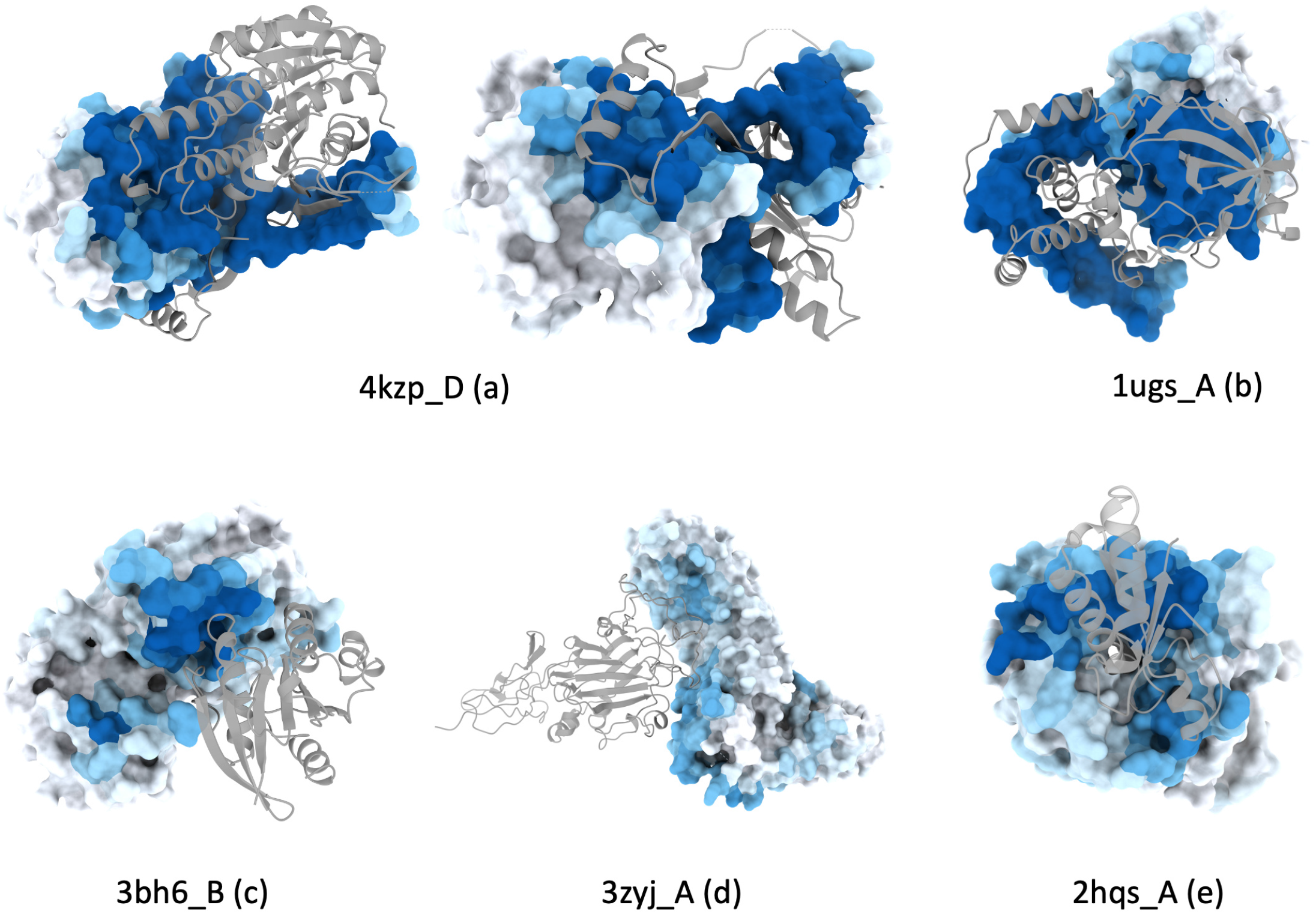
Visualization of ScanNet PPBS prediction for heterodimers. Typical test sets examples are shown, with delta likelihoood values close to the median performance of the test set.

**FIG. S11.**
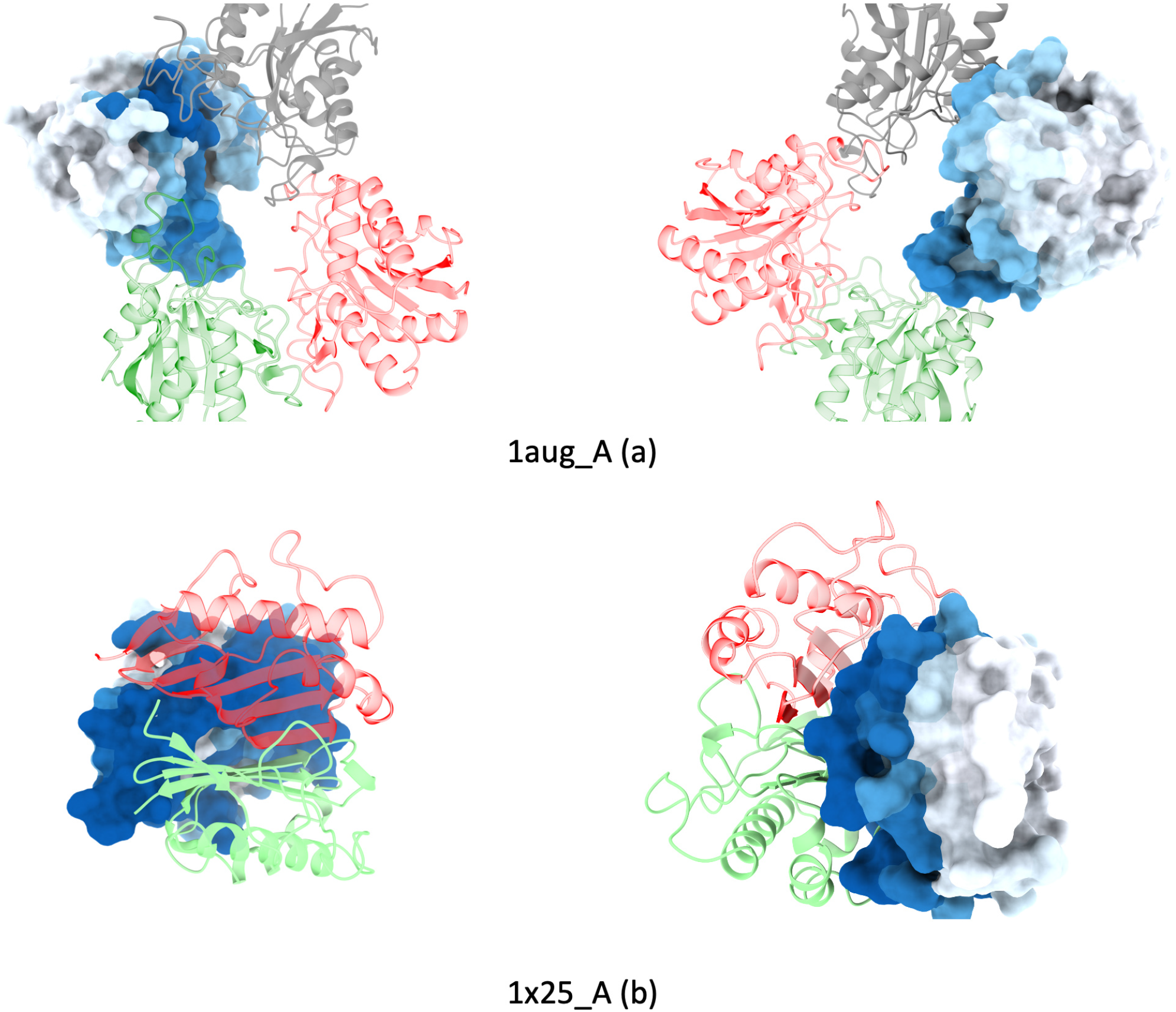
Visualization of ScanNet PPBS prediction for homomultimers. Typical test sets examples are shown, with delta likelihoood values close to the median performance of the test set.

**FIG. S12.**
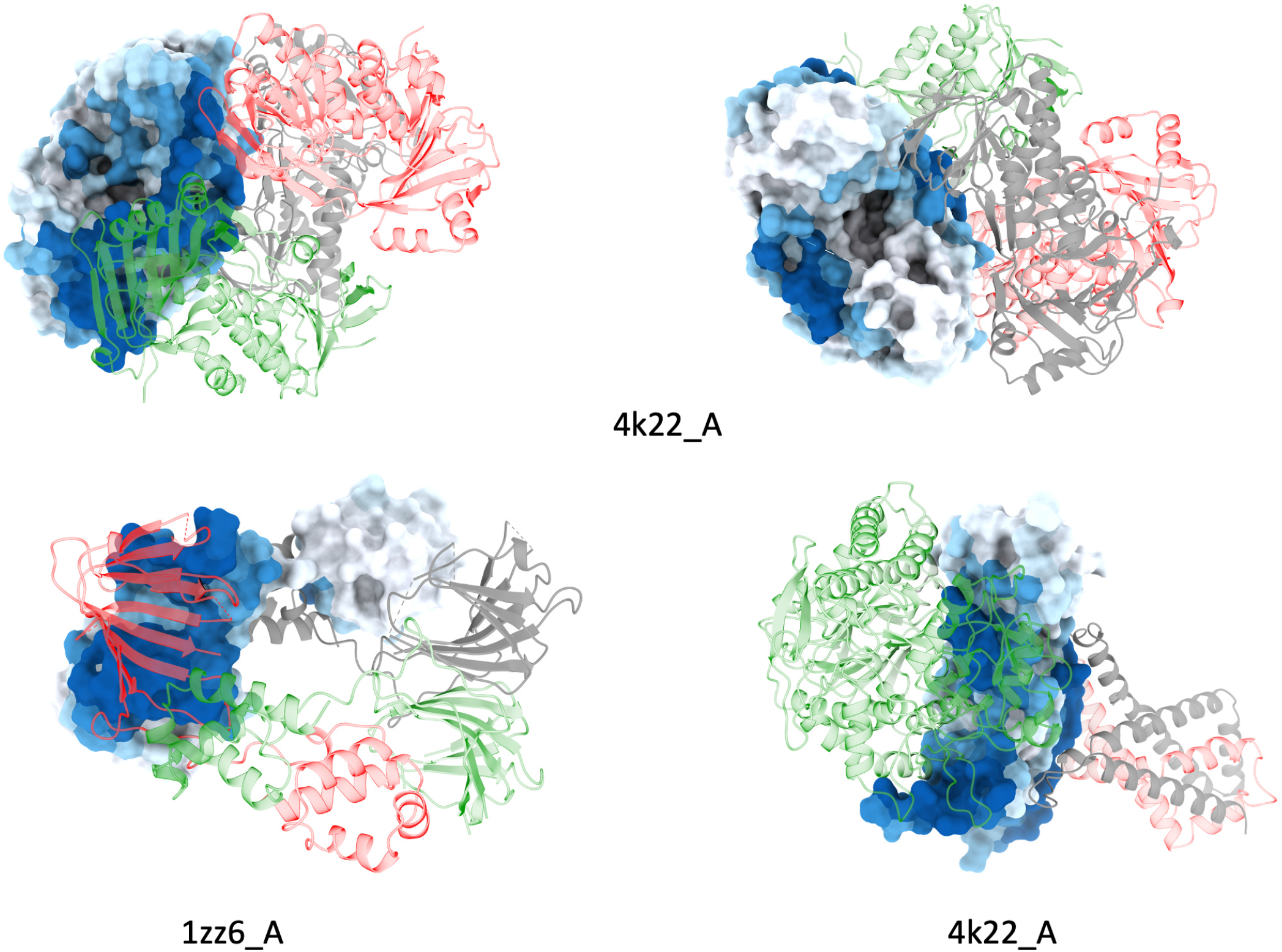
Visualization of ScanNet PPBS prediction for heteromultimers. Typical test sets examples are shown, with delta likelihoood values close to the median performance of the test set.

**FIG. S13.**
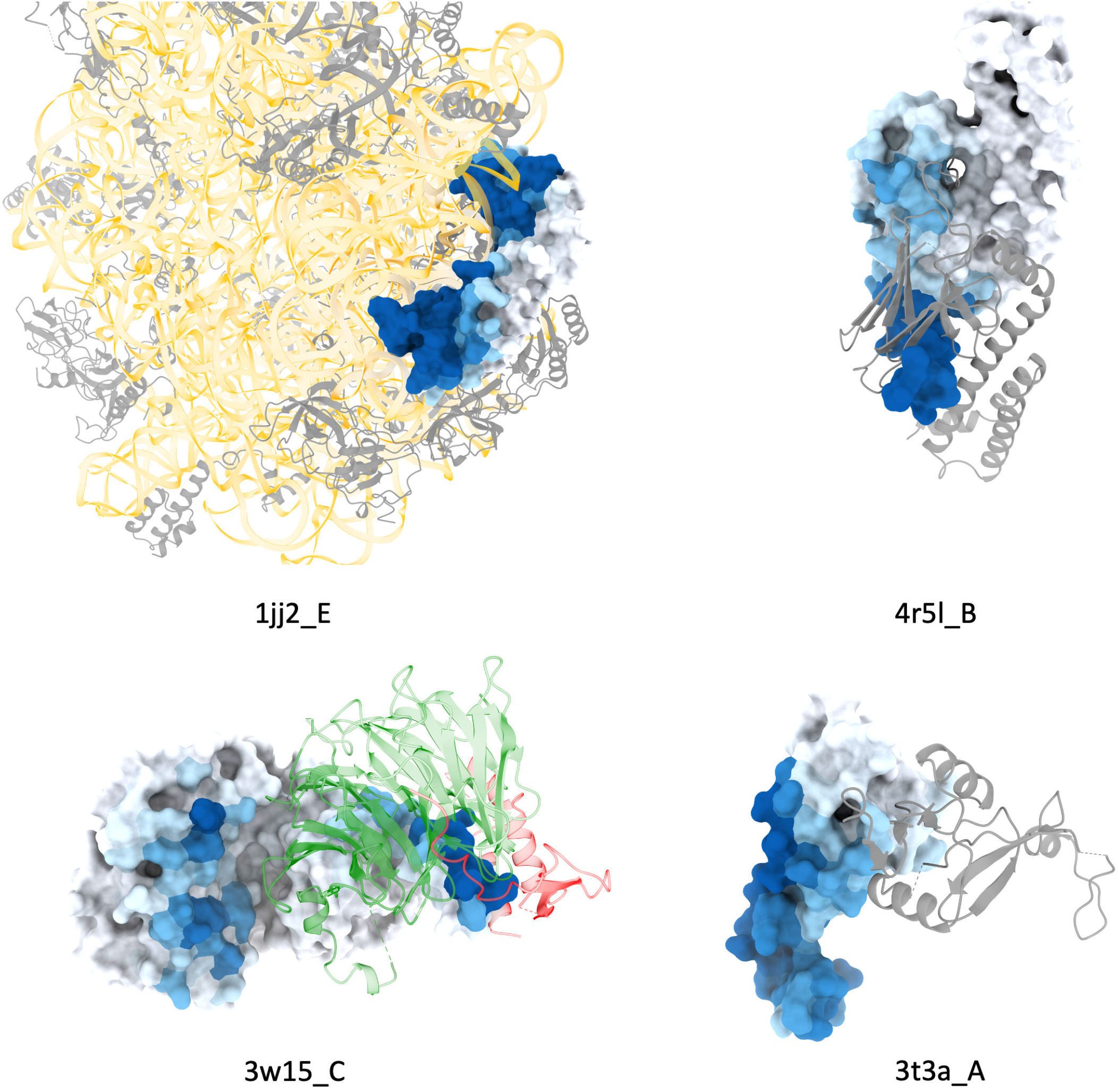
Example of incorrect PPBS predictions. Examples shown have delta likelihoood values in the lower decile of the test set. In the first instance, the network confuses an RNA binding site with a protein-protein binding site. In the second instance, the network fails to identify the cavity as a binding site for the histidine tag of the protein partner. The last two show misplaced binding sites

**FIG. S14.**
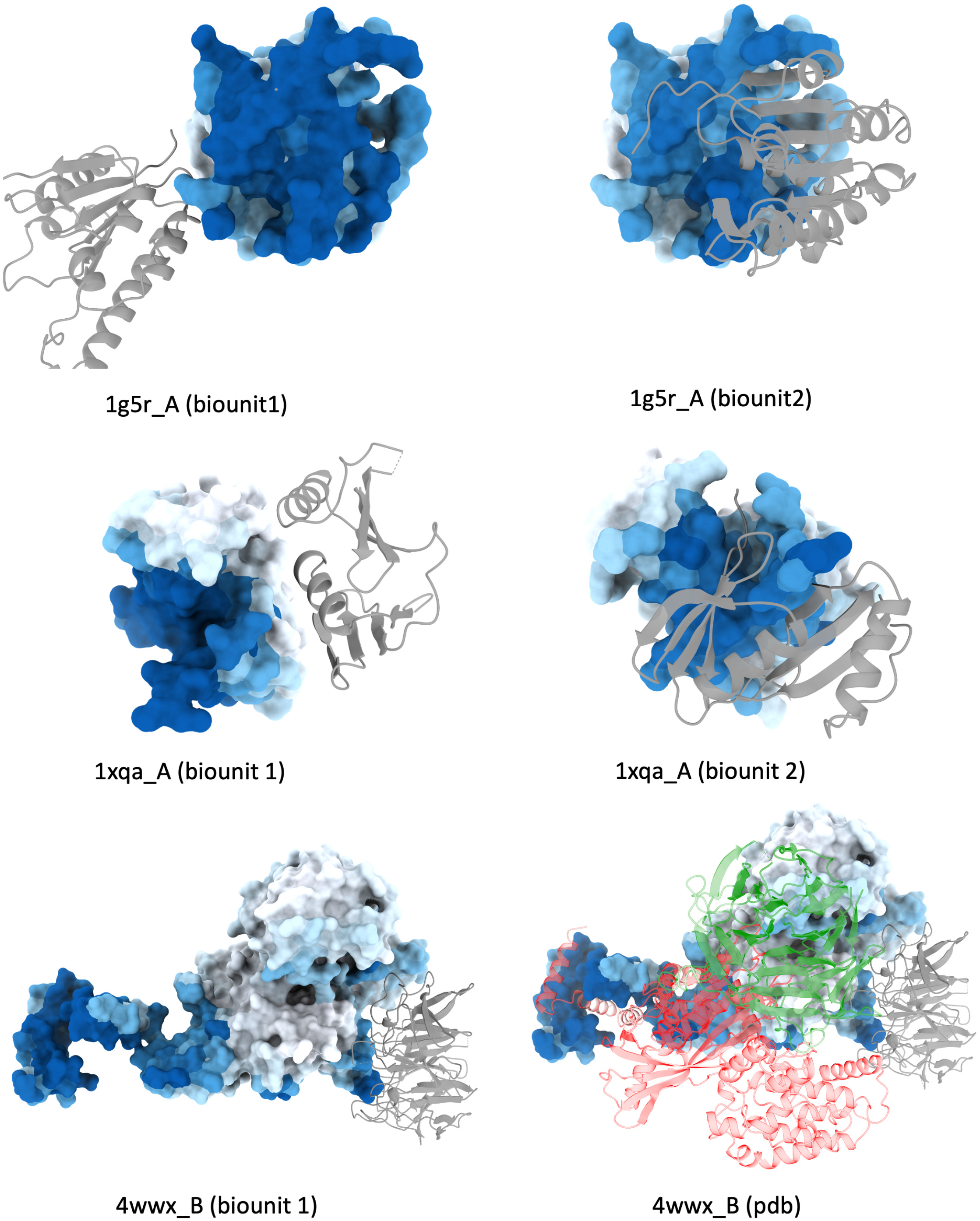
Example of correct PPBS predictions misclassified due to incorrect labels. Each example belongs to the train set, yet the network fails to learn the apparent native binding sites. The predicted labels match the binding sites of another biological assembly file.

**FIG. S15.**
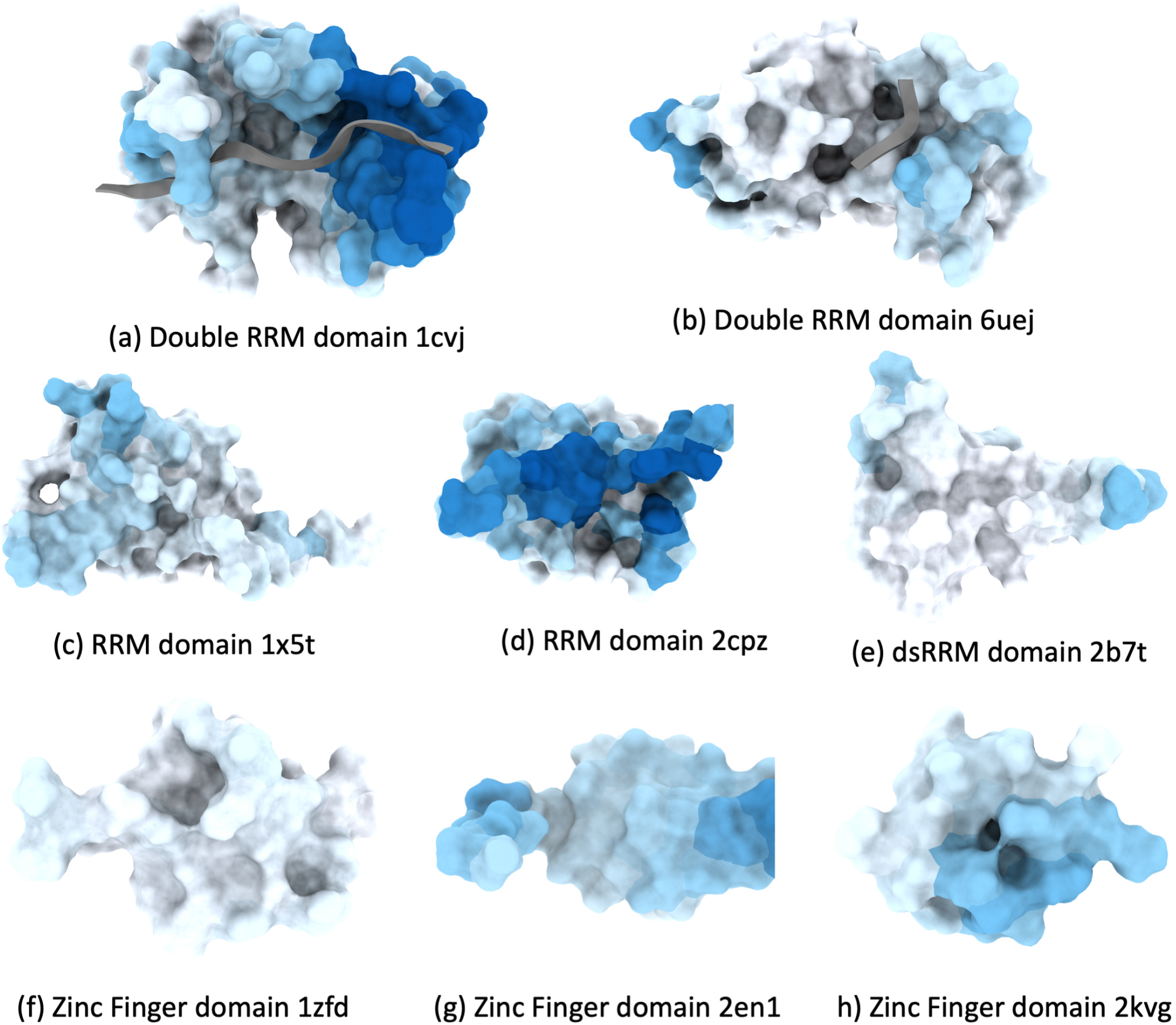
Evaluation of PPBS prediction on representative RNA-binding proteins. Each protein contains one or two RNA-binding domains. Two out of ten domains are misclassified as protein binding.

**FIG. S16.**
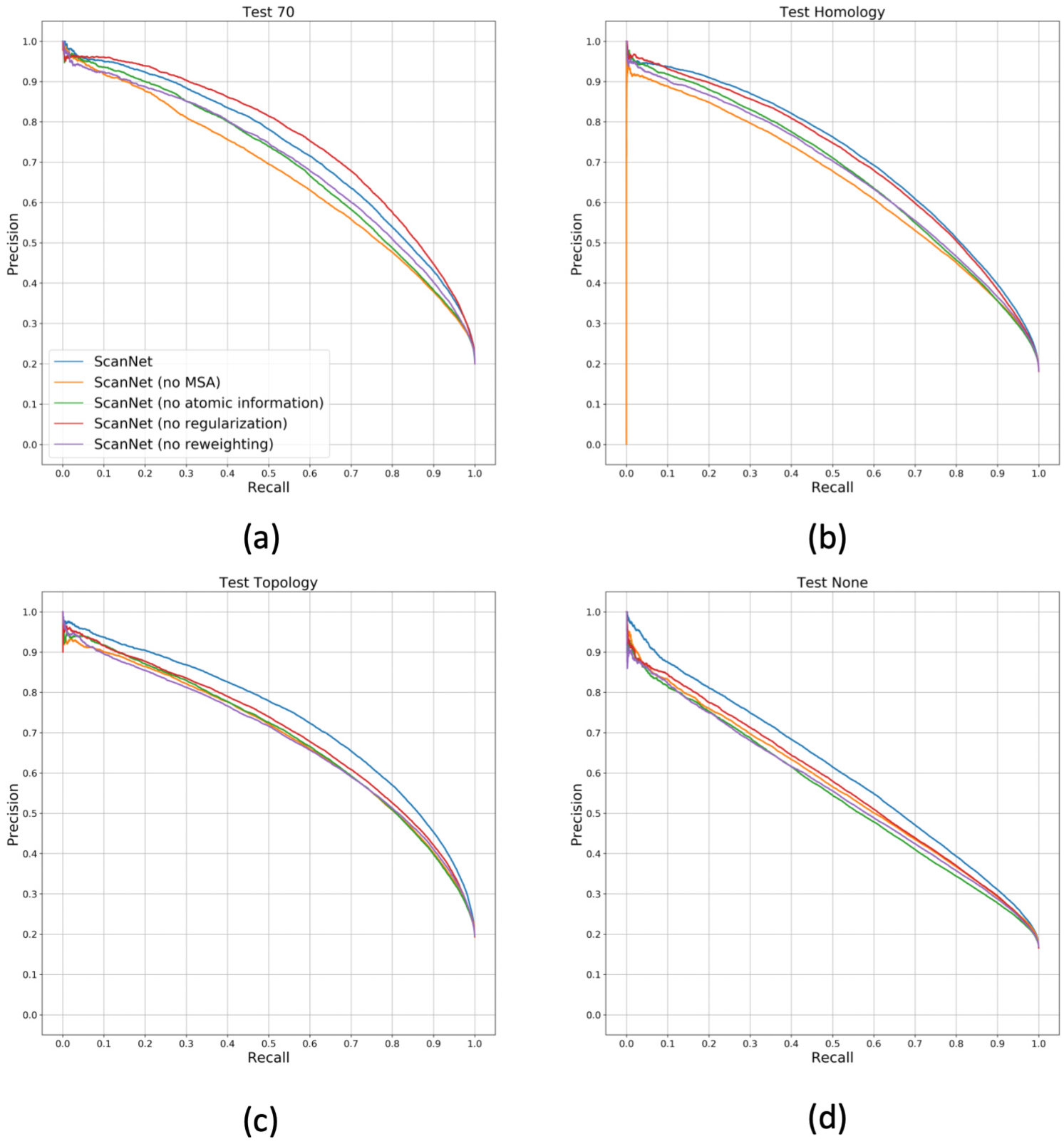
Performance of Protein-Protein Binding Sites (PPBS) prediction with ablated ScanNet, see description of ablations in main text. Precision-Recall curves on the train set and the four test subsets, see Fig. 2.

**FIG. S17.**
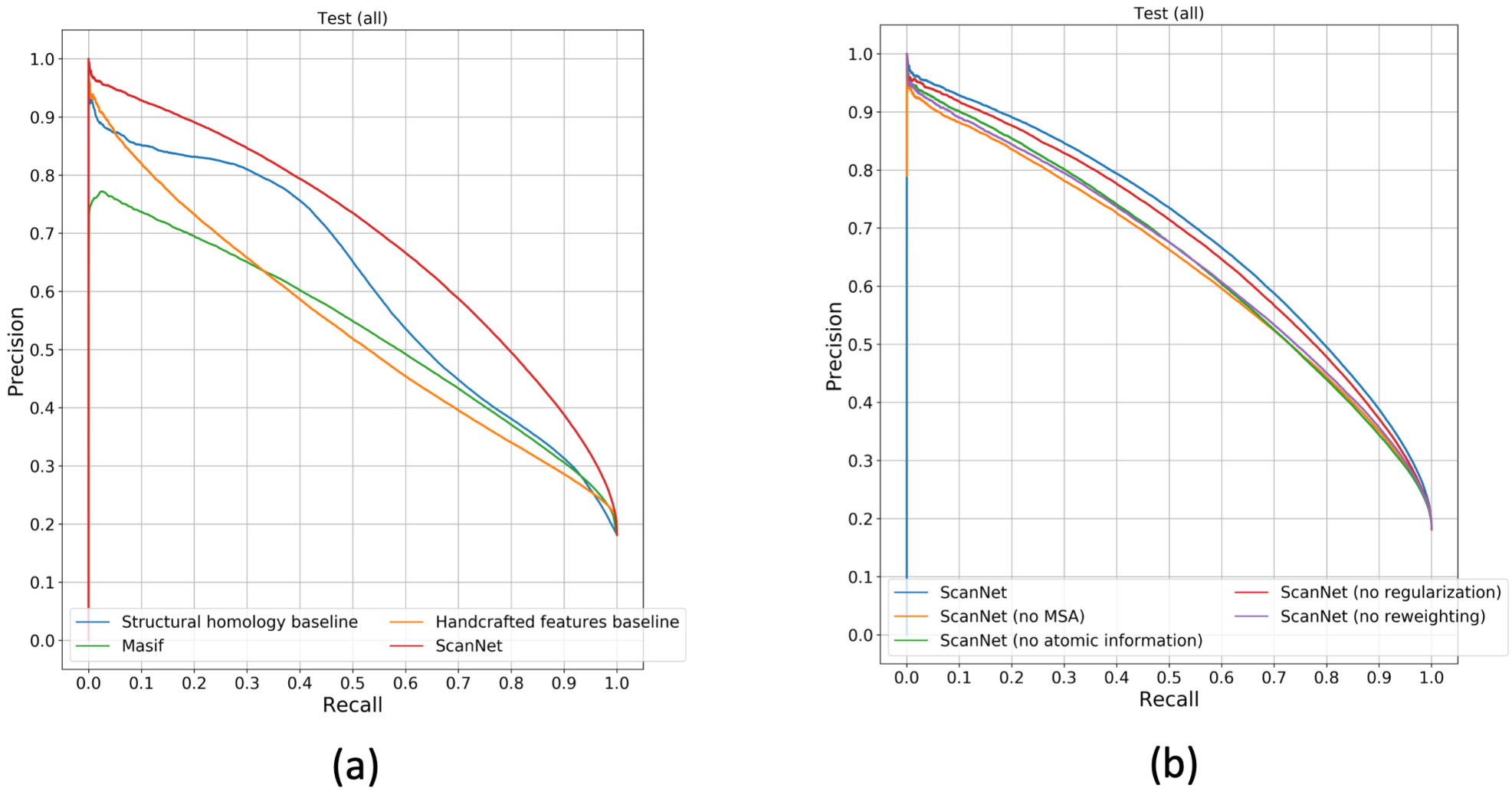
Performance of Protein-Protein Binding Sites (PPBS) prediction. Precision-Recall curves of PPBS prediction performance, across the entire test set.

**FIG. S18.**
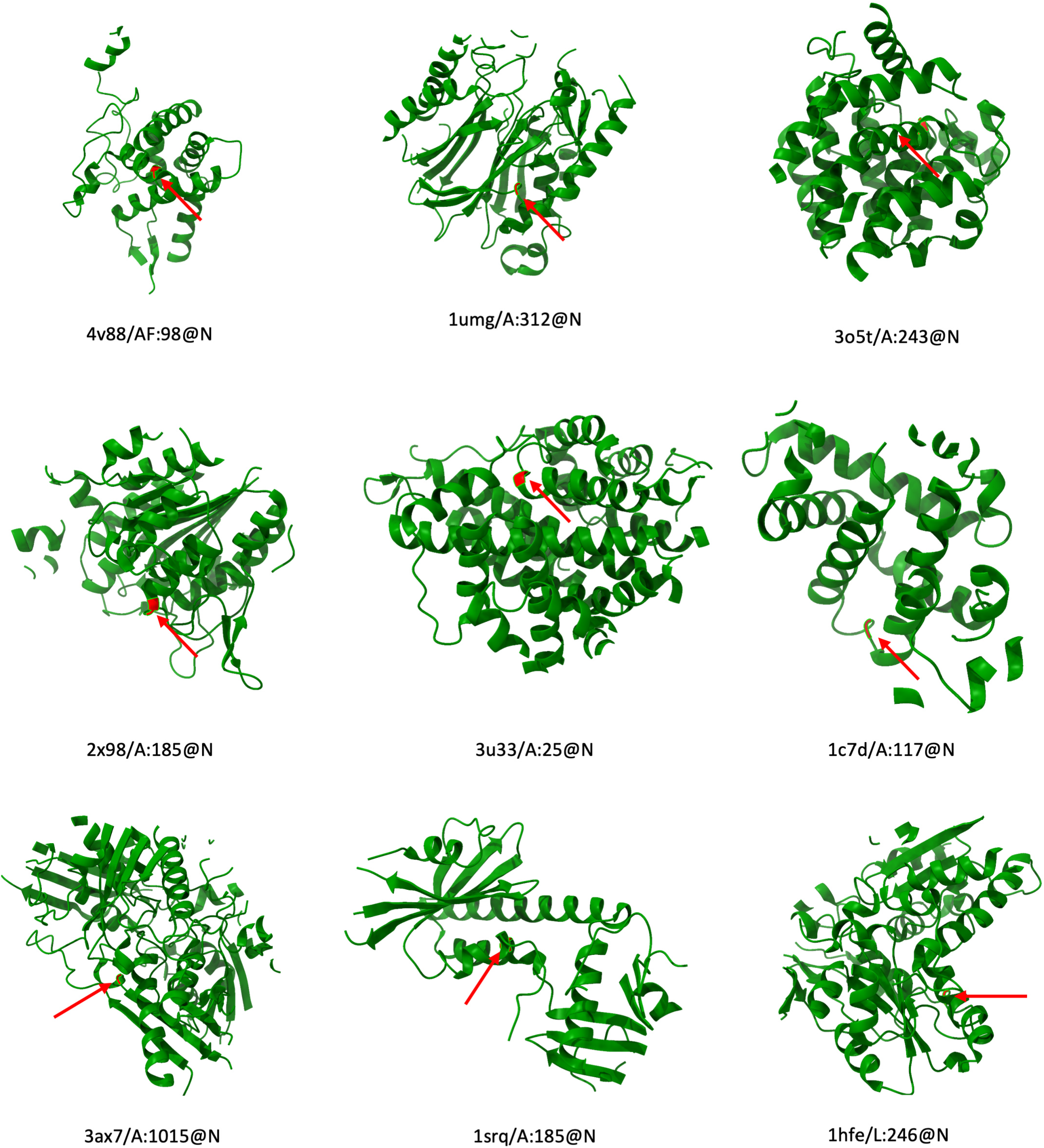
Visualization of the top nine activating neighborhoods of atomic filter (b) shown in Fig.3. The top-activating atom is the backbone nitrogen of the residue shown in red (zoom-in for clarity). Each of the top-activating nitrogens is located at a contact zone between two helical fragments.

**FIG. S19.**
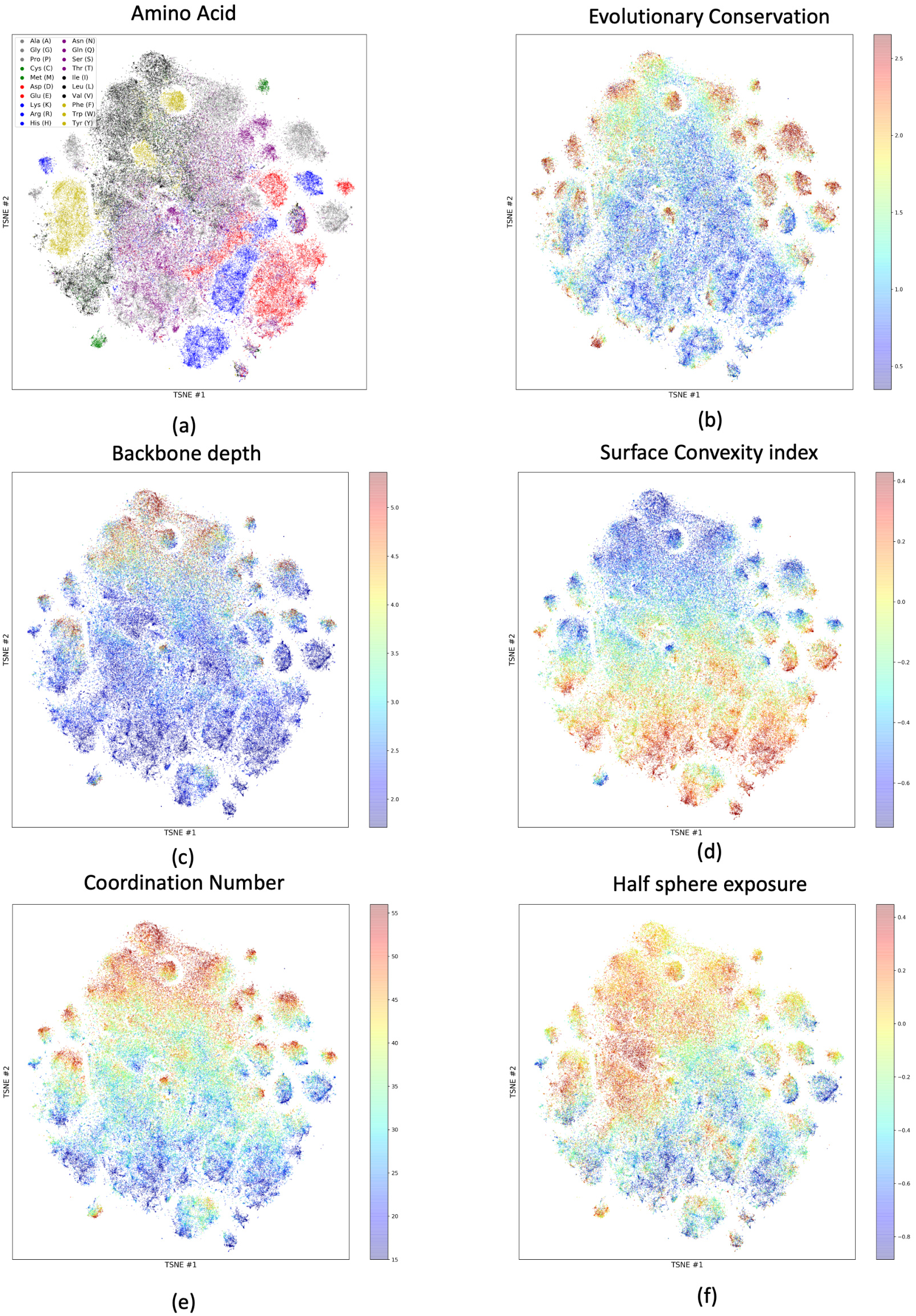
Two-dimensional projection of the learnt amino acid scale representation using T-SNE [60]. Each point corresponds to one amino acid of a representative set of proteins. Coloring based on (a) Amino acid type (b) evolutionary conservation (c) backbone depth (d) surface convexity index (e) coordination number (f) Half-sphere exposure

**FIG. S20.**
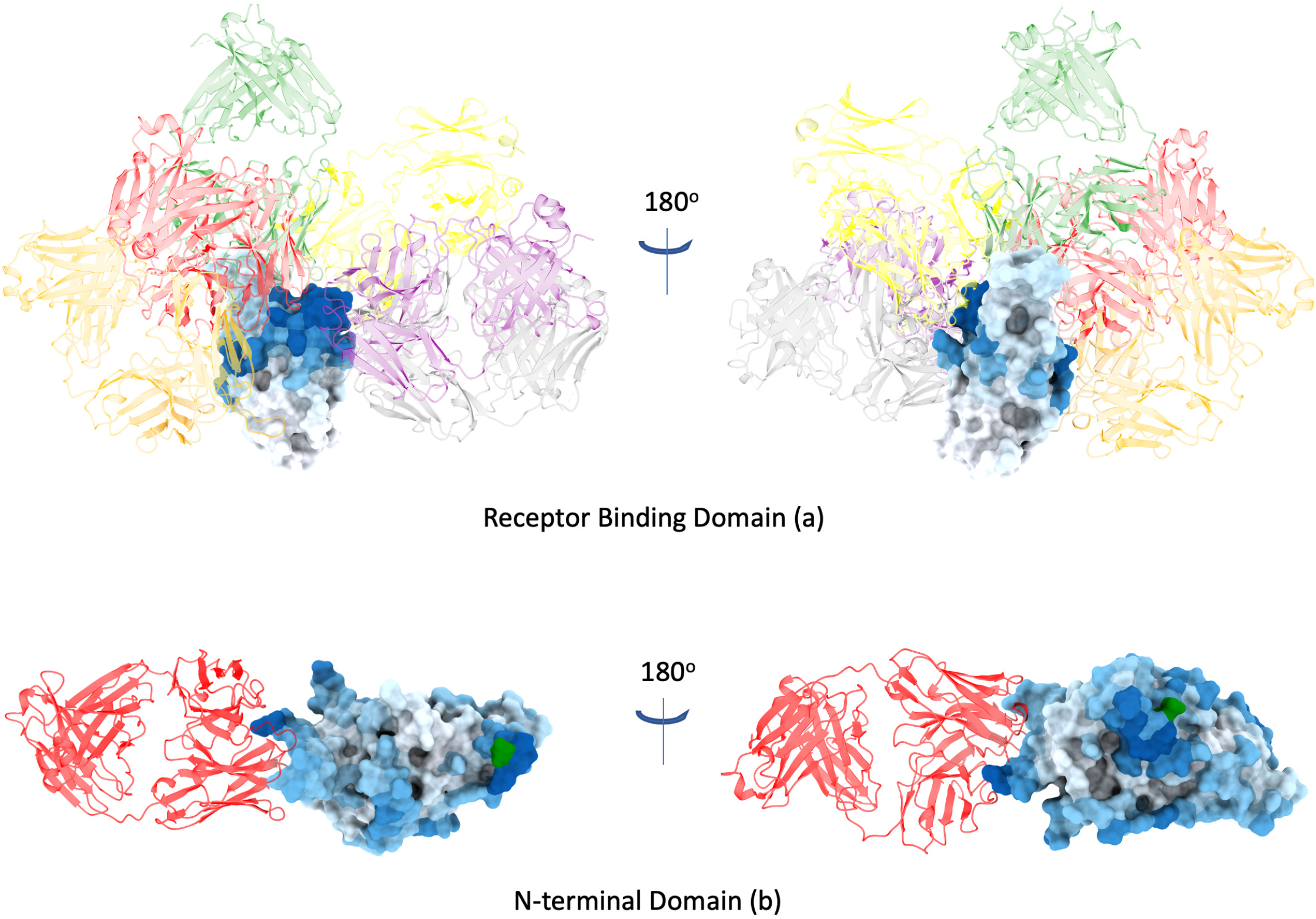
B-cell conformational epitope prediction on SARS-CoV2 Spike Protein. a) Receptor Binding Domain (PDB ID 6xkp [97]) N-terminal domain (PDB ID 7l2c [101]). Domains depicted as surfaces, Green residues indicate glycosylated asparagines.

**FIG. S21.**
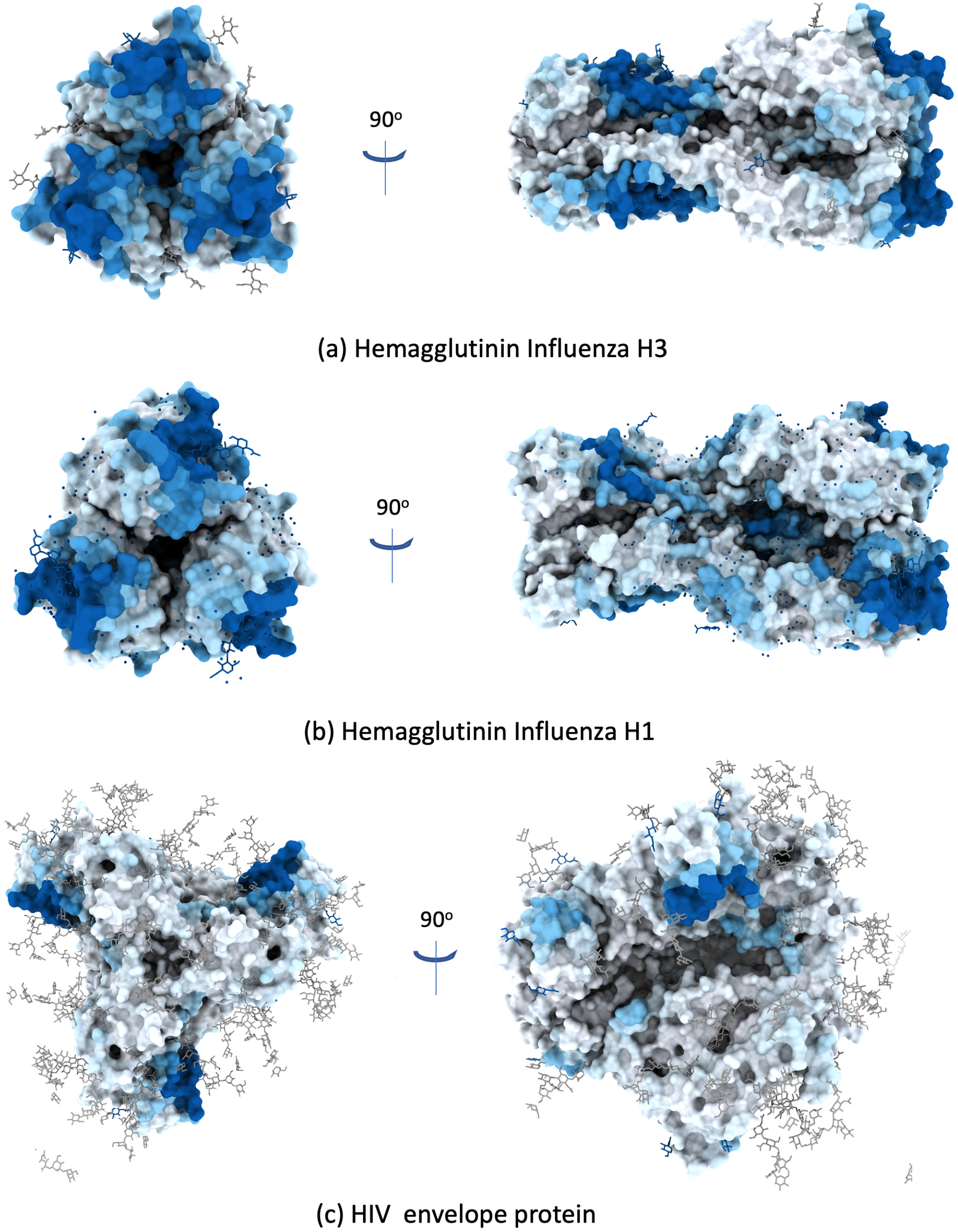
Additional B-cell conformational epitope predictions with ScanNet. a) Hemagglutinin trimer of Influenza H3 (PDB ID: 4o5n [102]) b) Hemagglutinin trimer of Influenza H1 (PDB ID: 1rvx [103]) c) HIV Envelope protein (PDB ID 5fyl [104]). Predictions are performed in cross-validation setting (for each protein, we use the network that was not trained on it). Examples selected from [105]

**FIG. S22.**
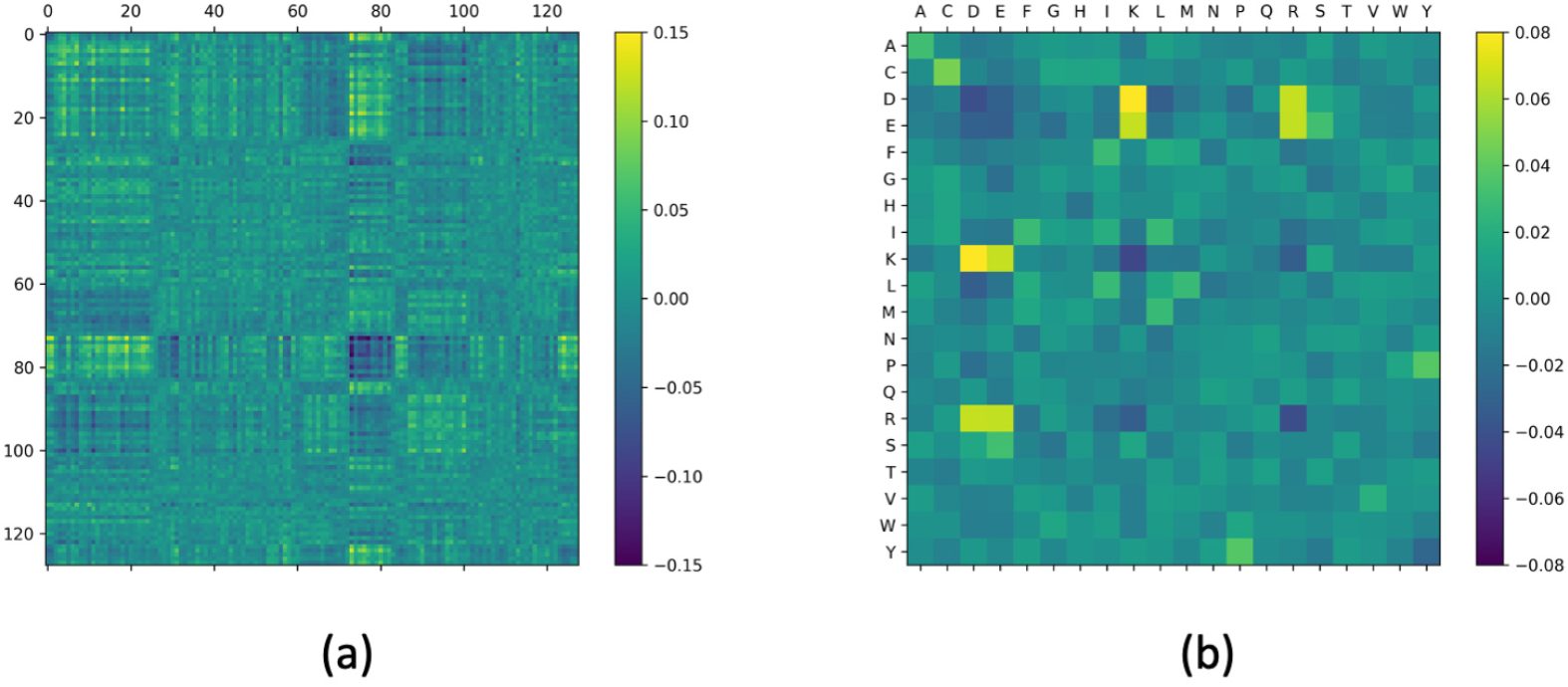
Correlation of ScanNet filter activities and amino acid types between interacting binding sites. For each of the 271 dimers of the benchmark 5.5 dataset [92], the pairs of interacting binding sites (defined as < 4 Å between any two heavy atoms) are identified. Next, the amino acid ScanNet filter activities are computed for each binding site (128-dimensional vector for each residue), and the cross-correlation is subsequently computed (Panel a). The cross-correlation matrix features significantly large entries (|*r*| ~ 0.15), suggesting that favorable interactions require complementary spatio-chemical patterns. As a control, the cross-correlation between amino acid types is similarly computed and features known complementary amino acid pairs such as K/R and D/E (of opposite electrostatic charge), but lower correlations (|*r*| ~ 0.08).

## References

[1] W. Kühlbrandt, The resolution revolution, Science 343, 1443 (2014).

[2] J. Jumper, R. Evans, A. Pritzel, T. Green, M. Figurnov, O. Ronneberger, K. Tunyasuvunakool, R. Bates, A. Žídek, A. Potapenko, et al., Highly accurate protein structure prediction with alphafold, Nature, 1 (2021).

[3] K. Tunyasuvunakool, J. Adler, Z. Wu, T. Green, M. Zielinski, A. Žídek, A. Bridgland, A. Cowie, C. Meyer, A. Laydon, et al., Highly accurate protein structure prediction for the human proteome, Nature, 1 (2021).

[4] M. Chruszcz, M. Domagalski, T. Osinski, A. Wlodawer, and W. Minor, Unmet challenges of structural genomics, Current opinion in structural biology 20, 587 (2010).

[5] A. Shulman-Peleg, R. Nussinov, and H. J. Wolfson, Siteengines: recognition and comparison of binding sites and protein–protein interfaces, Nucleic acids research 33, W337 (2005).

[6] N. Carl, J. Konc, B. Vehar, and D. Janezic, Protein-protein binding site prediction by local structural alignment, Journal of chemical information and modeling 50, 1906 (2010).

[7] Q. C. Zhang, D. Petrey, R. Norel, and B. H. Honig, Protein interface conservation across structure space, Proceedings of the National Academy of Sciences 107, 10896 (2010).

[8] L. C. Xue, D. Dobbs, and V. Honavar, Homppi: a class of sequence homology based protein-protein interface prediction methods, BMC bioinformatics 12, 1 (2011).

[9] B. A. Shoemaker, D. Zhang, M. Tyagi, R. R. Thangudu, J. H. Fong, A. Marchler-Bauer, S. H. Bryant, T. Madej, and A. R. Panchenko, Ibis (inferred biomolecular interaction server) reports, predicts and integrates multiple types of conserved interactions for proteins, Nucleic acids research 40, D834 (2012).

[10] R. A. Jordan, E.-M. Yasser, D. Dobbs, and V. Honavar, Predicting protein-protein interface residues using local surface structural similarity, BMC bioinformatics 13, 1 (2012).

[11] R. Esmaielbeiki and J.-C. Nebel, Unbiased protein interface prediction based on ligand diversity quantification, (2012).

[12] L. C. Xue, D. Dobbs, A. M. Bonvin, and V. Honavar, Computational prediction of protein interfaces: A review of data driven methods, FEBS letters 589, 3516 (2015).

[13] R. Esmaielbeiki, K. Krawczyk, B. Knapp, J.-C. Nebel, and C. M. Deane, Progress and challenges in predicting protein interfaces, Briefings in bioinformatics 17, 117 (2016).

[14] H. Neuvirth, R. Raz, and G. Schreiber, Promate: a structure based prediction program to identify the location of protein–protein binding sites, Journal of molecular biology 338, 181 (2004).

[15] J.-L. Chung, W. Wang, and P. E. Bourne, Exploiting sequence and structure homologs to identify protein–protein binding sites, Proteins: Structure, Function, and Bioinformatics 62, 630 (2006).

[16] A. Porollo and J. Meller, Prediction-based fingerprints of protein–protein interactions, Proteins: Structure, Function, and Bioinformatics 66, 630 (2007).

[17] M. J. Sweredoski and P. Baldi, Pepito: improved discontinuous b-cell epitope prediction using multiple distance thresholds and half sphere exposure, Bioinformatics 24, 1459 (2008).

[18] S. K. Mishra, G. Kandoi, and R. L. Jernigan, Coupling dynamics and evolutionary information with structure to identify protein regulatory and functional binding sites, Proteins: Structure, Function, and Bioinformatics 87, 850 (2019).

[19] A. Klug and D. Rhodes, ‘zinc fingers’: a novel protein motif for nucleic acid recognition, Trends in Biochemical Sciences 12, 464 (1987).

[20] A. A. Bogan and K. S. Thorn, Anatomy of hot spots in protein interfaces, Journal of molecular biology 280, 1 (1998).

[21] M. Wensien, F. R. von Pappenheim, L.-M. Funk, P. Kloskowski, U. Curth, U. Diederichsen, J. Uranga, J. Ye, P. Fang, K.-T. Pan, et al., A lysine–cysteine redox switch with an nos bridge regulates enzyme function, Nature 593, 460 (2021).

[22] A. Elnaggar, M. Heinzinger, C. Dallago, G. Rihawi, Y. Wang, L. Jones, T. Gibbs, T. Feher, C. Angerer, M. Steinegger, et al., Prottrans: towards cracking the language of life’s code through self-supervised deep learning and high performance computing, arXiv preprint arXiv:2007.06225 (2020).

[23] A. Rives, J. Meier, T. Sercu, S. Goyal, Z. Lin, J. Liu, D. Guo, M. Ott, C. L. Zitnick, J. Ma, et al., Biological structure and function emerge from scaling unsupervised learning to 250 million protein sequences, Proceedings of the National Academy of Sciences 118(2021).

[24] J. Ingraham, A. Riesselman, C. Sander, and D. Marks, Learning protein structure with a differentiable simulator, in International Conference on Learning Representations (2018).

[25] J. Ingraham, V. K. Garg, R. Barzilay, and T. Jaakkola, Generative models for graph-based protein design, (2019).

[26] X. Jing and J. Xu, Fast and effective protein model refinement by deep graph neural networks, bioRxiv (2020).

[27] F. Baldassarre, D. Menéndez Hurtado, A. Elofsson, and H. Azizpour, Graphqa: protein model quality assessment using graph convolutional networks, Bioinformatics 37, 360 (2021).

[28] I. Wallach, M. Dzamba, and A. Heifets, Atomnet: a deep convolutional neural network for bioactivity prediction in structure-based drug discovery, arXiv preprint arXiv:1510.02855 (2015).

[29] M. Ragoza, J. Hochuli, E. Idrobo, J. Sunseri, and D. R. Koes, Protein–ligand scoring with convolutional neural networks, Journal of chemical information and modeling 57, 942 (2017).

[30] G. Pagès, B. Charmettant, and S. Grudinin, Protein model quality assessment using 3d oriented convolutional neural networks, Bioinformatics 35, 3313 (2019).

[31] R. Townshend, R. Bedi, P. Suriana, and R. Dror, End-to-end learning on 3d protein structure for interface prediction, Advances in Neural Information Processing Systems 32, 15642 (2019).

[32] X. Wang, G. Terashi, C. W. Christoffer, M. Zhu, and D. Kihara, Protein docking model evaluation by 3d deep convolutional neural networks, Bioinformatics 36, 2113 (2020).

[33] I. Igashov, K. Olechnovic, M. Kadukova, Č. Venclovas, and S. Grudinin, Vorocnn: Deep convolutional neural network built on 3d voronoi tessellation of protein structures, bioRxiv (2020).

[34] N. Renaud, C. Geng, S. Georgievska, F. Ambrosetti, L. Ridder, D. F. Marzella, A. M. Bonvin, and L. C. Xue, Deeprank: A deep learning framework for data mining 3d protein-protein interfaces, Biorxiv (2021).

[35] S. Eismann, P. Suriana, B. Jing, R. J. Townshend, and R. O. Dror, Protein model quality assessment using rotation-equivariant, hierarchical neural networks, arXiv preprint arXiv:2011.13557 (2020).

[36] P. Gainza, F. Sverrisson, F. Monti, E. Rodola, D. Boscaini, M. Bronstein, and B. Correia, Deciphering interaction fingerprints from protein molecular surfaces using geometric deep learning, Nature Methods 17, 184 (2020).

[37] F. Sverrisson, J. Feydy, B. E. Correia, and M. M. Bronstein, Fast end-to-end learning on protein surfaces, in Proceedings of the IEEE/CVF Conference on Computer Vision and Pattern Recognition (2021) pp. 15272–15281.

[38] M. M. Bronstein, J. Bruna, Y. LeCun, A. Szlam, and P. Vandergheynst, Geometric deep learning: going beyond euclidean data, IEEE Signal Processing Magazine 34, 18 (2017).

[39] M. M. Bronstein, J. Bruna, T. Cohen, and P. Veličković, Geometric deep learning: Grids, groups, graphs, geodesics, and gauges, arXiv preprint arXiv:2104.13478 (2021).

[40] J. Gilmer, S. S. Schoenholz, P. F. Riley, O. Vinyals, and G. E. Dahl, Neural message passing for quantum chemistry, in International conference on machine learning (PMLR, 2017) pp. 1263–1272.

[41] P. Veličković, G. Cucurull, A. Casanova, A. Romero, P. Lio, and Y. Bengio, Graph attention networks, arXiv preprint arXiv:1710.10903 (2017).

[42] O. Keskin, B. Ma, and R. Nussinov, Hot regions in protein–protein interactions: the organization and contribution of structurally conserved hot spot residues, Journal of molecular biology 345, 1281 (2005).

[43] Y. Ofran and B. Rost, Protein–protein interaction hotspots carved into sequences, PLoS computational biology 3, e119 (2007).

[44] S. Dey, D. W. Ritchie, and E. D. Levy, Pdb-wide identification of biological assemblies from conserved quaternary structure geometry, Nature methods 15, 67 (2018).

[45] P. J. Kundrotas, I. Anishchenko, T. Dauzhenka, I. Kotthoff, D. Mnevets, M. M. Copeland, and I. A. Vakser, Dockground: a comprehensive data resource for modeling of protein complexes, Protein Science 27, 172 (2018).

[46] T. Chen and C. Guestrin, Xgboost: A scalable tree boosting system, in Proceedings of the 22nd acm sigkdd international conference on knowledge discovery and data mining (2016) pp. 785–794.

[47] M. Shatsky, R. Nussinov, and H. J. Wolfson, Multiprot—a multiple protein structural alignment algorithm, in International Workshop on Algorithms in Bioinformatics (Springer, 2002) pp. 235–250.

[48] I. Sillitoe, N. Bordin, N. Dawson, V. P. Waman, P. Ashford, H. M. Scholes, C. S. Pang, L. Woodridge, C. Rauer, N. Sen, et al., Cath: increased structural coverage of functional space, Nucleic acids research 49, D266 (2021).

[49] E. Jurrus, D. Engel, K. Star, K. Monson, J. Brandi, L. E. Felberg, D. H. Brookes, L. Wilson, J. Chen, K. Liles, et al., Improvements to the apbs biomolecular solvation software suite, Protein Science 27, 112 (2018).

[50] J. Dunbar, K. Krawczyk, J. Leem, T. Baker, A. Fuchs, G. Georges, J. Shi, and C. M. Deane, Sabdab: the structural antibody database, Nucleic acids research 42, D1140 (2014).

[51] J. V. Kringelum, C. Lundegaard, O. Lund, and M. Nielsen, Reliable b cell epitope predictions: impacts of method development and improved benchmarking, PLoS Comput Biol 8, e1002829 (2012).

[52] M. Yuan, D. Huang, C.-C. D. Lee, N. C. Wu, A. M. Jackson, X. Zhu, H. Liu, L. Peng, M. J. van Gils, R. W. Sanders, et al., Structural and functional ramifications of antigenic drift in recent sars-cov-2 variants, Science (2021).

[53] E. Shrock, E. Fujimura, T. Kula, R. T. Timms, I.-H. Lee, Y. Leng, M. L. Robinson, B. M. Sie, M. Z. Li, Y. Chen, et al., Viral epitope profiling of covid-19 patients reveals cross-reactivity and correlates of severity, Science 370(2020).

[54] M. M. Sauer, M. A. Tortorici, Y.-J. Park, A. C. Walls, L. Homad, O. J. Acton, J. E. Bowen, C. Wang, X. Xiong, W. de van der Schueren, et al., Structural basis for broad coronavirus neutralization, Nature Structural & Molecular Biology, 1 (2021).

[55] Y. Watanabe, J. D. Allen, D. Wrapp, J. S. McLellan, and M. Crispin, Site-specific glycan analysis of the sars-cov-2 spike, Science 369, 330 (2020).

[56] R. Evans, M. O’Neill, A. Pritzel, N. Antropova, A. W. Senior, T. Green, A. Žídek, R. Bates, S. Blackwell, J. Yim, et al., Protein complex prediction with alphafold-multimer, Biorxiv (2021).

[57] A. M. Buckle, G. Schreiber, and A. R. Fersht, Protein-protein recognition: Crystal structural analysis of a barnase-barstar complex at 2.0-. ang. resolution, Biochemistry 33, 8878 (1994).

[58] G. Fenalti, R. H. Law, A. M. Buckle, C. Langendorf, K. Tuck, C. J. Rosado, N. G. Faux, K. Mahmood, C. S. Hampe, J. P. Banga, et al., Gaba production by glutamic acid decarboxylase is regulated by a dynamic catalytic loop, Nature structural & molecular biology 14, 280 (2007).

[59] T. D. Goddard, C. C. Huang, E. C. Meng, E. F. Pettersen, G. S. Couch, J. H. Morris, and T. E. Ferrin, Ucsf chimerax: Meeting modern challenges in visualization and analysis, Protein Science 27, 14 (2018).

[60] L. Van der Maaten and G. Hinton, Visualizing data using t-sne., Journal of machine learning research 9(2008).

[61] W. Kabsch and C. Sander, Dictionary of protein secondary structure: pattern recognition of hydrogen-bonded and geometrical features, Biopolymers: Original Research on Biomolecules 22, 2577 (1983).

[62] R. Amaro and A. Mulholland, Biomolecular simulations in the time of covid19, and after, Computing in Science & Engineering (2020).

[63] P. J. Cock, T. Antao, J. T. Chang, B. A. Chapman, C. J. Cox, A. Dalke, I. Friedberg, T. Hamelryck, F. Kauff, B. Wilczynski, et al., Biopython: freely available python tools for computational molecular biology and bioinformatics, Bioinformatics 25, 1422 (2009).

[64] If both are equally far away e.g. for isoleucine, we choose the first one according to the residue id.

[65] M. Remmert, A. Biegert, A. Hauser, and J. Söding, Hhblits: lightning-fast iterative protein sequence searching by hmm-hmm alignment, Nature methods 9, 173 (2012).

[66] M. Mirdita, L. von den Driesch, C. Galiez, M. J. Martin, J. Söding, and M. Steinegger, Uniclust databases of clustered and deeply annotated protein sequences and alignments, Nucleic acids research 45, D170 (2017).

[67] S. Cocco, C. Feinauer, M. Figliuzzi, R. Monasson, and M. Weigt, Inverse statistical physics of protein sequences: a key issues review, Reports on Progress in Physics 81, 032601 (2018).

[68] L. Posani, Inference and modeling of biological networks: a statistical-physics approach to neural attractors and protein fitness landscapes, Ph.D. thesis, Université Paris sciences et lettres (2018).

[69] W. Chen, X. Han, G. Li, C. Chen, J. Xing, Y. Zhao, and H. Li, Deep rbfnet: Point cloud feature learning using radial basis functions, arXiv preprint arXiv:1812.04302 (2018).

[70] C. R. Qi, H. Su, K. Mo, and L. J. Guibas, Pointnet: Deep learning on point sets for 3d classification and segmentation, in Proceedings of the IEEE conference on computer vision and pattern recognition (2017) pp. 652–660.

[71] C. R. Qi, L. Yi, H. Su, and L. J. Guibas, Pointnet++: Deep hierarchical feature learning on point sets in a metric space, arXiv preprint arXiv:1706.02413 (2017).

[72] I. Igashov, N. Pavlichenko, and S. Grudinin, Spherical convolutions on molecular graphs for protein model quality assessment, Machine Learning: Science and Technology (2021).

[73] J. Tubiana, S. Cocco, and R. Monasson, Learning protein constitutive motifs from sequence data, Elife 8, e39397 (2019).

[74] S. Ioffe and C. Szegedy, Batch normalization: Accelerating deep network training by reducing internal covariate shift, in International conference on machine learning (PMLR, 2015) pp. 448–456.

[75] F. Pedregosa, G. Varoquaux, A. Gramfort, V. Michel, B. Thirion, O. Grisel, M. Blondel, P. Prettenhofer, R. Weiss, V. Dubourg, et al., Scikit-learn: Machine learning in python, the Journal of machine Learning research 12, 2825 (2011).

[76] J. Long, E. Shelhamer, and T. Darrell, Fully convolutional networks for semantic segmentation, in Proceedings of the IEEE conference on computer vision and pattern recognition (2015) pp. 3431–3440.

[77] D. P. Kingma and J. Ba, Adam: A method for stochastic optimization, arXiv preprint arXiv:1412.6980 (2014).

[78] M. Abadi, P. Barham, J. Chen, Z. Chen, A. Davis, J. Dean, M. Devin, S. Ghemawat, G. Irving, M. Isard, et al., Tensorflow: A system for large-scale machine learning, in 12th {USENIX} symposium on operating systems design and implementation ({OSDI} 16) (2016) pp. 265–283.

[79] F. Chollet, Deep learning with Python (Simon and Schuster, 2017).

[80] J. Song, H. Tan, K. Takemoto, and T. Akutsu, Hsepred: predict half-sphere exposure from protein sequences, Bioinformatics 24, 1489 (2008).

[81] S. Chakravarty and R. Varadarajan, Residue depth: a novel parameter for the analysis of protein structure and stability, Structure 7, 723 (1999).

[82] M. F. Sanner, A. J. Olson, and J.-C. Spehner, Reduced surface: an efficient way to compute molecular surfaces, Biopolymers 38, 305 (1996).

[83] M. L. Connolly, Shape complementarity at the hemoglobin *α1β1* subunit interface, Biopolymers 25, 1229 (1986).

[84] L. Fu, B. Niu, Z. Zhu, S. Wu, and W. Li, Cd-hit: accelerated for clustering the next-generation sequencing data, Bioinformatics 28, 3150 (2012).

[85] W. Li and A. Godzik, Cd-hit: a fast program for clustering and comparing large sets of protein or nucleotide sequences, Bioinformatics 22, 1658 (2006).

[86] T. Nakamura, K. D. Yamada, K. Tomii, and K. Katoh, Parallelization of mafft for large-scale multiple sequence alignments, Bioinformatics 34, 2490 (2018).

[87] For the PPBS data set, 8.9% of the binding site residues are actually in unbound conformation, as their label was inferred from another pdb file, see data preparation.

[88] T. Kirys, A. M. Ruvinsky, D. Singla, A. V. Tuzikov, P. J. Kundrotas, and I. A. Vakser, Simulated unbound structures for benchmarking of protein docking in the dockground resource, BMC bioinformatics 16, 1 (2015).

[89] U. Ghani, I. Desta, A. Jindal, O. Khan, G. Jones, S. Kotelnikov, D. Padhorny, S. Vajda, and D. Kozakov, Improved docking of protein models by a combination of alphafold2 and cluspro, BioRxiv (2021).

[90] M. Mirdita, K. Schütze, Y. Moriwaki, L. Heo, S. Ovchinnikov, and M. Steinegger, Colabfold-making protein folding accessible to all, (2021).

[91] J. Jankauskaitė, B. Jiménez-García, J. Dapkūnas, J. Fernández-Recio, and I. H. Moal, Skempi 2.0: an updated benchmark of changes in protein–protein binding energy, kinetics and thermodynamics upon mutation, Bioinformatics 35, 462 (2019).

[92] T. Vreven, I. H. Moal, A. Vangone, B. G. Pierce, P. L. Kastritis, M. Torchala, R. Chaleil, B. Jiménez-García, P. A. Bates, J. Fernandez-Recio, et al., Updates to the integrated protein–protein interaction benchmarks: docking benchmark version 5 and affinity benchmark version 2, Journal of molecular biology 427, 3031 (2015).

[93] R. F. Alford, A. Leaver-Fay, J. R. Jeliazkov, M. J. O’Meara, F. P. DiMaio, H. Park, M. V. Shapovalov, P. D. Renfrew, V. K. Mulligan, K. Kappel, et al., The rosetta all-atom energy function for macromolecular modeling and design, Journal of chemical theory and computation 13, 3031 (2017).

[94] S. Chaudhury, S. Lyskov, and J. J. Gray, Pyrosetta: a script-based interface for implementing molecular modeling algorithms using rosetta, Bioinformatics 26, 689 (2010).

[95] M. Yuan, H. Liu, N. C. Wu, C.-C. D. Lee, X. Zhu, F. Zhao, D. Huang, W. Yu, Y. Hua, H. Tien, et al., Structural basis of a public antibody response to sars-cov-2, BioRxiv (2020).

[96] N. C. Wu, M. Yuan, H. Liu, C.-C. D. Lee, X. Zhu, S. Bangaru, J. L. Torres, T. G. Caniels, P. J. Brouwer, M. J. Van Gils, et al., An alternative binding mode of ighv3-53 antibodies to the sars-cov-2 receptor binding domain, Cell reports 33, 108274 (2020).

[97] J. Kreye, S. M. Reincke, H.-C. Kornau, E. Sánchez-Sendin, V. M. Corman, H. Liu, M. Yuan, N. C. Wu, X. Zhu, C.-C. D. Lee, et al., A therapeutic non-self-reactive sars-cov-2 antibody protects from lung pathology in a covid-19 hamster model, Cell 183, 1058 (2020).

[98] J. Hansen, A. Baum, K. E. Pascal, V. Russo, S. Giordano, E. Wloga, B. O. Fulton, Y. Yan, K. Koon, K. Patel, et al., Studies in humanized mice and convalescent humans yield a sars-cov-2 antibody cocktail, Science 369, 1010 (2020).

[99] H. Liu, M. Yuan, D. Huang, S. Bangaru, F. Zhao, C.-C. D. Lee, L. Peng, S. Barman, X. Zhu, D. Nemazee, et al., A combination of cross-neutralizing antibodies synergizes to prevent sars-cov-2 and sars-cov pseudovirus infection, Cell host & microbe 29, 806 (2021).

[100] H. Liu, N. C. Wu, M. Yuan, S. Bangaru, J. L. Torres, T. G. Caniels, J. Van Schooten, X. Zhu, C.-C. D. Lee, P. J. Brouwer, et al., Cross-neutralization of a sars-cov-2 antibody to a functionally conserved site is mediated by avidity, Immunity 53, 1272 (2020).

[101] G. Cerutti, Y. Guo, T. Zhou, J. Gorman, M. Lee, M. Rapp, E. R. Reddem, J. Yu, F. Bahna, J. Bimela, et al., Potent sars-cov-2 neutralizing antibodies directed against spike n-terminal domain target a single supersite, Cell Host & Microbe 29, 819 (2021).

[102] P. S. Lee, N. Ohshima, R. L. Stanfield, W. Yu, Y. Iba, Y. Okuno, Y. Kurosawa, and I. A. Wilson, Receptor mimicry by antibody f045–092 facilitates universal binding to the h3 subtype of influenza virus, Nature communications 5, 1 (2014).

[103] S. Gamblin, L. Haire, R. Russell, D. Stevens, B. Xiao, Y. Ha, N. Vasisht, D. Steinhauer, R. Daniels, A. Elliot, et al., The structure and receptor binding properties of the 1918 influenza hemagglutinin, Science 303, 1838 (2004).

[104] G. B. Stewart-Jones, C. Soto, T. Lemmin, G.-Y. Chuang, A. Druz, R. Kong, P. V. Thomas, K. Wagh, T. Zhou, A.-J. Behrens, et al., Trimeric hiv-1-env structures define glycan shields from clades a, b, and g, Cell 165, 813 (2016).

[105] B. Hie, E. D. Zhong, B. Berger, and B. Bryson, Learning the language of viral evolution and escape, Science 371, 284 (2021).

