## Supplementary material for "ScanNet: An interpretable geometric deep learning model for structure-based protein binding site prediction": 3D visualisation and activation statistics of all the atomic filters of the protein-protein binding site network

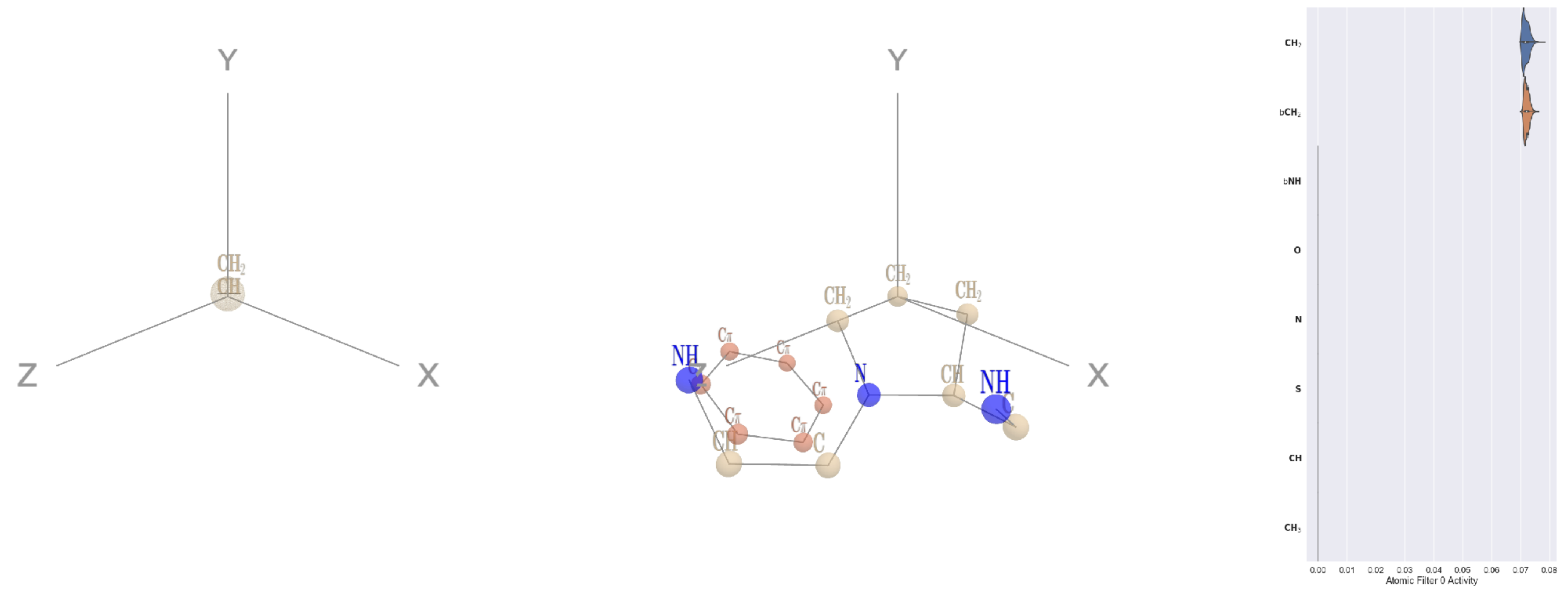


Filter 0
$R_{\mathrm{pred}}^{\mathrm{max}} = 0.04$
$R_{\mathrm{pred}}^{\mathrm{mean}} = 0.02$
Percent Active = 16
Pooling attention 0.00
Pooling feature 0.00
Top activators
1nn4:A:44 P CG
3kyh:C:265 P CD
4mex:C:894 Q CG
1r7r:A:616 N CB
2o3o:A:34 K CB
4buj:B:489 D CB
1rq0:A:64 K CE
4dag:A:64 P CD
5ui2:A:229 F CB
4ani:A:167 P CD

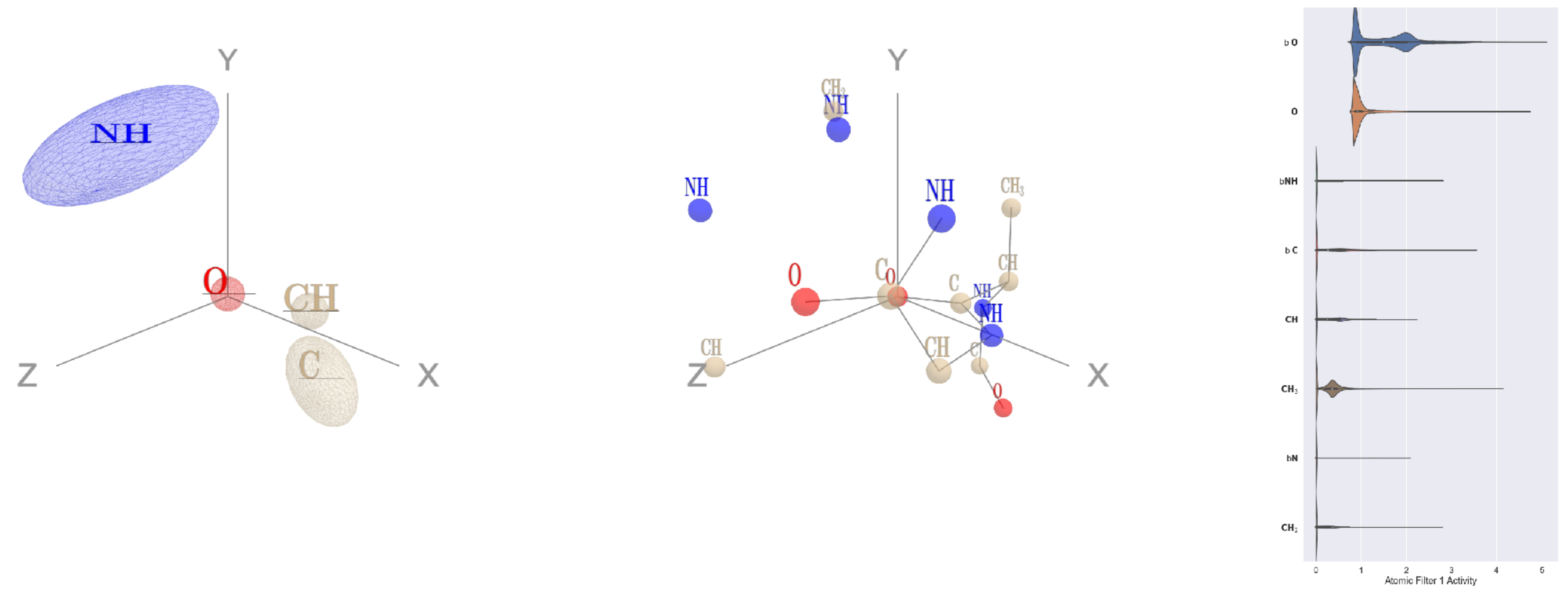


Filter 1
$R_{\mathrm{pred}}^{\mathrm{max}} = -0.00$
$R_{\mathrm{pred}}^{\mathrm{mean}} = -0.05$
Percent Active = 44
Pooling attention 0.14
Pooling feature 0.26
Top activators
4hh2:D:194 A O
2iwh:B:302 D O
1cm5:A:412 D O
4cy6:A:182 S O
1p32:A:84 D O
4pbw:B:75 D O
4cy6:A:458 D O
3bof:A:127 N O
1h4s:A:327 R O
4k1n:A:200 D O

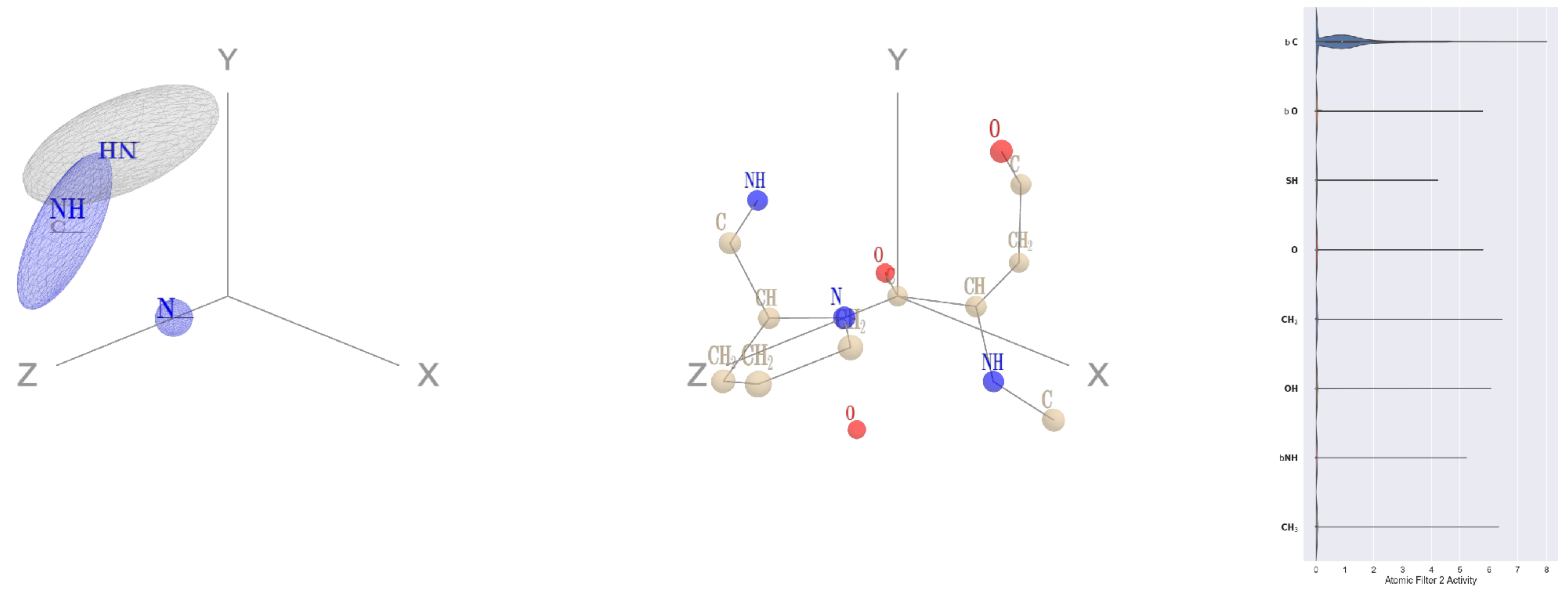


Filter 2
$R_{\mathrm{pred}}^{\mathrm{max}} = -0.04$
$R_{\mathrm{pred}}^{\mathrm{mean}} = -0.08$
Percent Active = 21
Pooling attention 0.10
Pooling feature 0.86
Top activators
2fq1:A:270 N C
2pmz:D:128 I C
1eg1:A:238 G C
1i3r:B:190 V C
1fx0:B:329 V C
1sl0:A:352 V C
1izl:D:330 A C
4qiw:A:155 F C
1adj:A:249 V C
4ldb:A:96 L C

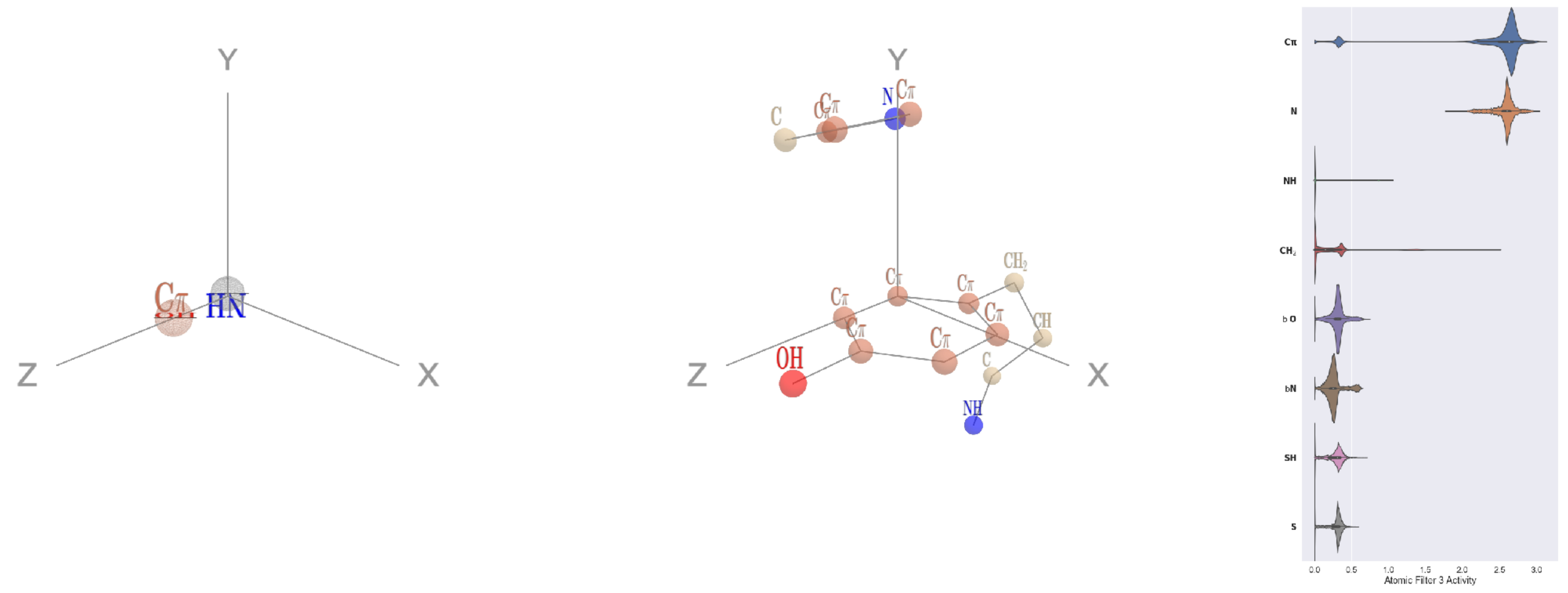


Filter 3
$R_{\mathrm{pred}}^{\mathrm{max}} = 0.05$
$R_{\mathrm{pred}}^{\mathrm{mean}} = 0.06$
Percent Active = 64
Pooling attention 0.08
Pooling feature 2.37
Top activators
1o7g:A:394 Y CD2
3h1s:B:190 F CD2
3h1s:B:185 F CD2
3u33:A:42 F CD1
4v8p:BC:42 F CD2
1cm5:A:594 Y CD1
1h4r:A:119 F CD2
1jqn:A:668 Y CD2
3i6x:A:96 Y CD2
3nh5:A:204 F CD2

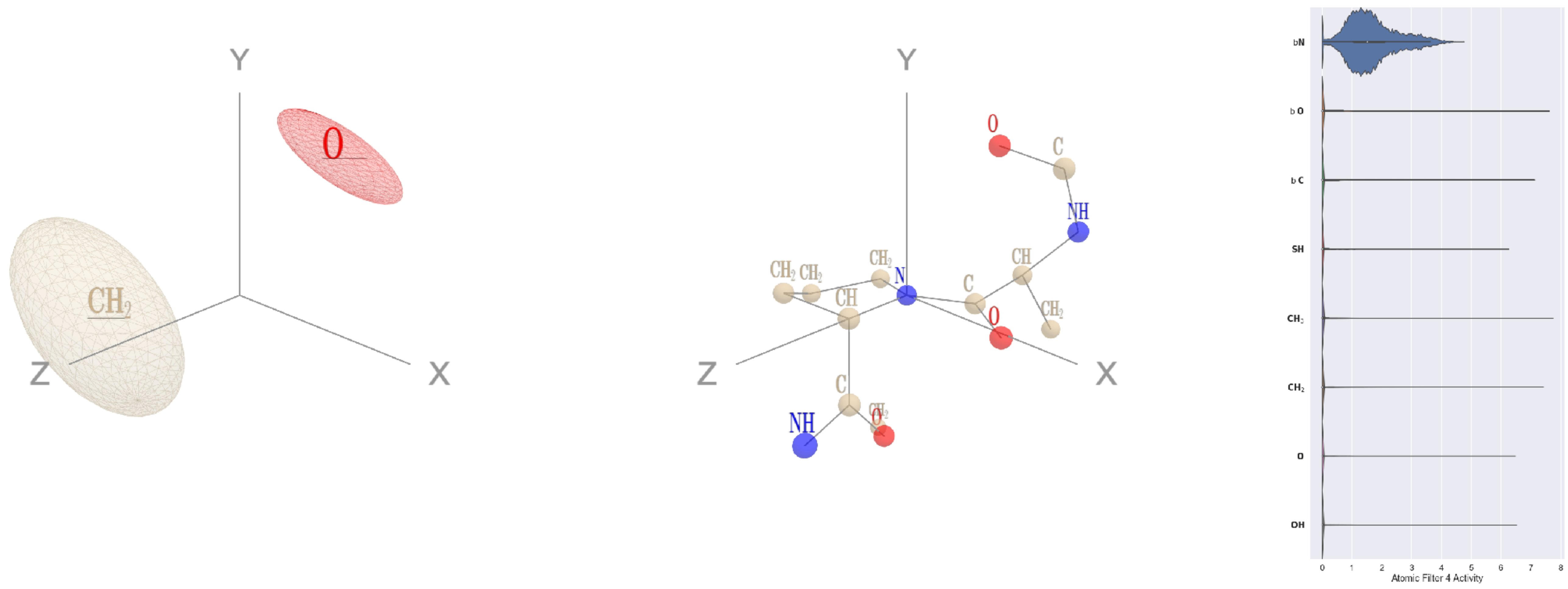


Filter 4
$R_{\text{pred}}^{\text{max}} = -0.03$
$R_{\text{pred}}^{\text{mean}} = -0.06$
Percent Active = 26
Pooling attention 0.13
Pooling feature 0.53
Top activators
4buj:B:1327 P N
1ppt:A:13 P N
2np9:A:223 P N
2iwh:B:922 P N
1v8f:A:67 P N
2pm9:A:374 P N
3b8e:A:217 P N
1d0n:A:240 P N
3b8e:A:504 P N
1bcp:A:4 P N

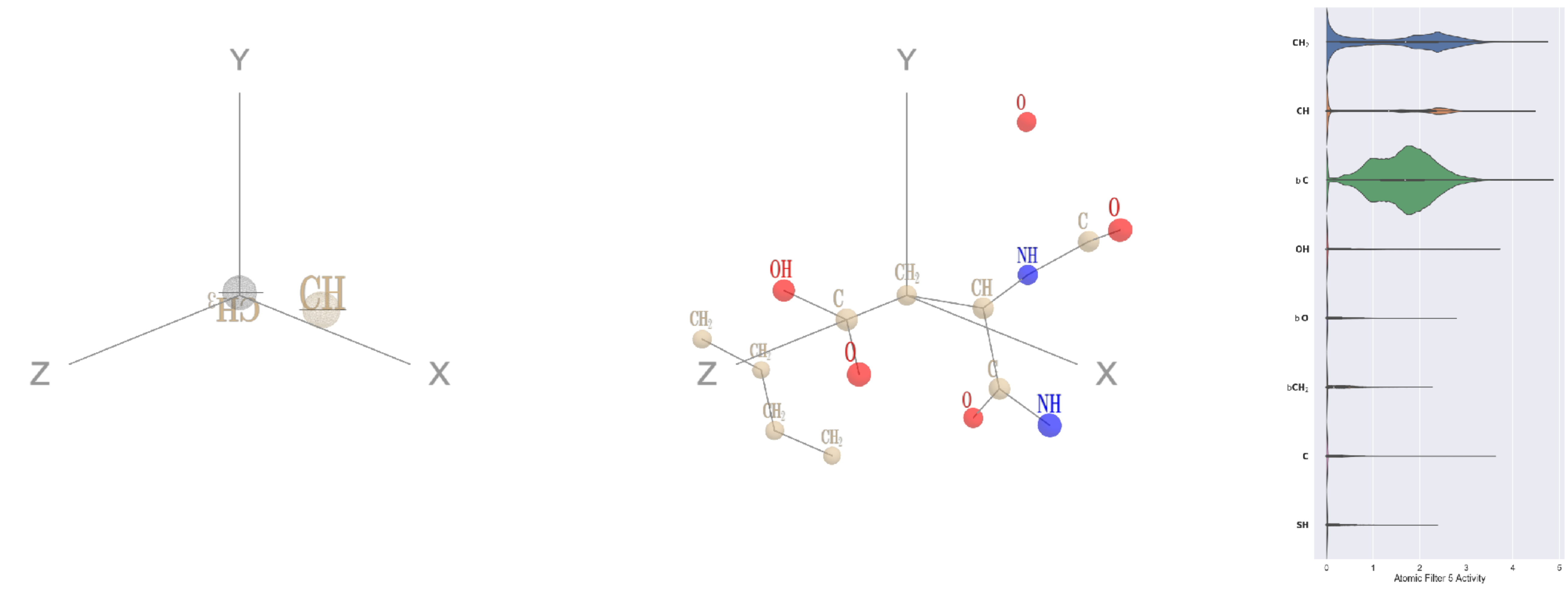


Filter 5
$R_{\text{pred}}^{\text{max}} = 0.02$
$R_{\text{pred}}^{\text{mean}} = -0.08$
Percent Active = 54
Pooling attention 0.43
Pooling feature 0.14
Top activators
2d73:A:356 D CB
1jqn:A:190 E CB
5ui2:A:229 F CB
4c2m:N:149 D CB
2v6e:A:234 D CB
3l0o:A:351 R CB
1u5e:A:125 K CB
1izl:D:183 L CB
4pk7:A:581 D CB
3wqa:A:3354 Y CB

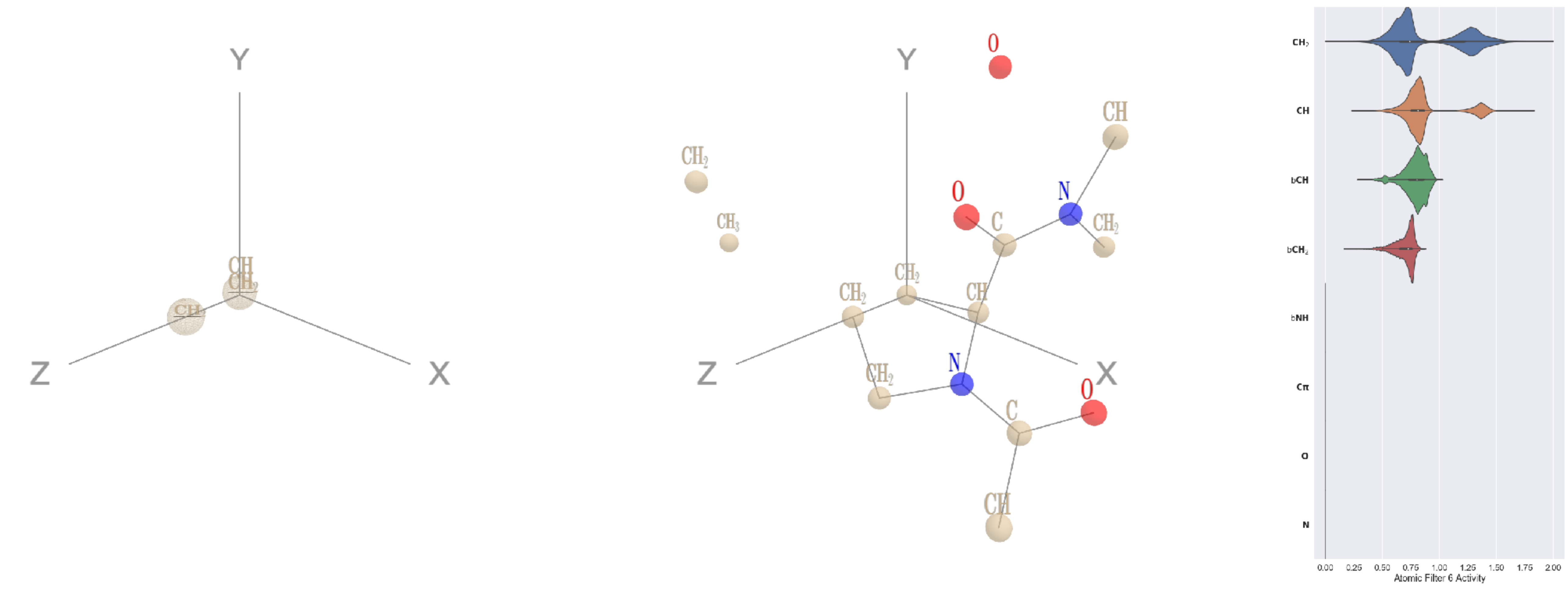


Filter 6
$R_{\text{pred}}^{\text{max}} = 0.05$
$R_{\text{pred}}^{\text{mean}} = 0.02$
Percent Active = 32
Pooling attention 0.03
Pooling feature 0.77
Top activators
1lnl:A:396 P CB
2hh1:H:217 P CB
3c24:A:134 P CB
3h8a:E:13 P CB
3dlb:A:28 R CG
2np9:A:321 K CD
2cge:A:328 P CB
4kzx:R:122 P CB
1lnl:A:206 P CB
1lnl:A:249 P CG

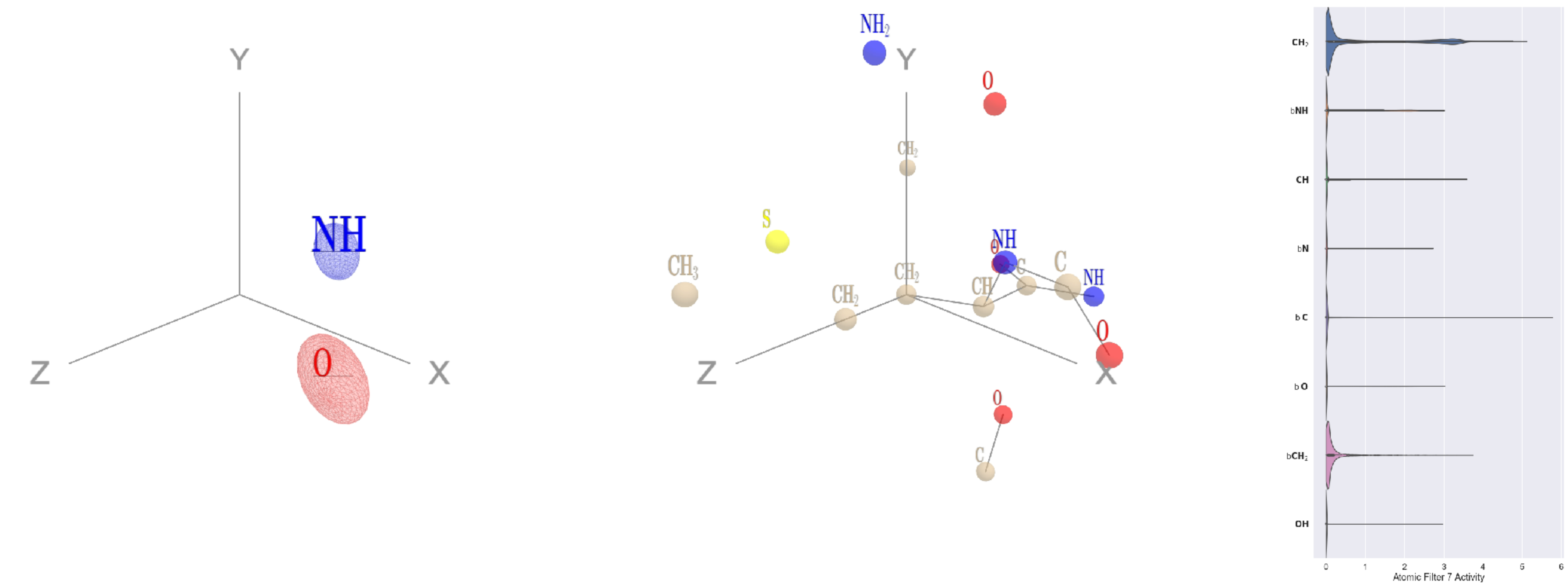


Filter 7
$R_{\text{pred}}^{\text{max}} = 0.04$
$R_{\text{pred}}^{\text{mean}} = 0.01$
Percent Active = 29
Pooling attention 0.13
Pooling feature 0.19
Top activators
4ny2:A:306 M CB
1qhd:A:300 M CB
4kzx:E:201 H CB
3ztv:A:335 E CB
2nog:A:859 K CB
1mj2:A:92 M CB
1w99:A:537 Y CB
3v4l:A:719 L CB
3b8e:A:683 F CB
5kqv:E:207 C CB

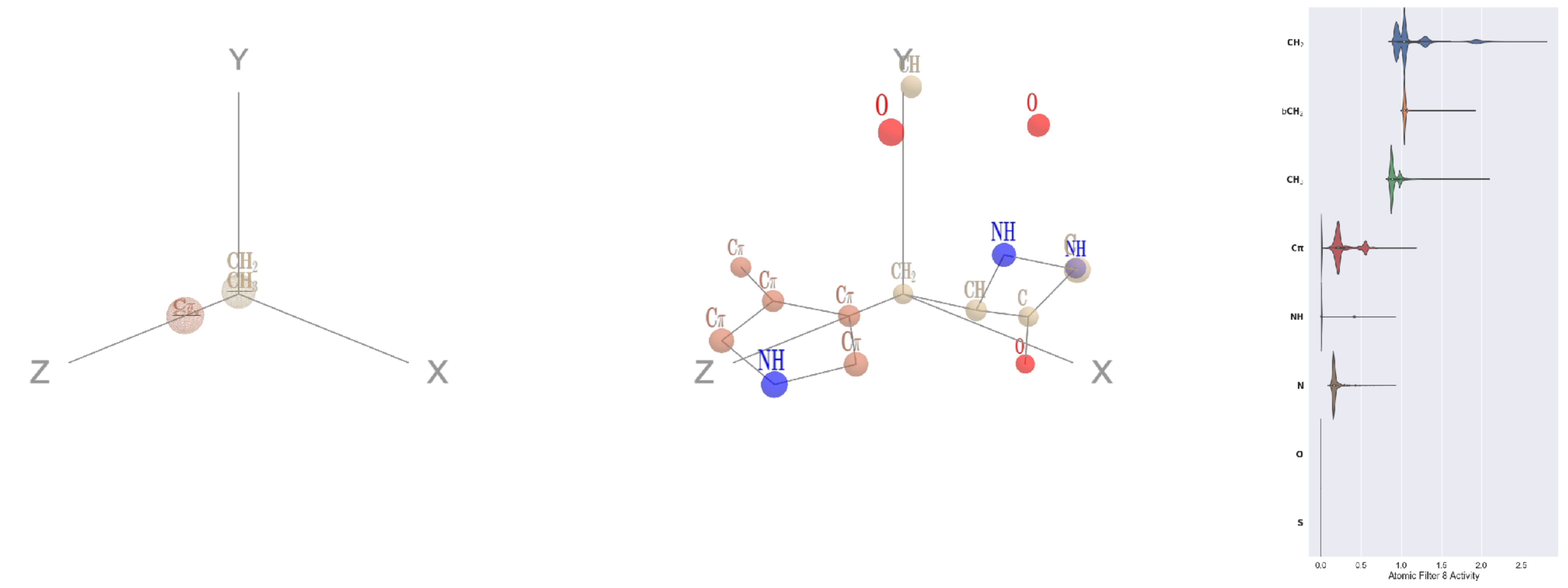


Filter 8
$R_{\mathrm{pred}}^{\max} = 0.04$
$R_{\mathrm{pred}}^{\mathrm{mean}} = 0.01$
Percent Active = 31
Pooling attention 0.02
Pooling feature 1.00
Top activators
1lnl:A:187 W CB
1lnl:A:93 W CB
3eag:A:283 W CB
1l7k:B:143 H CB
1ivu:A:592 H CB
4kzx:E:130 F CB
2wsc:B:557 F CB
3dlb:A:415 W CB
1lnl:A:222 W CB
2pm9:A:45 W CB

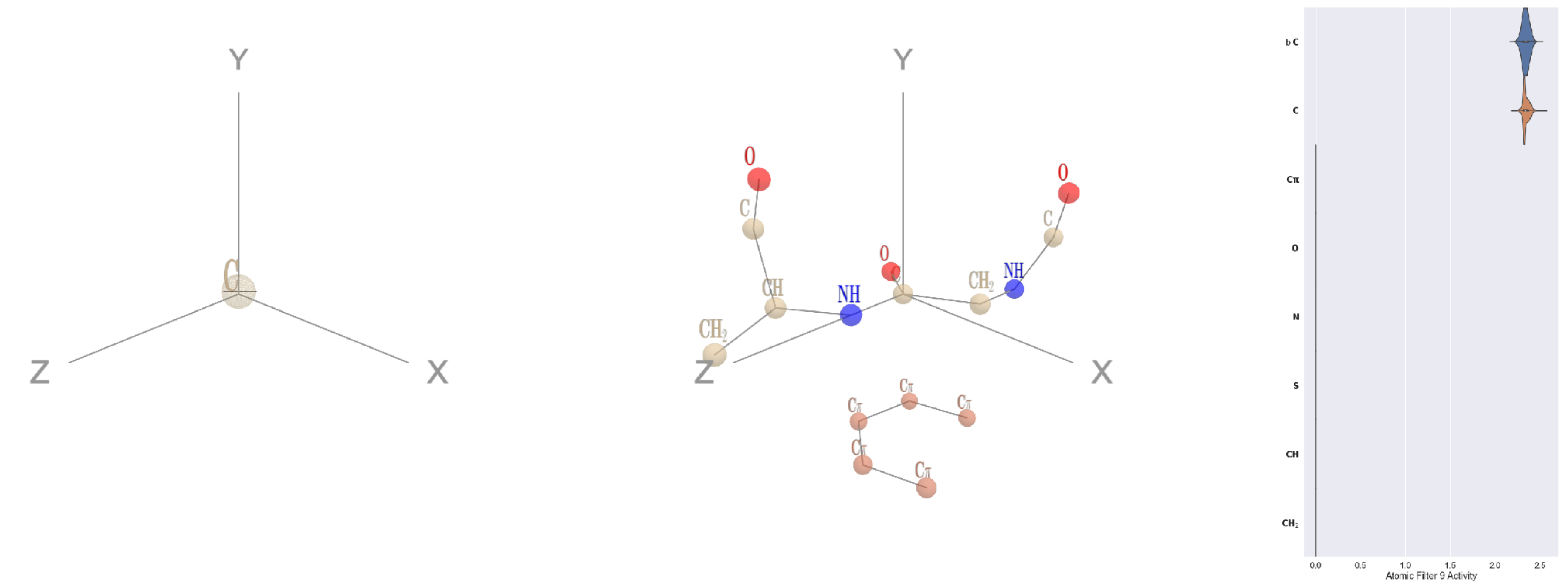


Filter 9
$R_{\mathrm{pred}}^{\max} = 0.03$
$R_{\mathrm{pred}}^{\mathrm{mean}} = -0.02$
Percent Active = 16
Pooling attention 0.22
Pooling feature 0.02
Top activators
1ghq:C:40 G C
1mu2:A:315 E C
2a62:A:190 G C
2gfq:A:108 S C
2b0o:F:613 G C
4u7u:M:402 G C
1y4j:B:209 G C
2pys:B:8 C C
1vpn:B:170 E C
1f4a:A:665 S C

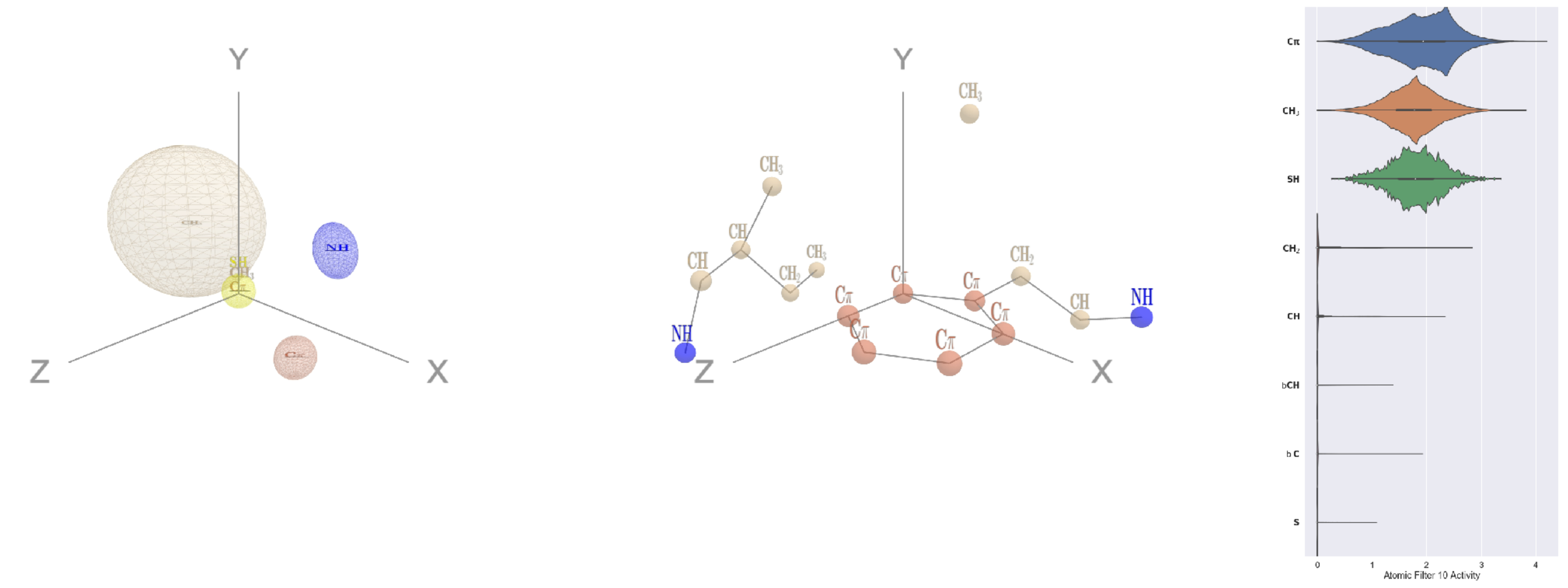


Filter 10
$R_{\mathrm{pred}}^{\max} = -0.01$
$R_{\mathrm{pred}}^{\mathrm{mean}} = -0.00$
Percent Active = 29
Pooling attention 0.30
Pooling feature 1.68
Top activators
2v6e:A:496 F CD2
1ewy:A:216 F CZ
3b8e:A:310 W CZ3
1am4:A:199 F CE1
4c8q:H:605 F CD2
4v4k:M:34 F CE2
3irm:A:52 F CE2
4v8p:BC:255 F CE1
1c3r:A:34 F CZ
1jqj:A:319 F CZ

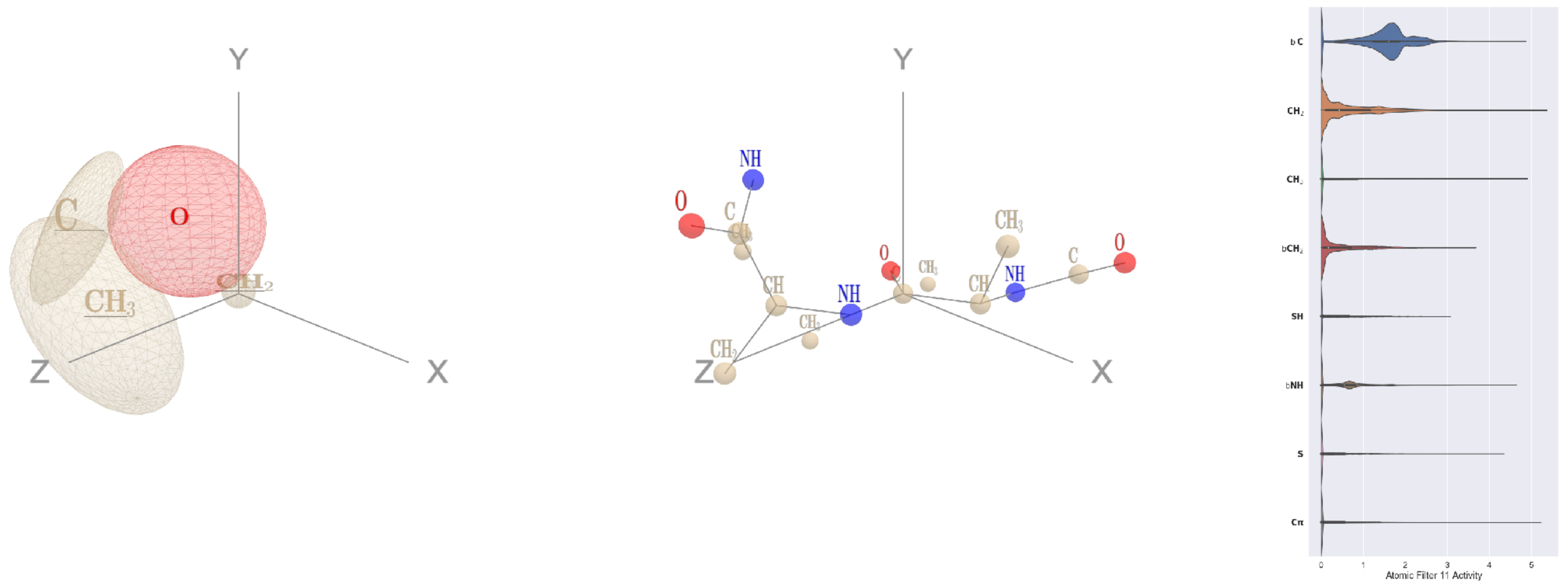


Filter 11
$R_{\mathrm{pred}}^{\max} = -0.09$
$R_{\mathrm{pred}}^{\mathrm{mean}} = -0.20$
Percent Active = 54
Pooling attention 0.29
Pooling feature 0.07
Top activators
2pmz:D:174 A C
4v8p:BC:154 E C
4g1m:B:400 E C
1fx0:B:201 G C
1aom:A:261 G C
4mex:C:13 K C
4s20:L:161 I C
2wsc:B:605 N C
3dlb:A:340 R C
2nyg:A:48 G C

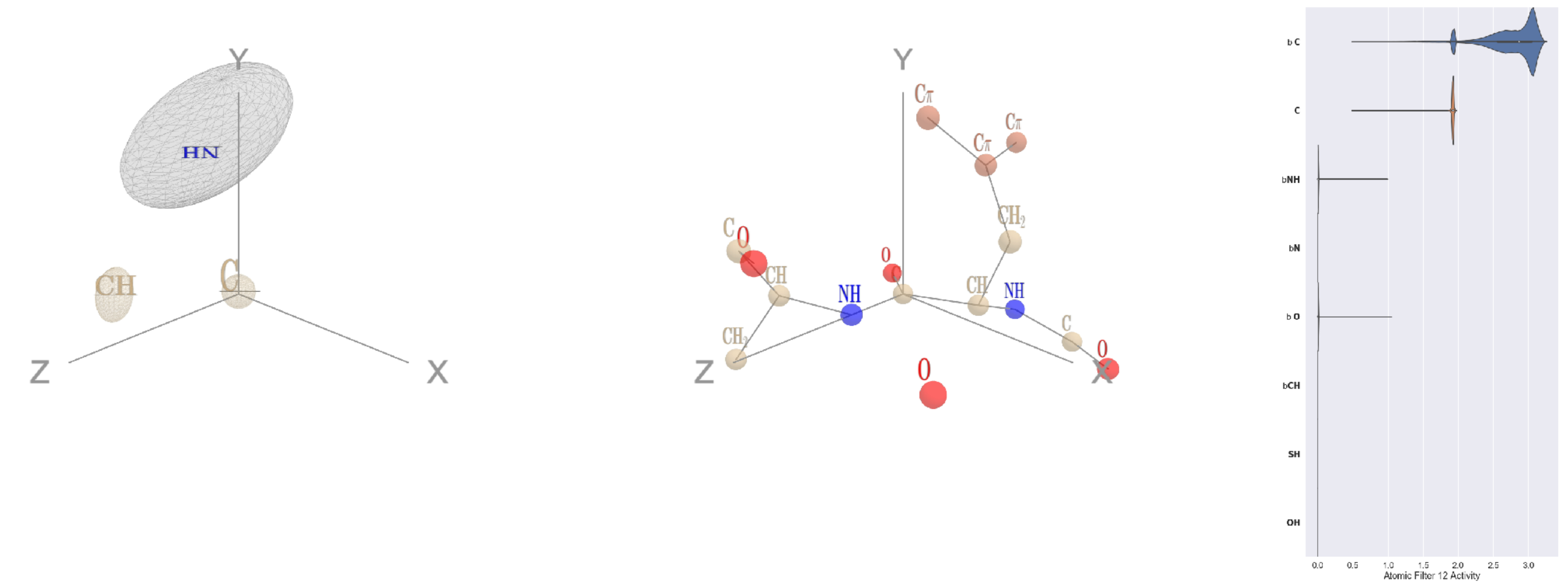


Filter 12
$R_{\text{pred}}^{\text{max}} = 0.00$
$R_{\text{pred}}^{\text{mean}} = -0.04$
Percent Active = 18
Pooling attention 0.21
Pooling feature 0.02
Top activators
3gku:A:31 Y C
4u4c:A:150 L C
2fa0:A:10 M C
1vl4:A:144 Y C
4asi:C:2055 W C
4d0l:A:694 K C
3d00:A:179 L C
2fa0:A:184 S C
4jdm:C:121 N C
2cge:A:337 K C

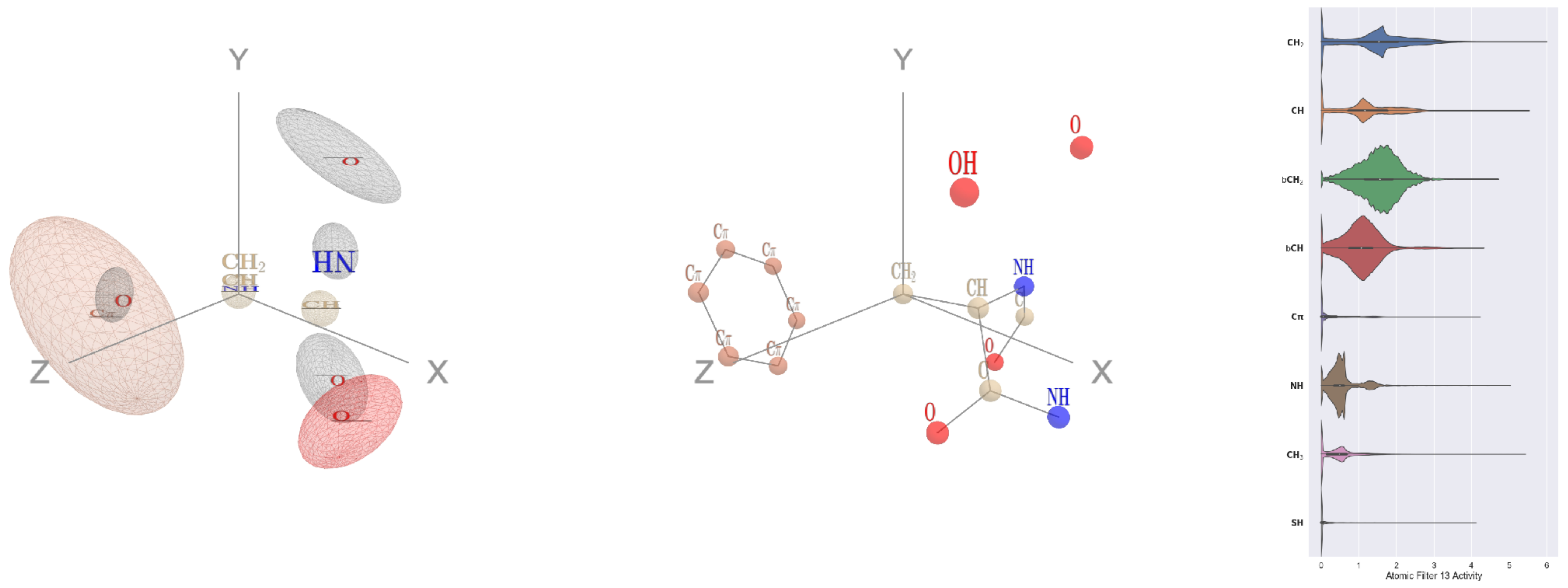


Filter 13
$R_{\text{pred}}^{\text{max}} = 0.00$
$R_{\text{pred}}^{\text{mean}} = -0.03$
Percent Active = 59
Pooling attention 0.34
Pooling feature 0.89
Top activators
4v4k:M:167 K CB
4d10:K:200 R CB
4u4c:A:599 I CG1
4buj:B:80 N CB
4hg6:A:306 H CB
2wk4:B:452 Q CG
1fpq:A:25 C CB
3qzy:B:170 I CG1
1qhd:A:1 M CB
1sl0:A:18 F CB

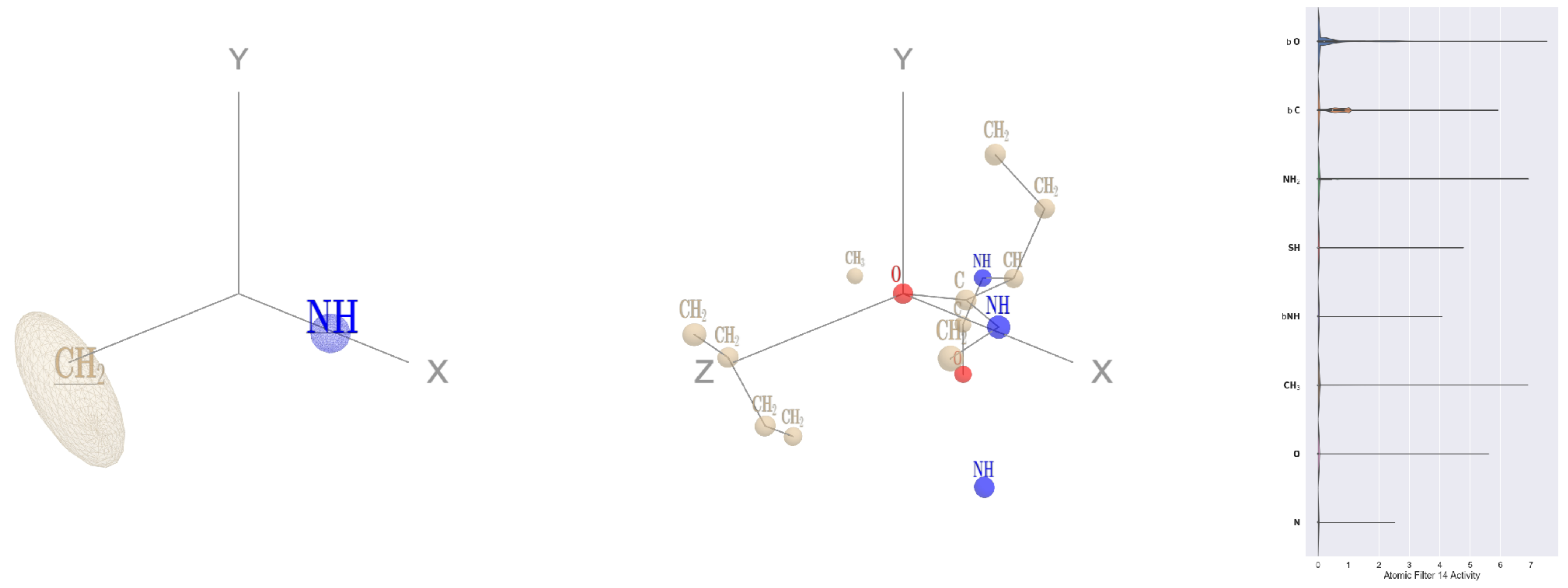


Filter 14
$R_{\text{pred}}^{\text{max}} = -0.01$
$R_{\text{pred}}^{\text{mean}} = -0.04$
Percent Active = 20
Pooling attention 0.12
Pooling feature 0.06
Top activators
1cm5:A:16 K O
3tj0:A:121 A O
1aom:A:529 S O
2hb6:A:415 K O
4r3z:C:429 G O
4s20:L:127 A O
1ewy:A:70 N O
3noy:B:41 N O
1lqb:C:124 T O
3ilk:A:71 D O

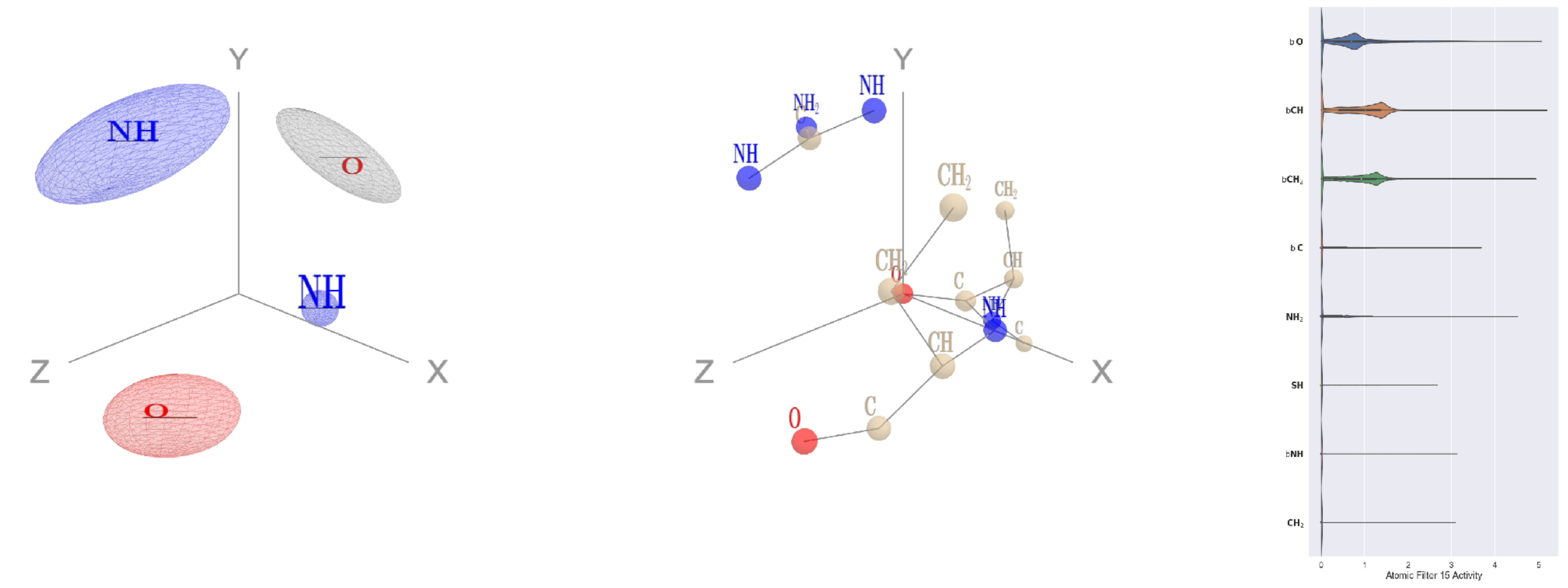


Filter 15
$R_{\text{pred}}^{\text{max}} = -0.03$
$R_{\text{pred}}^{\text{mean}} = -0.06$
Percent Active = 32
Pooling attention 0.09
Pooling feature 0.17
Top activators
4f52:E:433 K O
2pif:B:7 A O
4d10:K:11 S O
1izl:D:325 I O
1qvr:A:97 L O
1cm5:A:412 D O
4bxf:A:347 G O
3s9m:C:217 K O
1w36:B:721 N O
3bma:C:240 Y O

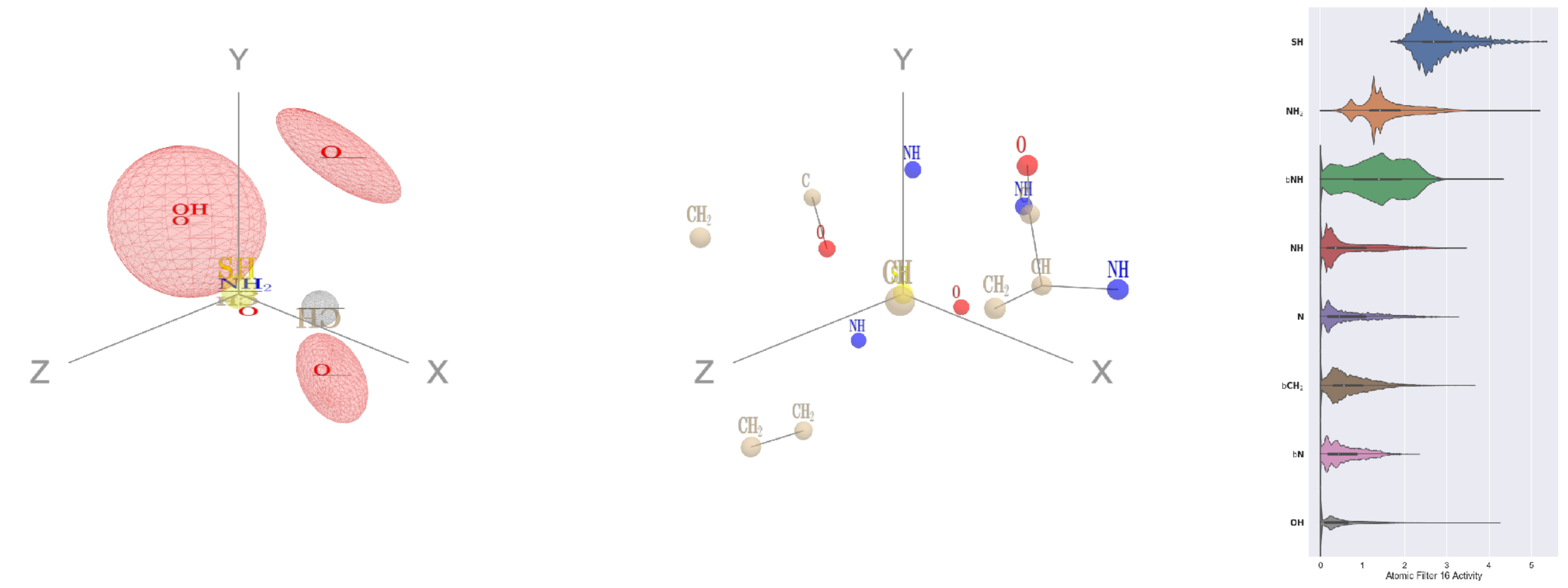

Filter 16
$R_{\text{pred}}^{\text{max}} = -0.05$
$R_{\text{pred}}^{\text{mean}} = -0.06$
Percent Active = 43
Pooling attention 0.22
Pooling feature 0.82
Top activators
1pgu:B:286 C SG
3kyh:C:192 C SG
4r3z:B:293 C SG
1k72:B:560 C SG
1r7r:A:105 C SG
1qha:A:672 C SG
2zy4:F:132 C SG
1f4a:A:500 C SG
4jds:A:288 C SG
4a7z:A:643 C SG

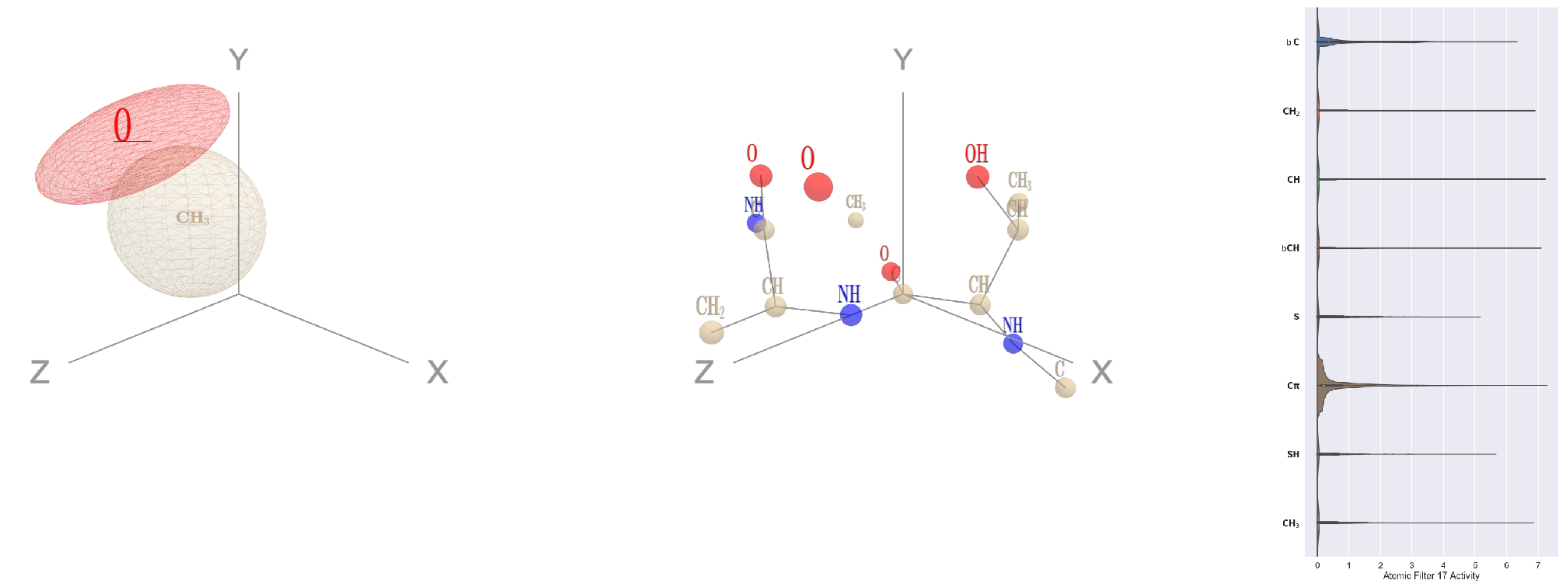

Filter 17
$R_{\text{pred}}^{\text{max}} = -0.07$
$R_{\text{pred}}^{\text{mean}} = -0.12$
Percent Active = 40
Pooling attention 0.40
Pooling feature 0.15
Top activators
3b8e:A:50 T C
4g1m:B:573 L C
1mp9:A:95 G C
4gu0:A:173 P C
2b0o:F:622 G C
4v88:AF:39 E C
3b8m:A:159 A C
4e0s:B:113 K C
3a5c:G:54 D C
4mex:C:438 G C

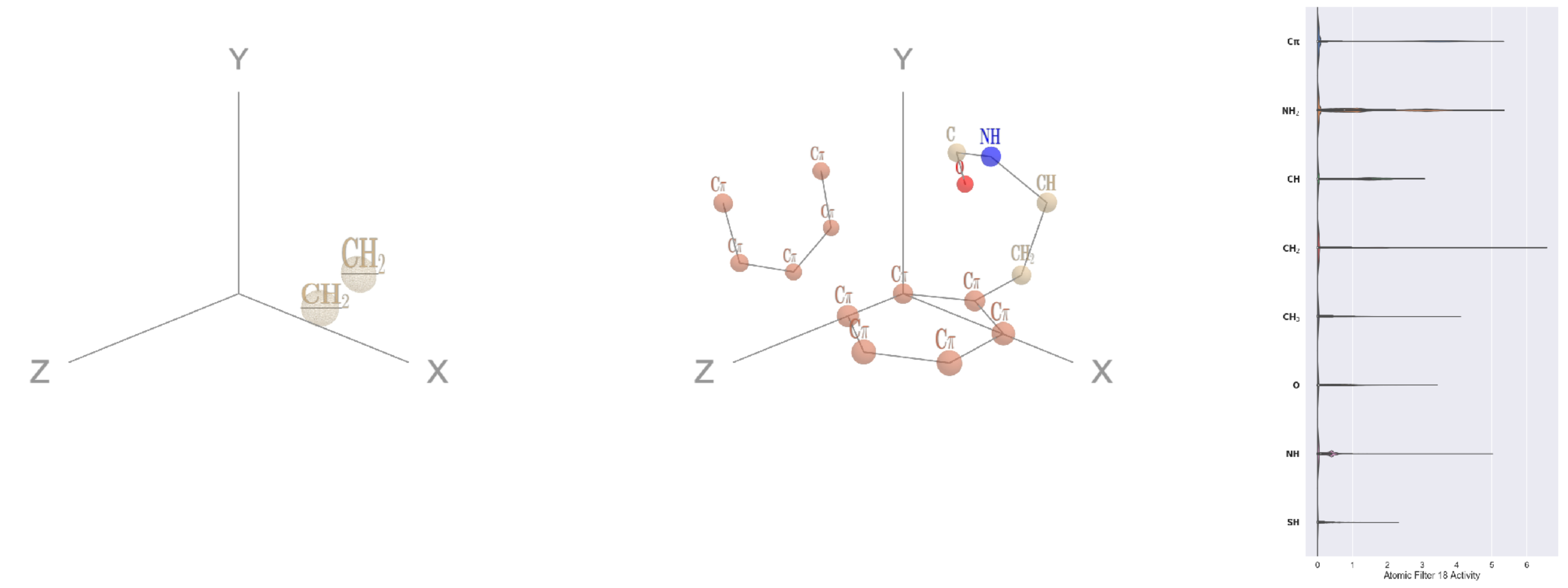

Filter 18
$R_{\text{pred}}^{\text{max}} = 0.08$
$R_{\text{pred}}^{\text{mean}} = 0.06$
Percent Active = 25
Pooling attention 0.19
Pooling feature 0.67
Top activators
1b8a:A:181 F CD2
1j5w:B:221 Y CD1
2fa0:A:190 Y CD2
3h1s:B:10 Y CD1
1ivu:A:512 Y CD1
1ko6:A:831 Y CD2
1q18:A:81 F CD1
1vr0:A:198 Y CD2
1ezv:C:184 Y CD2
4l8n:A:133 Y CD1

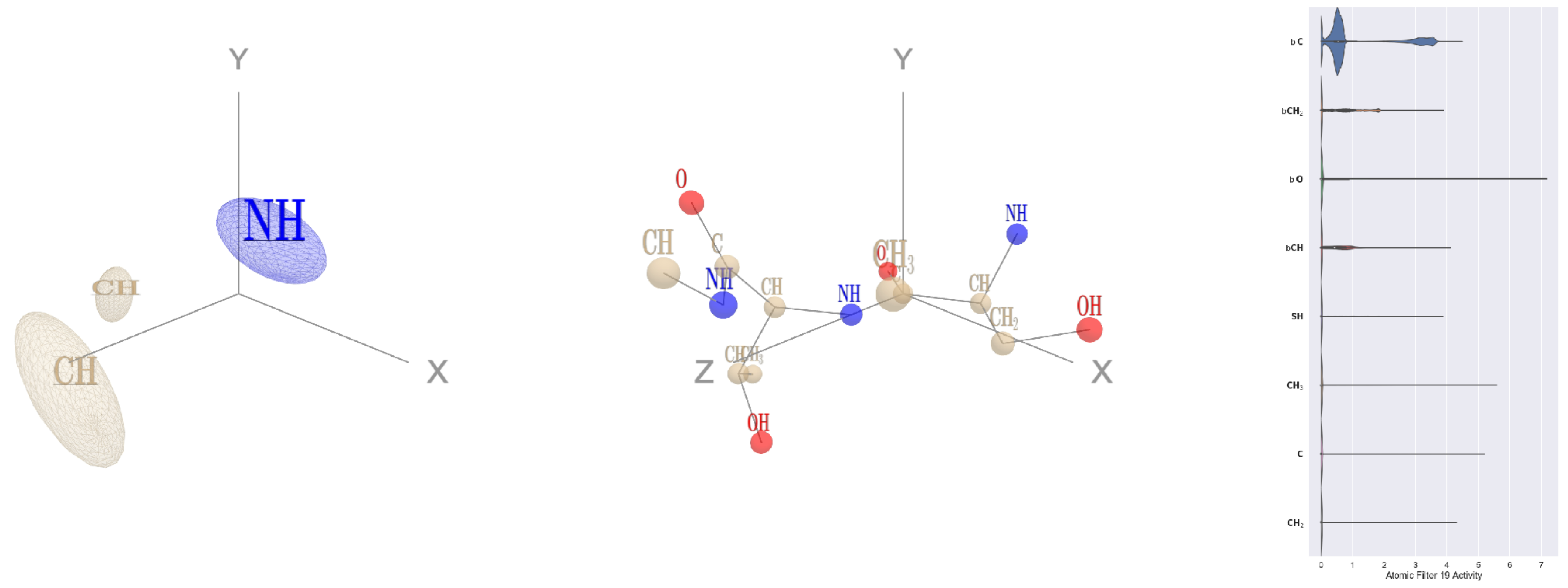

Filter 19
$R_{\text{pred}}^{\text{max}} = -0.03$
$R_{\text{pred}}^{\text{mean}} = -0.07$
Percent Active = 30
Pooling attention 0.13
Pooling feature 0.13
Top activators
1e1c:A:2 S C
3ax7:A:590 A C
2gy7:B:290 A C
2d73:A:55 V C
1cm5:A:100 Q C
2wsc:B:296 G C
4wj3:B:361 E C
1umg:A:42 I C
4pht:A:324 D C
4pbw:B:257 N C

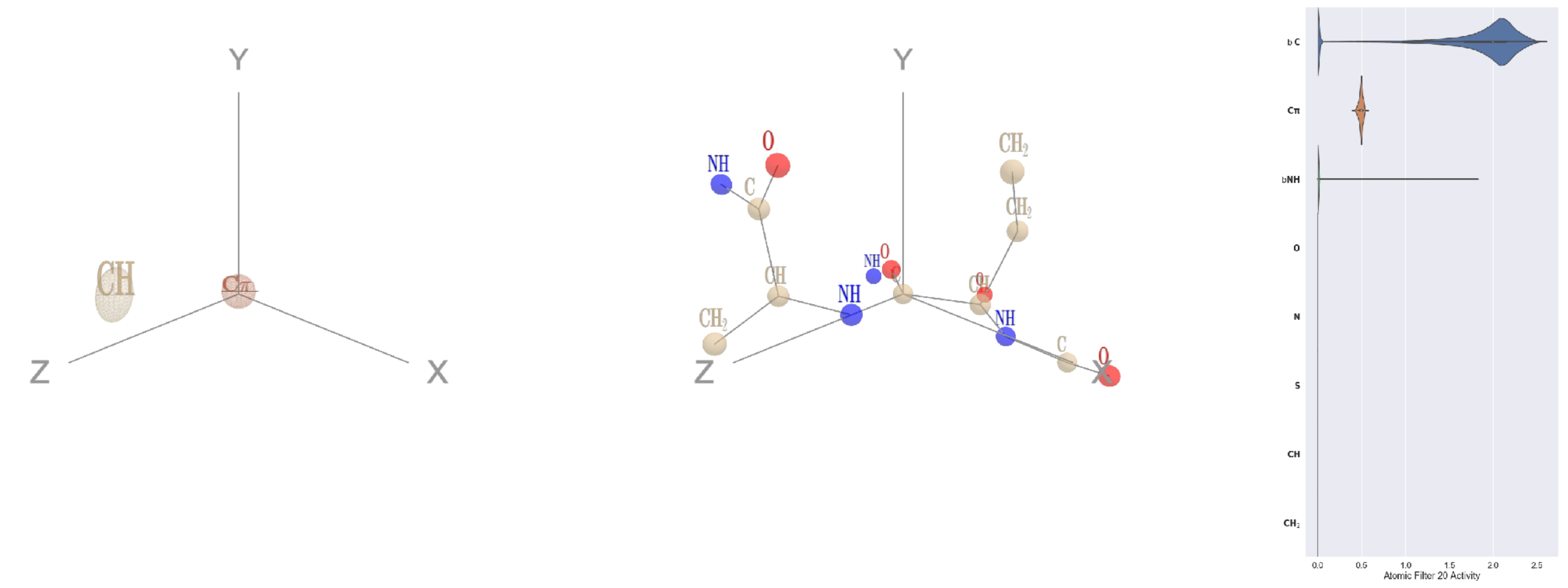

Filter 20
$R_{\text{pred}}^{\text{max}} = -0.01$
$R_{\text{pred}}^{\text{mean}} = -0.03$
Percent Active = 21
Pooling attention 0.10
Pooling feature 1.24
Top activators
1u5e:A:126 R C
3ako:B:161 I C
2fa0:A:10 M C
1u5e:A:46 E C
3gku:A:31 Y C
4u0u:A:180 R C
4cy6:A:486 S C
1vl4:A:144 Y C
1ivu:A:276 S C
1vl4:A:116 V C

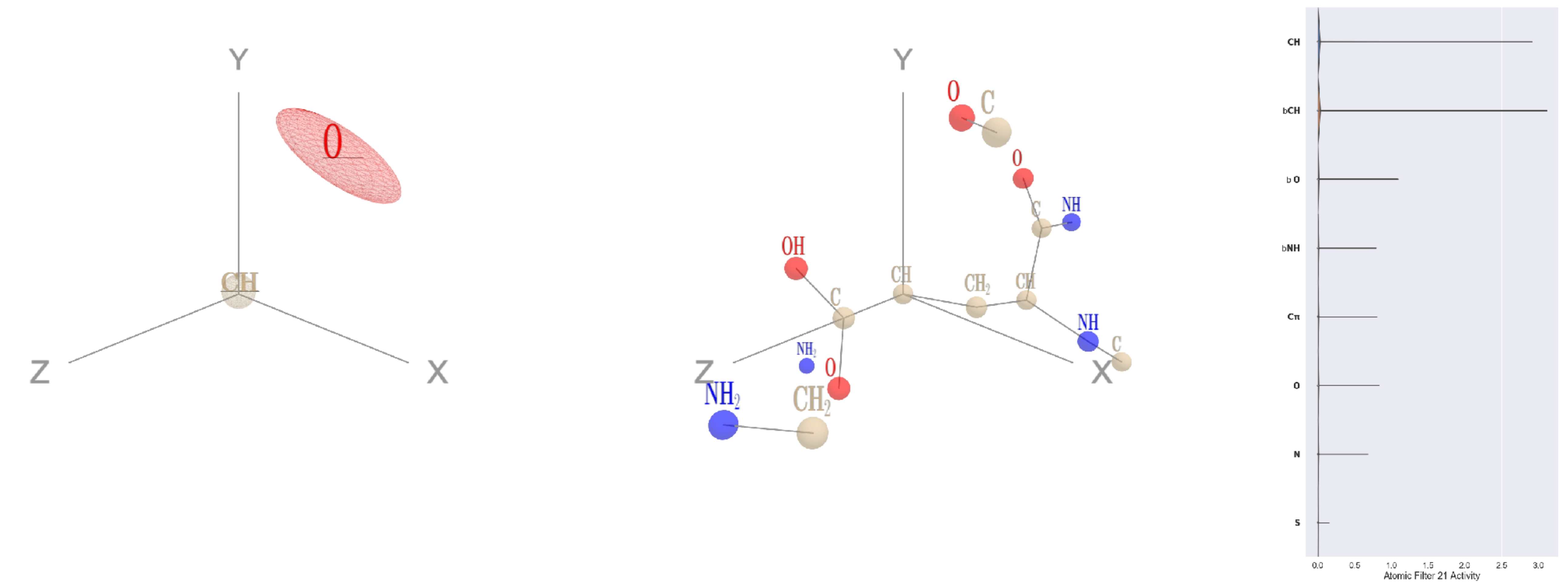

Filter 21
$R_{\text{pred}}^{\text{max}} = -0.01$
$R_{\text{pred}}^{\text{mean}} = -0.02$
Percent Active = 7
Pooling attention 0.01
Pooling feature 0.01
Top activators
1p4e:B:221 E CG
1p32:A:101 T CB
2pmz:D:121 V CB
2o3o:A:191 T CB
2qh9:A:36 T CB
4ny2:A:294 T CB
2x98:A:344 T CB
1e5d:A:67 I CB
3n99:G:194 T CB
1umg:A:134 T CB

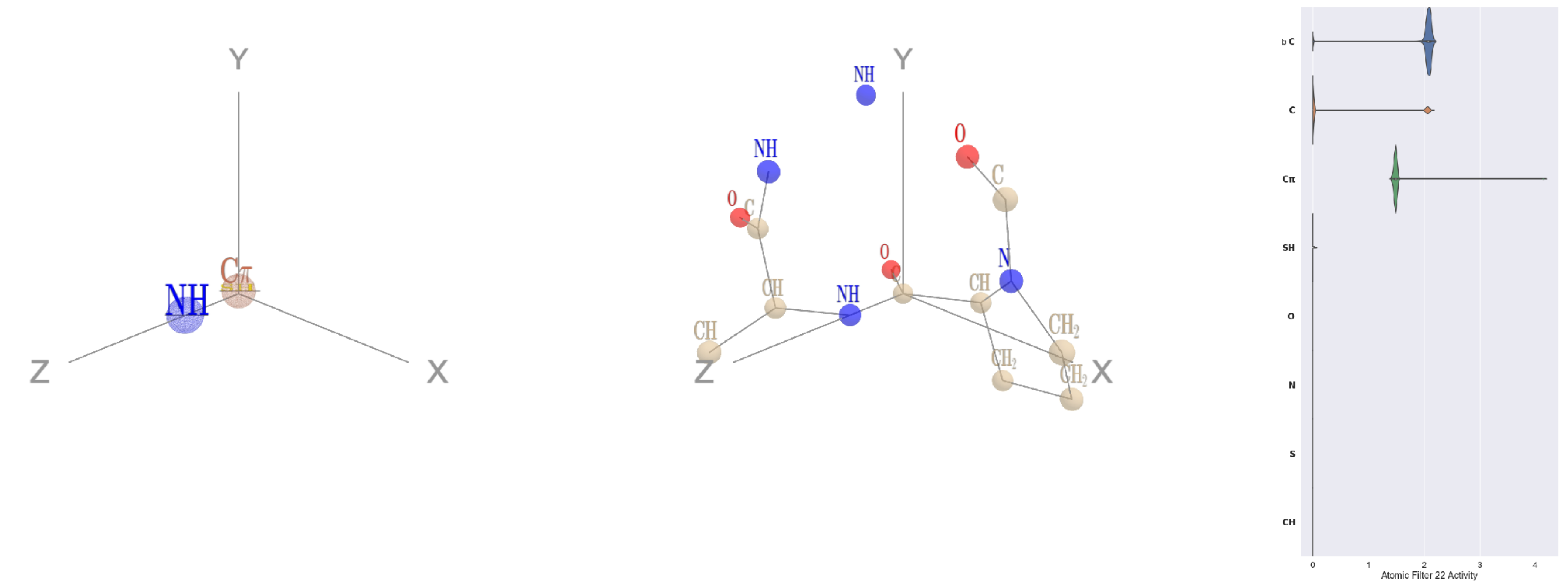

Filter 22
$R_{\text{pred}}^{\text{max}} = 0.02$
$R_{\text{pred}}^{\text{mean}} = 0.04$
Percent Active = 21
Pooling attention 0.15
Pooling feature 0.99
Top activators
4nnq:B:243 P C
4u0u:A:371 R C
4c0b:B:242 Q C
4pfp:A:163 F C
5ui2:A:93 P C
4ehi:B:181 Y C
1e1c:A:219 R C
4ehi:B:81 H C
3o47:A:114 K C
3l09:B:206 R C

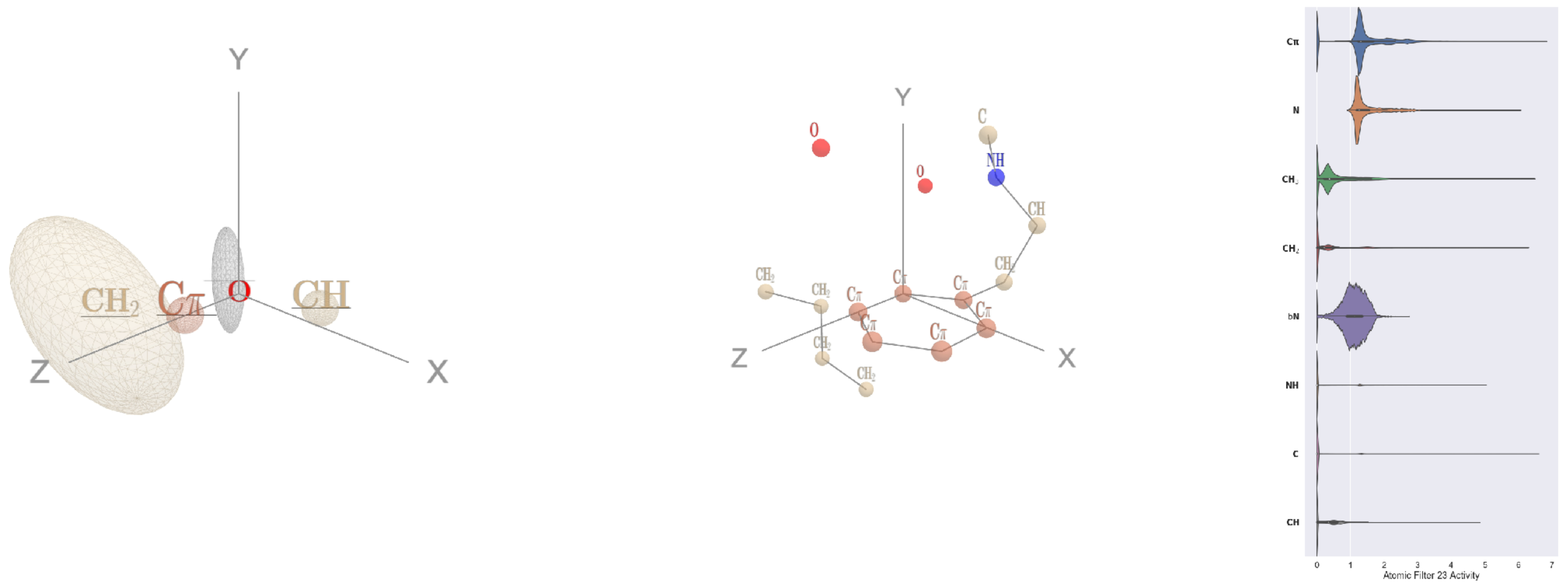

Filter 23
$R_{\text{pred}}^{\text{max}} = -0.04$
$R_{\text{pred}}^{\text{mean}} = -0.01$
Percent Active = 37
Pooling attention 0.05
Pooling feature 0.35
Top activators
4oph:A:150 F CD2
1ii7:A:331 W CZ2
4kzx:E:130 F CD1
3bof:A:254 F CE1
1he8:A:713 F CZ
1fx0:B:298 Y CD2
2b5d:X:212 W CZ2
4v8p:BC:29 F CD1
3hfd:A:49 Y CZ
2iwh:B:811 Y CZ

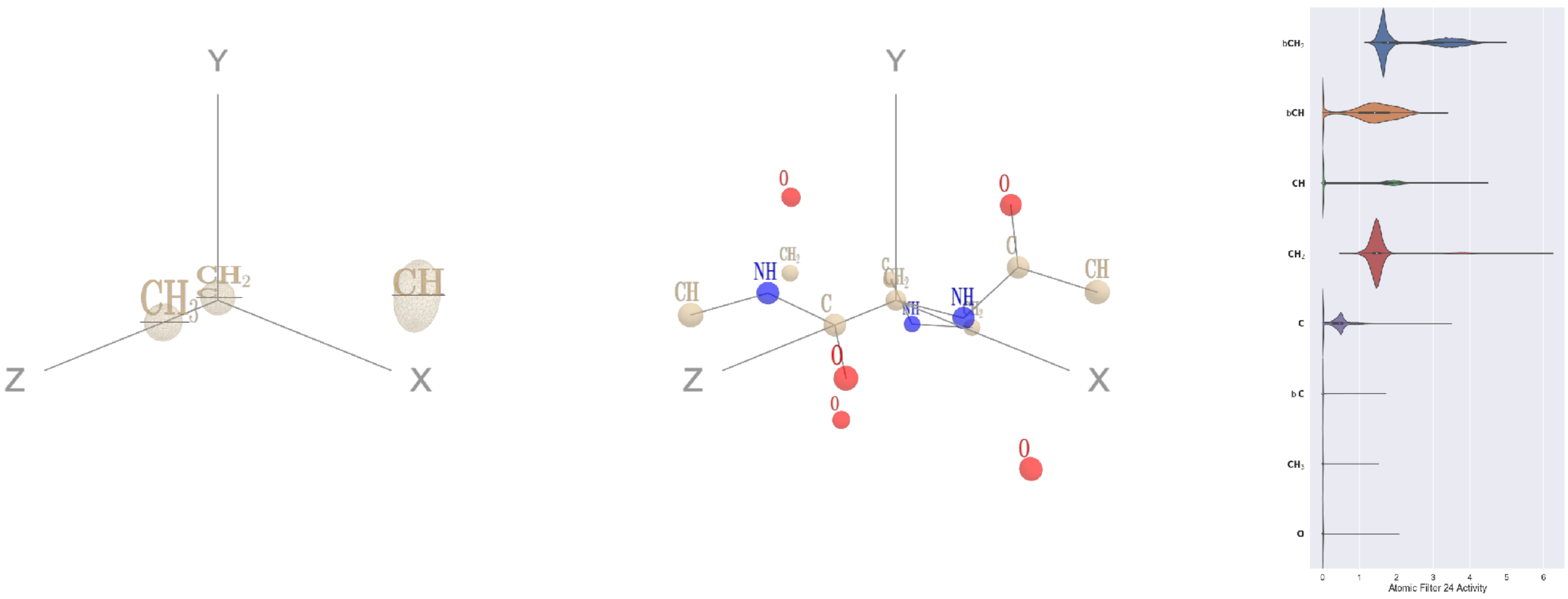

Filter 24
$R_{\mathrm{pred}}^{\mathrm{max}} = -0.02$
$R_{\mathrm{pred}}^{\mathrm{mean}} = -0.02$
Percent Active = 34
Pooling attention 0.22
Pooling feature 1.18
Top activators
1h2i:A:125 G CA
1lnl:A:33 G CA
4a7z:A:56 G CA
1ei6:A:350 G CA
1bxr:A:432 G CA
1f4a:A:605 G CA
3zxu:A:136 G CA
2nyg:A:156 G CA
3drw:A:285 G CA
3n6r:G:434 G CA

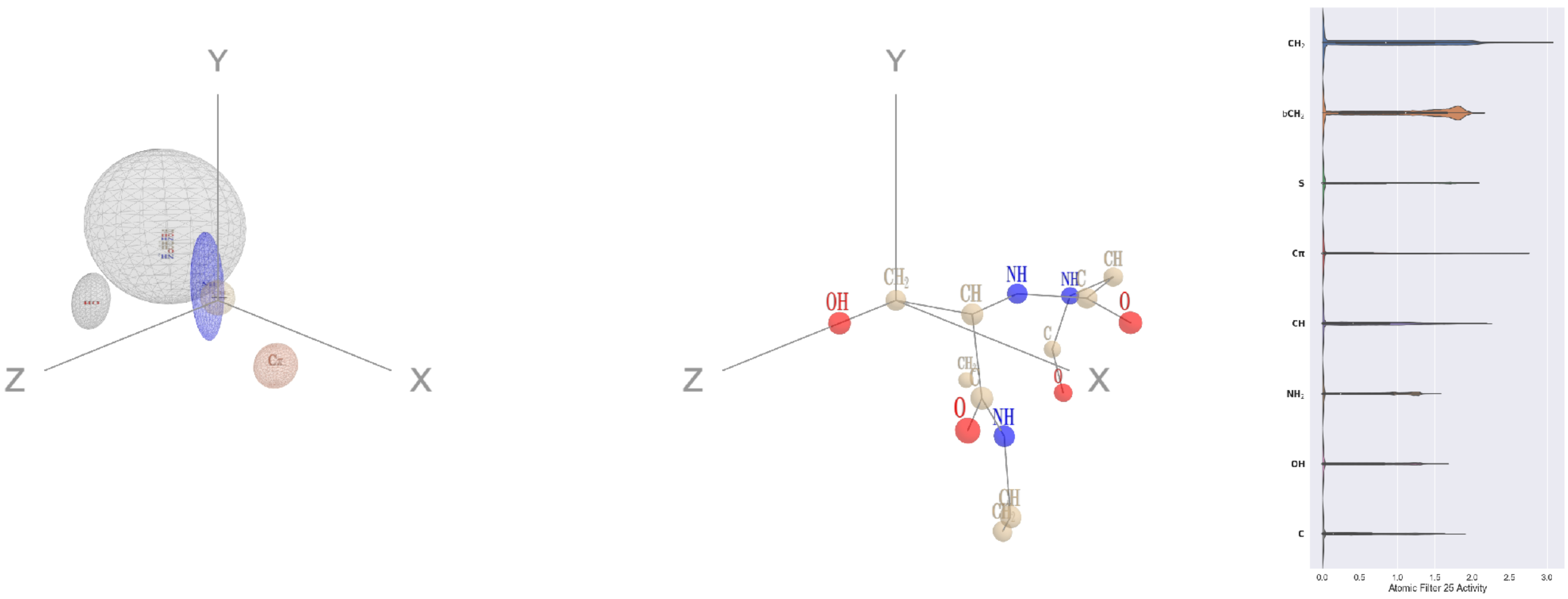

Filter 25
$R_{\mathrm{pred}}^{\mathrm{max}} = 0.28$
$R_{\mathrm{pred}}^{\mathrm{mean}} = 0.32$
Percent Active = 40
Pooling attention 0.83
Pooling feature 1.47
Top activators
4u7u:M:224 S CB
1qvr:A:85 S CB
1v8f:A:249 S CB
4ddg:A:150 S CB
3dlb:A:328 S CB
3h1s:B:17 S CB
2iwh:B:1 S CB
2cjr:A:313 S CB
4f52:E:397 S CB
4c2m:M:104 S CB

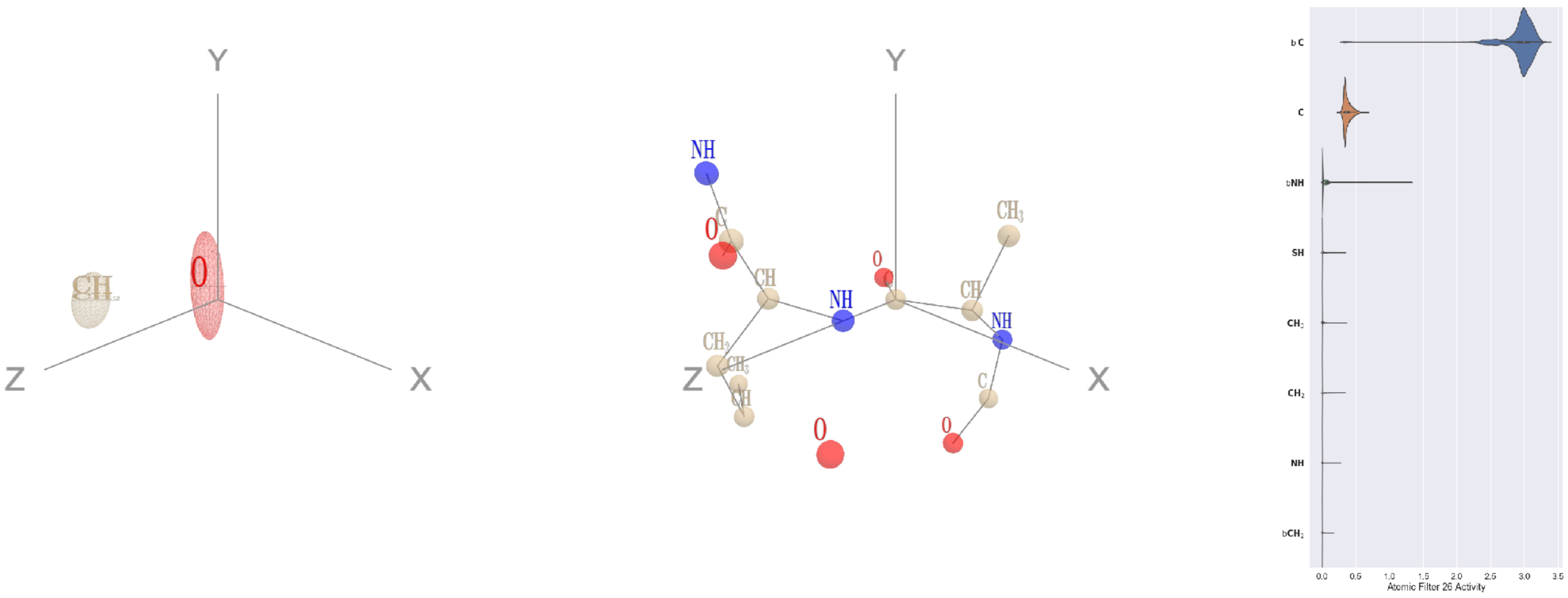

Filter 26
$R_{\mathrm{pred}}^{\mathrm{max}} = -0.02$
$R_{\mathrm{pred}}^{\mathrm{mean}} = -0.08$
Percent Active = 32
Pooling attention 0.27
Pooling feature 1.25
Top activators
2wgq:B:56 A C
3fbv:D:823 R C
4cy6:A:454 G C
2x0q:A:89 K C
3dlb:A:126 G C
3emf:A:160 T C
1ivu:A:276 S C
4asi:C:1920 C C
3rby:A:223 N C
3tj0:A:469 A C

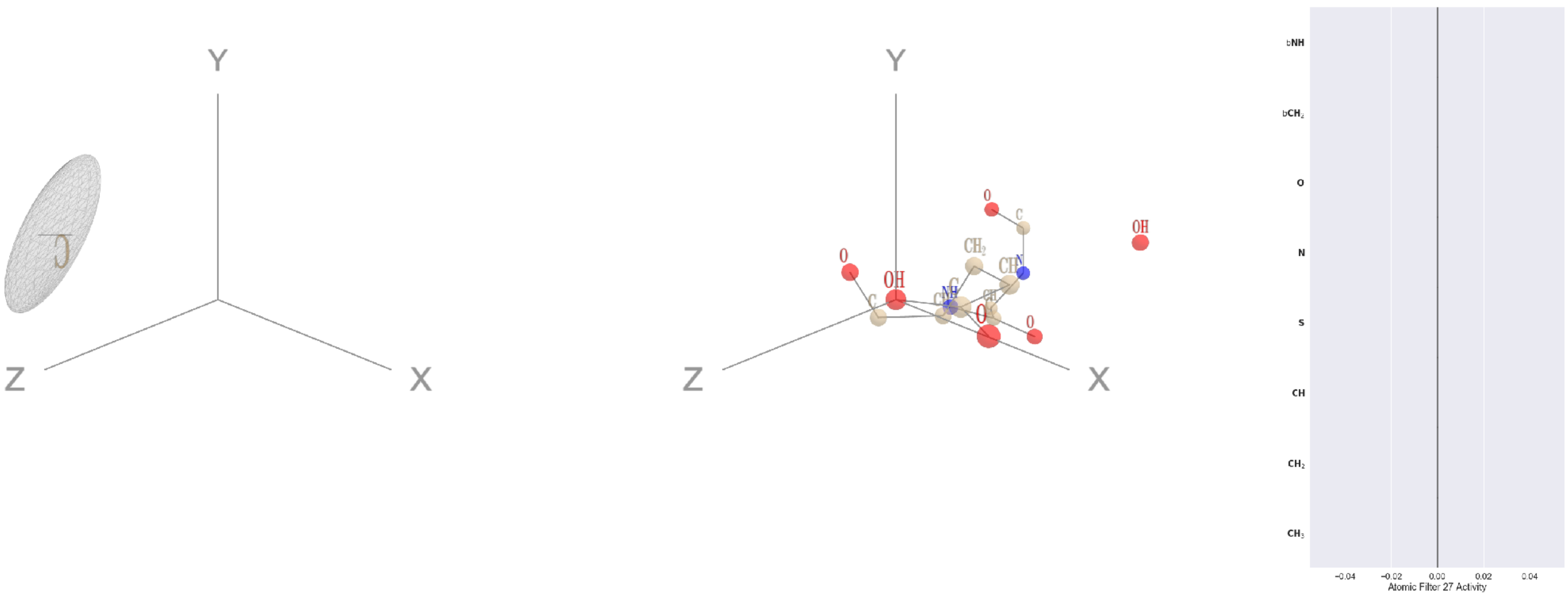

Filter 27
$R_{\mathrm{pred}}^{\mathrm{max}} = nan$
$R_{\mathrm{pred}}^{\mathrm{mean}} = nan$
Percent Active = 0
Pooling attention 0.00
Pooling feature 0.00
Top activators
5wy5:A:246 E OE2
2b0o:F:573 F N
2b0o:F:574 G O
2b0o:F:574 G C
2b0o:F:574 G CA
2b0o:F:574 G N
2b0o:F:573 F CZ
2b0o:F:573 F CE2
2b0o:F:573 F CE1
2b0o:F:573 F CD2

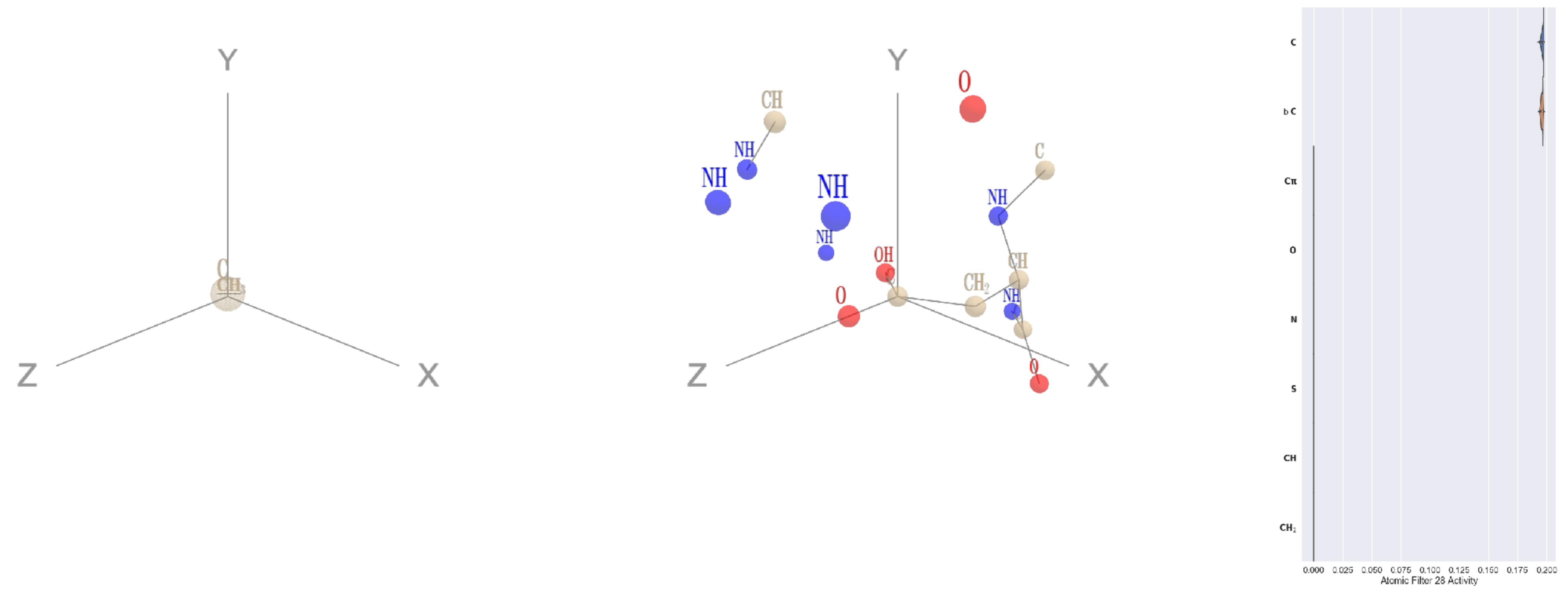


Filter 28
$R_{\mathrm{pred}}^{\mathrm{max}} = 0.03$
$R_{\mathrm{pred}}^{\mathrm{mean}} = -0.03$
Percent Active = 16
Pooling attention 0.00
Pooling feature 0.00
Top activators
1ii7:A:86 D CG
4wj3:B:356 N CG
4u4c:A:446 N CG
2iut:A:433 D CG
1ghq:C:106 N CG
3t5v:C:25 N CG
1lnl:A:53 N CG
1k90:C:684 D CG
4mex:C:23 D CG
1bxr:A:133 D CG

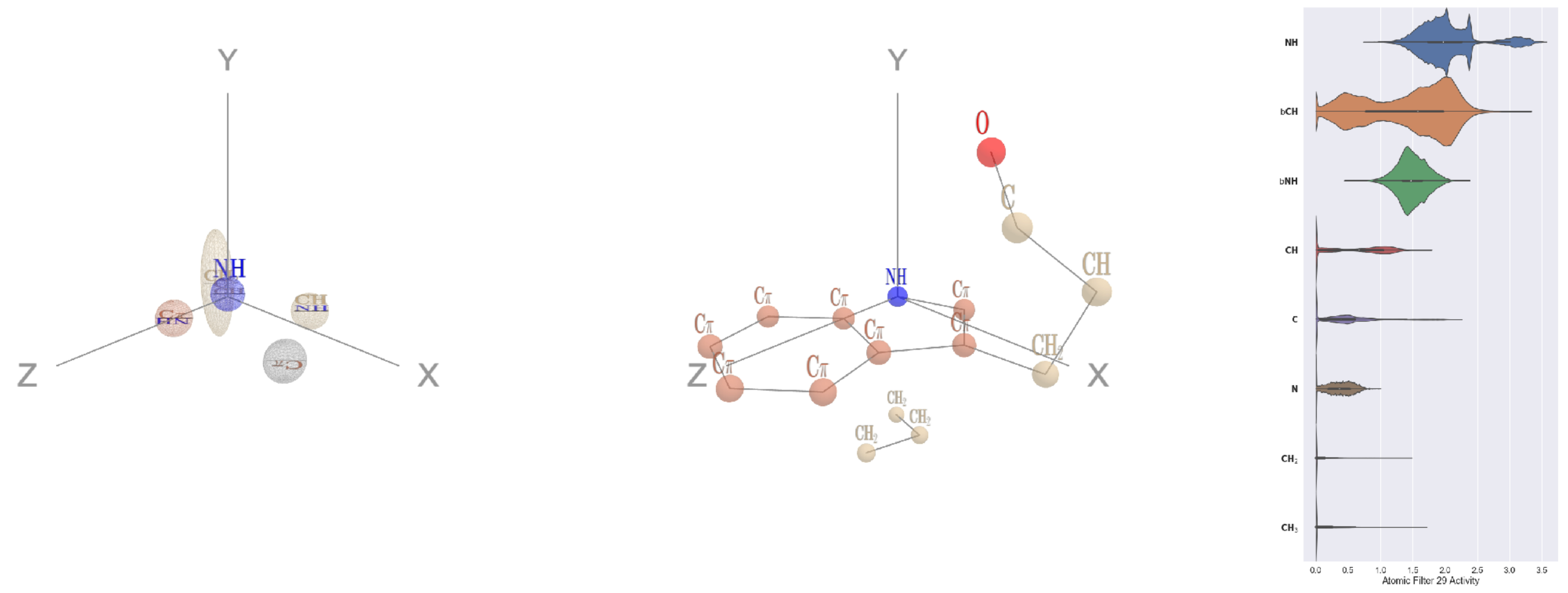


Filter 29
$R_{\mathrm{pred}}^{\mathrm{max}} = 0.06$
$R_{\mathrm{pred}}^{\mathrm{mean}} = 0.07$
Percent Active = 42
Pooling attention 0.09
Pooling feature 1.34
Top activators
3d00:A:115 W NE1
2idb:A:459 W NE1
1k90:C:513 W NE1
4a7z:A:585 W NE1
1jdw:A:280 W NE1
3c24:A:238 W NE1
2cge:A:300 W NE1
1aom:A:226 W NE1
1b8p:A:217 W NE1
4pk7:A:168 W NE1

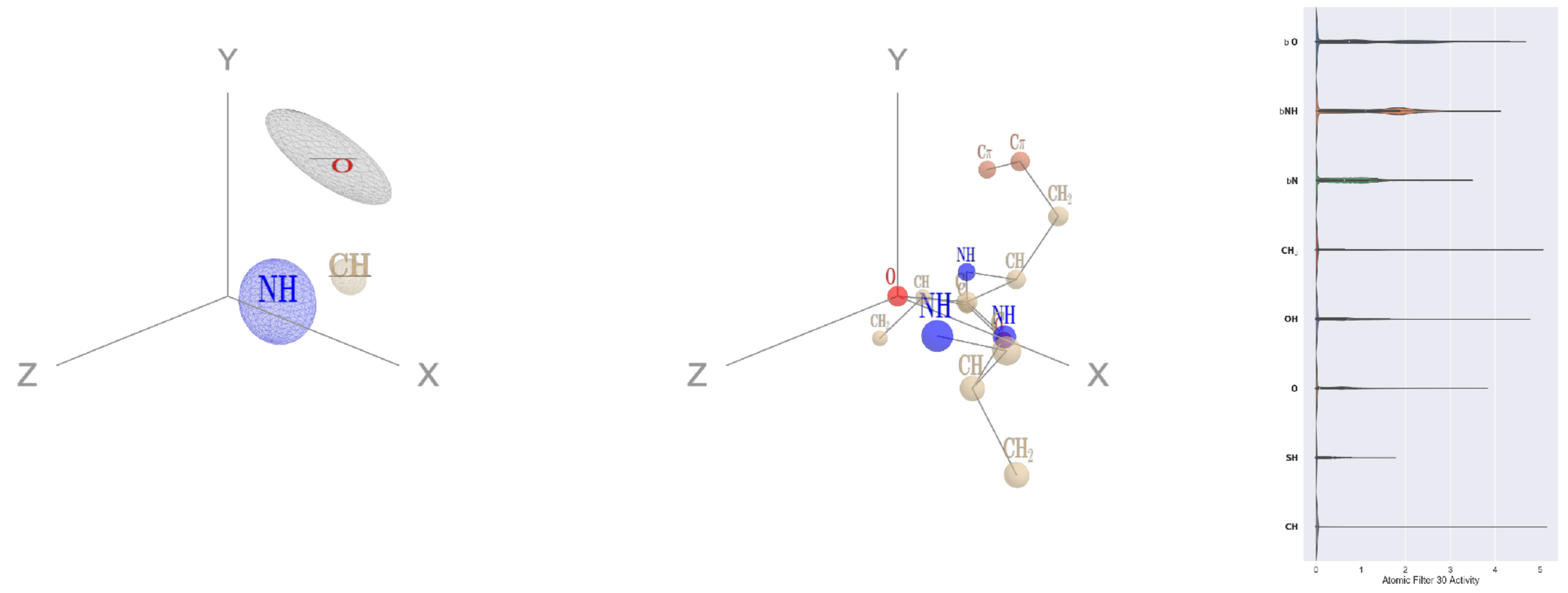


Filter 30
$R_{\mathrm{pred}}^{\mathrm{max}} = 0.03$
$R_{\mathrm{pred}}^{\mathrm{mean}} = 0.01$
Percent Active = 33
Pooling attention 0.20
Pooling feature 0.43
Top activators
4kzx:L:102 F O
1fx0:B:225 Q O
1u5e:A:139 K O
1p35:A:89 F O
1hfe:L:176 T O
1izn:B:164 T O
4a7z:A:176 V O
1vbk:A:117 T O
1p32:A:167 N O
4kzx:R:94 E O

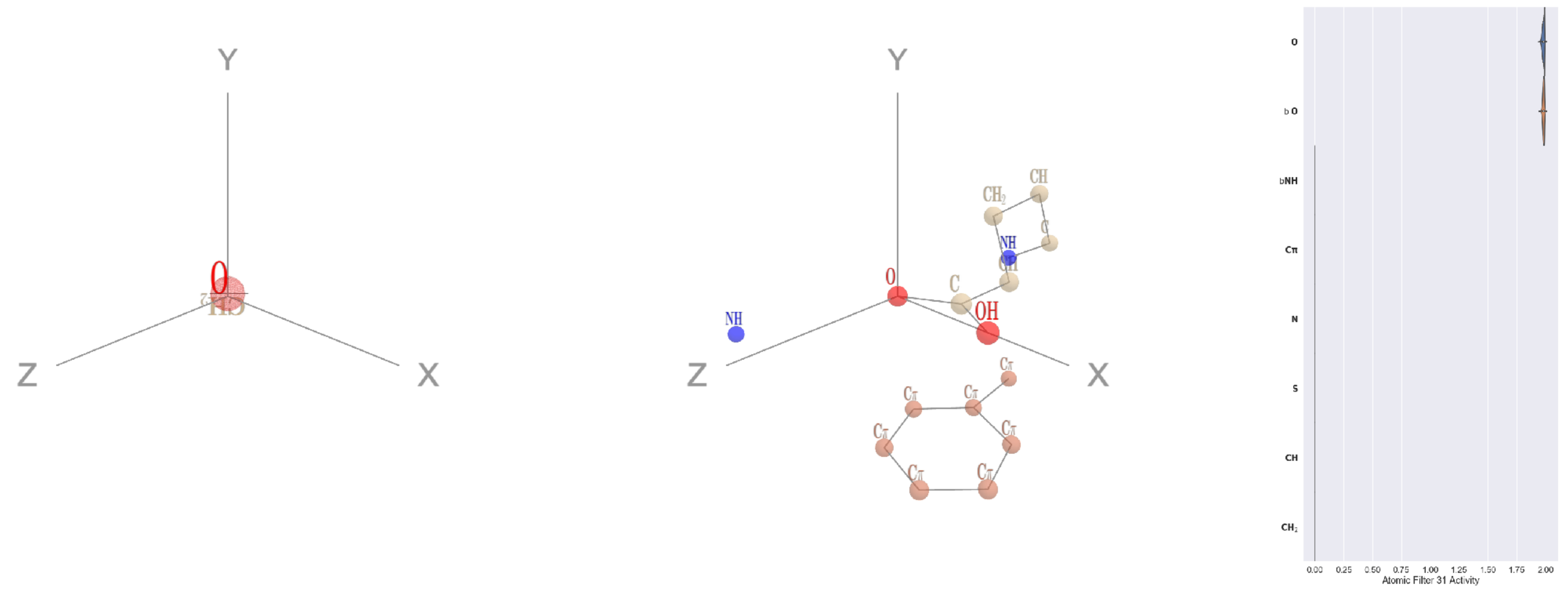


Filter 31
$R_{\mathrm{pred}}^{\mathrm{max}} = 0.05$
$R_{\mathrm{pred}}^{\mathrm{mean}} = -0.05$
Percent Active = 15
Pooling attention 0.08
Pooling feature 0.05
Top activators
2gy7:B:81 E OE1
4whj:A:502 E OE1
3o5t:A:28 E OE1
3di4:A:227 E OE1
4cy6:A:459 Q OE1
3bma:C:218 Q OE1
1d0n:A:679 Q OE1
1lnl:A:386 D OD1
2cge:A:301 E OE1
1sl0:A:159 E OE1

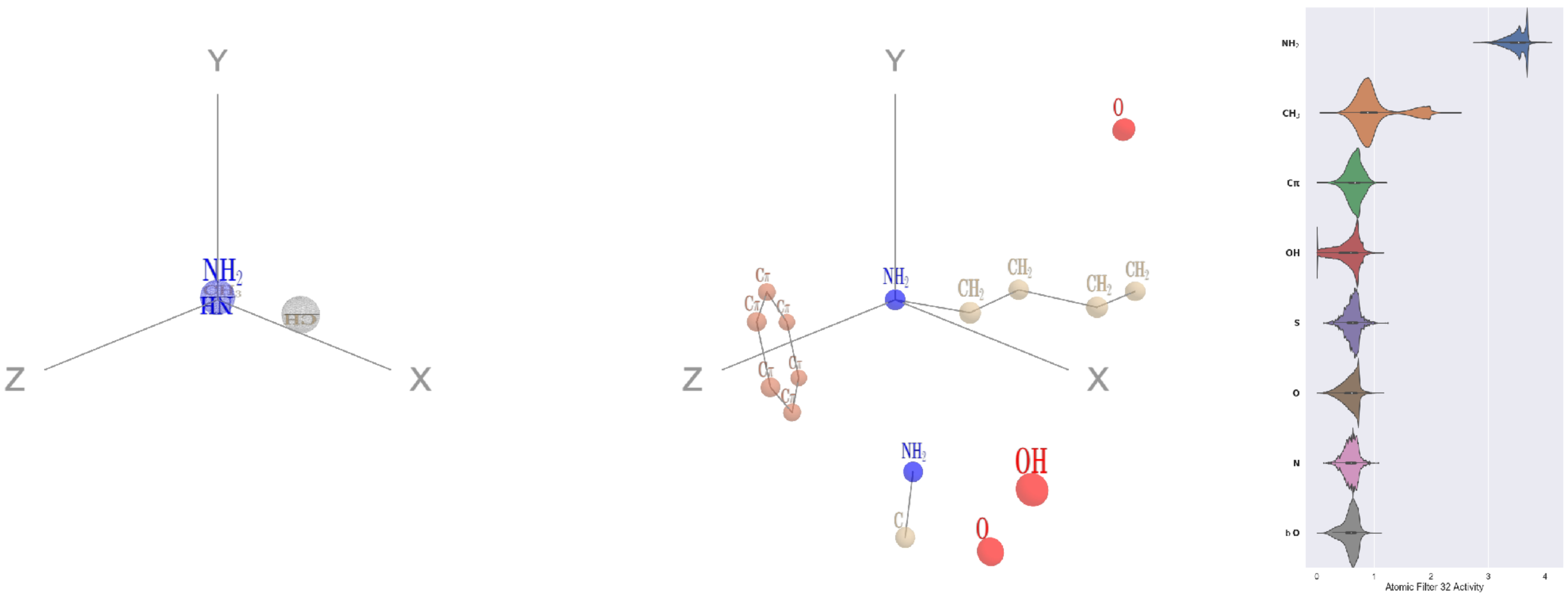


Filter 32
$R_{\text{pred}}^{\text{max}} = 0.07$
$R_{\text{pred}}^{\text{mean}} = 0.12$
Percent Active = 67
Pooling attention 0.25
Pooling feature 0.51
Top activators
4v4k:M:582 K NZ
1aom:A:285 R NH2
2dyo:A:208 R NH2
2x7x:A:172 R NH2
5hu6:C:228 K NZ
3gw6:F:837 R NH2
5kqv:E:409 R NH2
4gt2:A:249 Q NE2
4g1m:B:611 K NZ
3ne5:C:368 R NH2

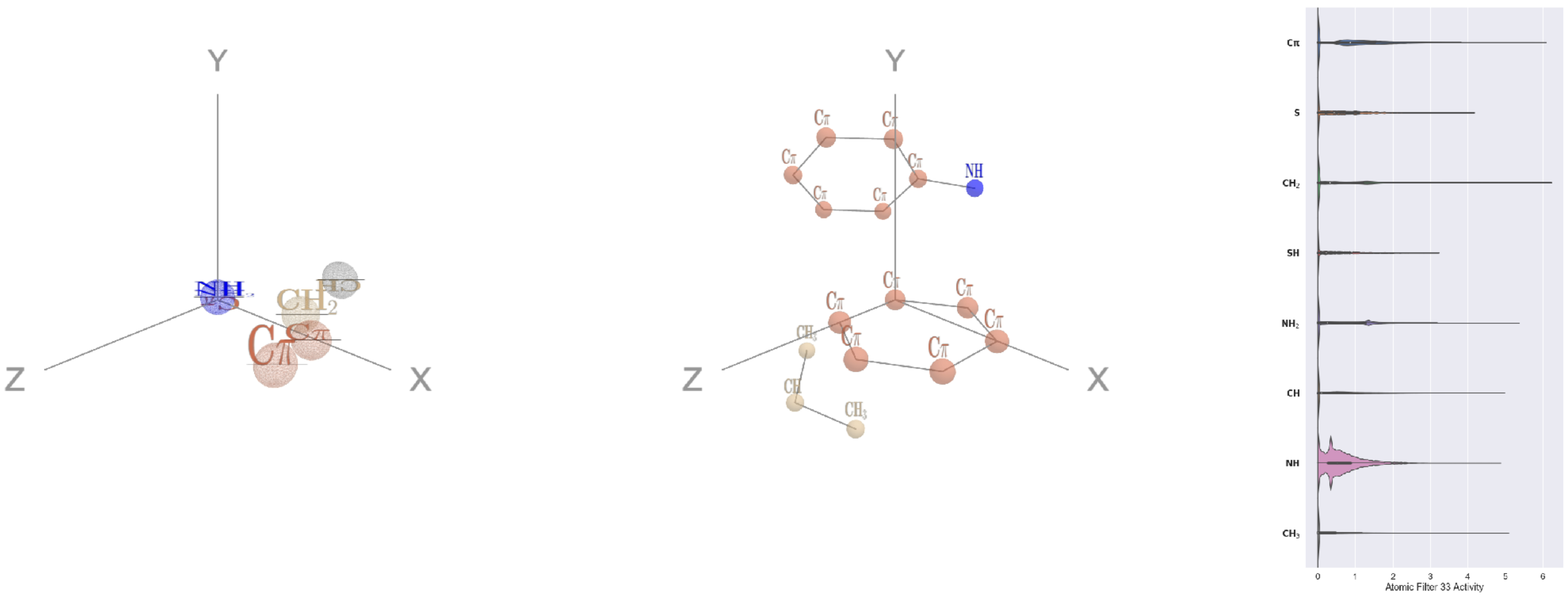


Filter 33
$R_{\text{pred}}^{\text{max}} = -0.06$
$R_{\text{pred}}^{\text{mean}} = -0.08$
Percent Active = 39
Pooling attention 0.06
Pooling feature 0.81
Top activators
4k1n:A:423 F CZ
3irm:A:345 F CE2
1lnl:A:340 F CE1
1srq:A:236 F CZ
3tj0:A:390 F CE2
4rap:E:327 F CD2
4kzx:E:86 F CE2
4buj:B:264 F CE1
1bcp:A:178 Y CD1
3pq1:A:445 F CE2

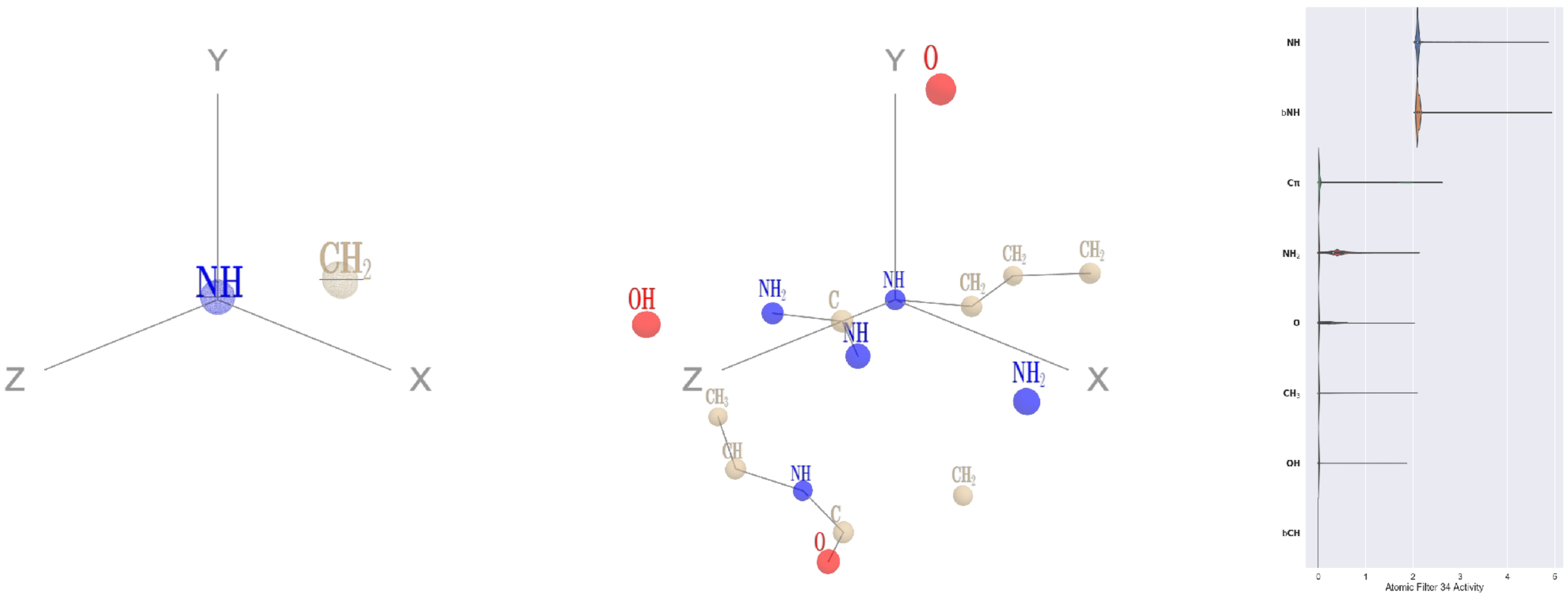


Filter 34
$R_{\text{pred}}^{\text{max}} = 0.01$
$R_{\text{pred}}^{\text{mean}} = 0.01$
Percent Active = 22
Pooling attention 0.15
Pooling feature 0.03
Top activators
4rcn:B:608 R NE
3dlb:A:486 R NE
4v88:AF:102 R NE
3do6:A:72 R NE
4d0l:A:163 R NE
1h4r:A:198 R NE
4asi:C:2258 R NE
3drw:A:392 R NE
4qiw:A:446 R NE
2hh1:H:154 R NE

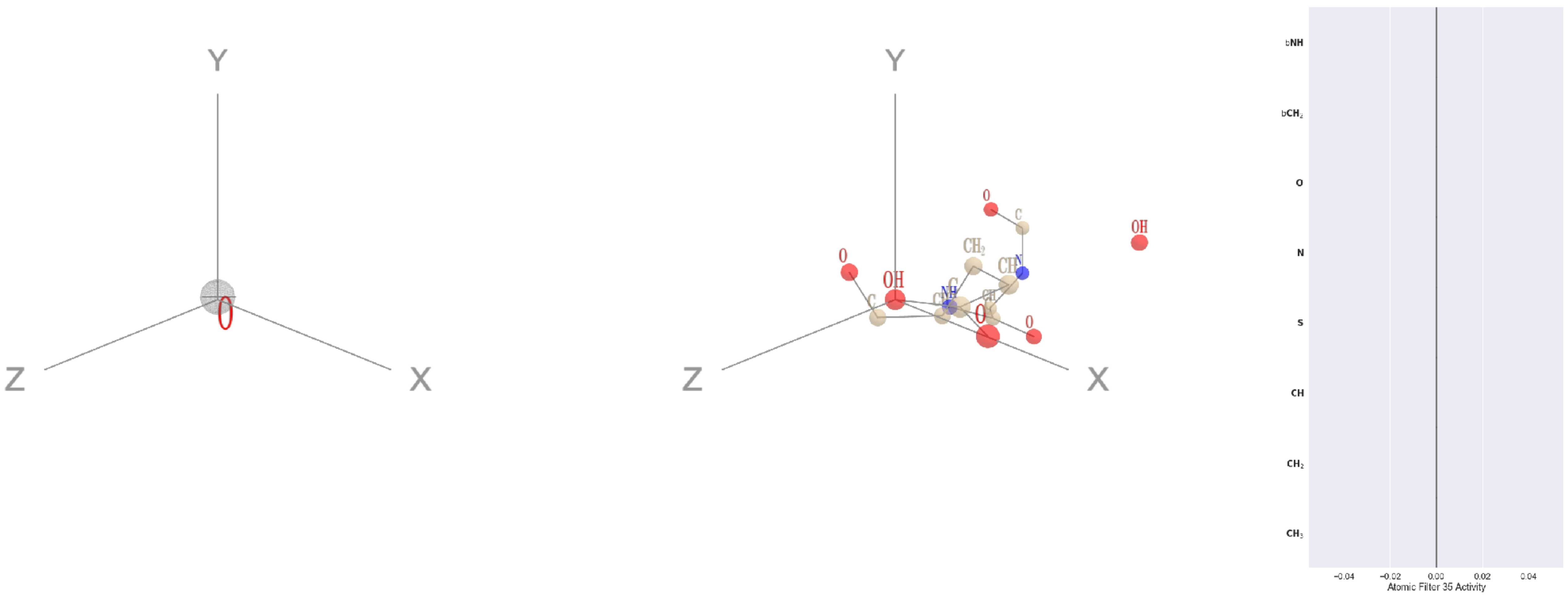


Filter 35
$R_{\text{pred}}^{\text{max}} = nan$
$R_{\text{pred}}^{\text{mean}} = nan$
Percent Active = 0
Pooling attention 0.00
Pooling feature 0.00
Top activators
5wy5:A:246 E OE2
2b0o:F:573 F N
2b0o:F:574 G O
2b0o:F:574 G C
2b0o:F:574 G CA
2b0o:F:574 G N
2b0o:F:573 F CZ
2b0o:F:573 F CE2
2b0o:F:573 F CE1
2b0o:F:573 F CD2

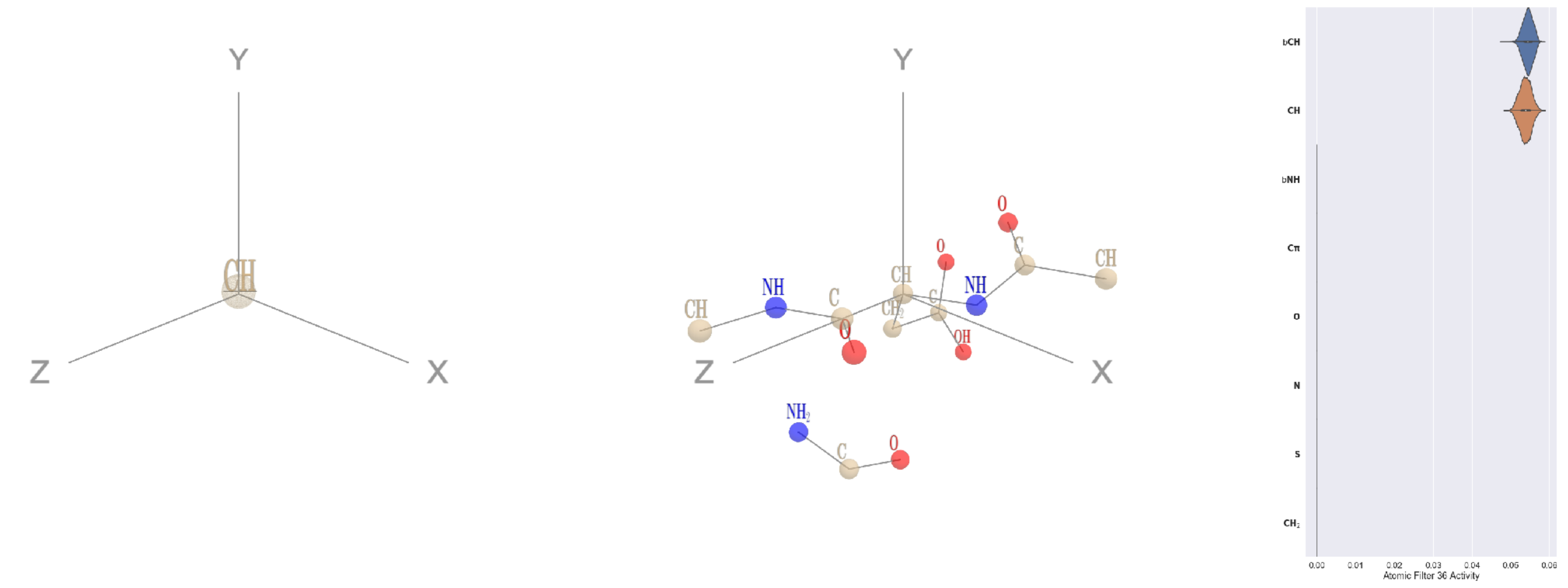


Filter 36
$R_{\text{pred}}^{\text{max}} = 0.03$
$R_{\text{pred}}^{\text{mean}} = -0.04$
Percent Active = 16
Pooling attention 0.00
Pooling feature 0.00
Top activators
3b8e:A:893 D CA
3l0o:A:224 R CA
1i3r:B:176 N CA
4cvw:A:274 D CA
1bh3:A:289 F CA
1a92:A:47 D CA
2idb:A:143 N CA
4r3z:B:385 D CA
4kwb:A:278 D CA
2qn5:B:42 R CA

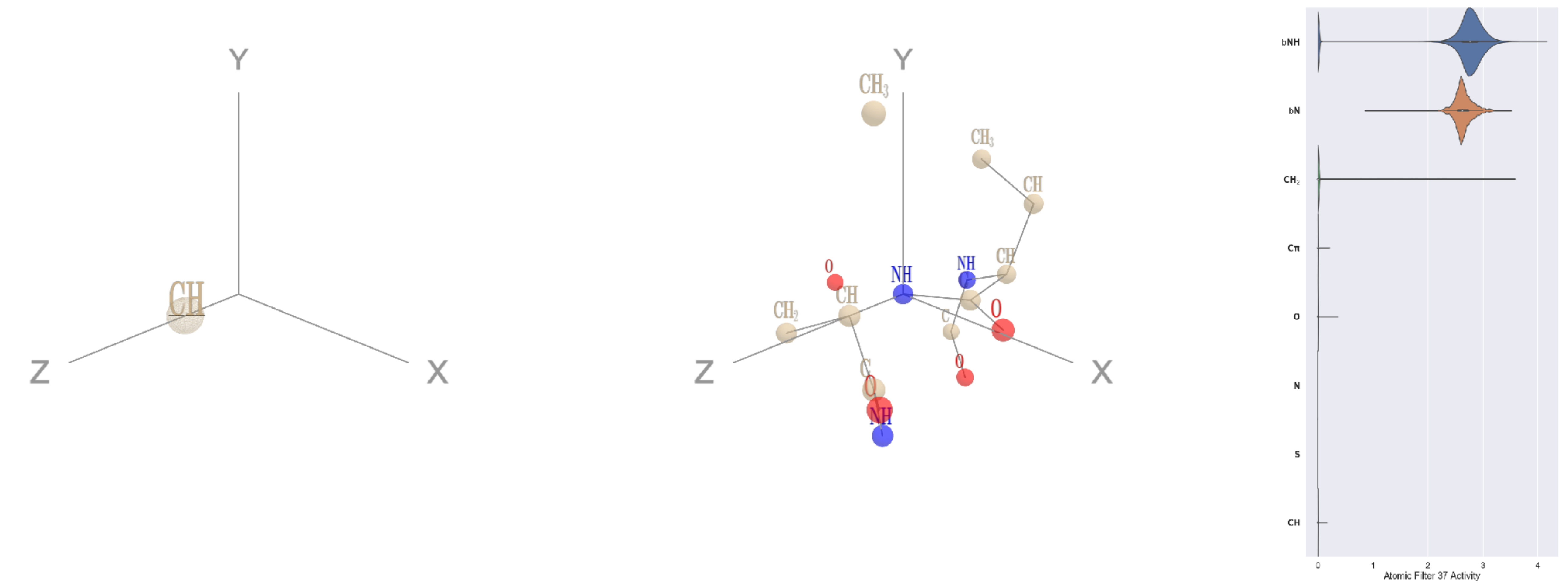


Filter 37
$R_{\text{pred}}^{\text{max}} = 0.02$
$R_{\text{pred}}^{\text{mean}} = -0.04$
Percent Active = 14
Pooling attention 0.23
Pooling feature 0.04
Top activators
1lnl:A:208 F N
1u5e:A:48 L N
1u5e:A:17 I N
4kzx:L:20 K N
3a5c:G:175 Q N
3a5c:G:144 E N
1qc9:A:146 S N
1e1c:A:679 D N
1u5e:A:40 K N
1lnl:A:47 C N

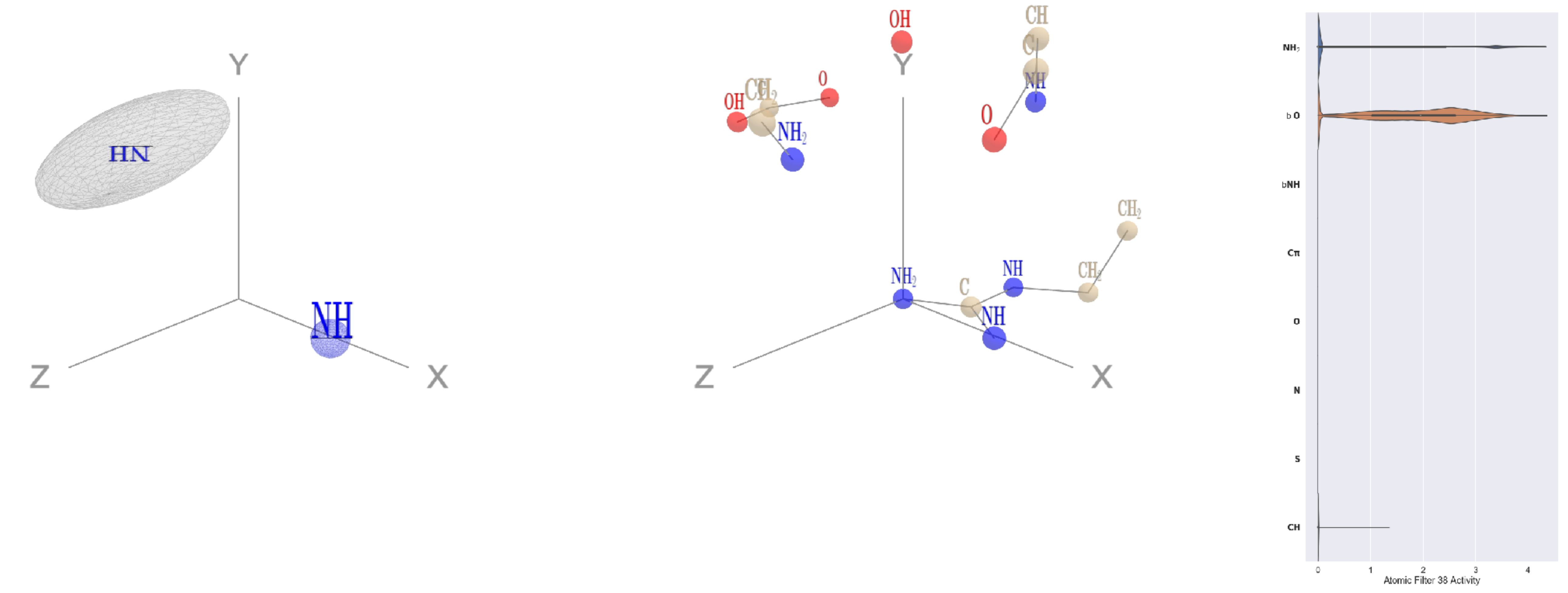


Filter 38
$R_{\text{pred}}^{\text{max}} = 0.02$
$R_{\text{pred}}^{\text{mean}} = -0.01$
Percent Active = 12
Pooling attention 0.21
Pooling feature 0.03
Top activators
1dtn:A:10 R NH2
1bxr:A:294 R NH2
1bxr:A:131 R NH2
3oru:A:195 R NH2
1f4a:A:425 R NH2
1bxr:A:509 R NH2
1q18:A:16 R NH2
3drw:A:369 R NH2
1bxr:A:998 R NH2
1ei6:A:130 R NH2

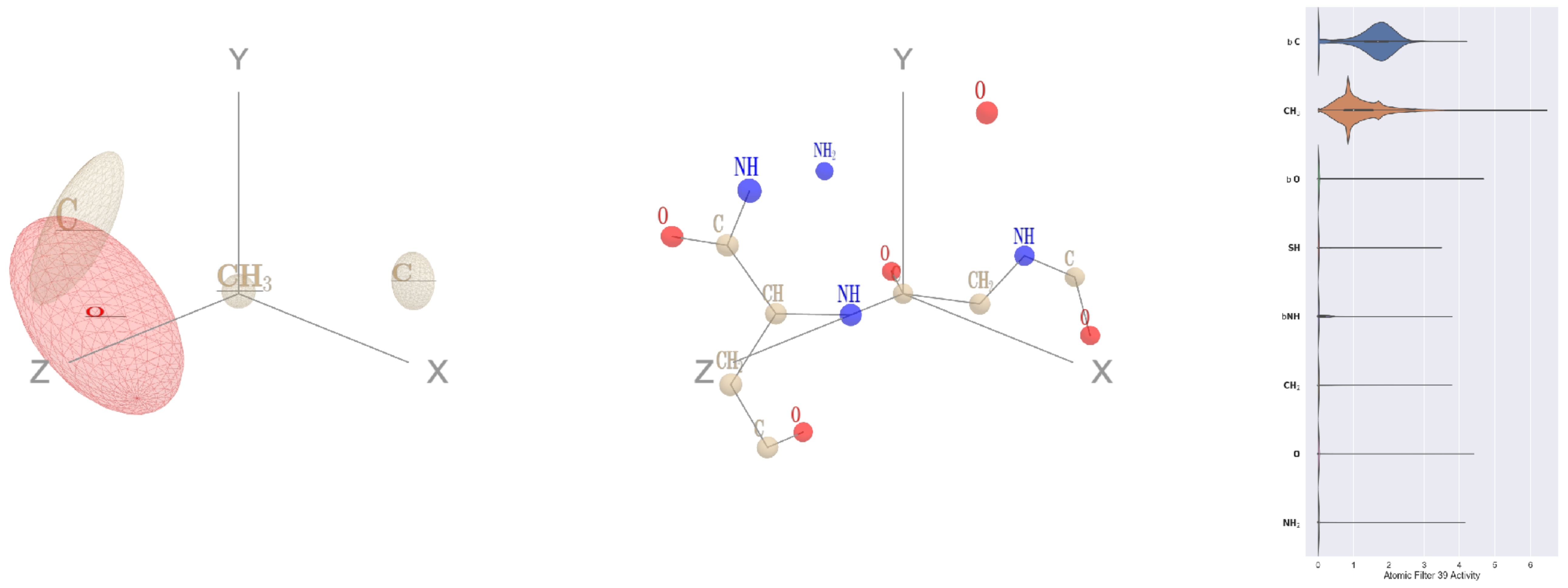


Filter 39
$R_{\text{pred}}^{\text{max}} = -0.06$
$R_{\text{pred}}^{\text{mean}} = -0.11$
Percent Active = 35
Pooling attention 0.28
Pooling feature 0.20
Top activators
2gfq:A:211 G C
2zm2:A:206 G C
1vqv:B:141 G C
2nr6:A:186 G C
2wgq:B:538 G C
4a7z:A:238 G C
1r7r:A:624 N C
2w27:A:230 G C
2dyo:A:123 G C
3tsy:A:723 E C

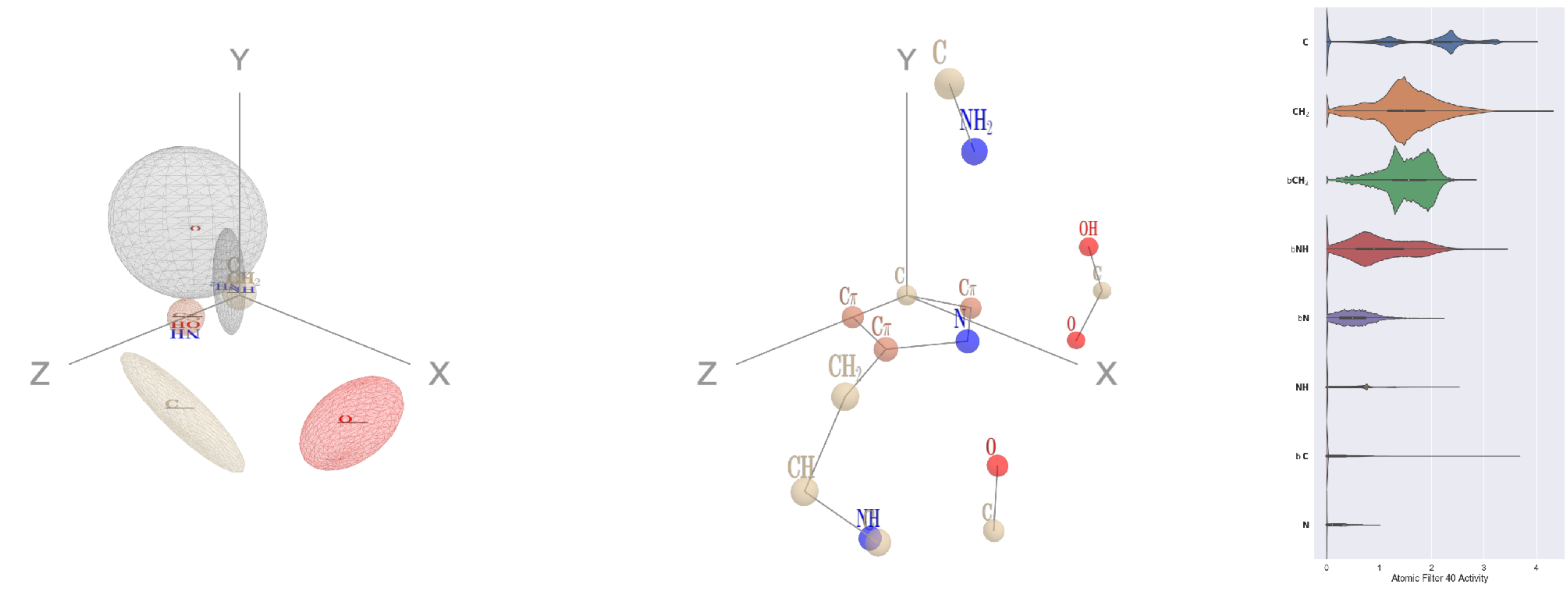

Filter 40
$R_{\text{pred}}^{\text{max}} = 0.07$
$R_{\text{pred}}^{\text{mean}} = 0.05$
Percent Active = 40
Pooling attention 0.25
Pooling feature 1.04
Top activators
4qiw:A:179 H NE2
4mex:C:1244 H NE2
4gfh:A:937 H NE2
2mas:A:288 H NE2
1l7k:B:43 H NE2
3rwr:D:158 H NE2
3rby:A:91 H NE2
4r3z:C:490 H NE2
3drw:A:258 H NE2
1e5d:A:115 H NE2

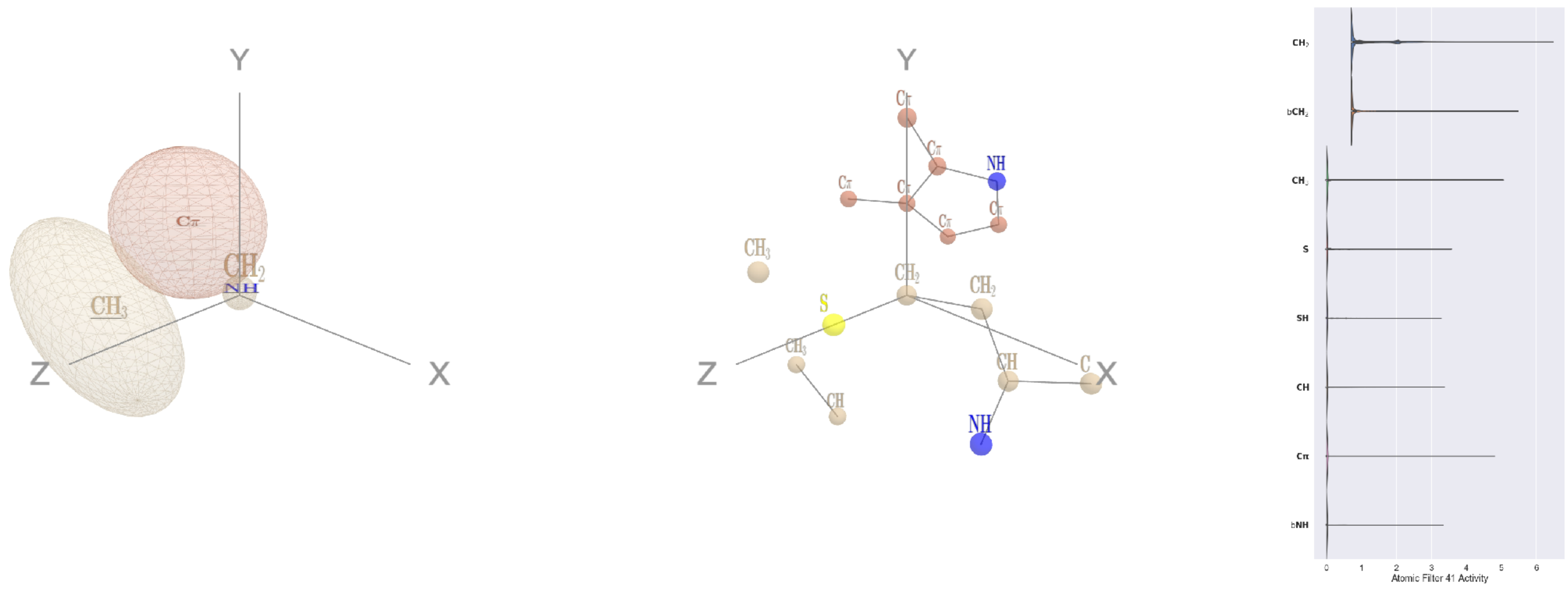

Filter 41
$R_{\text{pred}}^{\text{max}} = -0.07$
$R_{\text{pred}}^{\text{mean}} = -0.10$
Percent Active = 25
Pooling attention 0.06
Pooling feature 0.02
Top activators
1dtn:A:193 M CG
3bsd:A:170 I CG1
1sl0:A:336 P CB
1r6m:A:30 L CB
1mp9:A:30 L CB
2hh1:H:232 K CE
2pm9:A:374 P CB
2qn5:B:25 C CB
1l7k:B:124 L CB
2zm2:A:27 Q CG

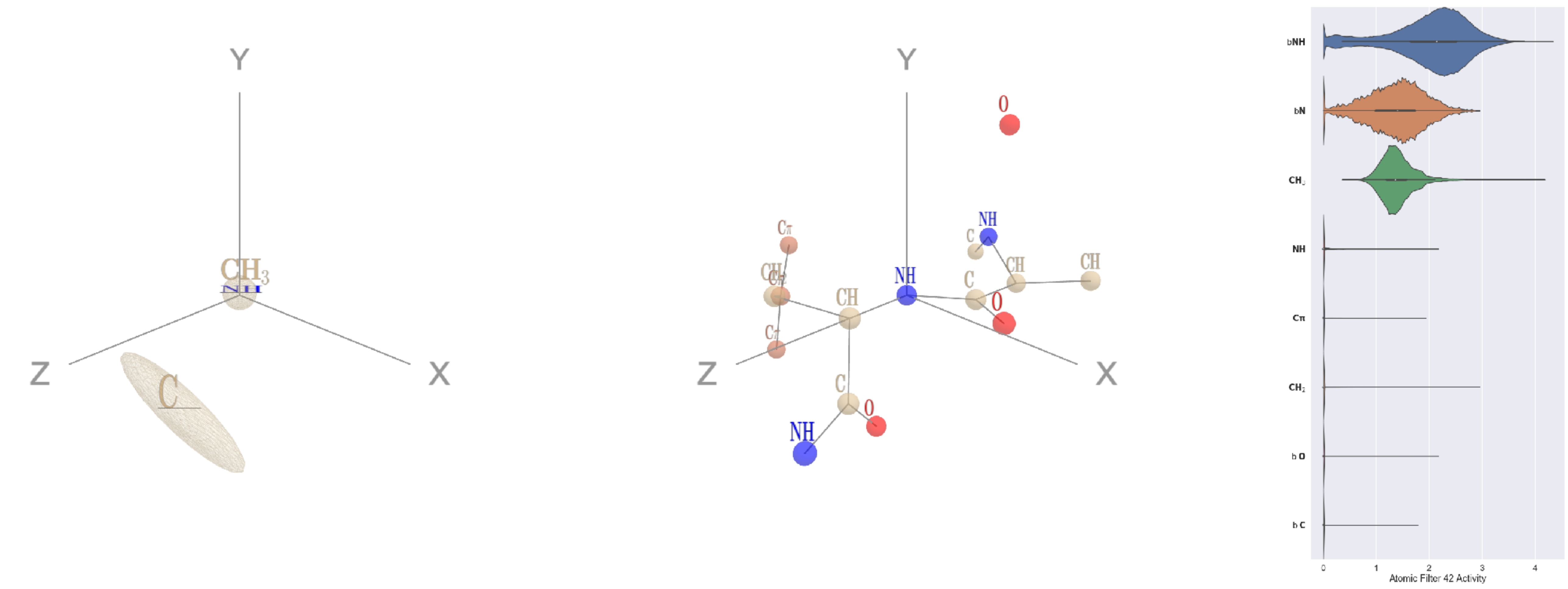

Filter 42
$R_{\text{pred}}^{\text{max}} = -0.01$
$R_{\text{pred}}^{\text{mean}} = -0.06$
Percent Active = 26
Pooling attention 0.04
Pooling feature 0.59
Top activators
1e1c:A:210 Y N
3din:E:43 F N
3bsd:A:330 Y N
2cge:A:300 W N
1iwa:A:76 A N
4bxf:A:250 H N
1ivu:A:201 H N
2pm9:A:376 W N
4r3z:C:511 W N
3o65:A:4 F N

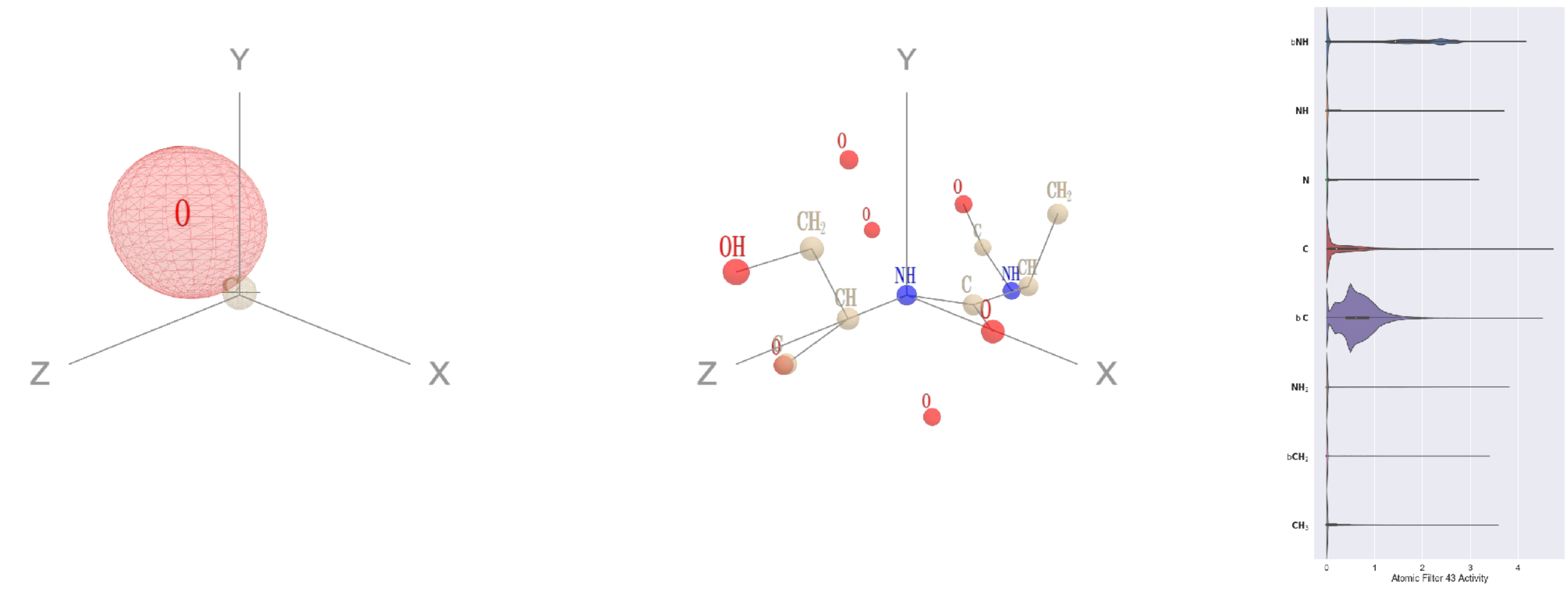

Filter 43
$R_{\text{pred}}^{\text{max}} = -0.07$
$R_{\text{pred}}^{\text{mean}} = -0.14$
Percent Active = 40
Pooling attention 0.14
Pooling feature 0.35
Top activators
1w36:B:449 S N
3b8e:A:373 T N
3tu3:B:683 N N
2wsc:B:190 W N
4c8q:H:623 F N
4r3z:C:473 A N
4gxg:A:155 A N
3n6r:G:106 A N
3n6r:G:600 D N
1e79:I:7 A N

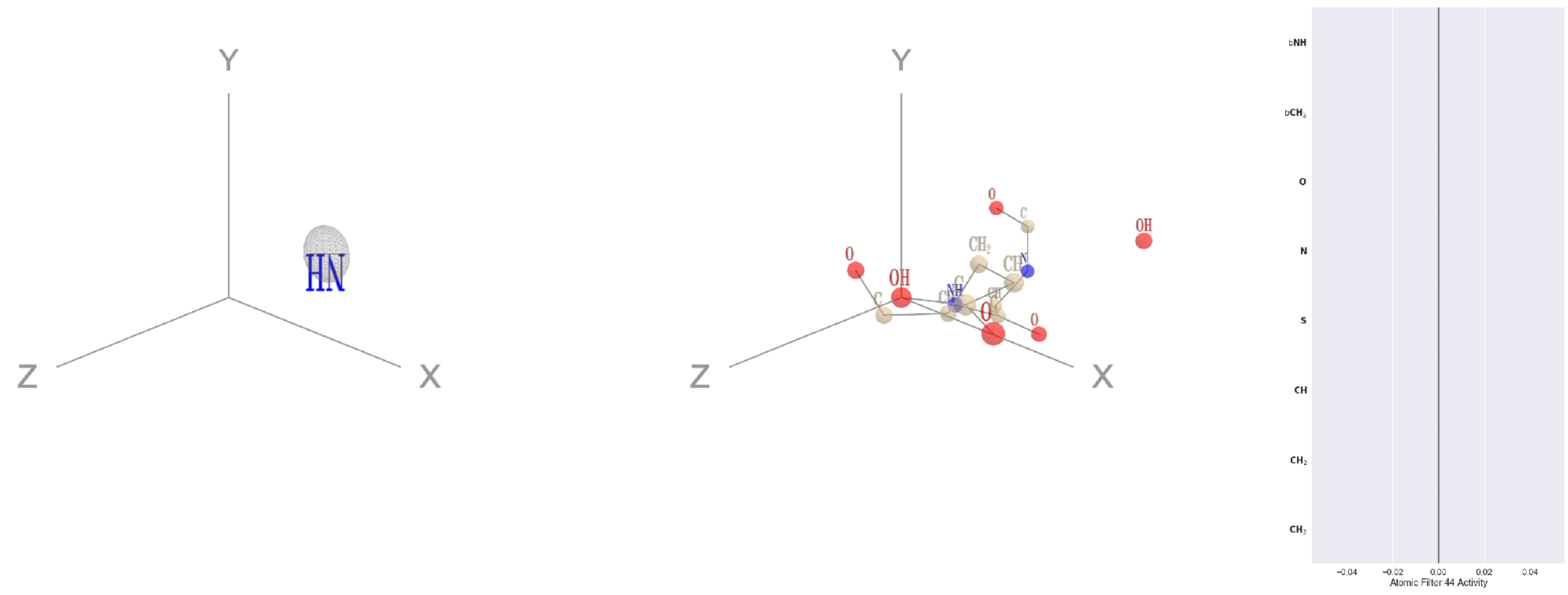

Filter 44
$R_{\text{pred}}^{\text{max}} = nan$
$R_{\text{pred}}^{\text{mean}} = nan$
Percent Active = 0
Pooling attention 0.00
Pooling feature 0.00
Top activators
5wy5:A:246 E OE2
2b0o:F:573 F N
2b0o:F:574 G O
2b0o:F:574 G C
2b0o:F:574 G CA
2b0o:F:574 G N
2b0o:F:573 F CZ
2b0o:F:573 F CE2
2b0o:F:573 F CE1
2b0o:F:573 F CD2

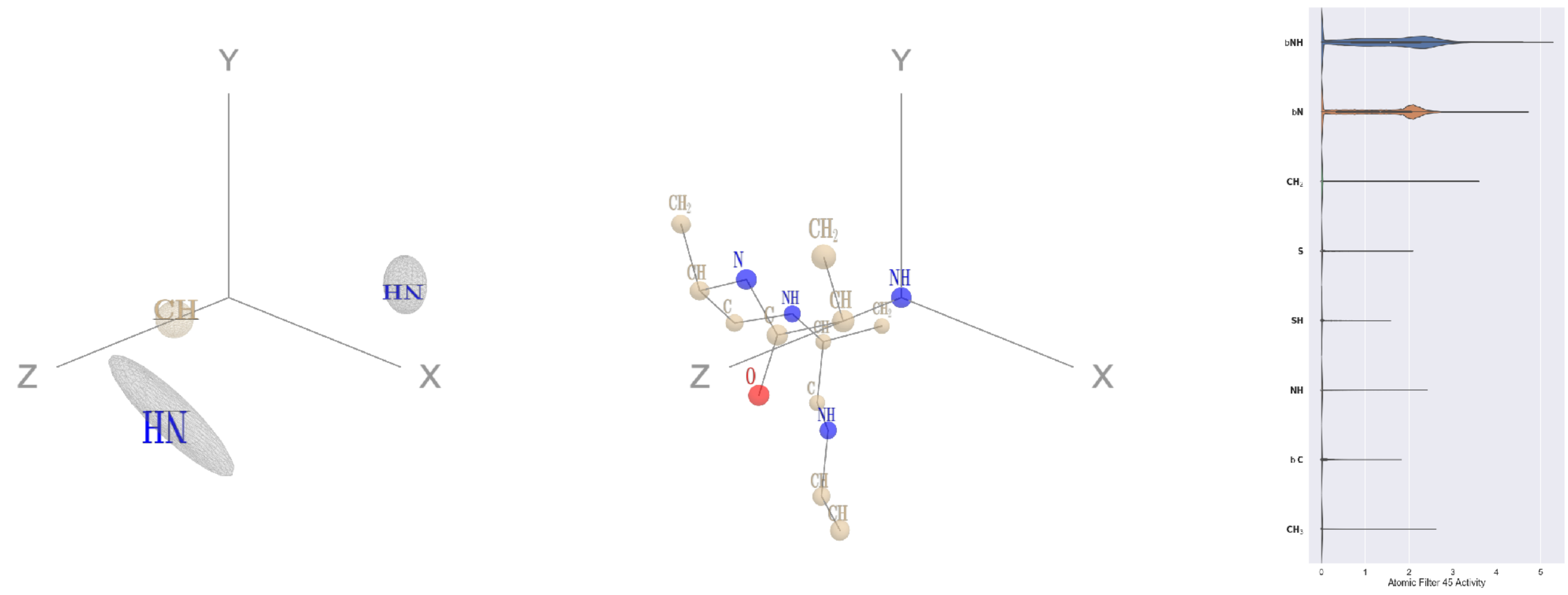

Filter 45
$R_{\text{pred}}^{\text{max}} = -0.01$
$R_{\text{pred}}^{\text{mean}} = -0.08$
Percent Active = 27
Pooling attention 0.14
Pooling feature 0.40
Top activators
4c8q:H:656 S N
2b5d:X:475 Q N
4gfh:A:7 S N
3vie:B:116 S N
3zdl:B:7 D N
4f52:E:1 M N
3kyh:C:284 Y N
4qiw:A:6 K N
1mu2:A:72 R N
3d00:A:127 S N

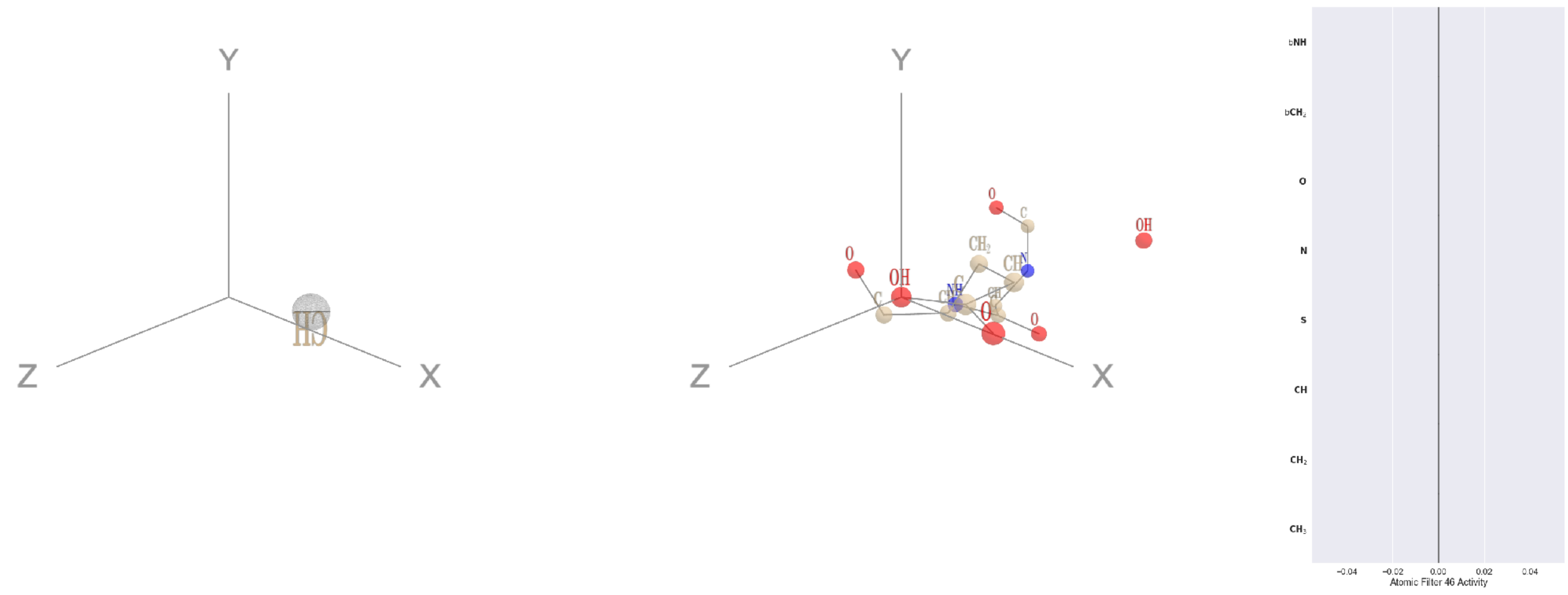

Filter 46
$R_{\text{pred}}^{\text{max}} = nan$
$R_{\text{pred}}^{\text{mean}} = nan$
Percent Active = 0
Pooling attention 0.00
Pooling feature 0.00
Top activators
5wy5:A:246 E OE2
2b0o:F:573 F N
2b0o:F:574 G O
2b0o:F:574 G C
2b0o:F:574 G CA
2b0o:F:574 G N
2b0o:F:573 F CZ
2b0o:F:573 F CE2
2b0o:F:573 F CE1
2b0o:F:573 F CD2

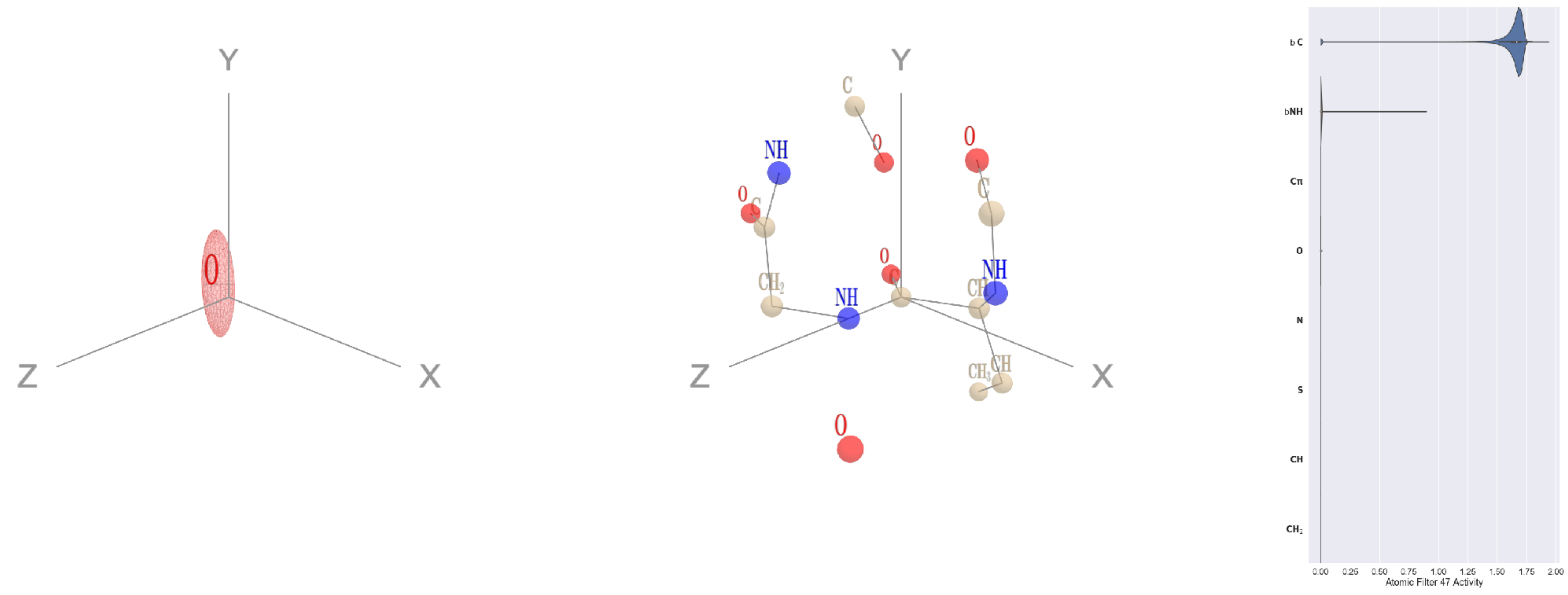

Filter 47
$R_{\text{pred}}^{\text{max}} = -0.02$
$R_{\text{pred}}^{\text{mean}} = -0.08$
Percent Active = 13
Pooling attention 0.02
Pooling feature 0.01
Top activators
1e5d:A:213 V C
4buj:B:520 E C
4oph:A:183 G C
2fml:A:188 E C
1u5e:A:164 F C
1u5e:A:159 Q C
3kyh:C:389 N C
1u5e:A:189 I C
3drw:A:218 A C
1ppt:A:24 L C

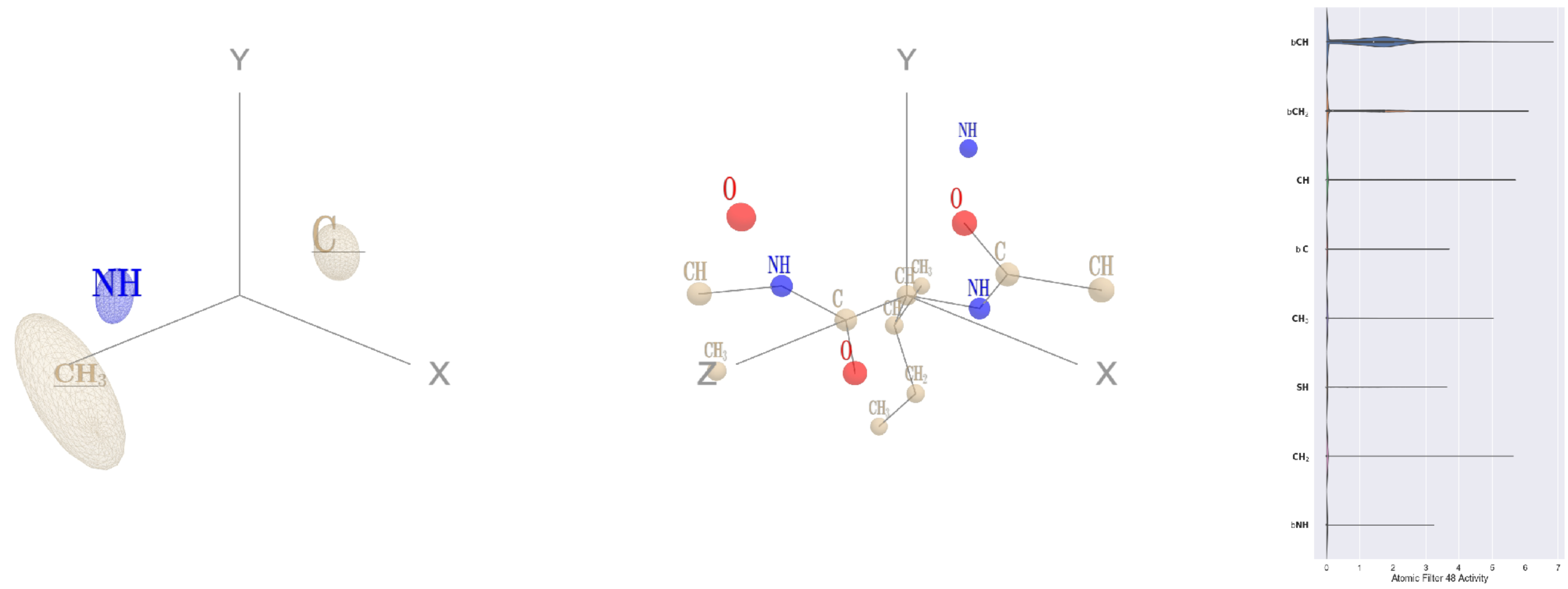


Filter 48
$R_{pred}^{max} = -0.01$
$R_{pred}^{mean} = -0.05$
Percent Active = 19
Pooling attention 0.12
Pooling feature 0.34
Top activators
2wsc:B:254 I CA
3bof:A:501 N CA
3rzz:A:142 D CA
1am4:A:45 F CA
1lnl:A:108 D CA
3ffk:A:254 P CA
1f4a:A:789 L CA
2hb6:A:231 P CA
1k90:C:296 L CA
2x98:A:435 H CA

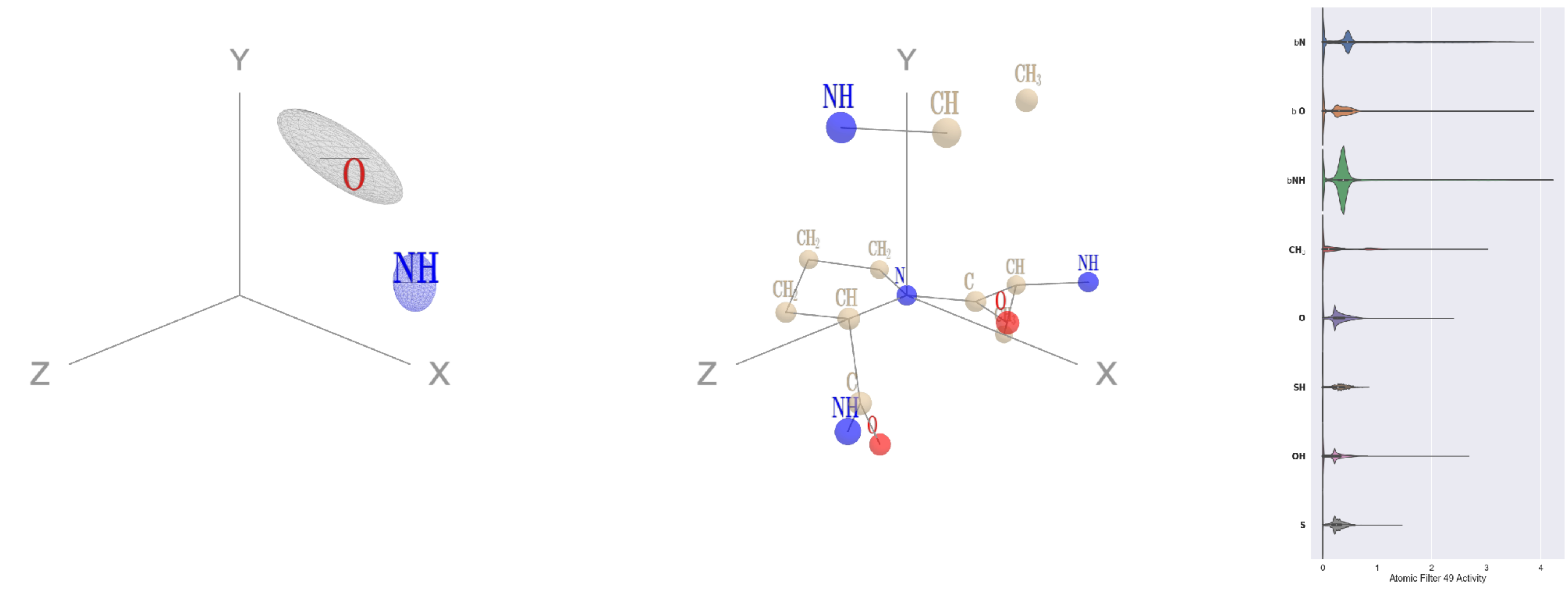


Filter 49
$R_{pred}^{max} = 0.01$
$R_{pred}^{mean} = -0.01$
Percent Active = 58
Pooling attention 0.27
Pooling feature 0.07
Top activators
4ny2:A:235 P N
1ivu:A:228 P N
1iri:A:484 P N
2wk4:B:428 P N
1qha:A:326 P N
3c24:A:157 P N
2hh1:H:78 P N
4ny2:A:112 P N
4nqj:A:305 P N
1ei6:A:167 P N

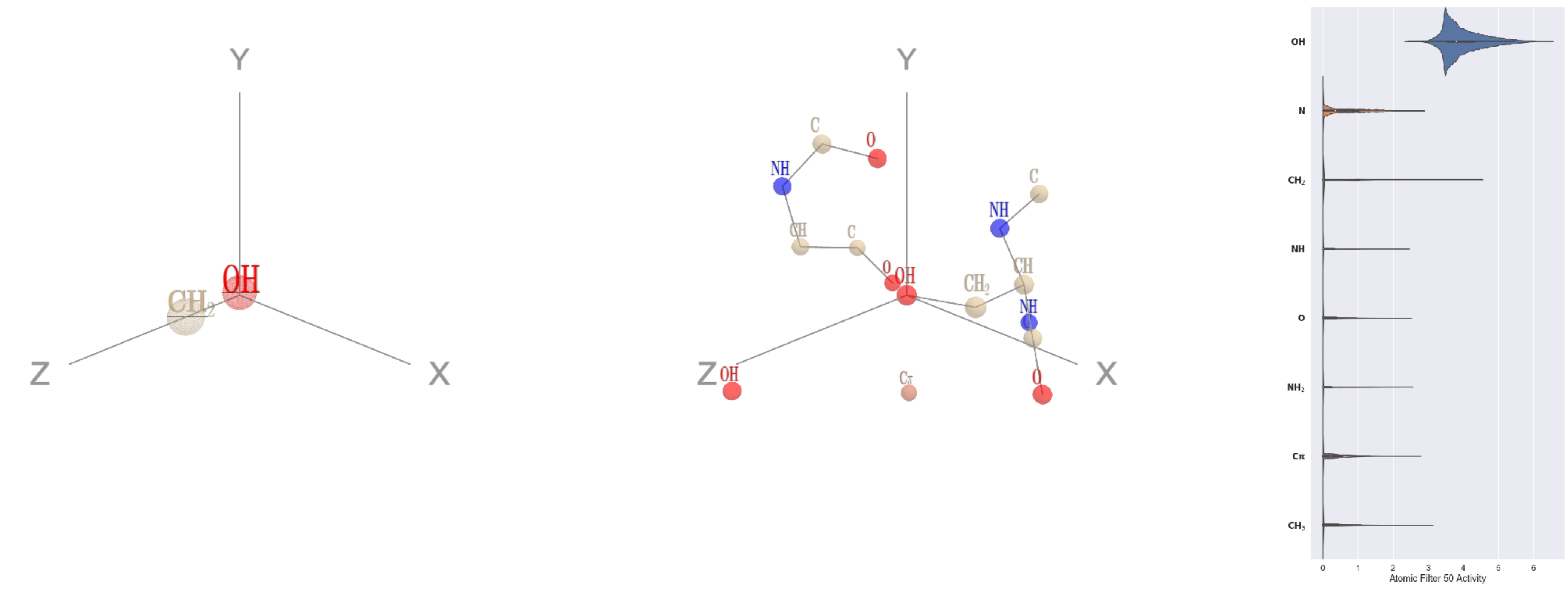


Filter 50
$R_{pred}^{max} = -0.01$
$R_{pred}^{mean} = -0.04$
Percent Active = 42
Pooling attention 0.15
Pooling feature 0.66
Top activators
2wk4:B:327 S OG
1c0i:A:1268 S OG
4hg6:A:385 T OG1
2qn5:B:104 D OD2
3hot:A:236 E OE2
1s4k:A:86 S OG
1efu:B:43 S OG
2pm9:A:53 E OE2
1e1c:A:209 T OG1
3b8e:A:636 D OD2

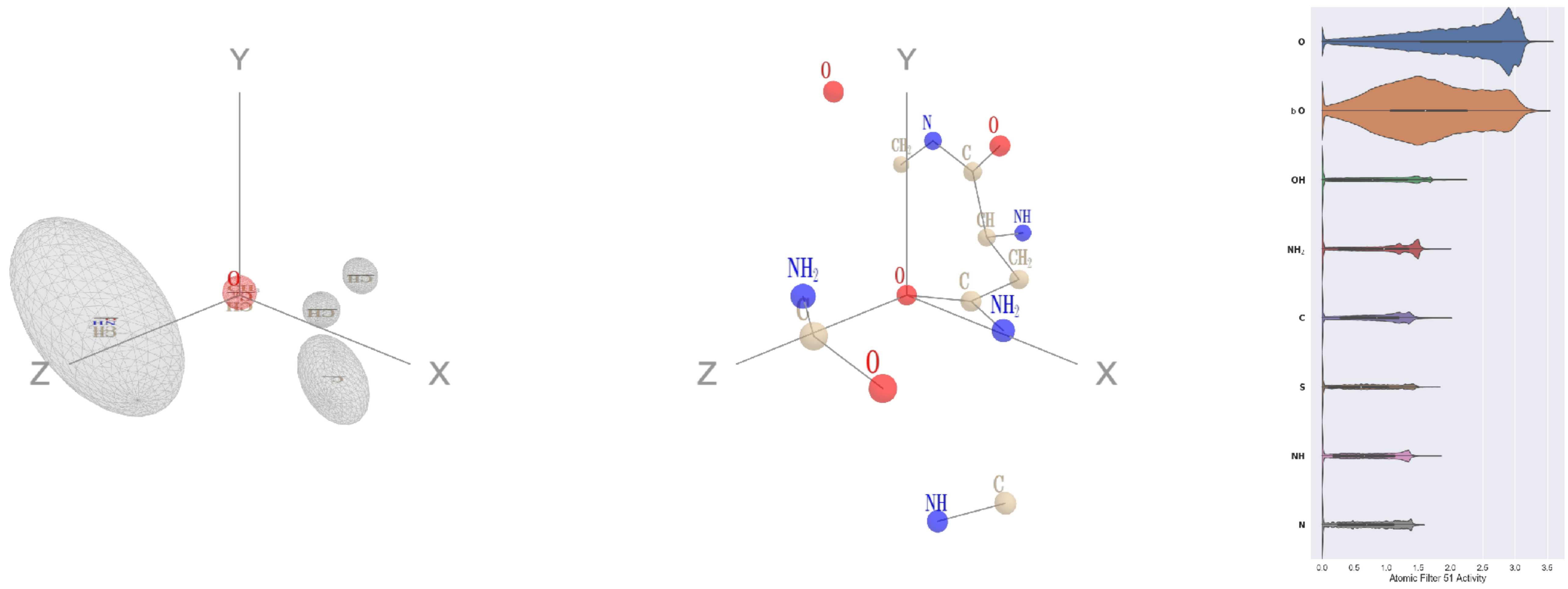


Filter 51
$R_{pred}^{max} = 0.18$
$R_{pred}^{mean} = 0.21$
Percent Active = 47
Pooling attention 0.73
Pooling feature 0.98
Top activators
3bsd:A:136 N OD1
3n6r:G:203 D OD1
1lk5:A:188 D OD1
1lnl:A:354 D OD1
1izn:B:11 D OD1
4c2m:A:1199 Q OE1
4c8q:H:493 E OE1
1q18:A:229 D OD1
1pgu:B:446 N OD1
1bo1:A:228 D OD1

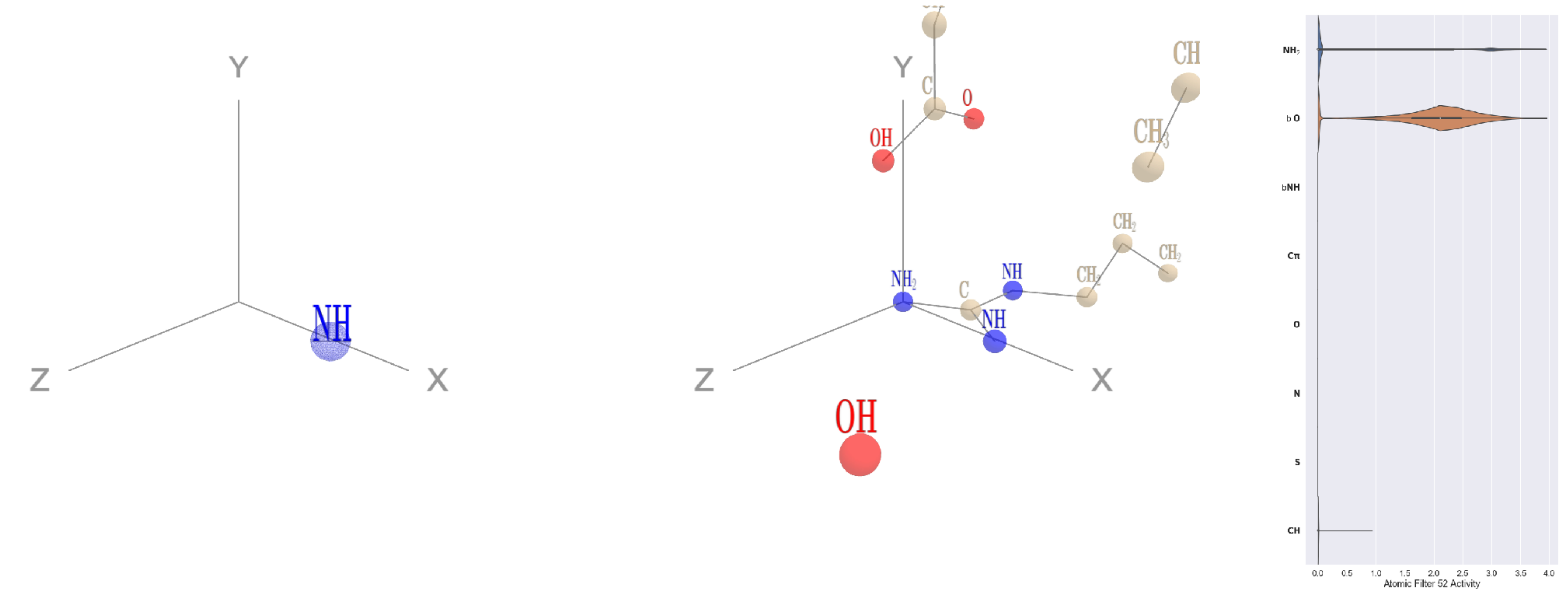


Filter 52
$R_{\text{pred}}^{\text{max}} = 0.01$
$R_{\text{pred}}^{\text{mean}} = -0.03$
Percent Active = 12
Pooling attention 0.10
Pooling feature 0.05
Top activators
1bxr:A:294 R NH2
1bxr:A:426 R NH2
1bxr:A:509 R NH2
1dtn:A:10 R NH2
1f4a:A:425 R NH2
1bxr:A:131 R NH2
1c0i:A:1261 R NH2
3do6:A:82 R NH2
3drw:A:369 R NH2
1a0j:A:222 R NH2

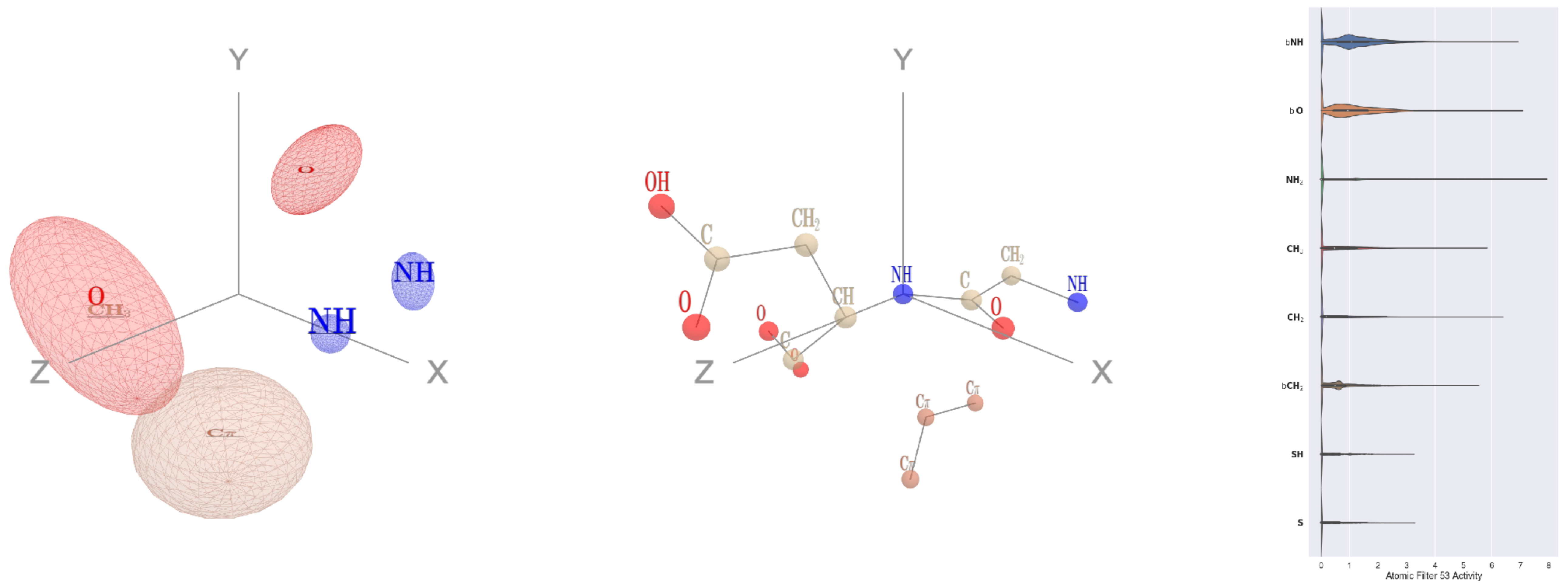


Filter 53
$R_{\text{pred}}^{\text{max}} = -0.06$
$R_{\text{pred}}^{\text{mean}} = -0.17$
Percent Active = 57
Pooling attention 0.45
Pooling feature 0.26
Top activators
1fpq:A:260 D N
1qha:A:79 D N
4jdm:C:151 K N
2pm9:A:329 T N
4f5w:A:193 A N
4v4k:M:56 Q N
4asi:C:1978 R N
3i5d:B:74 I N
3kov:A:58 A N
4a7z:A:547 R N

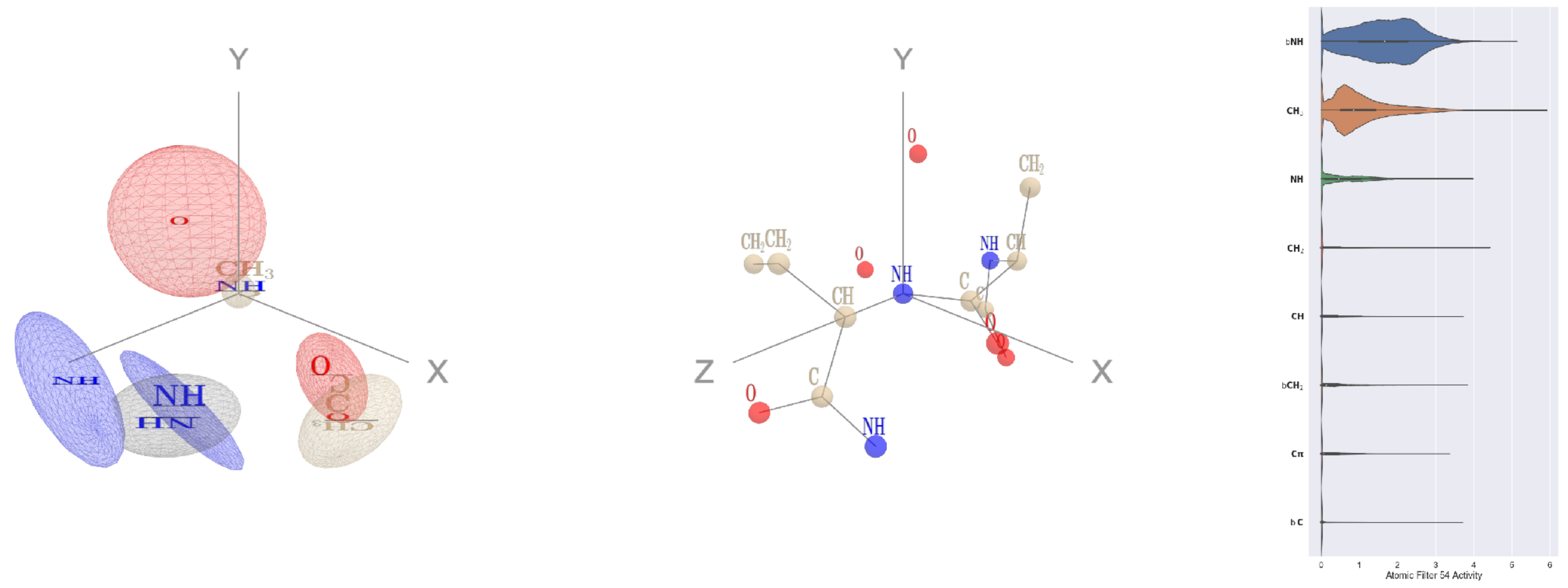


Filter 54
$R_{\text{pred}}^{\text{max}} = -0.05$
$R_{\text{pred}}^{\text{mean}} = -0.12$
Percent Active = 45
Pooling attention 0.45
Pooling feature 1.21
Top activators
4v88:AF:98 M N
1umg:A:312 D N
3o5t:A:243 D N
2x98:A:185 N N
3u33:A:25 G N
1c7d:A:117 F N
3ax7:A:1015 N N
1srq:A:185 D N
1hfe:L:246 L N
1lqb:C:190 D N

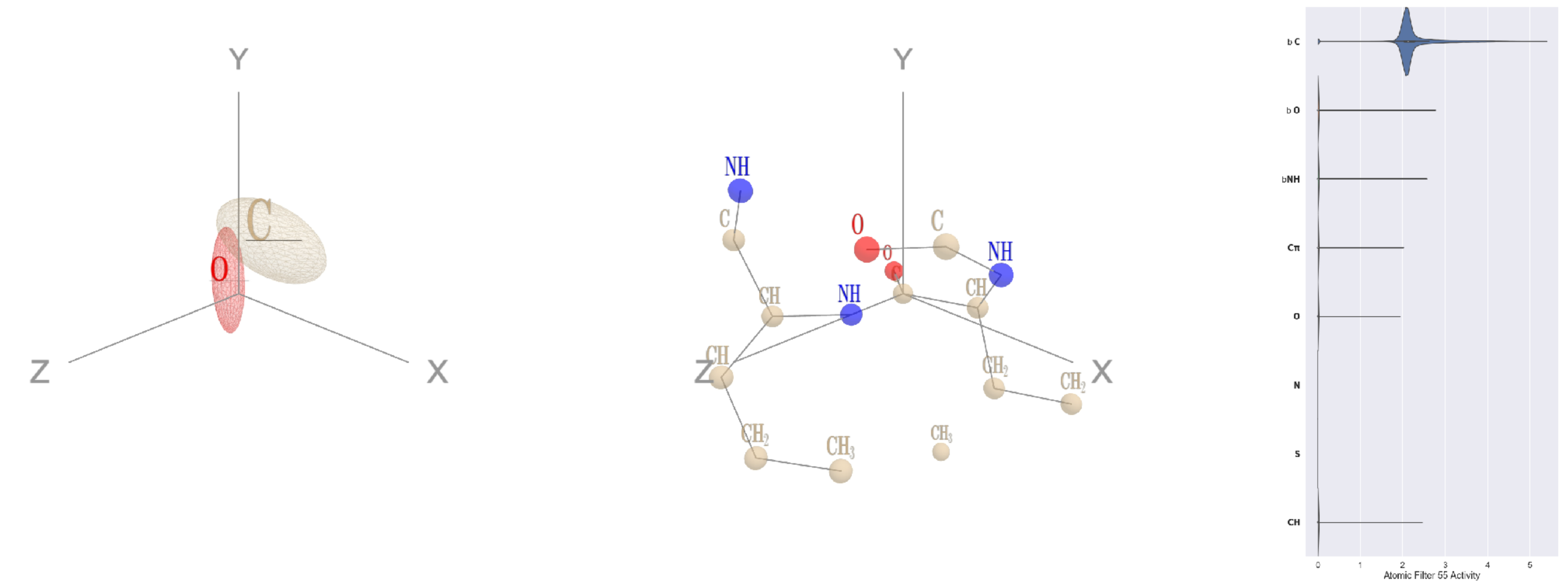


Filter 55
$R_{\text{pred}}^{\text{max}} = 0.01$
$R_{\text{pred}}^{\text{mean}} = -0.06$
Percent Active = 15
Pooling attention 0.18
Pooling feature 0.04
Top activators
4s20:L:189 M C
3b8e:A:294 F C
2wgq:B:218 K C
3b8e:A:1007 G C
4s20:L:375 E C
2wsc:B:367 T C
3b8e:A:626 I C
1qc9:A:63 K C
3dlb:A:426 L C
2wsc:B:217 P C

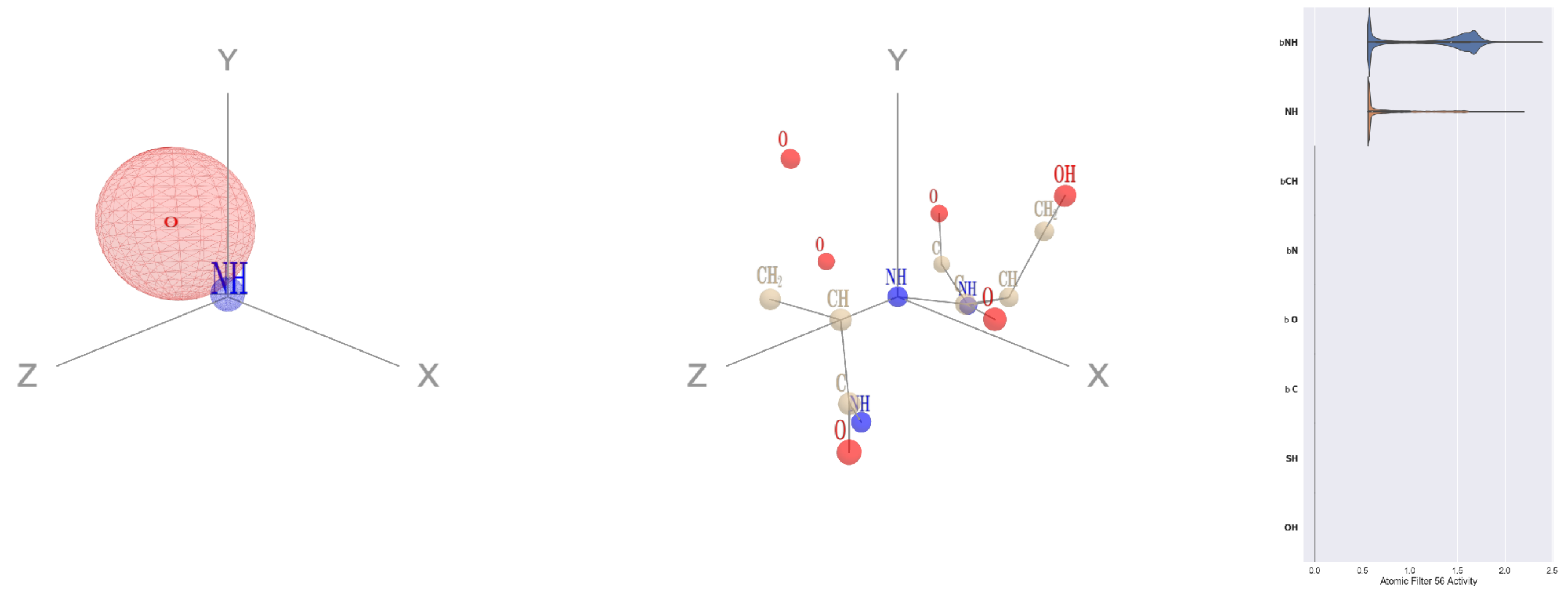


Filter 56
$R_{\text{pred}}^{\text{max}} = -0.07$
$R_{\text{pred}}^{\text{mean}} = -0.09$
Percent Active = 14
Pooling attention 0.03
Pooling feature 0.01
Top activators
4c8q:H:623 F N
2iut:A:505 S N
3bsd:A:172 S N
2r3s:A:127 G N
1e79:I:7 A N
1w36:B:449 S N
4wj3:B:205 Q N
2wsc:B:190 W N
4r3z:C:473 A N
2iut:A:352 V N

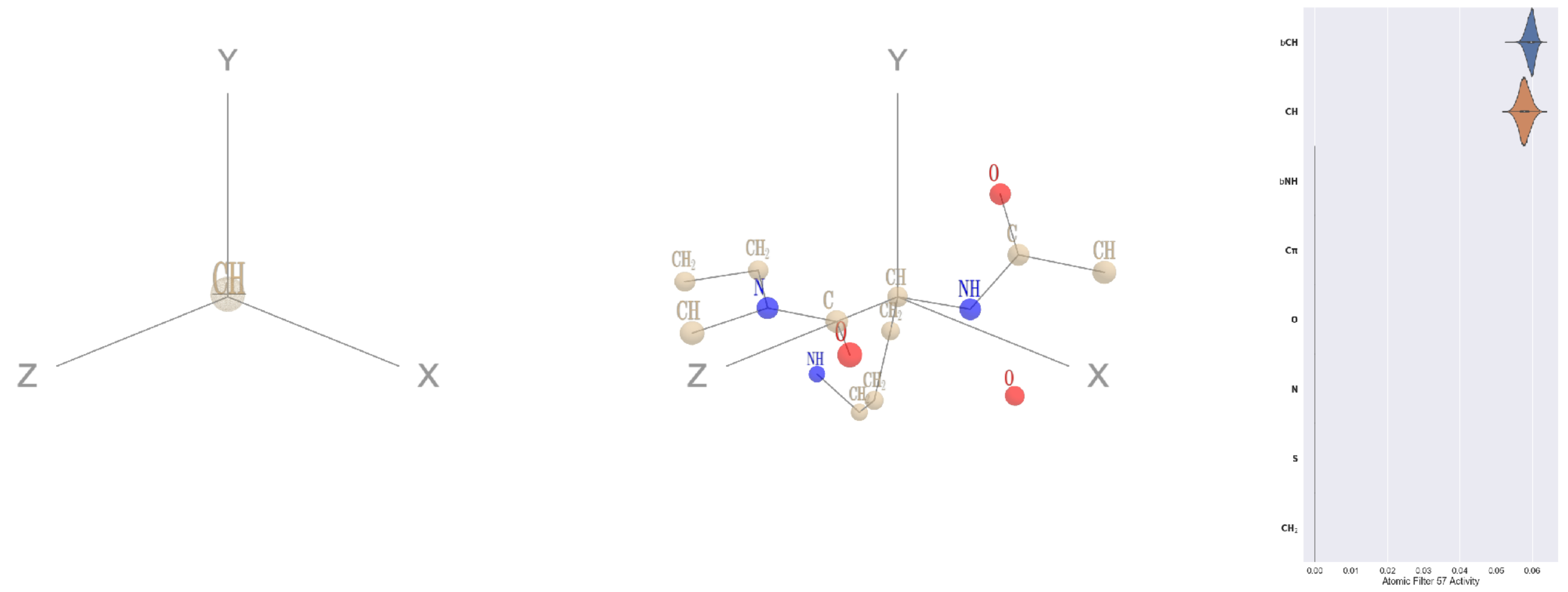


Filter 57
$R_{\text{pred}}^{\text{max}} = 0.03$
$R_{\text{pred}}^{\text{mean}} = -0.04$
Percent Active = 16
Pooling attention 0.00
Pooling feature 0.00
Top activators
1qc9:A:183 R CA
2akf:A:430 V CA
4cvw:A:792 L CA
2fq1:A:215 K CA
4k1n:A:310 E CA
2w27:A:185 S CA
2np9:A:324 I CA
2hb6:A:429 M CA
1lnl:A:264 E CA
2qn5:B:97 E CA

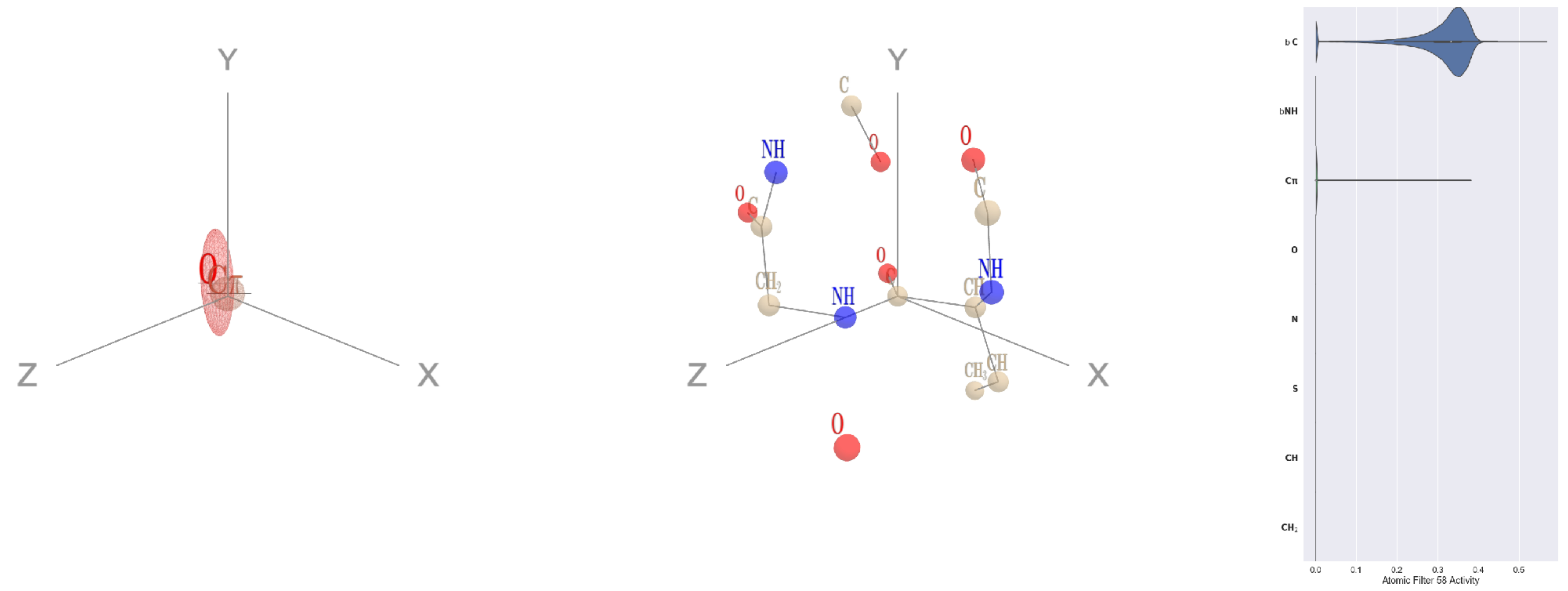


Filter 58
$R_{\text{pred}}^{\text{max}} = -0.03$
$R_{\text{pred}}^{\text{mean}} = -0.08$
Percent Active = 12
Pooling attention 0.01
Pooling feature 0.00
Top activators
1e5d:A:213 V C
4buj:B:520 E C
4oph:A:183 G C
2fml:A:188 E C
1u5e:A:159 Q C
1u5e:A:164 F C
3kyh:C:389 N C
1u5e:A:189 I C
1f4a:A:350 L C
3drw:A:218 A C

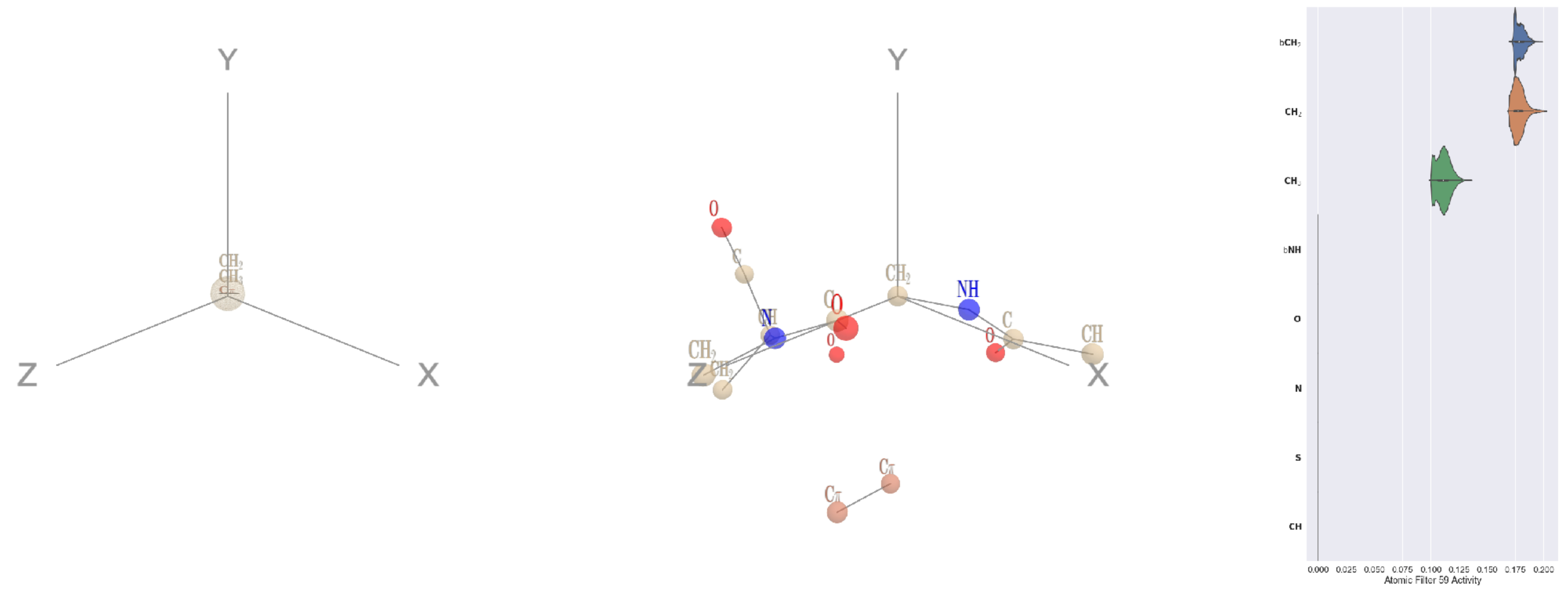


Filter 59
$R_{\text{pred}}^{\text{max}} = 0.03$
$R_{\text{pred}}^{\text{mean}} = -0.01$
Percent Active = 23
Pooling attention 0.01
Pooling feature 0.00
Top activators
1umg:A:306 G CA
4whj:A:494 G CA
4wj3:B:138 G CA
1ppt:A:1 G CA
4v81:A:16 G CA
1q18:A:140 G CA
2nyg:A:228 G CA
1k72:B:442 G CA
2hh1:H:8 G CA
4cy6:A:194 G CA

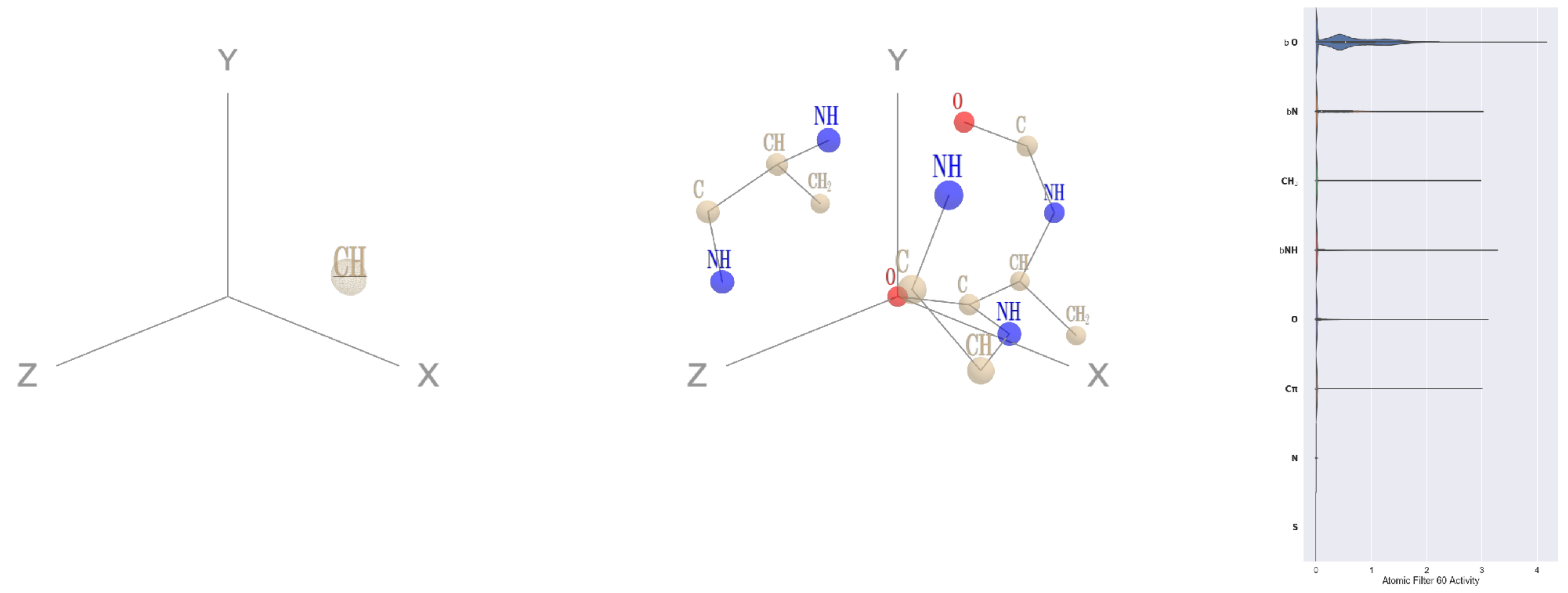


Filter 60
$R_{\text{pred}}^{\text{max}} = -0.01$
$R_{\text{pred}}^{\text{mean}} = -0.04$
Percent Active = 18
Pooling attention 0.07
Pooling feature 0.02
Top activators
3a5c:G:183 L O
3a5c:G:32 R O
3a5c:G:22 Q O
3a5c:G:185 Q O
3a5c:G:184 E O
3a5c:G:14 R O
3a5c:G:6 P O
3a5c:G:37 A O
3a5c:G:36 V O
3a5c:G:181 Q O

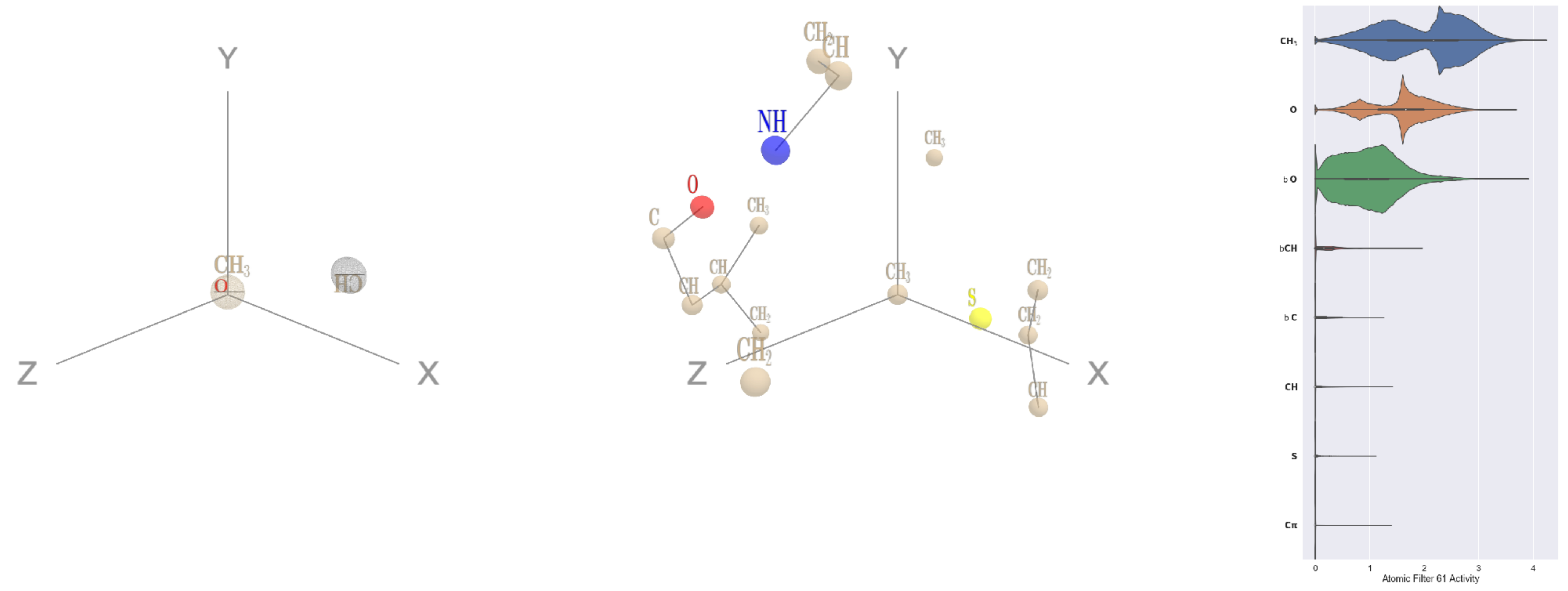


Filter 61
$R_{\text{pred}}^{\text{max}} = -0.08$
$R_{\text{pred}}^{\text{mean}} = -0.09$
Percent Active = 42
Pooling attention 0.12
Pooling feature 0.47
Top activators
1izn:B:13 M CE
3fbv:D:906 M CE
2zy4:F:471 M CE
3o47:A:152 M CE
3hh0:A:28 L CD2
1w36:B:15 L CD1
4nm9:B:152 L CD1
4buj:B:1143 L CD2
3kq4:A:183 M CE
4buj:B:859 L CD2

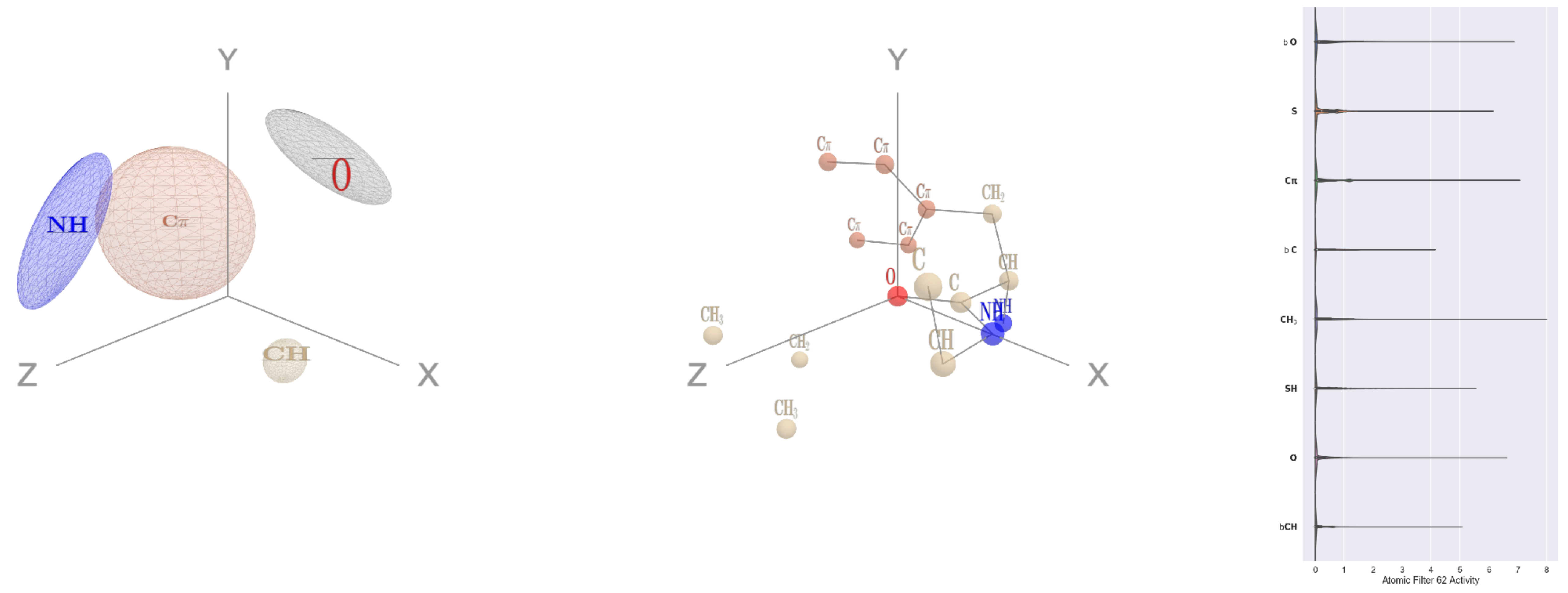


Filter 62
$R_{\text{pred}}^{\text{max}} = -0.06$
$R_{\text{pred}}^{\text{mean}} = -0.12$
Percent Active = 46
Pooling attention 0.05
Pooling feature 0.27
Top activators
3fbv:D:842 F O
1w36:B:265 W O
2np9:A:274 G O
4qiw:A:656 L O
4hg6:A:503 F O
4l8n:A:291 Y O
2a62:A:152 Q O
4hh2:D:50 L O
2f8x:C:276 L O
3ucq:A:520 R O

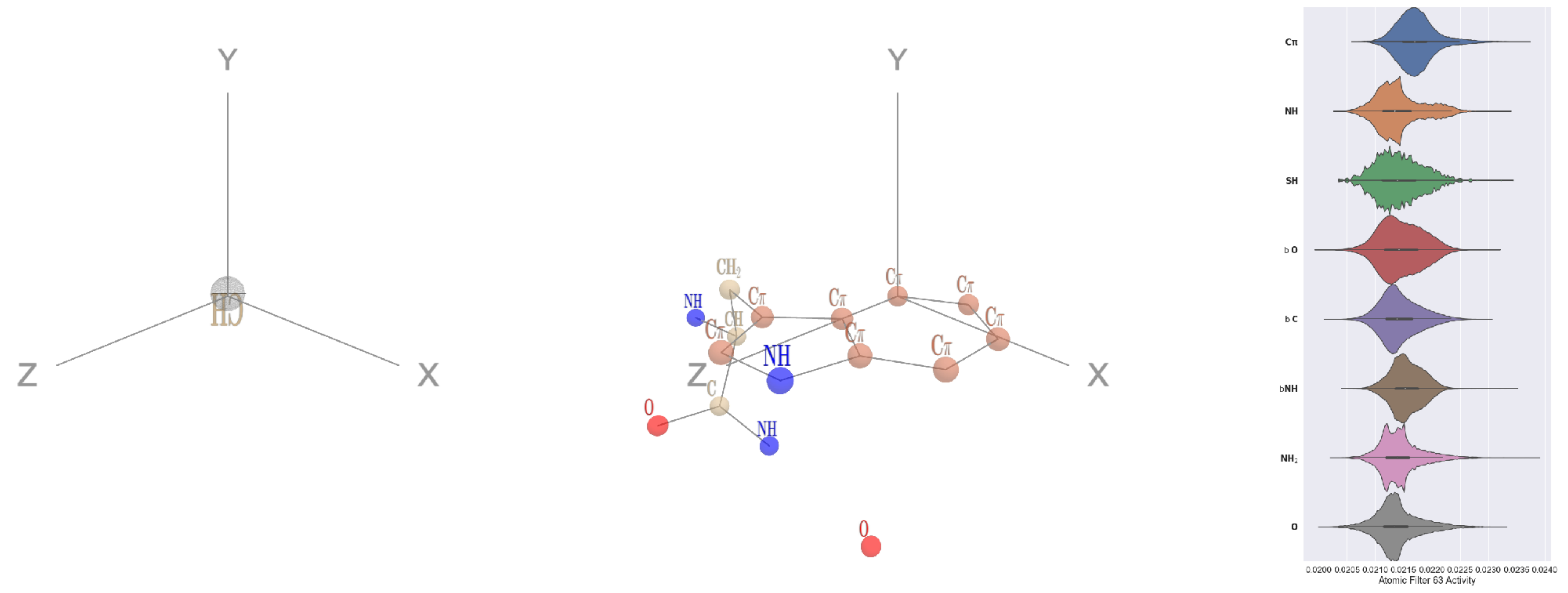


Filter 63
$R_{\text{pred}}^{\text{max}} = 0.00$
$R_{\text{pred}}^{\text{mean}} = 0.04$
Percent Active = 84
Pooling attention 0.00
Pooling feature 0.00
Top activators
3i0t:B:38 W CE3
5kqv:E:551 W CE3
2cge:A:300 W CE3
4v88:AF:46 W CE3
2eo4:A:148 W CE3
1mj2:A:102 W CE3
4nqj:A:311 W CE3
5ajd:B:466 W CE3
2gy7:B:163 H CD2
1r6m:A:59 W CE3

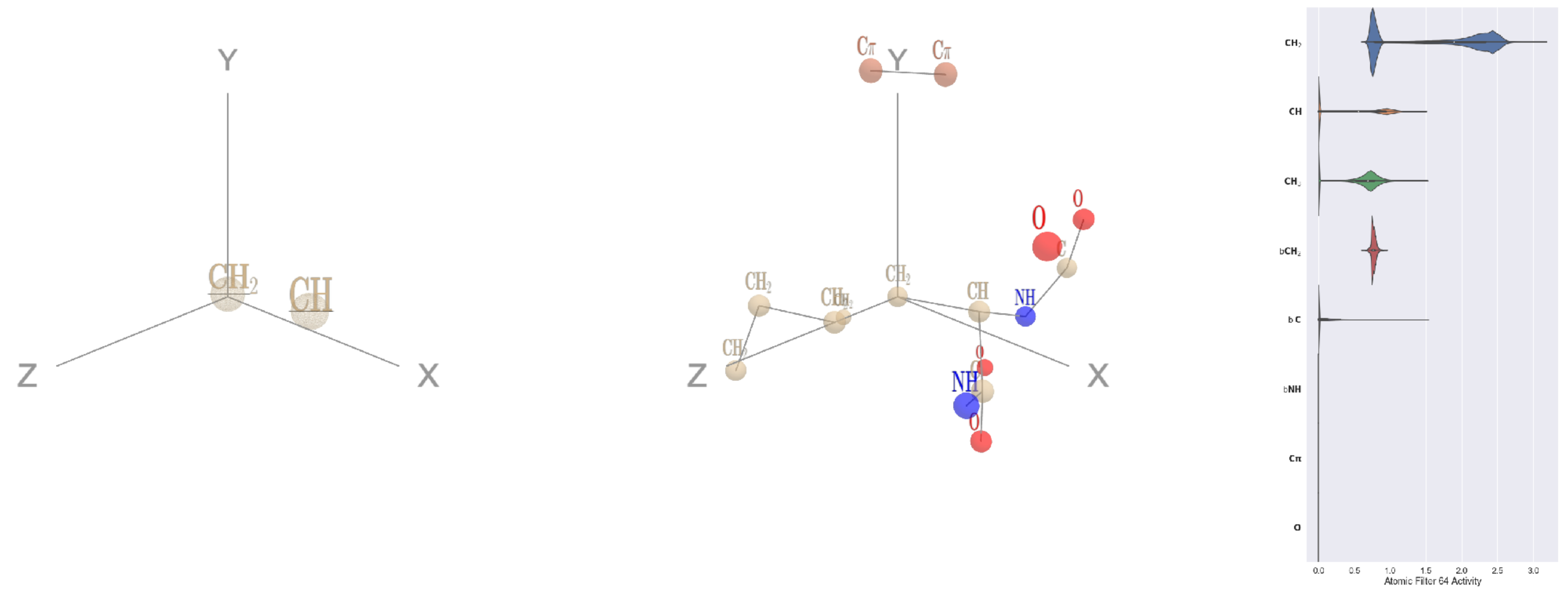

Filter 64
$R_{\mathrm{pred}}^{\max} = 0.05$
$R_{\mathrm{pred}}^{\mathrm{mean}} = -0.02$
Percent Active = 31
Pooling attention 0.26
Pooling feature 0.02
Top activators
1u5e:A:125 K CB
1ghq:C:33 S CB
1bh3:A:213 Y CB
1e1c:A:84 W CB
3a5c:G:183 L CB
1bh3:A:96 Y CB
3a5c:G:135 E CB
1qha:A:293 F CB
1ei6:A:242 H CB
1e1c:A:212 Y CB

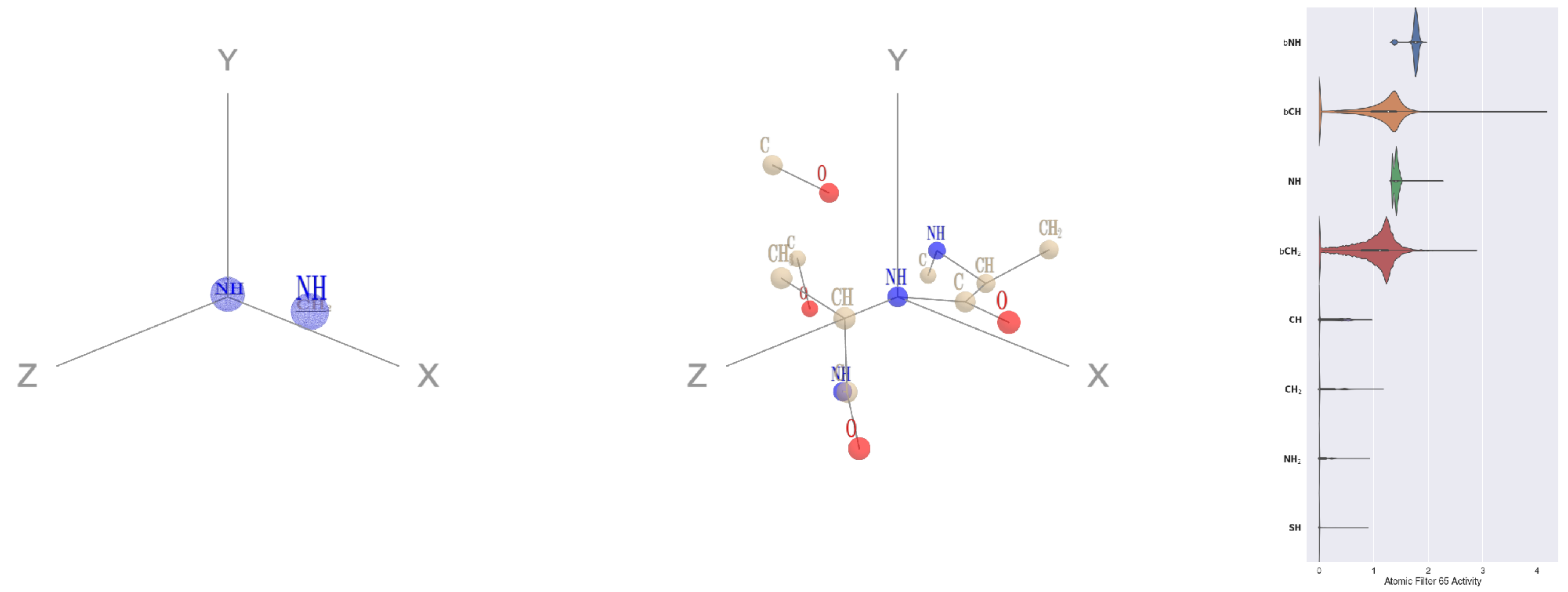

Filter 65
$R_{\mathrm{pred}}^{\max} = 0.00$
$R_{\mathrm{pred}}^{\mathrm{mean}} = -0.04$
Percent Active = 33
Pooling attention 0.03
Pooling feature 1.00
Top activators
3a5c:G:203 A N
3a5c:G:144 E N
1u5e:A:188 E N
2np9:A:303 V N
1vl4:A:31 F N
1u5e:A:186 C N
1bgv:A:260 S N
4kzx:R:91 L N
4kzx:L:18 Q N
1f4a:A:651 L N

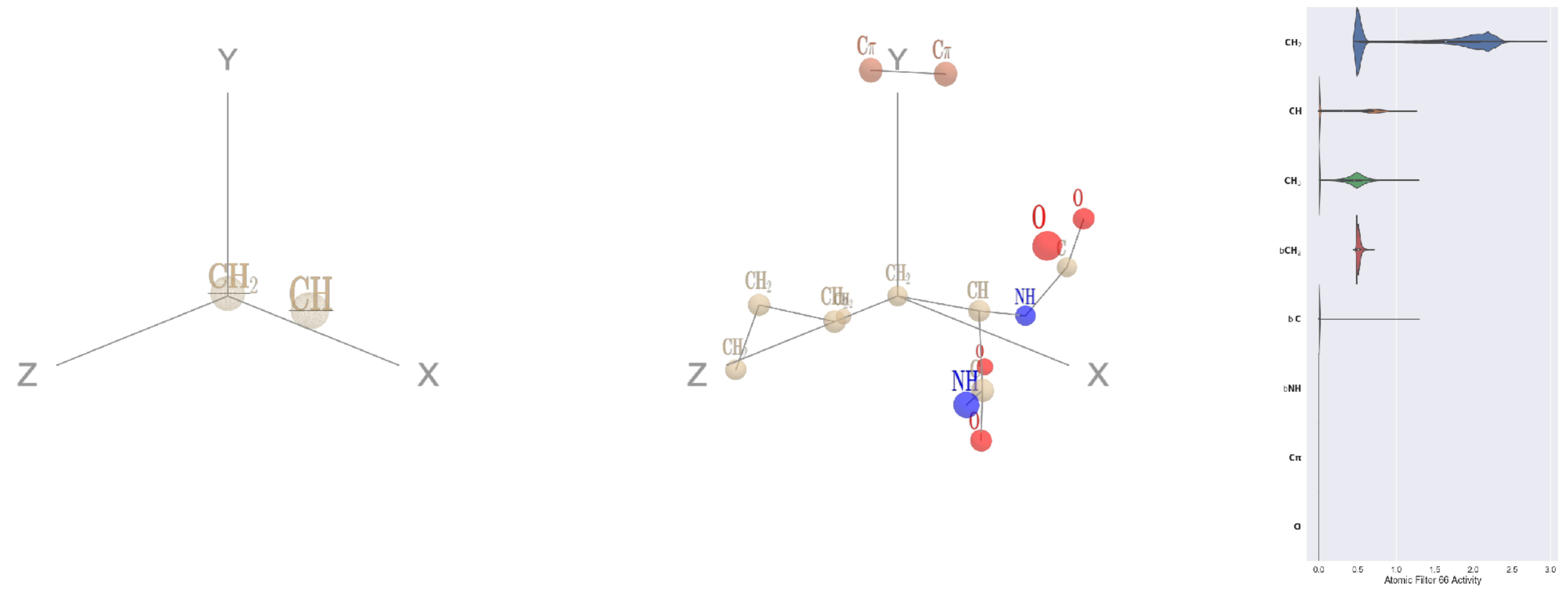

Filter 66
$R_{\mathrm{pred}}^{\max} = 0.04$
$R_{\mathrm{pred}}^{\mathrm{mean}} = -0.01$
Percent Active = 25
Pooling attention 0.17
Pooling feature 0.02
Top activators
1u5e:A:125 K CB
1ghq:C:33 S CB
1bh3:A:213 Y CB
1e1c:A:84 W CB
3a5c:G:183 L CB
1bh3:A:96 Y CB
3a5c:G:135 E CB
1qha:A:293 F CB
1e1c:A:212 Y CB
1bgv:A:113 K CB

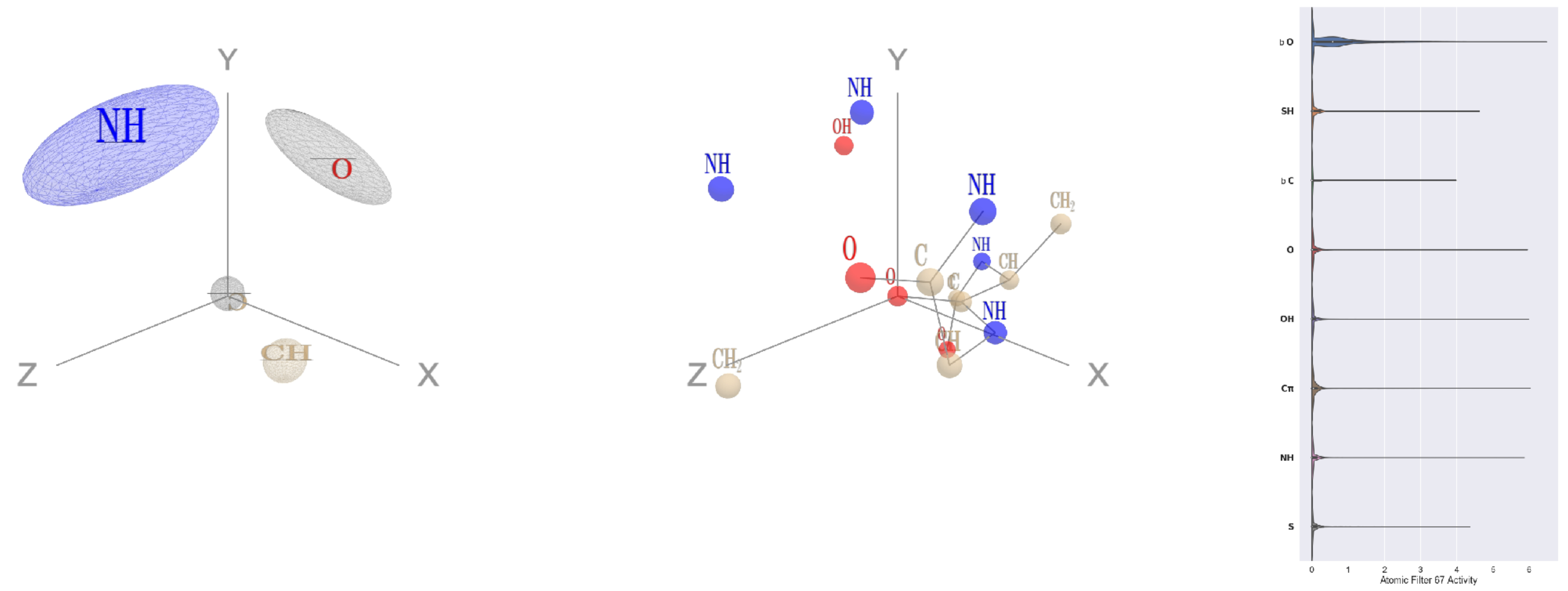

Filter 67
$R_{\mathrm{pred}}^{\max} = -0.05$
$R_{\mathrm{pred}}^{\mathrm{mean}} = -0.07$
Percent Active = 39
Pooling attention 0.12
Pooling feature 0.48
Top activators
1w36:B:721 N O
2pif:B:7 A O
3lnm:D:169 S O
1he8:A:621 A O
1cm5:A:412 D O
4f52:E:433 K O
1izl:D:34 G O
4rcn:B:359 G O
3s9m:C:217 K O
1izl:D:325 I O

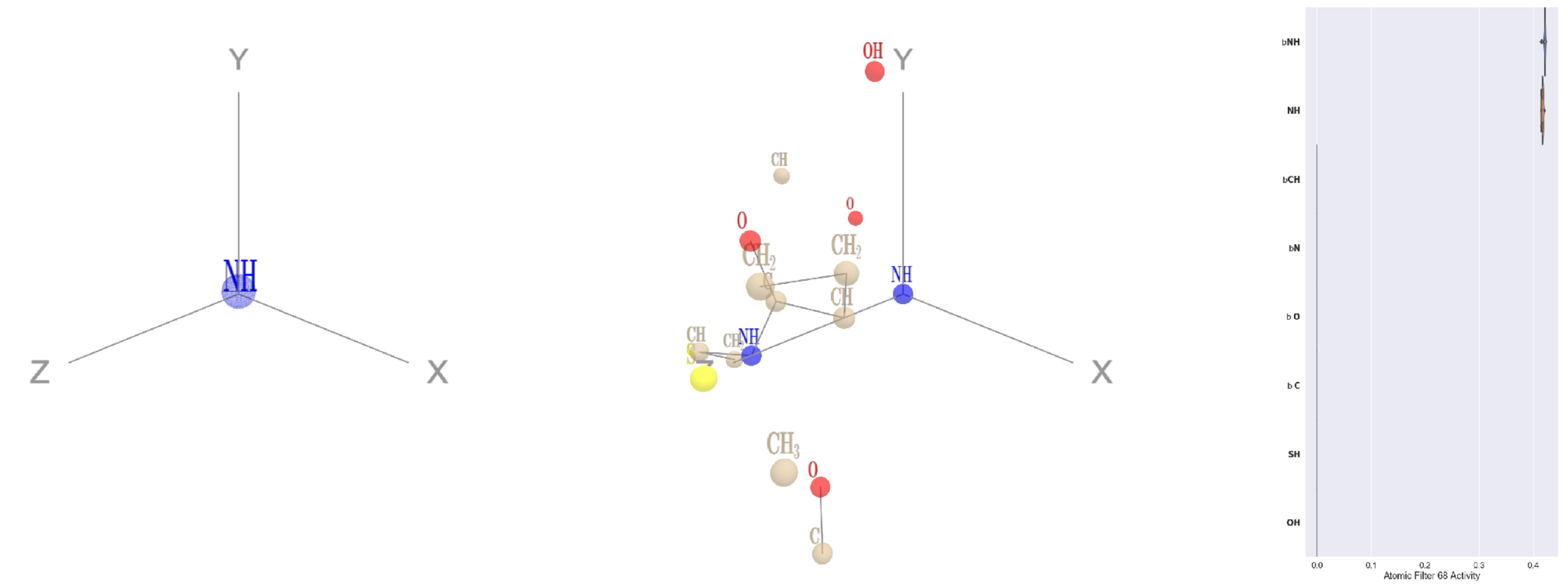


Filter 68
$R_{\text{pred}}^{\text{max}} = 0.00$
$R_{\text{pred}}^{\text{mean}} = -0.02$
Percent Active = 14
Pooling attention 0.01
Pooling feature 0.00
Top activators
1ii7:A:1 M N
4cht:B:2 N N
4pfp:A:5 K N
3kyh:C:284 Y N
1iwa:A:293 L N
3fbv:D:1095 V N
3ilk:A:100 V N
4pk7:A:181 V N
1ivu:A:26 I N
2wsw:A:24 I N

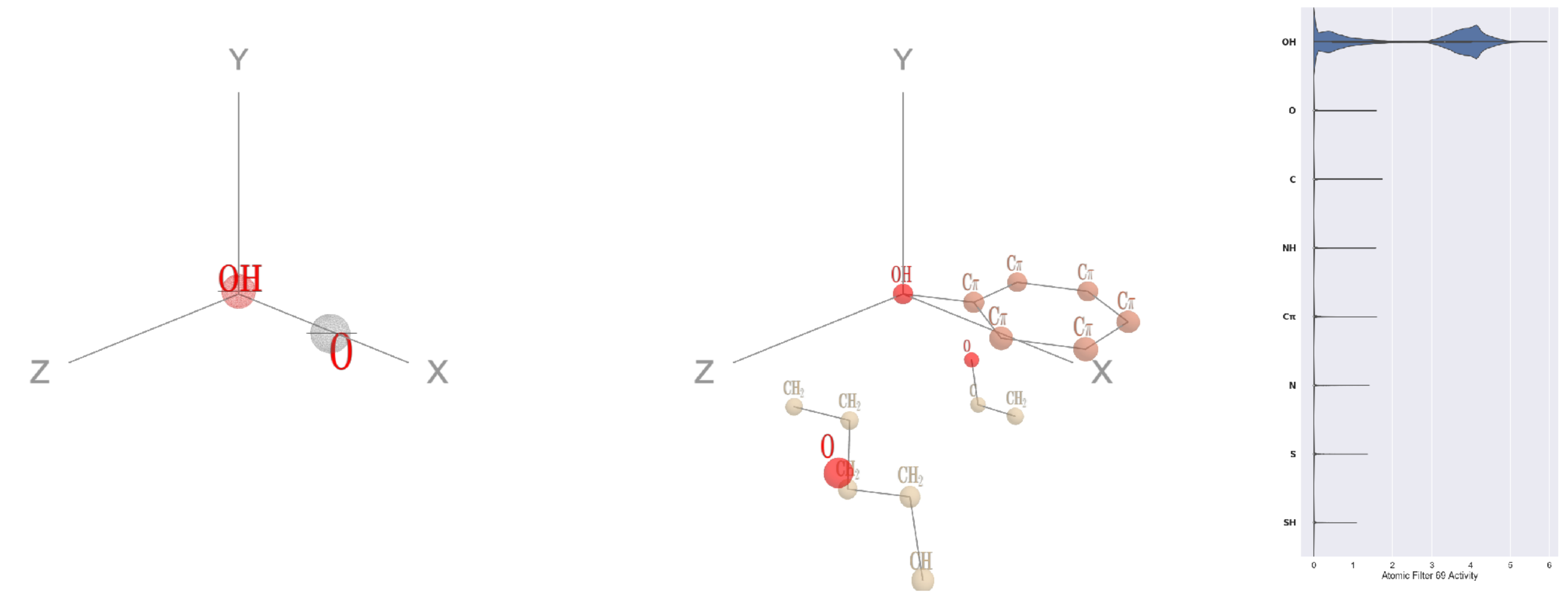


Filter 69
$R_{\text{pred}}^{\text{max}} = 0.03$
$R_{\text{pred}}^{\text{mean}} = 0.02$
Percent Active = 17
Pooling attention 0.13
Pooling feature 0.06
Top activators
2i71:B:363 Y OH
2x3l:B:230 Y OH
2q13:A:181 Y OH
3g8q:A:95 Y OH
4cy6:A:299 Y OH
1mu2:A:144 Y OH
4pba:A:281 S OG
4gfh:A:887 Y OH
3wrx:C:1084 Y OH
4pht:A:466 Y OH

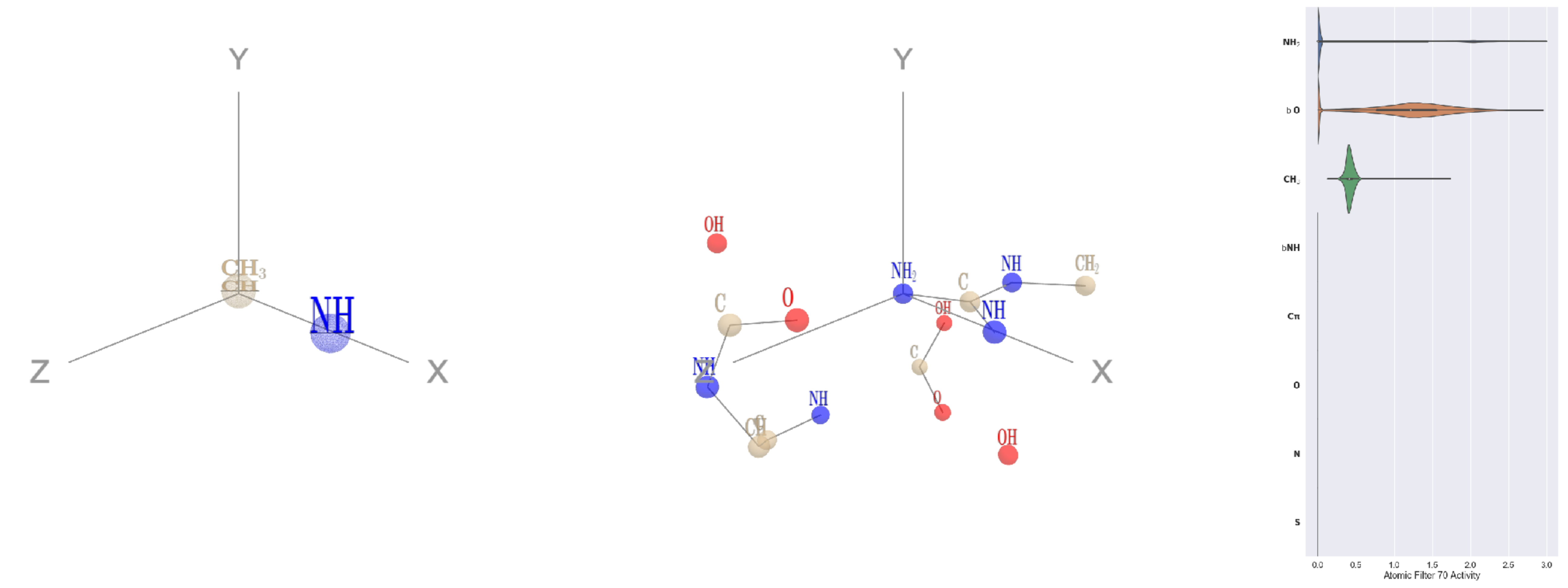


Filter 70
$R_{\text{pred}}^{\text{max}} = 0.01$
$R_{\text{pred}}^{\text{mean}} = -0.04$
Percent Active = 20
Pooling attention 0.02
Pooling feature 0.15
Top activators
1b8p:A:199 R NH2
1bxr:A:1027 R NH2
1bxr:A:343 R NH2
3g8q:A:270 R NH2
1qha:A:381 R NH2
3ksr:A:77 R NH2
1f4a:A:561 R NH2
2akf:A:453 R NH2
4rg1:A:93 R NH2
1e1c:A:584 R NH2

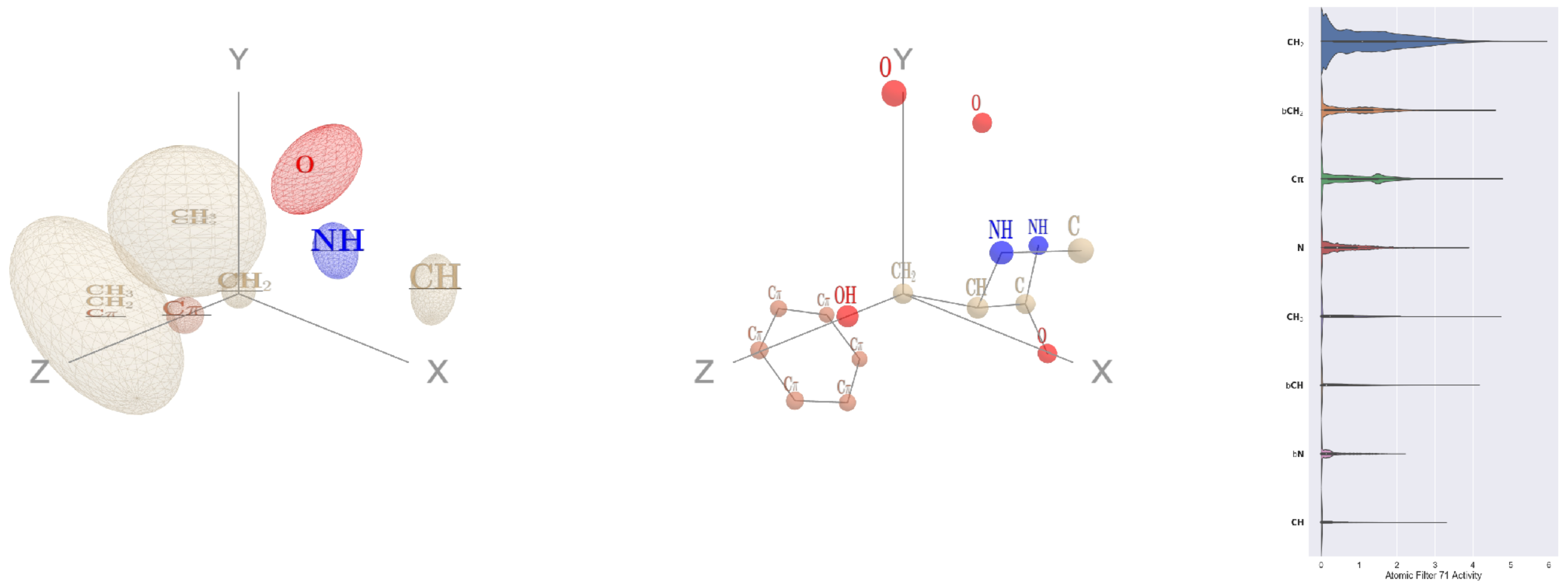


Filter 71
$R_{\text{pred}}^{\text{max}} = -0.04$
$R_{\text{pred}}^{\text{mean}} = -0.11$
Percent Active = 43
Pooling attention 0.39
Pooling feature 0.82
Top activators
4c2m:A:675 S CB
3ucq:A:145 Y CB
4ldb:A:64 H CB
2d32:A:180 Y CB
1o7g:A:352 F CB
2i71:B:128 H CB
2wk4:B:452 Q CG
4v4k:M:223 Y CB
1d0n:A:638 H CB
1lnl:A:77 F CB

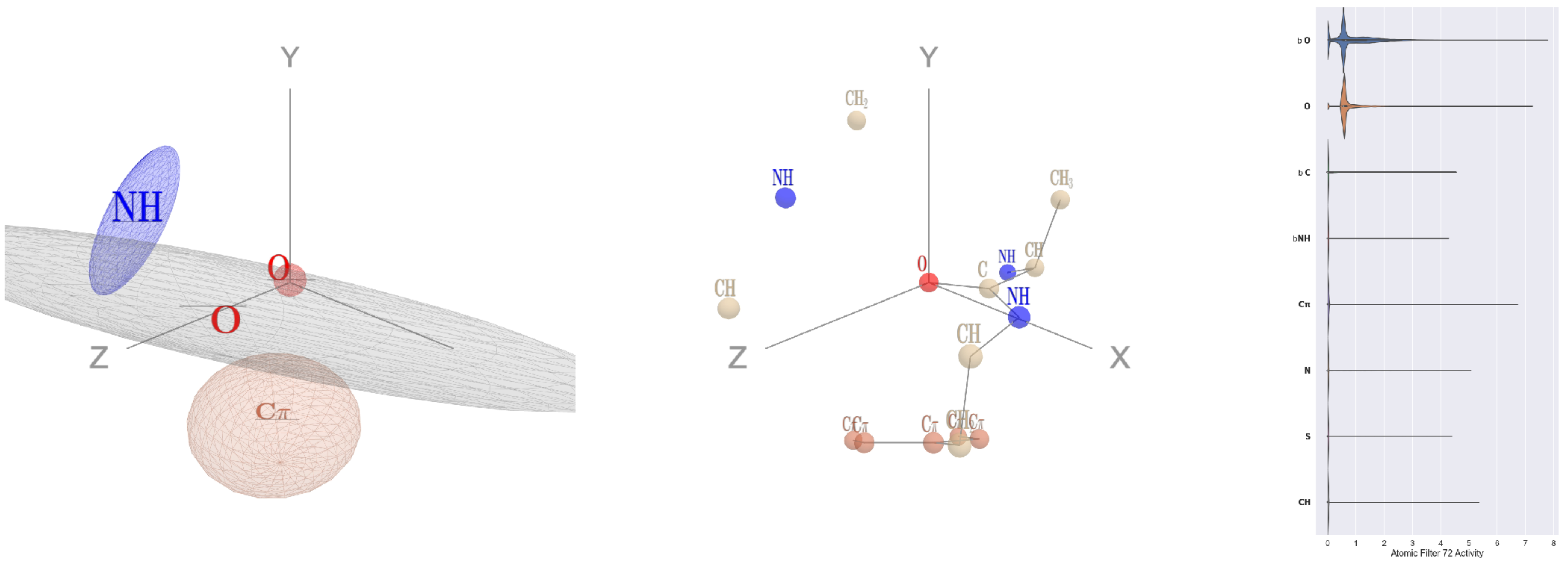


Filter 72
$R_{\text{pred}}^{\text{max}} = -0.03$
$R_{\text{pred}}^{\text{mean}} = -0.06$
Percent Active = 23
Pooling attention 0.10
Pooling feature 0.40
Top activators
2wk4:B:161 A O
3b8e:A:858 G O
4d10:K:4 A O
1bcp:A:167 V O
3kyh:C:249 K O
4e0s:B:788 D O
3wr9:A:202 N O
2pys:B:8 C O
2x0q:A:309 G O
1ewy:A:161 P O

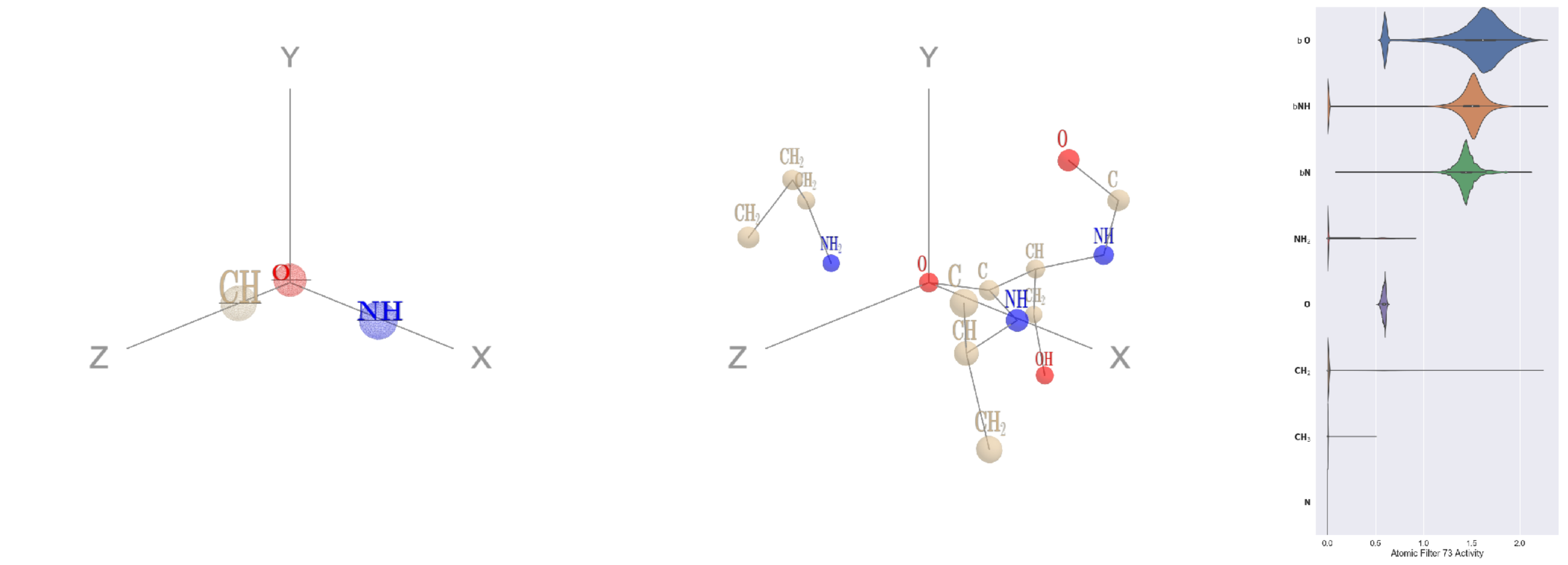


Filter 73
$R_{\text{pred}}^{\text{max}} = 0.01$
$R_{\text{pred}}^{\text{mean}} = -0.08$
Percent Active = 30
Pooling attention 0.18
Pooling feature 0.02
Top activators
3n99:G:264 S O
4nnq:B:180 A O
3ucq:A:198 Y O
4cht:B:126 P O
4c0b:B:231 Q O
4pfp:A:12 P O
4u0u:A:434 T O
1l7k:B:59 A O
3l09:B:52 E O
4nnq:B:153 P O

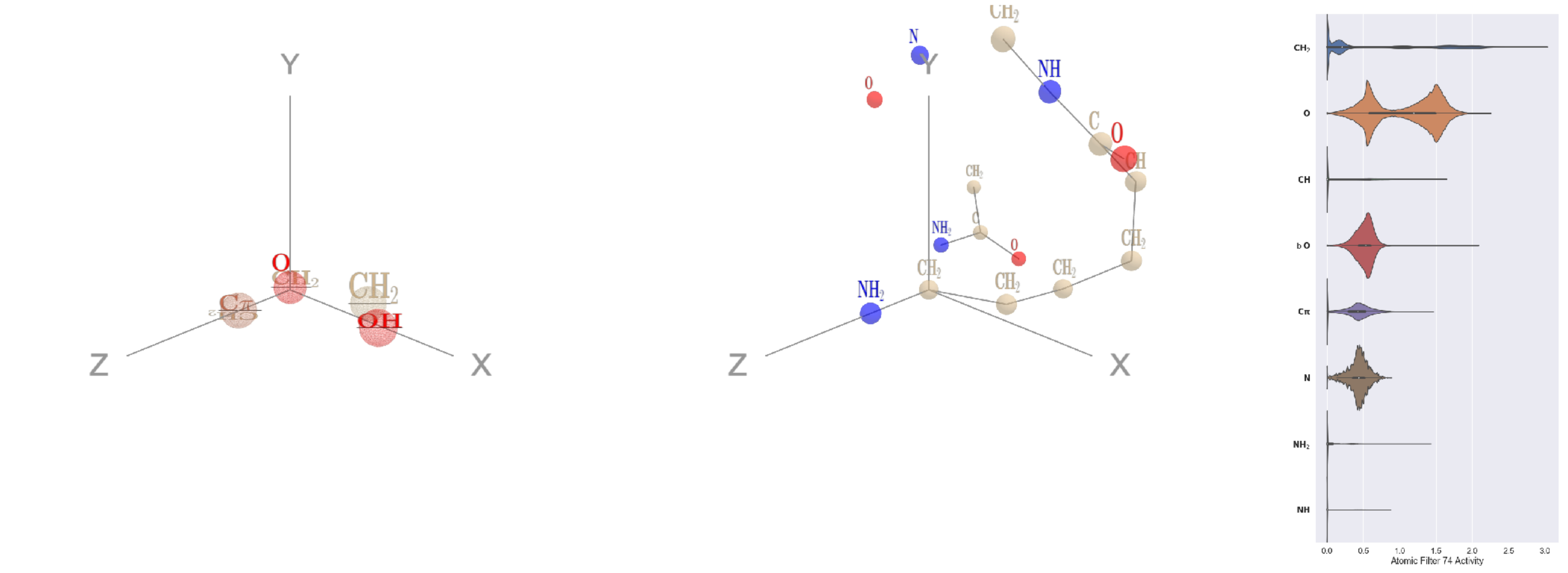


Filter 74
$R_{\text{pred}}^{\text{max}} = 0.07$
$R_{\text{pred}}^{\text{mean}} = 0.06$
Percent Active = 38
Pooling attention 0.09
Pooling feature 0.90
Top activators
1lnl:A:275 K CE
3o5t:A:81 R CD
3ucq:A:126 M CG
3ax7:A:779 M CG
1lnl:A:229 R CD
3o5t:A:198 R CD
3ucq:A:87 Q CG
3ucq:A:551 R CD
3hgi:A:33 R CD
3n99:G:71 R CD

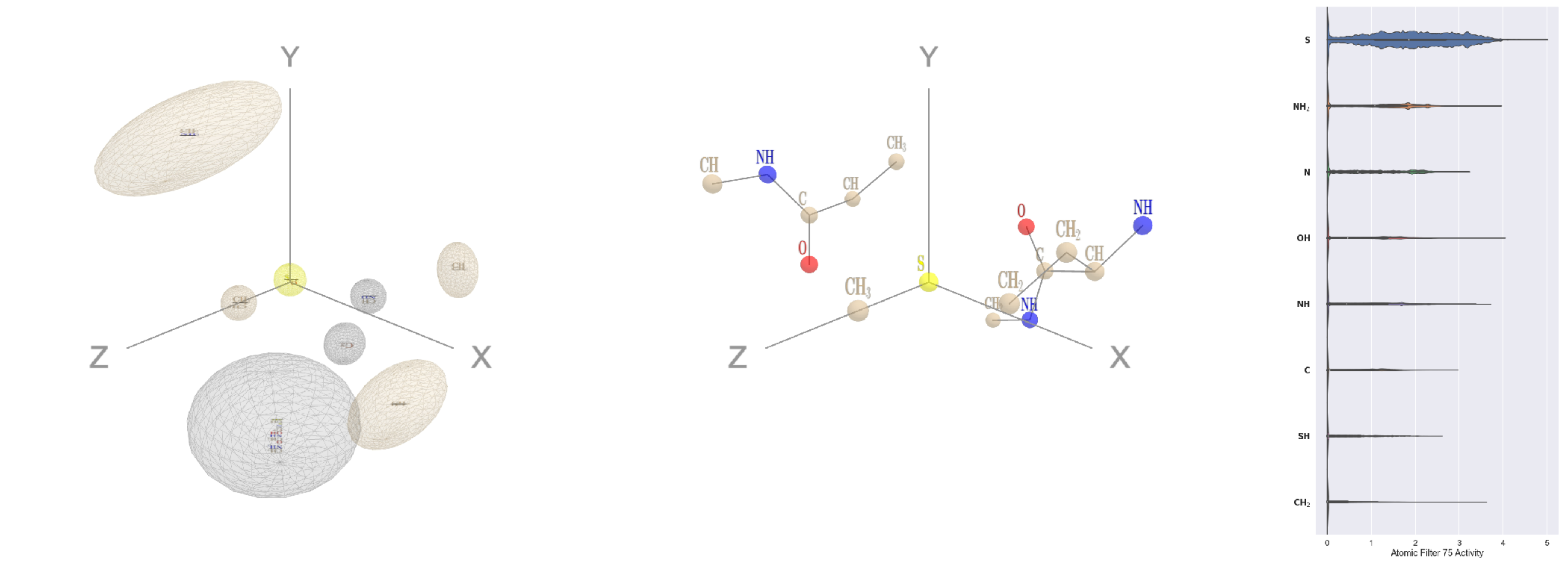


Filter 75
$R_{\text{pred}}^{\text{max}} = 0.20$
$R_{\text{pred}}^{\text{mean}} = 0.21$
Percent Active = 39
Pooling attention 0.61
Pooling feature 0.90
Top activators
1fyz:C:35 M SD
4g59:C:111 M SD
4pht:A:309 M SD
2nyg:A:68 M SD
1wnl:A:1 M SD
1o7g:A:366 M SD
4c2m:A:471 M SD
4o93:A:1 M SD
3nh5:A:212 M SD
1lnl:A:63 M SD

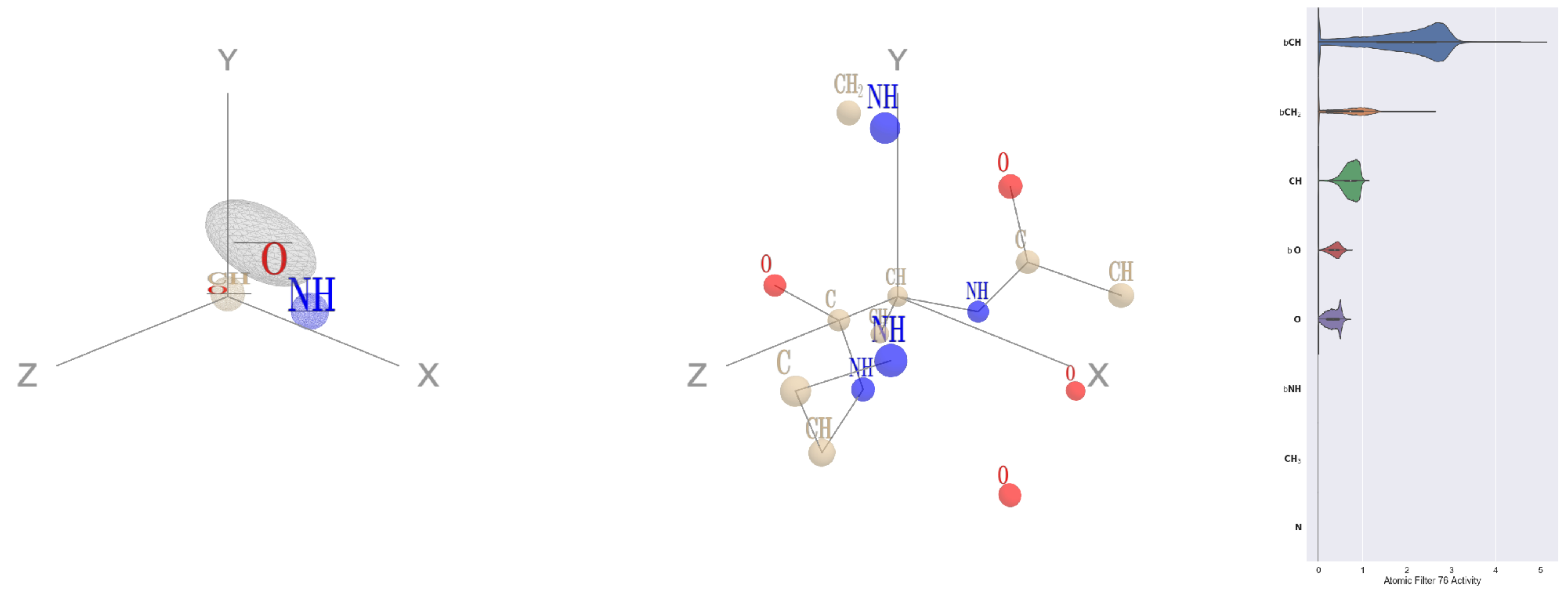


Filter 76
$R_{\mathrm{pred}}^{\mathrm{max}} = -0.01$
$R_{\mathrm{pred}}^{\mathrm{mean}} = -0.05$
Percent Active = 30
Pooling attention 0.14
Pooling feature 0.14
Top activators
3a5c:G:182 V CA
3a5c:G:184 E CA
3a5c:G:164 L CA
3a5c:G:25 V CA
3a5c:G:143 T CA
2b0o:F:619 A CA
1u5e:A:124 E CA
3nqr:D:67 Q CA
3a5c:G:17 Q CA
2wsc:B:398 Y CA

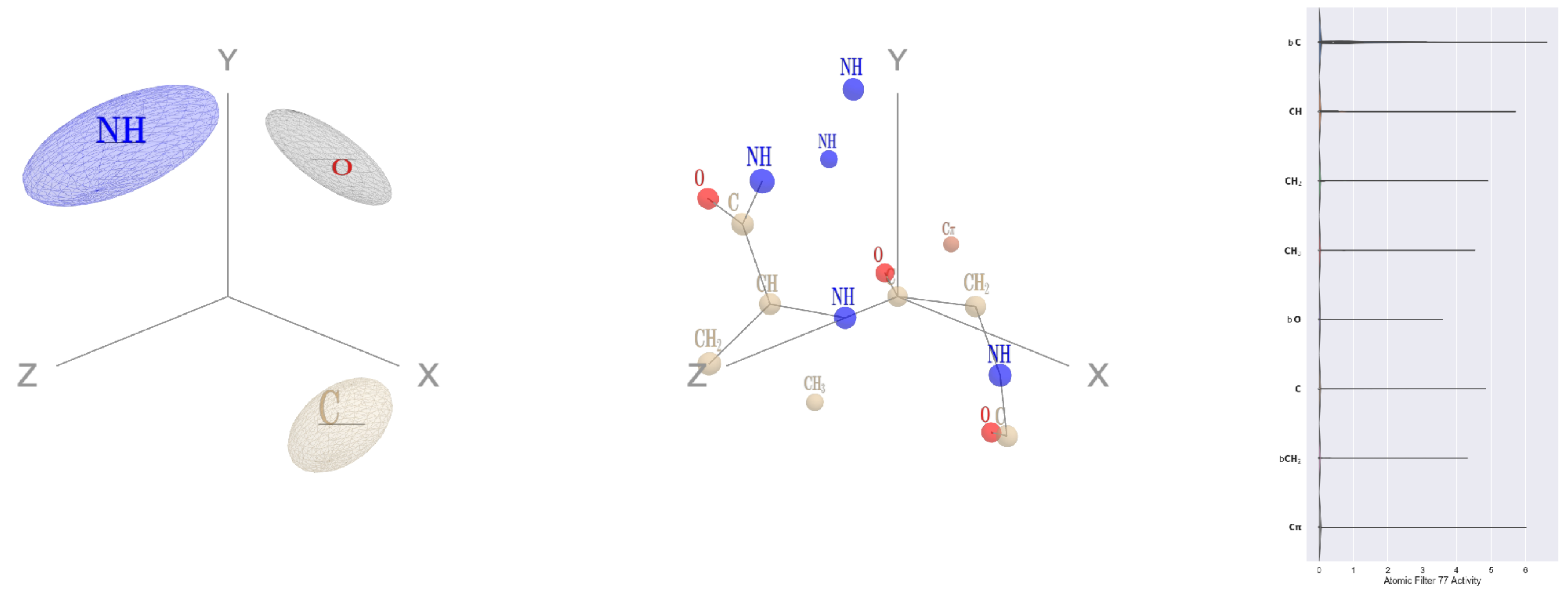


Filter 77
$R_{\mathrm{pred}}^{\mathrm{max}} = -0.02$
$R_{\mathrm{pred}}^{\mathrm{mean}} = -0.05$
Percent Active = 19
Pooling attention 0.18
Pooling feature 0.09
Top activators
4nmn:A:211 G C
3o47:A:159 G C
4pht:A:265 G C
2d73:A:443 S C
3ax7:A:1041 G C
3tsy:A:636 M C
3u33:A:444 G C
1vpn:B:273 C C
1jqn:A:540 G C
1r7r:A:129 N C

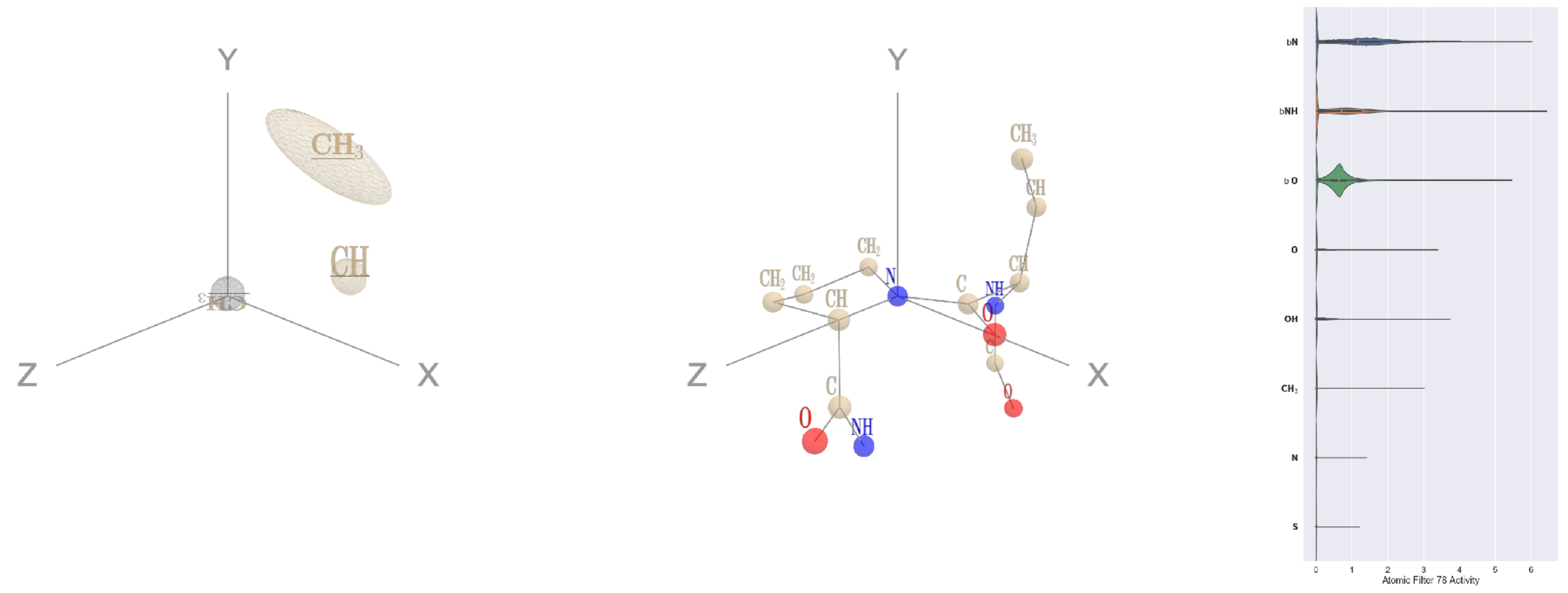


Filter 78
$R_{\mathrm{pred}}^{\mathrm{max}} = -0.01$
$R_{\mathrm{pred}}^{\mathrm{mean}} = -0.05$
Percent Active = 26
Pooling attention 0.09
Pooling feature 0.05
Top activators
4l8n:A:87 P N
4l8n:A:302 P N
2nr6:A:253 P N
2iwh:B:223 P N
3t3o:A:44 P N
4asi:C:1639 P N
4u4c:A:619 P N
4rnd:A:179 P N
2zy4:F:211 P N
4ot9:A:590 P N

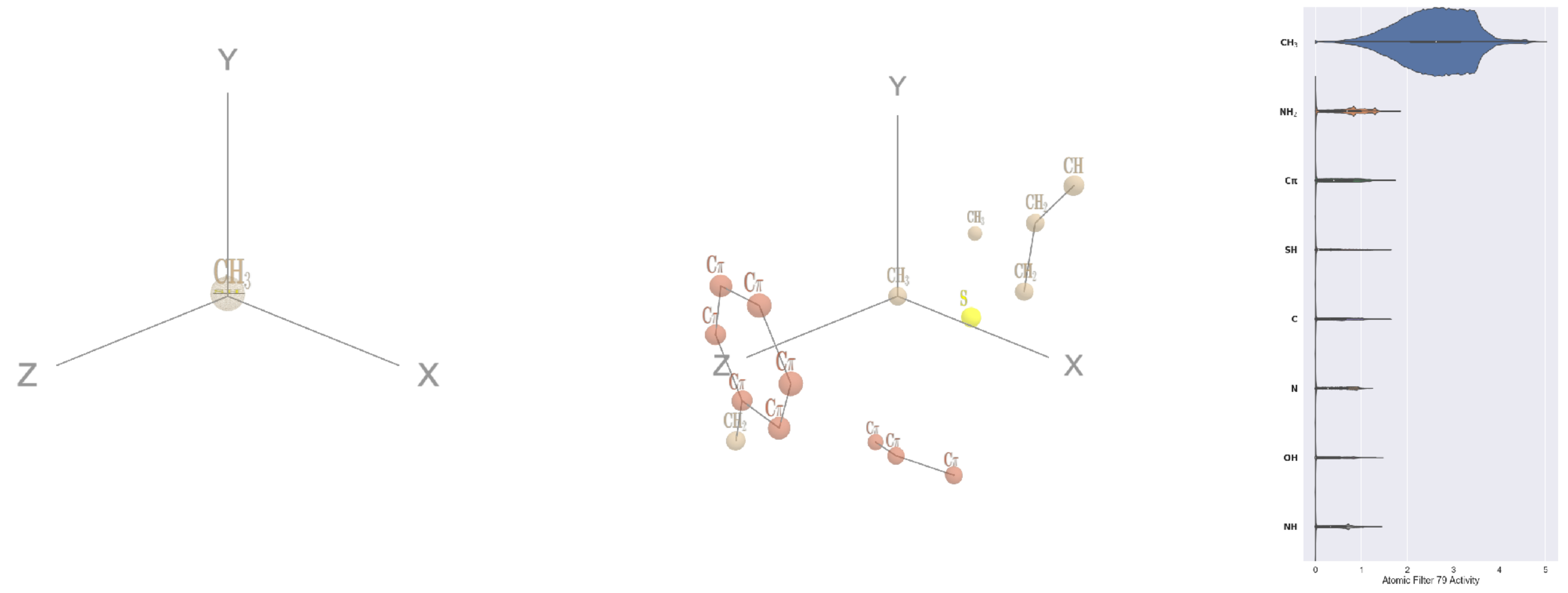


Filter 79
$R_{\mathrm{pred}}^{\mathrm{max}} = 0.08$
$R_{\mathrm{pred}}^{\mathrm{mean}} = 0.12$
Percent Active = 40
Pooling attention 0.18
Pooling feature 2.06
Top activators
1lk5:A:193 M CE
4c2m:A:18 I CD1
4c2m:A:1144 L CD2
1f4a:A:968 M CE
2zbk:B:409 M CE
3i0t:B:71 M CE
4buj:B:1252 L CD1
2iwh:B:857 M CE
1lql:A:1 M CE
4c2m:A:1185 V CG1

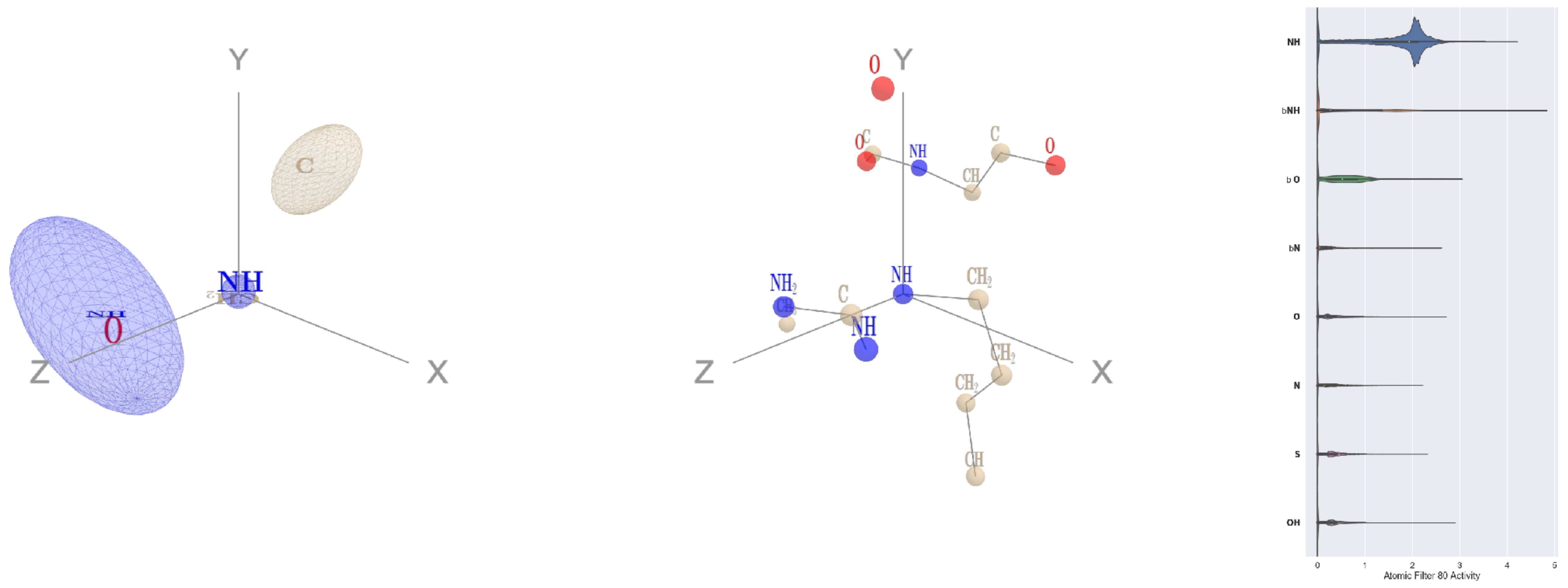


Filter 80
$R_{\text{pred}}^{\max} = 0.05$
$R_{\text{pred}}^{\text{mean}} = 0.03$
Percent Active = 46
Pooling attention 0.09
Pooling feature 0.90
Top activators
2hye:B:143 R NE
3hwp:A:124 W NE1
4mex:C:758 R NE
2wgq:B:696 W NE1
2f8x:C:281 R NE
3h8a:E:23 R NE
1dtn:A:315 W NE1
4cy6:A:250 W NE1
4kzx:E:235 W NE1
2wgq:B:69 W NE1

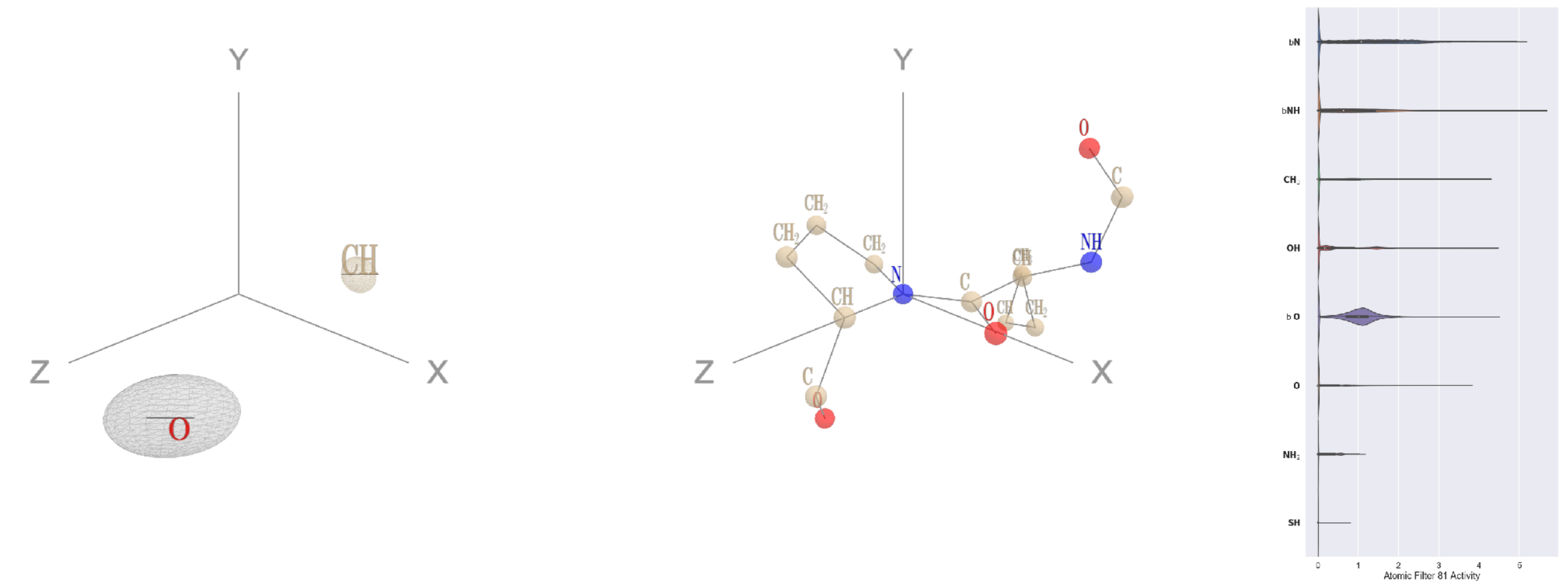


Filter 81
$R_{\text{pred}}^{\max} = -0.00$
$R_{\text{pred}}^{\text{mean}} = -0.03$
Percent Active = 29
Pooling attention 0.34
Pooling feature 0.06
Top activators
4u0u:A:441 P N
4pfp:A:10 P N
3t3o:A:44 P N
4nnq:B:17 P N
3t3o:A:99 P N
4cht:B:153 P N
3t3o:A:391 P N
3jsb:A:138 P N
3o5t:A:265 P N
4cht:B:124 P N

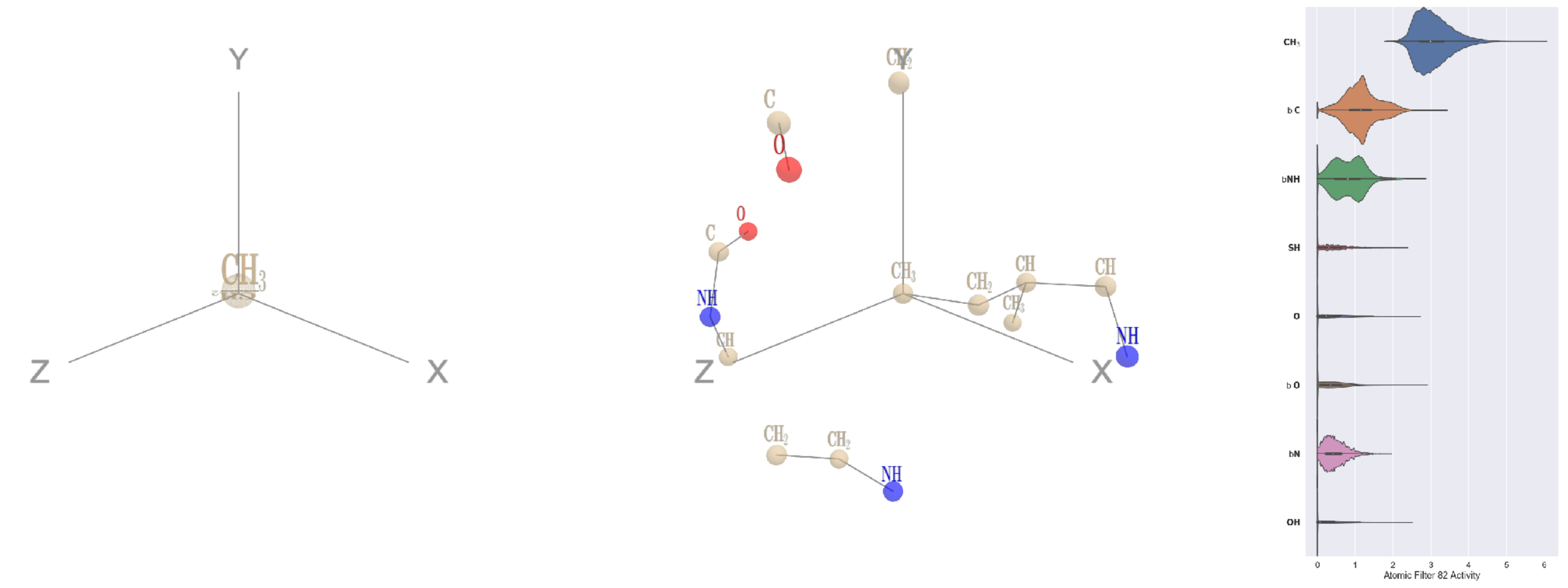


Filter 82
$R_{\text{pred}}^{\max} = -0.07$
$R_{\text{pred}}^{\text{mean}} = -0.12$
Percent Active = 64
Pooling attention 0.14
Pooling feature 2.33
Top activators
3tsy:A:196 I CD1
3a1m:A:80 M CE
1bh3:A:100 M CE
1o7g:A:366 M CE
1ewy:A:234 M CE
1vl4:A:238 M CE
1cmx:A:206 L CD1
1q18:A:196 V CG2
1wsr:A:334 M CE
4buj:B:873 V CG1

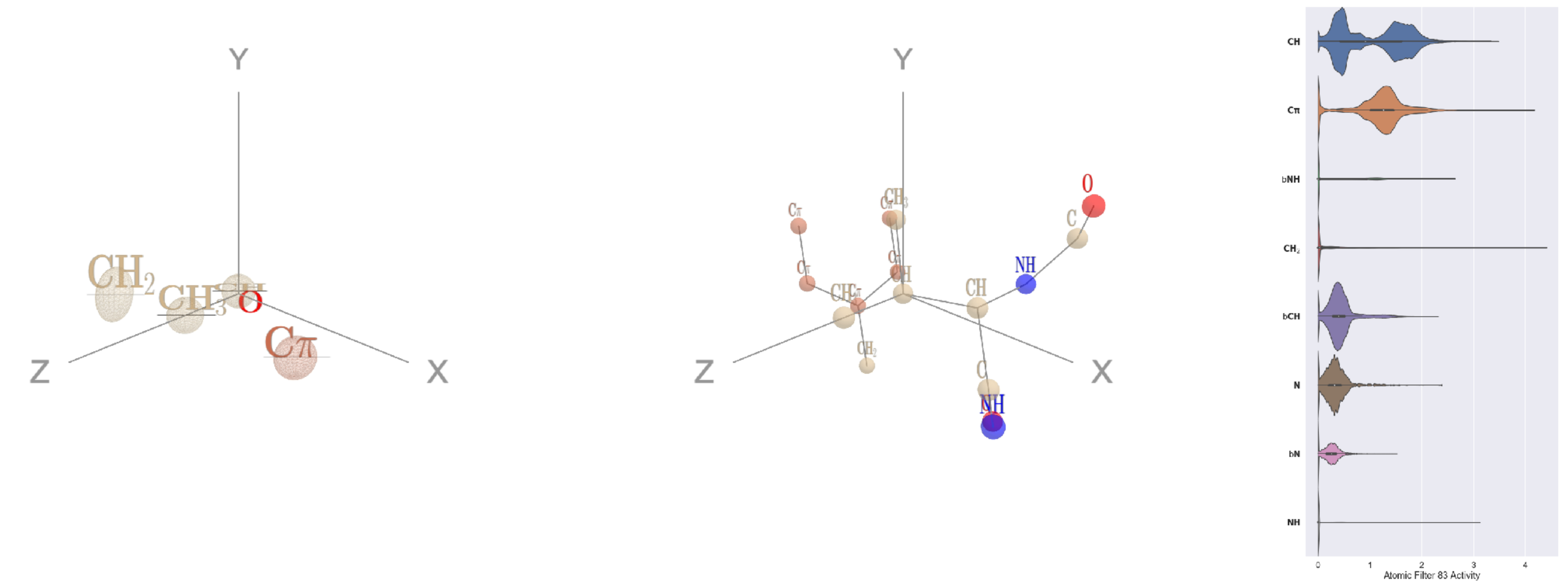


Filter 83
$R_{\text{pred}}^{\max} = 0.03$
$R_{\text{pred}}^{\text{mean}} = 0.04$
Percent Active = 44
Pooling attention 0.11
Pooling feature 1.63
Top activators
1ko6:A:784 V CB
1pgu:B:598 L CG
1f4a:A:478 V CB
2wsc:B:584 L CG
2f8x:C:34 V CB
2cjr:A:340 L CG
1o7g:A:182 L CG
1p4e:B:38 L CG
1lnl:A:75 V CB
1bo1:A:79 V CB

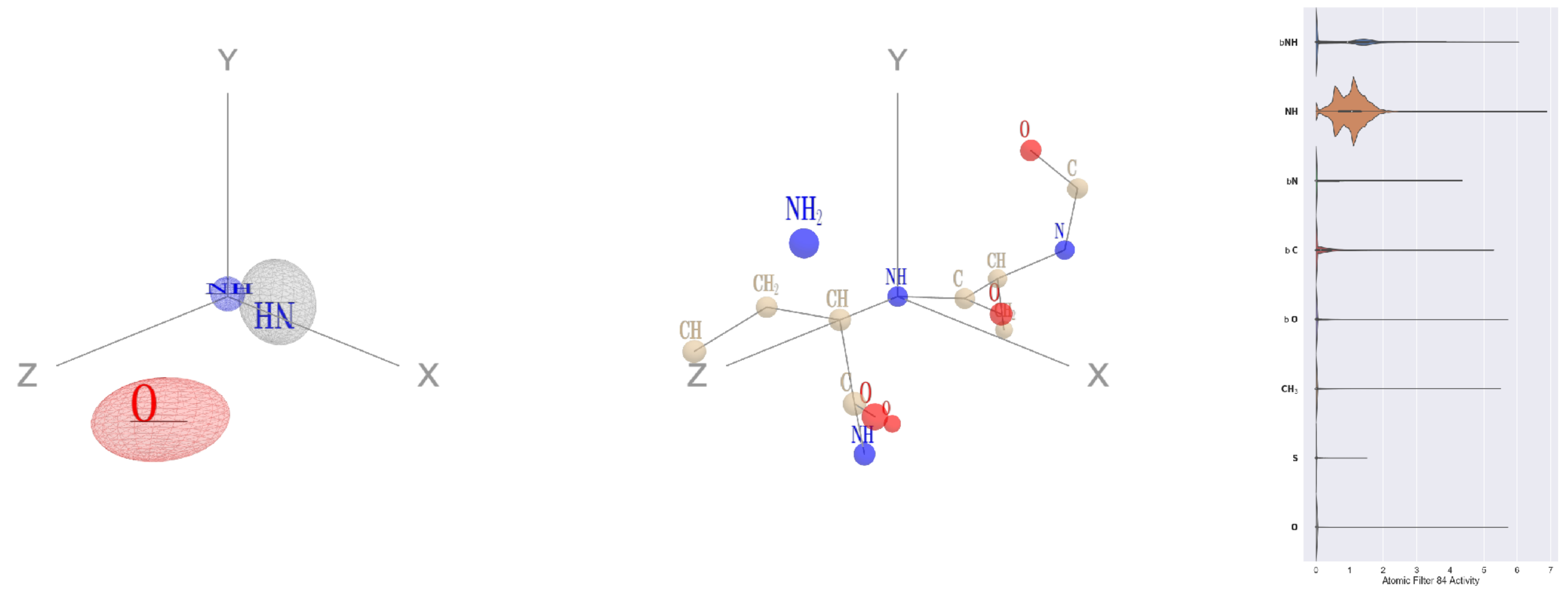


Filter 84
$R_{\mathrm{pred}}^{\mathrm{max}} = 0.01$
$R_{\mathrm{pred}}^{\mathrm{mean}} = -0.04$
Percent Active = 29
Pooling attention 0.39
Pooling feature 0.25
Top activators
1am4:A:132 L N
3wrx:C:1020 F N
4gu0:A:177 I N
4v81:H:268 H N
3igh:X:31 F N
4dag:A:178 N N
1qc9:A:82 F N
3hzu:A:294 V N
2pmz:D:150 G N
1lnl:A:292 V N

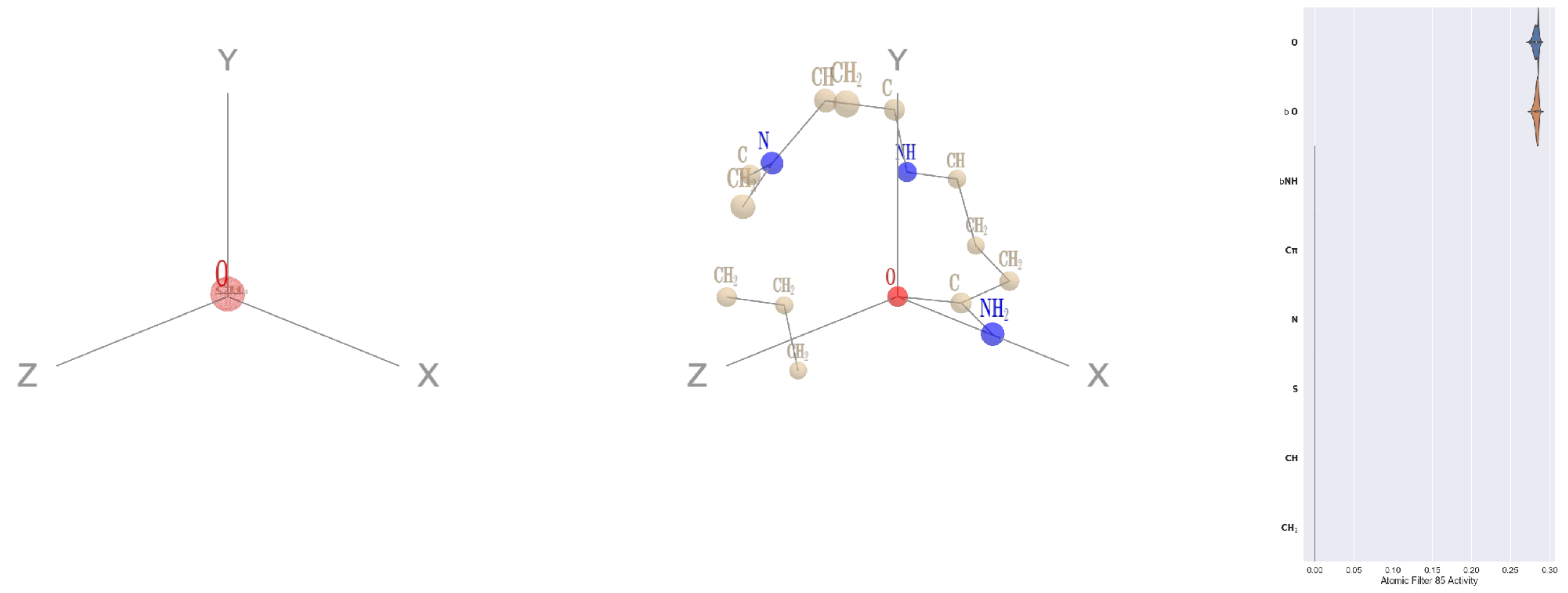


Filter 85
$R_{\mathrm{pred}}^{\mathrm{max}} = 0.06$
$R_{\mathrm{pred}}^{\mathrm{mean}} = -0.05$
Percent Active = 15
Pooling attention 0.00
Pooling feature 0.00
Top activators
3fbv:D:801 Q OE1
1dvk:B:153 N OD1
1cm5:A:515 D OD1
1efu:B:183 E OE1
4whj:A:285 E OE1
4ri0:A:210 Q OE1
2o35:B:7 E OE1
1p32:A:177 D OD1
1fps:A:308 E OE1
3ztv:A:305 E OE1

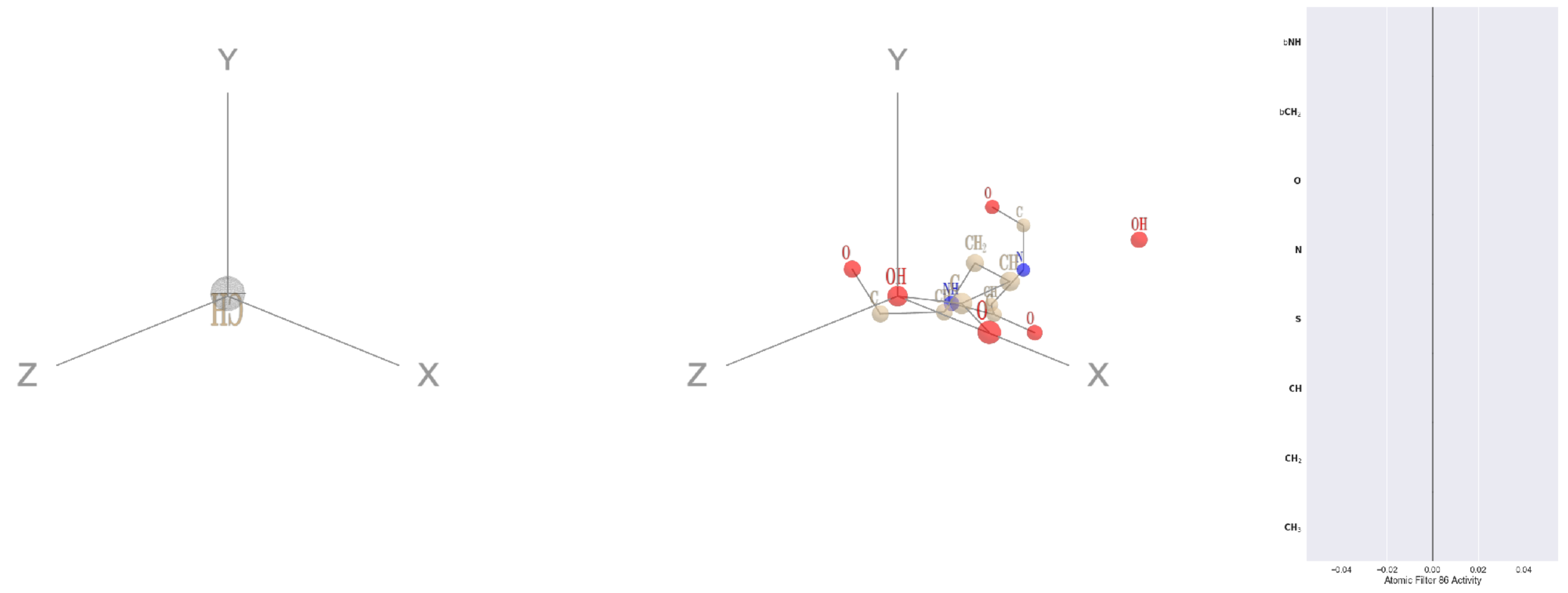


Filter 86
$R_{\mathrm{pred}}^{\mathrm{max}} = nan$
$R_{\mathrm{pred}}^{\mathrm{mean}} = nan$
Percent Active = 0
Pooling attention 0.00
Pooling feature 0.00
Top activators
5wy5:A:246 E OE2
2b0o:F:573 F N
2b0o:F:574 G O
2b0o:F:574 G C
2b0o:F:574 G CA
2b0o:F:574 G N
2b0o:F:573 F CZ
2b0o:F:573 F CE2
2b0o:F:573 F CE1
2b0o:F:573 F CD2

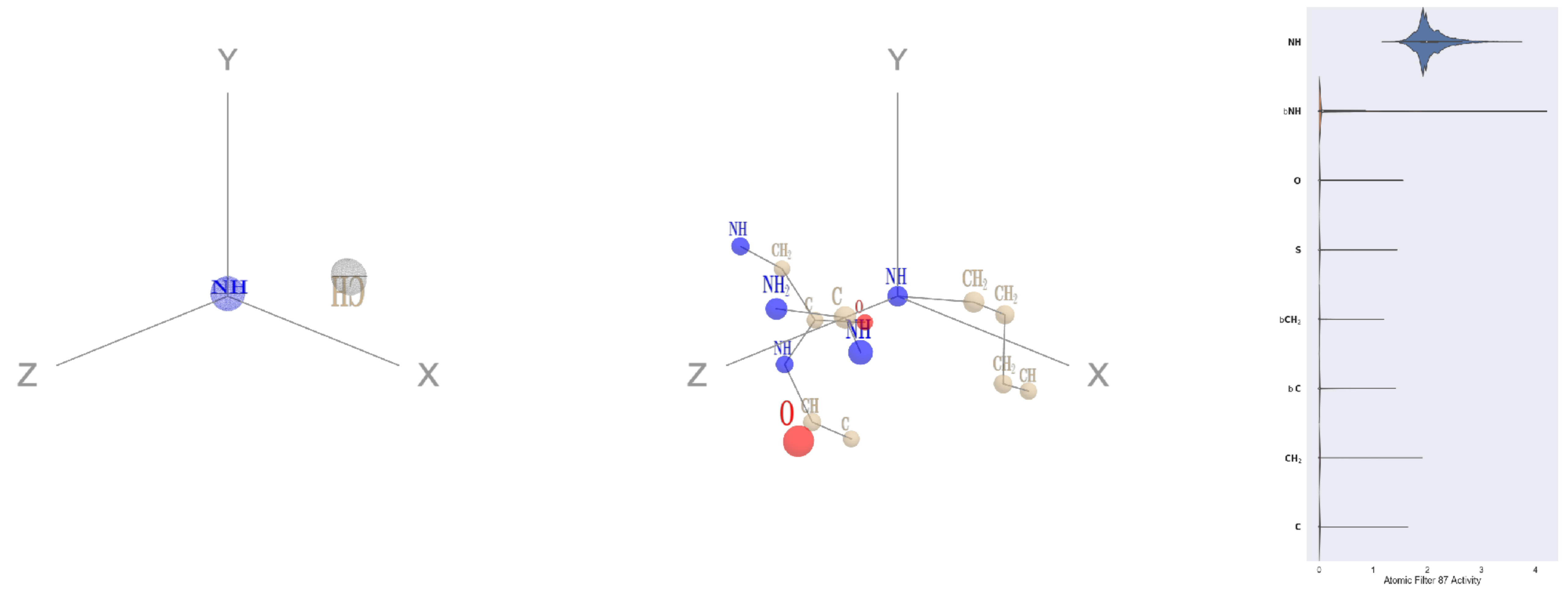


Filter 87
$R_{\mathrm{pred}}^{\mathrm{max}} = 0.04$
$R_{\mathrm{pred}}^{\mathrm{mean}} = 0.03$
Percent Active = 17
Pooling attention 0.32
Pooling feature 0.23
Top activators
2ba0:B:208 R NE
2zbk:B:193 R NH1
4u9c:B:156 R NH1
1h4s:A:168 R NH1
3qzy:B:159 R NH1
2wsc:B:292 R NH1
4v81:H:380 R NH1
1fyz:C:122 R NH1
3drw:A:227 R NH1
1f4a:A:561 R NH1

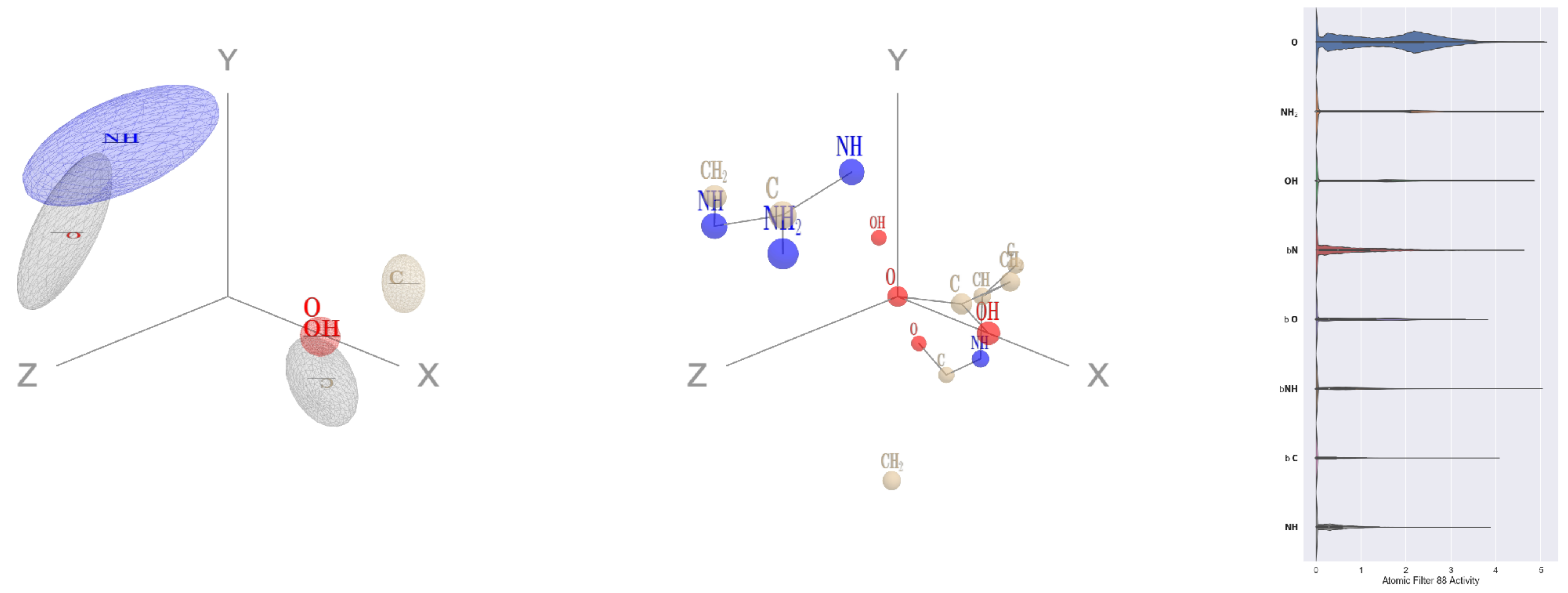


Filter 88
$R_{\text{pred}}^{\text{max}} = 0.00$
$R_{\text{pred}}^{\text{mean}} = -0.03$
Percent Active = 35
Pooling attention 0.25
Pooling feature 0.85
Top activators
4j4w:A:123 D OD1
1w36:B:348 E OE1
2ba0:B:107 D OD1
1qha:A:76 E OE1
5uqy:A:46 D OD1
4wj3:B:165 E OE1
4hh2:D:279 D OD1
3b8m:A:283 E OE1
1pgu:B:393 D OD1
1vpn:B:121 E OE1

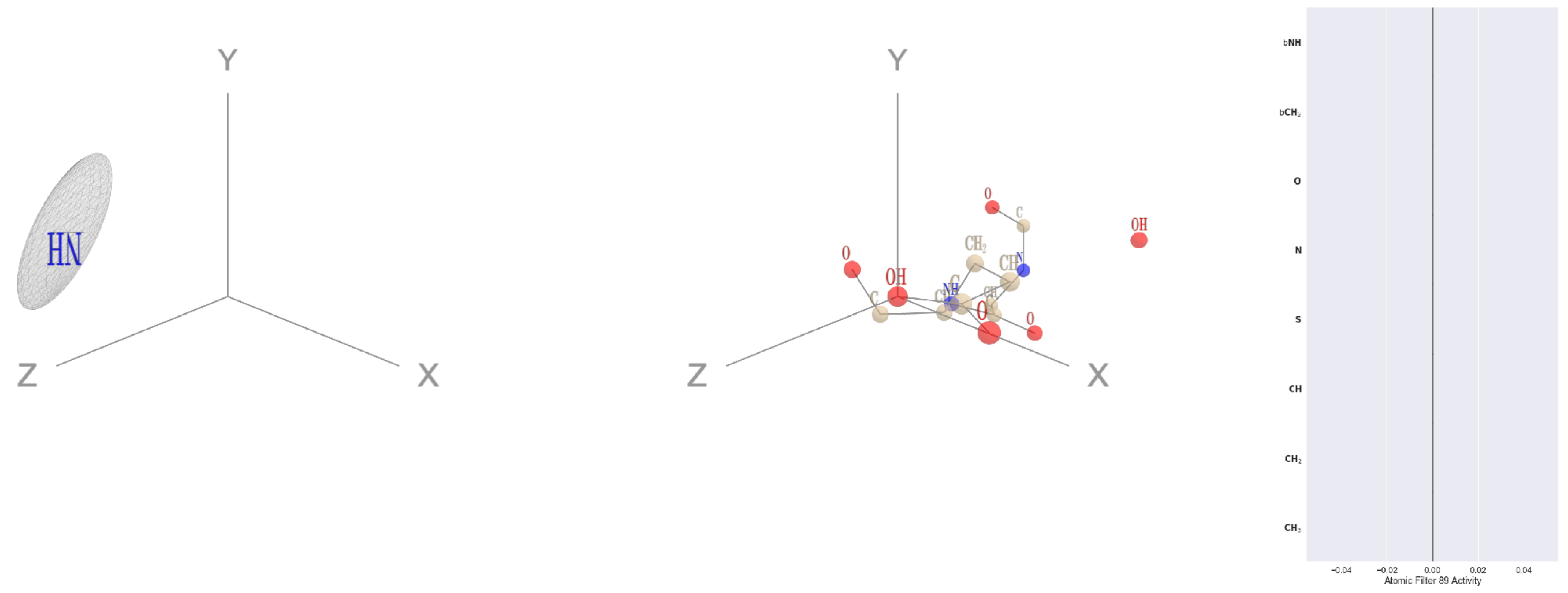


Filter 89
$R_{\text{pred}}^{\text{max}} = nan$
$R_{\text{pred}}^{\text{mean}} = nan$
Percent Active = 0
Pooling attention 0.00
Pooling feature 0.00
Top activators
5wy5:A:246 E OE2
2b0o:F:573 F N
2b0o:F:574 G O
2b0o:F:574 G C
2b0o:F:574 G CA
2b0o:F:574 G N
2b0o:F:573 F CZ
2b0o:F:573 F CE2
2b0o:F:573 F CE1
2b0o:F:573 F CD2

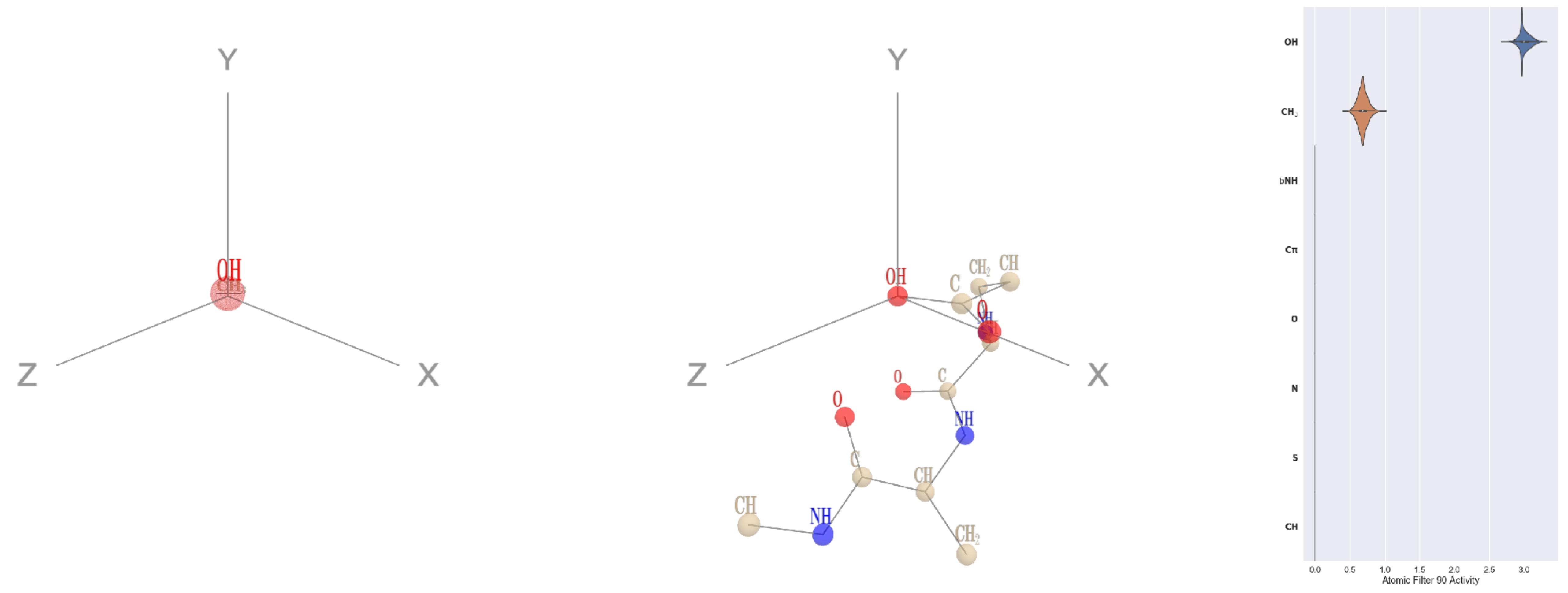


Filter 90
$R_{\text{pred}}^{\text{max}} = 0.01$
$R_{\text{pred}}^{\text{mean}} = -0.01$
Percent Active = 11
Pooling attention 0.04
Pooling feature 0.04
Top activators
2qn5:B:97 E OE2
1y4j:B:246 E OE2
3wrx:C:782 E OE2
2x98:A:77 E OE2
4itr:A:3749 E OE2
1efu:B:188 E OE2
3kyh:C:399 E OE2
3v4l:A:632 E OE2
4pfp:A:540 D OD2
1sl0:A:13 E OE2

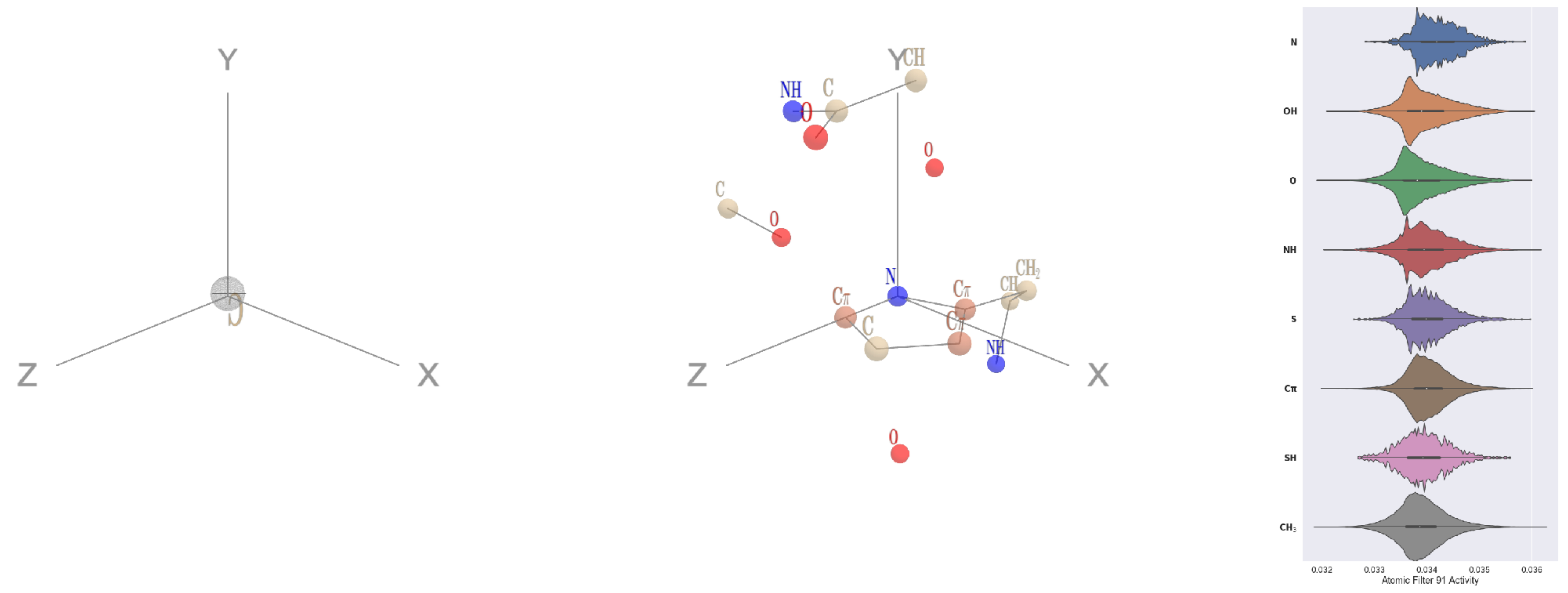


Filter 91
$R_{\text{pred}}^{\text{max}} = -0.04$
$R_{\text{pred}}^{\text{mean}} = 0.02$
Percent Active = 84
Pooling attention 0.00
Pooling feature 0.00
Top activators
2z67:A:397 H ND1
1yqg:A:158 H ND1
1jdw:A:325 H ND1
1vbk:A:187 H ND1
4pfp:A:11 H ND1
1ivu:A:517 H ND1
1iwa:A:238 H ND1
2iut:A:625 H ND1
1he8:A:1022 H ND1
2pm9:A:72 H ND1

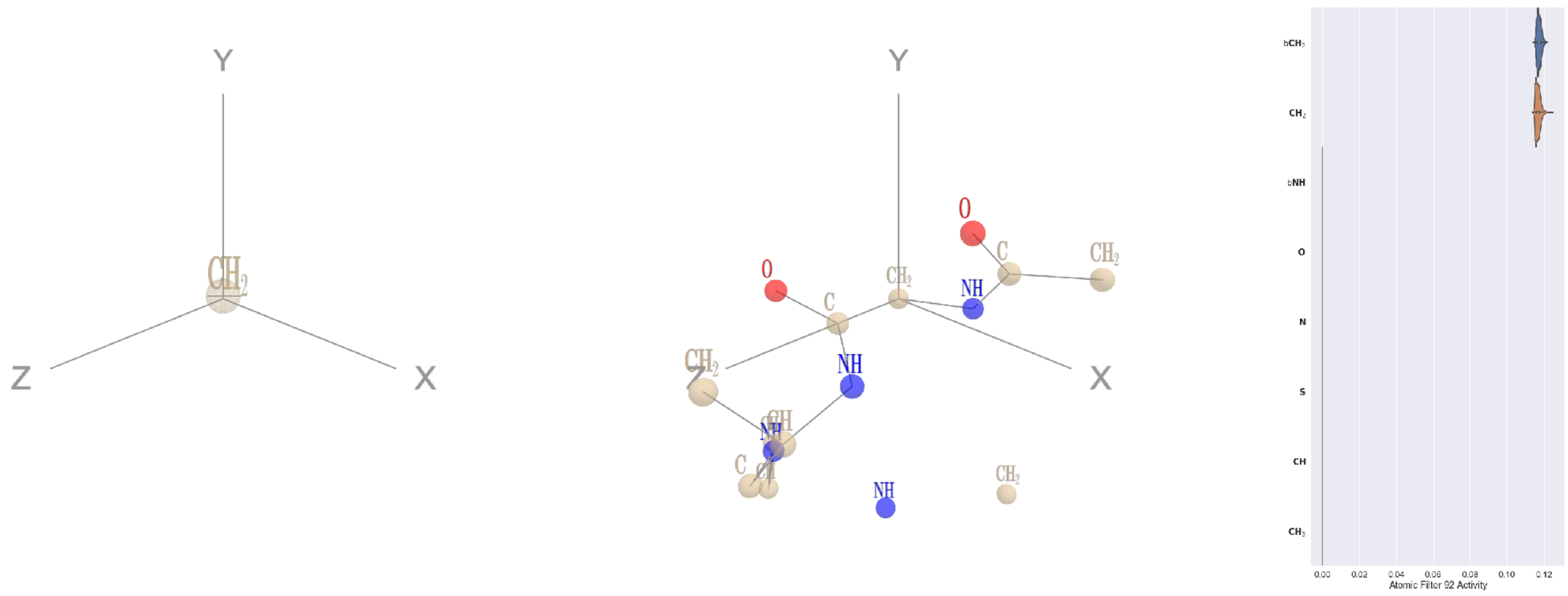


Filter 92
$R_{\text{pred}}^{\text{max}} = 0.04$
$R_{\text{pred}}^{\text{mean}} = 0.02$
Percent Active = 16
Pooling attention 0.00
Pooling feature 0.00
Top activators
4c8q:H:474 G CA
3dlb:A:388 G CA
1ppt:A:1 G CA
3irm:A:131 G CA
3c57:A:149 G CA
1umg:A:306 G CA
4v81:A:16 G CA
1fx0:B:28 G CA
4r3z:B:528 G CA
1bh3:A:221 G CA

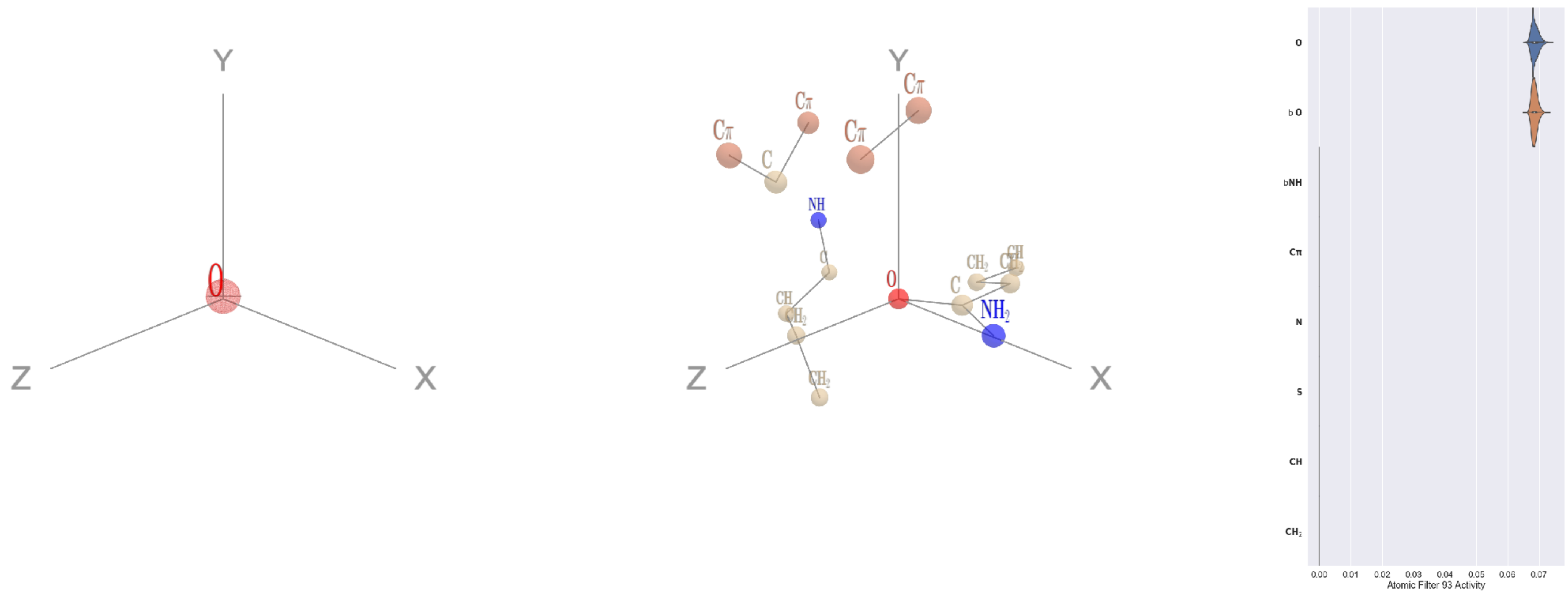


Filter 93
$R_{\text{pred}}^{\text{max}} = -0.07$
$R_{\text{pred}}^{\text{mean}} = -0.05$
Percent Active = 15
Pooling attention 0.00
Pooling feature 0.00
Top activators
4ldb:A:159 Q OE1
3t5v:C:88 N OD1
1e1c:A:357 Q OE1
3fbv:D:801 Q OE1
1jr7:A:78 Q OE1
1qha:A:783 E OE1
4c2m:A:874 E OE1
3pq1:A:511 Q OE1
4ny2:A:349 E OE1
1bxr:A:334 E OE1

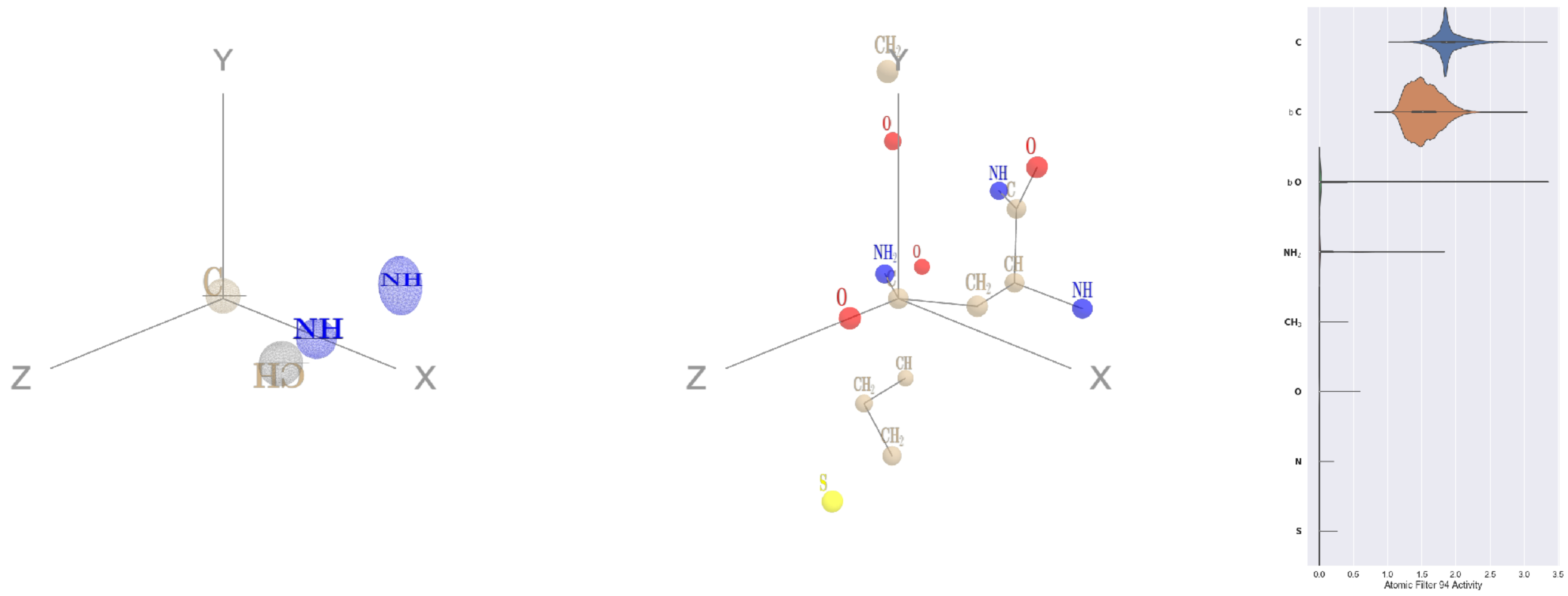


Filter 94
$R_{\text{pred}}^{\text{max}} = 0.03$
$R_{\text{pred}}^{\text{mean}} = 0.01$
Percent Active = 23
Pooling attention 0.06
Pooling feature 0.68
Top activators
2gy7:B:427 N CG
2iwh:B:432 N CG
4gfh:A:723 Q CD
1e1c:A:504 D CG
1ivu:A:592 H NE2
1bxr:A:996 E CD
4qiw:A:488 Q CD
1k72:B:605 N CG
3l0o:A:263 D CG
1h2i:A:117 D CG

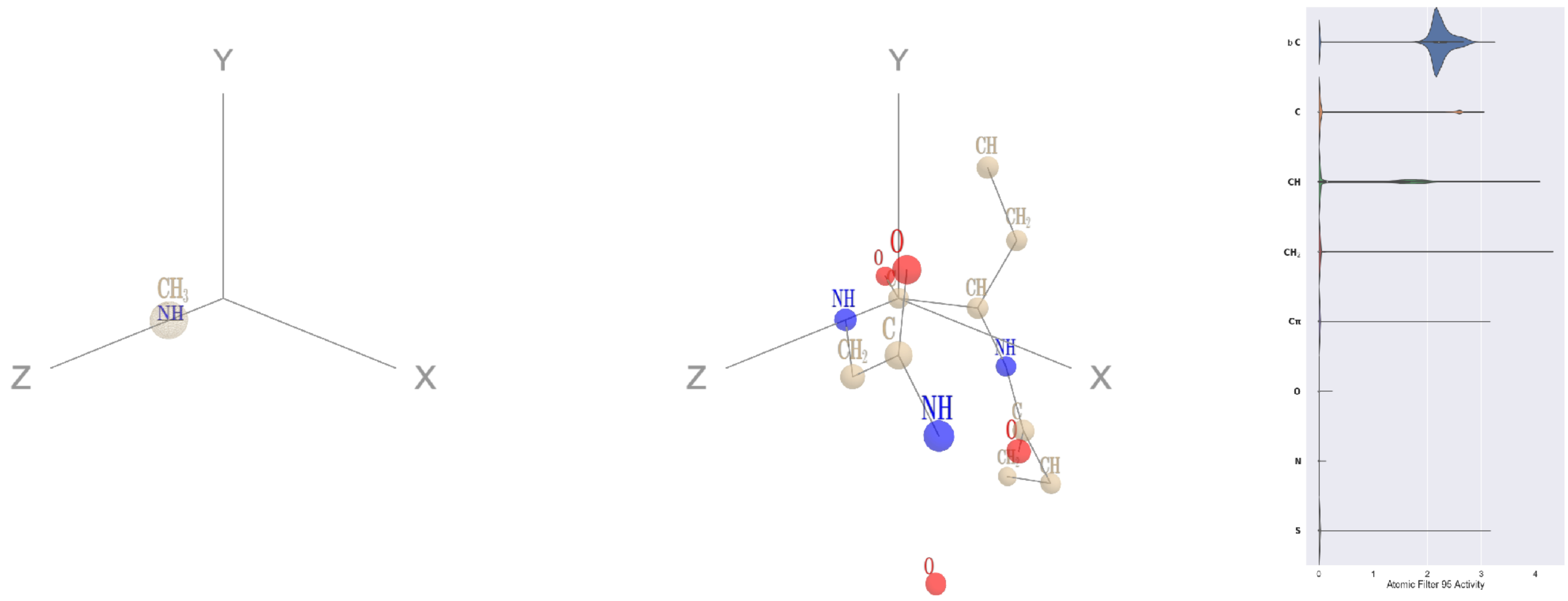


Filter 95
$R_{\text{pred}}^{\text{max}} = 0.01$
$R_{\text{pred}}^{\text{mean}} = -0.02$
Percent Active = 18
Pooling attention 0.13
Pooling feature 0.19
Top activators
4mex:C:626 E C
3l0o:A:269 D C
4r3z:B:452 F C
3l0o:A:79 L C
3hfd:A:89 N C
4v8p:BF:174 K C
4c0b:D:499 S C
4r3z:B:527 R C
4r3z:B:479 E C
3l0o:A:185 K C

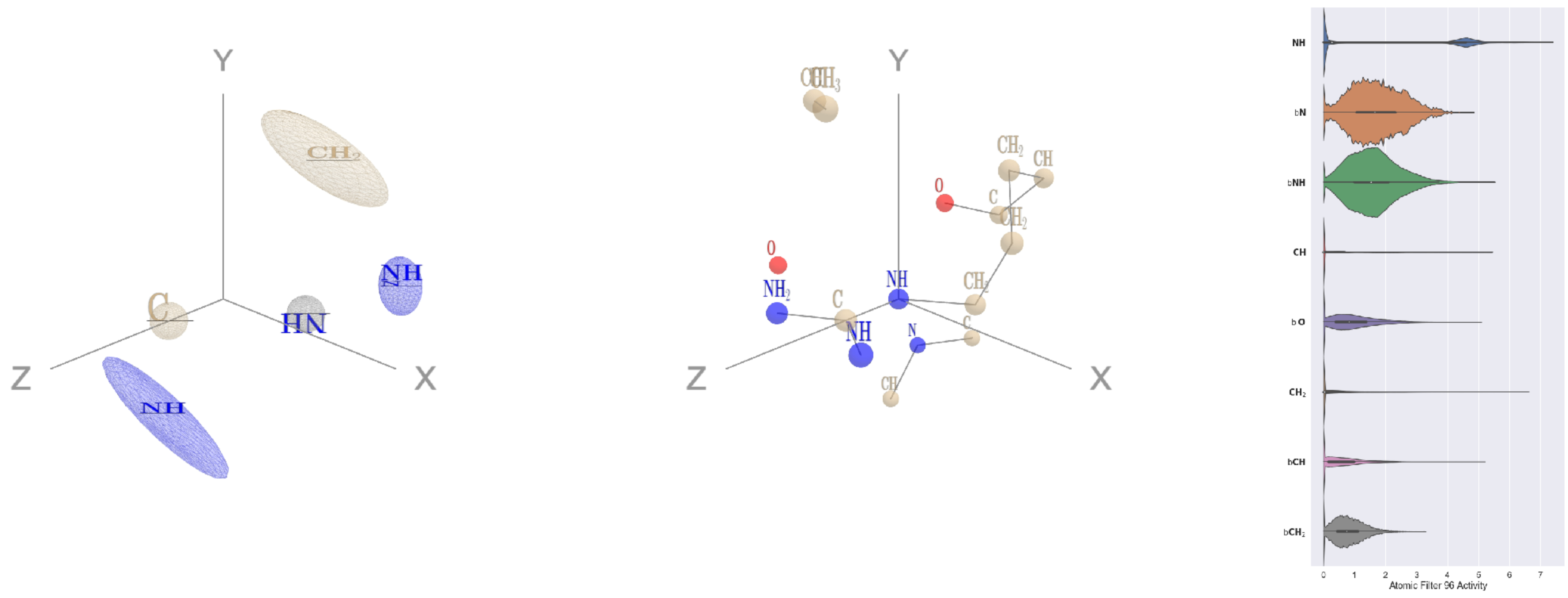


Filter 96
$R_{\text{pred}}^{\max} = 0.07$
$R_{\text{pred}}^{\text{mean}} = 0.01$
Percent Active = 63
Pooling attention 0.42
Pooling feature 0.75
Top activators
1iwa:A:139 R NE
3irm:A:118 R NE
3o47:A:112 R NE
2dm9:A:92 R NE
3drw:A:41 R NE
4u0u:A:374 R NE
3o5t:A:107 R NE
1v8f:A:227 R NE
1o7g:A:414 R NE
3zxu:A:247 R NE

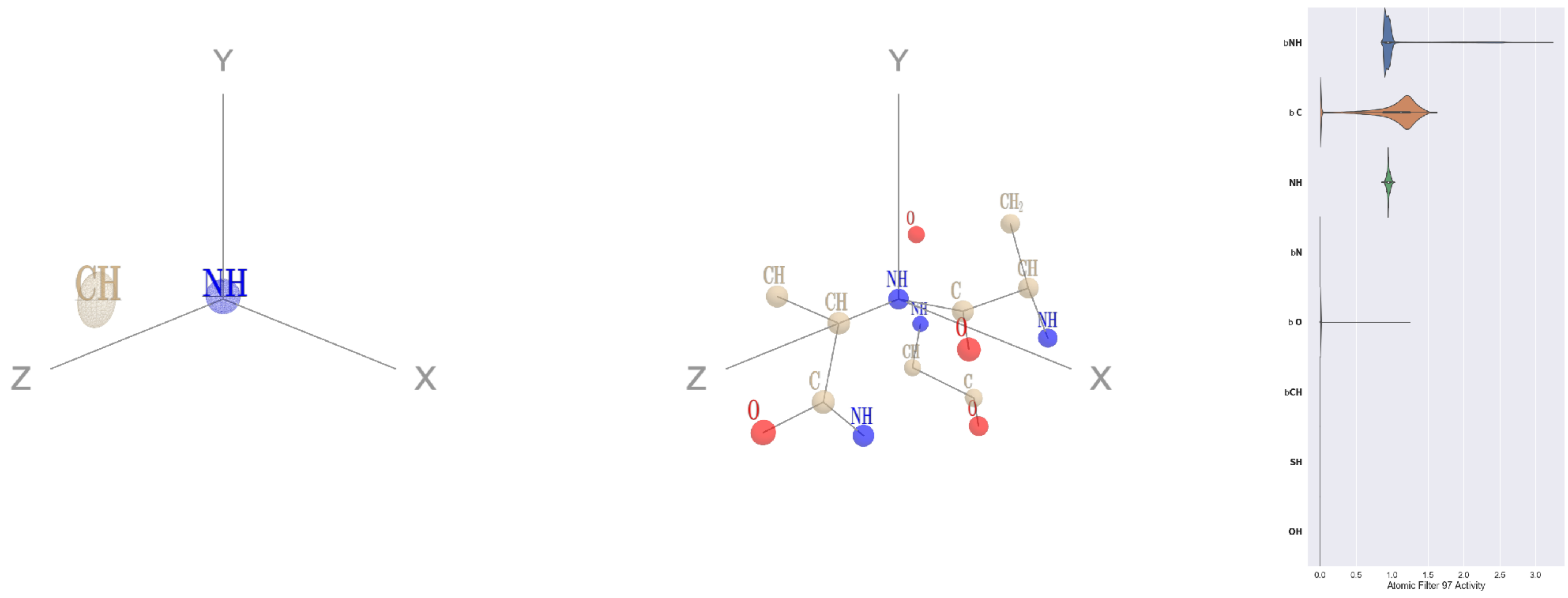


Filter 97
$R_{\text{pred}}^{\max} = -0.00$
$R_{\text{pred}}^{\text{mean}} = -0.05$
Percent Active = 25
Pooling attention 0.18
Pooling feature 0.01
Top activators
3a5c:G:138 I N
1lnl:A:7 V N
1bgv:A:209 T N
3a5c:G:168 V N
1fgu:A:251 V N
4kzx:L:122 I N
1w36:B:454 V N
4kzx:L:120 V N
1w36:B:787 V N
1lnl:A:74 T N

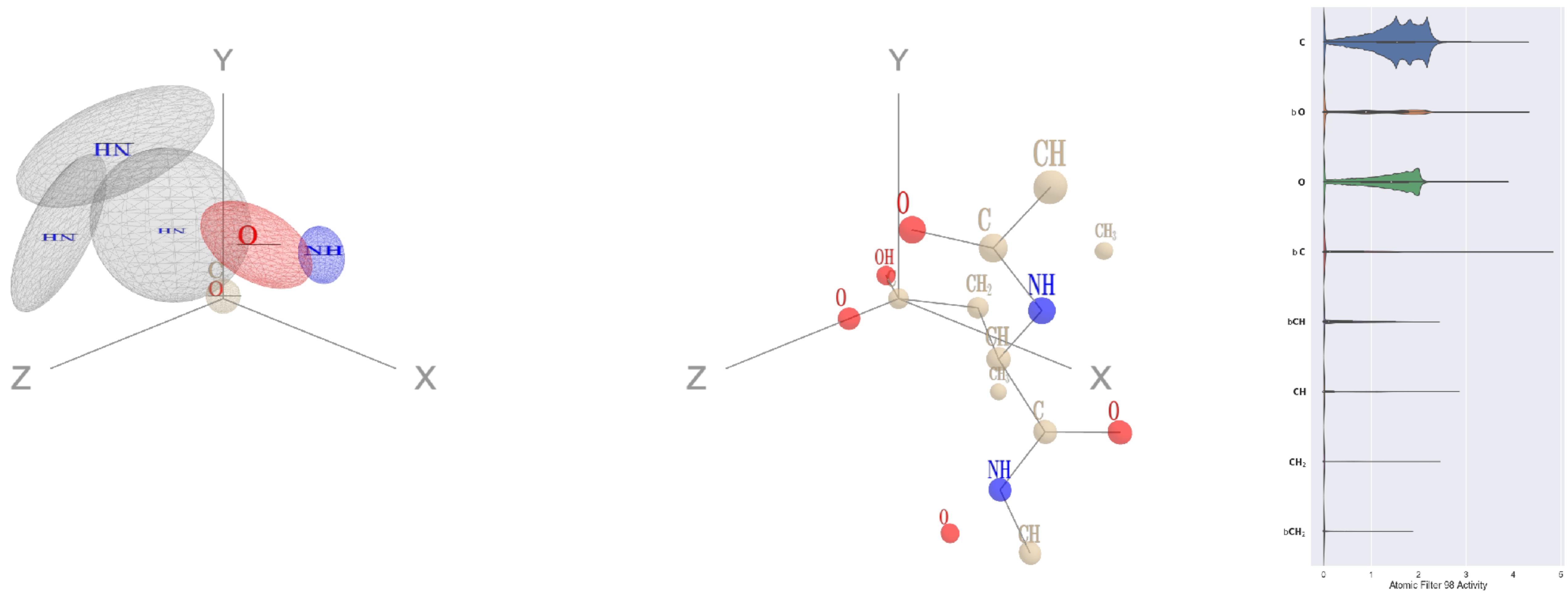


Filter 98
$R_{\text{pred}}^{\max} = 0.09$
$R_{\text{pred}}^{\text{mean}} = 0.05$
Percent Active = 34
Pooling attention 0.22
Pooling feature 1.43
Top activators
2zbk:B:514 D CG
1usv:B:101 D CG
2iut:A:433 D CG
4pba:A:82 N CG
4jdm:C:51 D CG
4ny2:A:417 Q CD
1lnl:A:128 D CG
4iqj:M:489 D CG
1qha:A:881 E CD
4e0s:B:736 N CG

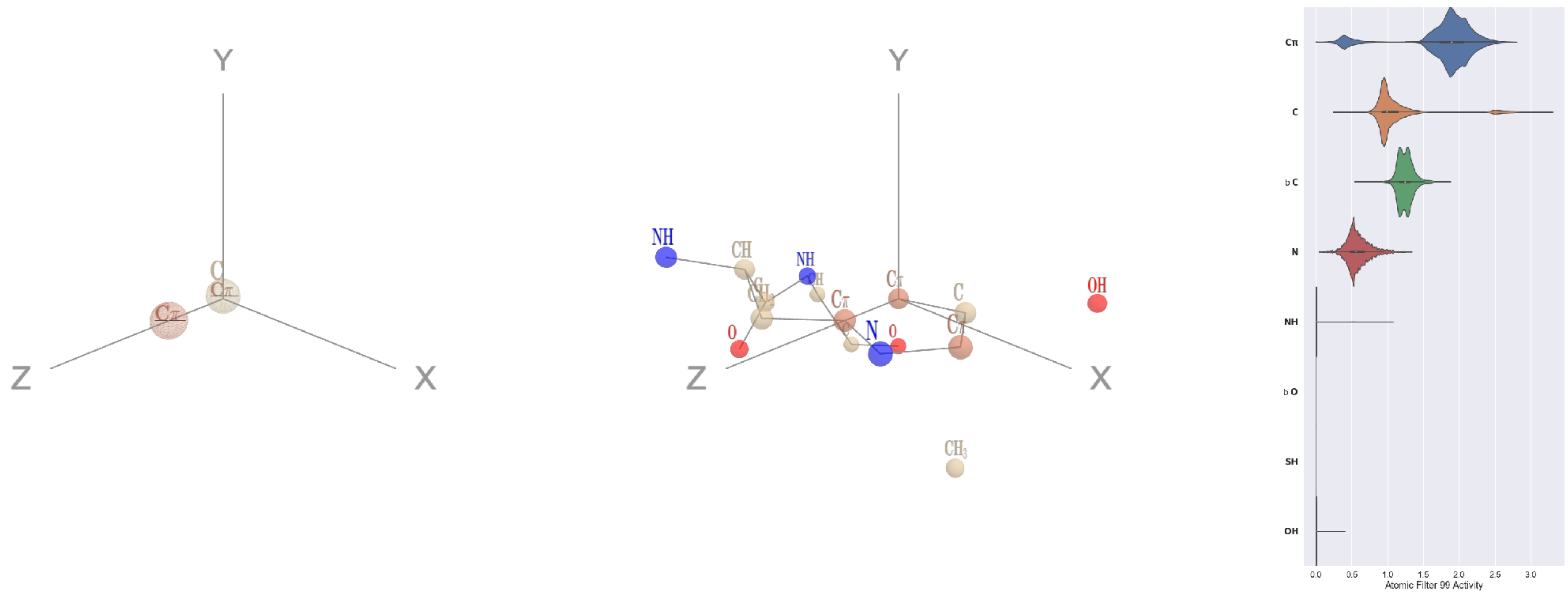


Filter 99
$R_{\text{pred}}^{\max} = 0.04$
$R_{\text{pred}}^{\text{mean}} = 0.04$
Percent Active = 25
Pooling attention 0.03
Pooling feature 1.34
Top activators
3ax7:A:665 H CD2
4hg6:A:479 Y CE1
1c0i:A:1338 Y CE1
4ot9:A:513 H CD2
4qiw:A:605 Y CE2
1eg1:A:356 H CD2
2wgq:B:568 Y CE1
1e1c:A:266 Y CE2
1r7r:A:173 Y CE1
3nh5:A:291 H CD2

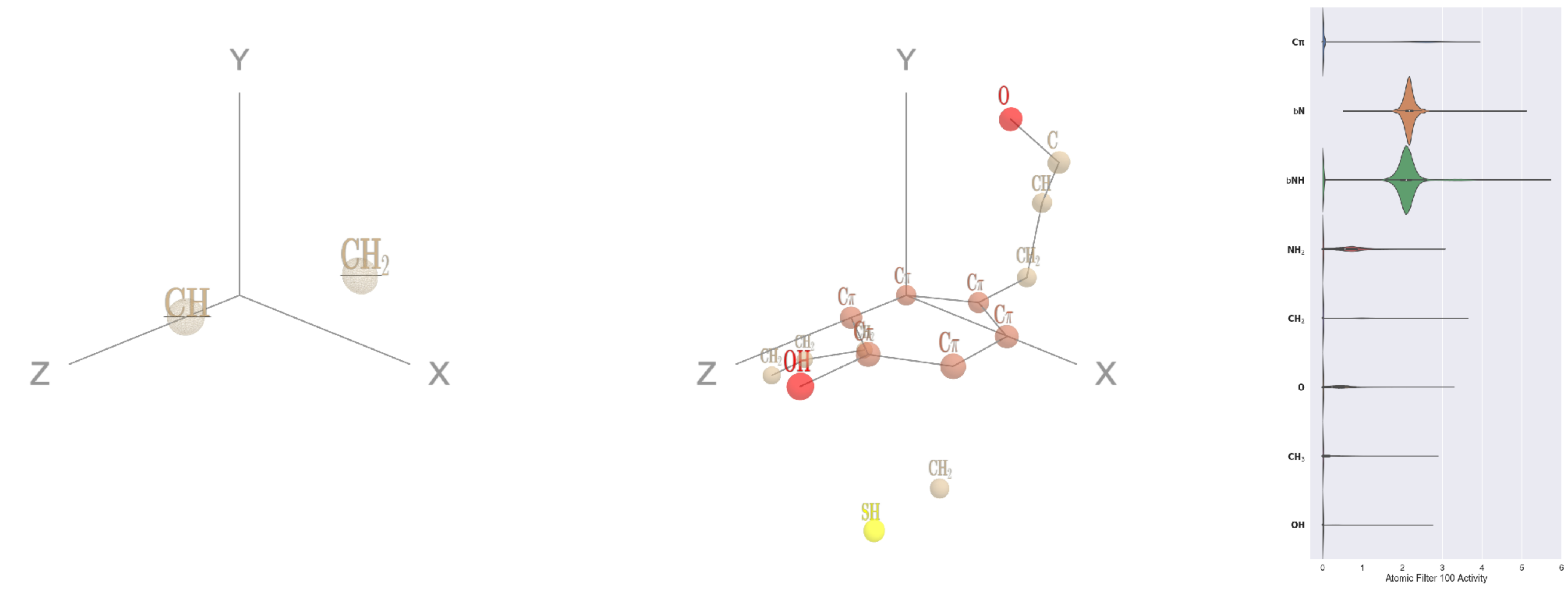


Filter 100
$R_{\text{pred}}^{\text{max}} = 0.05$
$R_{\text{pred}}^{\text{mean}} = 0.01$
Percent Active = 24
Pooling attention 0.27
Pooling feature 0.15
Top activators
1h2i:A:171 Y CD1
1lnl:A:106 F CD2
2np9:A:258 Y CD1
1bkj:A:231 Y CD1
1sl0:A:139 Y CD1
1bkj:A:199 Y CD1
1bkj:A:192 Y CD1
2dyo:A:264 Y CD1
1iwa:A:29 Y CD1
1a0j:A:29 Y CD1

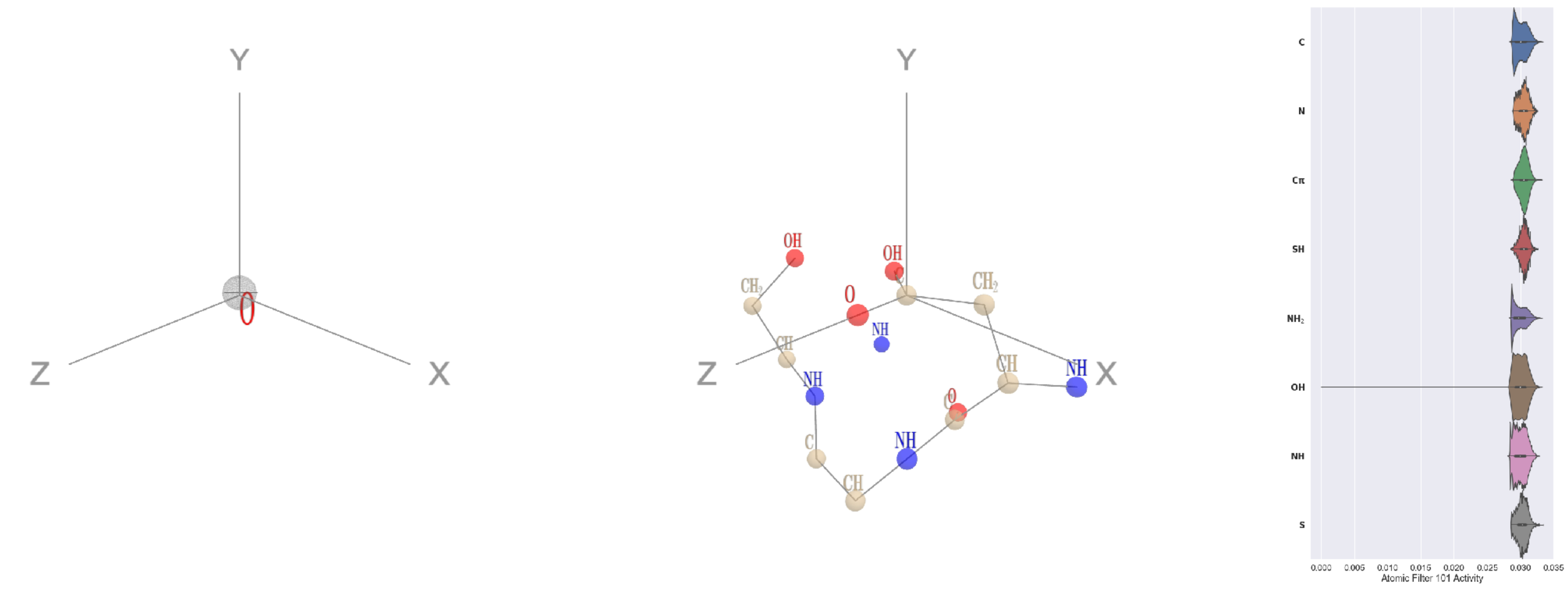


Filter 101
$R_{\text{pred}}^{\text{max}} = -0.04$
$R_{\text{pred}}^{\text{mean}} = 0.01$
Percent Active = 85
Pooling attention 0.00
Pooling feature 0.00
Top activators
1b8c:A:53 D CG
3ucq:A:111 R CZ
4g1m:B:143 R CZ
1r7r:A:319 E CD
1usv:B:119 E CD
2x98:A:77 E CD
1bgv:A:353 E CD
1qvr:A:576 R CZ
1efu:B:188 E CD
3bf0:C:501 E CD

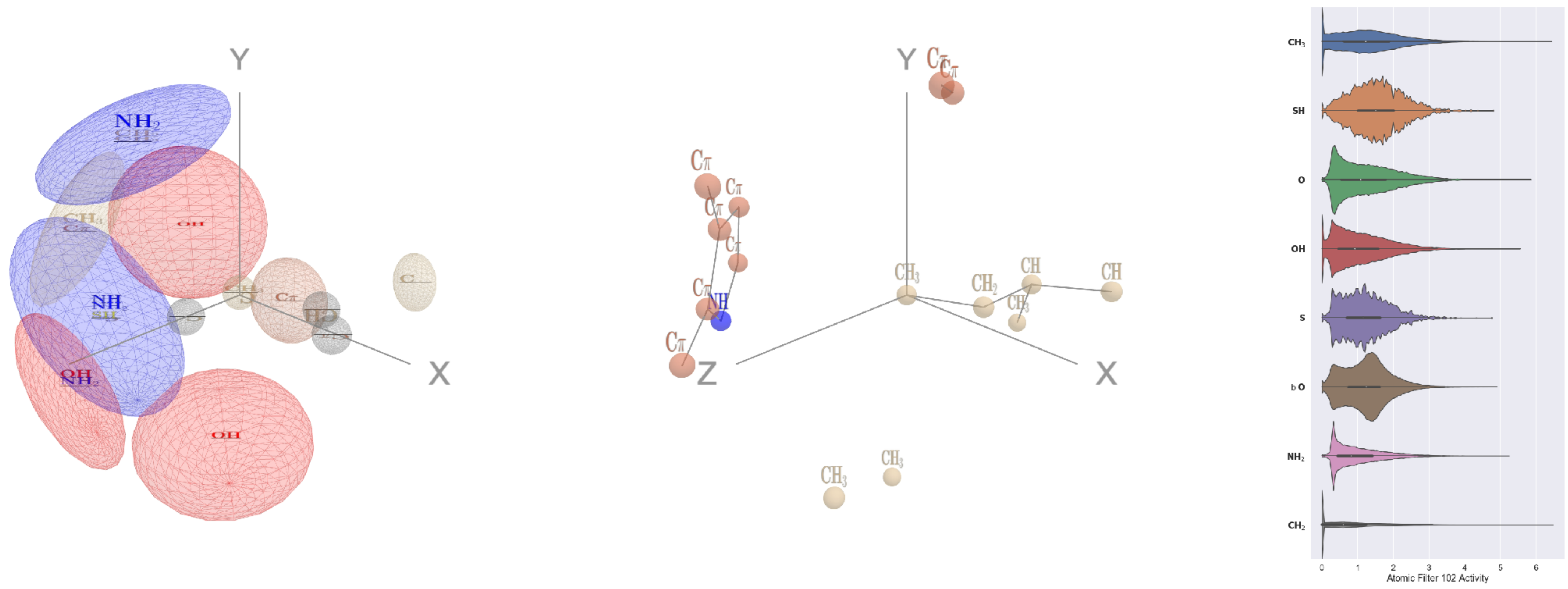


Filter 102
$R_{\text{pred}}^{\text{max}} = -0.10$
$R_{\text{pred}}^{\text{mean}} = -0.21$
Percent Active = 78
Pooling attention 0.52
Pooling feature 1.02
Top activators
1v8f:A:258 I CD1
4g59:C:199 M CE
3hot:A:203 I CD1
3ucq:A:126 M CE
4c2m:A:238 M CE
1mu2:A:179 I CD1
2x3l:B:277 I CD1
4rcn:B:154 M CE
2pmz:D:109 I CD1
2o3o:A:109 I CD1

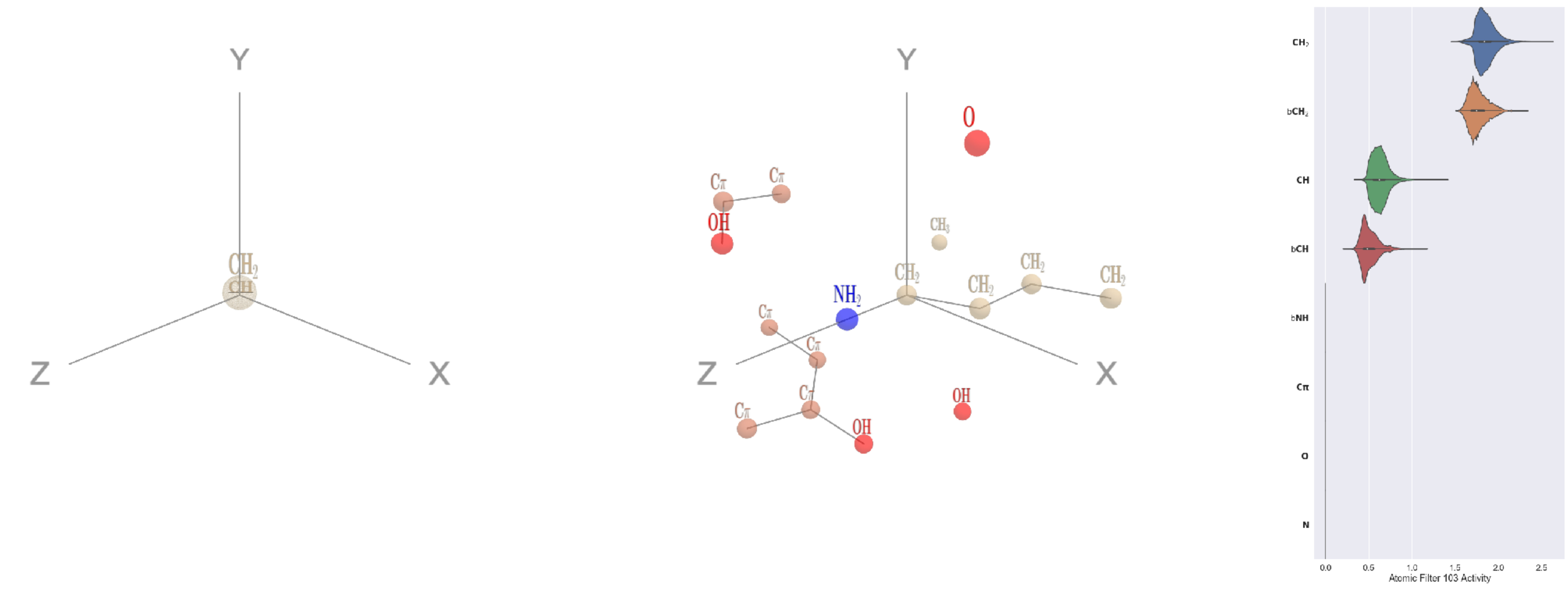


Filter 103
$R_{\text{pred}}^{\text{max}} = 0.04$
$R_{\text{pred}}^{\text{mean}} = 0.01$
Percent Active = 32
Pooling attention 0.16
Pooling feature 0.70
Top activators
1p35:A:263 K CE
3l0o:A:177 R CG
4kzx:L:152 K CE
3b8e:A:694 I CG1
4gu0:A:549 S CB
4v4k:M:167 K CB
1w36:B:555 S CB
3n6r:G:294 R CD
4pk7:A:563 I CG1
3qsq:A:433 R CD

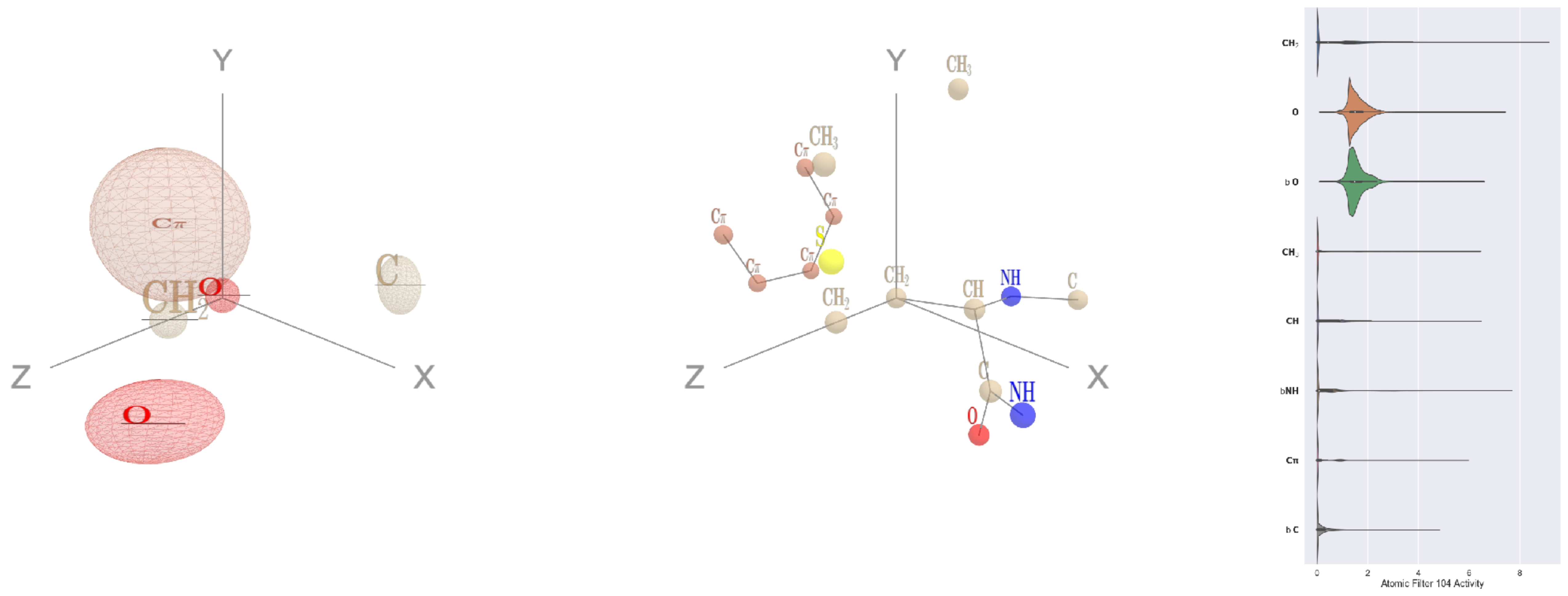

Filter 104
$R_{\text{pred}}^{\text{max}} = -0.04$
$R_{\text{pred}}^{\text{mean}} = -0.08$
Percent Active = 50
Pooling attention 0.36
Pooling feature 1.20
Top activators
1he8:A:514 M CB
1o7g:A:196 K CG
3b8m:A:100 R CG
1umg:A:265 M CB
2wsw:A:146 P CB
1h4s:A:238 R CG
2fml:A:233 R CG
3c24:A:267 R CG
2f8x:C:55 P CB
1ogl:A:60 K CG

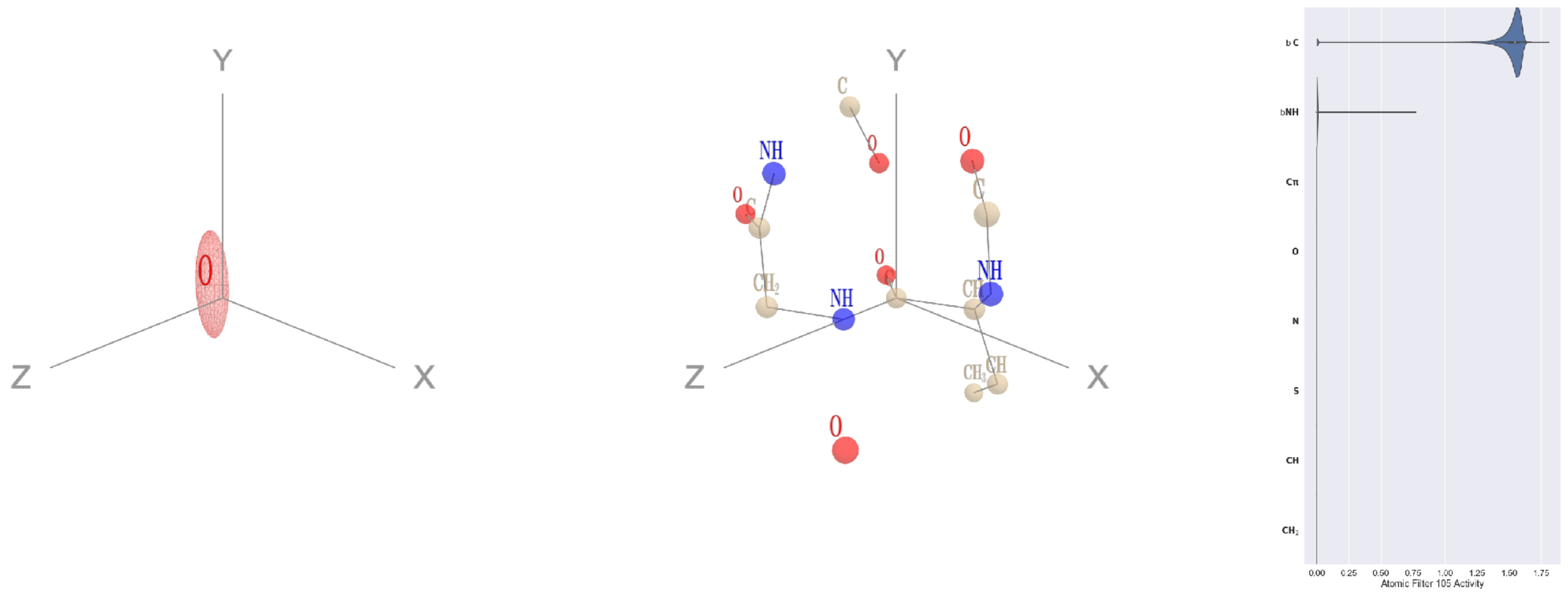

Filter 105
$R_{\text{pred}}^{\text{max}} = -0.02$
$R_{\text{pred}}^{\text{mean}} = -0.08$
Percent Active = 13
Pooling attention 0.02
Pooling feature 0.01
Top activators
1e5d:A:213 V C
4buj:B:520 E C
4oph:A:183 G C
2fml:A:188 E C
1u5e:A:189 I C
1u5e:A:159 Q C
3kyh:C:389 N C
1u5e:A:164 F C
3drw:A:218 A C
1f4a:A:350 L C

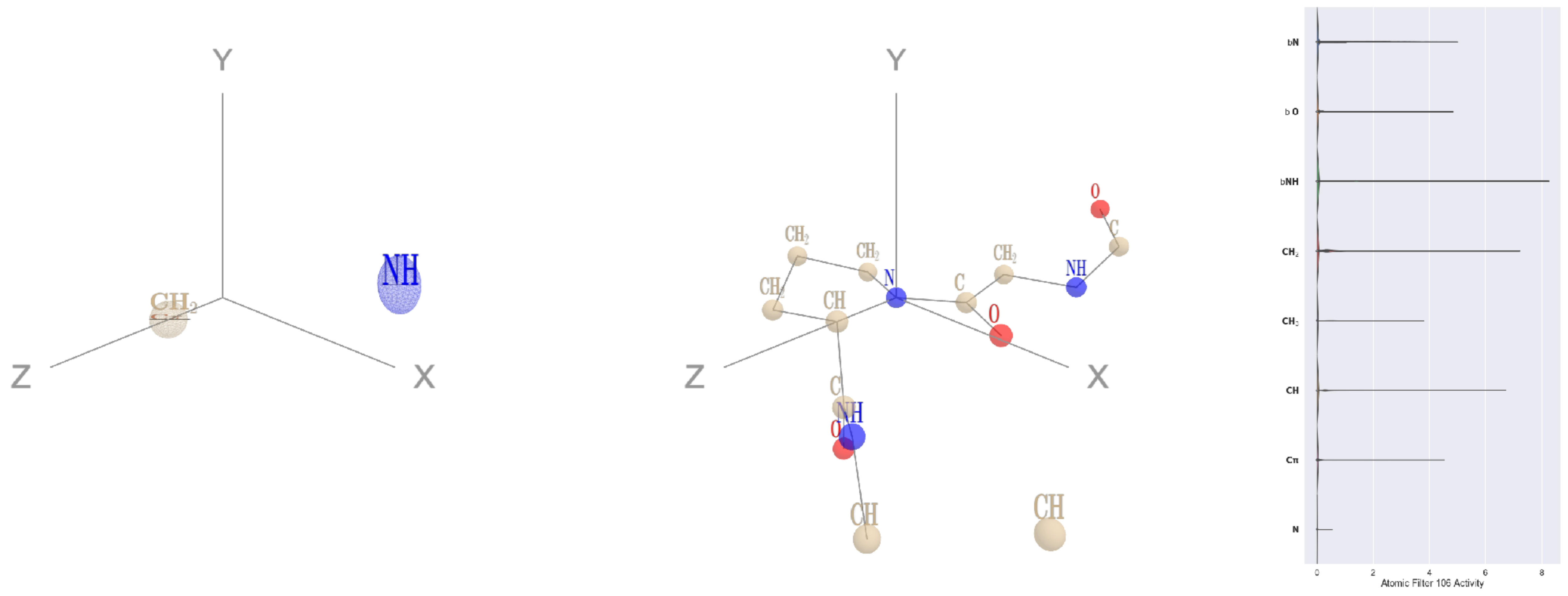

Filter 106
$R_{\text{pred}}^{\text{max}} = 0.01$
$R_{\text{pred}}^{\text{mean}} = -0.00$
Percent Active = 18
Pooling attention 0.16
Pooling feature 0.22
Top activators
2hh1:H:78 P N
4ny2:A:235 P N
1ivu:A:228 P N
3do6:A:469 P N
1ewy:A:73 P N
1iri:A:484 P N
2wk4:B:428 P N
3c24:A:157 P N
2f8x:C:332 P N
1ei6:A:167 P N

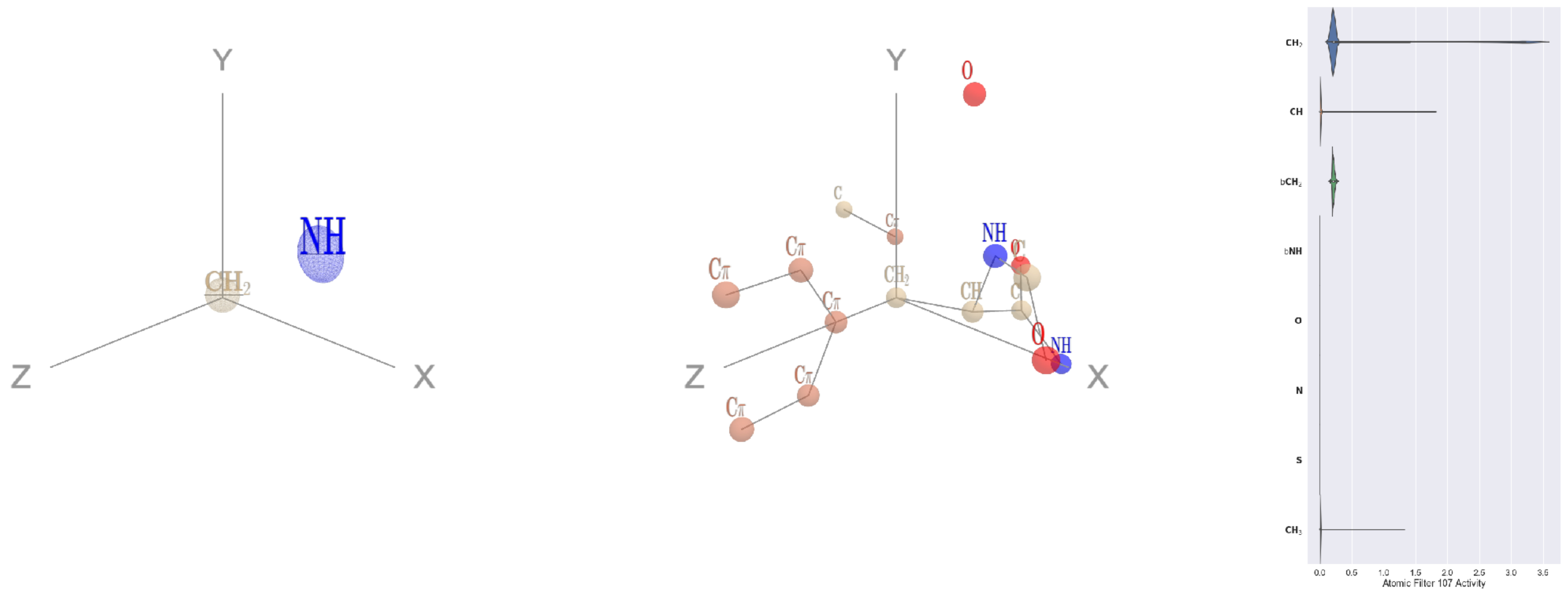

Filter 107
$R_{\text{pred}}^{\text{max}} = 0.04$
$R_{\text{pred}}^{\text{mean}} = 0.03$
Percent Active = 18
Pooling attention 0.14
Pooling feature 0.03
Top activators
2x3l:B:318 Y CB
3bof:A:22 Y CB
1a0j:A:151 Y CB
4d0l:A:661 N CB
1bxr:A:493 K CB
2nyg:A:235 H CB
1e5d:A:196 N CB
1qha:A:890 F CB
2wsc:B:671 W CB
4c2m:A:675 S CB

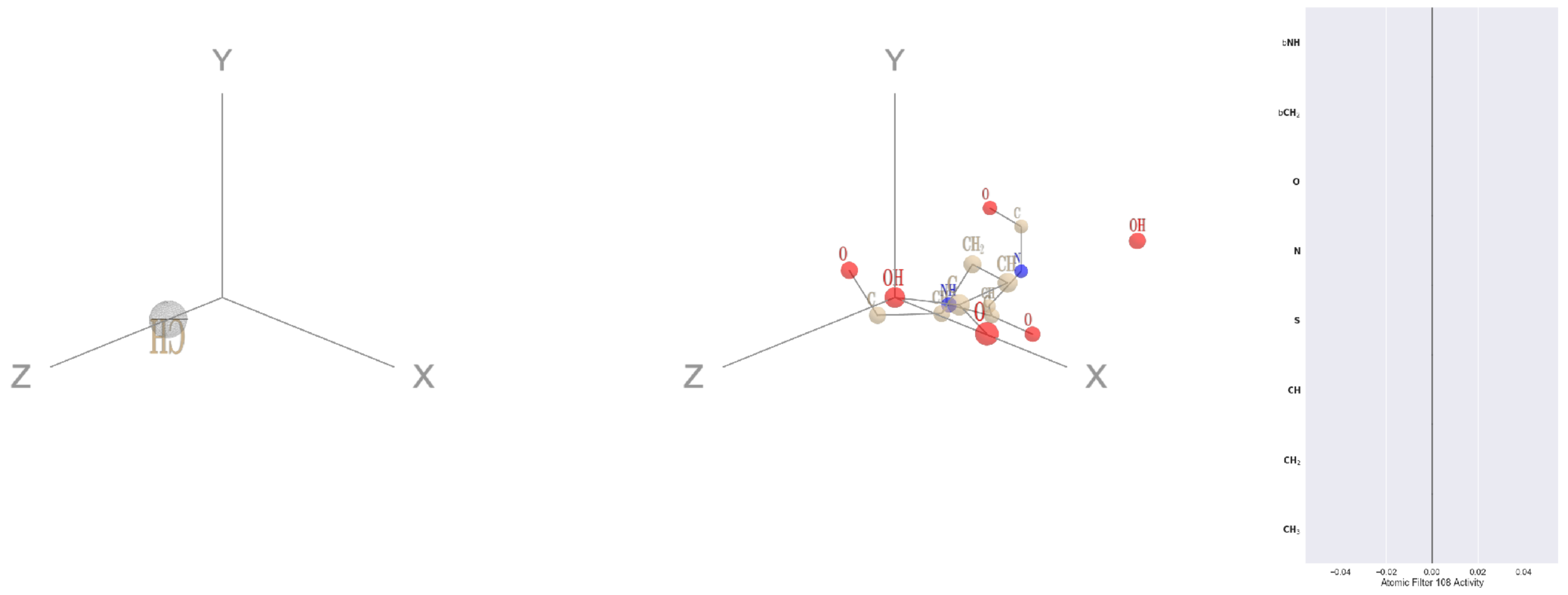

Filter 108
$R_{\text{pred}}^{\max} = nan$
$R_{\text{pred}}^{\text{mean}} = nan$
Percent Active = 0
Pooling attention 0.00
Pooling feature 0.00
Top activators
5wy5:A:246 E OE2
2b0o:F:573 F N
2b0o:F:574 G O
2b0o:F:574 G C
2b0o:F:574 G CA
2b0o:F:574 G N
2b0o:F:573 F CZ
2b0o:F:573 F CE2
2b0o:F:573 F CE1
2b0o:F:573 F CD2

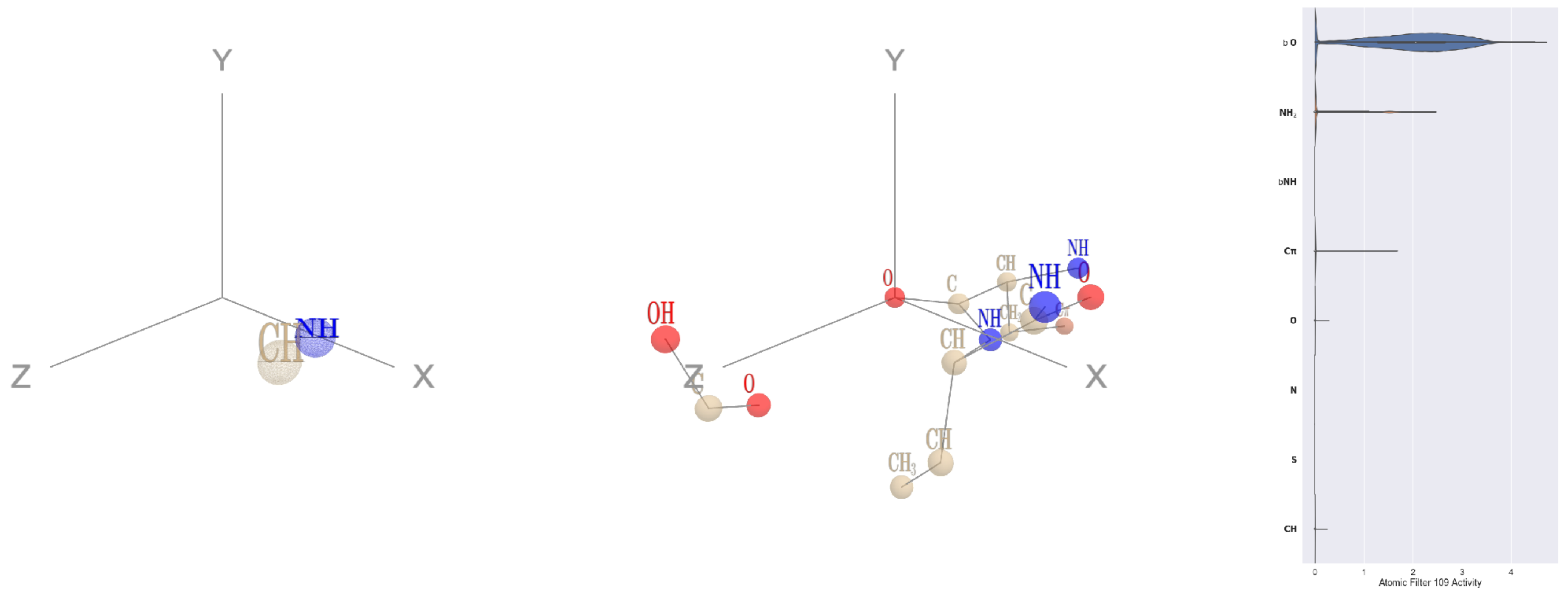

Filter 109
$R_{\text{pred}}^{\max} = -0.04$
$R_{\text{pred}}^{\text{mean}} = -0.07$
Percent Active = 12
Pooling attention 0.14
Pooling feature 0.25
Top activators
1rq0:A:212 Y O
3dlb:A:354 A O
1e1c:A:163 V O
1f4a:A:748 C O
1e1c:A:16 P O
1f4a:A:5 D O
1bxr:A:797 P O
1f4a:A:257 T O
3b8e:A:183 L O
1f4a:A:916 D O

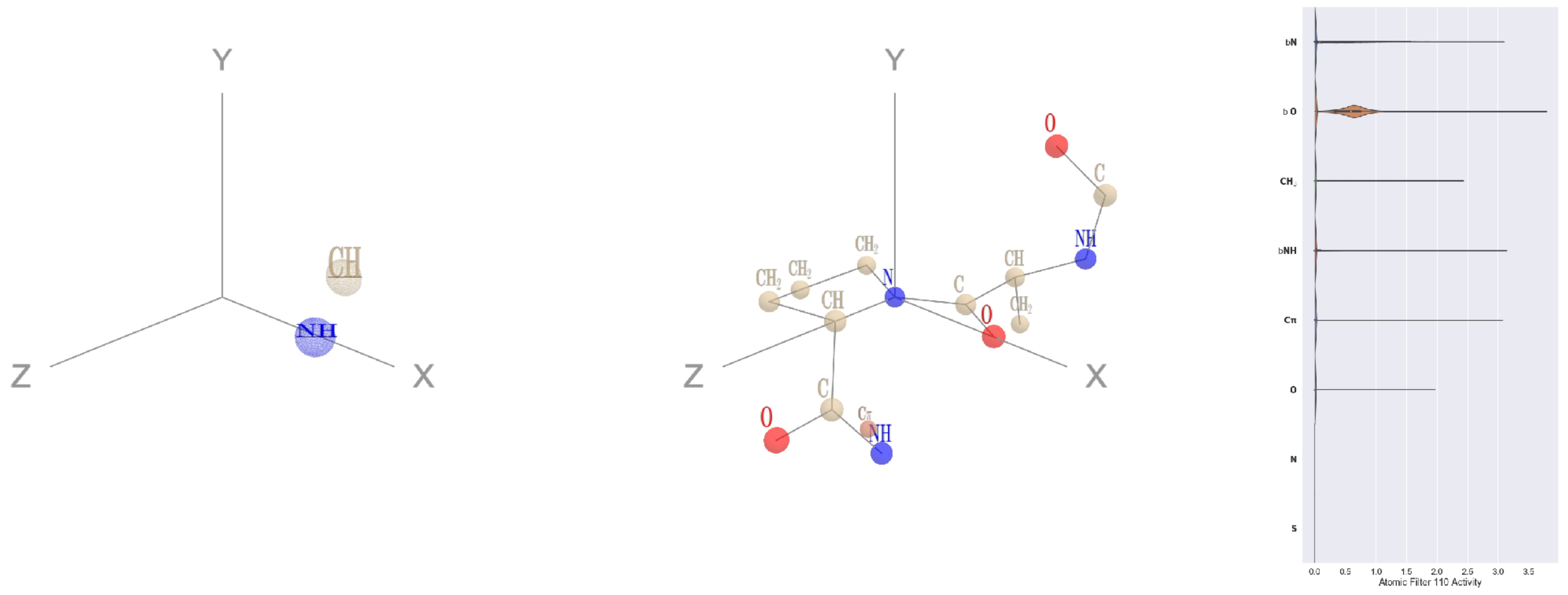

Filter 110
$R_{\text{pred}}^{\max} = -0.01$
$R_{\text{pred}}^{\text{mean}} = -0.03$
Percent Active = 17
Pooling attention 0.06
Pooling feature 0.02
Top activators
4cht:B:126 P N
4pfp:A:10 P N
4geq:A:190 P N
4nnq:B:243 P N
3t3o:A:99 P N
2fa0:A:316 P N
1pkp:A:82 P N
1a0j:A:124 P N
4pfp:A:774 P N
3ucq:A:196 P N

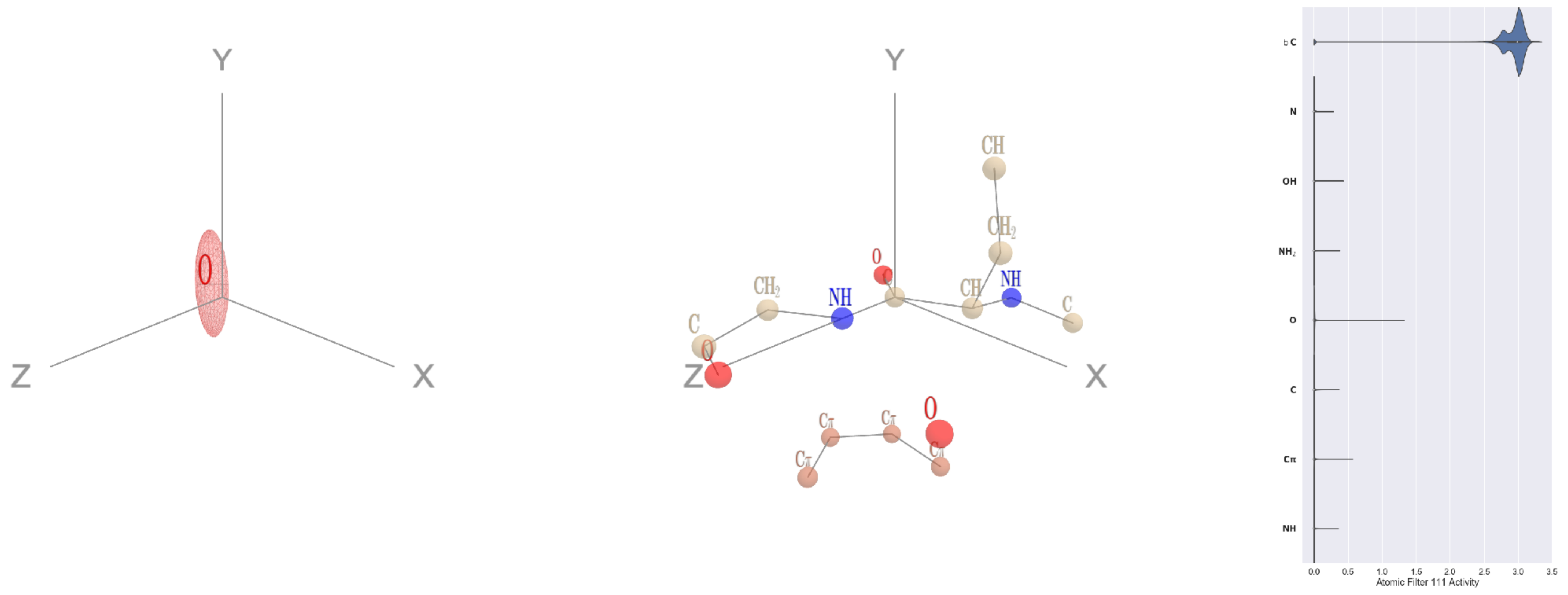

Filter 111
$R_{\text{pred}}^{\max} = -0.02$
$R_{\text{pred}}^{\text{mean}} = -0.08$
Percent Active = 29
Pooling attention 0.35
Pooling feature 0.09
Top activators
1mu2:A:315 E C
1ghq:C:40 G C
2zbk:B:168 H C
2d32:A:191 G C
1sl0:A:20 C C
1fx0:B:194 G C
1e5d:A:213 V C
4u7u:M:402 G C
1d0n:A:201 C C
2cge:A:300 W C

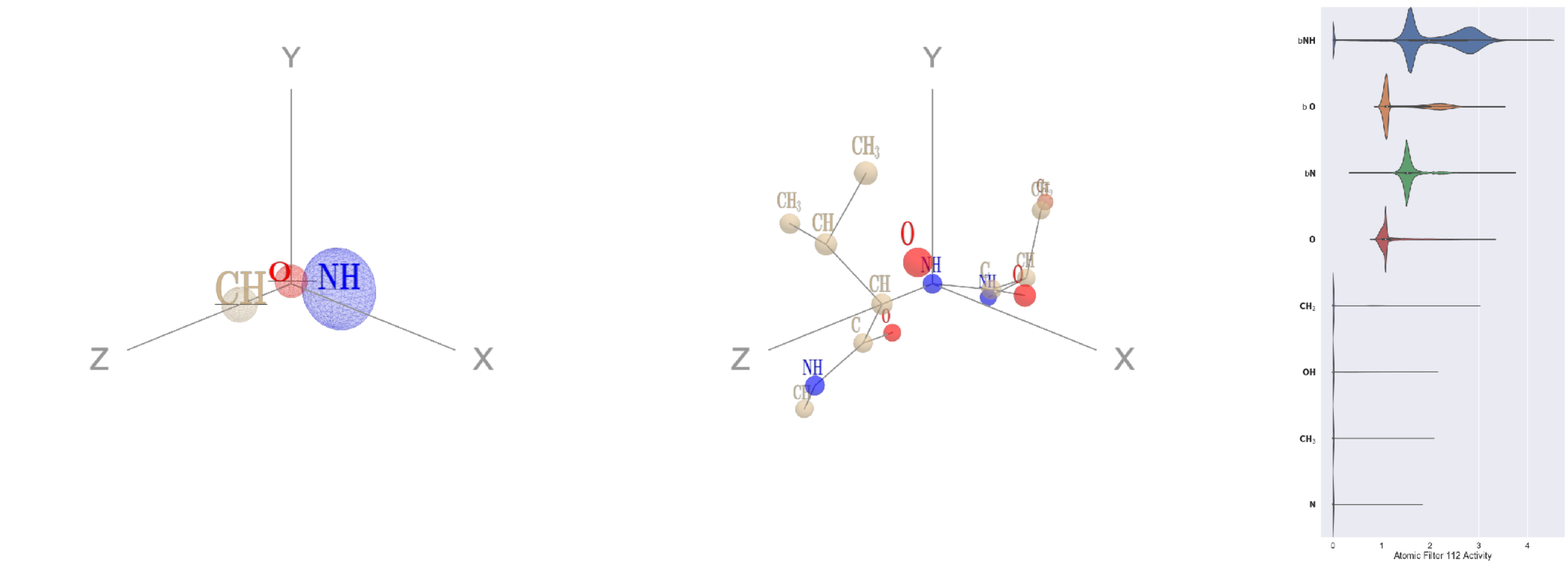


Filter 112
$R_{\text{pred}}^{\text{max}} = 0.02$
$R_{\text{pred}}^{\text{mean}} = -0.06$
Percent Active = 33
Pooling attention 0.14
Pooling feature 0.11
Top activators
1rq0:A:213 V N
3a5c:G:196 R N
1qha:A:720 D N
2wsc:B:629 S N
1w36:B:909 L N
1fx0:B:313 I N
4o93:A:61 E N
3b8e:A:456 C N
2wsc:B:337 A N
1u5e:A:44 K N

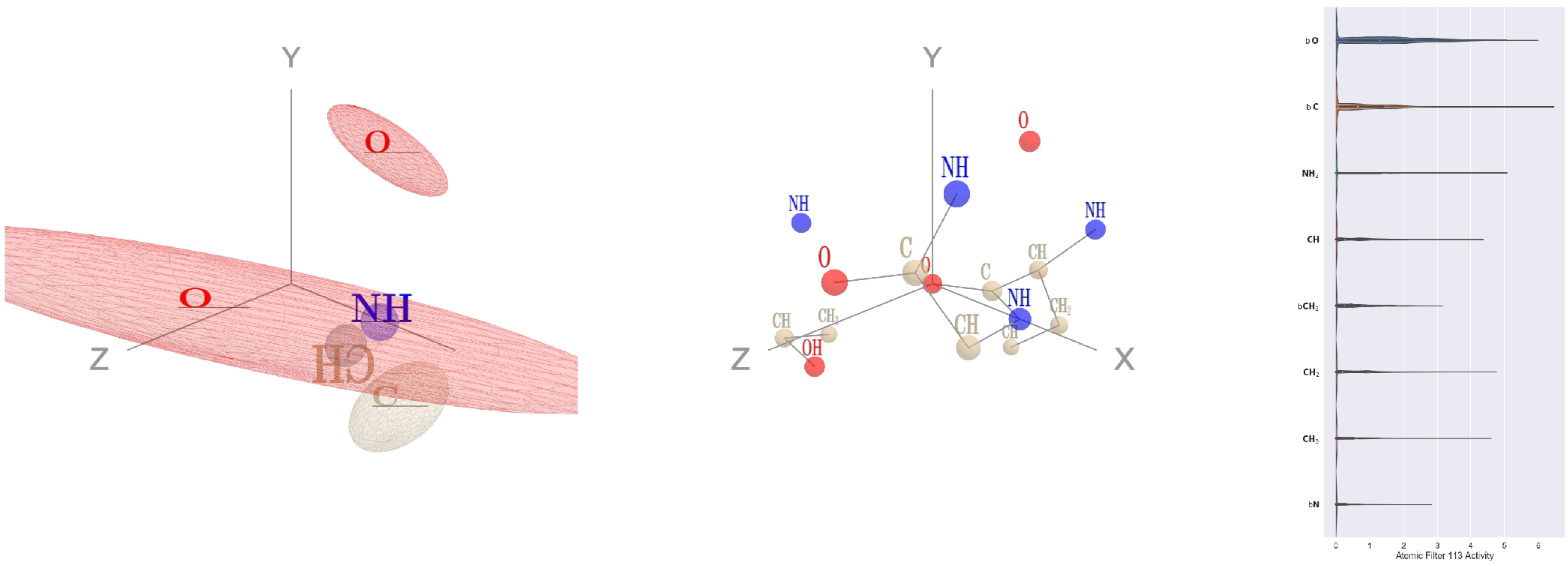


Filter 113
$R_{\text{pred}}^{\text{max}} = 0.00$
$R_{\text{pred}}^{\text{mean}} = -0.06$
Percent Active = 48
Pooling attention 0.27
Pooling feature 0.63
Top activators
4l8n:A:188 E O
1qha:A:314 A O
3vie:B:134 I O
1y88:A:100 A O
4u0u:A:430 L O
1ppt:A:28 L O
1e1c:A:612 R O
1e5d:A:356 A O
3fbv:D:858 P O
1b8a:A:390 E O

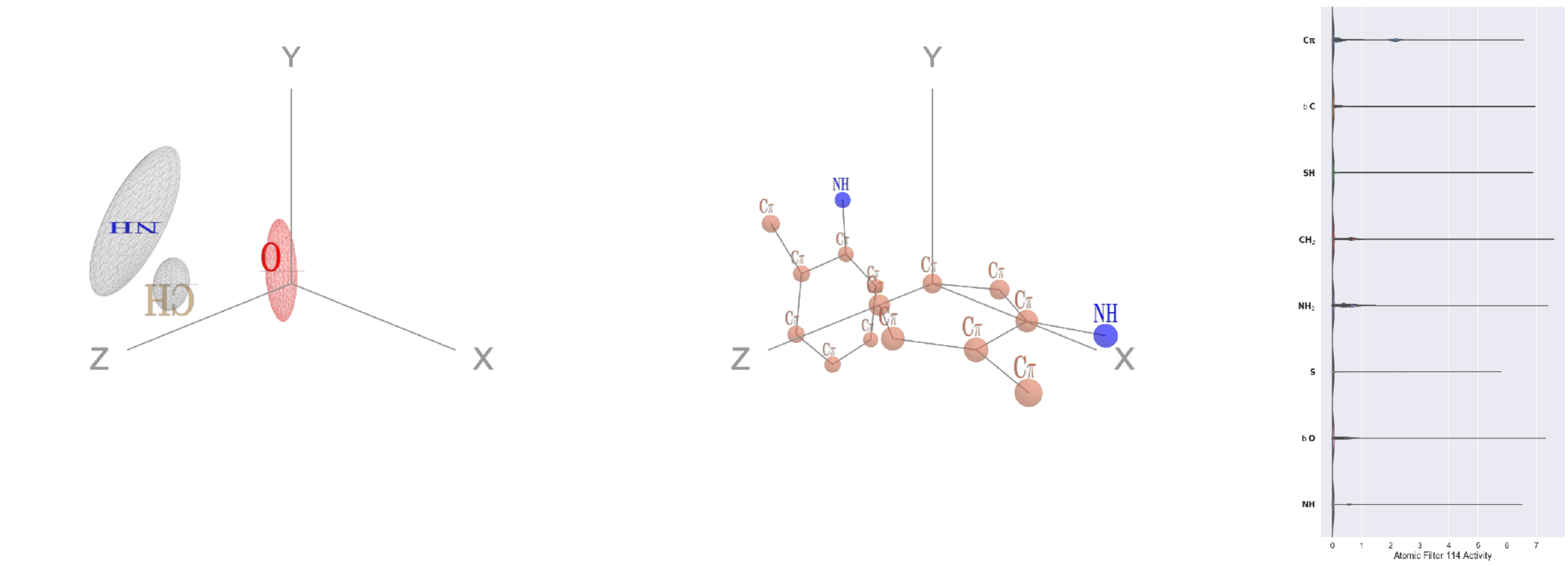


Filter 114
$R_{\text{pred}}^{\text{max}} = 0.02$
$R_{\text{pred}}^{\text{mean}} = 0.00$
Percent Active = 27
Pooling attention 0.20
Pooling feature 1.44
Top activators
3a1m:A:76 W CH2
1lnl:A:312 F CE1
1e1c:A:286 F CE1
1k72:B:498 F CZ
1bcp:A:50 F CE1
1oxw:C:2278 W CZ3
4v4k:M:15 F CD2
3jsb:A:154 W CZ2
2prv:A:132 F CE1
2z67:A:122 F CE1

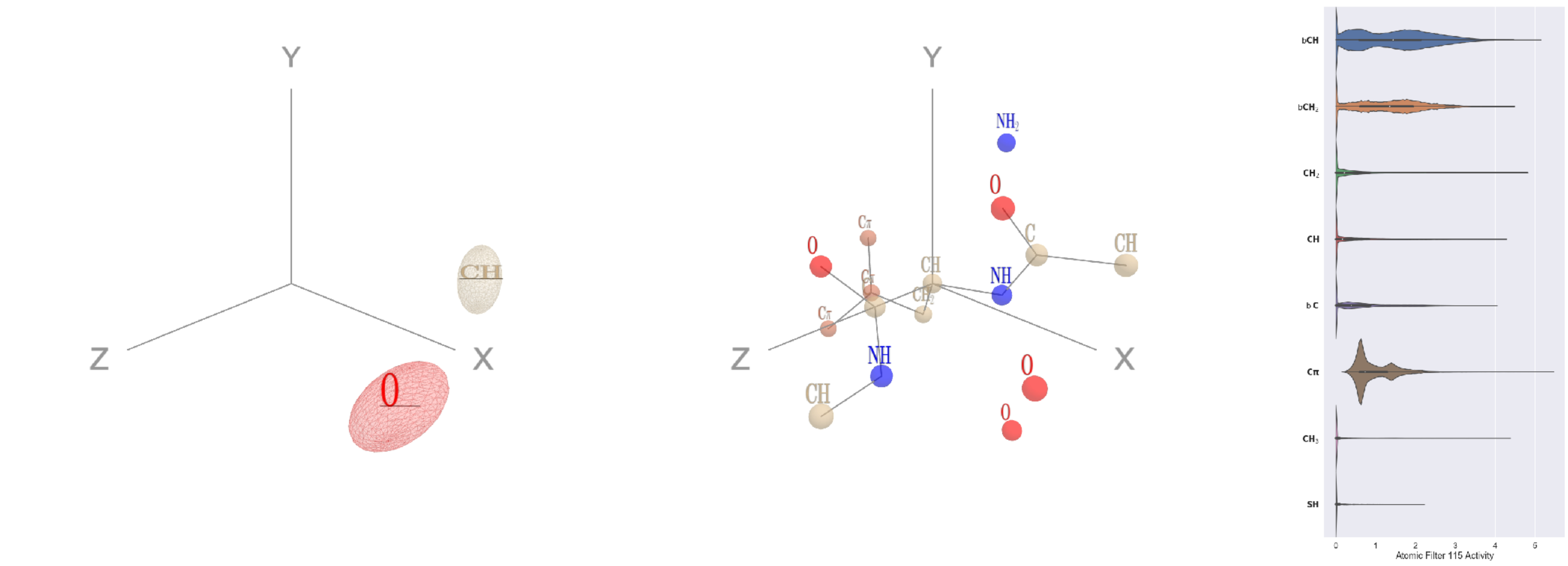


Filter 115
$R_{\text{pred}}^{\text{max}} = -0.04$
$R_{\text{pred}}^{\text{mean}} = -0.08$
Percent Active = 48
Pooling attention 0.22
Pooling feature 0.39
Top activators
2cge:A:664 F CA
4v8p:BC:369 H CA
3r9j:C:51 C CA
2iwh:B:420 F CA
4ani:A:89 F CA
1d0n:A:318 W CA
4oph:A:14 F CA
1jqn:A:578 F CA
4hh2:D:341 Y CA
2wsc:B:168 F CA

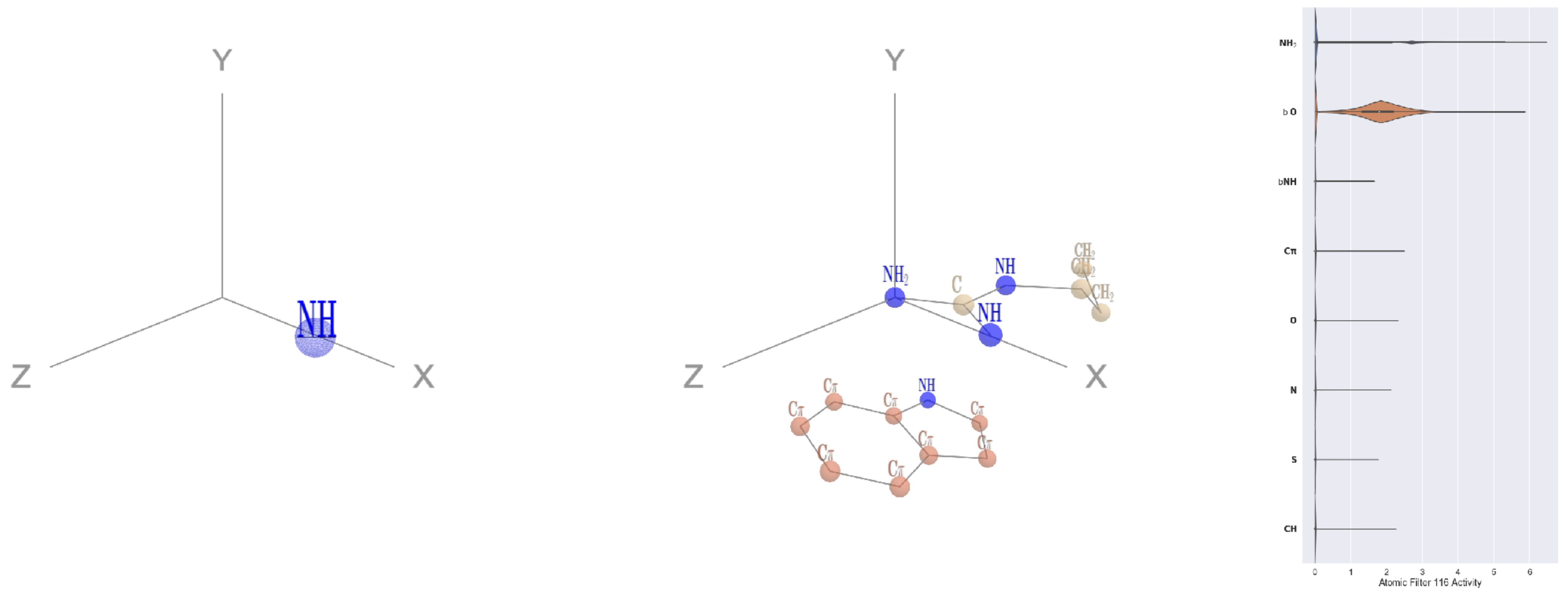


Filter 116
$R_{\text{pred}}^{\text{max}} = 0.01$
$R_{\text{pred}}^{\text{mean}} = -0.03$
Percent Active = 14
Pooling attention 0.11
Pooling feature 0.03
Top activators
1vqv:B:187 R NH2
1yqg:A:135 R NH2
1qvr:A:727 R NH2
1vl4:A:330 R NH2
4pht:A:422 R NH2
3v4l:A:559 R NH2
2eo4:A:9 R NH2
4ehi:B:339 R NH2
4u4c:A:747 R NH2
4gfh:A:83 R NH2

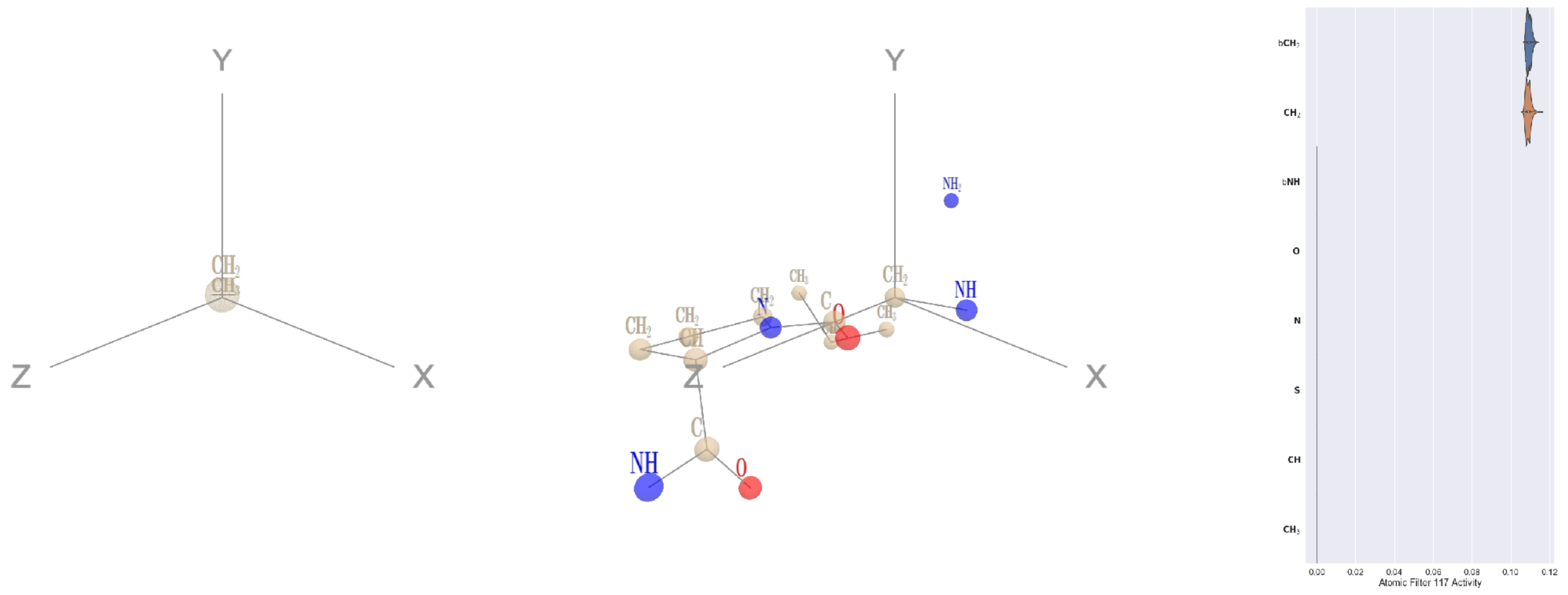


Filter 117
$R_{\text{pred}}^{\text{max}} = 0.04$
$R_{\text{pred}}^{\text{mean}} = 0.02$
Percent Active = 16
Pooling attention 0.00
Pooling feature 0.00
Top activators
1ppt:A:1 G CA
1umg:A:306 G CA
1fx0:B:28 G CA
4whj:A:125 G CA
3bsd:A:216 G CA
3ucq:A:-4 G CA
3c57:A:149 G CA
3dlb:A:388 G CA
4c8q:H:474 G CA
4v4k:M:427 G CA

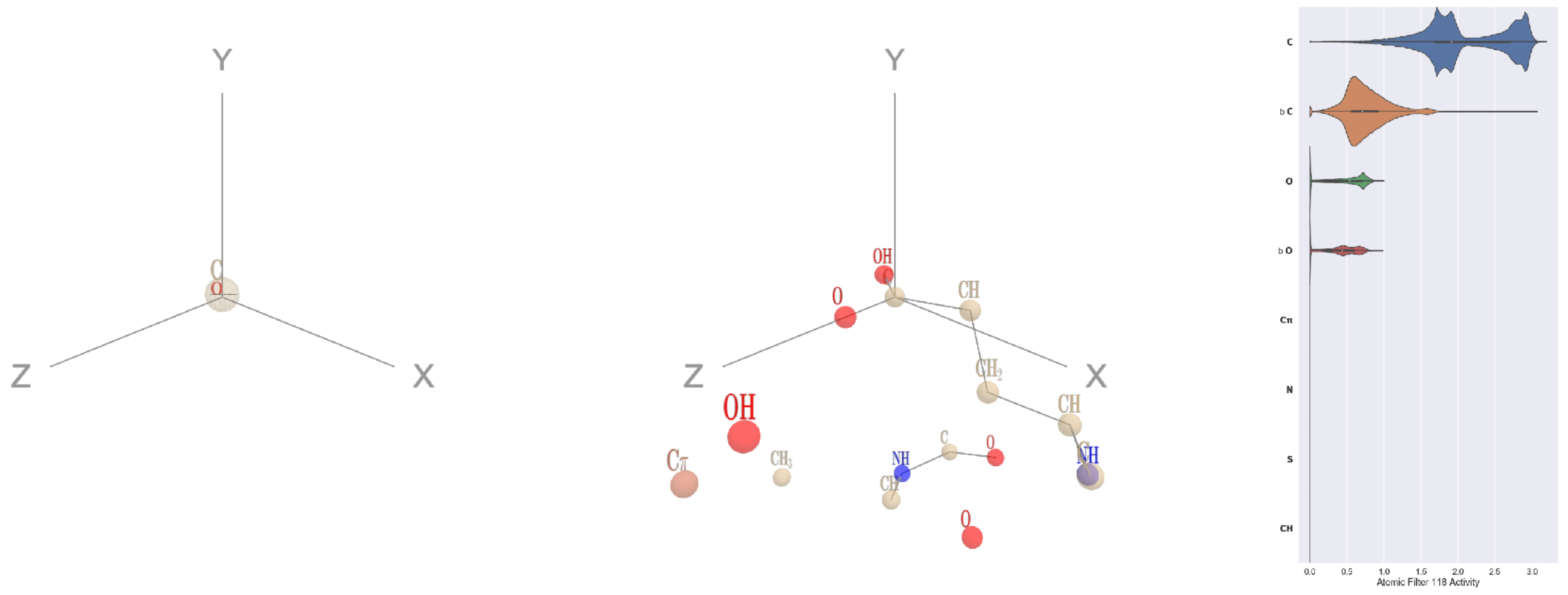


Filter 118
$R_{\text{pred}}^{\text{max}} = 0.05$
$R_{\text{pred}}^{\text{mean}} = 0.05$
Percent Active = 29
Pooling attention 0.19
Pooling feature 0.84
Top activators
1am4:A:95 E CD
1qha:A:731 E CD
1bxr:A:683 E CD
1dtn:A:333 E CD
1bxr:A:478 E CD
3t3o:A:438 E CD
1bxr:A:591 E CD
4f52:E:108 E CD
3hfd:A:267 E CD
1bgv:A:12 E CD

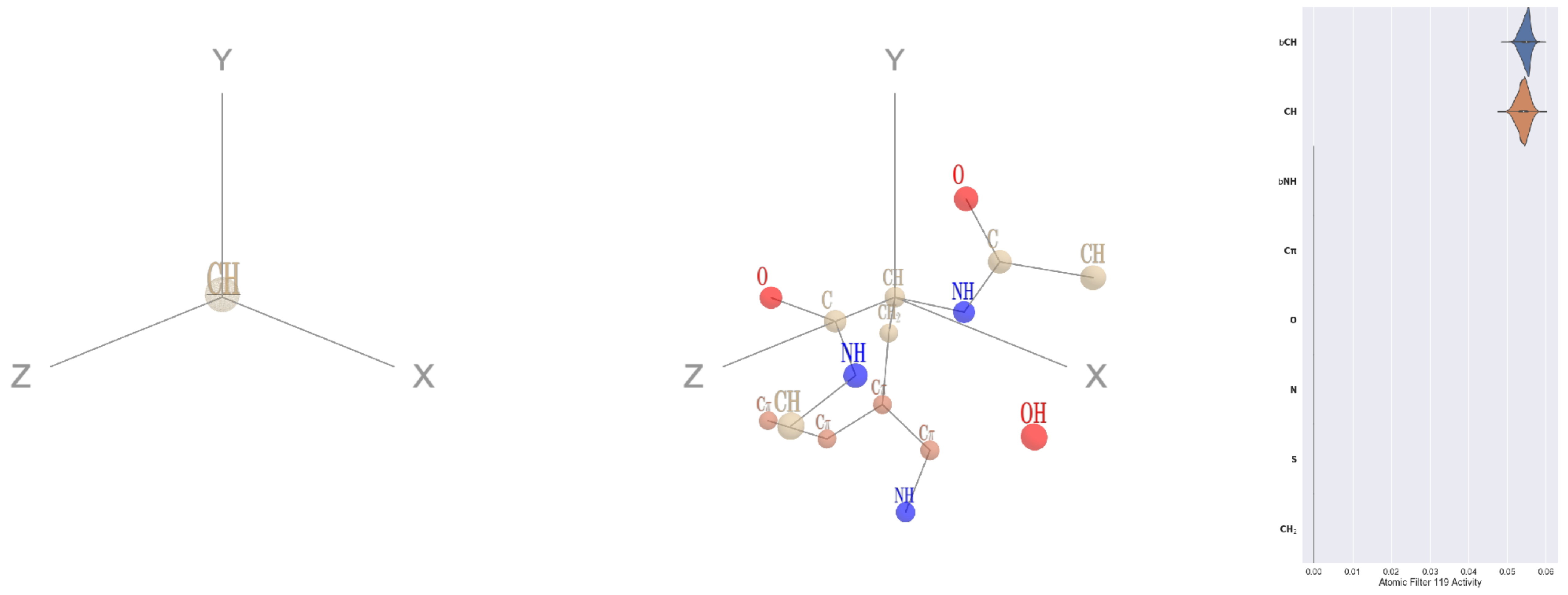


Filter 119
$R_{\text{pred}}^{\text{max}} = 0.03$
$R_{\text{pred}}^{\text{mean}} = -0.04$
Percent Active = 16
Pooling attention 0.00
Pooling feature 0.00
Top activators
2cge:A:300 W CA
1bh3:A:121 Y CA
1c0i:A:1107 W CA
4qiw:A:659 W CA
1w99:A:89 W CA
4rcn:B:903 H CA
1c3r:A:91 Y CA
3bma:C:144 Y CA
4rcn:B:837 W CA
4gu0:A:559 W CA

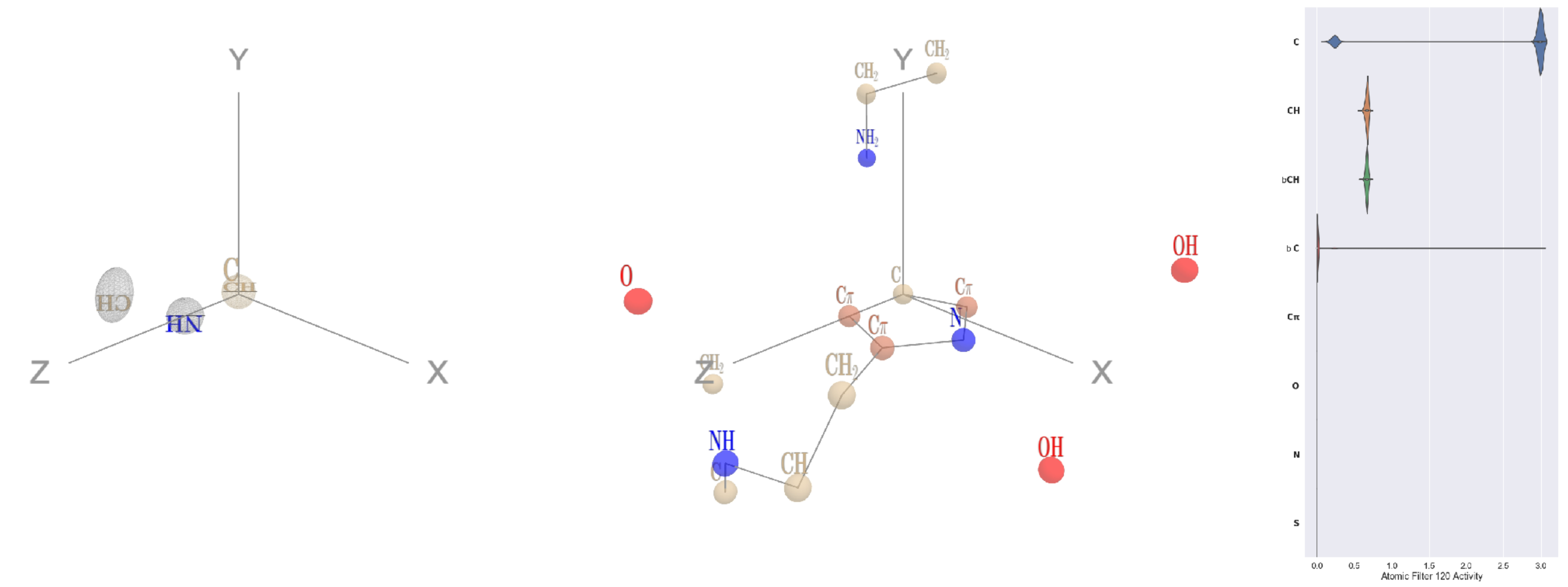

Filter 120
$R_{\mathrm{pred}}^{\mathrm{max}} = 0.03$
$R_{\mathrm{pred}}^{\mathrm{mean}} = 0.01$
Percent Active = 21
Pooling attention 0.10
Pooling feature 0.02
Top activators
4buj:B:1102 H NE2
1a71:A:234 E CD
4nm9:B:587 E CD
2wgq:B:651 E CD
1oqo:C:24 N CG
3i5d:B:237 D CG
2iwh:B:528 E CD
1u6z:B:356 H NE2
3hwp:A:117 E CD
1vbk:A:142 E CD

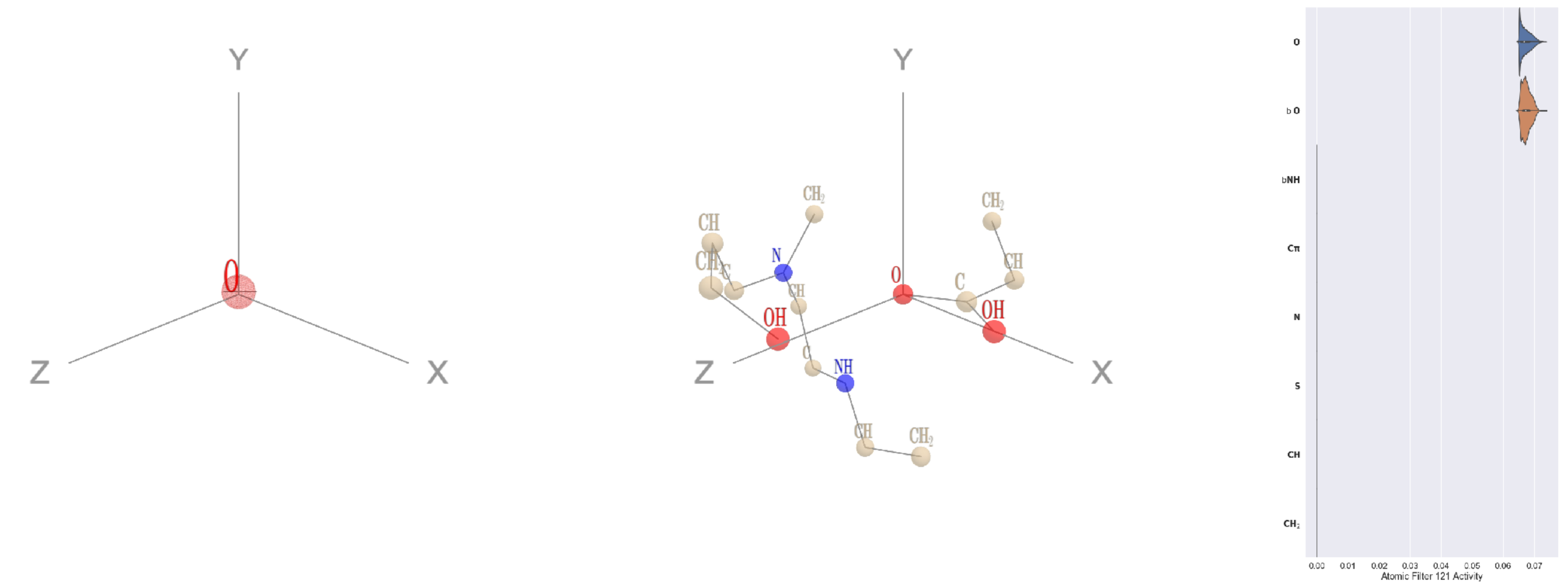

Filter 121
$R_{\mathrm{pred}}^{\mathrm{max}} = -0.14$
$R_{\mathrm{pred}}^{\mathrm{mean}} = -0.06$
Percent Active = 15
Pooling attention 0.00
Pooling feature 0.00
Top activators
4nm9:B:74 E OE1
4hg6:A:389 Q OE1
1mr1:D:297 Q OE1
1he8:A:814 E OE1
1a71:A:96 Q OE1
4mex:C:1114 E OE1
1e1c:A:223 E OE1
2f8x:C:328 E OE1
1qhd:A:189 Q OE1
2v6e:A:184 Q OE1

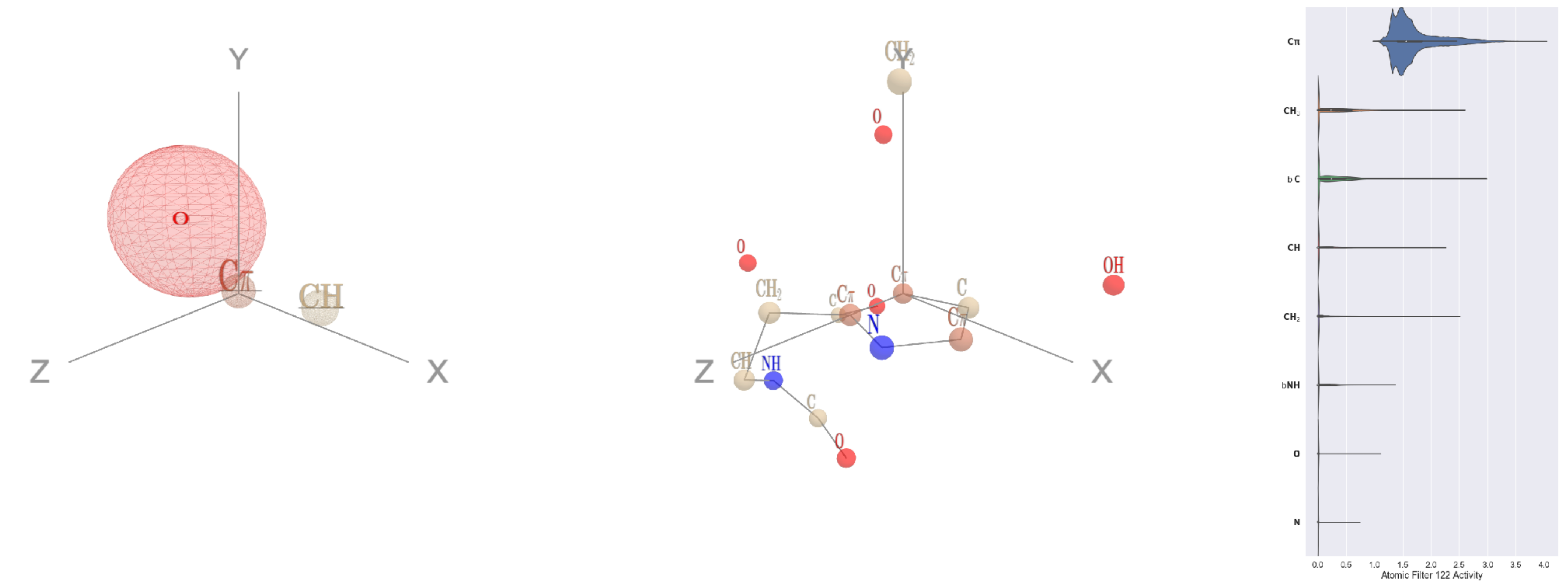

Filter 122
$R_{\mathrm{pred}}^{\mathrm{max}} = -0.02$
$R_{\mathrm{pred}}^{\mathrm{mean}} = -0.01$
Percent Active = 36
Pooling attention 0.37
Pooling feature 1.76
Top activators
5hu6:C:208 H CD2
1d0n:A:567 F CE1
3hot:A:293 H CD2
4cvw:A:404 H CD2
1qha:A:631 H CD2
1iri:A:389 H CD2
4gfh:A:123 H CD2
2cge:A:308 H CD2
3n99:G:170 H CD2
3n6r:G:455 W CZ3

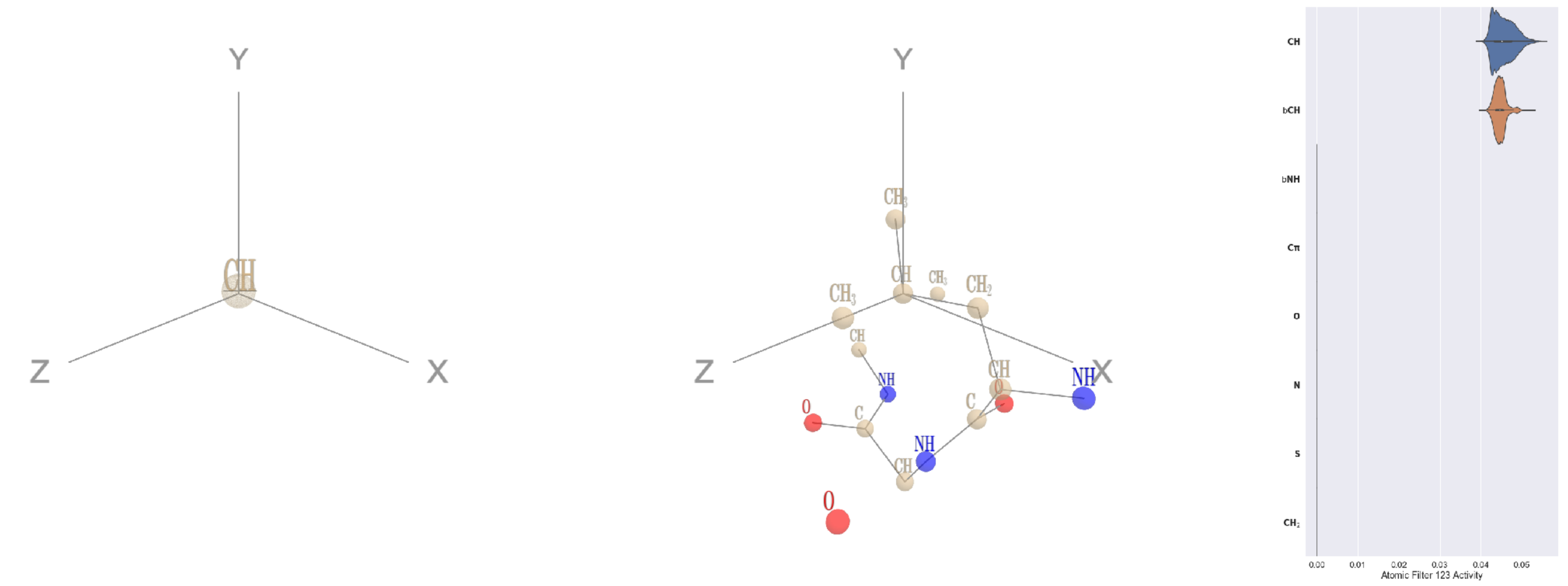

Filter 123
$R_{\mathrm{pred}}^{\mathrm{max}} = 0.02$
$R_{\mathrm{pred}}^{\mathrm{mean}} = -0.04$
Percent Active = 16
Pooling attention 0.00
Pooling feature 0.00
Top activators
1umg:A:171 L CG
4qiw:A:552 L CG
4uzi:A:357 L CG
2d73:A:258 L CG
2b0o:F:452 L CG
1w36:B:987 E CG
2fq1:A:34 L CG
1vio:A:65 L CG
2nyg:A:172 L CG
4rcn:B:709 L CG

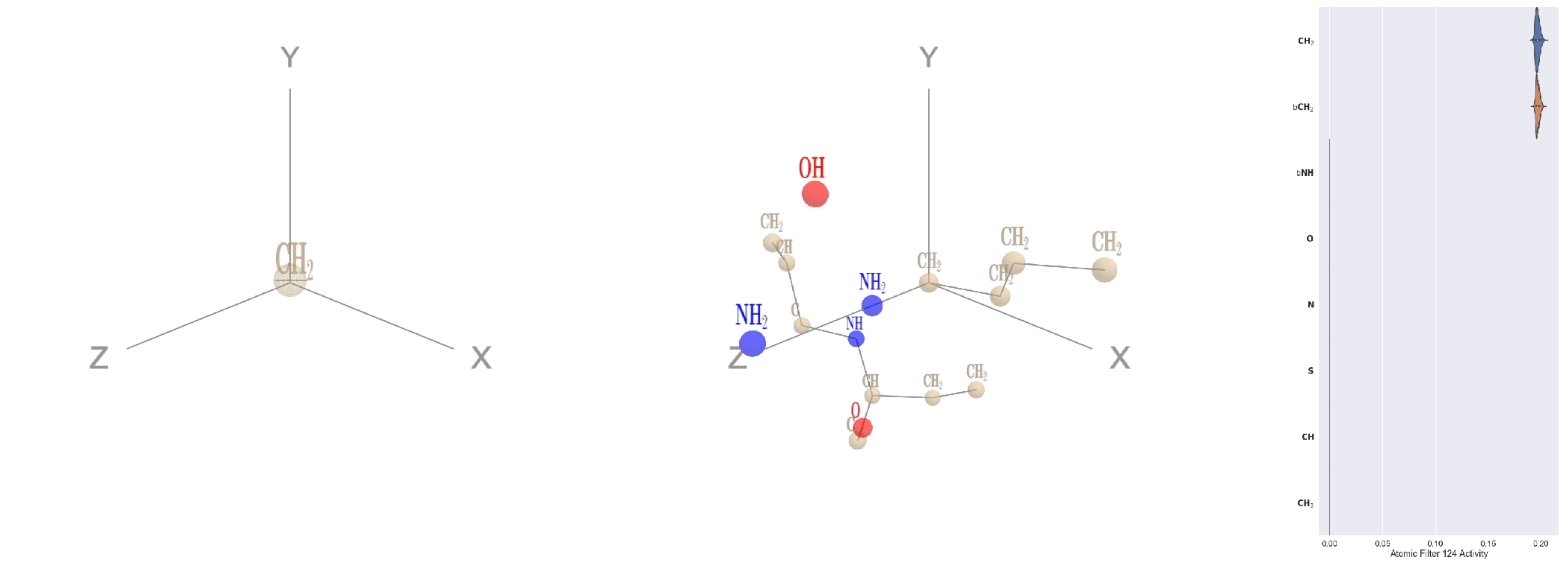


Filter 124
$R_{\text{pred}}^{\text{max}} = 0.04$
$R_{\text{pred}}^{\text{mean}} = 0.02$
Percent Active = 16
Pooling attention 0.01
Pooling feature 0.01
Top activators
2a6q:B:28 K CE
3mk7:B:155 K CE
4e0s:B:16 K CE
4kzx:E:242 K CE
1c3r:A:325 K CE
1l7k:B:150 K CE
4gu0:A:168 R CD
3ilk:A:232 K CE
4c2m:A:261 I CG1
4nmn:A:212 K CE

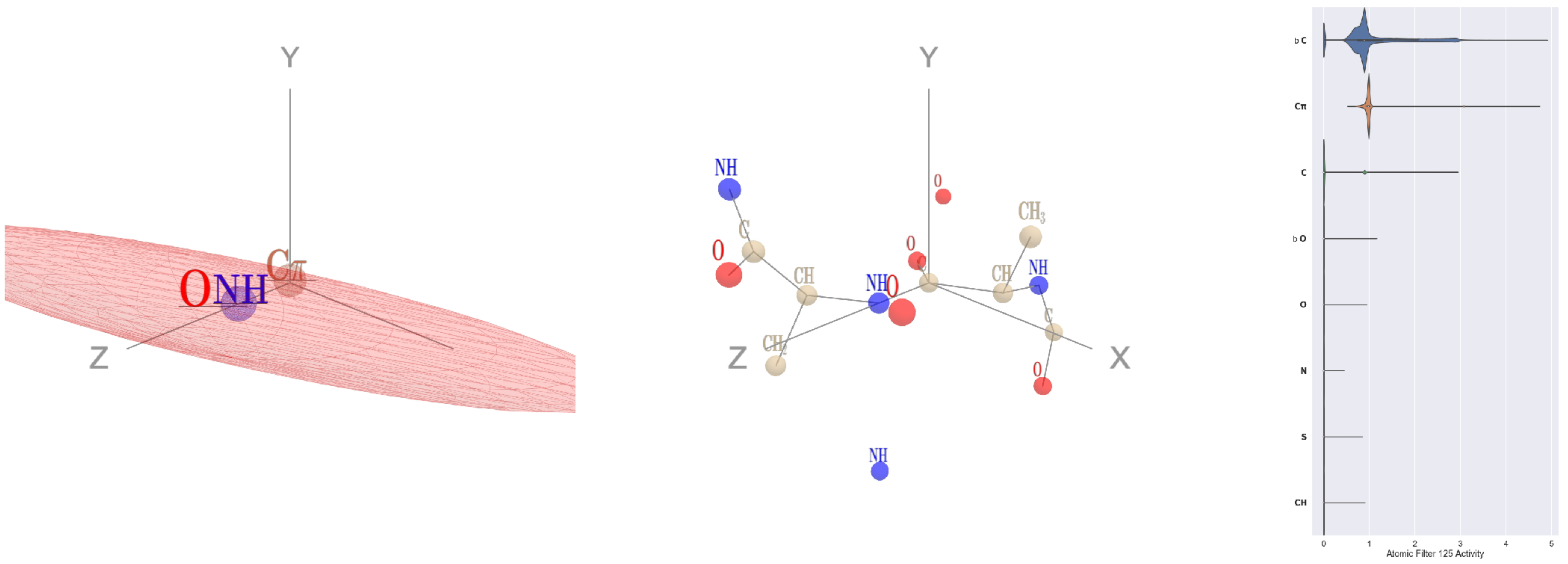


Filter 125
$R_{\text{pred}}^{\text{max}} = -0.01$
$R_{\text{pred}}^{\text{mean}} = 0.02$
Percent Active = 25
Pooling attention 0.06
Pooling feature 1.41
Top activators
3rwr:D:103 A C
2i8d:A:38 F C
1f34:B:81 V C
1lnl:A:339 Y C
4pht:A:189 P C
4hh2:D:184 G C
1fpq:A:305 V C
4c0b:B:280 L C
4cbv:D:152 K C
4rap:E:58 D C

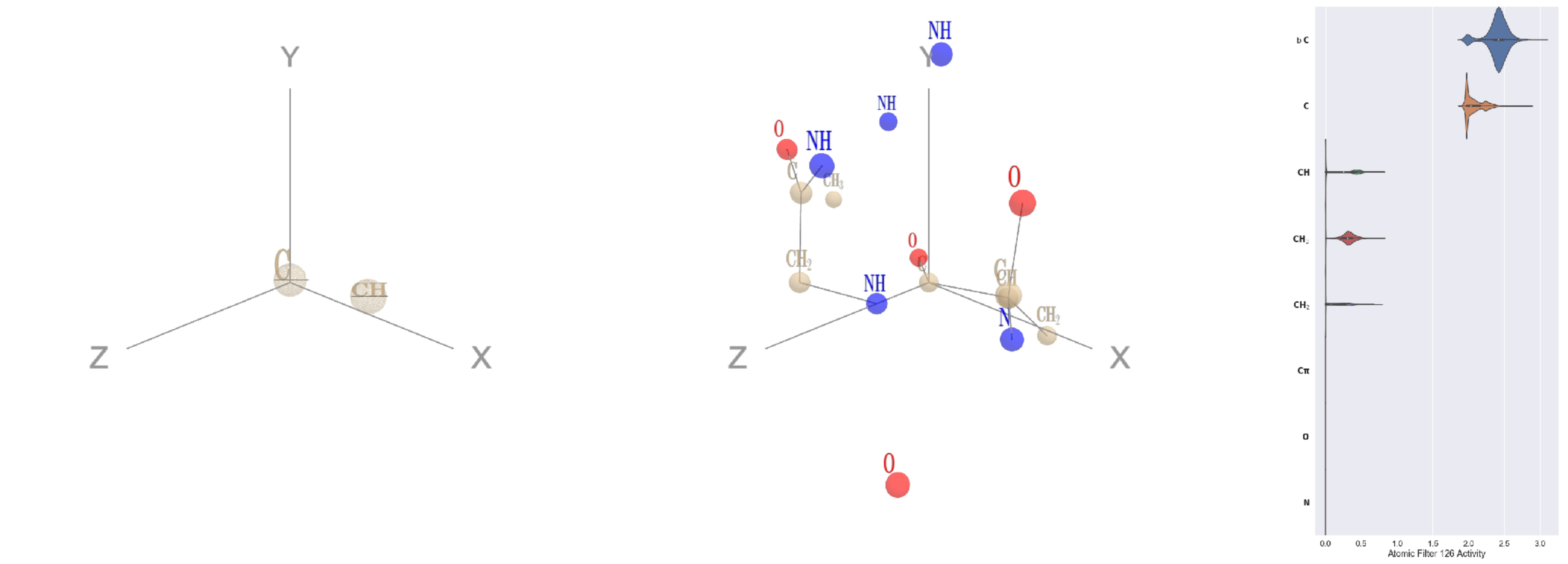


Filter 126
$R_{\text{pred}}^{\text{max}} = 0.01$
$R_{\text{pred}}^{\text{mean}} = -0.05$
Percent Active = 33
Pooling attention 0.45
Pooling feature 0.02
Top activators
3a5c:G:170 P C
4kzx:L:151 T C
3a5c:G:51 K C
3sf5:C:254 S C
1u5e:A:188 E C
2cjr:A:273 Q C
1lnl:A:317 I C
3b8e:A:1014 T C
1qha:A:568 Q C
1u5e:A:13 S C

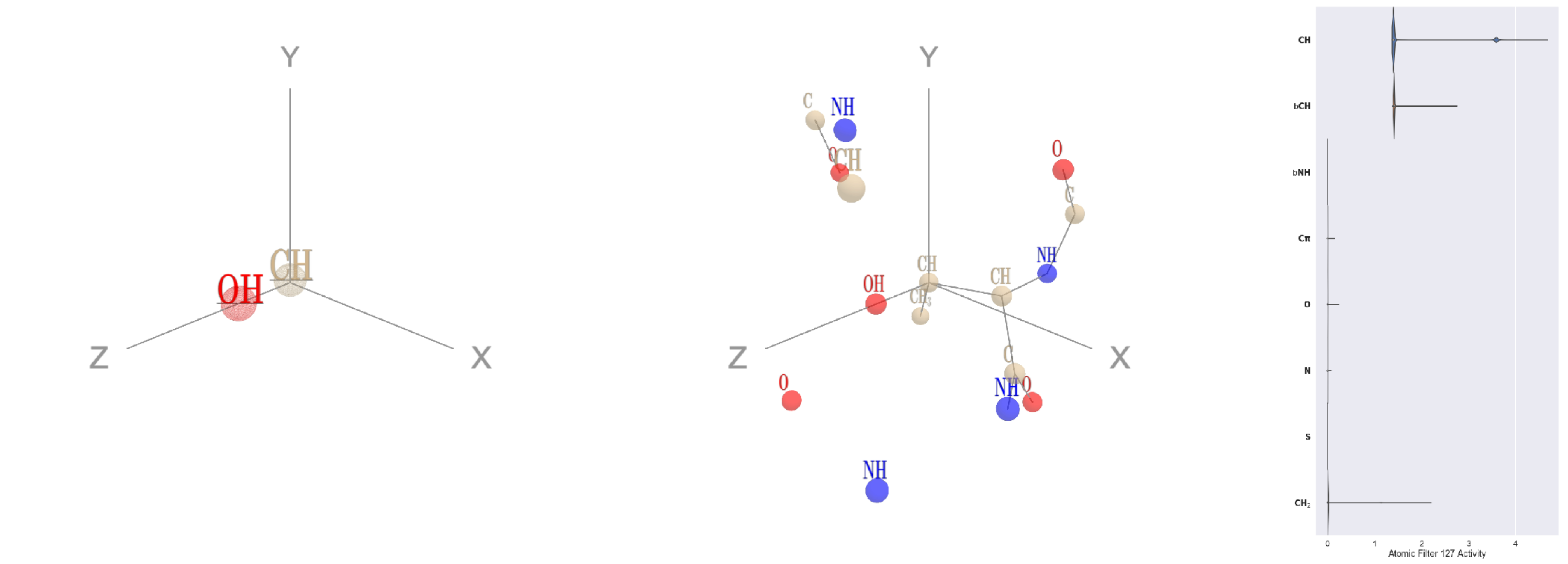


Filter 127
$R_{\text{pred}}^{\text{max}} = 0.01$
$R_{\text{pred}}^{\text{mean}} = -0.03$
Percent Active = 17
Pooling attention 0.11
Pooling feature 0.95
Top activators
1f4a:A:496 T CB
1bxr:A:923 T CB
1dtn:A:218 T CB
2d73:A:545 T CB
1ei6:A:108 T CB
1y4j:B:260 T CB
3l0o:A:311 T CB
2hb6:A:364 T CB
2v6e:A:349 T CB
5kqv:E:461 T CB
