## Supplementary material for "ScanNet: An interpretable geometric deep learning model for structure-based protein binding site prediction": 3D visualisation and activation statistics of all the amino acid filters of the protein-protein binding site network

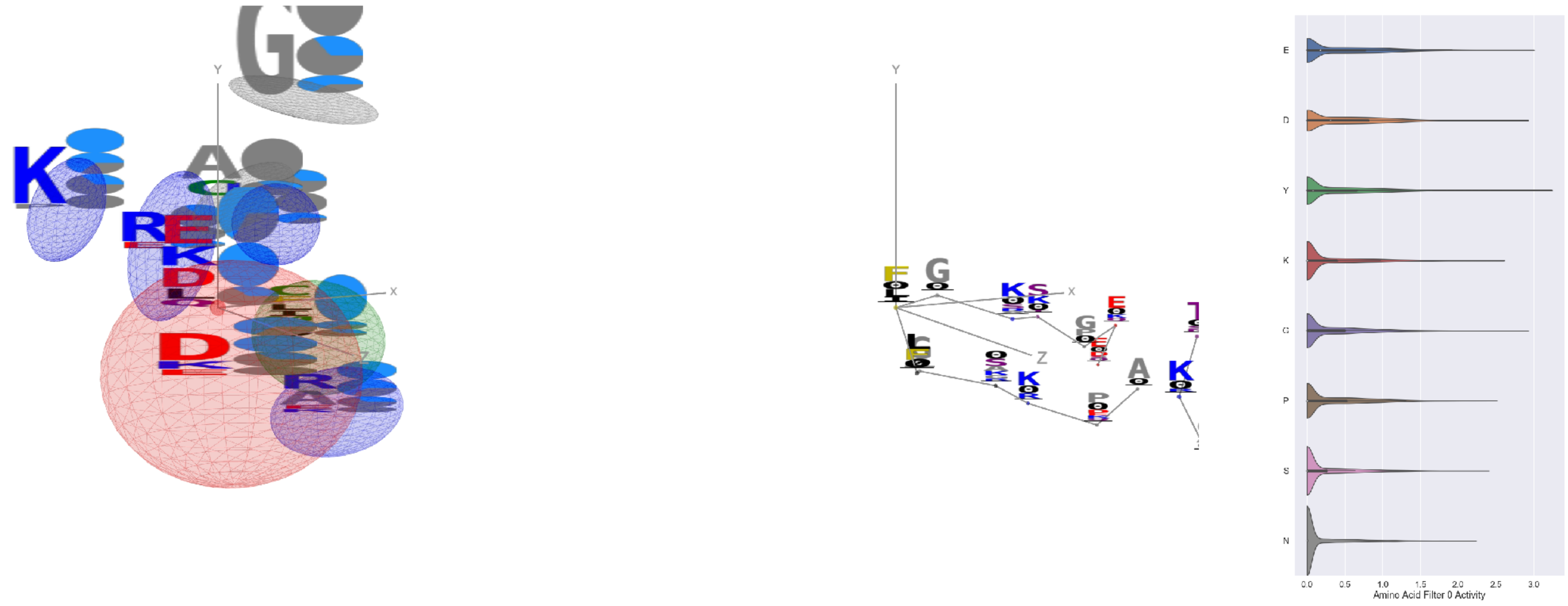

Filter 0
$R_{\mathrm{pred}} = -0.09$
Percent Active = 36
Pooling attention 0.50
Pooling feature 1.43
Top activators
3gku:A:41 F
2yma:A:399 Y
4rg1:A:374 Y
4pht:A:497 Y
4buj:B:191 E
4buj:B:239 E
2prv:A:106 Y
4asi:C:2200 E
2fq1:A:283 Y
4buj:B:190 G

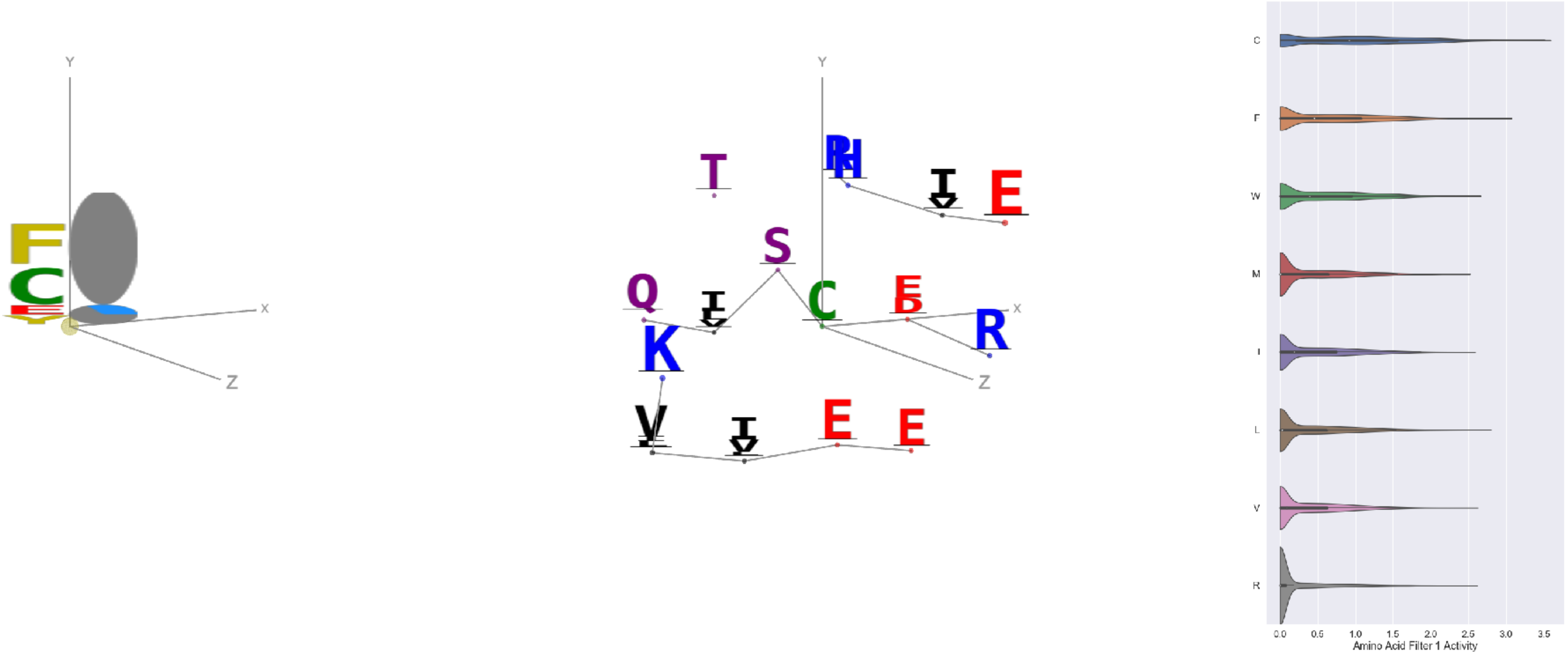

Filter 1
$R_{\mathrm{pred}} = -0.24$
Percent Active = 33
Pooling attention 0.38
Pooling feature 2.63
Top activators
3n6r:G:290 C
3bof:A:207 C
5kqv:E:237 C
5kqv:E:212 C
4cy6:A:359 E
1u6z:B:23 F
5kqv:E:208 C
3ax7:A:113 C
2x98:A:210 F
3qzy:B:50 C

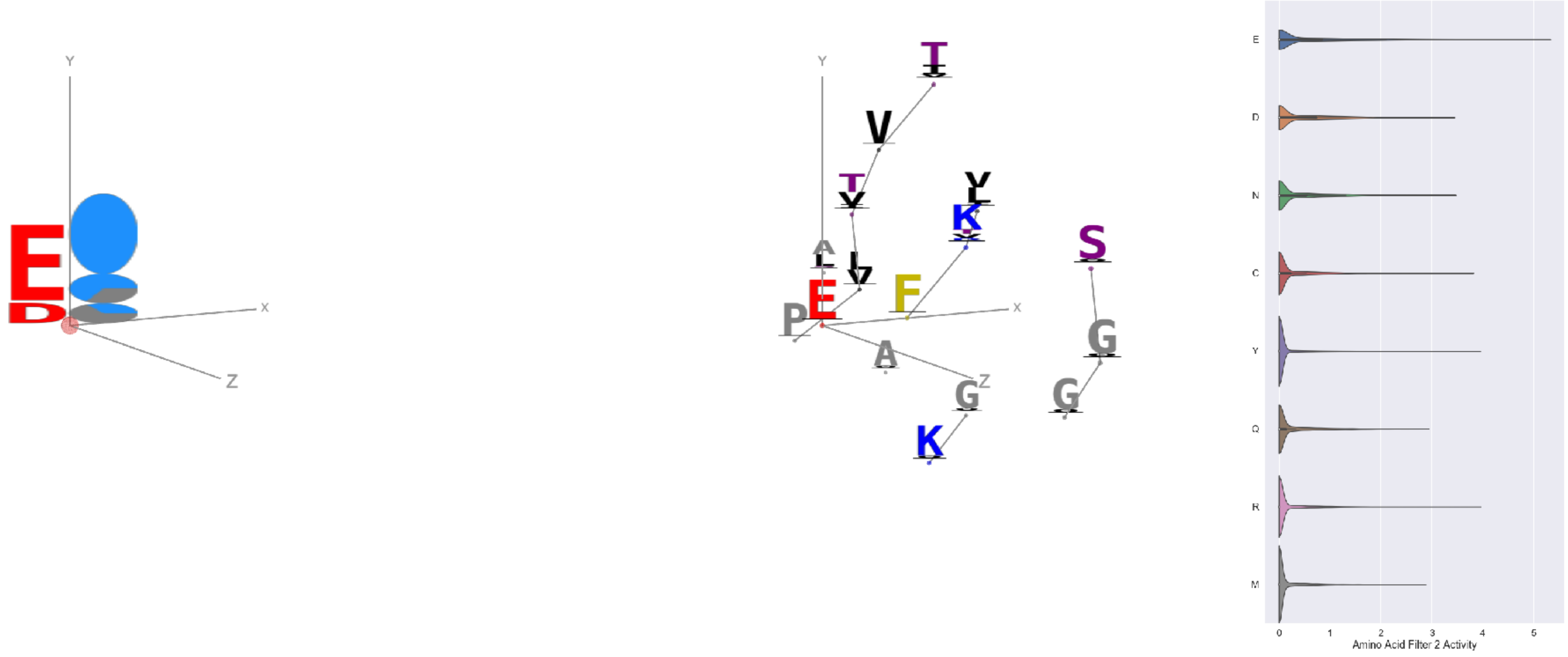

Filter 2
$R_{\mathrm{pred}} = 0.09$
Percent Active = 16
Pooling attention 0.47
Pooling feature 1.65
Top activators
3tu3:B:377 E
4l2w:D:902 E
4c2m:A:646 E
1e1c:A:393 E
1gvn:A:16 E
3tsy:A:395 E
1bxr:A:81 E
1bxr:A:349 E
1qvr:A:271 E
4oph:A:71 E

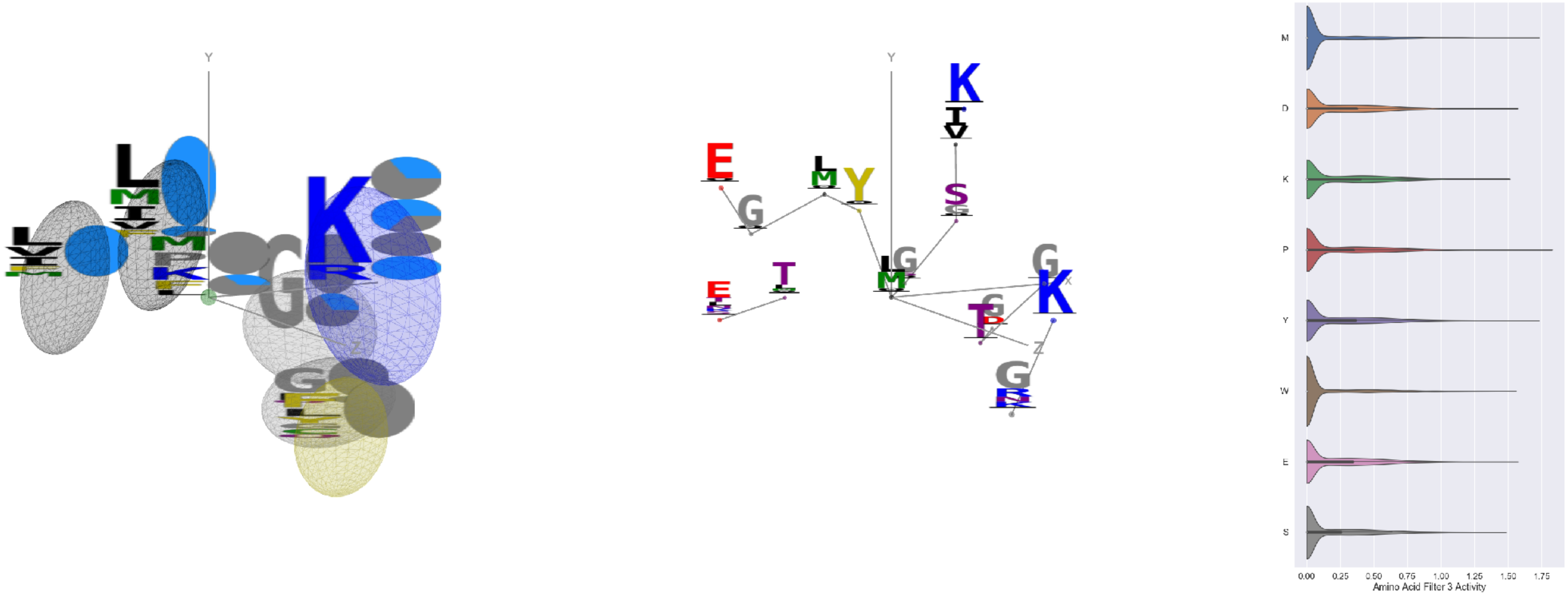

Filter 3
$R_{\mathrm{pred}} = 0.20$
Percent Active = 33
Pooling attention 0.02
Pooling feature 2.39
Top activators
1qha:A:300 L
4f52:E:480 F
4a7z:A:262 P
4oph:A:65 V
3t3o:A:495 V
1mu2:A:157 P
1qha:A:748 M
2iwh:B:439 Y
4kzx:R:66 V
2iwh:B:831 M

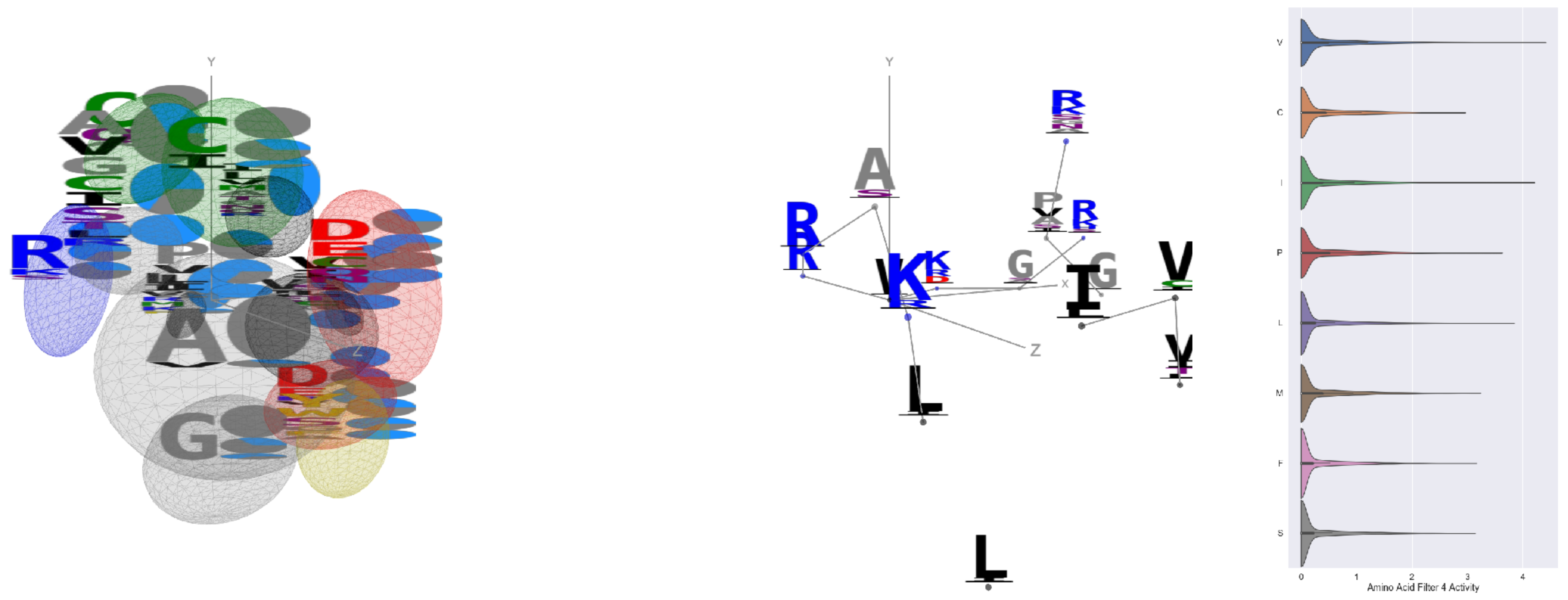


Filter 4
$R_{pred} = 0.00$
Percent Active = 27
Pooling attention 0.36
Pooling feature 3.15
Top activators
3pio:M:99 V
4e0s:B:158 V
3tsy:A:57 I
3tsy:A:399 V
3s9m:C:624 Y
2d4z:B:547 V
3do6:A:515 I
3tsy:A:600 A
2iwh:B:573 L
1v8f:A:171 T

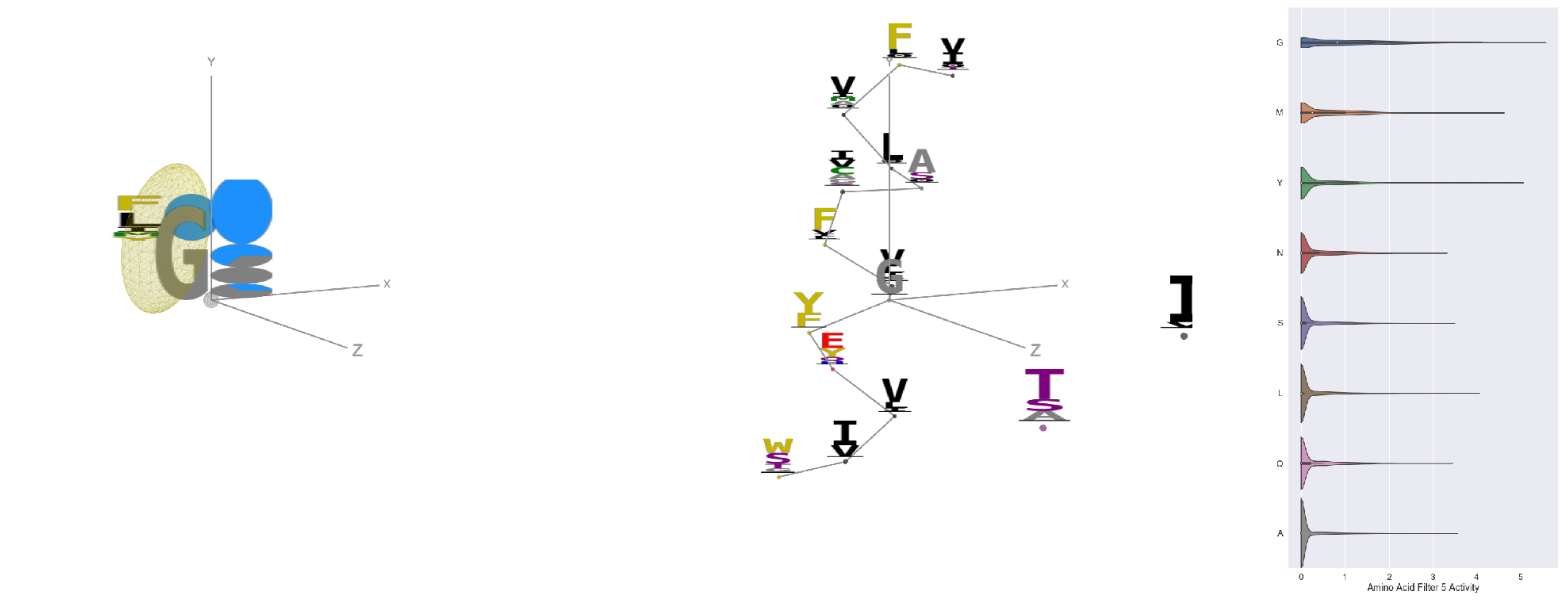


Filter 5
$R_{pred} = 0.18$
Percent Active = 28
Pooling attention 0.07
Pooling feature 0.69
Top activators
4o93:A:19 G
5uqy:A:141 G
4ehi:B:364 G
4d0l:A:704 G
1vl4:A:158 G
1ezv:C:381 Y
3nqr:D:173 G
1mu2:A:461 G
4e0s:B:92 G
3tsy:A:943 G

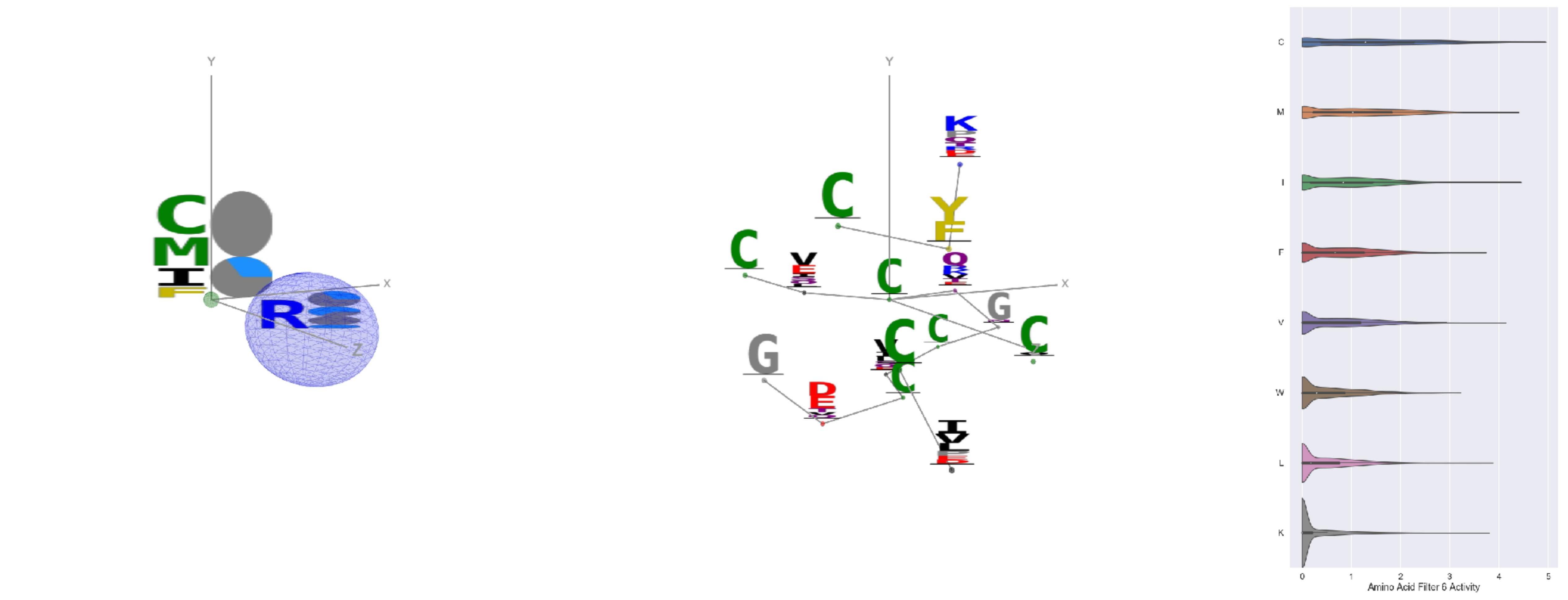


Filter 6
$R_{pred} = -0.11$
Percent Active = 29
Pooling attention 0.07
Pooling feature 0.33
Top activators
4g1m:B:506 C
1h8p:A:69 C
1h8p:A:83 C
4e0s:B:520 C
2gy7:B:224 C
1i3r:B:41 C
1h8p:A:24 C
2iwh:B:796 I
2gy7:B:227 C
4oph:A:104 M

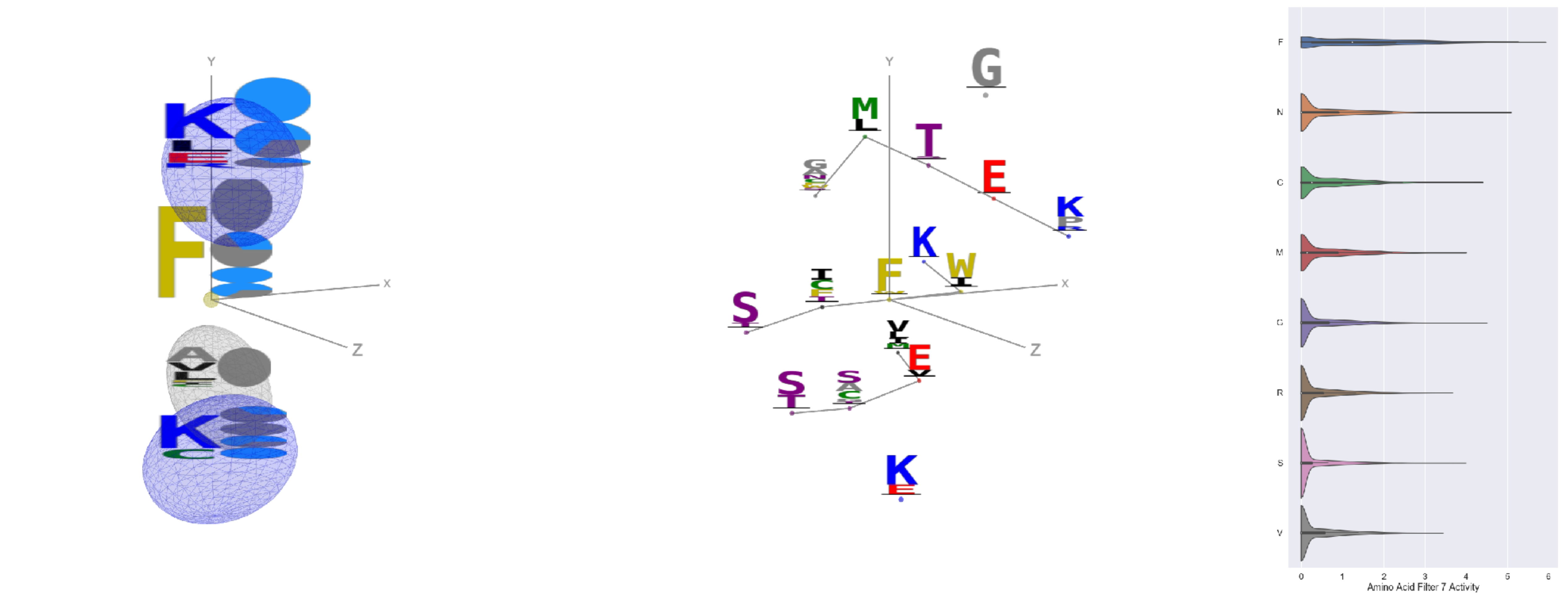


Filter 7
$R_{pred} = -0.15$
Percent Active = 34
Pooling attention 0.04
Pooling feature 0.61
Top activators
3u33:A:216 F
1bxr:A:618 F
4cvw:A:470 F
1qvr:A:591 F
4nm9:B:613 F
4ny2:A:297 F
1e1c:A:117 F
4v8p:BC:29 F
3tsy:A:716 F
2iwh:B:383 F

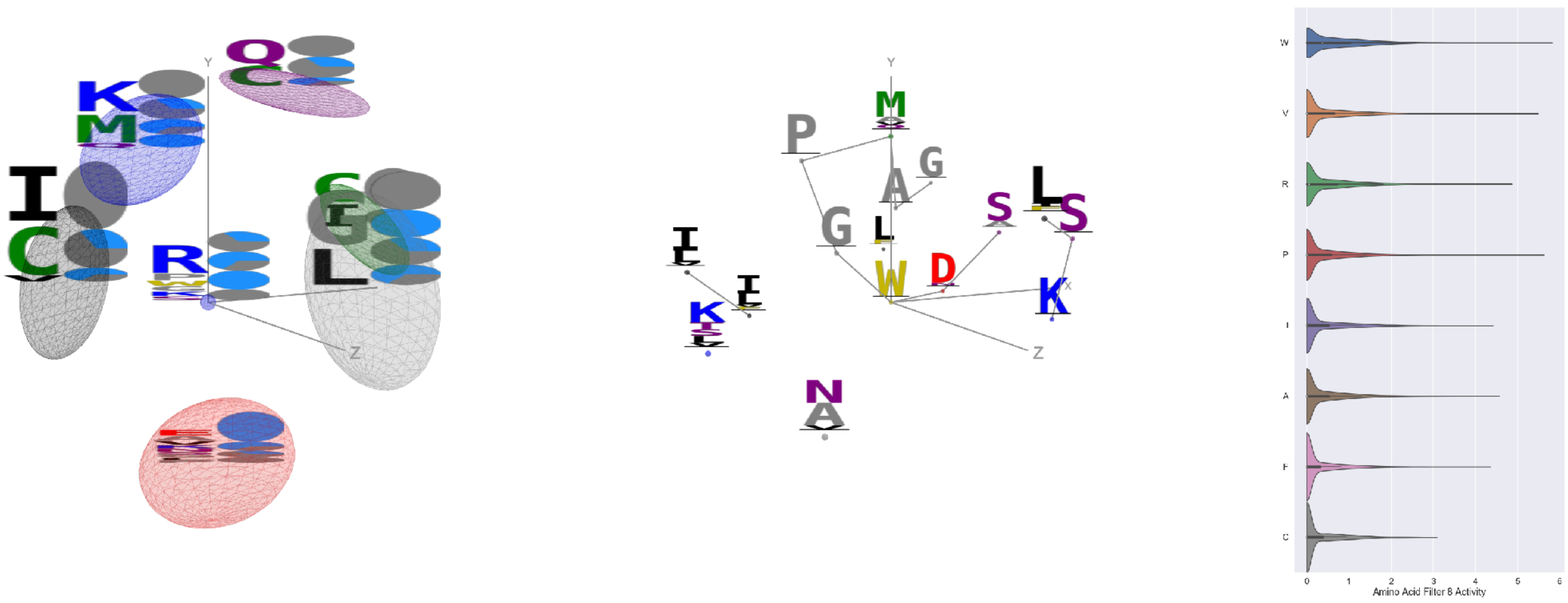


Filter 8
$R_{\text{pred}} = -0.12$
Percent Active = 31
Pooling attention 0.42
Pooling feature 0.23
Top activators
1k90:C:357 W
4ri0:A:308 P
3tsy:A:775 V
1jb0:K:70 G
2iwh:B:291 P
3g8q:A:65 R
1izl:D:280 W
4pht:A:323 P
4v8p:BC:256 V
2gy7:B:284 P

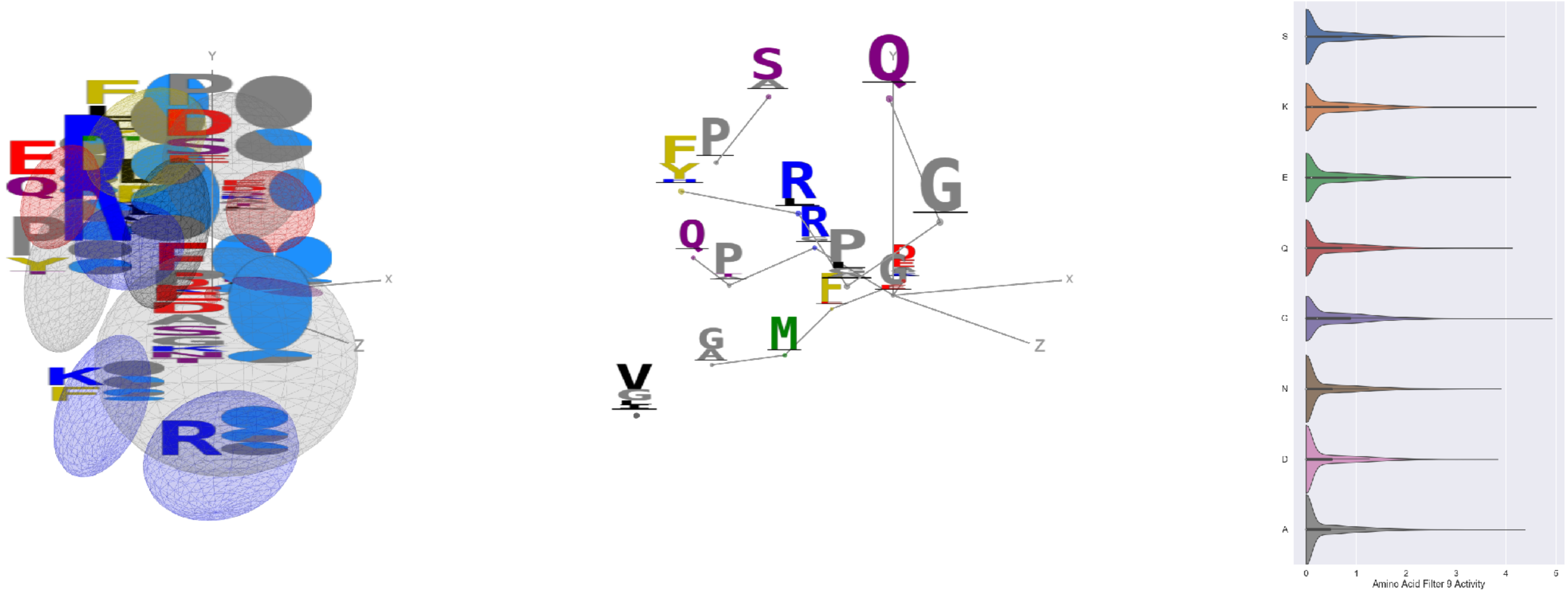


Filter 9
$R_{\text{pred}} = -0.01$
Percent Active = 39
Pooling attention 0.05
Pooling feature 1.50
Top activators
3tu3:B:292 G
1qvr:A:191 K
1ezv:C:288 K
3noy:B:38 A
1n0e:B:39 G
4rg1:A:138 P
3s9m:C:262 L
4v81:H:158 Q
3wb2:A:14 E
1bxr:A:748 H

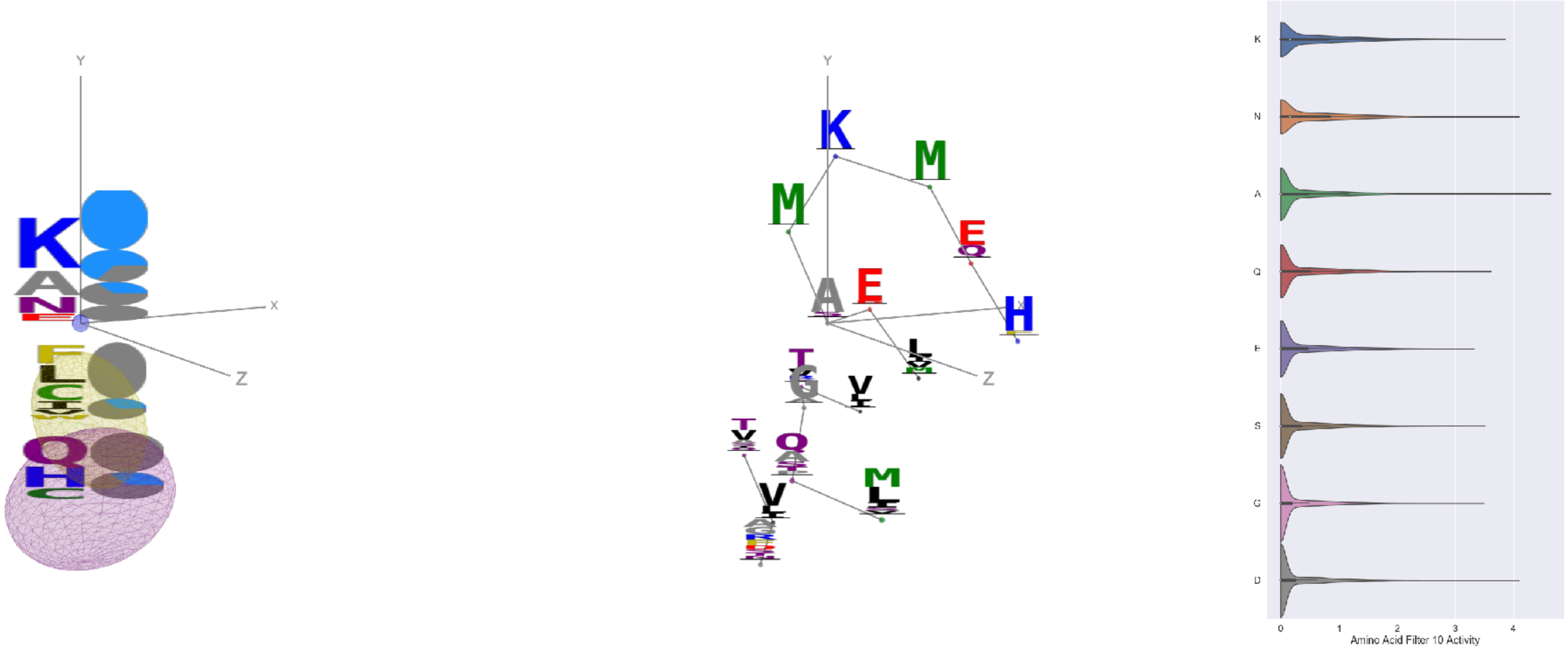


Filter 10
$R_{\text{pred}} = -0.04$
Percent Active = 27
Pooling attention 0.54
Pooling feature 0.44
Top activators
4rcn:B:520 A
1eg1:A:173 A
1iwa:A:241 N
3qzy:B:194 D
5hu6:C:201 A
1ezv:C:260 A
1qha:A:69 R
3igh:X:278 K
4v81:A:323 K
1eg1:A:221 K

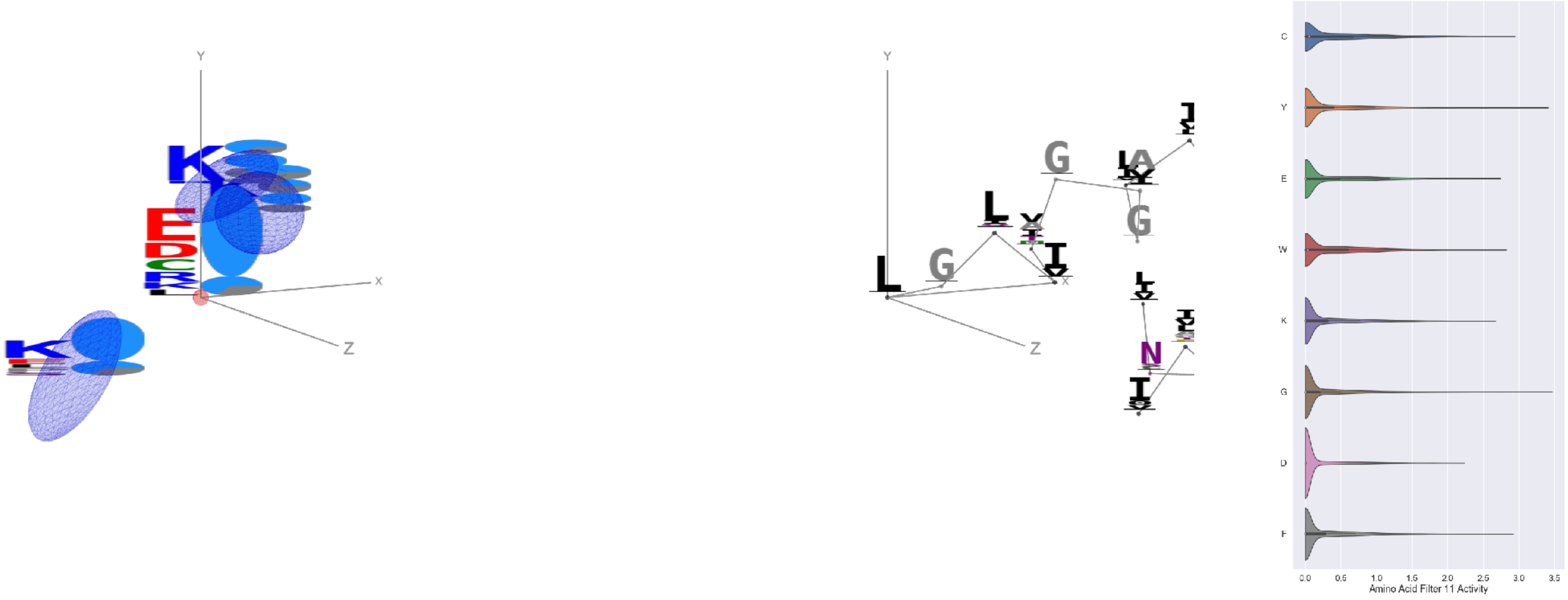


Filter 11
$R_{\text{pred}} = -0.01$
Percent Active = 30
Pooling attention 0.31
Pooling feature 0.93
Top activators
1jb0:K:77 L
4qiw:A:310 G
3gku:A:208 Y
2dyo:A:285 Y
5wy5:A:246 Y
3vr6:G:207 Y
4pba:A:318 Y
1qc9:A:130 I
3di4:A:0 Y
1srq:A:413 Y

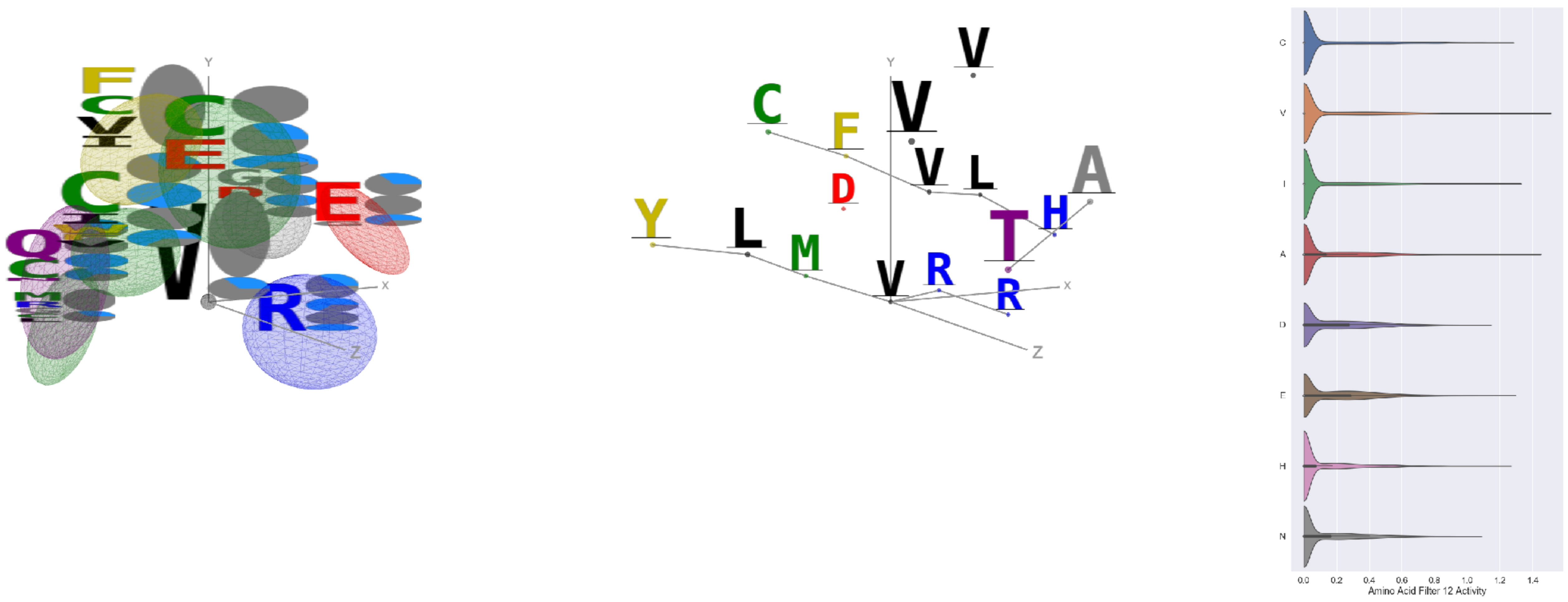


Filter 12
$R_{pred} = 0.11$
Percent Active = 32
Pooling attention 0.35
Pooling feature 3.56
Top activators
3tsy:A:744 V
2iut:A:400 V
1r7r:A:694 A
4g1m:B:579 G
3a5c:G:147 L
1r7r:A:569 A
4e0s:B:59 I
1hfe:L:40 R
2gy7:B:300 E
4e0s:B:106 C

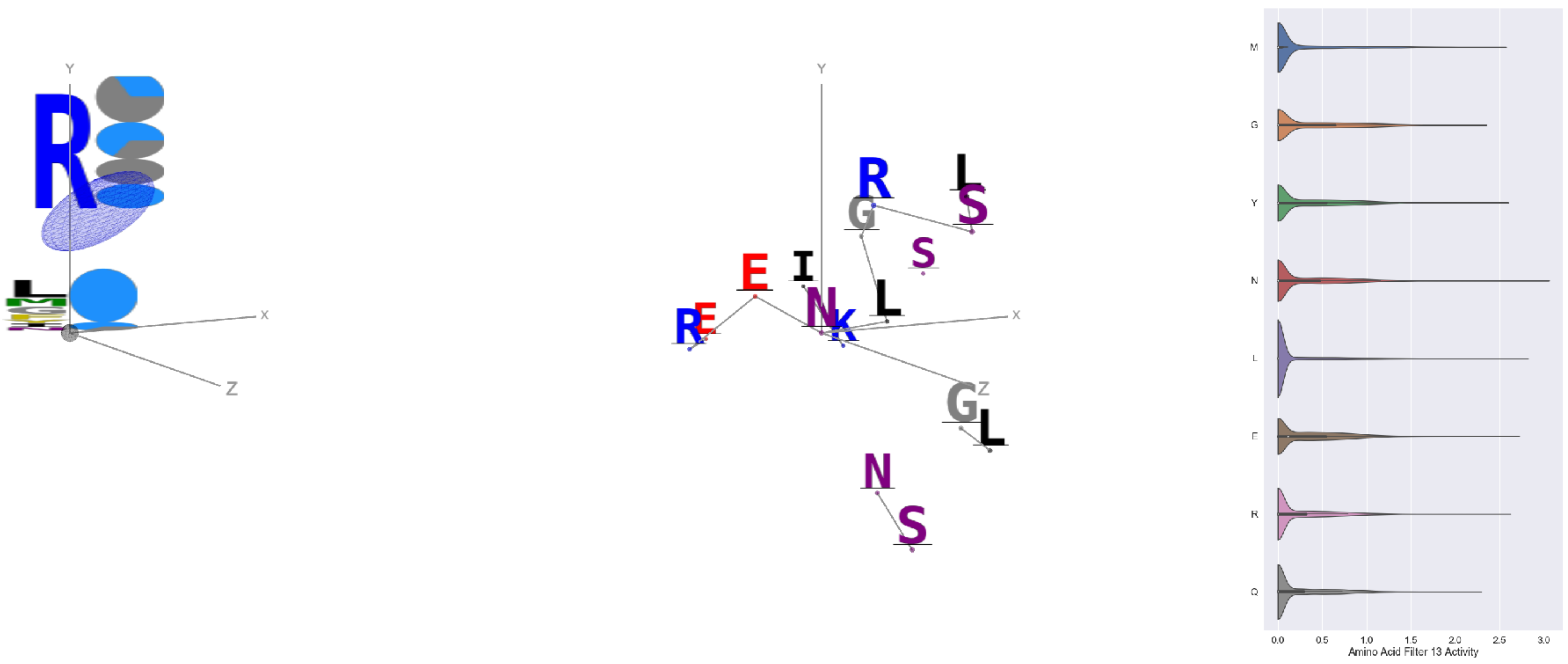


Filter 13
$R_{pred} = 0.28$
Percent Active = 35
Pooling attention 0.12
Pooling feature 0.17
Top activators
2iwh:B:255 N
4c2m:A:1006 L
4u4c:A:549 L
3g8q:A:210 E
4mex:C:473 R
2q5d:C:58 Y
3u33:A:415 Y
2iwh:B:768 M
2np9:A:395 L
3n6r:G:695 M

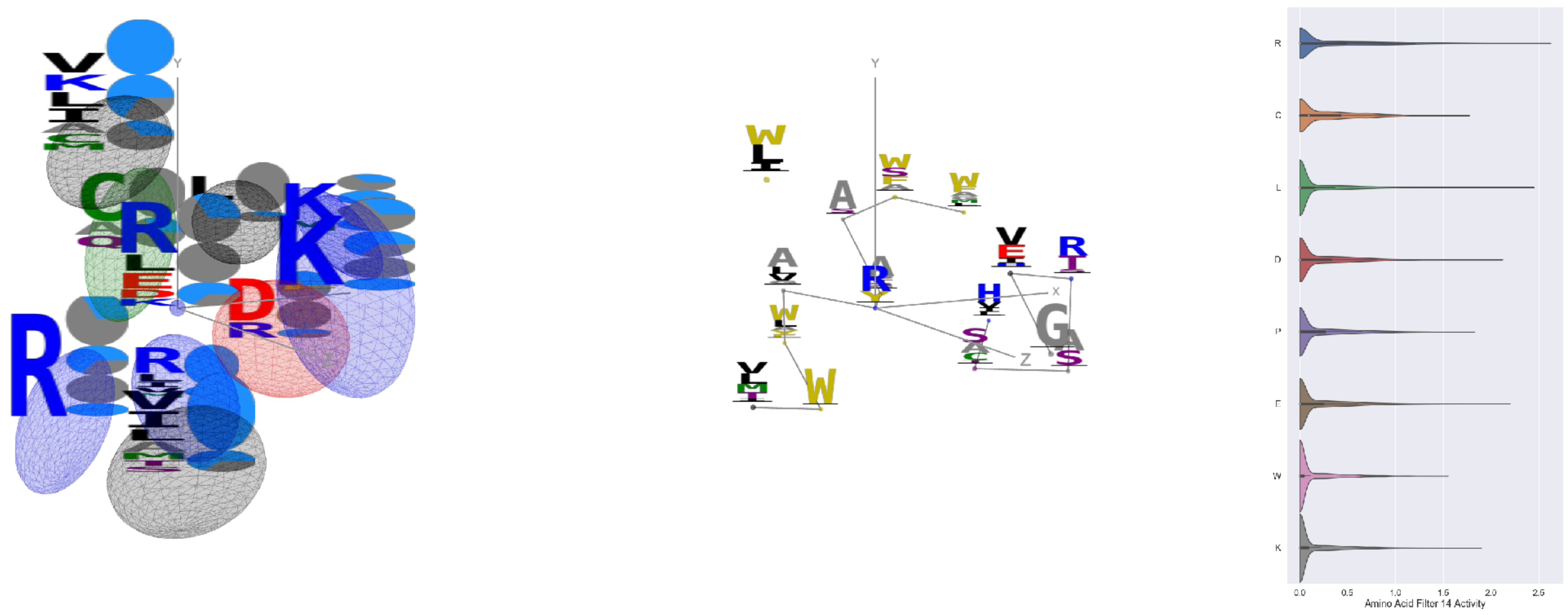


Filter 14
$R_{pred} = -0.15$
Percent Active = 26
Pooling attention 0.09
Pooling feature 0.12
Top activators
3hzu:A:106 R
4pk7:A:296 R
4nm9:B:385 L
4nm9:B:615 R
4cvw:A:548 R
2v6e:A:274 R
2d73:A:517 R
1iwa:A:319 R
4ny2:A:174 R
4r3z:C:267 R

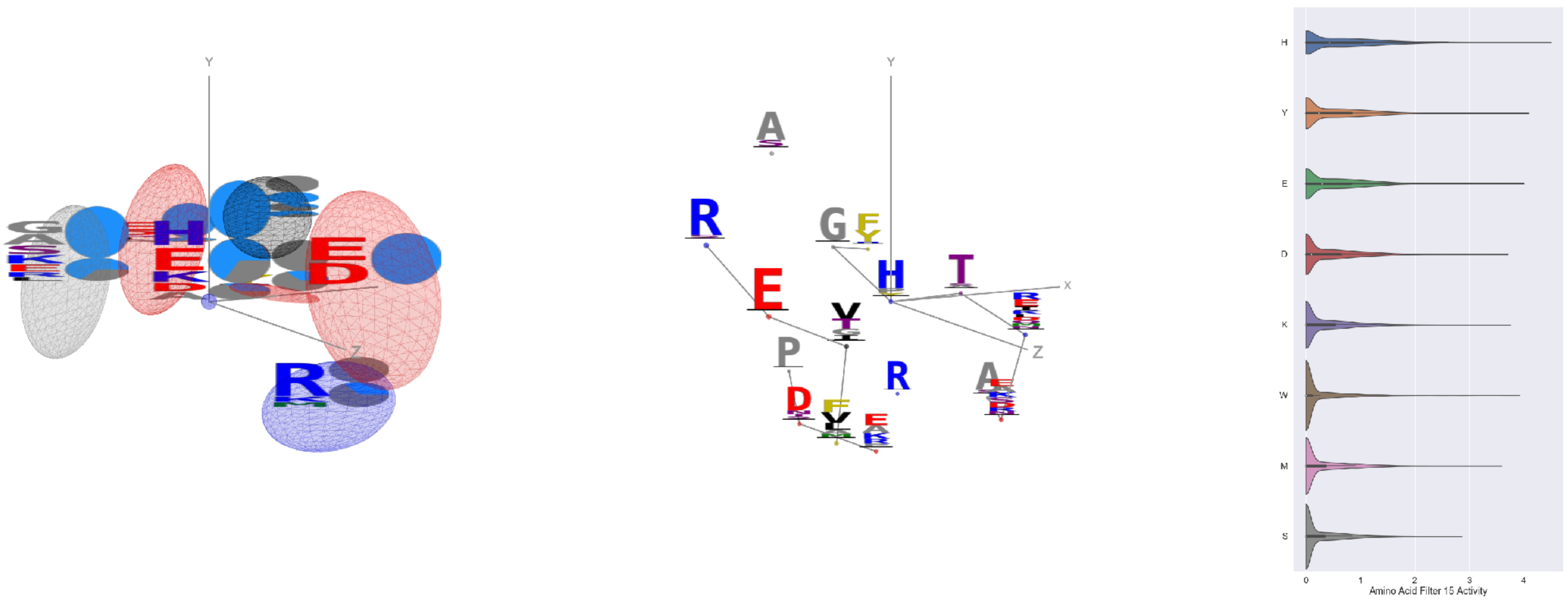


Filter 15
$R_{pred} = -0.07$
Percent Active = 38
Pooling attention 0.47
Pooling feature 1.14
Top activators
1r7r:A:404 H
1izl:D:304 R
1izl:D:196 F
1izl:D:348 Y
1izl:D:311 Y
4buj:B:154 P
1sl0:A:498 H
1r7r:A:501 E
3kio:B:277 Y
1izl:D:61 Y

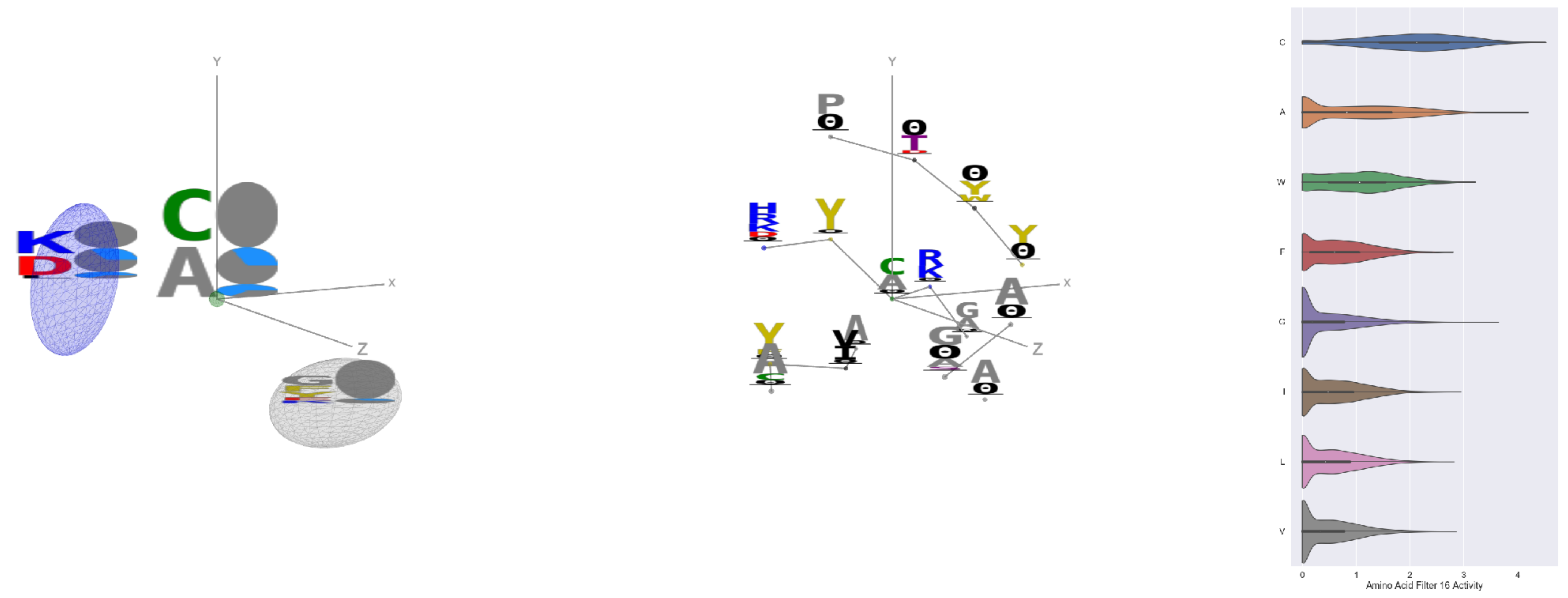

Filter 16
$R_{\mathrm{pred}} = -0.21$
Percent Active = 42
Pooling attention 0.12
Pooling feature 0.35
Top activators
1iwa:A:84 C
2gy7:B:240 C
1eg1:A:193 C
3bof:A:207 C
3v4l:A:472 C
1dci:A:170 A
2iwh:B:369 A
2gy7:B:227 C
1qha:A:834 C
1eg1:A:144 A

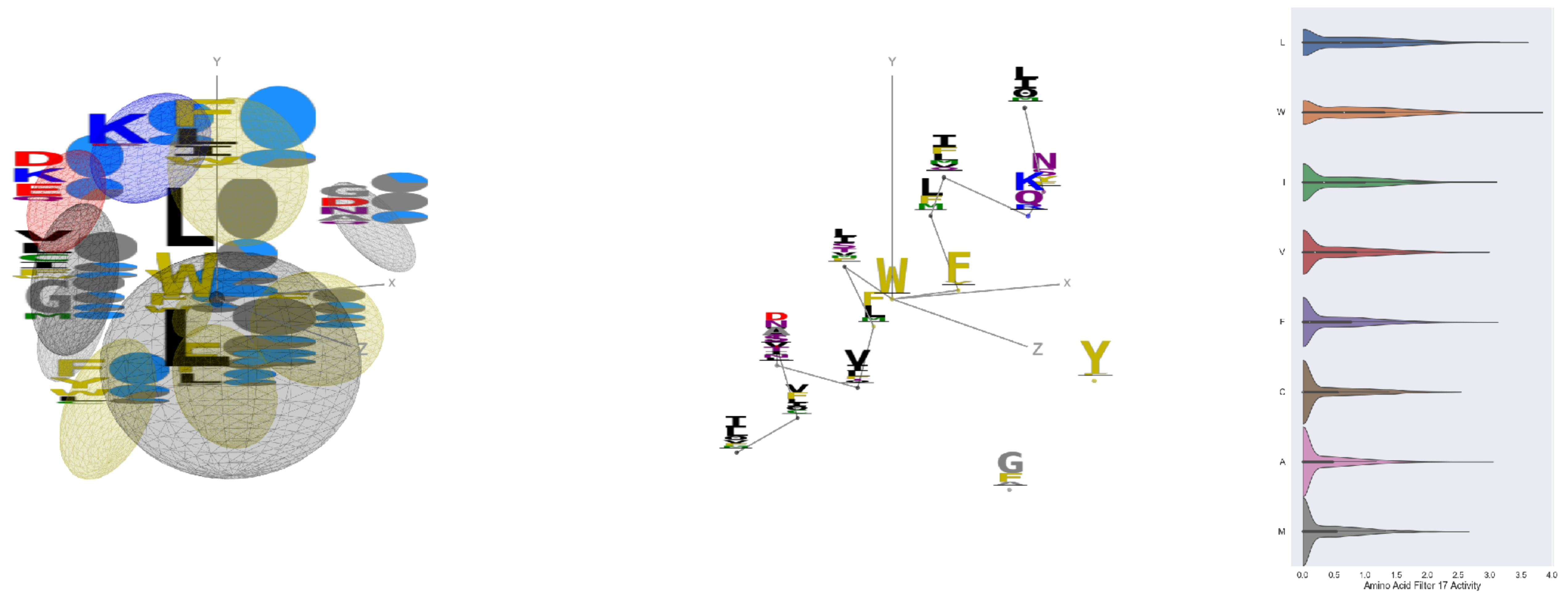

Filter 17
$R_{\mathrm{pred}} = 0.19$
Percent Active = 44
Pooling attention 0.21
Pooling feature 1.39
Top activators
1ezv:C:327 W
3b8e:A:989 L
5kqv:E:62 L
4o93:A:14 L
2hh1:H:27 L
4pk7:A:464 L
2hh1:H:21 W
4buj:B:1022 W
1jqn:A:356 L
4o93:A:9 L

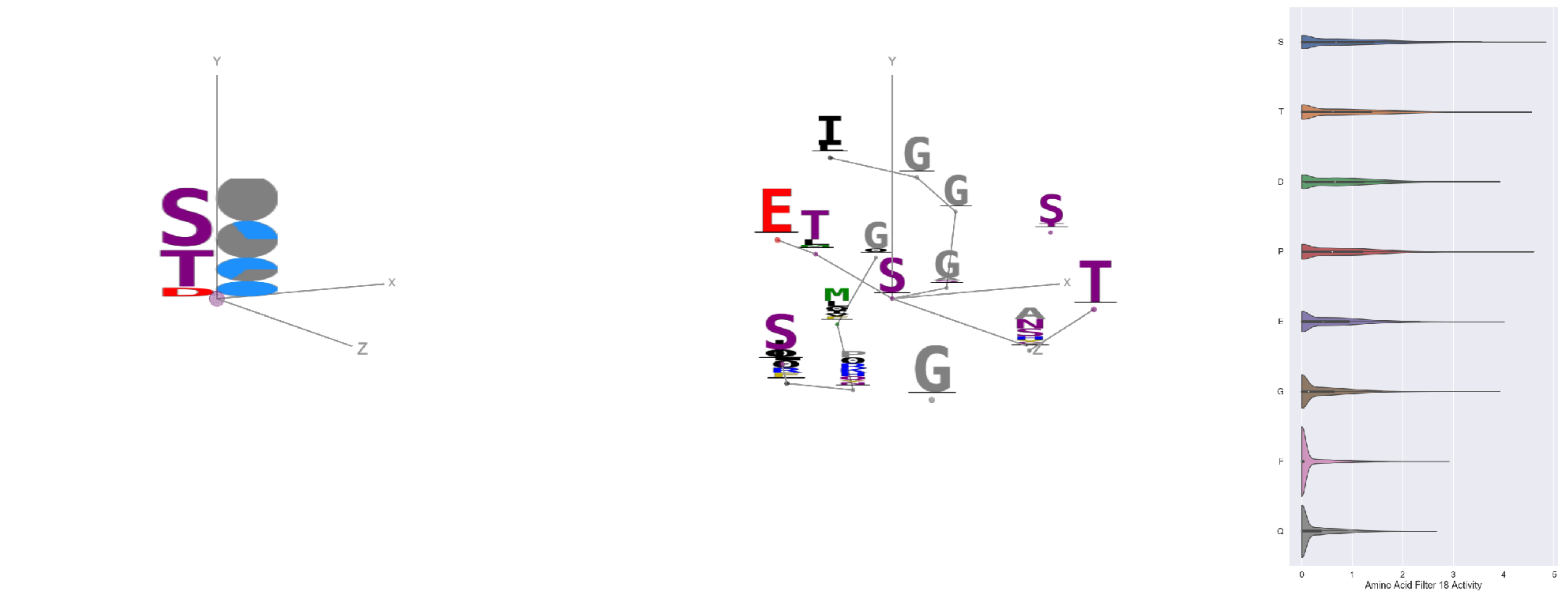

Filter 18
$R_{\mathrm{pred}} = -0.06$
Percent Active = 32
Pooling attention 0.02
Pooling feature 0.73
Top activators
1u6z:B:148 S
2pif:B:236 P
1p4e:B:185 T
2zy4:F:314 S
1cm5:A:535 S
1bo1:A:280 S
1bxr:A:312 S
2iwh:B:707 S
1bxr:A:792 S
1bxr:A:531 T

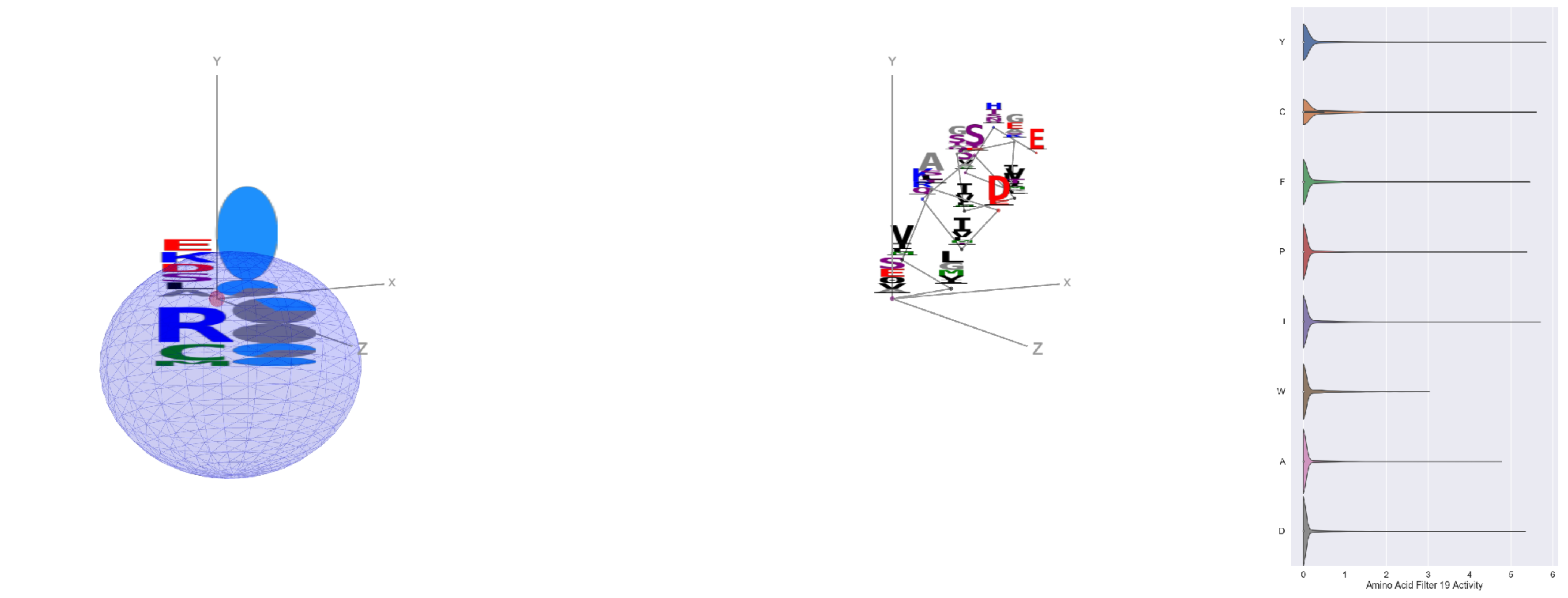

Filter 19
$R_{\mathrm{pred}} = -0.03$
Percent Active = 14
Pooling attention 0.09
Pooling feature 1.96
Top activators
3te3:A:733 S
1am4:A:234 Y
3u33:A:540 Y
1mu2:A:555 I
4gu0:A:822 Y
3bf0:C:551 Y
1h8p:A:109 C
4pk7:A:1047 Y
4kbj:B:425 F
4cyj:E:406 Y

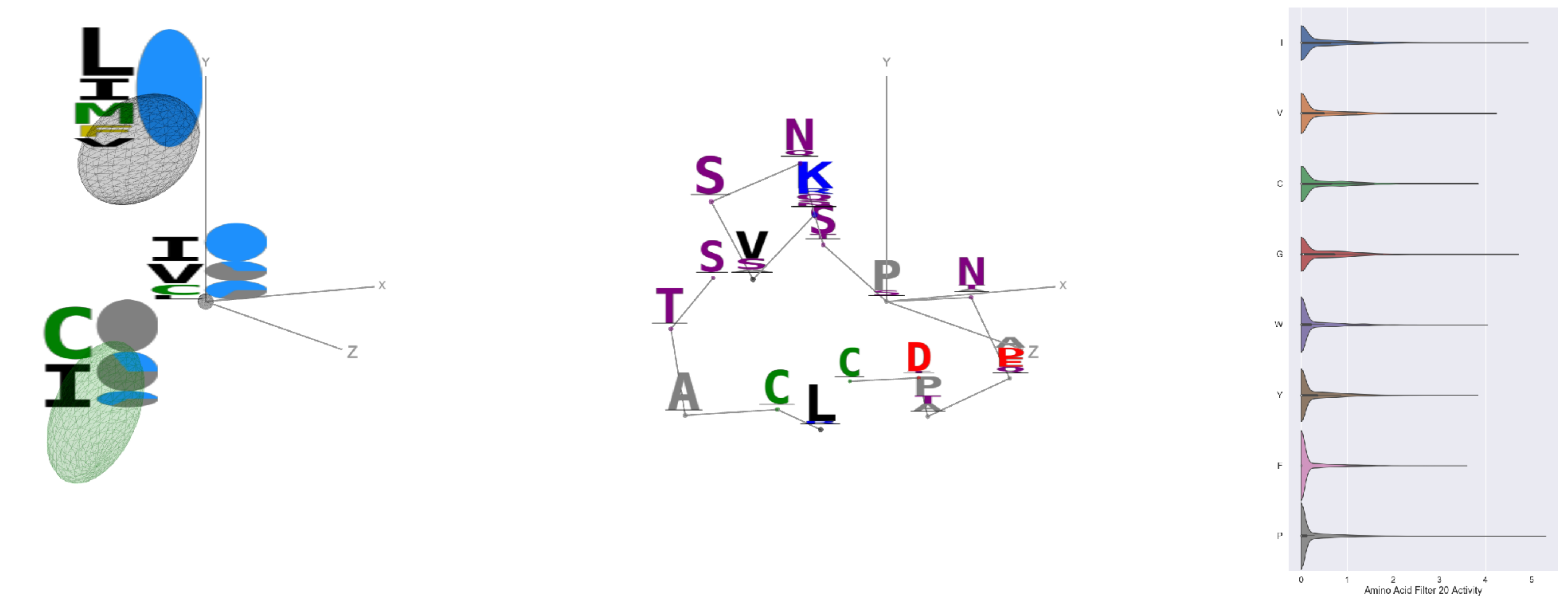

Filter 20
$R_{\text{pred}} = 0.20$
Percent Active = 28
Pooling attention 0.19
Pooling feature 0.31
Top activators
4ny2:A:205 P
4e0s:B:905 I
2gy7:B:320 G
1eg1:A:218 V
4pfp:A:95 V
3d00:A:179 L
3lnm:D:300 I
2pu9:A:72 W
1i3r:B:184 L
1ezv:C:290 G

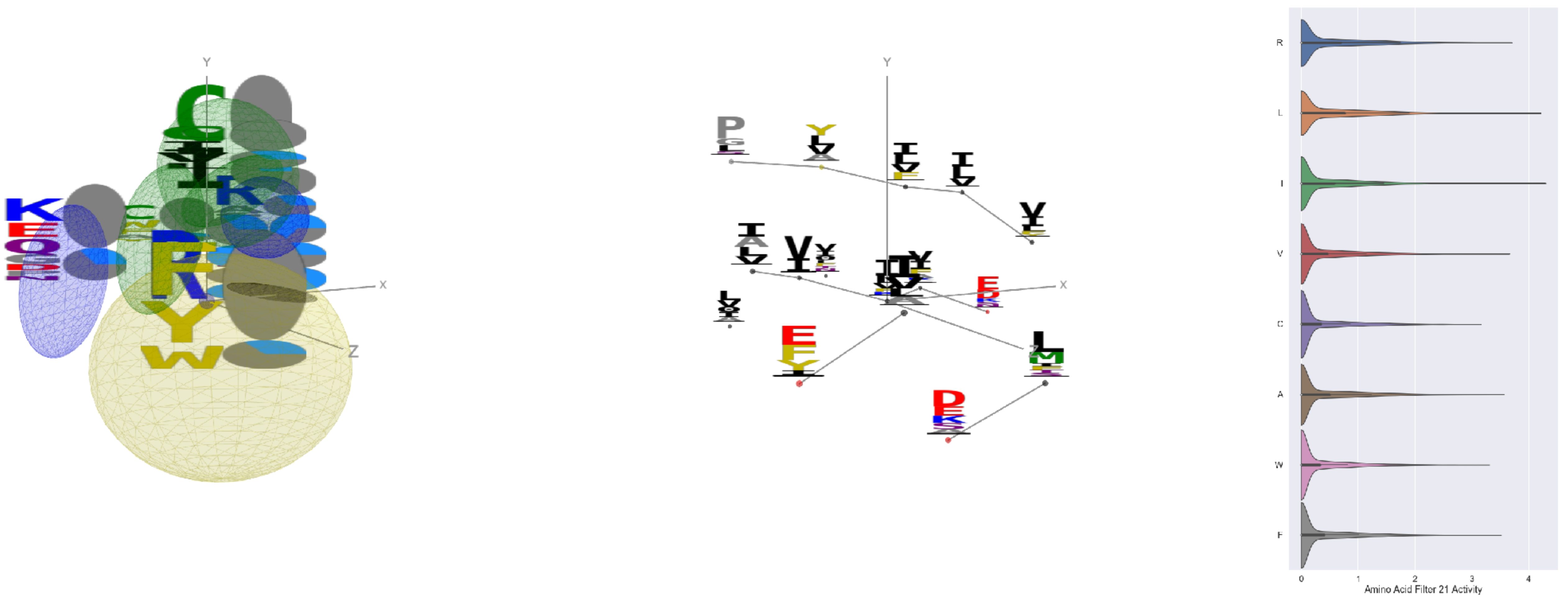

Filter 21
$R_{\text{pred}} = -0.13$
Percent Active = 33
Pooling attention 0.15
Pooling feature 0.23
Top activators
2i71:B:86 I
3tsy:A:746 L
1u6z:B:161 L
4jds:A:291 L
4gu0:A:420 R
5ui2:A:218 L
3s9m:C:369 V
2ba0:B:140 V
1he8:A:482 L
1w36:B:54 L

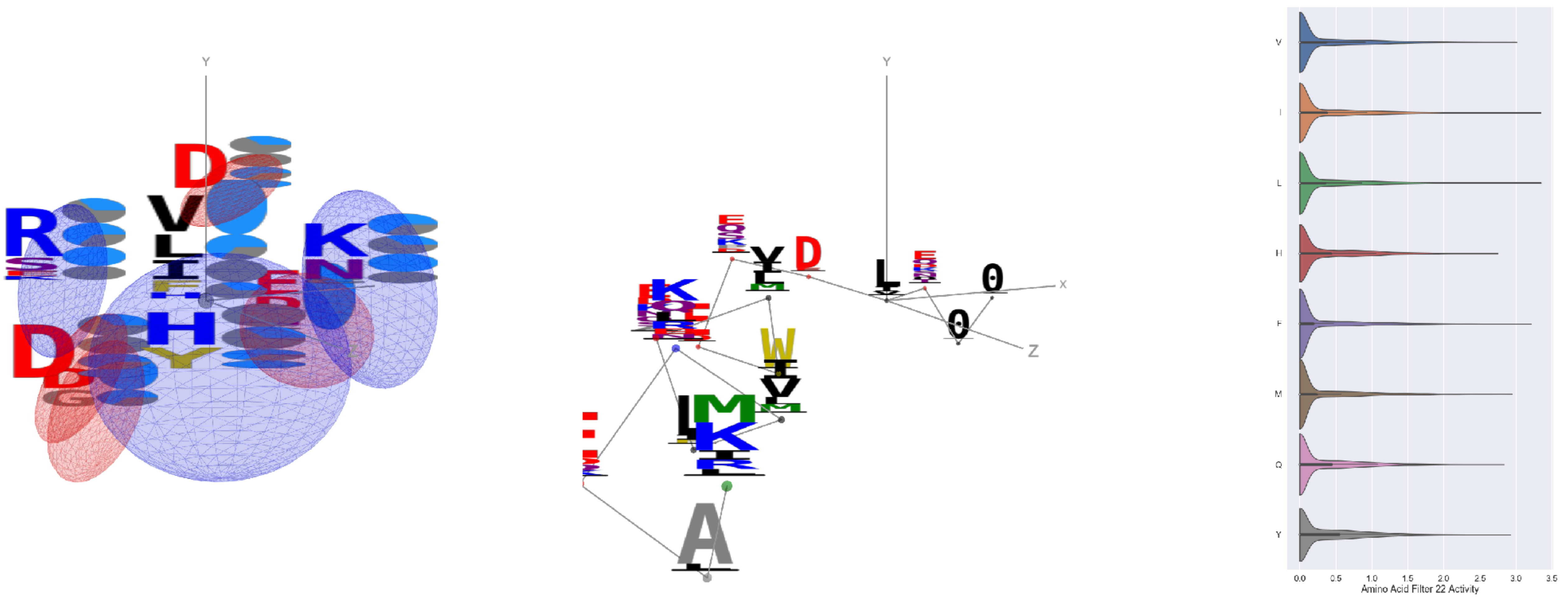

Filter 22
$R_{\text{pred}} = 0.29$
Percent Active = 34
Pooling attention 0.60
Pooling feature 1.51
Top activators
2yal:A:74 L
1n0w:B:1534 I
2iwh:B:498 I
3t5v:C:35 F
1cm5:A:547 I
3r3p:B:219 R
1efu:B:271 F
3wqa:A:3347 L
3klt:A:309 L
2iwh:B:779 D

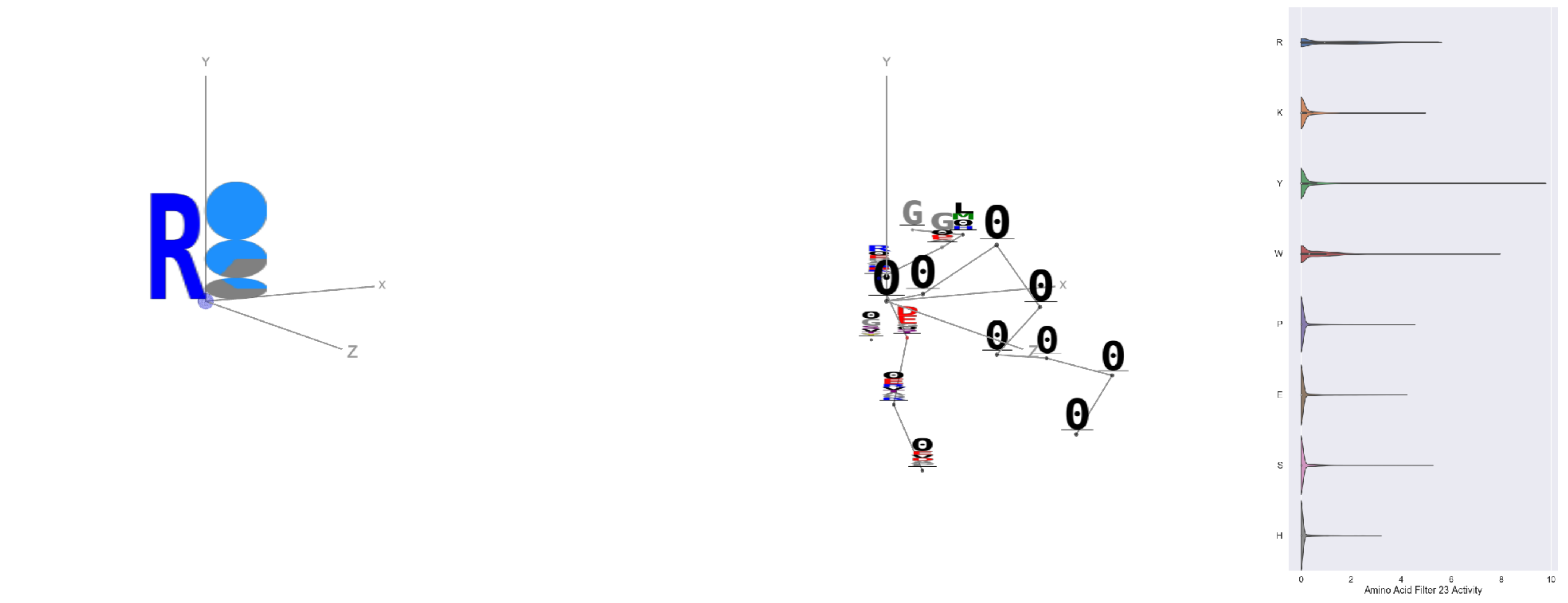

Filter 23
$R_{\text{pred}} = 0.16$
Percent Active = 23
Pooling attention 0.19
Pooling feature 0.27
Top activators
1o7g:A:448 Y
2v6e:A:535 Y
2iwh:B:977 W
2iut:A:723 Y
4g1m:B:692 Y
1hfe:L:398 Y
1lqb:C:210 Y
1f75:A:242 R
4v8p:BC:205 R
3g8q:A:278 R

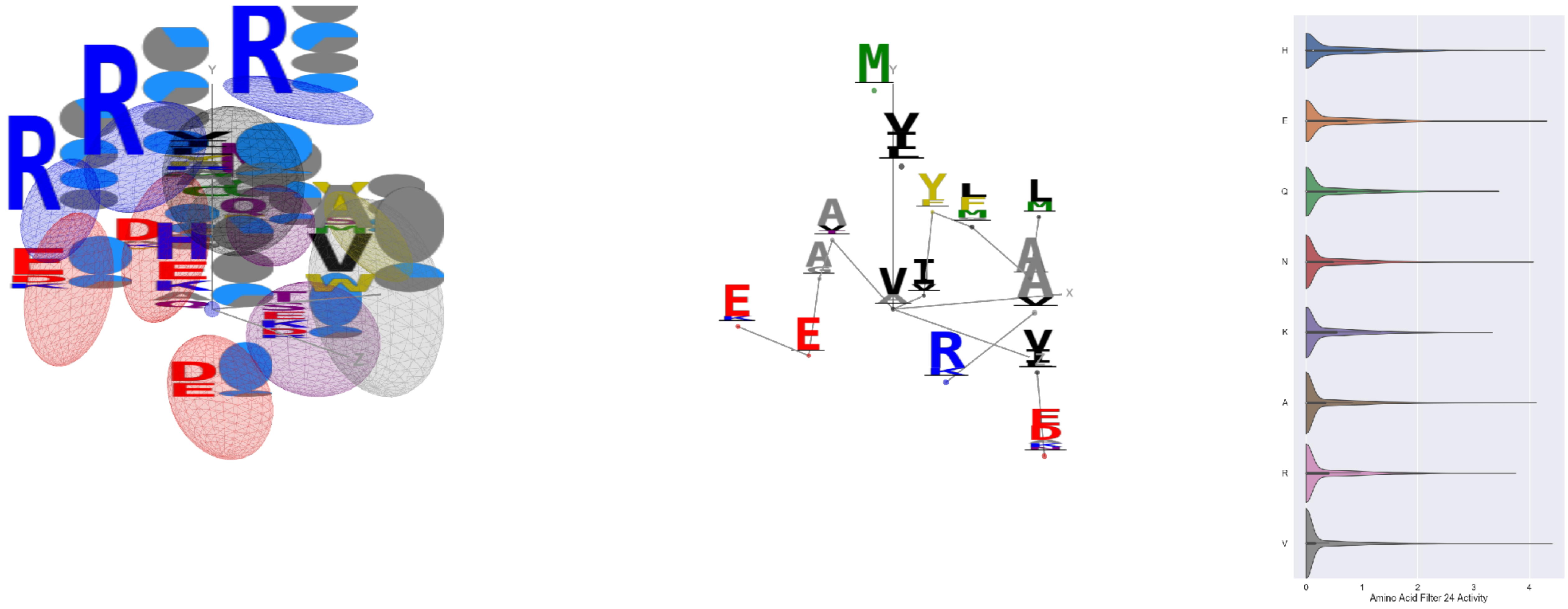


Filter 24
$R_{\text{pred}} = -0.04$
Percent Active = 33
Pooling attention 0.43
Pooling feature 1.93
Top activators
1e1c:A:180 V
3tsy:A:189 D
1iri:A:94 E
2iwh:B:975 H
3g8q:A:35 V
5ui2:A:182 T
3tsy:A:675 A
1mj2:A:46 N
3g8q:A:200 D
3g8q:A:155 A

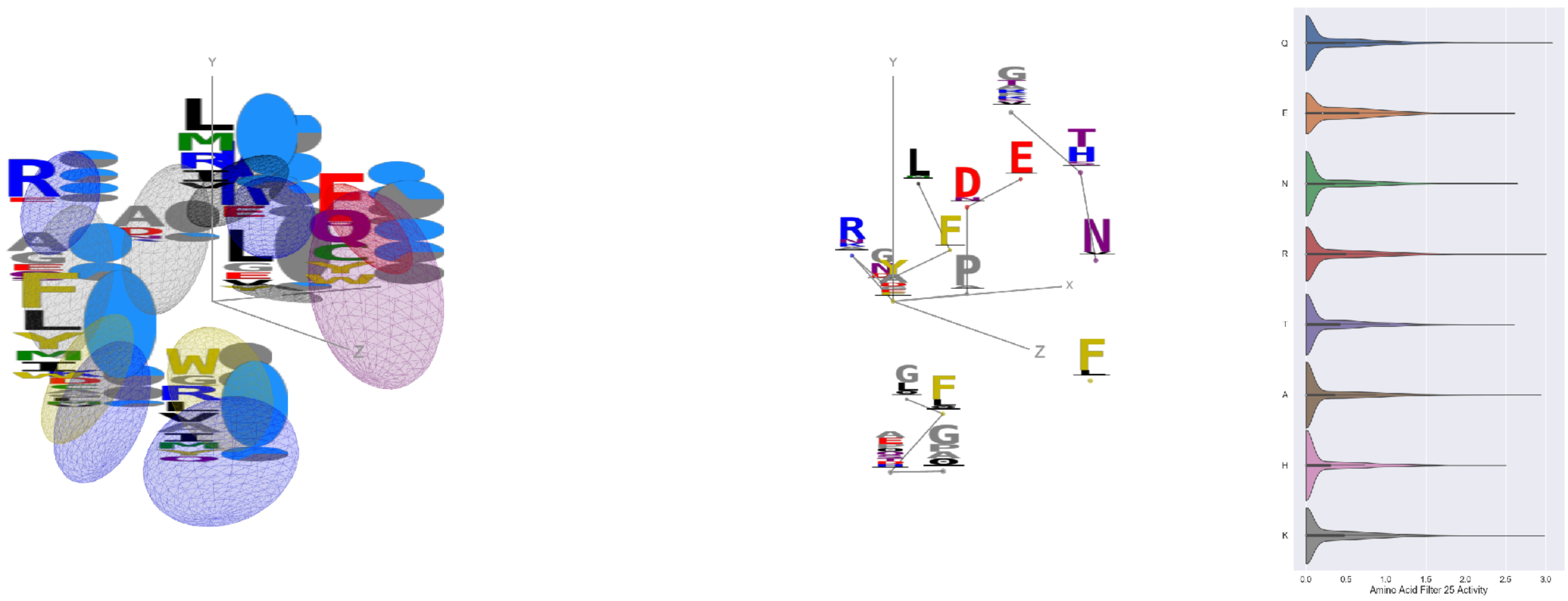


Filter 25
$R_{\text{pred}} = 0.06$
Percent Active = 39
Pooling attention 0.50
Pooling feature 3.37
Top activators
3n6r:G:406 Y
1ivu:A:119 Q
1w36:B:138 R
2iwh:B:186 K
1izl:D:133 A
3l09:B:235 A
4r3z:C:500 L
3qzy:B:86 I
3l0o:A:149 D
1ppt:A:17 L

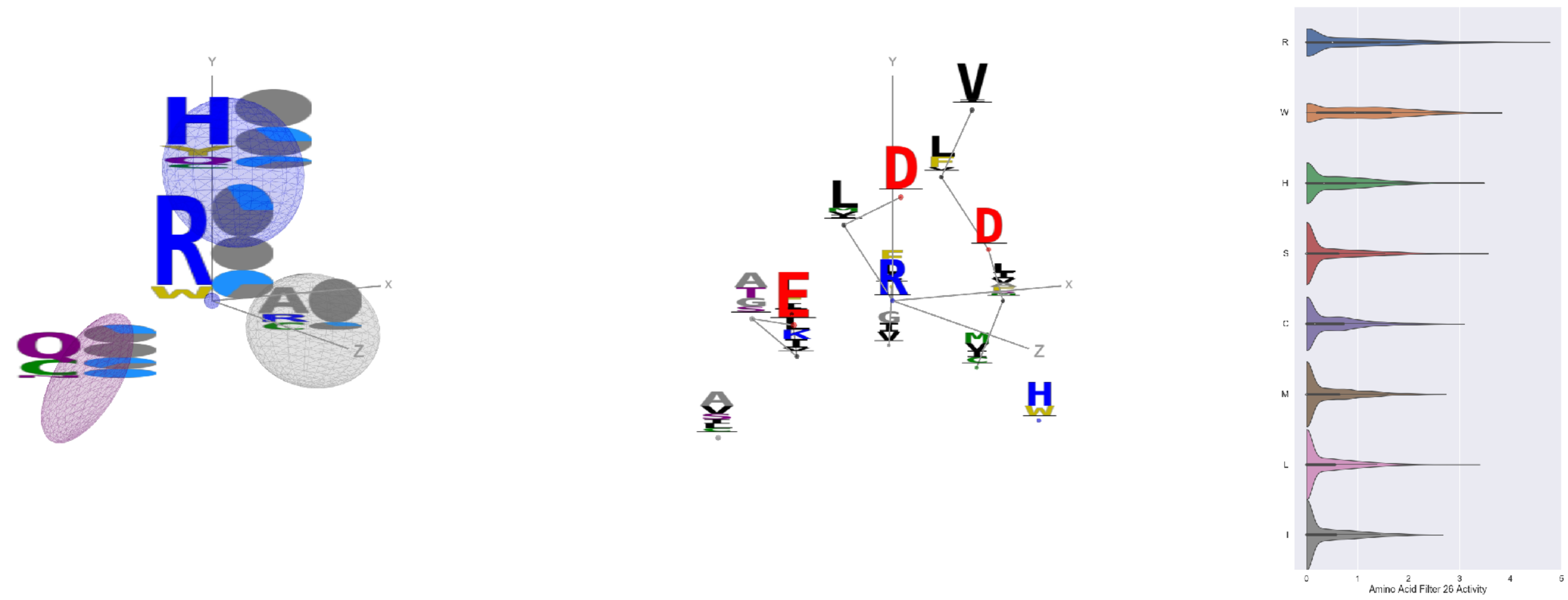


Filter 26
$R_{\text{pred}} = -0.18$
Percent Active = 35
Pooling attention 0.26
Pooling feature 0.25
Top activators
3ucq:A:282 R
3noy:B:101 R
1k90:C:329 R
5ui2:A:291 R
1k72:B:375 R
2iwh:B:457 R
4cvw:A:471 R
1iwa:A:41 R
4gu0:A:311 R
1cm5:A:176 R

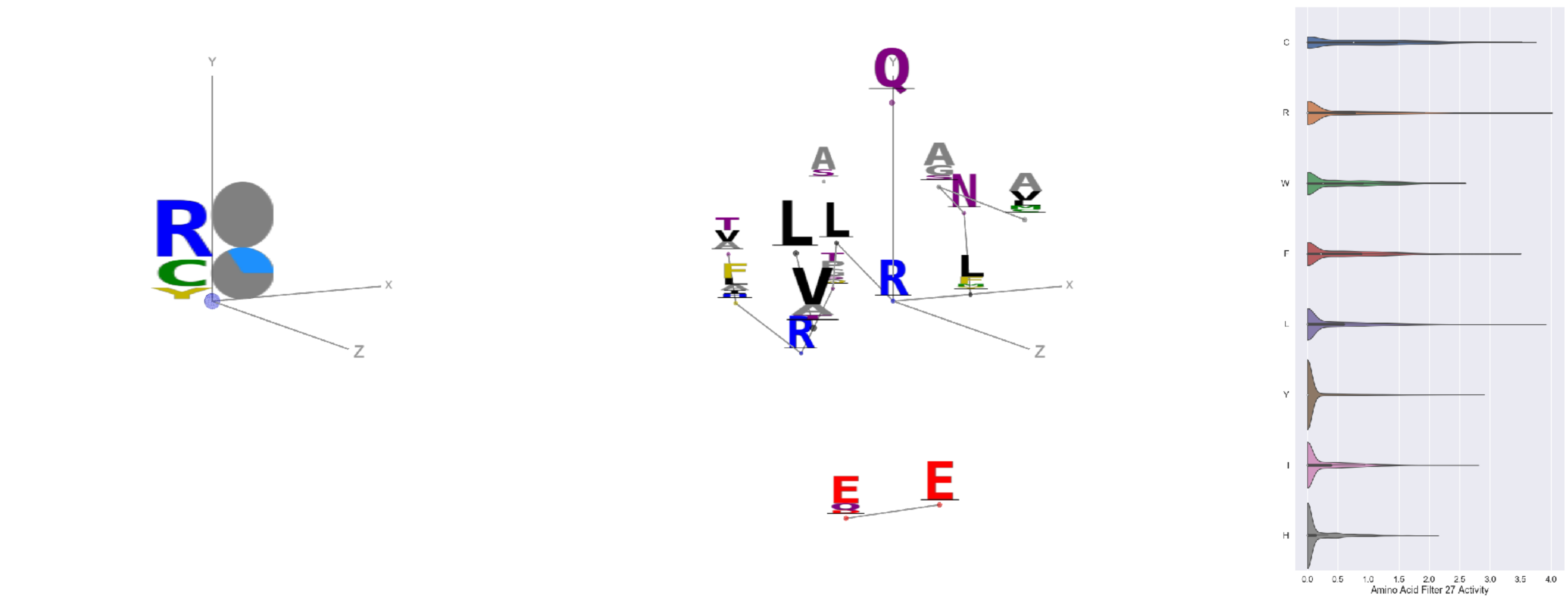


Filter 27
$R_{\text{pred}} = -0.16$
Percent Active = 24
Pooling attention 0.02
Pooling feature 0.82
Top activators
2np9:A:332 R
2iwh:B:452 L
2np9:A:254 R
1e1c:A:302 R
3tsy:A:291 C
3n6r:G:288 R
1bxr:A:715 R
2b0o:F:464 C
2d73:A:642 R
1jku:A:177 R

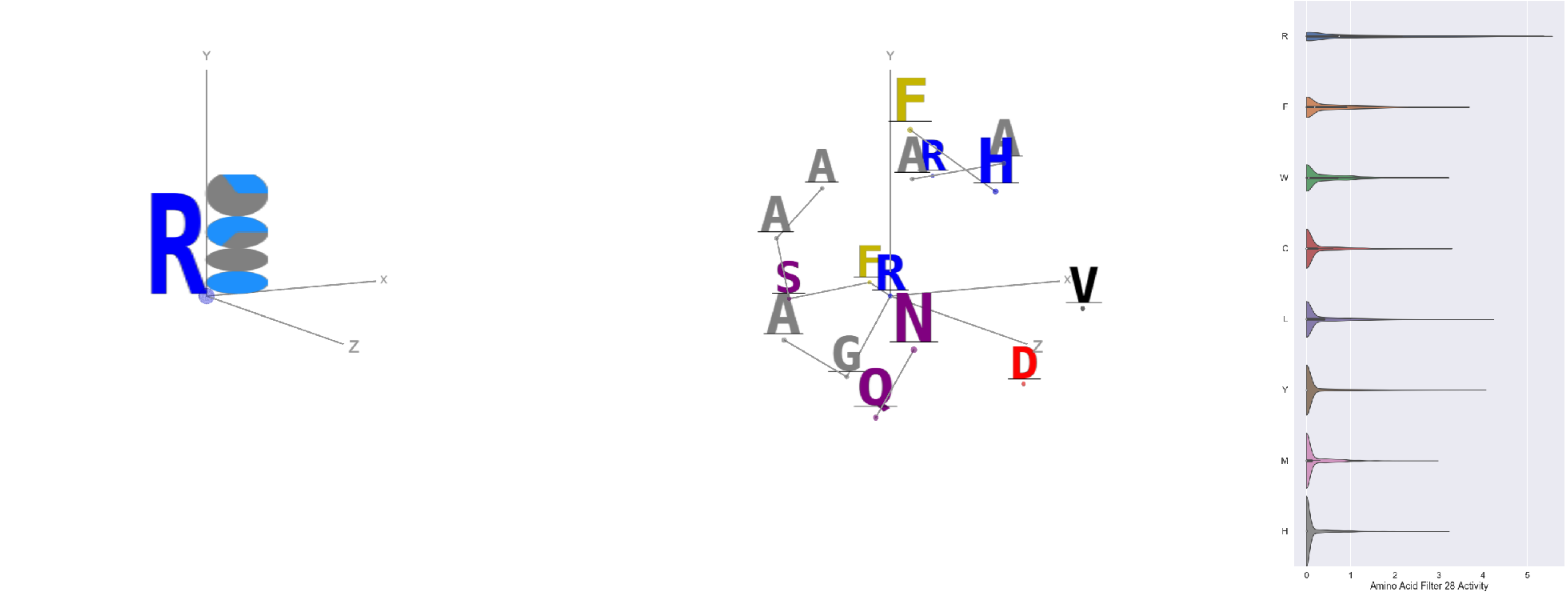


Filter 28
$R_{\text{pred}} = 0.12$
Percent Active = 21
Pooling attention 0.30
Pooling feature 0.32
Top activators
3tsy:A:136 R
3wrx:C:968 R
1r6m:A:7 R
3wb2:A:104 R
3tsy:A:945 R
3g8q:A:251 R
1v8f:A:117 R
3o47:A:281 R
4pht:A:320 R
3bf0:C:496 R

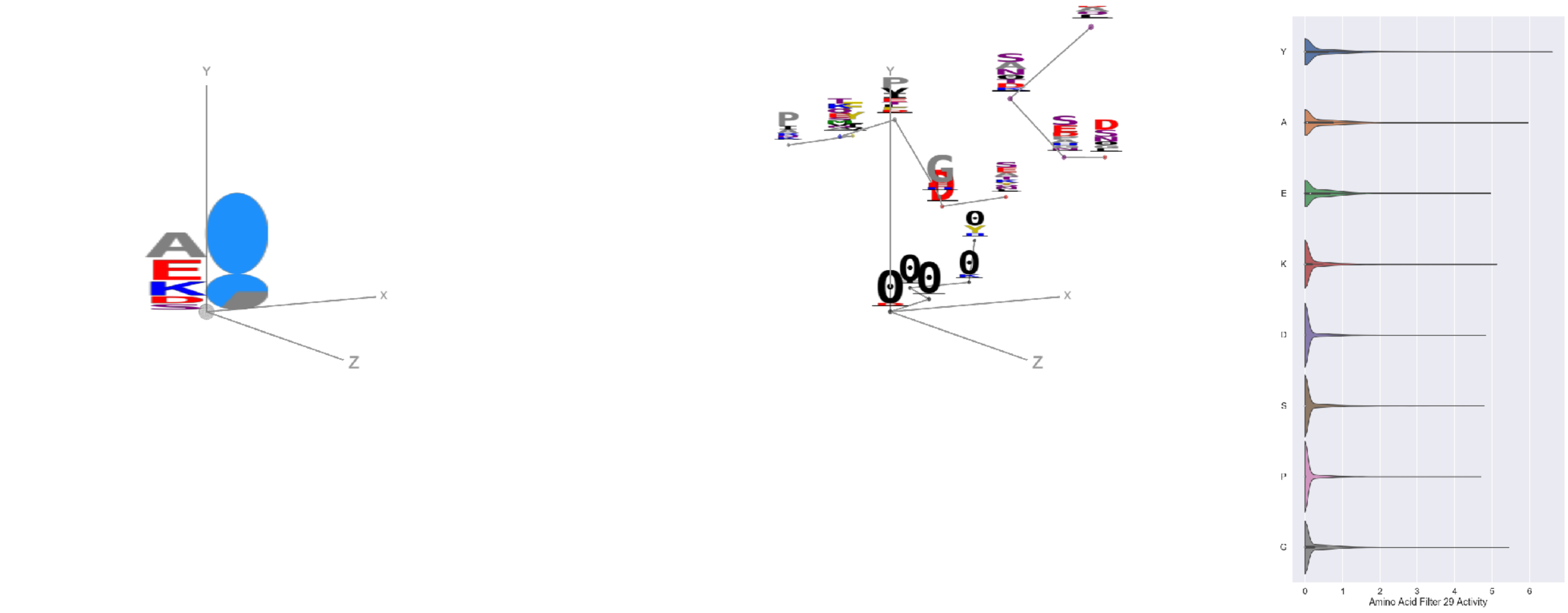


Filter 29
$R_{\text{pred}} = -0.01$
Percent Active = 27
Pooling attention 0.12
Pooling feature 0.75
Top activators
2yma:A:399 Y
3bf0:C:551 Y
4rg1:A:374 Y
2dyo:A:285 Y
3qzy:B:260 Y
2fq1:A:283 Y
4pht:A:497 Y
2hye:B:54 A
4ny2:A:433 Y
1c3r:A:373 Y

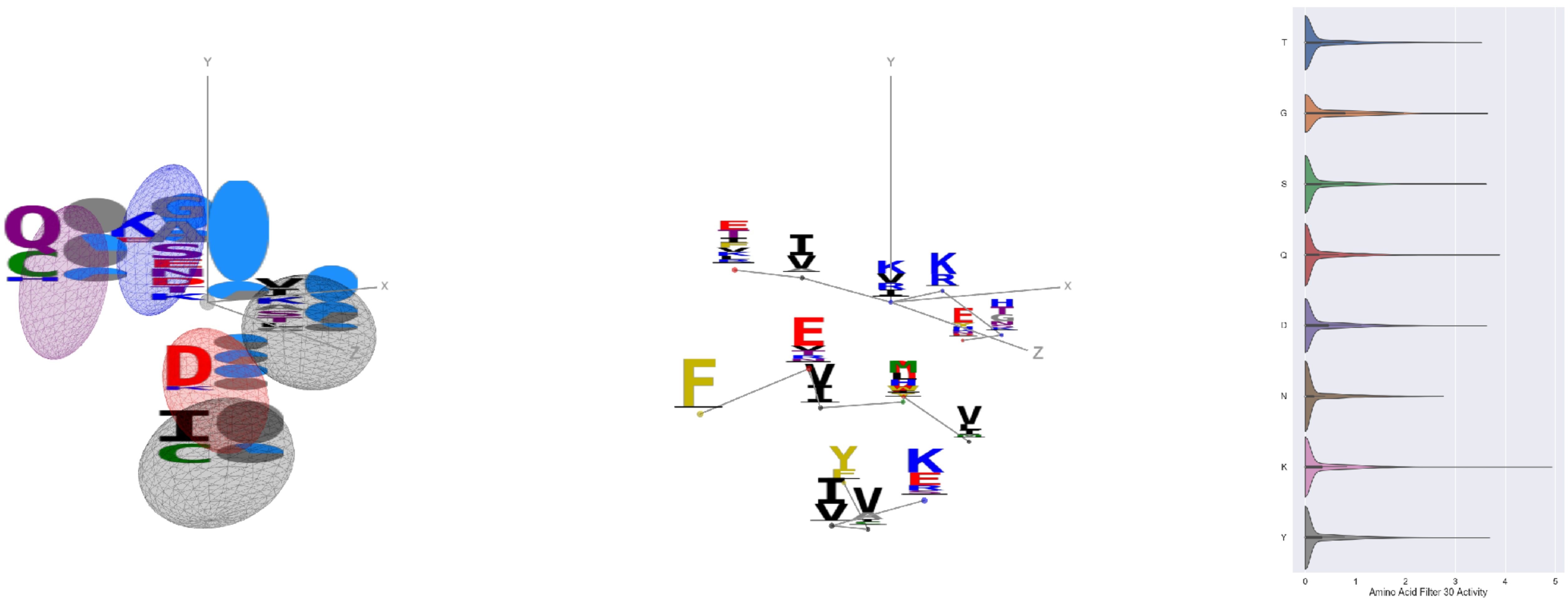


Filter 30
$R_{\text{pred}} = 0.16$
Percent Active = 29
Pooling attention 0.43
Pooling feature 0.84
Top activators
1umg:A:178 K
4rcn:B:524 E
3pio:M:47 I
1i3r:B:137 H
4dag:A:402 Q
2pg1:L:133 E
4u4c:A:952 Y
3tsy:A:182 G
1v8f:A:146 D
3s9m:C:280 S

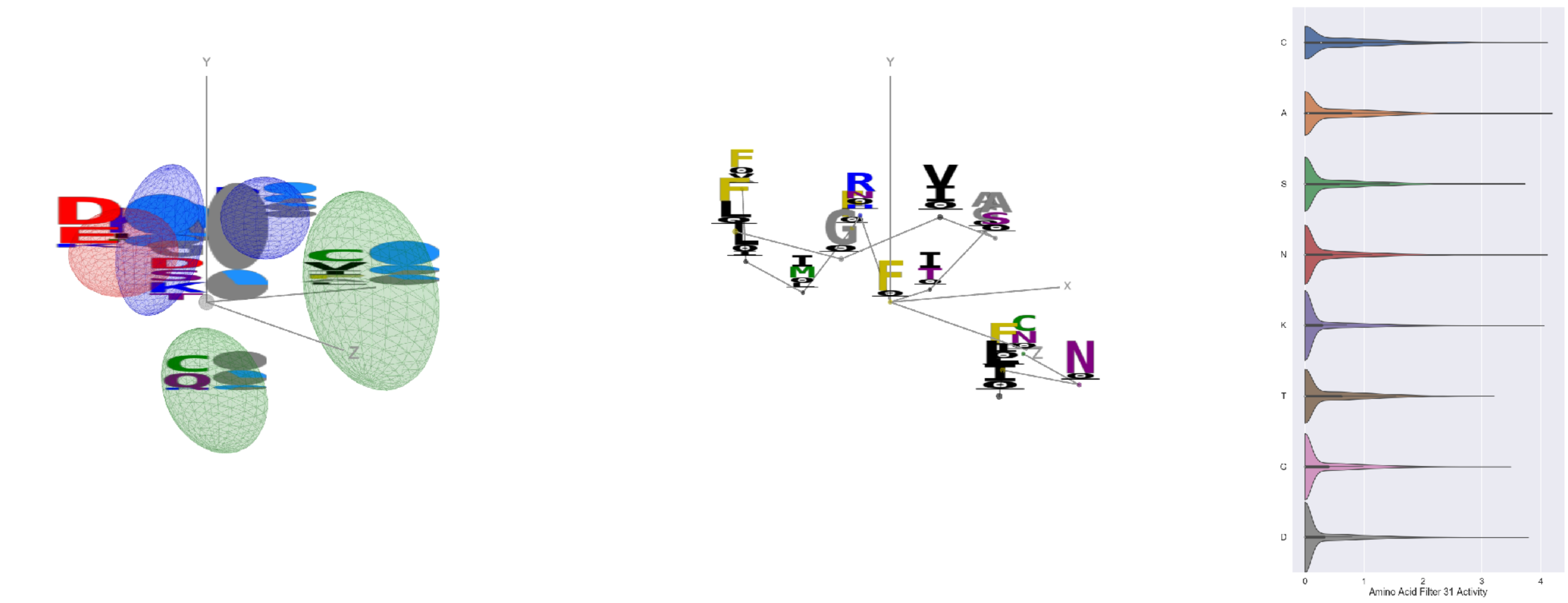


Filter 31
$R_{\text{pred}} = -0.18$
Percent Active = 35
Pooling attention 0.34
Pooling feature 1.28
Top activators
1izl:D:179 F
1izl:D:64 A
1r7r:A:728 Y
3d00:A:165 C
1izl:D:194 N
3s9m:C:233 K
1r7r:A:629 I
1fx0:B:329 V
2iut:A:498 D
1izl:D:338 Y

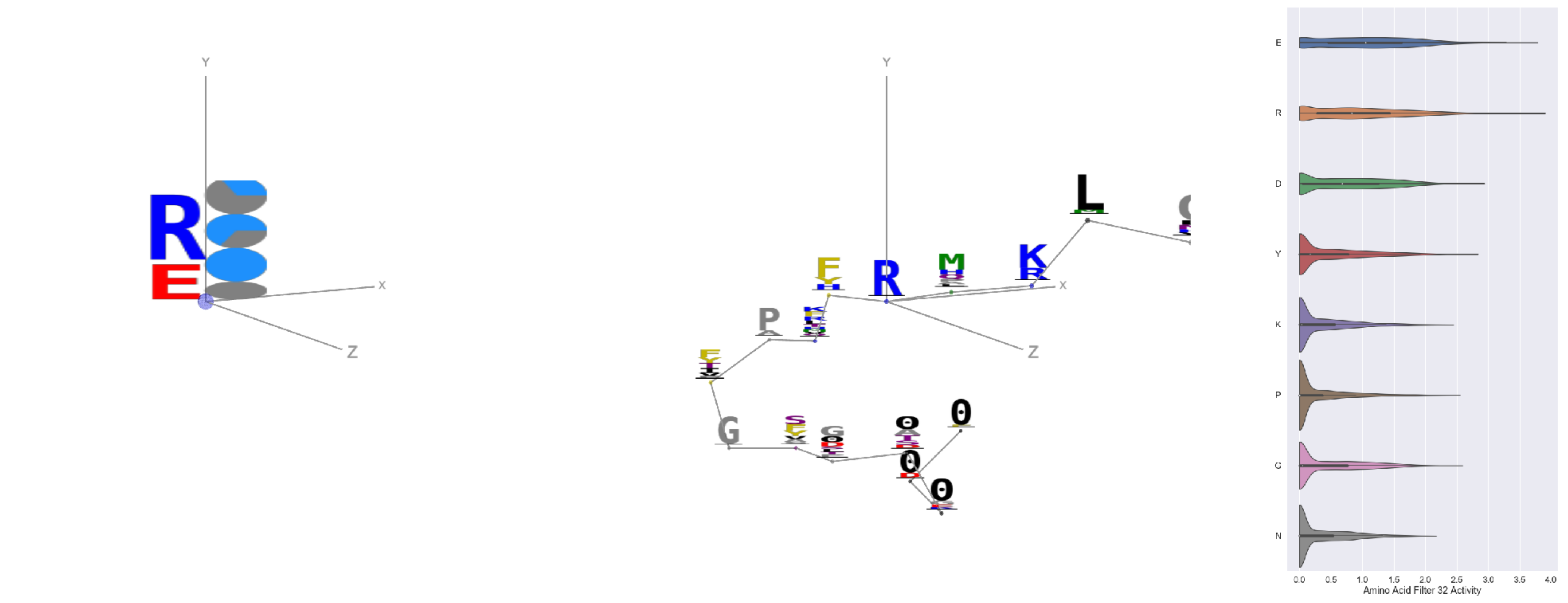


Filter 32
$R_{\text{pred}} = 0.11$
Percent Active = 44
Pooling attention 0.22
Pooling feature 0.57
Top activators
4c2m:N:169 R
1kf6:C:29 R
3tsy:A:452 E
3tsy:A:944 E
3g8q:A:252 E
1j5w:B:112 E
3g8q:A:207 R
4uzi:A:332 R
3t5v:C:36 E
4ehi:B:118 E

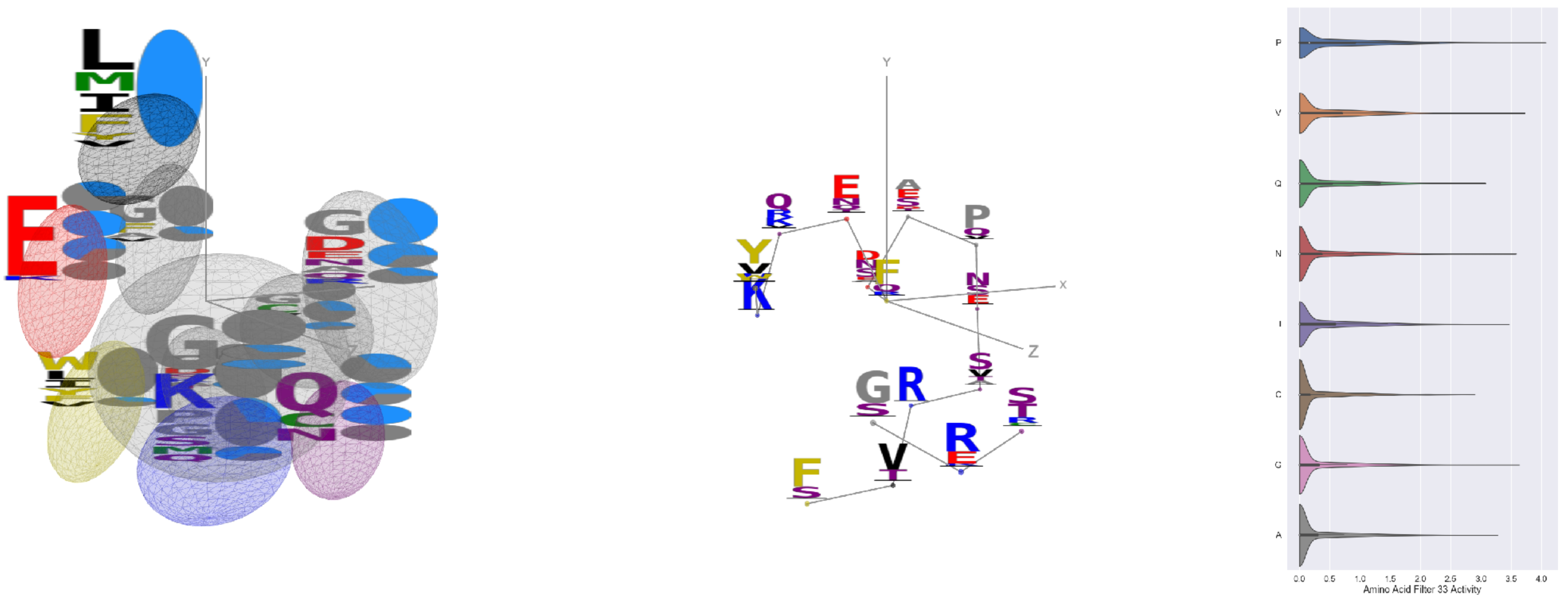


Filter 33
$R_{\text{pred}} = -0.06$
Percent Active = 34
Pooling attention 0.45
Pooling feature 1.96
Top activators
3jsb:A:111 F
1y4j:B:166 E
3tsy:A:815 P
3l0o:A:148 P
4r3z:C:514 P
1k90:C:380 V
1cmx:A:188 P
1is7:K:53 P
4kbj:B:164 G
1zt2:A:83 P

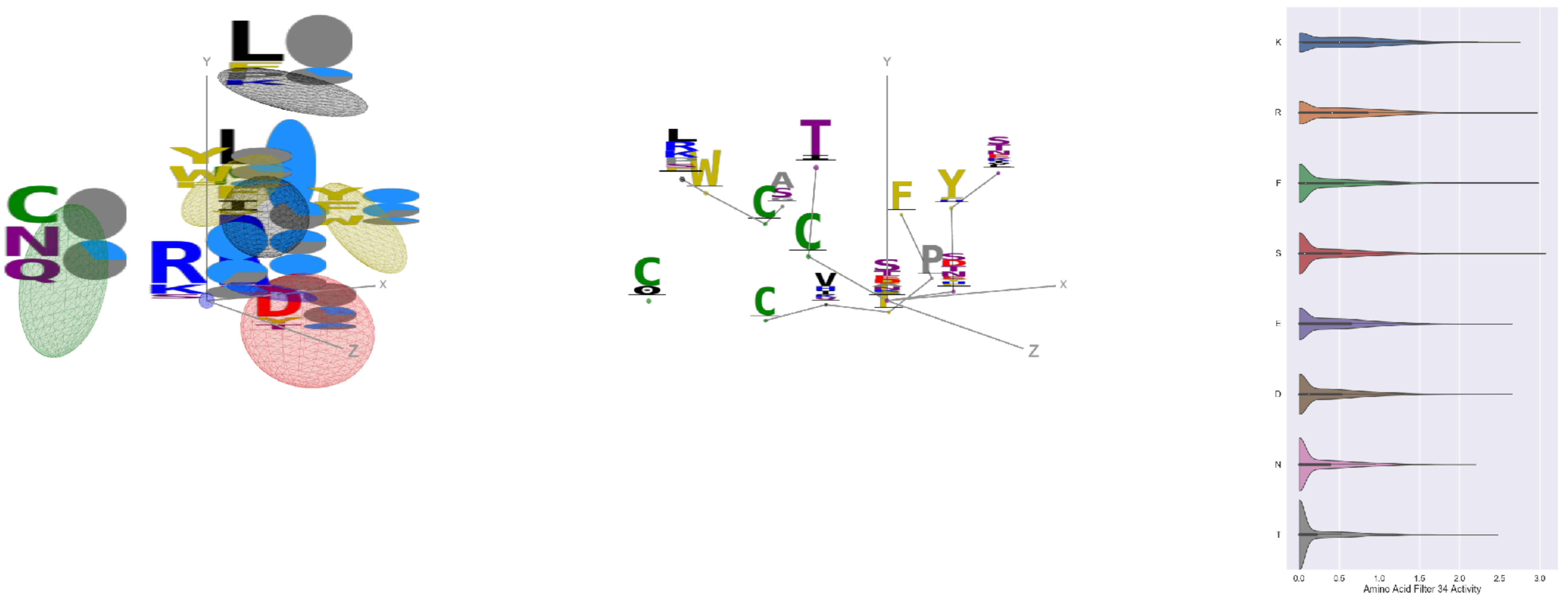


Filter 34
$R_{\text{pred}} = 0.05$
Percent Active = 36
Pooling attention 0.05
Pooling feature 0.70
Top activators
1h8p:A:82 S
4a7z:A:211 F
1rq0:A:237 R
4e0s:B:567 R
1j5w:B:117 S
2o35:B:41 F
3vie:B:122 R
1v2z:A:278 S
4jdm:C:230 S
1h8p:A:57 K

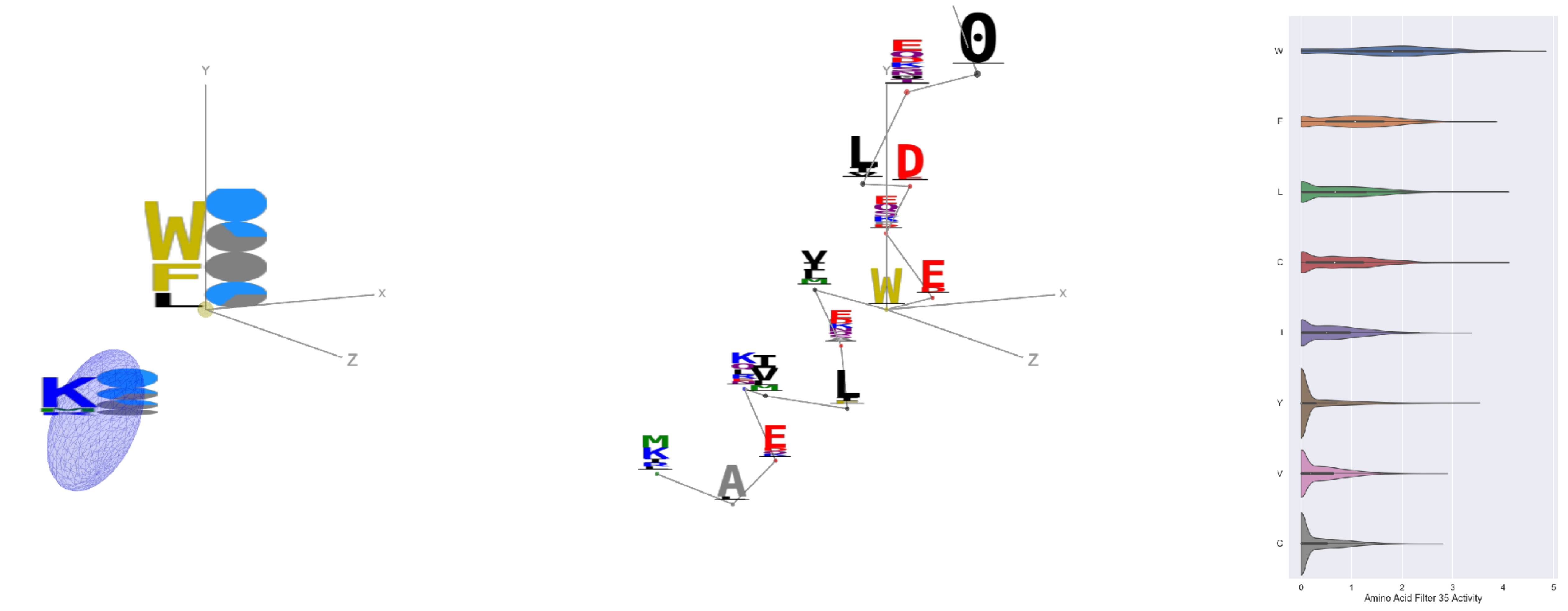


Filter 35
$R_{\text{pred}} = 0.15$
Percent Active = 34
Pooling attention 0.11
Pooling feature 0.15
Top activators
2yal:A:78 W
4c0b:D:482 W
4ny2:A:370 W
3vie:B:117 W
1mu2:A:266 W
2wsc:B:60 W
2iwh:B:878 C
1iri:A:162 L
4r3z:C:511 W
1a92:A:20 W

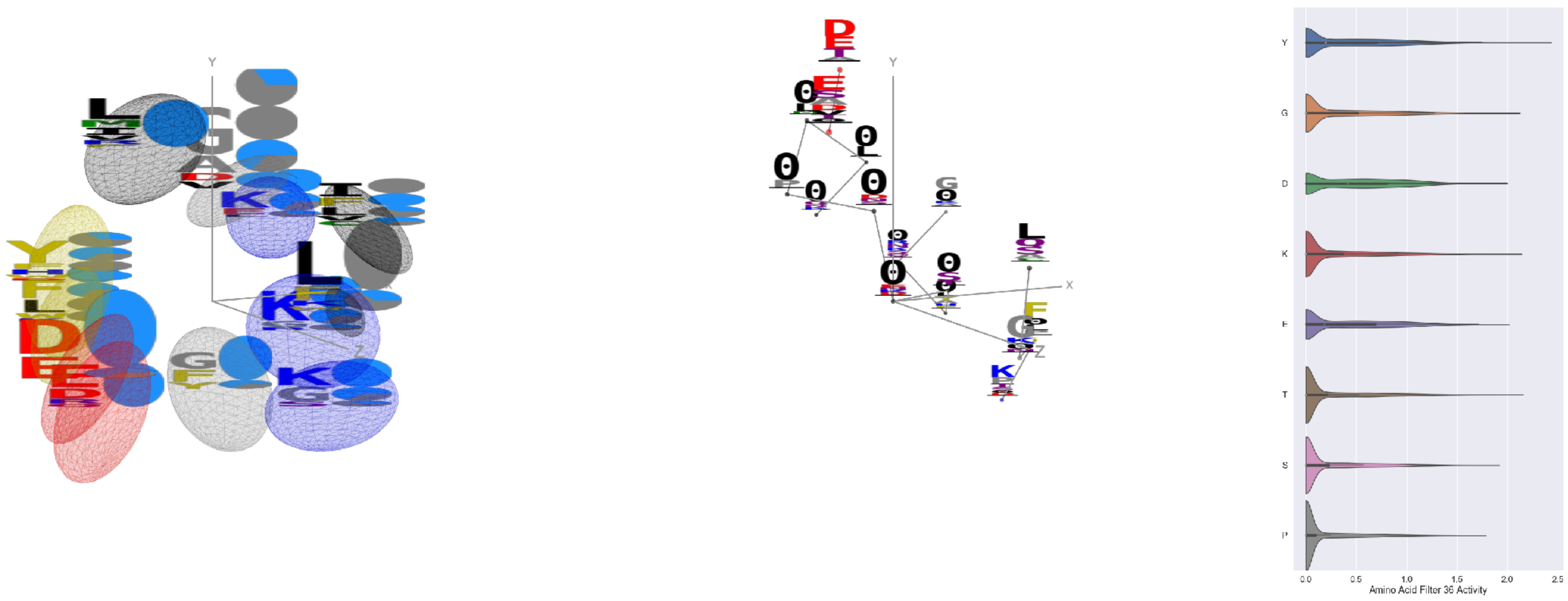

Filter 36
$R_{\text{pred}} = -0.07$
Percent Active = 36
Pooling attention 0.25
Pooling feature 0.22
Top activators
2hb6:A:485 Y
4cy6:A:209 R
2o3o:A:248 V
3hfd:A:161 T
4v81:H:151 K
1lk5:A:49 Y
2prv:A:82 G
4gt2:A:330 Y
3ne5:C:100 Y
1sl0:A:291 Y

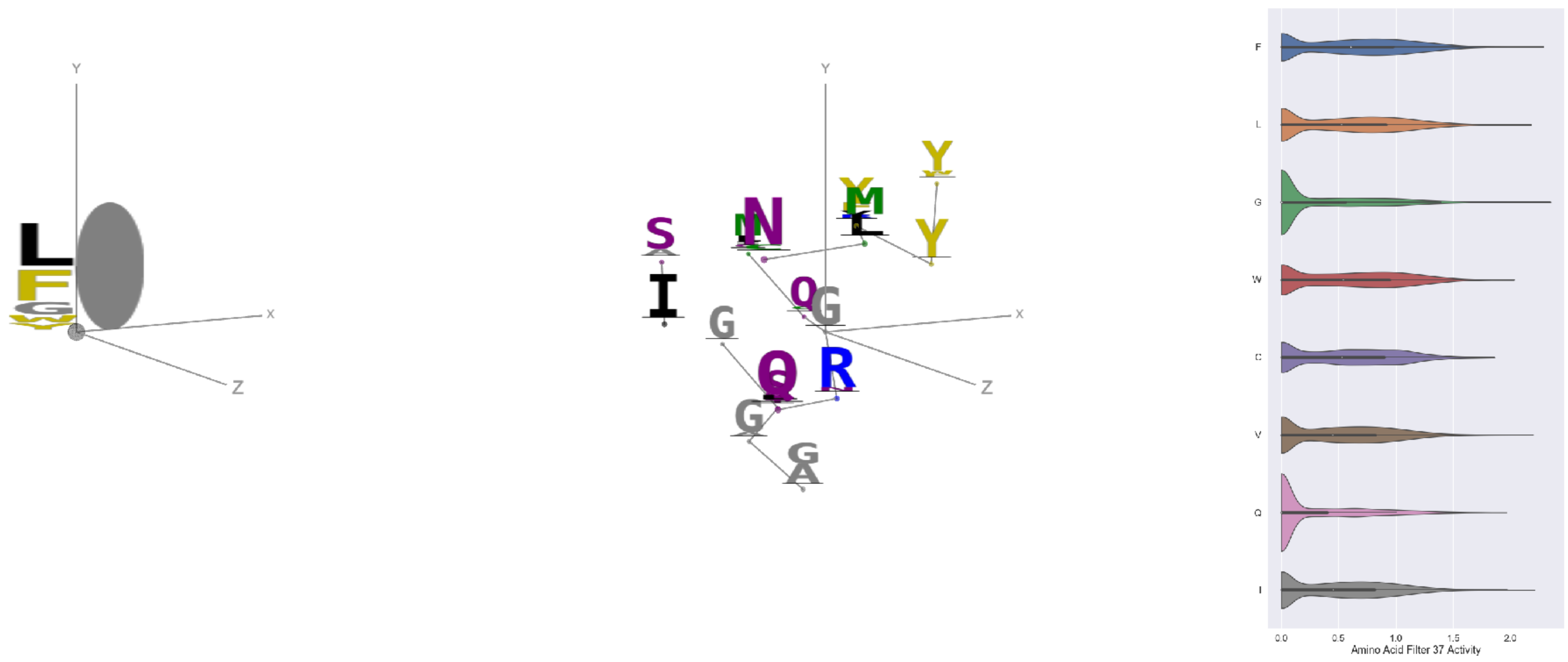

Filter 37
$R_{\text{pred}} = -0.32$
Percent Active = 44
Pooling attention 0.14
Pooling feature 0.92
Top activators
3bsd:A:142 G
3o47:A:181 F
4r3z:C:521 G
4r3z:C:747 G
1qvr:A:185 I
2cge:A:345 V
1f4a:A:992 G
1r7r:A:357 L
1cm5:A:433 G
3g8q:A:92 G

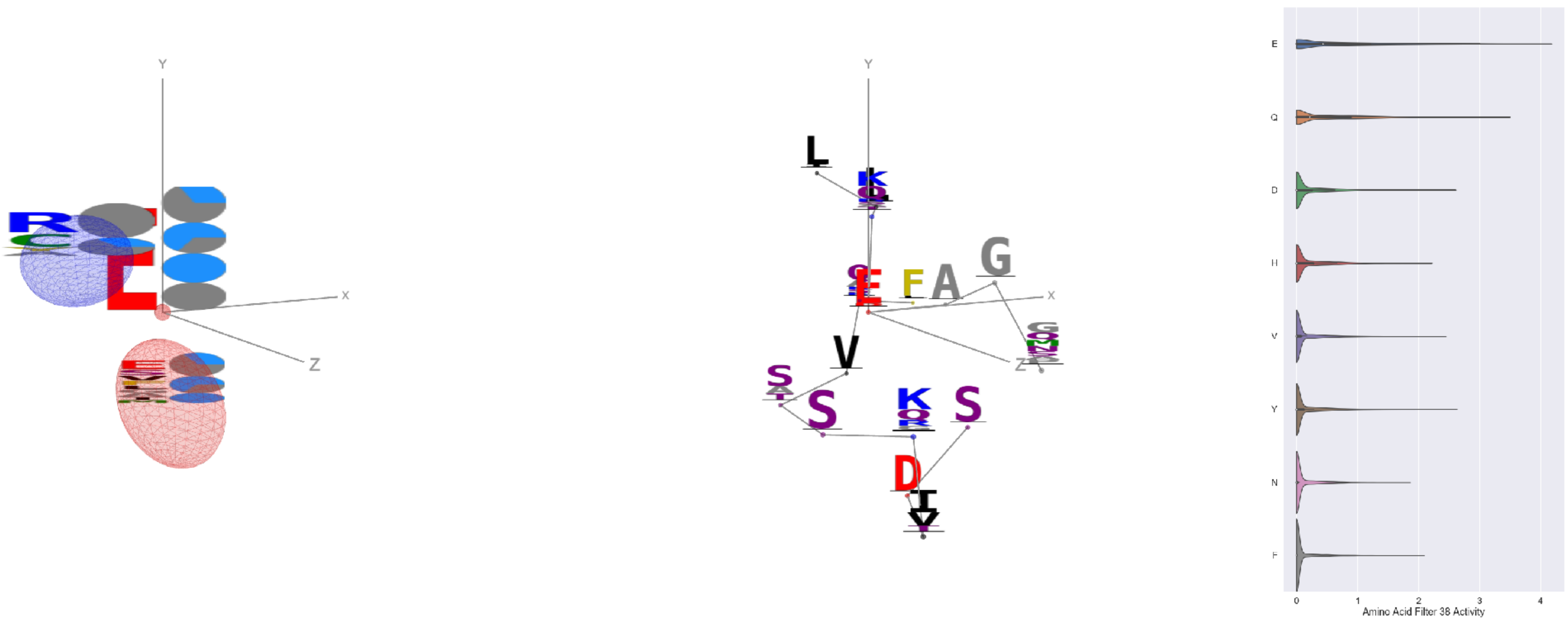

Filter 38
$R_{\text{pred}} = 0.01$
Percent Active = 23
Pooling attention 0.05
Pooling feature 0.66
Top activators
2iut:A:670 E
3g8q:A:46 E
1iwa:A:60 E
1e1c:A:203 E
1bxr:A:403 E
1fx0:B:439 E
3ax7:A:89 E
1k72:B:132 E
1f4a:A:198 E
3l0o:A:259 E

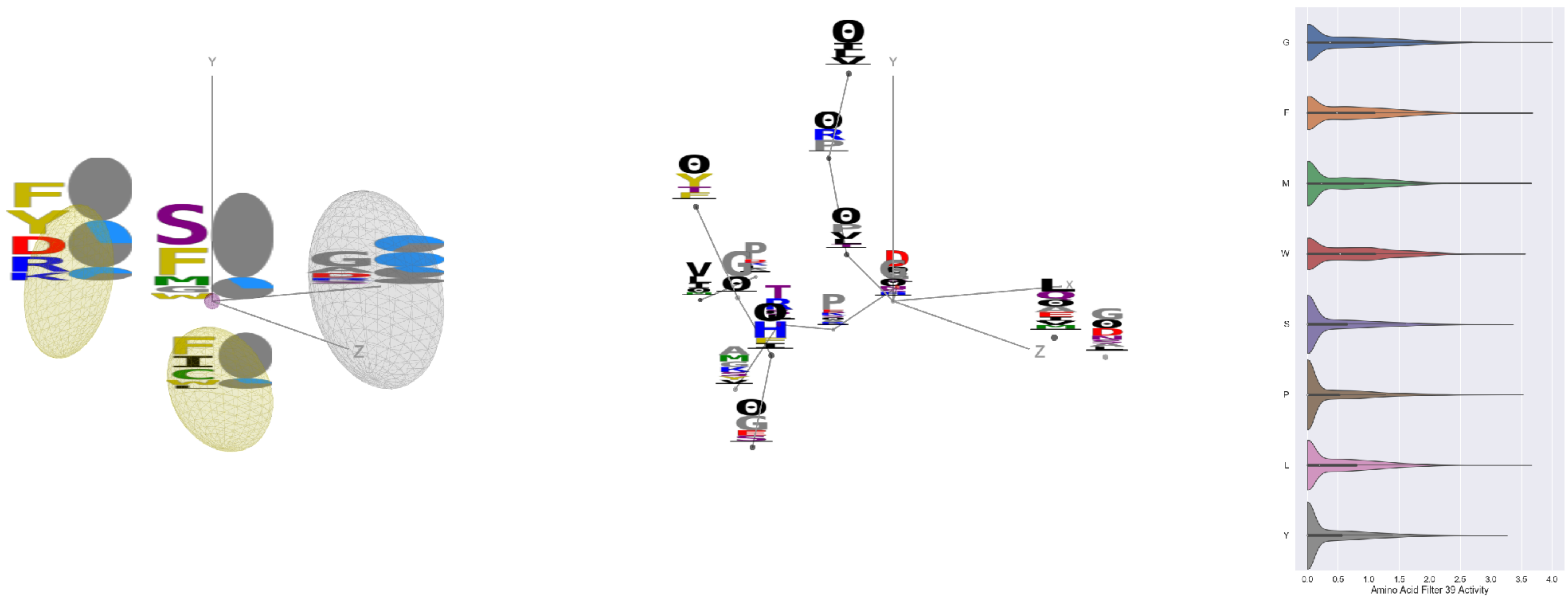

Filter 39
$R_{\text{pred}} = -0.19$
Percent Active = 45
Pooling attention 0.30
Pooling feature 0.34
Top activators
1c0i:A:1291 G
1e5d:A:232 K
1w99:A:398 F
1c0i:A:1094 L
1jdw:A:302 M
4u9c:B:222 D
2x98:A:301 G
2d73:A:331 W
2x3l:B:24 P
5ui2:A:161 M

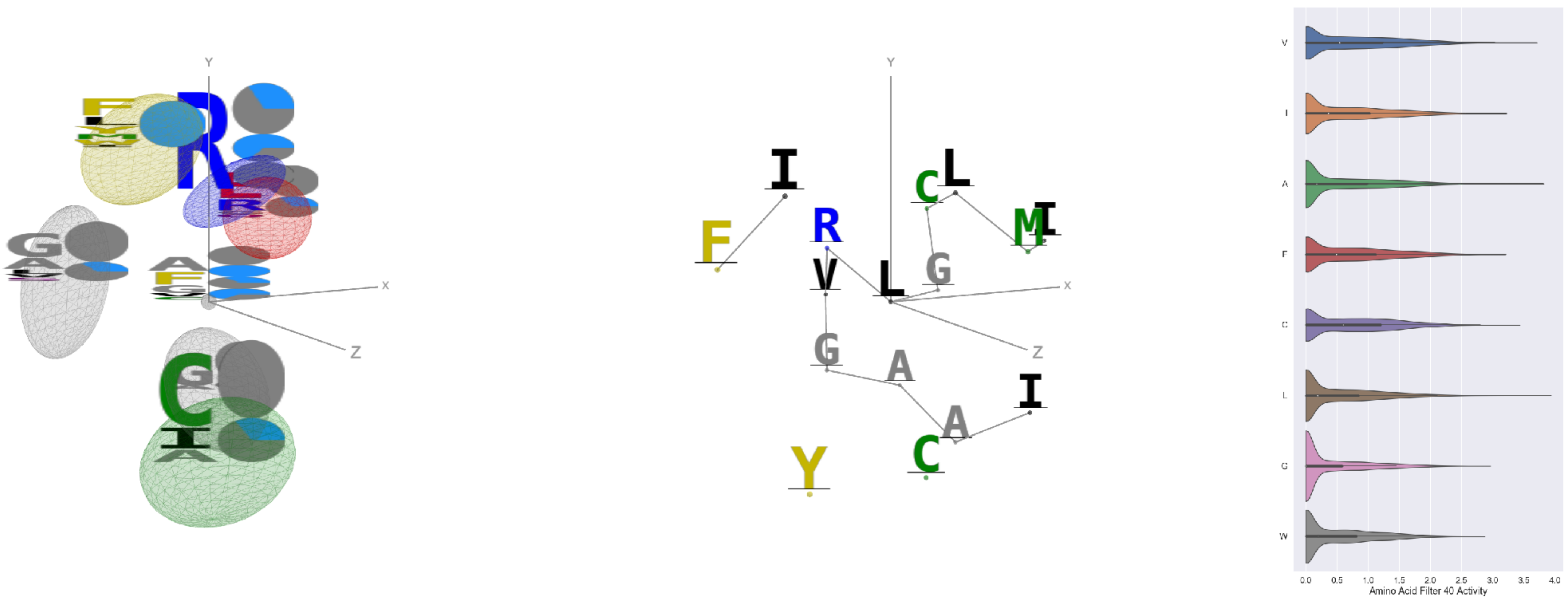

Filter 40
$R_{\text{pred}} = -0.25$
Percent Active = 38
Pooling attention 0.11
Pooling feature 0.24
Top activators
3tsy:A:293 L
1cm5:A:417 A
5ui2:A:271 V
3wr9:A:9 A
3tsy:A:776 V
2zm2:A:277 A
3bof:A:207 C
3bof:A:303 L
4rcn:B:750 V
1r6m:A:134 A

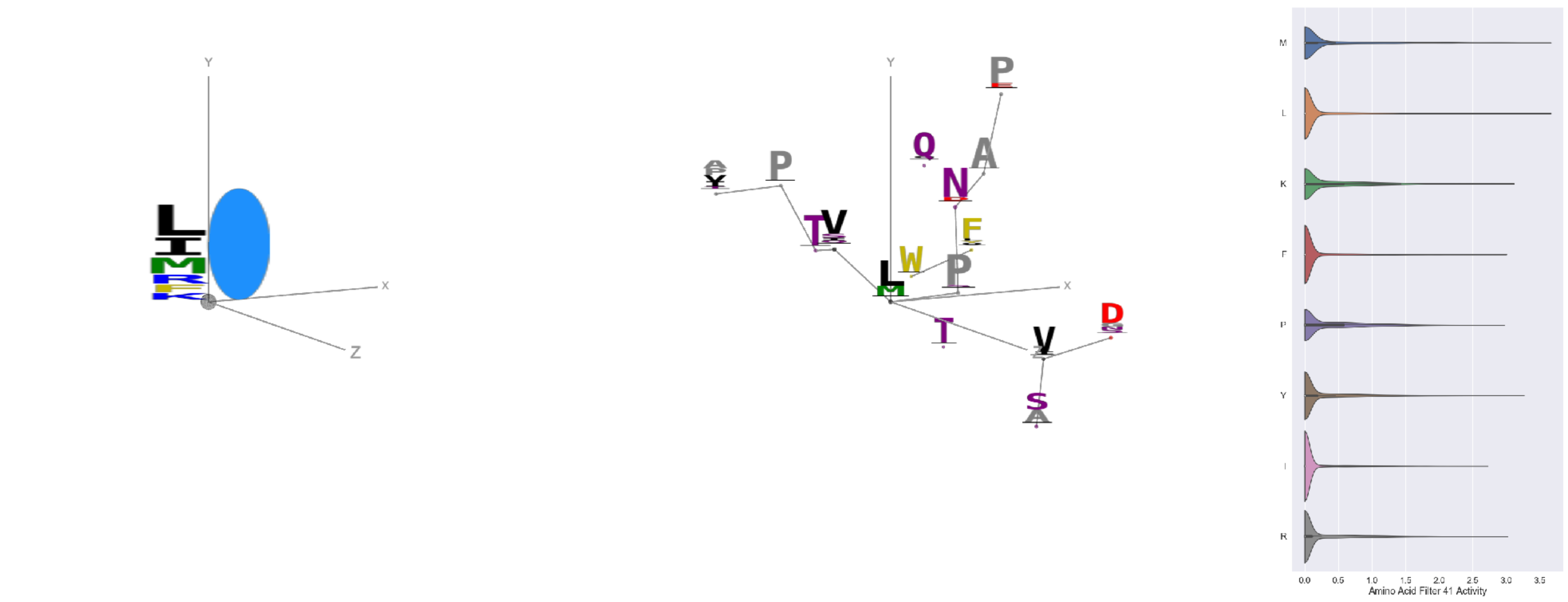

Filter 41
$R_{\text{pred}} = 0.31$
Percent Active = 25
Pooling attention 0.05
Pooling feature 0.16
Top activators
1ezv:C:263 L
3n6r:G:693 M
3tsy:A:724 M
3irm:A:519 M
4jdm:C:245 L
2iwh:B:823 M
1n0e:B:162 Y
4u4c:A:549 L
4c0b:B:239 E
2v6e:A:535 Y

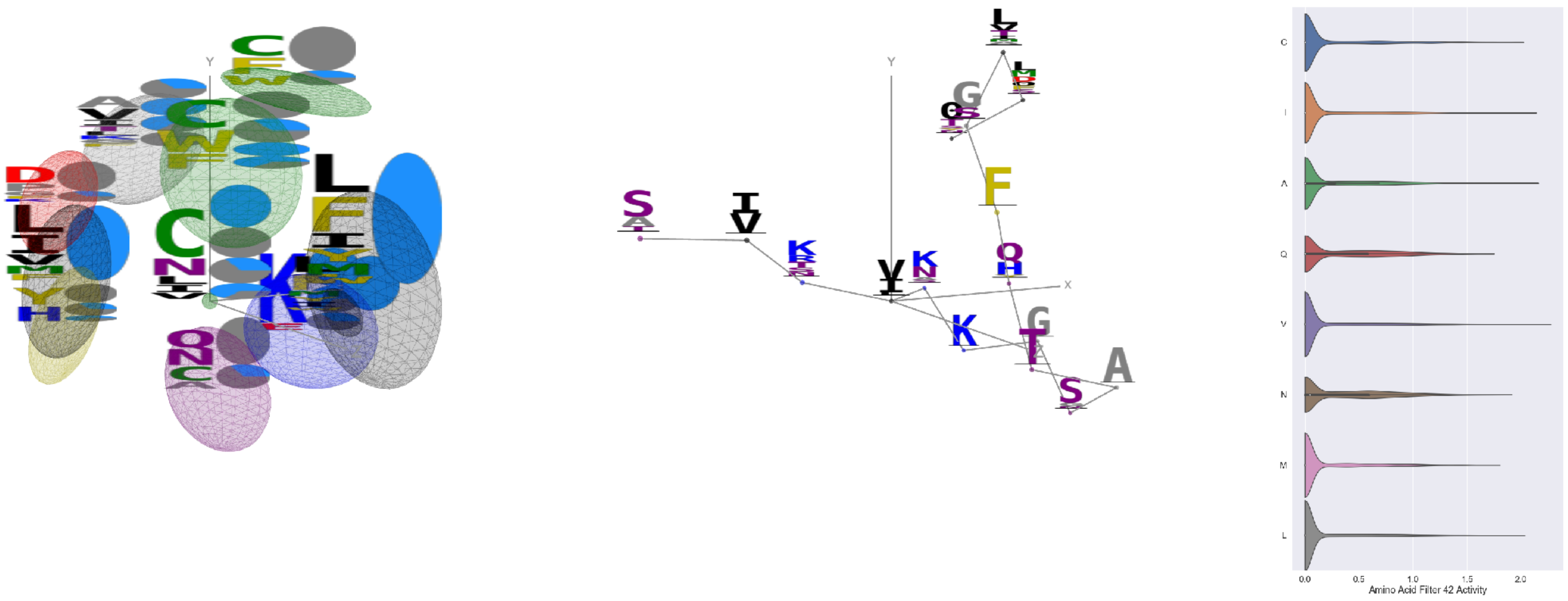

Filter 42
$R_{\text{pred}} = 0.33$
Percent Active = 38
Pooling attention 0.57
Pooling feature 2.55
Top activators
1n0w:B:1532 V
3din:D:50 G
2hd0:A:128 A
3te3:A:724 I
3b8e:A:102 I
2zbk:B:478 I
4hg6:A:24 Y
3vr6:G:169 A
3k6g:D:291 A
3din:E:68 L

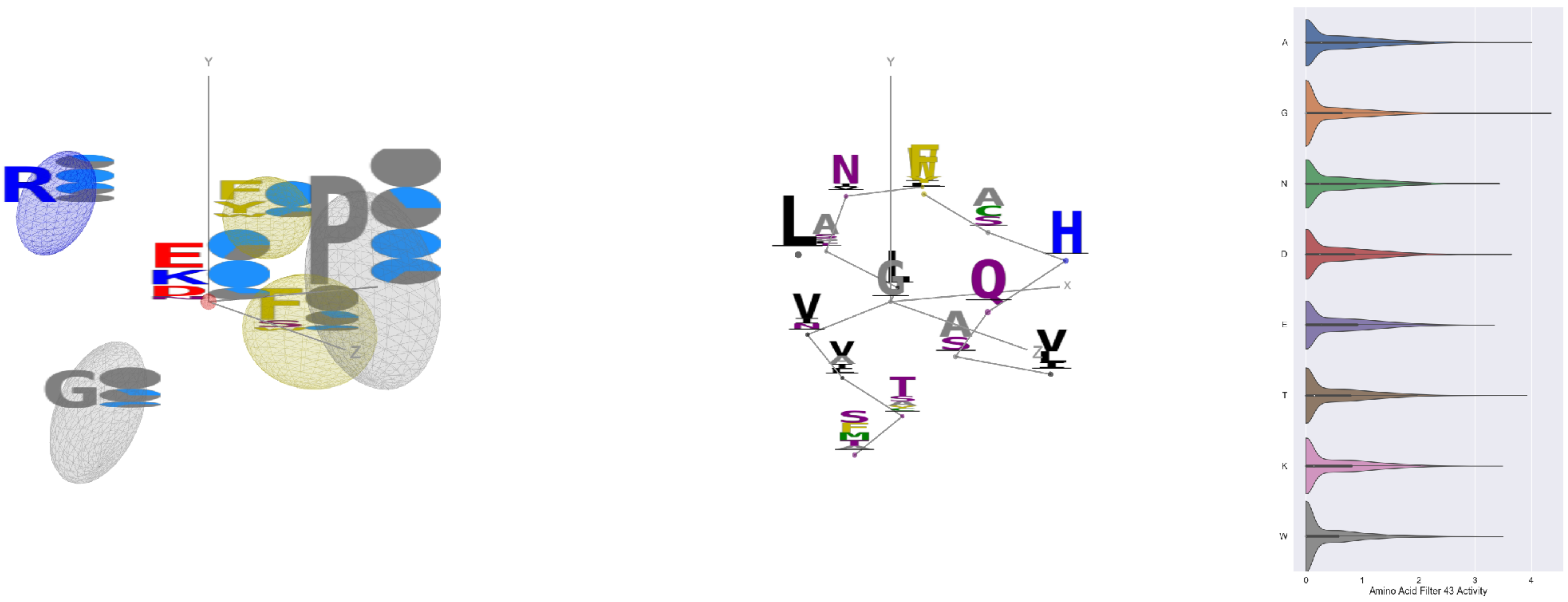

Filter 43
$R_{\text{pred}} = -0.02$
Percent Active = 47
Pooling attention 0.23
Pooling feature 0.82
Top activators
4buj:B:993 G
3kov:A:41 A
1e5d:A:150 T
4c8q:H:782 Y
1he8:A:963 I
3hwp:A:287 A
3tsy:A:477 V
4jdm:C:258 V
3tsy:A:755 G
2v6e:A:535 Y

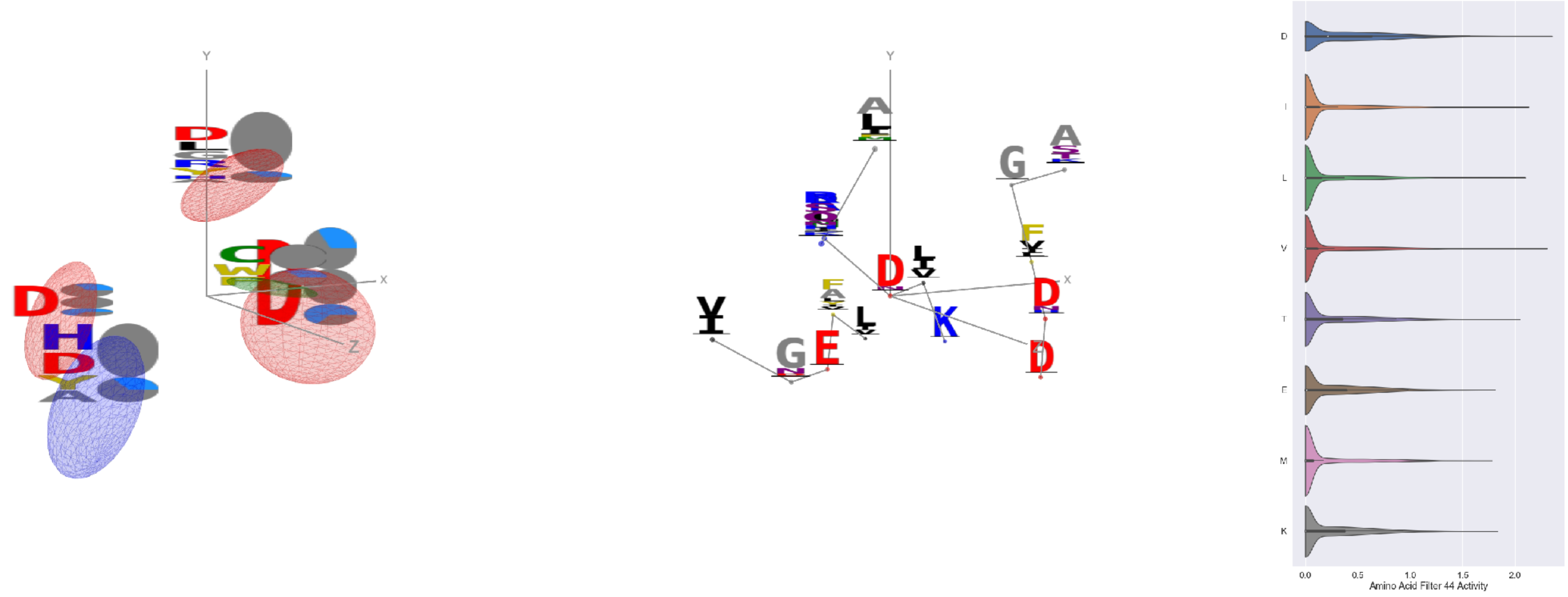

Filter 44
$R_{\mathrm{pred}} = 0.08$
Percent Active = 34
Pooling attention 0.28
Pooling feature 0.28
Top activators
2w27:A:175 D
3tsy:A:228 V
4e0s:B:344 Y
2wgq:B:429 I
1eg1:A:172 D
2iut:A:463 L
1bxr:A:614 D
1e5d:A:75 I
1ii7:A:52 D
4cbv:D:87 T

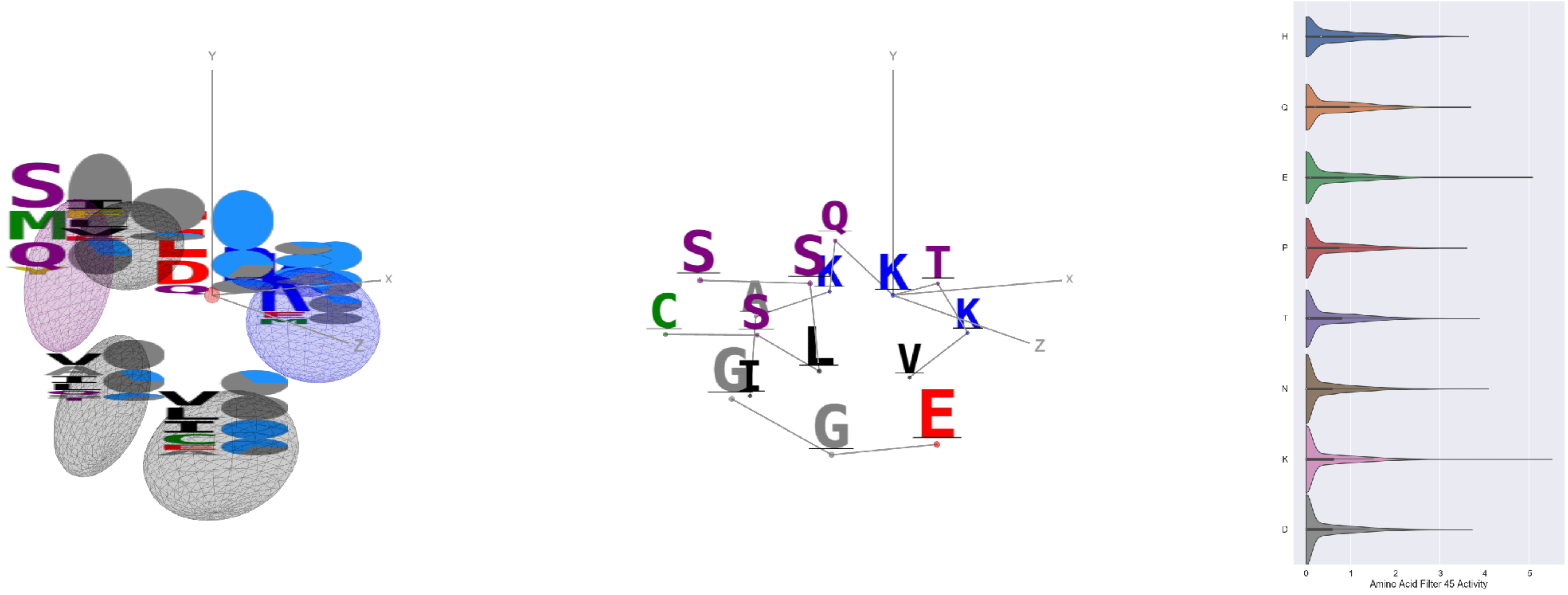

Filter 45
$R_{\mathrm{pred}} = -0.07$
Percent Active = 36
Pooling attention 0.46
Pooling feature 1.13
Top activators
2iwh:B:661 K
3n6r:G:696 E
4rcn:B:524 E
1p4e:B:394 Y
4whj:A:327 N
1f4a:A:487 E
1hfe:L:26 Y
3ucq:A:136 S
2f8x:C:323 E
1p4e:B:229 F

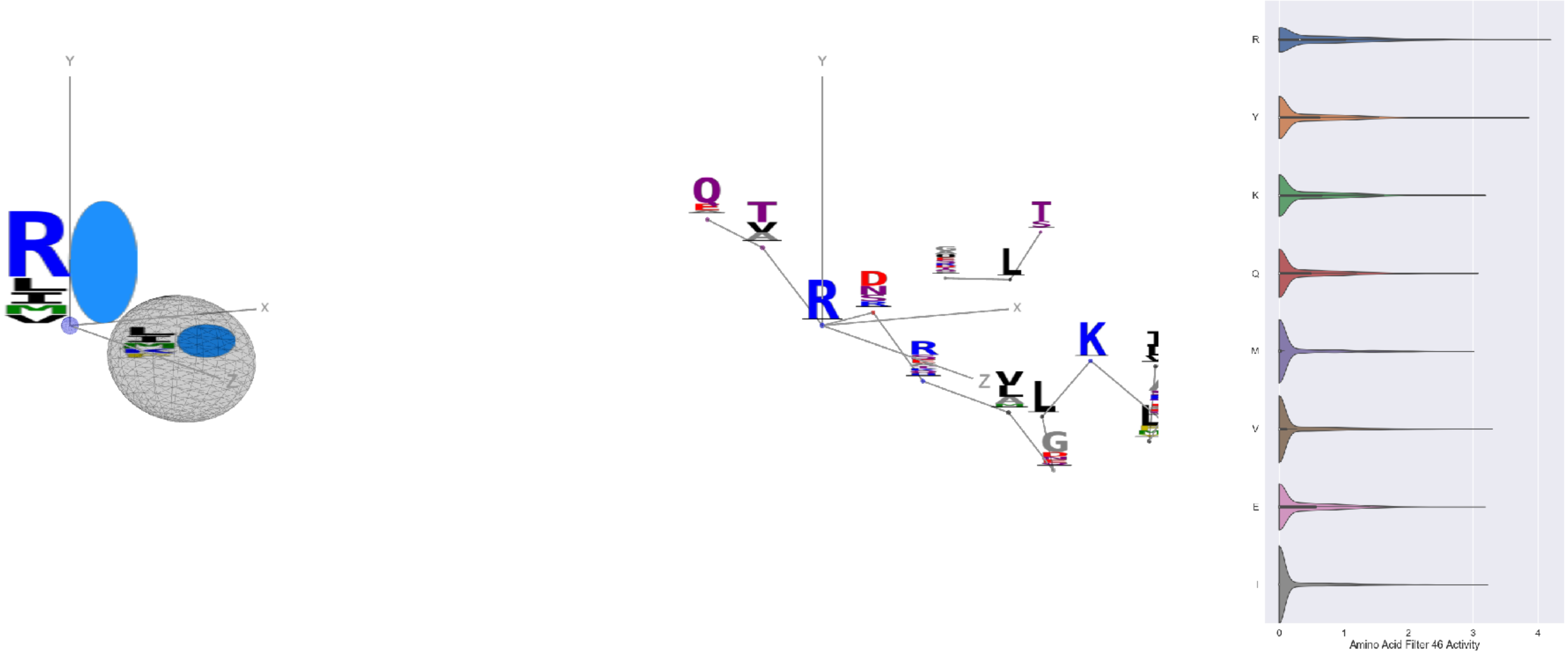

Filter 46
$R_{\mathrm{pred}} = 0.32$
Percent Active = 34
Pooling attention 0.26
Pooling feature 0.33
Top activators
3c57:A:197 R
1v2z:A:276 R
1ezv:C:70 R
2v6e:A:535 Y
3qsq:A:558 R
4c0b:D:480 R
2iwh:B:257 R
4kzx:R:67 R
3tsy:A:479 R
1o7g:A:448 Y

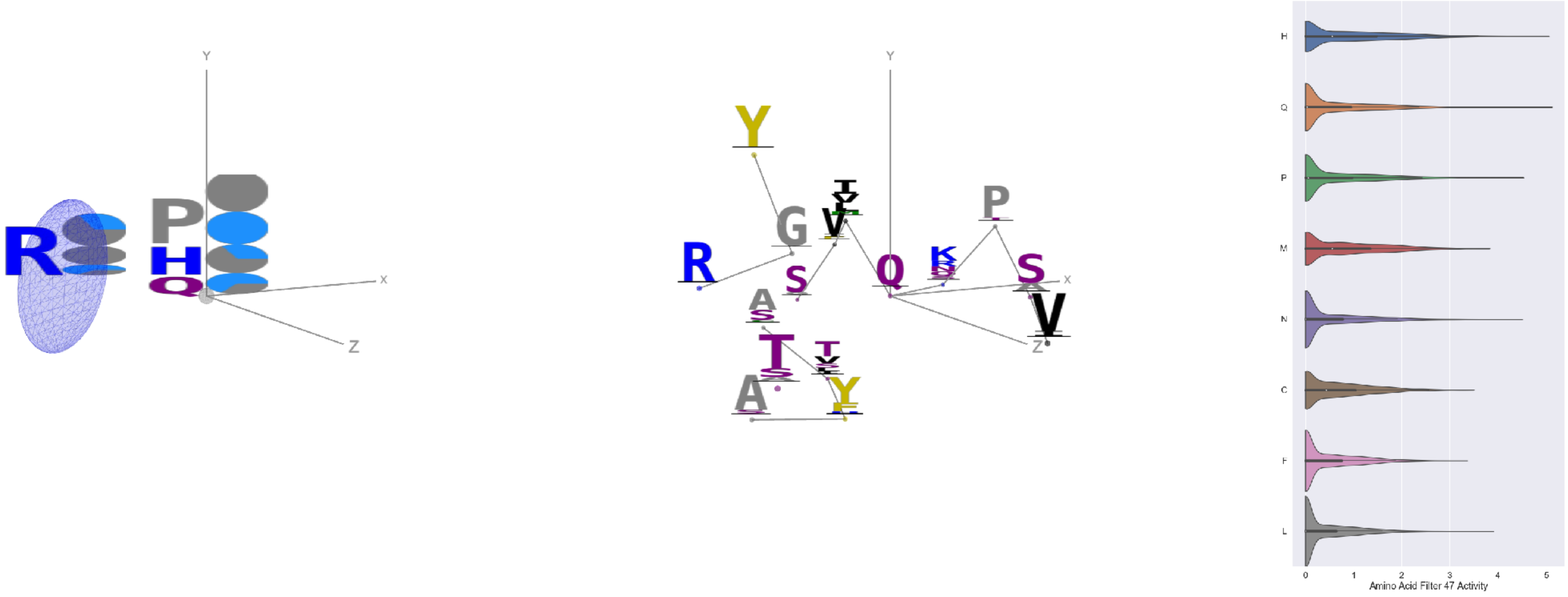

Filter 47
$R_{\mathrm{pred}} = -0.15$
Percent Active = 25
Pooling attention 0.05
Pooling feature 0.80
Top activators
4mex:C:659 Q
4uzi:A:304 H
1bxr:A:641 Q
2dyo:A:86 P
4kbj:B:369 N
4u4c:A:512 P
3di4:A:121 P
4u4c:A:539 Q
1bgv:A:162 P
1l7k:B:96 H

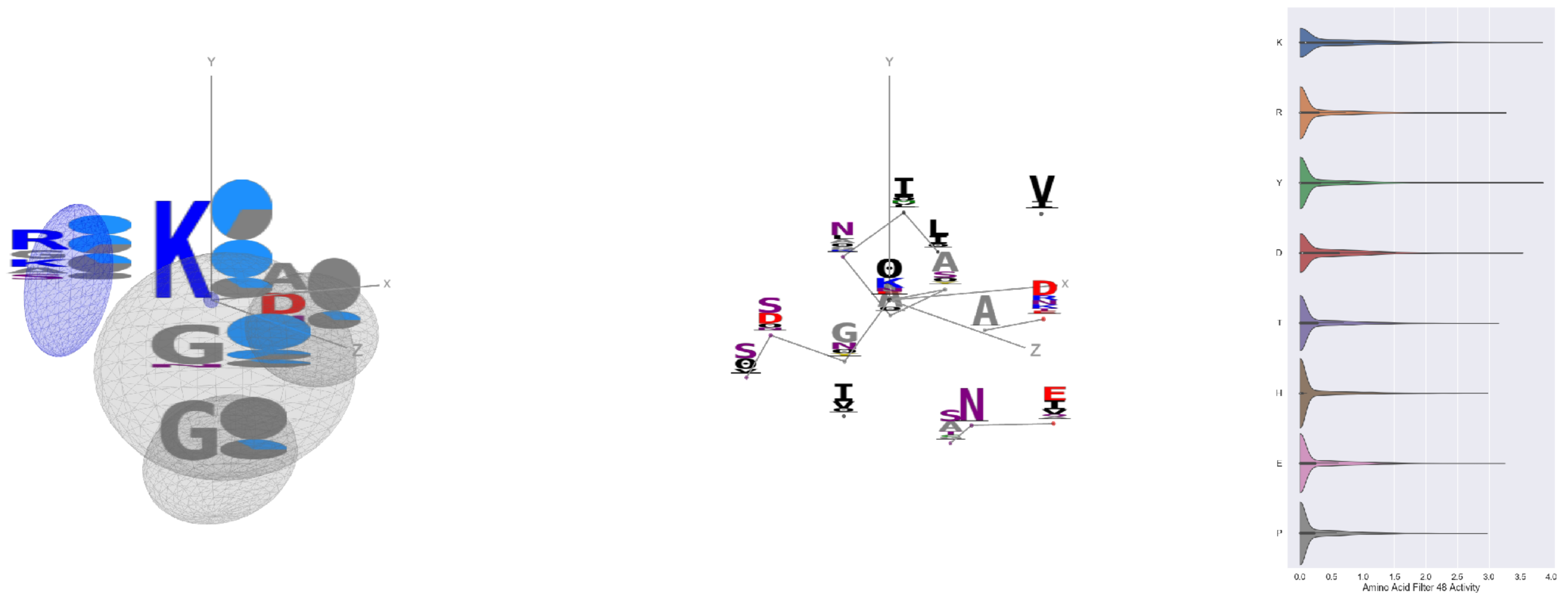


Filter 48
$R_{\text{pred}} = -0.08$
Percent Active = 30
Pooling attention 0.24
Pooling feature 0.39
Top activators
4v88:AF:203 Y
4cvw:A:745 K
4ddg:A:1271 K
3mk7:B:202 Y
2gfq:A:100 A
2ba1:A:81 Y
2pm9:A:75 G
3ztv:A:313 Y
1qha:A:282 D
1bh3:A:201 D

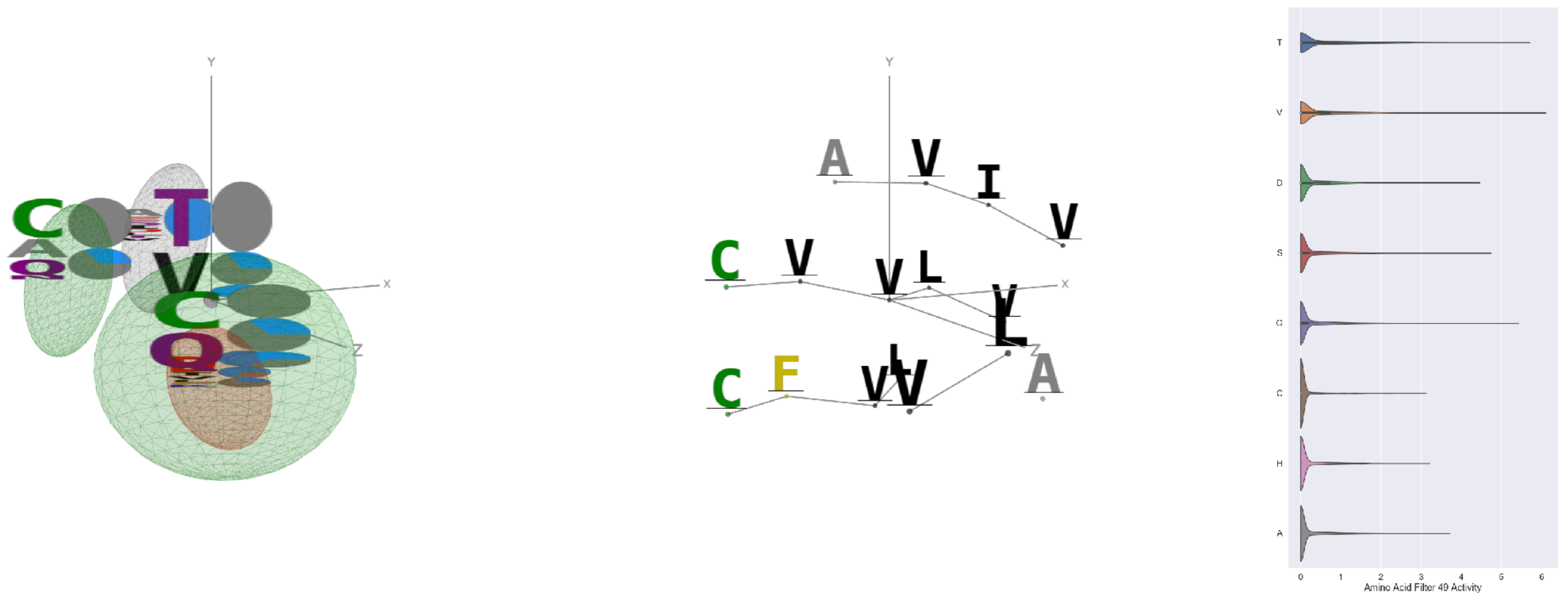


Filter 49
$R_{\text{pred}} = -0.03$
Percent Active = 20
Pooling attention 0.30
Pooling feature 0.48
Top activators
3tsy:A:775 V
4e0s:B:516 T
2iwh:B:91 V
3tsy:A:602 G
2iwh:B:361 V
4bxf:A:169 T
4pht:A:425 G
1eg1:A:23 G
3tsy:A:757 V
3tsy:A:598 Q

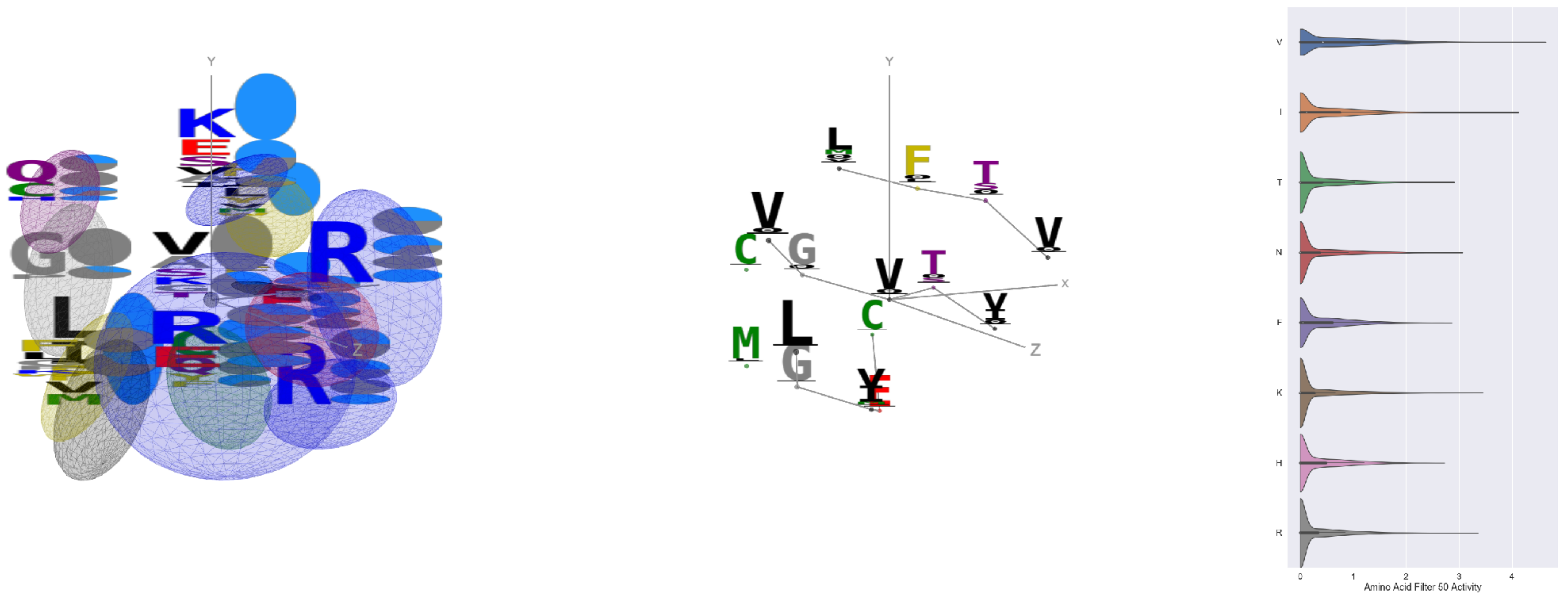


Filter 50
$R_{\text{pred}} = -0.20$
Percent Active = 34
Pooling attention 0.10
Pooling feature 1.11
Top activators
1qha:A:858 V
1wnl:A:125 V
3n6r:G:270 I
3t3o:A:475 V
2iwh:B:668 V
1qha:A:410 V
4hg6:A:242 V
1eg1:A:72 L
3noy:B:297 V
4oph:A:130 L

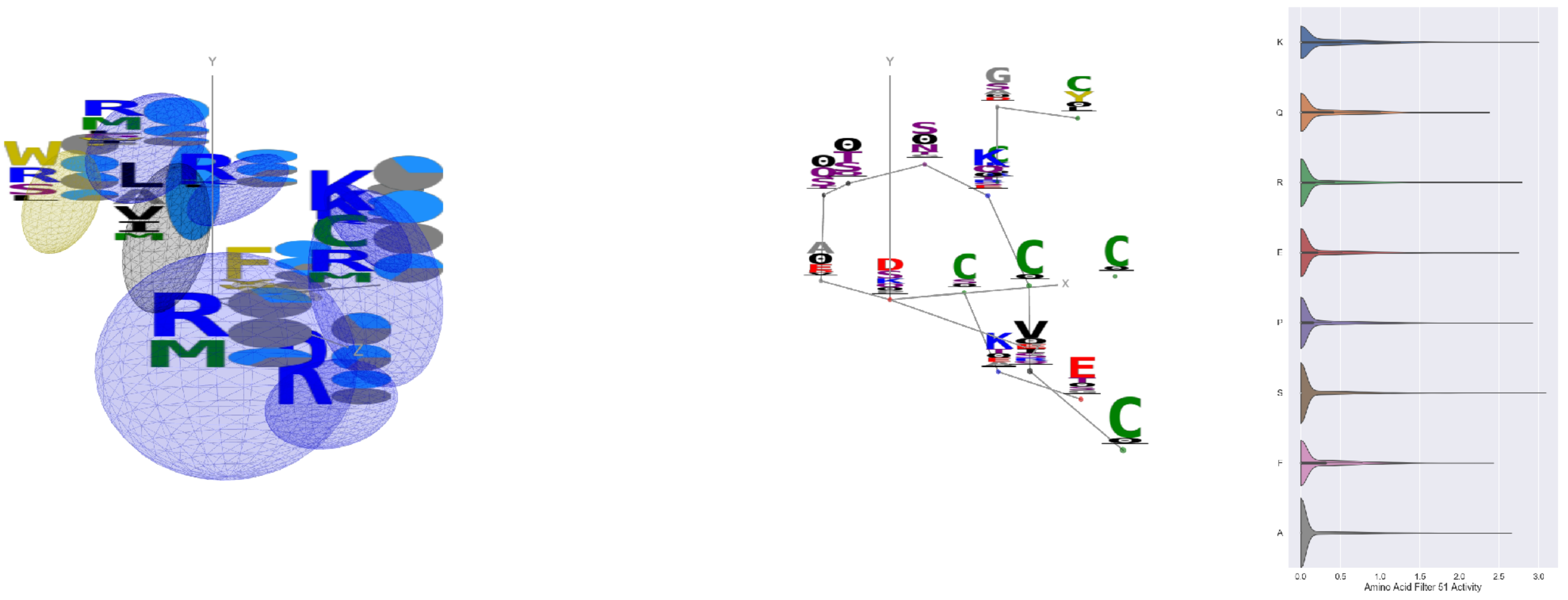


Filter 51
$R_{\text{pred}} = -0.07$
Percent Active = 24
Pooling attention 0.47
Pooling feature 0.43
Top activators
4e0s:B:853 D
4k1n:A:133 S
3a5c:G:203 Y
4kzx:R:107 K
1r7r:A:732 Y
3hwp:A:78 P
4qiw:A:153 Y
3s9m:C:258 G
1sl0:A:589 R
2akf:A:449 K

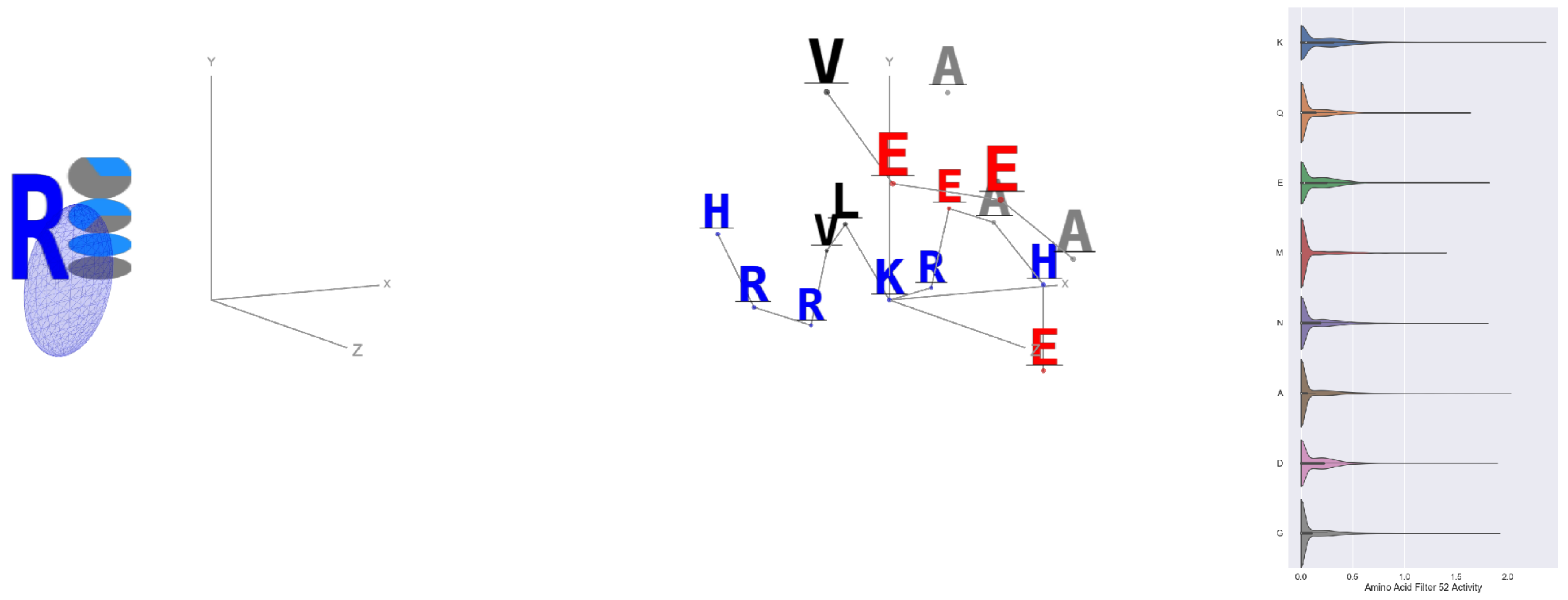

Filter 52
$R_{\mathrm{pred}} = 0.20$
Percent Active = 31
Pooling attention 0.15
Pooling feature 1.65
Top activators
3g8q:A:48 K
2v6e:A:380 K
2o35:B:15 A
2iwh:B:314 K
4rcn:B:606 G
1j5w:B:61 D
4s20:L:217 V
3tsy:A:710 K
3g8q:A:257 D
4r3z:C:513 D

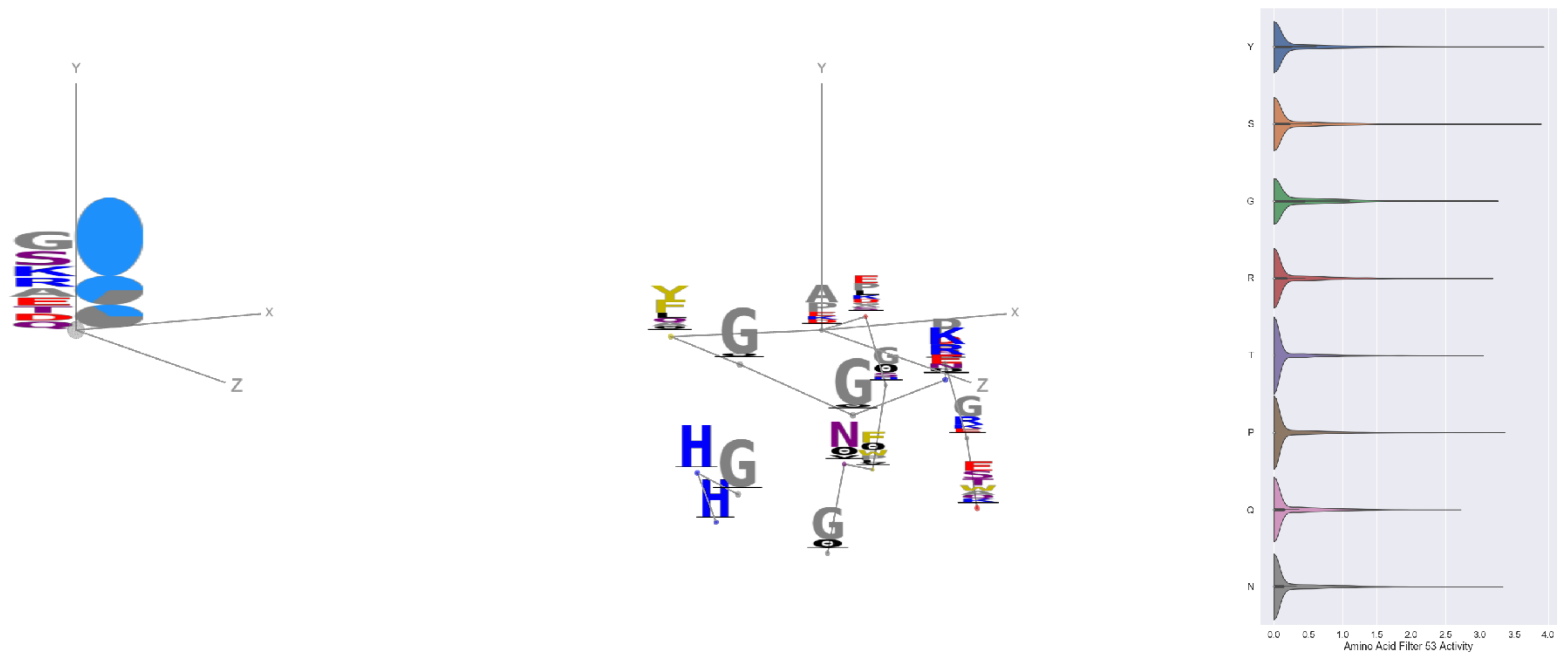

Filter 53
$R_{\mathrm{pred}} = 0.04$
Percent Active = 22
Pooling attention 0.58
Pooling feature 0.10
Top activators
2gfq:A:100 A
2d4z:B:686 Y
1e5d:A:261 S
2wgq:B:79 Y
2d73:A:660 A
2fml:A:4 Y
3drw:A:450 Y
4rap:E:19 Y
1jv1:A:220 P
1f75:A:225 P

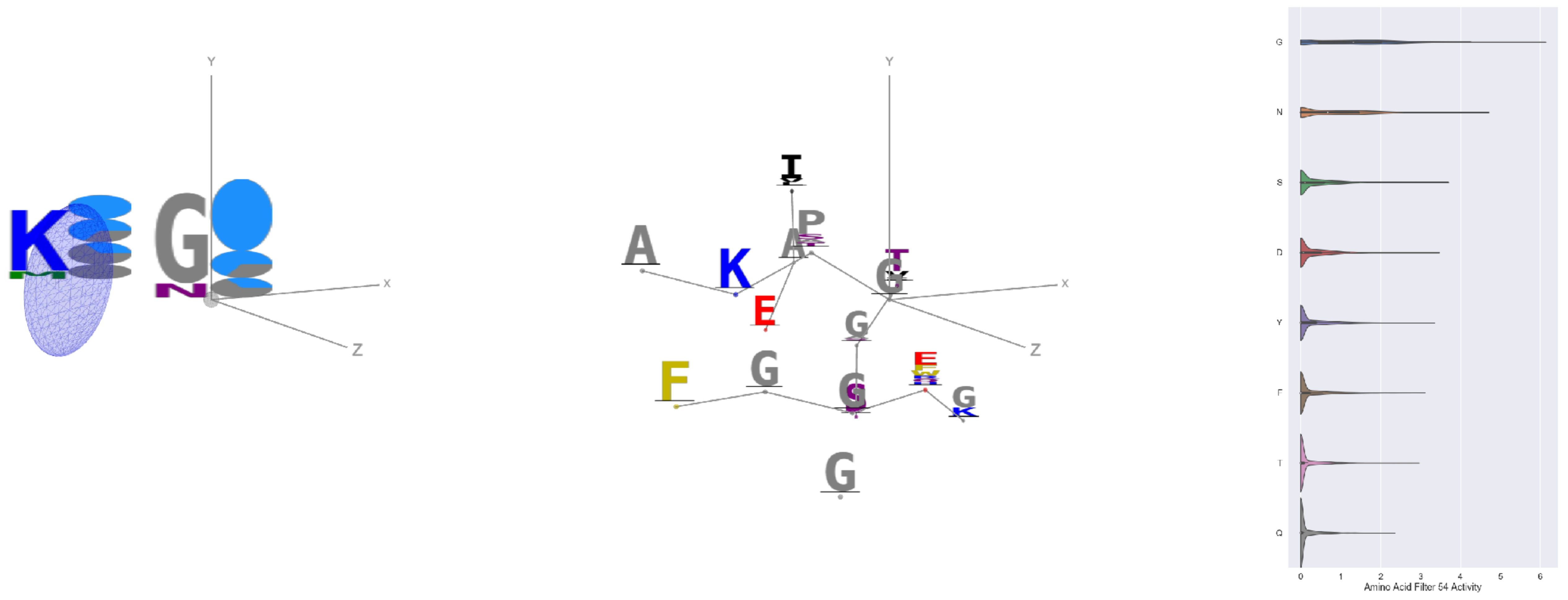

Filter 54
$R_{\mathrm{pred}} = 0.04$
Percent Active = 31
Pooling attention 0.02
Pooling feature 0.44
Top activators
4nm9:B:966 G
4ldb:A:273 G
4c2m:A:932 G
3tsy:A:160 N
1qha:A:701 G
1qha:A:590 G
3do6:A:74 G
4mex:C:703 G
4u4c:A:249 G
4gfh:A:698 G

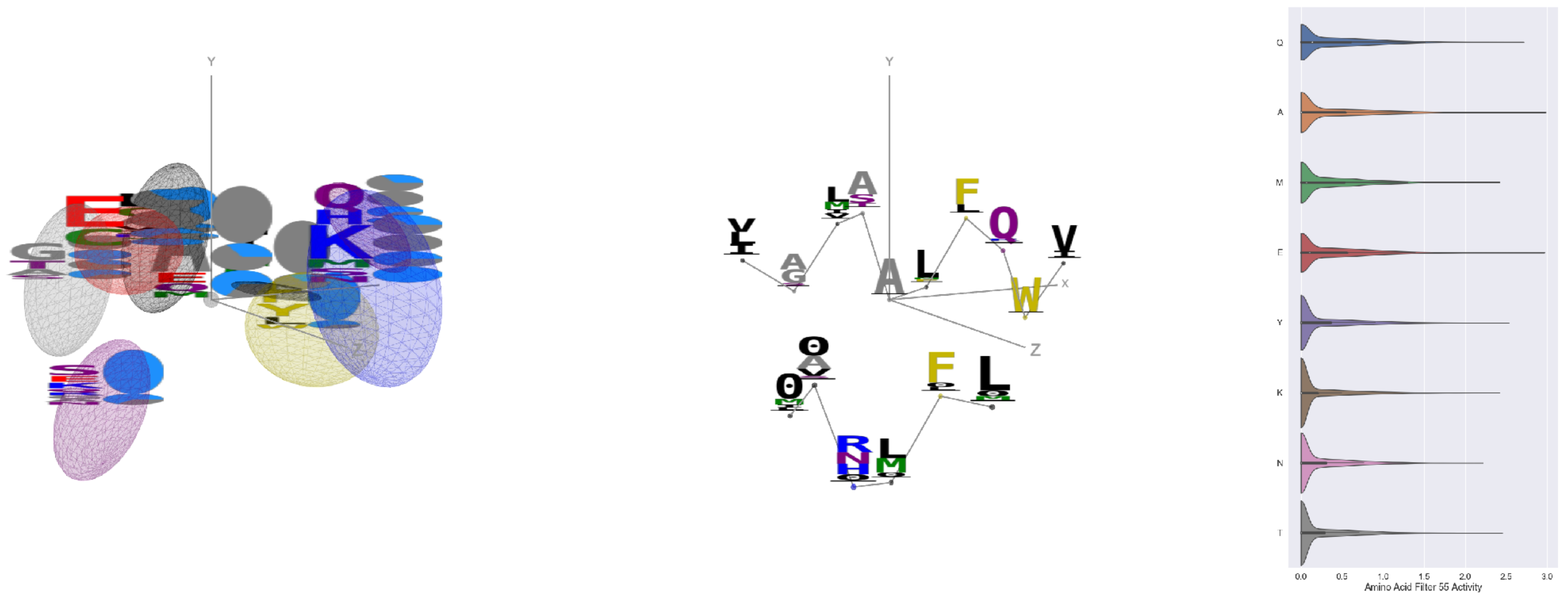

Filter 55
$R_{\mathrm{pred}} = 0.06$
Percent Active = 29
Pooling attention 0.10
Pooling feature 0.37
Top activators
4c8q:H:709 A
1bkj:A:6 E
3klt:A:296 E
1lql:A:43 E
2iwh:B:301 A
4oph:A:142 E
3te3:A:729 D
4mex:C:904 A
4oph:A:199 Q
3seo:B:234 A

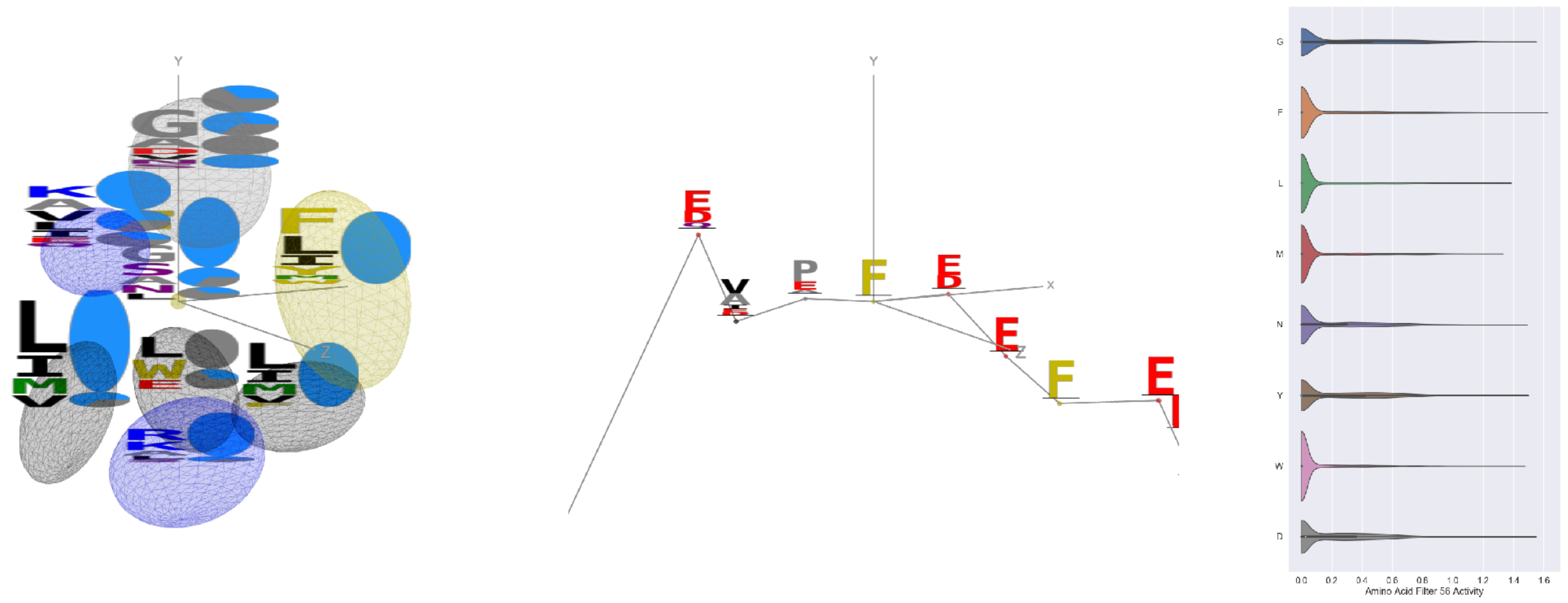


Filter 56
$R_{\text{pred}} = 0.35$
Percent Active = 35
Pooling attention 0.00
Pooling feature 2.02
Top activators
3t5v:C:38 F
4asi:C:2088 G
3din:D:48 D
2iwh:B:515 G
1izl:D:269 F
5uqy:A:180 Y
3qzy:B:237 N
1mu2:A:383 G
3o47:A:121 F
2yal:A:78 W

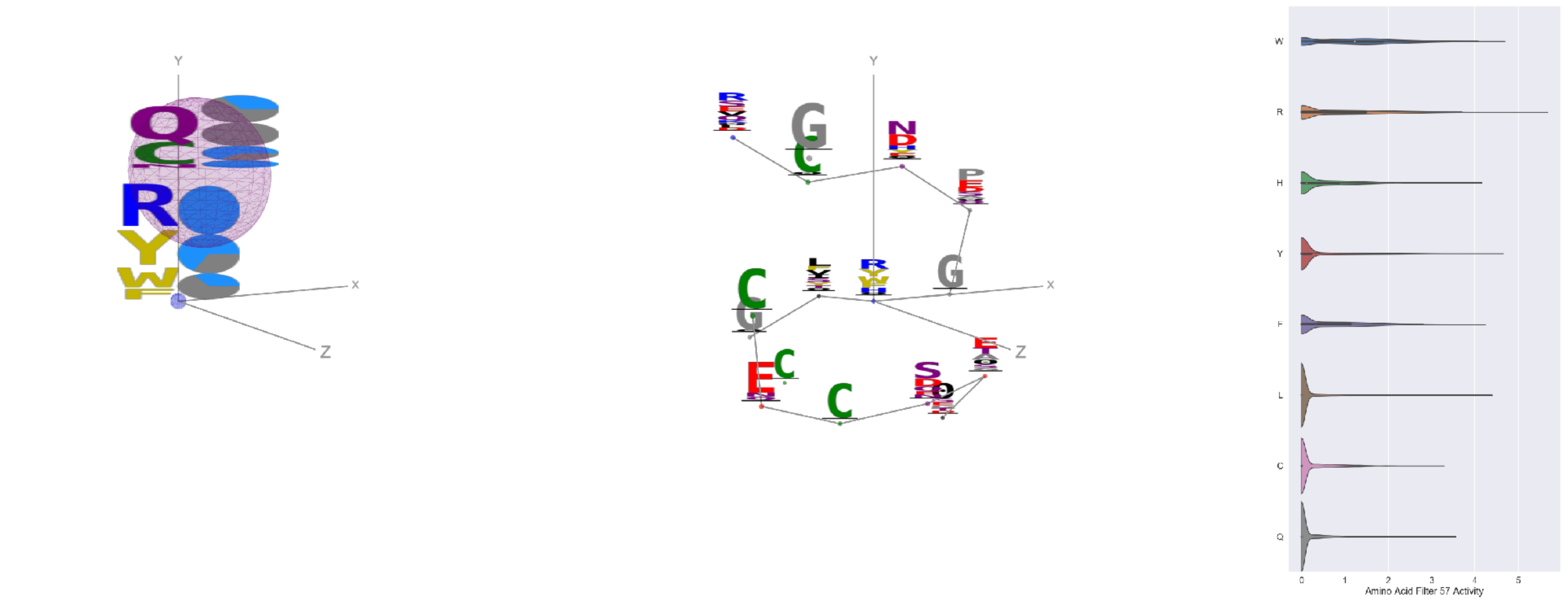


Filter 57
$R_{\text{pred}} = 0.12$
Percent Active = 21
Pooling attention 0.14
Pooling feature 1.37
Top activators
4g1m:B:466 R
1k72:B:375 R
3g8q:A:119 R
3g8q:A:65 R
4cvw:A:705 W
1umg:A:91 Y
4gfh:A:144 Y
2wgq:B:69 Y
2dsq:G:204 W
3gku:A:143 R

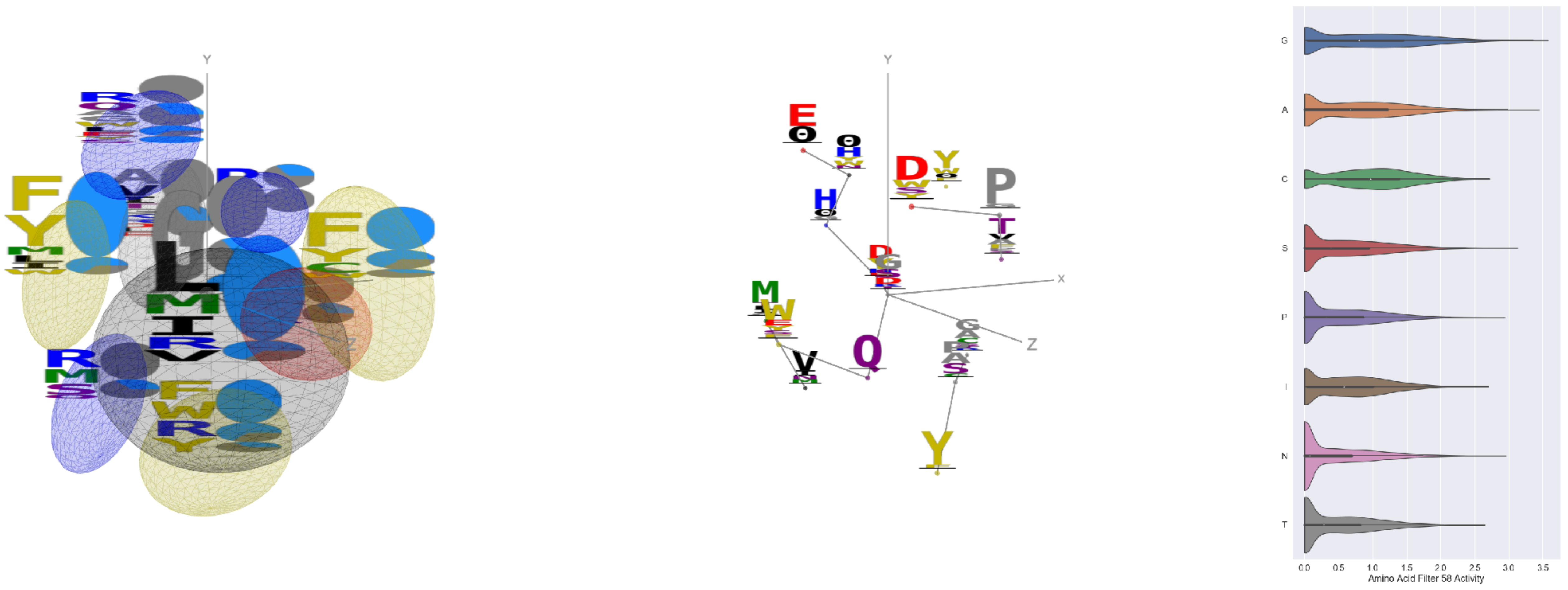


Filter 58
$R_{\text{pred}} = -0.25$
Percent Active = 50
Pooling attention 0.15
Pooling feature 0.44
Top activators
2d73:A:483 G
4uzi:A:303 A
3bma:C:117 G
1qvr:A:152 A
3b8e:A:858 G
2z67:A:248 A
1w36:B:418 A
4kbj:B:386 G
1bxr:A:92 G
4v88:AF:141 G

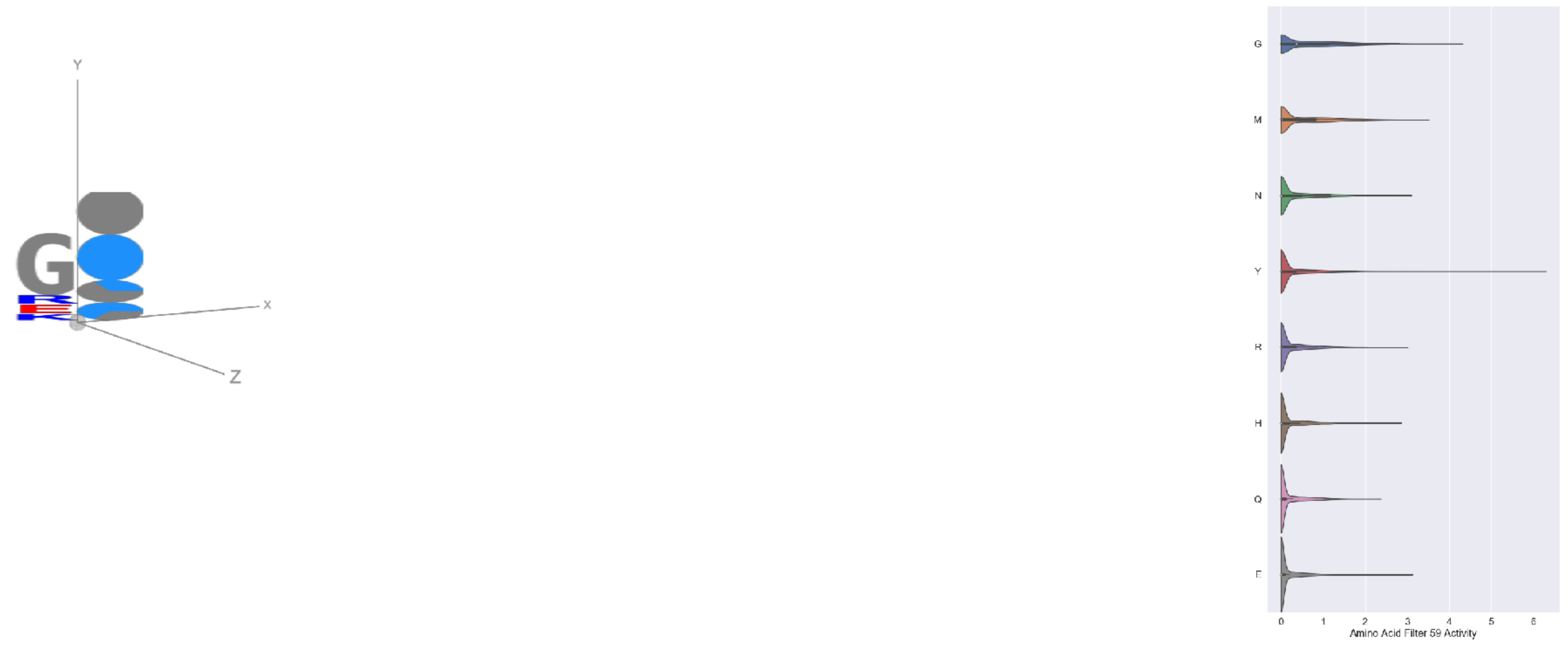


Filter 59
$R_{\text{pred}} = -0.09$
Percent Active = 24
Pooling attention 0.05
Pooling feature 0.58
Top activators
2v6e:A:535 Y
1o7g:A:448 Y
1hfe:L:398 Y
1gvn:A:90 Y
2np9:A:257 G
2iut:A:723 Y
2yma:A:399 Y
1ii2:A:221 G
2dyo:A:285 Y
3qzy:B:260 Y

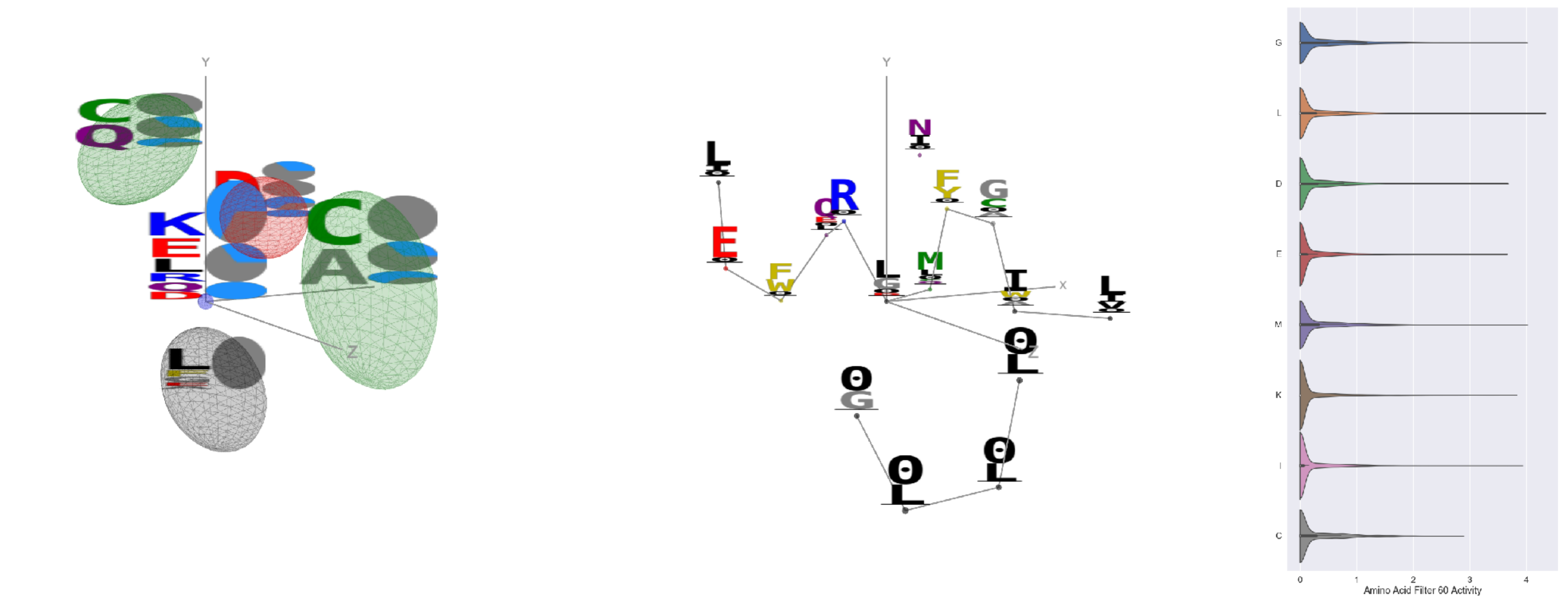

Filter 60
$R_{pred} = -0.20$
Percent Active = 28
Pooling attention 0.11
Pooling feature 0.21
Top activators
1izl:D:127 L
4v81:H:106 L
1r7r:A:642 L
1r7r:A:575 F
4ny2:A:267 T
1jb0:K:69 L
1izl:D:197 H
1izl:D:185 F
1izl:D:199 M
1r7r:A:616 G

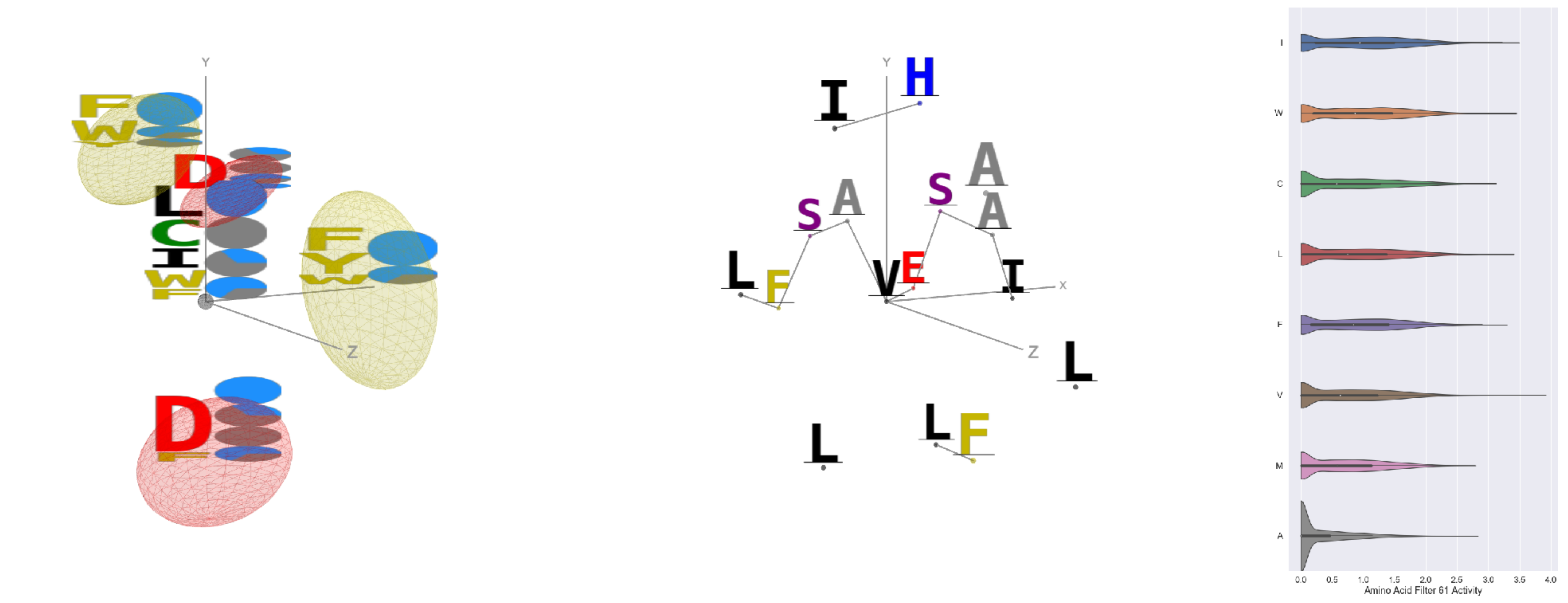

Filter 61
$R_{pred} = 0.05$
Percent Active = 45
Pooling attention 0.53
Pooling feature 1.11
Top activators
2iwh:B:32 V
2iwh:B:73 I
1mu2:A:229 W
1qha:A:531 L
1p4e:B:180 F
4mex:C:176 I
4buj:B:633 L
3bsd:A:18 I
3bsd:A:148 I
1ezv:C:154 I

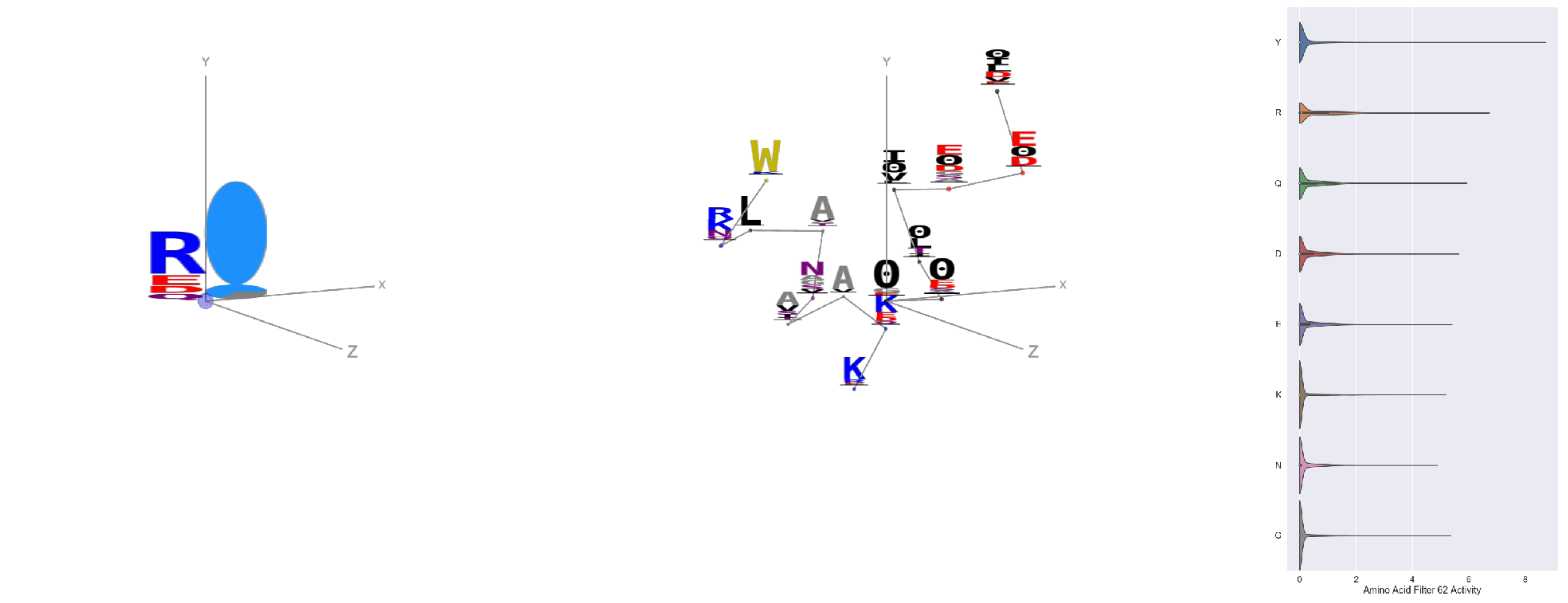

Filter 62
$R_{pred} = 0.08$
Percent Active = 20
Pooling attention 0.18
Pooling feature 0.93
Top activators
2iut:A:723 Y
2nog:A:896 Y
1oqo:C:39 Y
2v6e:A:535 Y
1o7g:A:448 Y
1f75:A:242 R
1hfe:L:398 Y
2i71:B:336 R
2iwh:B:977 W
5ajd:B:469 I

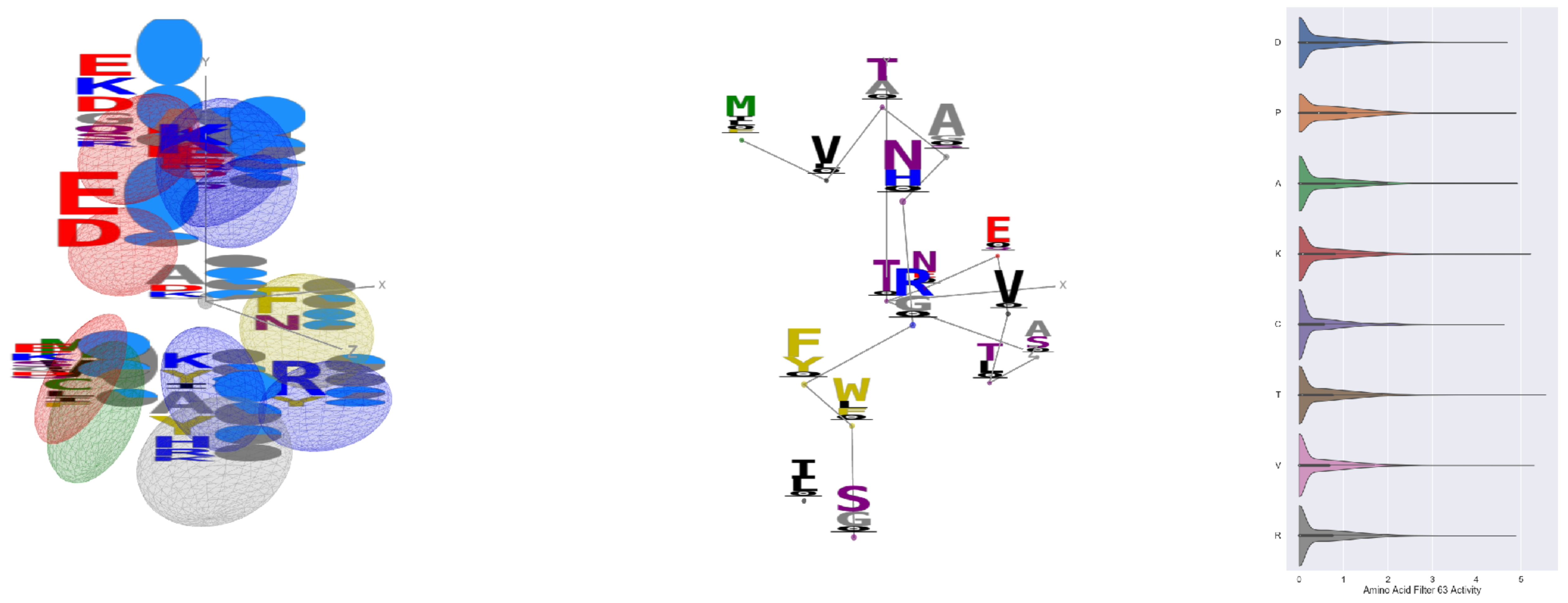

Filter 63
$R_{pred} = -0.18$
Percent Active = 46
Pooling attention 0.83
Pooling feature 1.76
Top activators
1izl:D:221 T
1izl:D:78 V
4mex:C:163 K
1izl:D:77 A
1izl:D:194 N
4v81:H:324 P
1r7r:A:641 R
3kio:B:271 Y
1r7r:A:577 D
1r7r:A:522 C

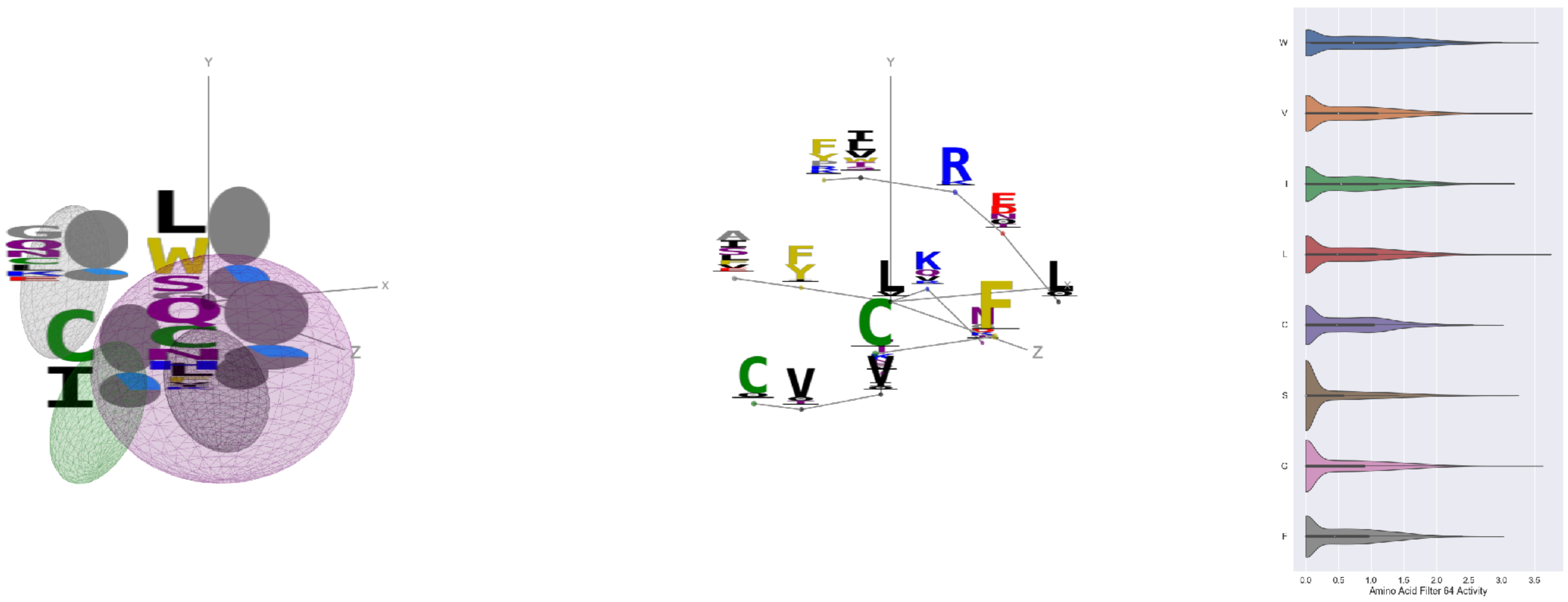

Filter 64
$R_{\mathrm{pred}} = -0.27$
Percent Active = 39
Pooling attention 0.17
Pooling feature 0.56
Top activators
1f34:B:20 L
1p35:A:44 L
2iut:A:626 L
3ax7:A:46 G
4rcn:B:934 L
4ny2:A:229 W
3hzu:A:248 G
4e0s:B:313 L
4y97:A:420 V
1iwa:A:170 L

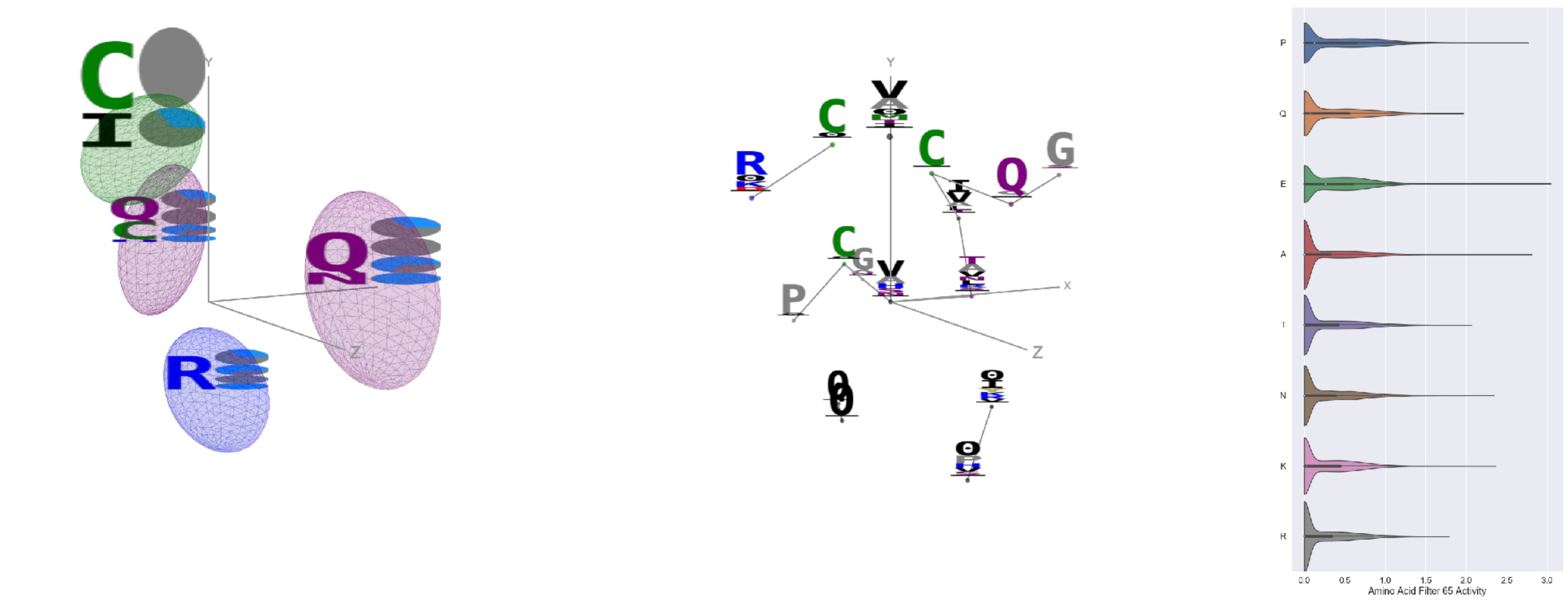

Filter 65
$R_{\mathrm{pred}} = 0.12$
Percent Active = 33
Pooling attention 0.16
Pooling feature 0.46
Top activators
1hfe:L:75 V
4ny2:A:117 E
4e0s:B:503 A
3noy:B:266 P
1jqn:A:700 P
4ehi:B:458 A
4v8p:BC:200 K
1qvr:A:192 N
2np9:A:233 A
3ilk:A:20 A

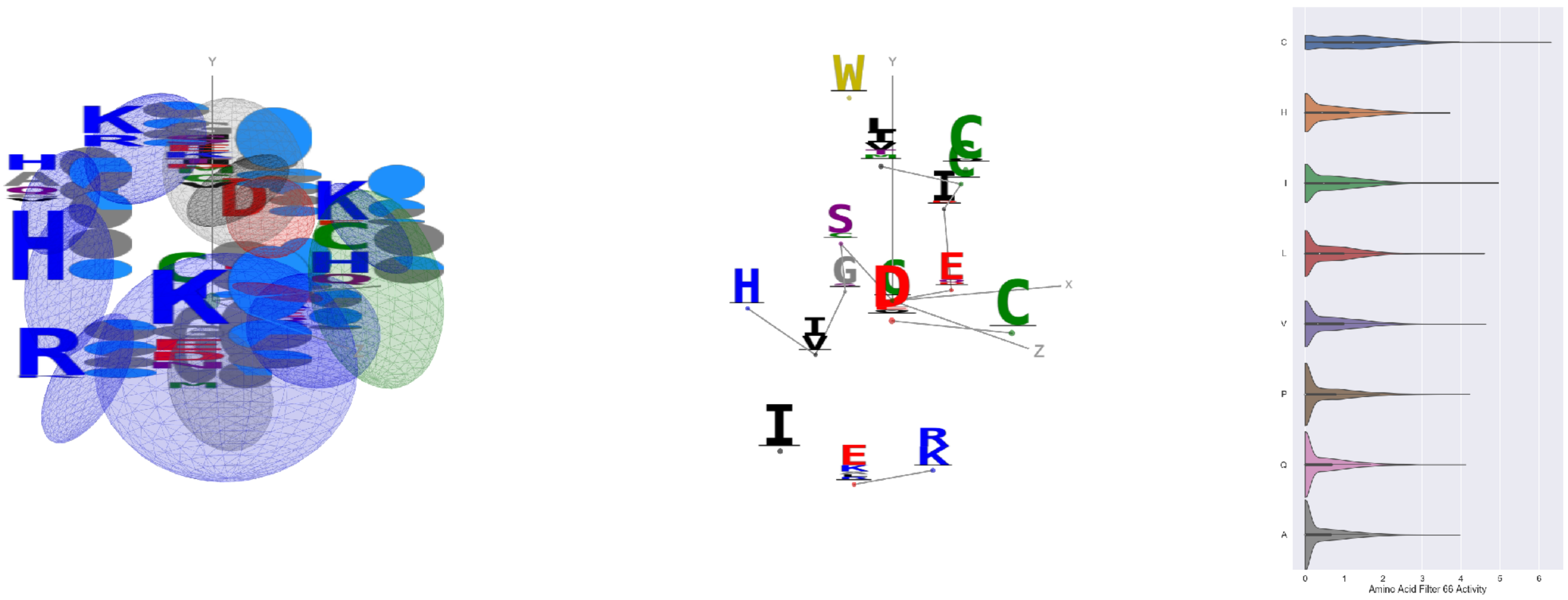

Filter 66
$R_{\mathrm{pred}} = -0.24$
Percent Active = 43
Pooling attention 0.66
Pooling feature 0.63
Top activators
2b0o:F:464 C
1r7r:A:540 I
4n4n:A:196 C
4e0s:B:531 C
1r7r:A:562 V
1r7r:A:642 L
2qn5:B:118 C
4n4n:A:283 C
1r7r:A:680 E
3i5d:B:135 C

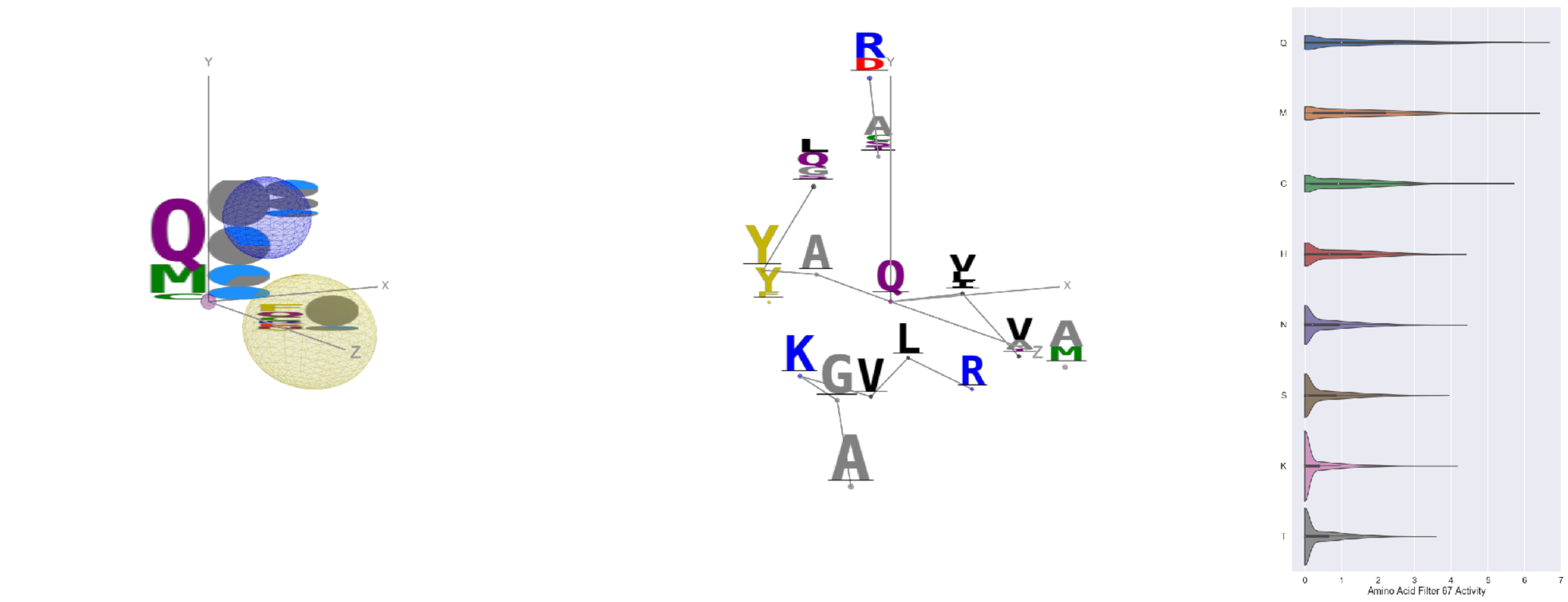

Filter 67
$R_{\mathrm{pred}} = -0.09$
Percent Active = 31
Pooling attention 0.22
Pooling feature 0.25
Top activators
4nm9:B:276 Q
1mj2:A:44 Q
3n6r:G:273 Q
3tsy:A:388 M
4rcn:B:210 Q
2iwh:B:760 Q
1e1c:A:330 Q
5ui2:A:226 Q
4nmn:A:357 Q
2w27:A:229 Q

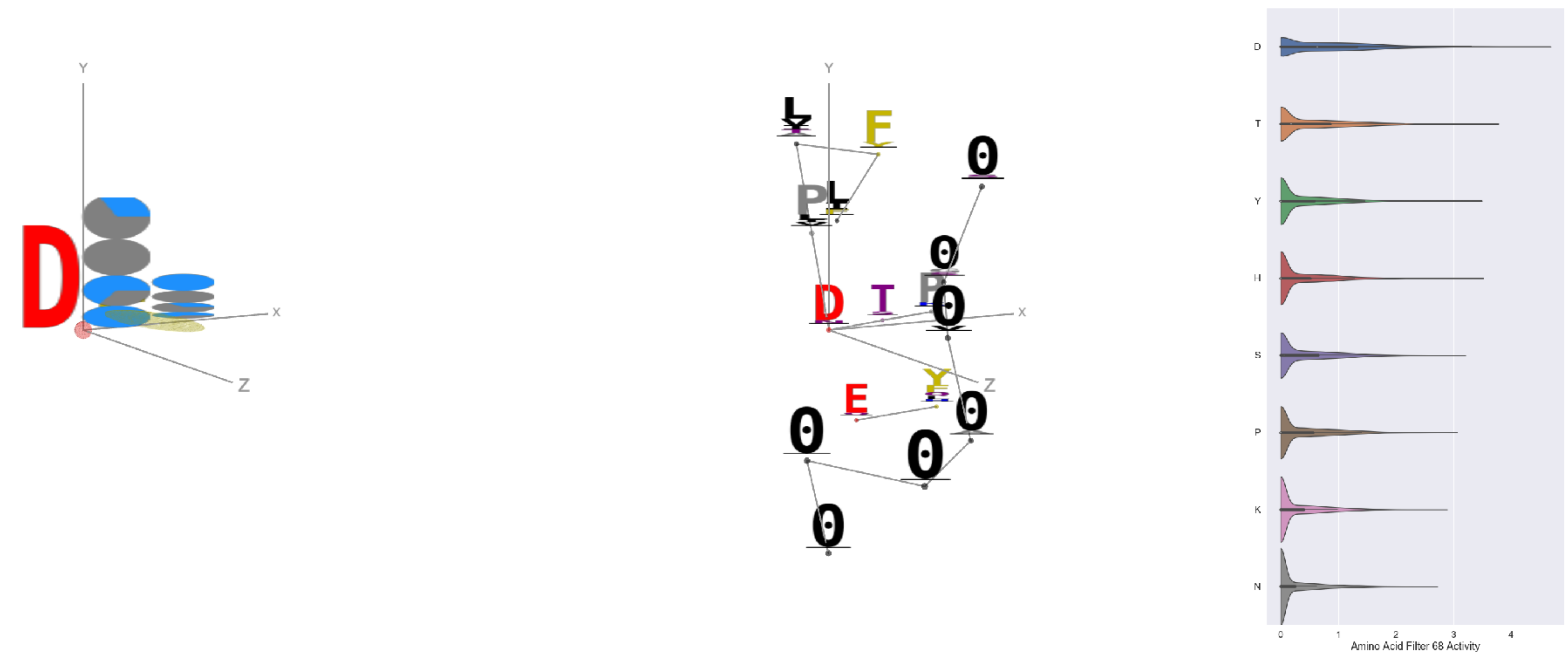

Filter 68
$R_{pred} = 0.11$
Percent Active = 33
Pooling attention 0.39
Pooling feature 0.40
Top activators
3kio:B:92 D
3kyh:C:371 D
4mex:C:624 D
4jdm:C:18 D
1n0e:B:107 D
4gfh:A:776 D
4nnq:B:71 D
1iri:A:511 D
1pgu:B:562 D
4v88:AF:55 D

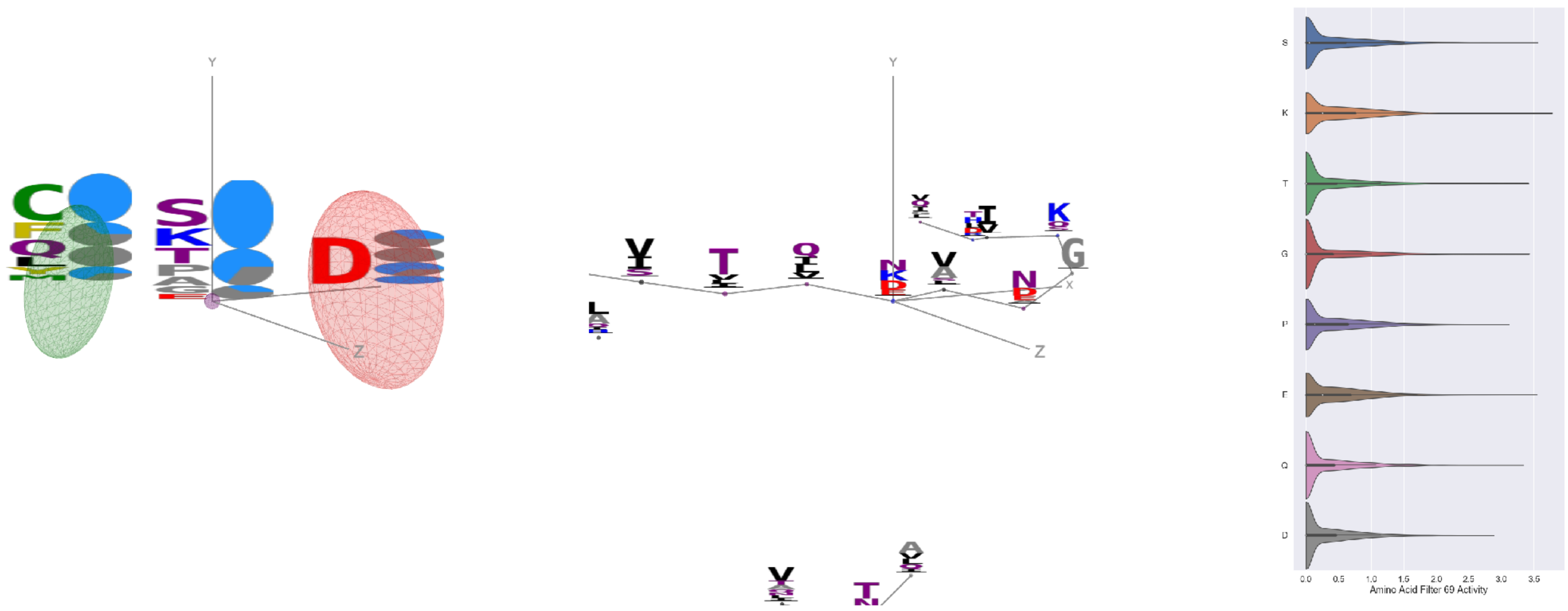

Filter 69
$R_{pred} = 0.21$
Percent Active = 33
Pooling attention 0.19
Pooling feature 0.35
Top activators
3a1m:A:29 K
2hd0:A:173 K
3kio:B:72 S
4c2m:N:52 E
2hd0:A:172 K
1a71:A:98 G
1qhd:A:169 S
5dzd:B:504 T
1k90:C:589 Q
2h4o:A:48 S

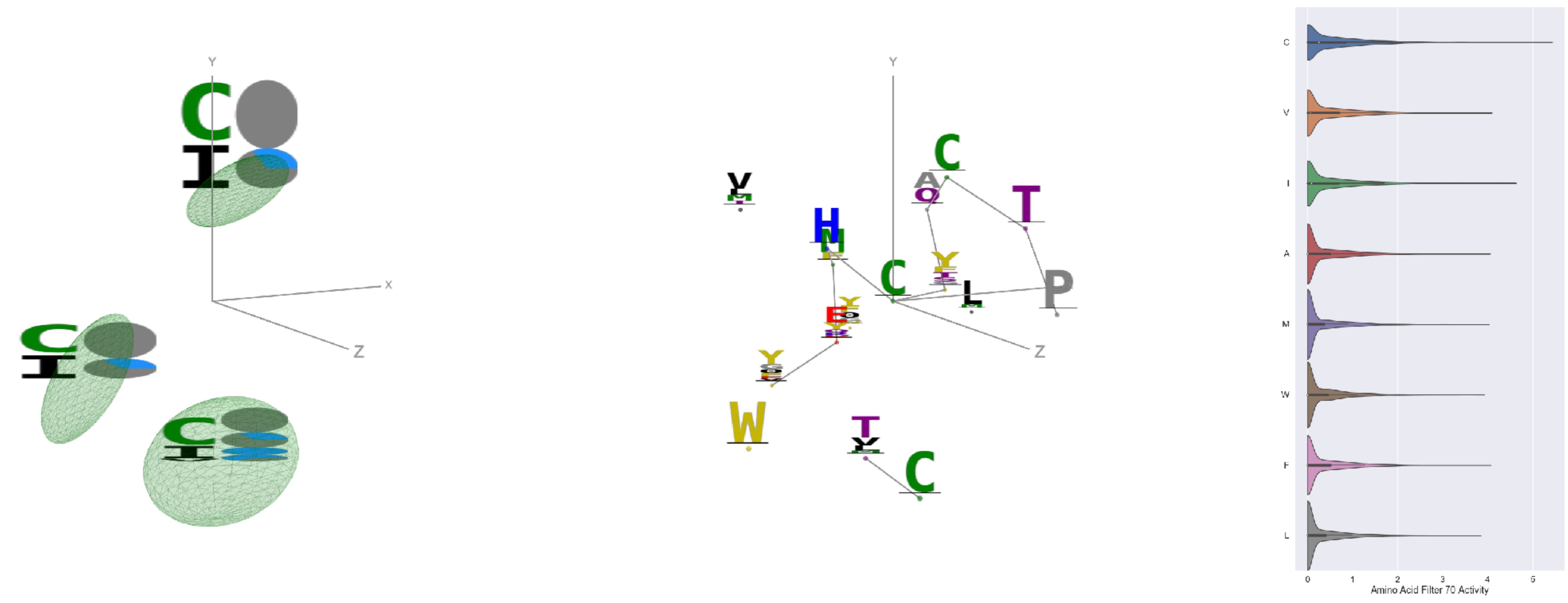

Filter 70
$R_{pred} = -0.17$
Percent Active = 25
Pooling attention 0.19
Pooling feature 1.03
Top activators
4n4n:A:337 C
1hfe:L:386 G
1hfe:L:233 P
3s9m:C:158 C
2iwh:B:665 I
4n4n:A:286 C
1hfe:L:385 G
2gy7:B:113 Y
1h8p:A:59 G
1eg1:A:290 I

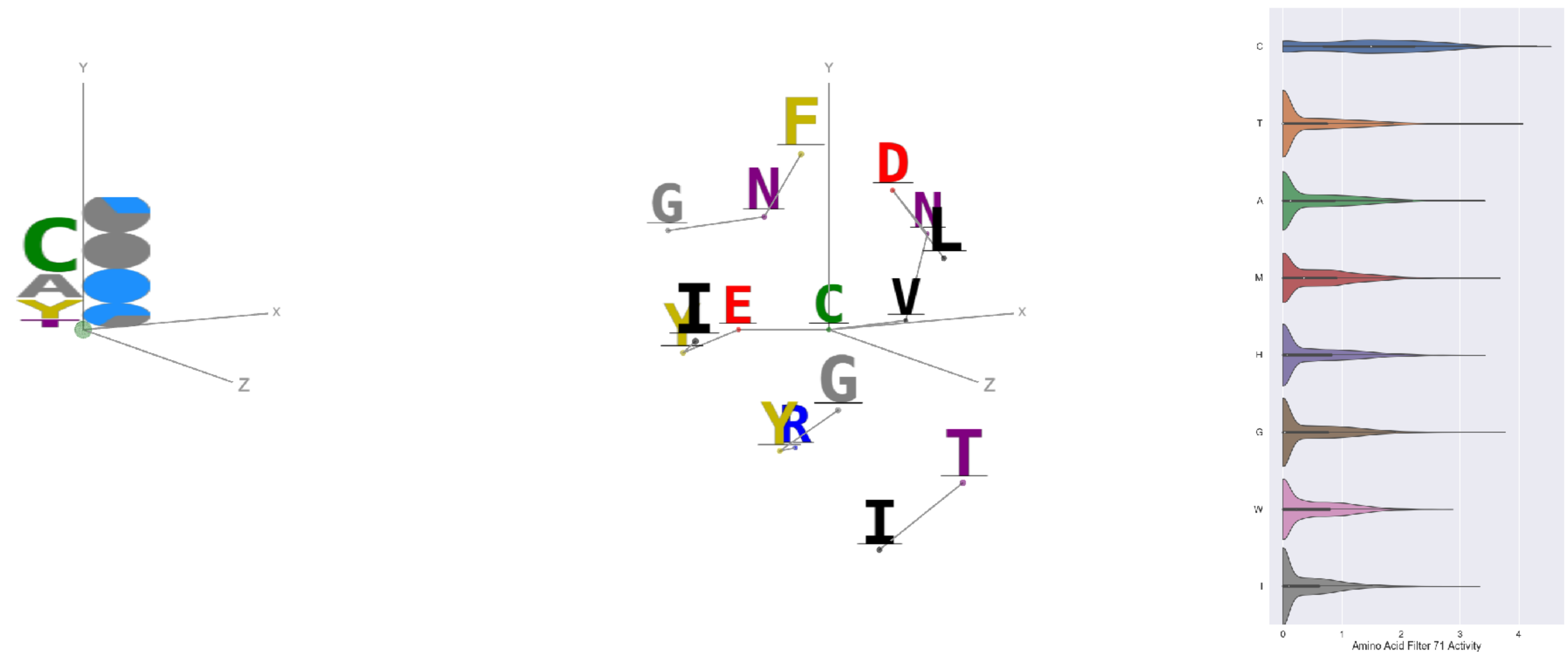

Filter 71
$R_{pred} = 0.04$
Percent Active = 34
Pooling attention 0.28
Pooling feature 1.10
Top activators
2iwh:B:606 C
2iwh:B:433 C
4n4n:A:283 C
4ny2:A:35 Y
2iwh:B:501 T
2hd0:A:184 C
4e0s:B:125 C
4e0s:B:531 C
1eg1:A:175 C
2zy4:F:318 G

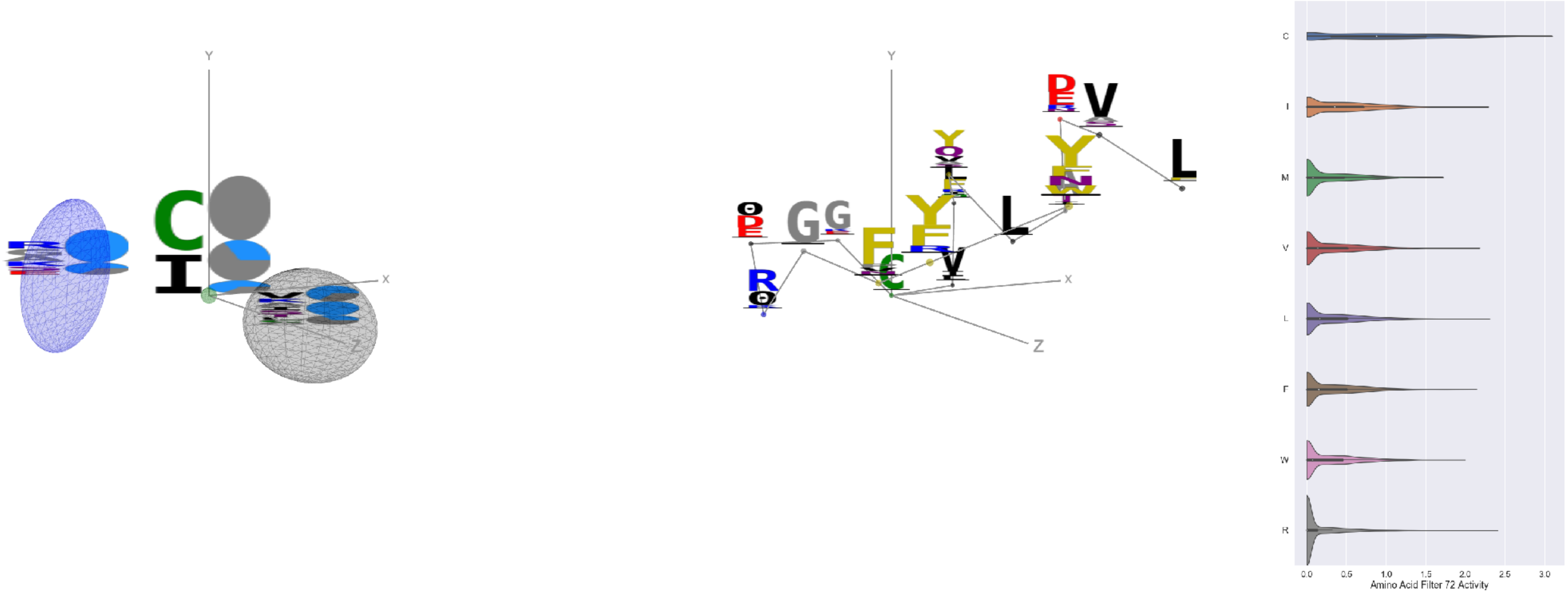

Filter 72
$R_{\text{pred}} = 0.13$
Percent Active = 31
Pooling attention 0.06
Pooling feature 0.06
Top activators
1b9e:B:19 C
3s9m:C:194 C
3noy:B:268 C
4n4n:A:106 C
2iwh:B:489 C
3tsy:A:647 C
4v8p:BF:163 C
5uqy:A:37 C
3mk7:B:68 C
3qzy:B:155 C

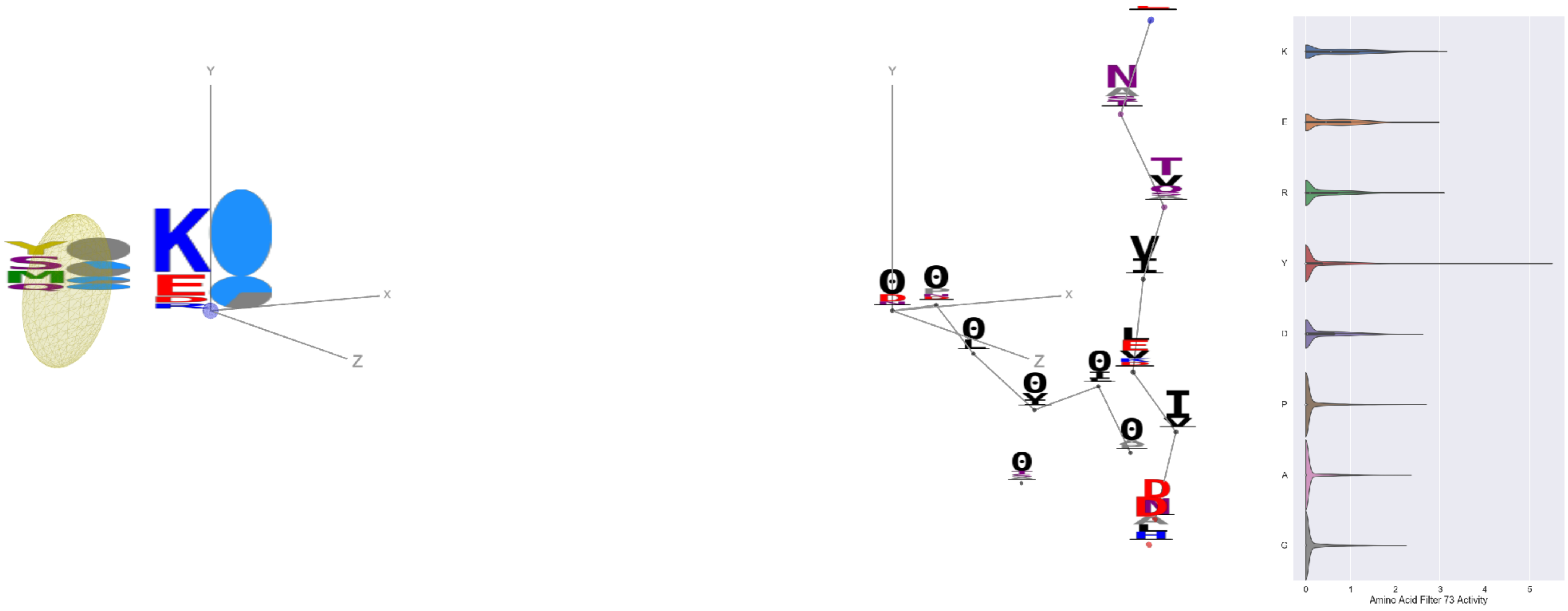

Filter 73
$R_{\text{pred}} = 0.15$
Percent Active = 32
Pooling attention 0.14
Pooling feature 0.18
Top activators
2v6e:A:535 Y
2iwh:B:977 W
1o7g:A:448 Y
2d32:A:7 Y
3tsy:A:702 K
2iwh:B:288 K
3l0o:A:330 R
2ovc:A:25 K
2iwh:B:663 K
1vpn:B:204 K

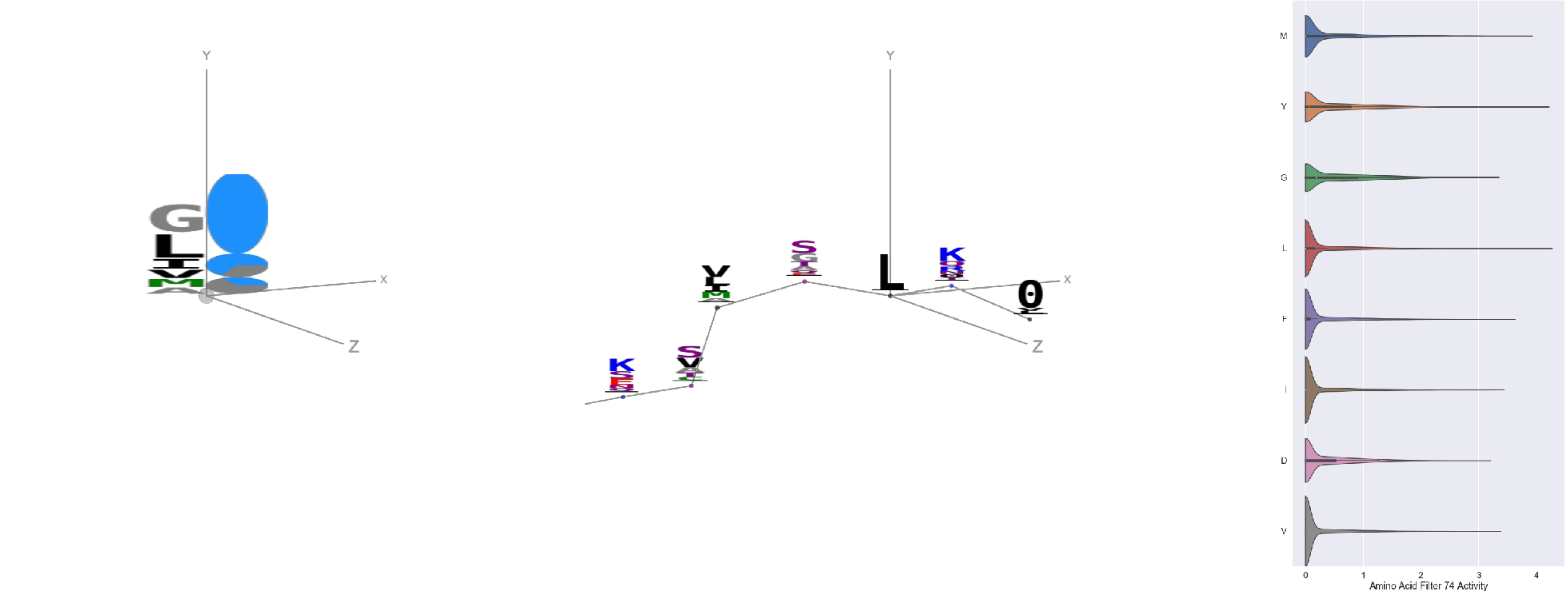

Filter 74
$R_{\text{pred}} = 0.36$
Percent Active = 33
Pooling attention 0.42
Pooling feature 0.78
Top activators
2pg1:L:112 L
1e79:I:1 Y
3t5v:C:24 Y
1jku:A:1 M
1ezv:C:1 Y
1iwa:A:8 Y
4bxf:A:290 L
2wgq:B:306 L
3irm:A:517 M
3g9w:A:193 Y

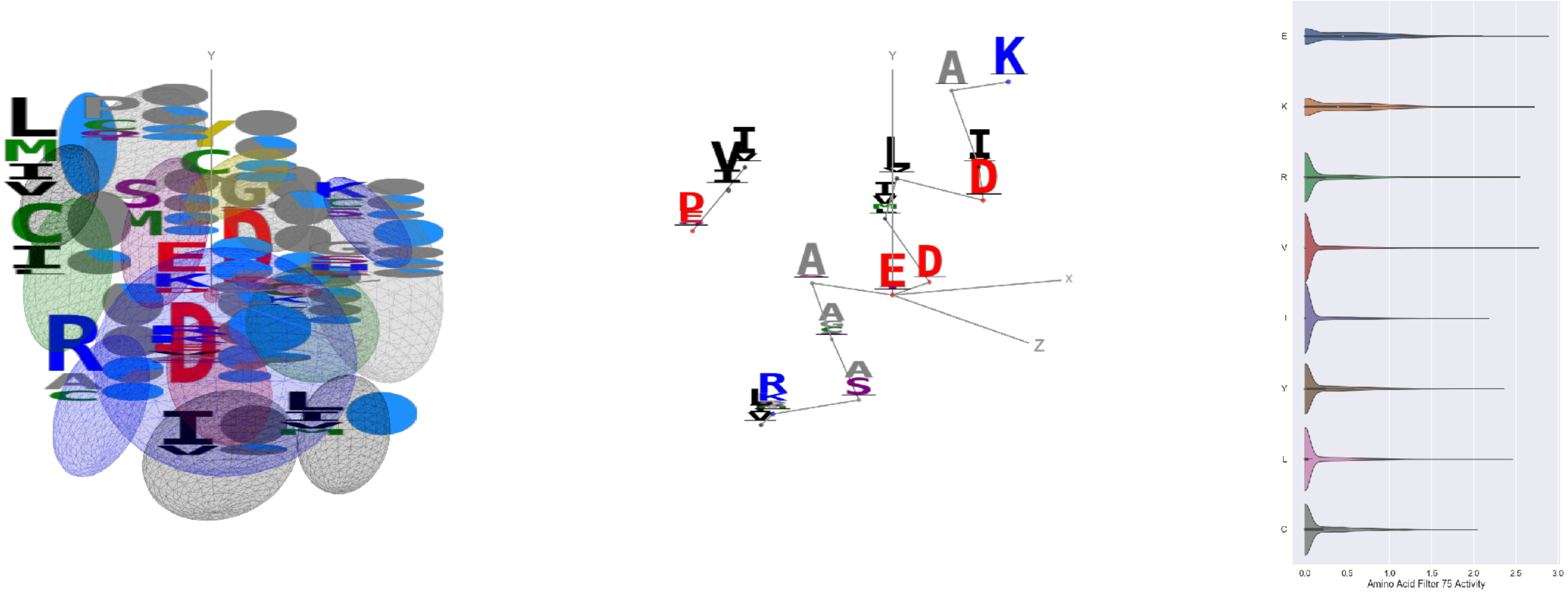

Filter 75
$R_{\text{pred}} = -0.07$
Percent Active = 32
Pooling attention 0.21
Pooling feature 1.23
Top activators
1qvr:A:387 E
4v81:A:296 V
3tsy:A:632 K
1r7r:A:529 K
1r7r:A:543 K
3qzy:B:12 K
2q6b:A:840 E
1r7r:A:678 R
1r7r:A:578 E
3drw:A:161 L

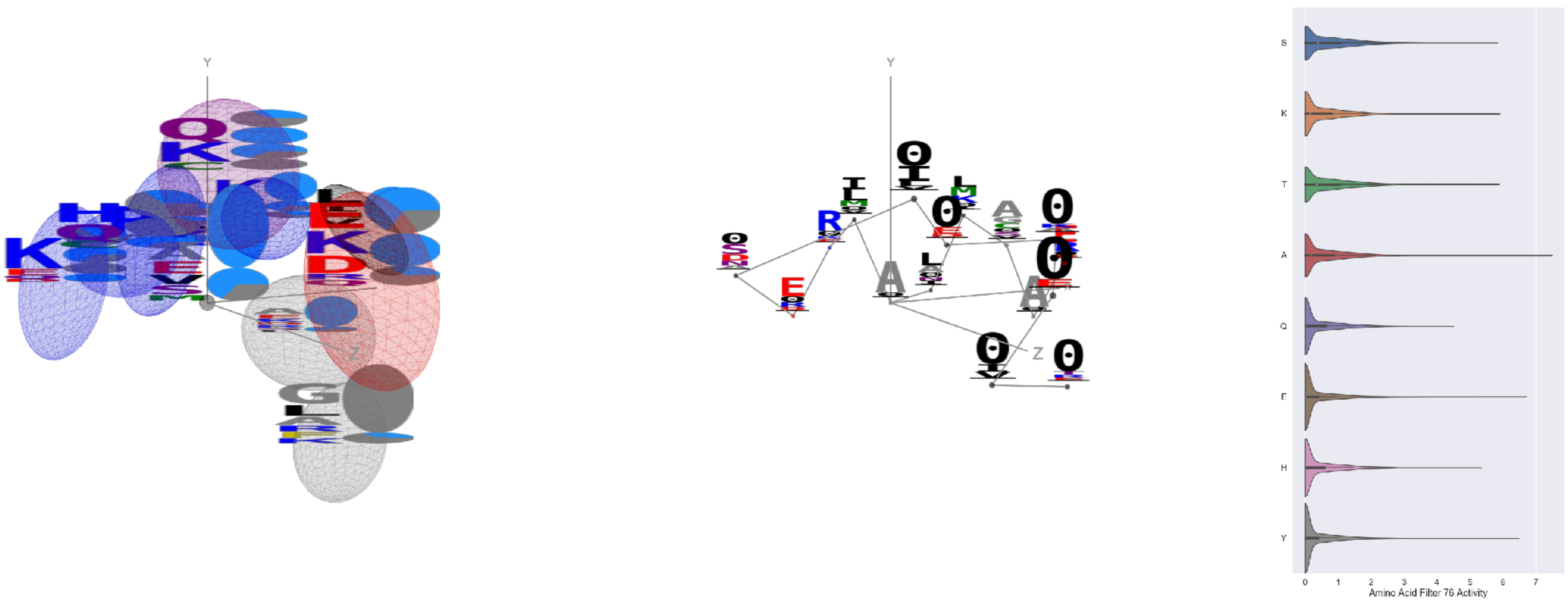

Filter 76
$R_{\text{pred}} = -0.20$
Percent Active = 41
Pooling attention 0.39
Pooling feature 1.13
Top activators
1r7r:A:698 A
1r7r:A:558 A
1r7r:A:483 E
1r7r:A:732 Y
1izl:D:338 Y
1r7r:A:676 A
1r7r:A:473 N
3a5c:G:17 R
1izl:D:312 Y
1r7r:A:529 K

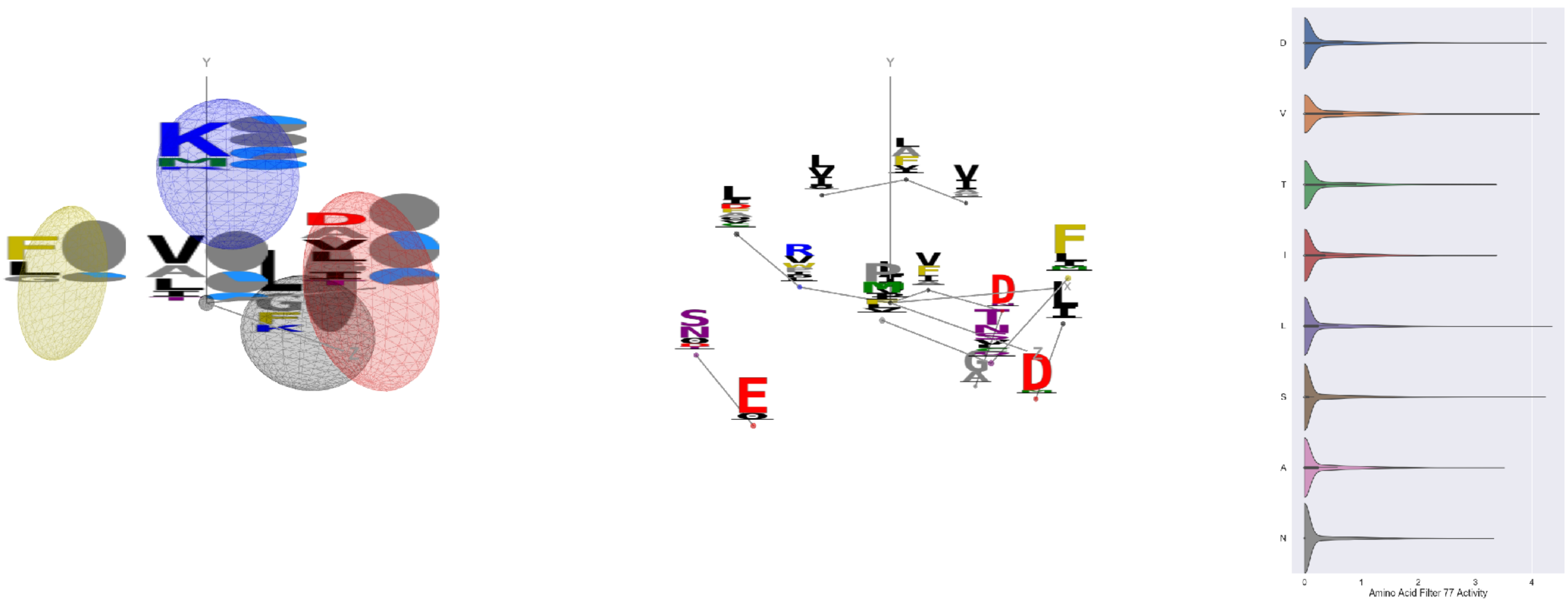

Filter 77
$R_{\text{pred}} = -0.17$
Percent Active = 27
Pooling attention 0.31
Pooling feature 1.37
Top activators
1jdw:A:259 L
1wsr:A:77 D
2d73:A:267 S
4nm9:B:274 V
1iwa:A:202 D
2gy7:B:101 Y
5hu6:C:392 V
3ucq:A:284 D
4y97:A:598 Y
4v81:A:223 Q

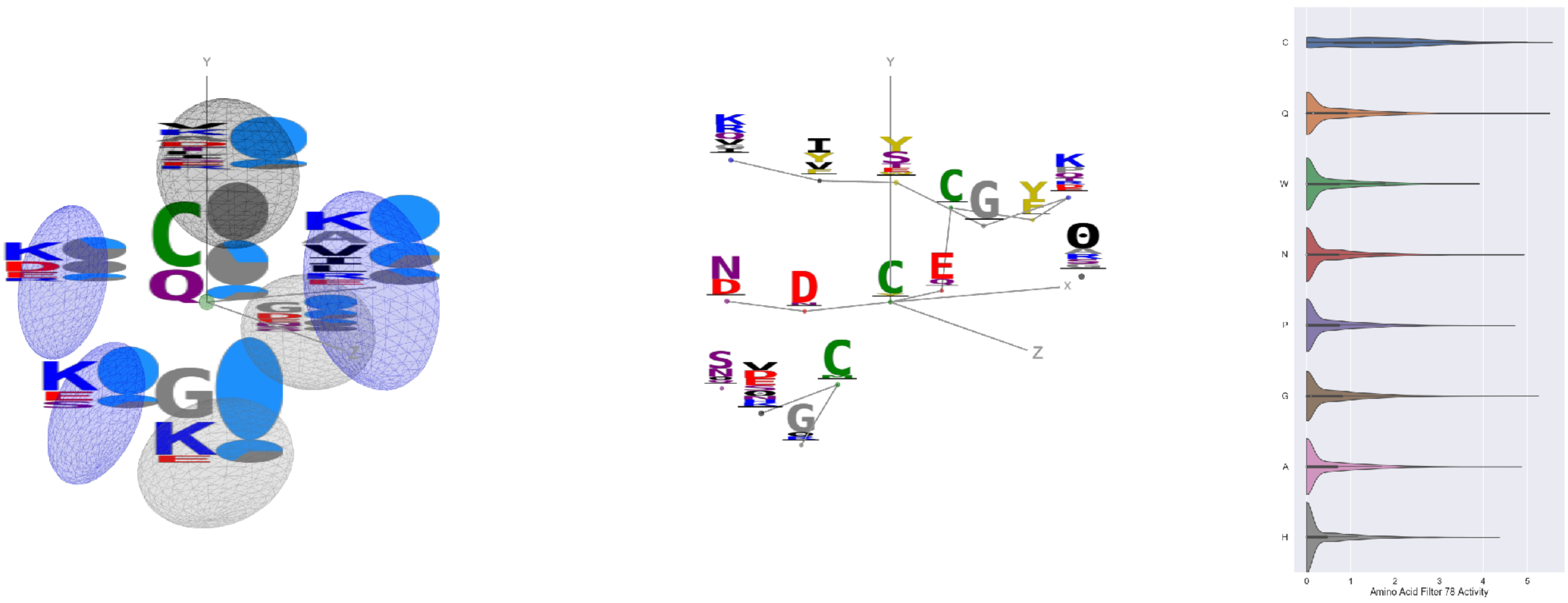

Filter 78
$R_{\text{pred}} = -0.13$
Percent Active = 36
Pooling attention 0.35
Pooling feature 0.37
Top activators
4g1m:B:523 C
3wr9:A:78 Y
1jku:A:45 Q
2hye:B:208 C
1a0j:A:191 C
1ogl:A:58 G
3b8e:A:447 G
3wrx:C:837 C
3s9m:C:283 Q
4e0s:B:137 C

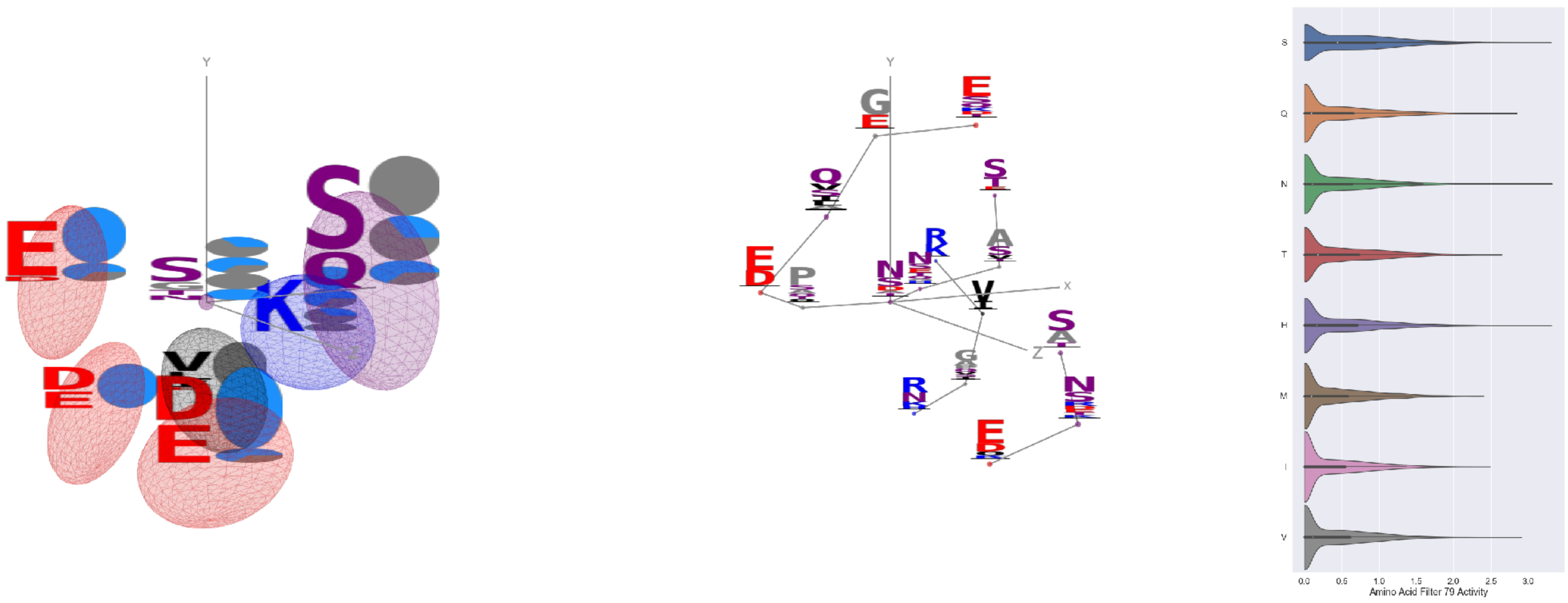

Filter 79
$R_{\text{pred}} = -0.09$
Percent Active = 37
Pooling attention 0.06
Pooling feature 0.51
Top activators
1jqj:A:296 N
4itr:A:3717 H
4c2m:A:485 S
1r6m:A:43 S
3o47:A:199 G
4l8n:A:223 S
4asi:C:1729 G
2cge:A:616 N
4a7z:A:422 E
2iwh:B:703 G

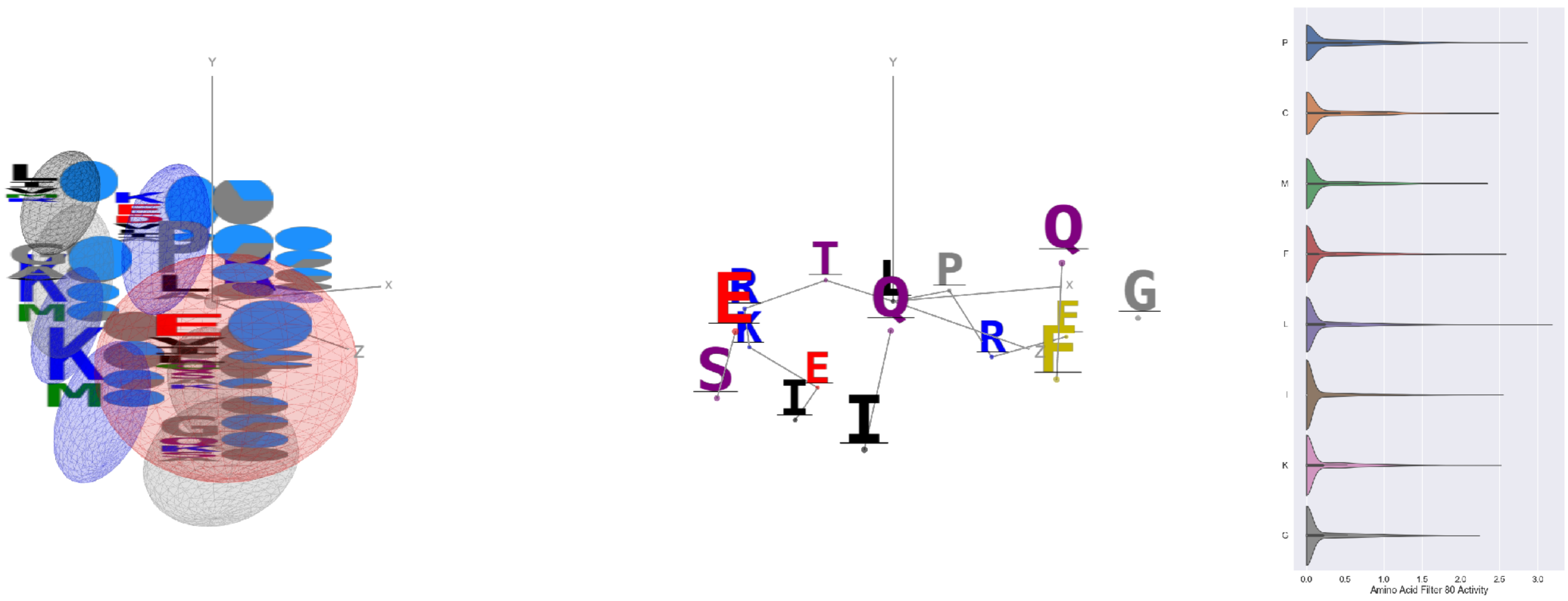


Filter 80
$R_{\text{pred}} = 0.09$
Percent Active = 30
Pooling attention 0.30
Pooling feature 3.01
Top activators
2iwh:B:869 L
4e0s:B:573 P
3bof:A:238 P
1umg:A:156 L
1bgv:A:132 P
1e7n:A:11 F
4f2n:B:71 N
3o47:A:241 L
1cm5:A:559 I
4a7z:A:477 P

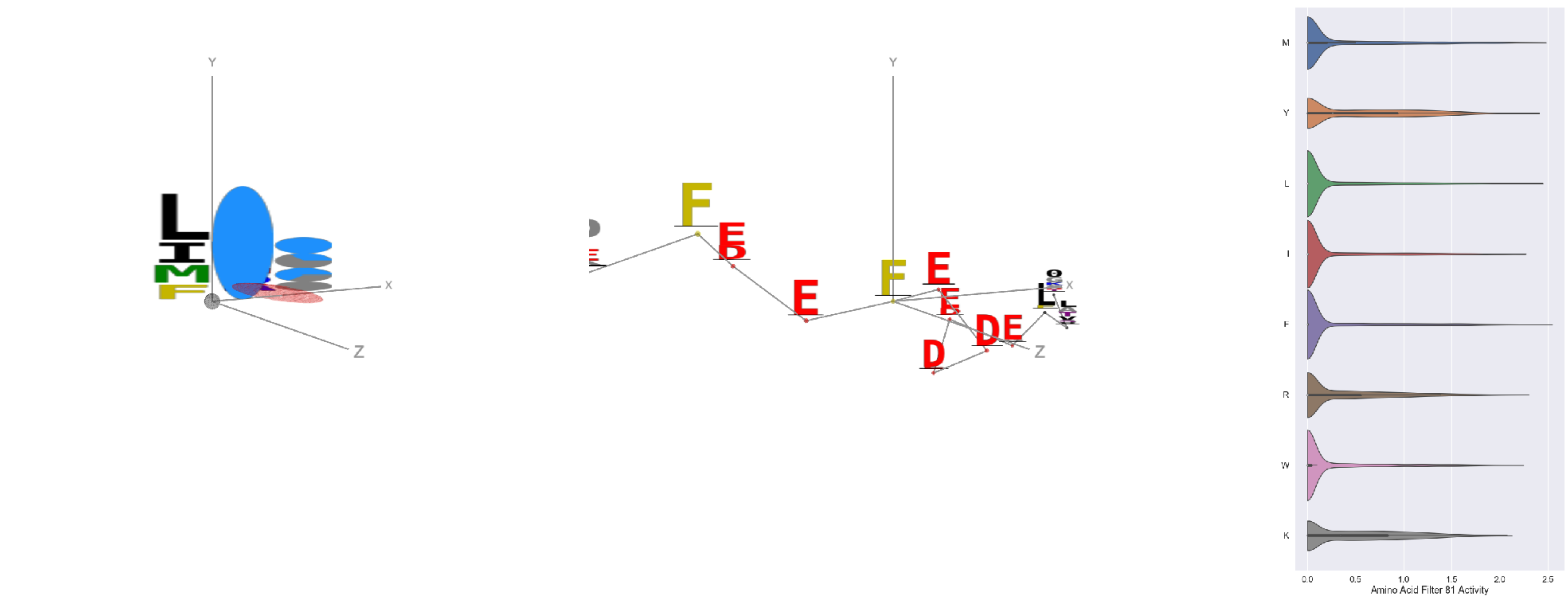


Filter 81
$R_{\text{pred}} = 0.47$
Percent Active = 43
Pooling attention 0.35
Pooling feature 0.77
Top activators
3t5v:C:35 F
3t5v:C:38 F
3n6r:G:693 M
1jyo:F:115 L
3te3:A:717 L
1mp9:A:196 Y
2h4o:A:57 L
1fx0:B:292 M
2pg1:L:114 V
2b0o:F:686 Y

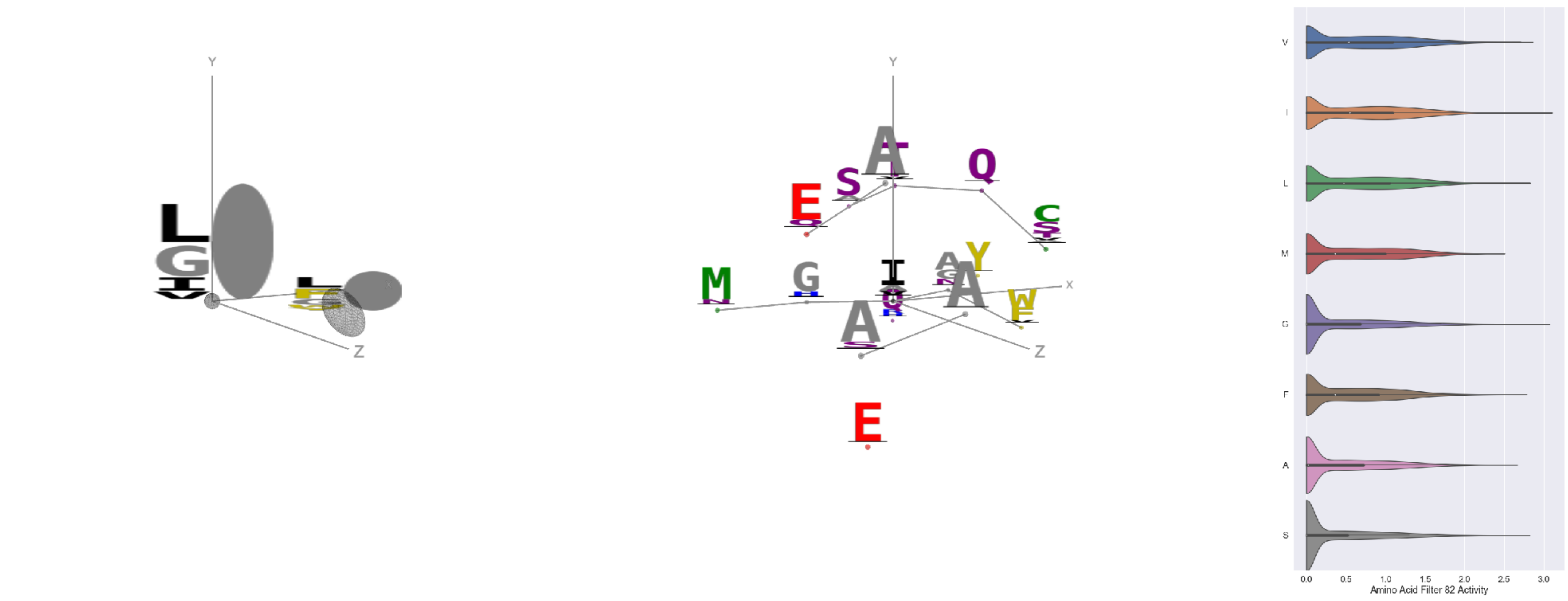


Filter 82
$R_{\text{pred}} = -0.29$
Percent Active = 41
Pooling attention 0.42
Pooling feature 1.00
Top activators
1e1c:A:290 I
1adj:A:286 G
1bgv:A:168 V
1lk5:A:120 I
4uzi:A:39 L
3rby:A:27 S
1ko6:A:777 F
2z67:A:164 Q
4a7z:A:37 V
3igh:X:414 G

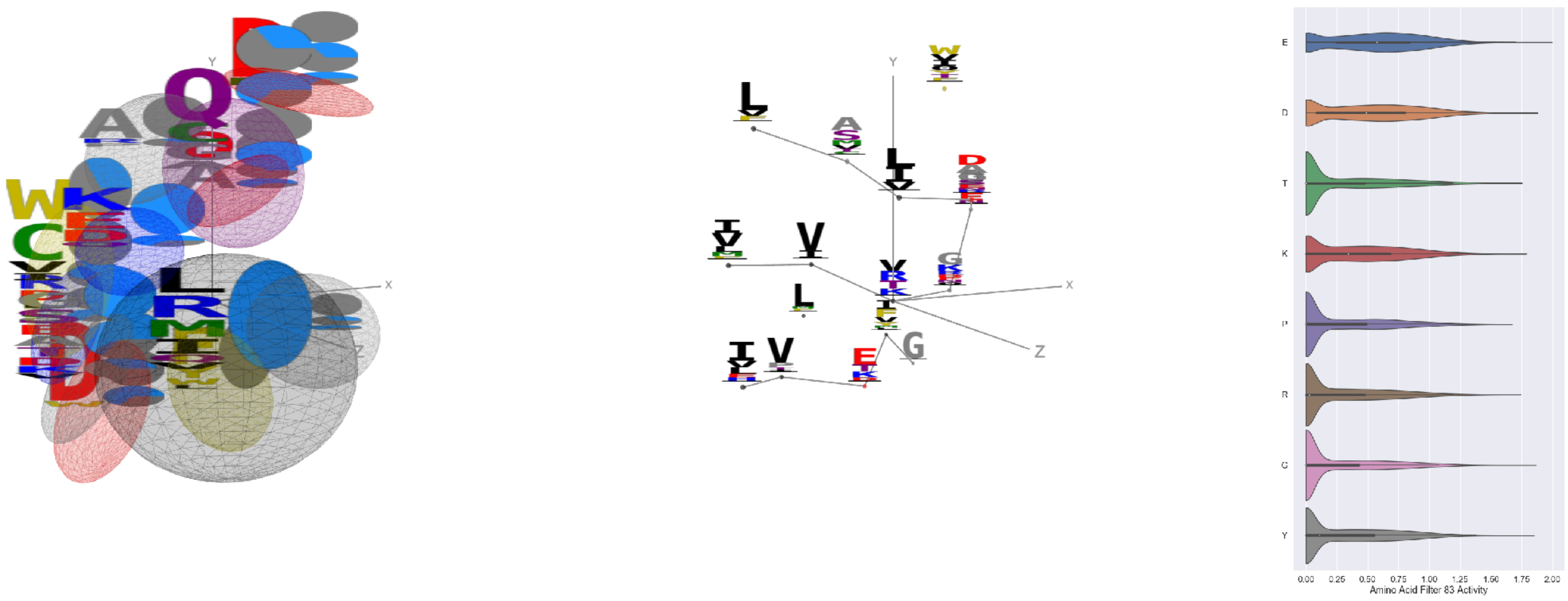


Filter 83
$R_{\text{pred}} = 0.01$
Percent Active = 43
Pooling attention 0.04
Pooling feature 0.96
Top activators
1jdw:A:369 V
1vqv:B:284 E
1k72:B:113 V
3l0o:A:84 D
5kqv:E:139 S
4u4c:A:893 G
4rcn:B:375 E
4cvw:A:6 Y
4asi:C:2124 E
4e0s:B:885 E

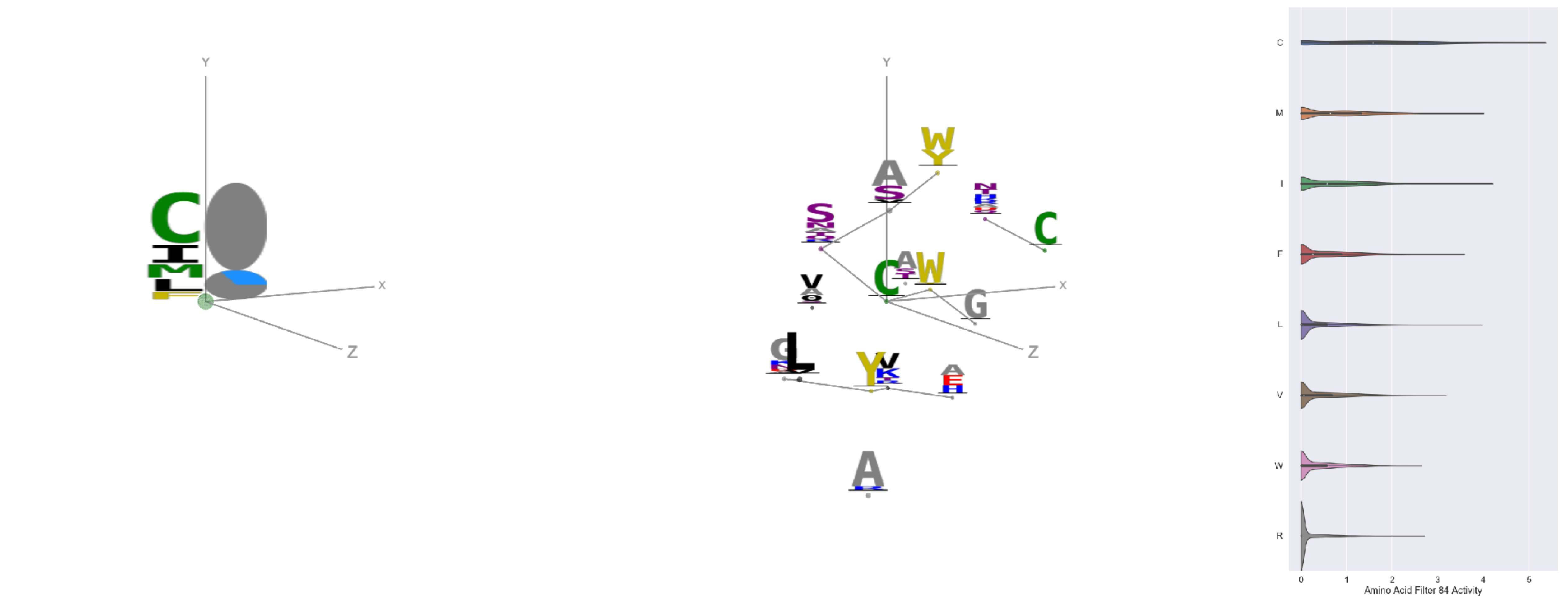

Filter 84
$R_{\text{pred}} = -0.12$
Percent Active = 24
Pooling attention 0.11
Pooling feature 0.50
Top activators
1hlq:A:68 C
3noy:B:300 C
1hlq:A:54 C
3d00:A:58 C
2iwh:B:273 C
2iwh:B:690 C
2wsc:B:559 C
2qn5:B:75 C
3tsy:A:187 C
3qzy:B:155 C

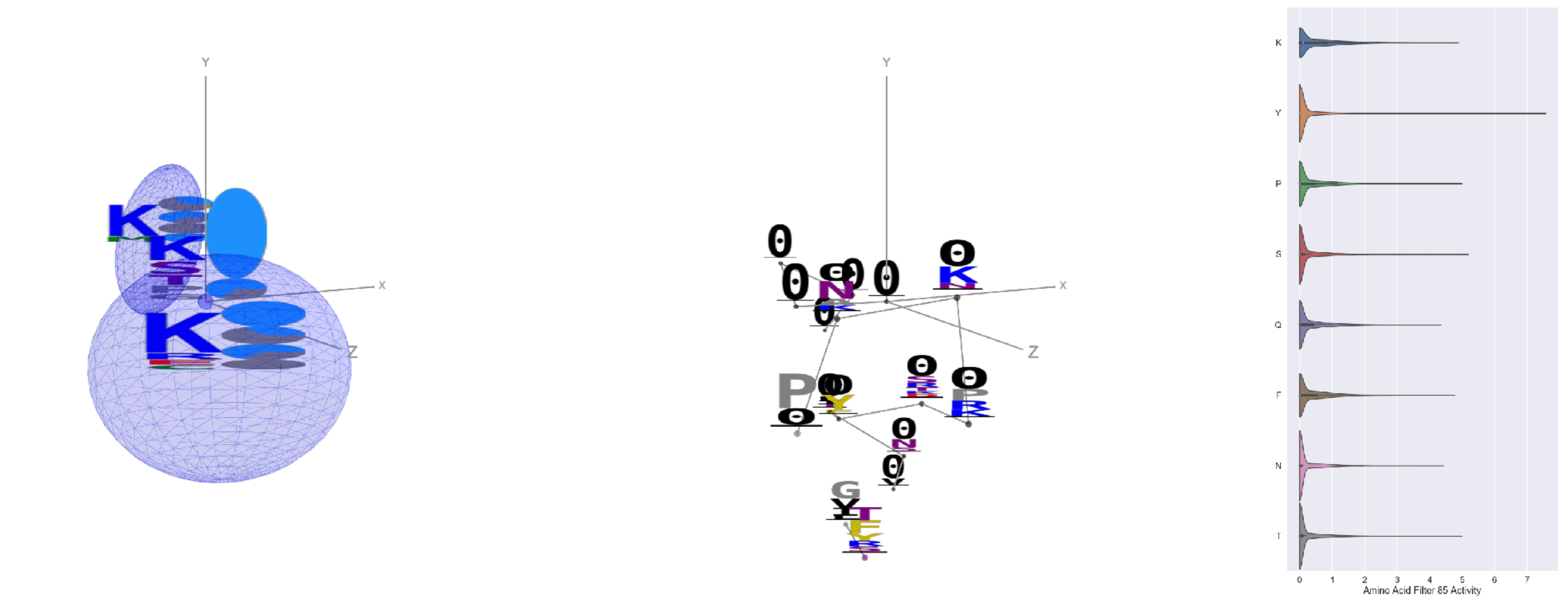

Filter 85
$R_{\text{pred}} = -0.10$
Percent Active = 27
Pooling attention 0.01
Pooling feature 0.18
Top activators
1ewy:A:1 Y
3o47:A:5 Y
4d10:K:3 Y
4pk7:A:83 Y
4c8q:H:471 Y
1jb0:K:20 I
4f52:E:1 Y
3dlb:A:3 Y
3d00:A:127 D
1bxr:A:9 S

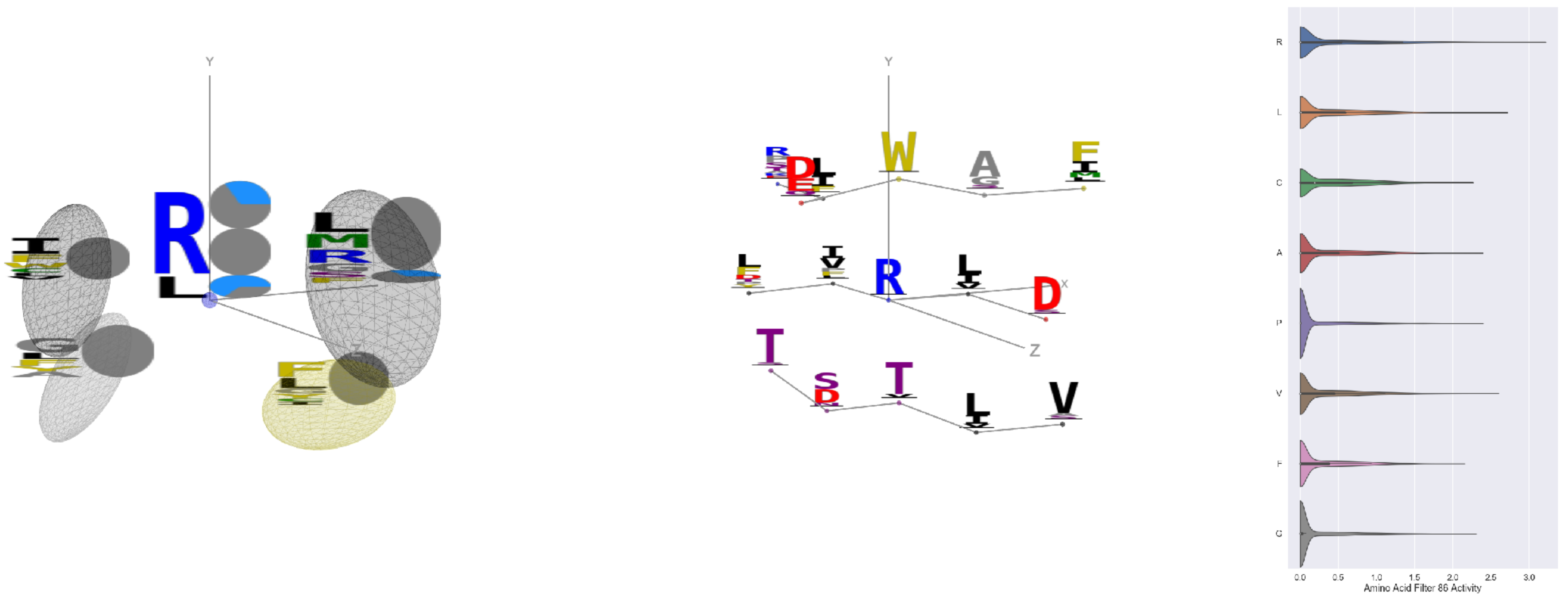

Filter 86
$R_{\text{pred}} = -0.24$
Percent Active = 28
Pooling attention 0.49
Pooling feature 1.61
Top activators
3rby:A:36 R
1iri:A:104 R
1w36:B:800 R
4rcn:B:228 R
1gyt:A:356 R
3g8q:A:65 R
1sl0:A:452 R
2iwh:B:695 R
1b8a:A:368 R
2d73:A:289 R

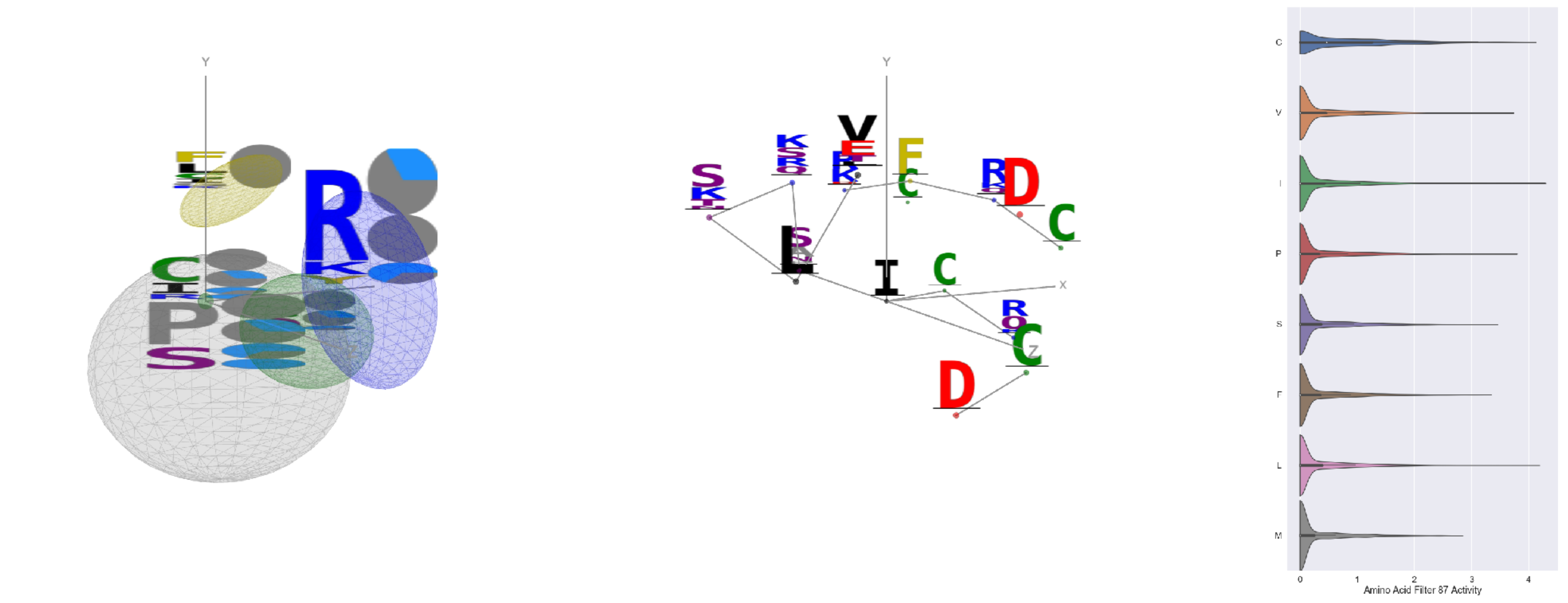

Filter 87
$R_{\text{pred}} = -0.05$
Percent Active = 32
Pooling attention 0.27
Pooling feature 0.95
Top activators
4e0s:B:131 I
4kzx:E:114 I
2hd0:A:217 L
4e0s:B:152 C
4uzi:A:324 Y
2np9:A:335 R
3tsy:A:55 I
1p35:A:21 I
1fx0:B:69 L
3g8q:A:231 A

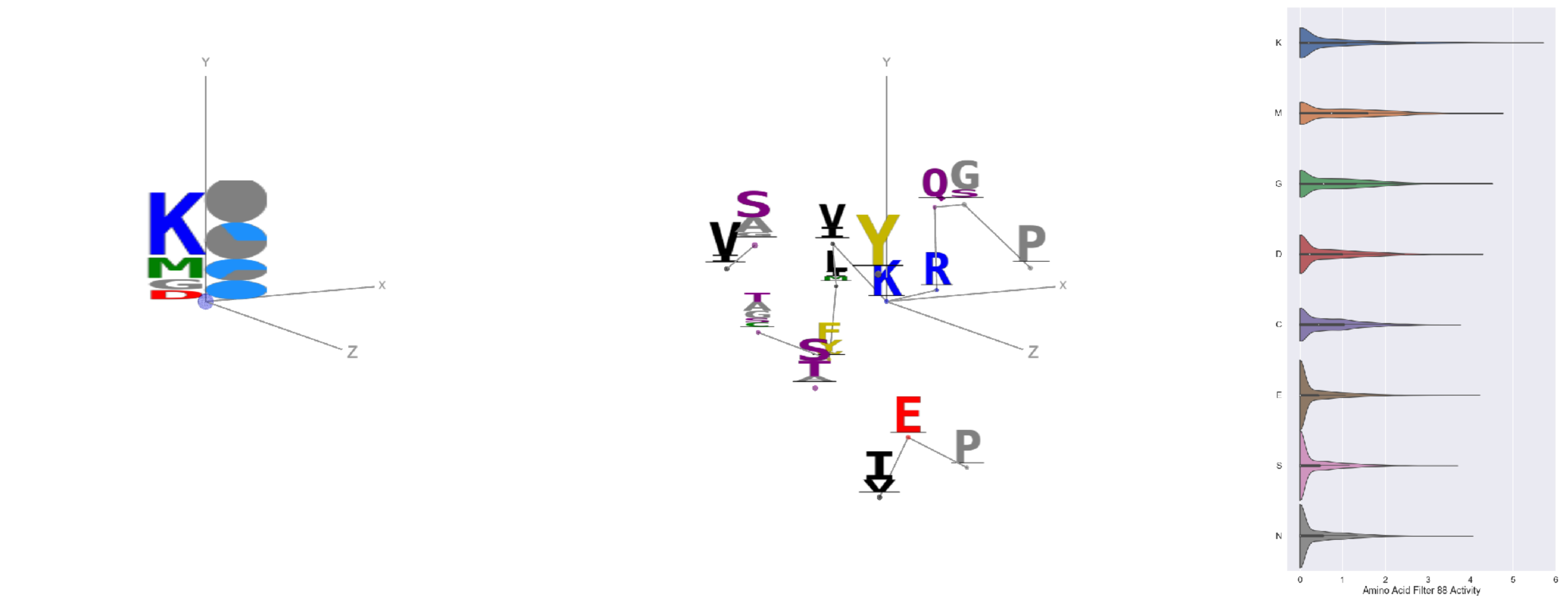

Filter 88
$R_{\mathrm{pred}} = -0.11$
Percent Active = 37
Pooling attention 0.36
Pooling feature 0.68
Top activators
4gfh:A:705 K
1qvr:A:204 K
1r7r:A:251 K
3wb2:A:86 K
1r7r:A:524 K
4nmn:A:212 K
1e1c:A:300 K
4cjn:A:406 K
1lk5:A:7 K
1w36:B:29 K

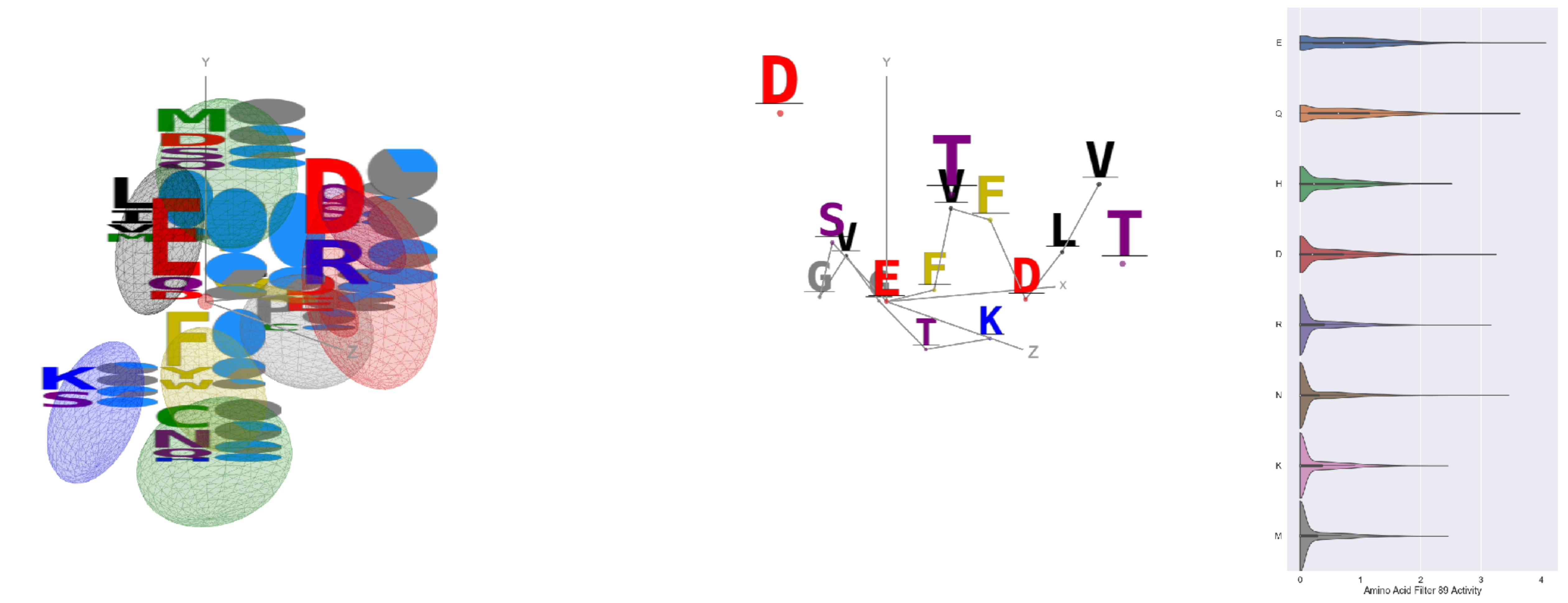

Filter 89
$R_{\mathrm{pred}} = 0.12$
Percent Active = 32
Pooling attention 0.04
Pooling feature 0.07
Top activators
2iwh:B:513 E
3g8q:A:179 E
1a92:A:18 E
4rp5:A:13 E
2wqi:A:366 E
2iwh:B:15 Q
4l2w:D:850 Q
3te3:A:716 Q
2fq1:A:92 N
2pnv:A:505 E

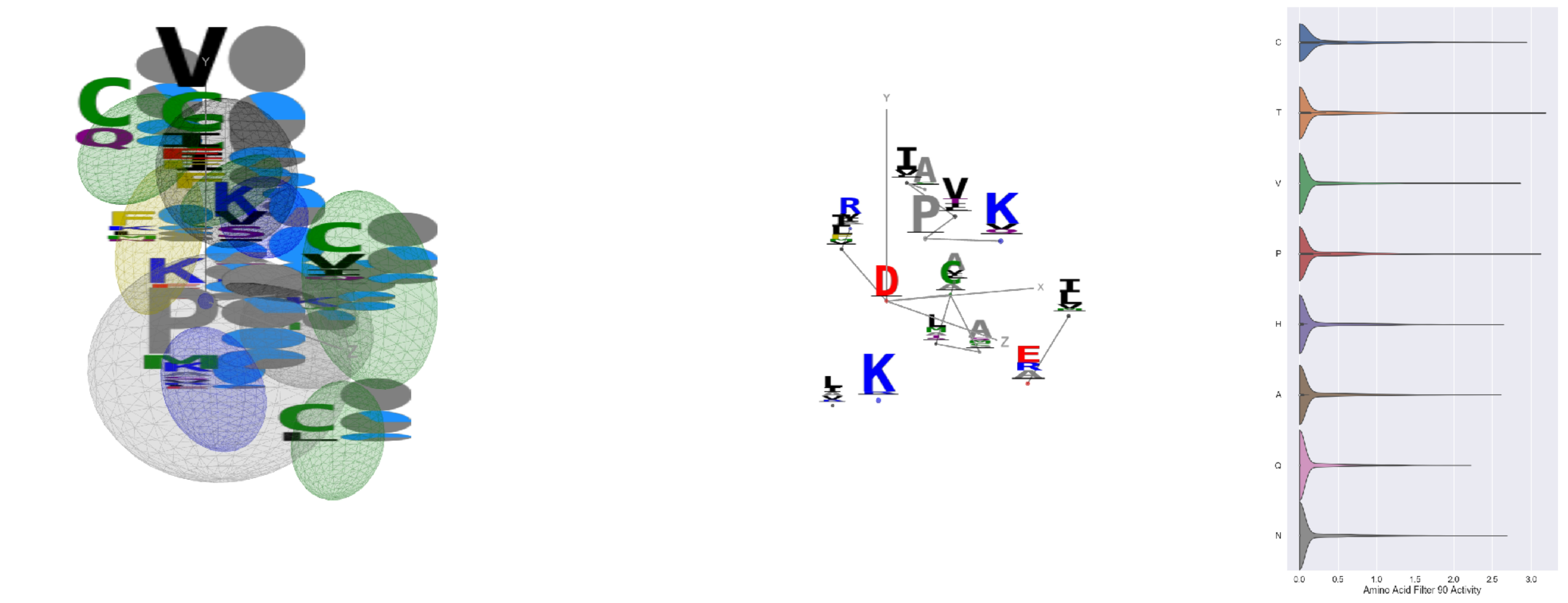

Filter 90
$R_{\mathrm{pred}} = 0.01$
Percent Active = 23
Pooling attention 0.91
Pooling feature 5.03
Top activators
1dci:A:182 D
1h8p:A:59 G
1h8p:A:107 G
3g8q:A:5 L
3tsy:A:832 T
4e0s:B:506 P
4e0s:B:131 I
3tsy:A:614 P
4gu0:A:147 C
5wy5:A:207 C

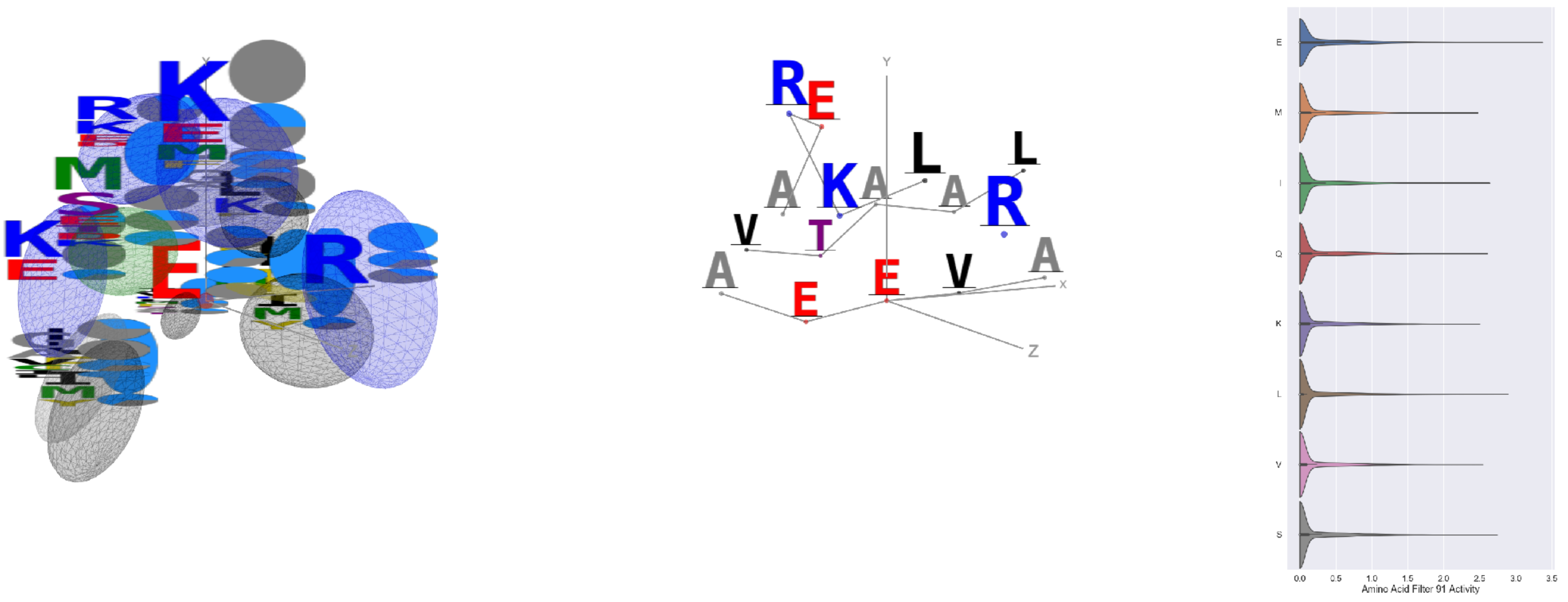

Filter 91
$R_{\mathrm{pred}} = 0.10$
Percent Active = 26
Pooling attention 0.72
Pooling feature 3.64
Top activators
3g8q:A:38 E
1k90:C:379 A
3o47:A:232 E
4qiw:A:15 L
3g8q:A:43 E
3ax7:A:338 A
3hgi:A:84 E
4kzx:E:29 P
2f8x:C:185 S
3sf5:C:119 E

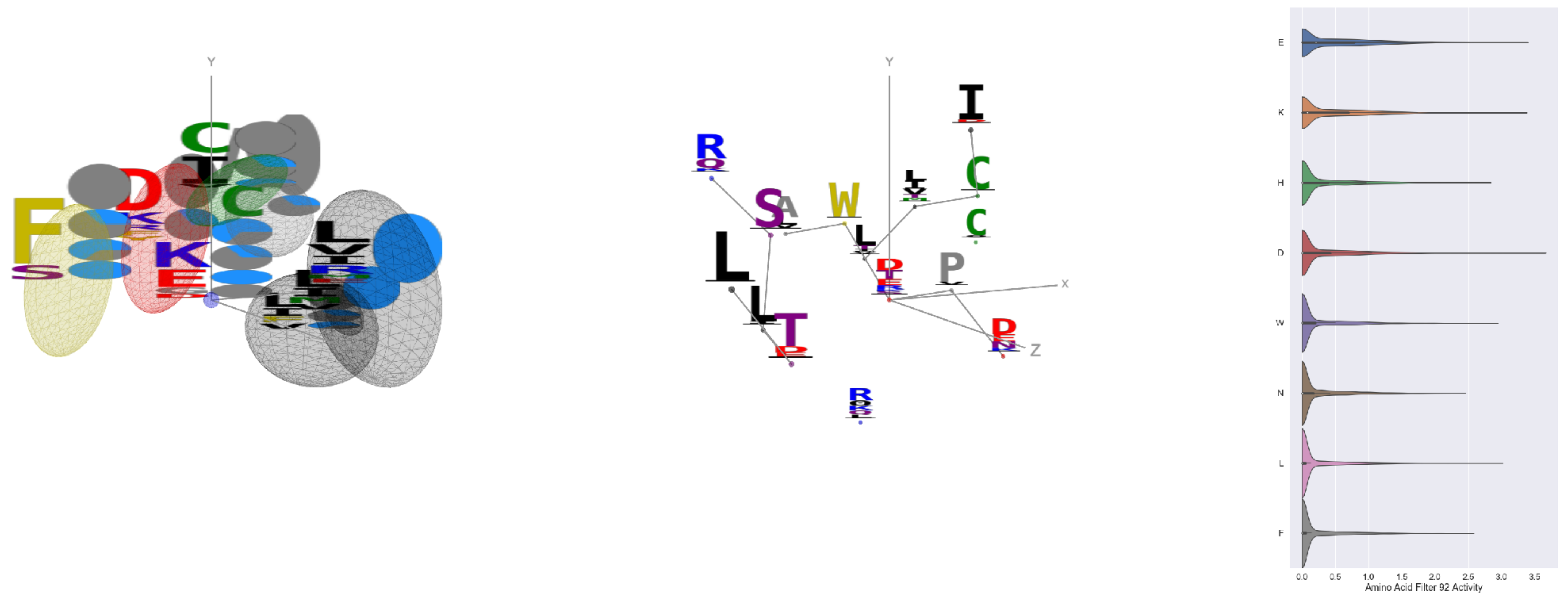


Filter 92
$R_{\mathrm{pred}} = -0.03$
Percent Active = 29
Pooling attention 0.26
Pooling feature 1.08
Top activators
2b0o:F:450 D
4r3z:B:602 Y
3tsy:A:935 E
1izl:D:178 I
2iwh:B:314 K
2iwh:B:307 E
2iwh:B:405 E
3irm:A:295 E
1izl:D:180 R
2b5d:X:185 E

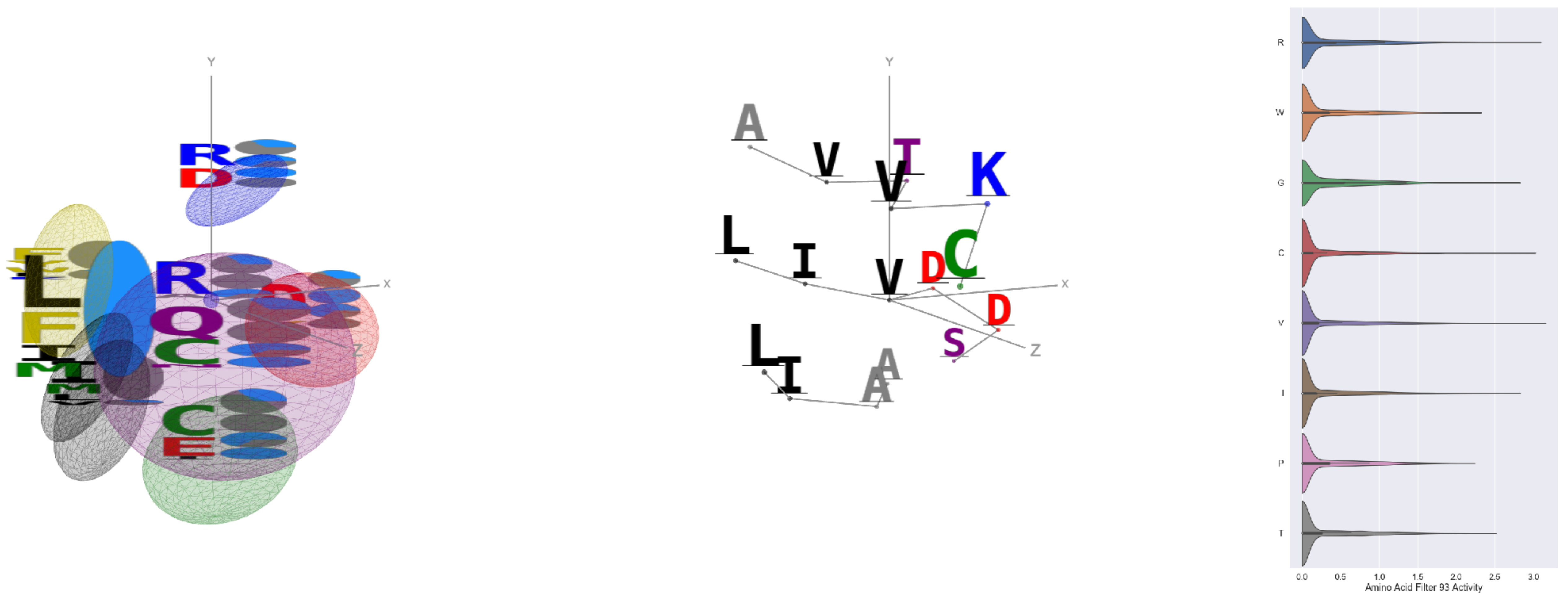


Filter 93
$R_{\mathrm{pred}} = 0.03$
Percent Active = 31
Pooling attention 0.44
Pooling feature 1.82
Top activators
3tsy:A:272 V
3g8q:A:189 R
4e0s:B:326 L
1h8p:A:69 C
3tsy:A:49 R
3tsy:A:202 R
1jb0:K:21 I
2gy7:B:305 G
3tsy:A:839 G
4u4c:A:243 R

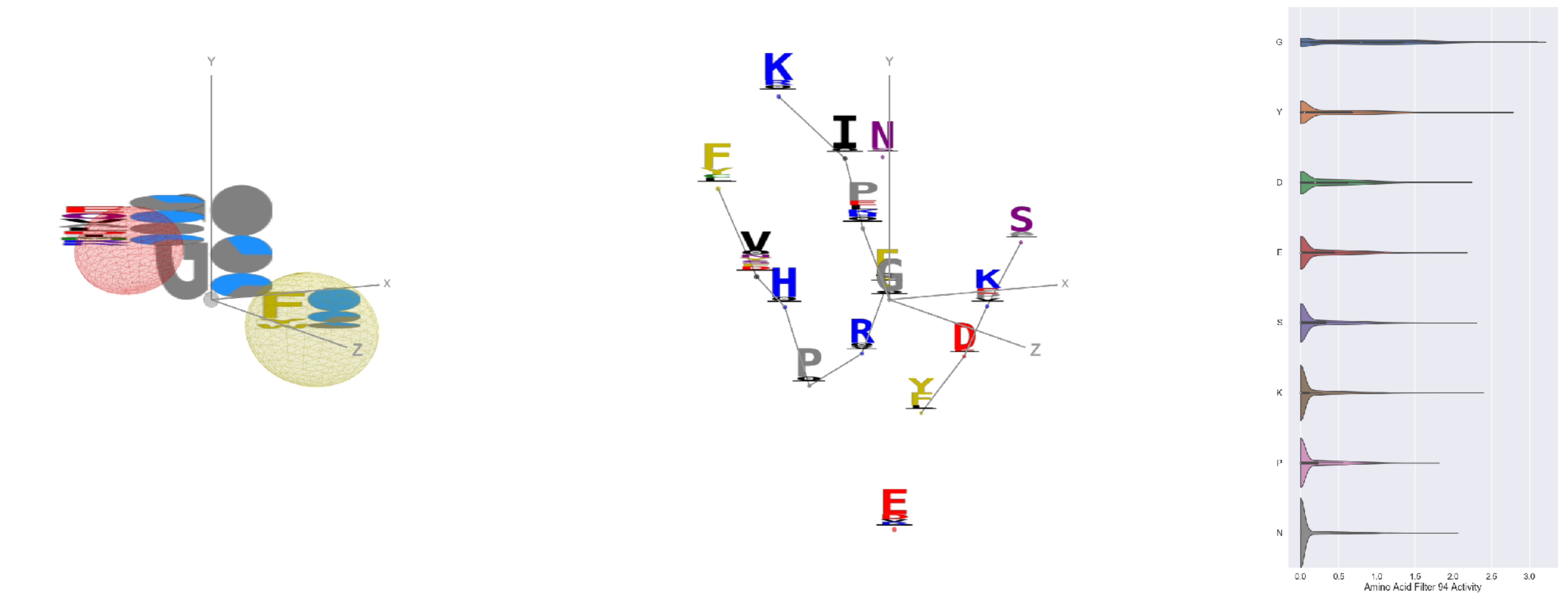


Filter 94
$R_{\mathrm{pred}} = 0.02$
Percent Active = 31
Pooling attention 0.07
Pooling feature 0.47
Top activators
4jds:A:314 G
4asi:C:1830 G
4v8p:BF:231 G
2iwh:B:323 G
1vqv:B:259 G
3pq1:A:231 G
1w36:B:28 G
4v8p:BC:382 Y
1wnl:A:47 G
3hzu:A:115 G

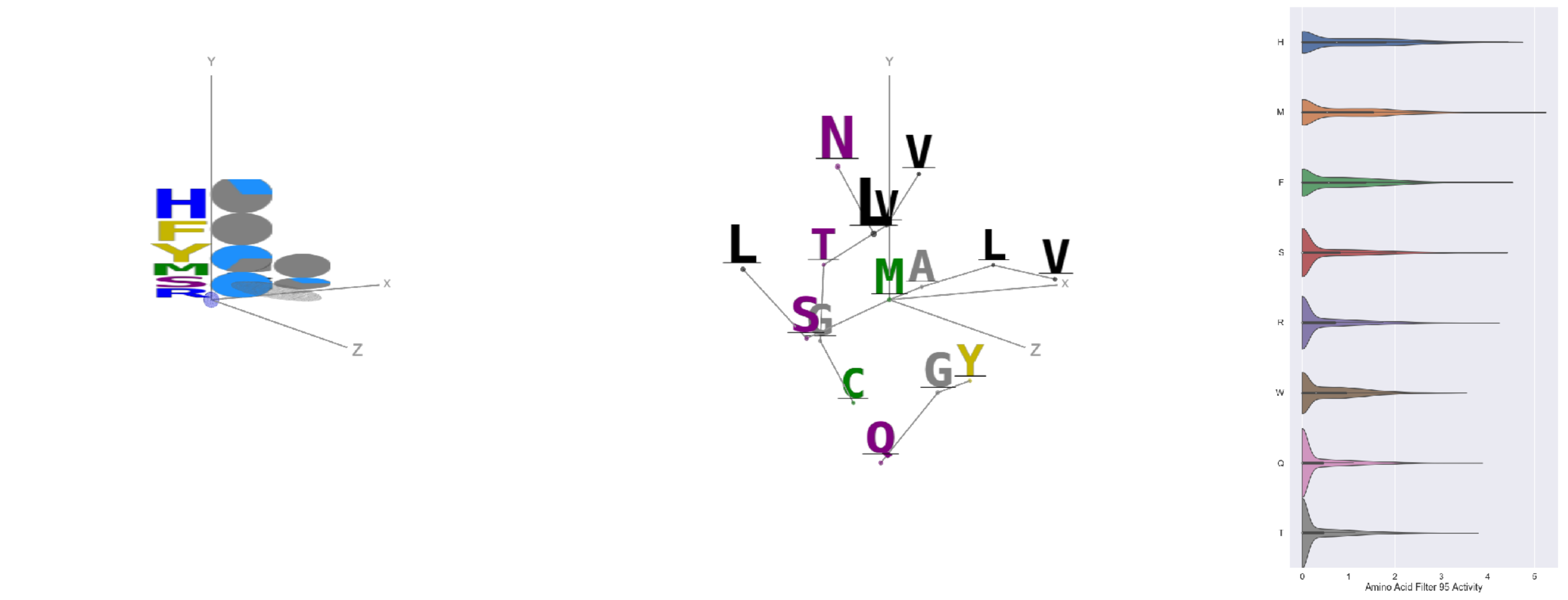


Filter 95
$R_{\mathrm{pred}} = -0.06$
Percent Active = 24
Pooling attention 0.27
Pooling feature 0.91
Top activators
3tsy:A:388 M
1umg:A:274 M
1ii7:A:208 H
4ddg:A:1270 Y
1iri:A:100 H
3nh5:A:116 H
4mex:C:586 F
1c3r:A:170 H
4hg6:A:276 H
4n4n:A:267 H

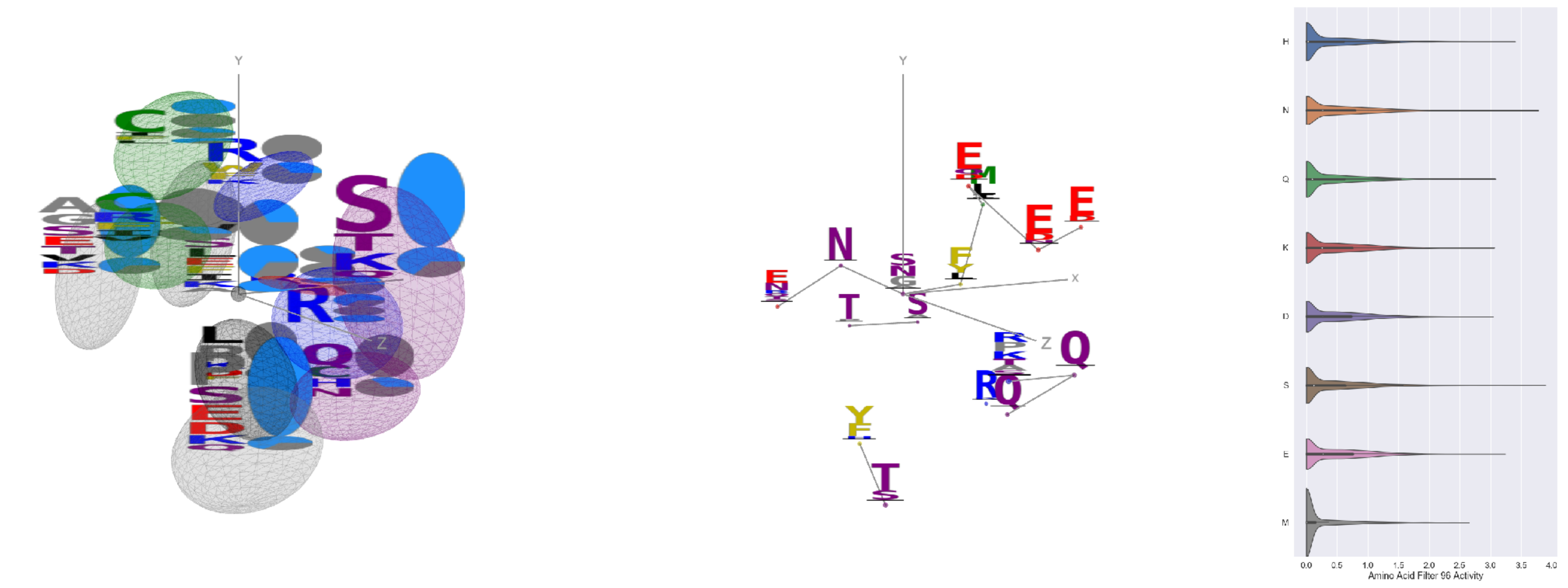

Filter 96
$R_{pred} = 0.01$
Percent Active = 41
Pooling attention 0.01
Pooling feature 0.17
Top activators
1srq:A:212 S
4ddg:A:77 N
2iwh:B:693 S
2f8x:C:185 S
2zbk:B:315 S
3qsq:A:533 H
4g1m:B:74 Y
1fx0:B:143 S
1cmx:A:12 E
4ued:B:69 R

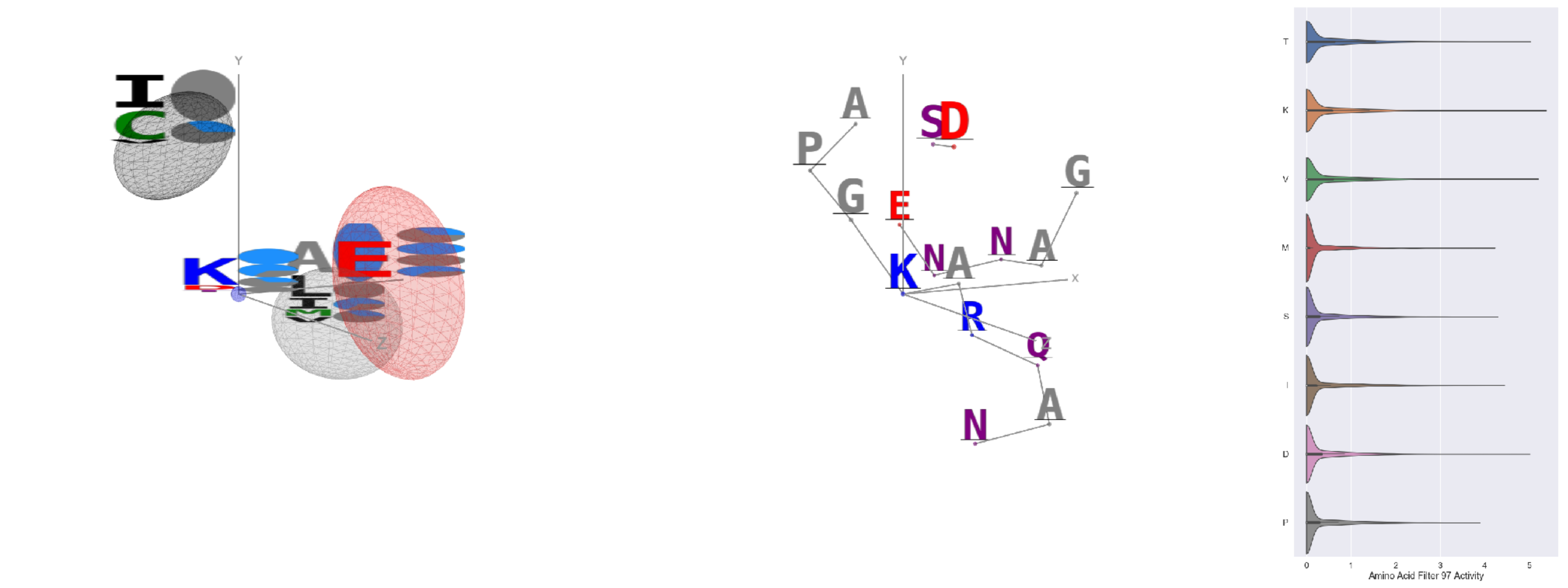

Filter 97
$R_{pred} = 0.04$
Percent Active = 28
Pooling attention 0.24
Pooling feature 0.30
Top activators
3tsy:A:601 K
1j5w:B:129 V
3i5d:B:316 K
4wj3:B:130 K
3n6r:G:397 E
1n0w:B:1524 F
1vpn:B:307 N
4kzx:L:127 T
4asi:C:2134 D
3tsy:A:300 L

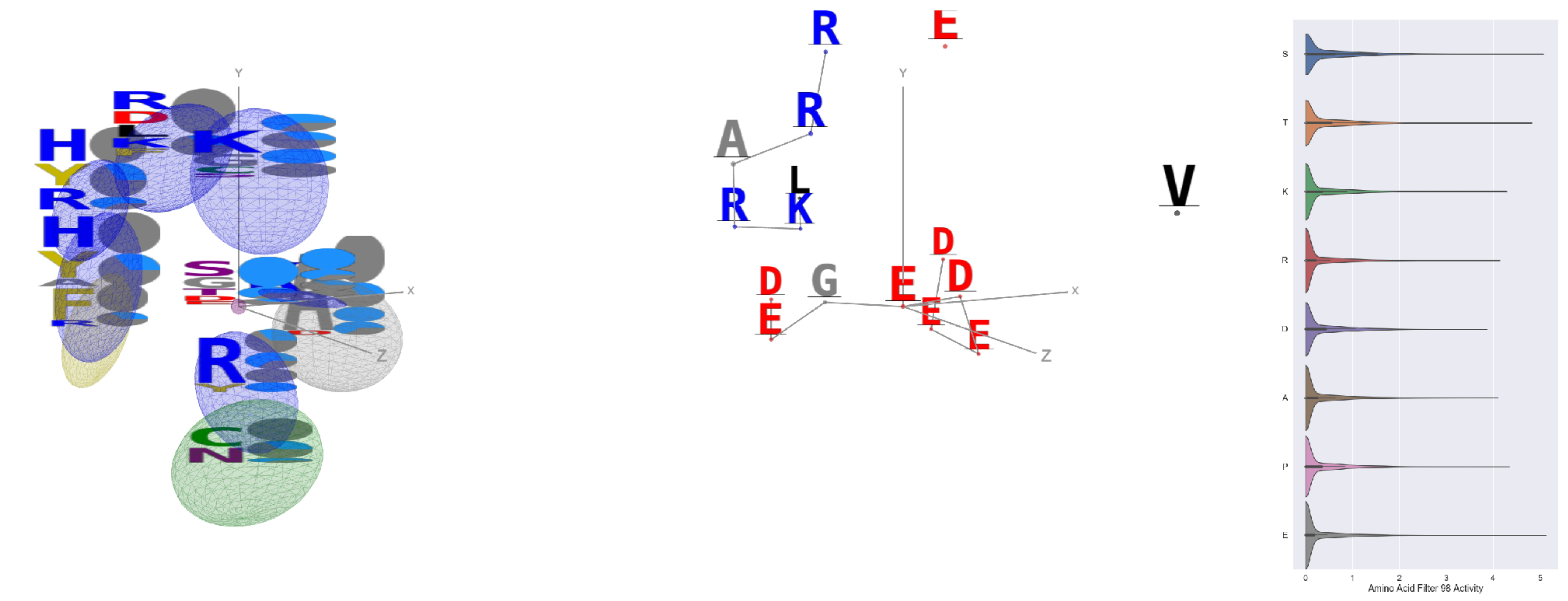

Filter 98
$R_{pred} = -0.12$
Percent Active = 30
Pooling attention 0.23
Pooling feature 2.04
Top activators
2iwh:B:426 E
4nmn:A:356 S
5uqy:A:152 T
1w99:A:290 E
3tsy:A:401 S
3tsy:A:806 S
4pfp:A:124 P
4ddg:A:1080 S
1iwa:A:19 Y
3n6r:G:298 K

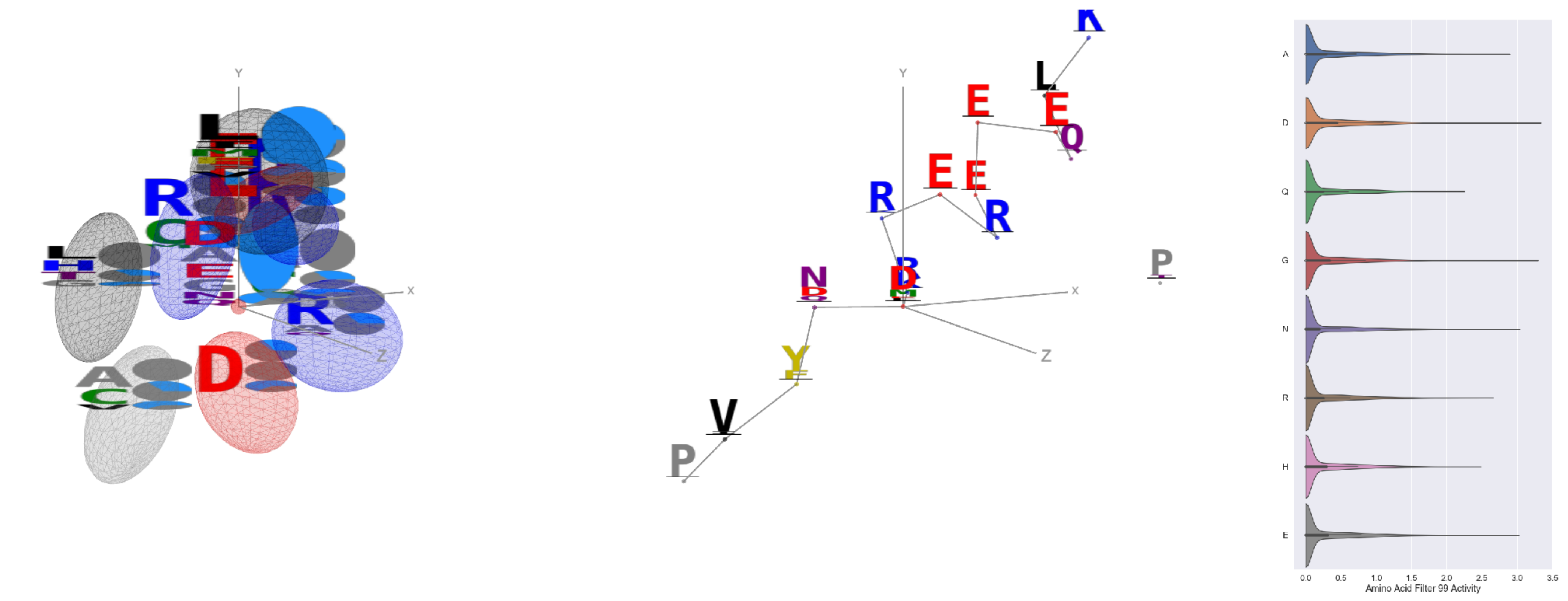

Filter 99
$R_{pred} = 0.19$
Percent Active = 25
Pooling attention 0.27
Pooling feature 0.41
Top activators
4kzx:R:82 D
2o35:B:72 G
3tsy:A:615 D
3tu3:B:306 D
2d4z:B:770 Y
1s4k:A:15 G
4c2m:A:649 N
4uzi:A:448 Y
3l0o:A:218 E
3vie:B:151 E

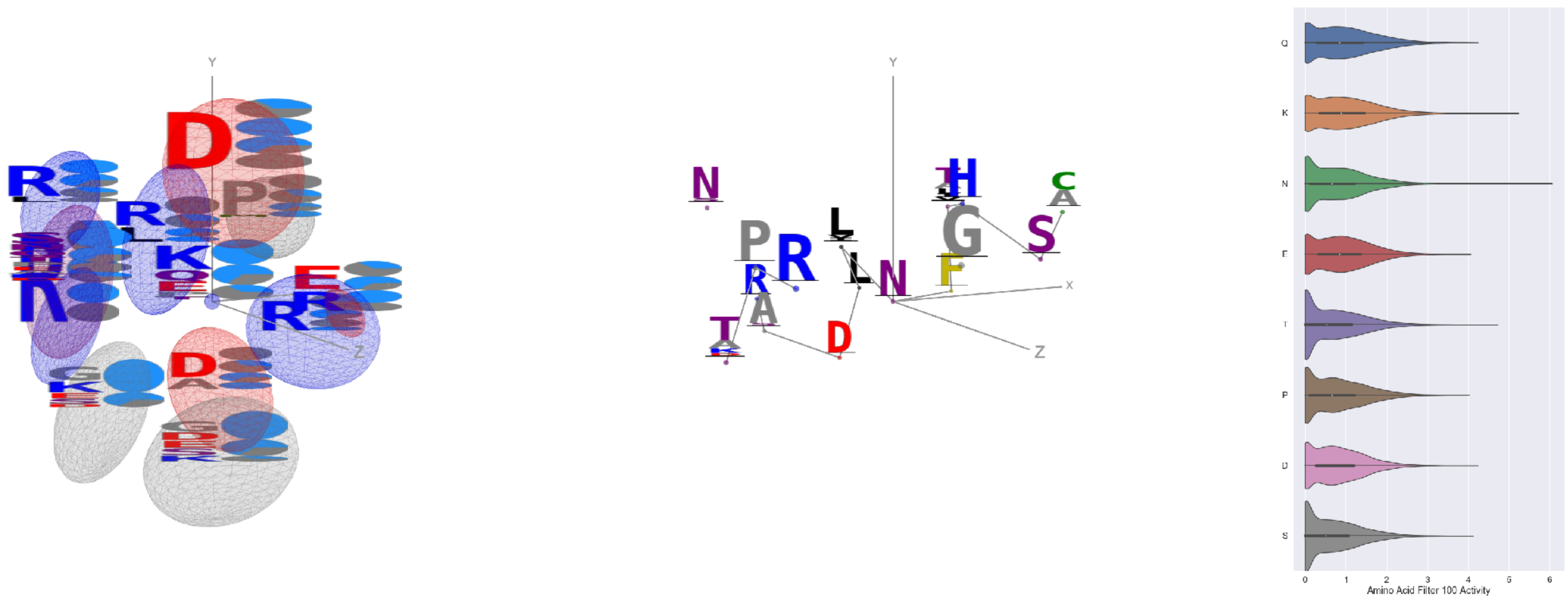


Filter 100
$R_{\text{pred}} = -0.12$
Percent Active = 55
Pooling attention 0.27
Pooling feature 1.40
Top activators
1j5w:B:244 N
3tsy:A:939 K
4oph:A:41 K
1r7r:A:623 T
2iwh:B:453 K
1qha:A:290 K
1umg:A:133 K
1r7r:A:624 N
3n6r:G:355 Q
1mj2:A:38 D

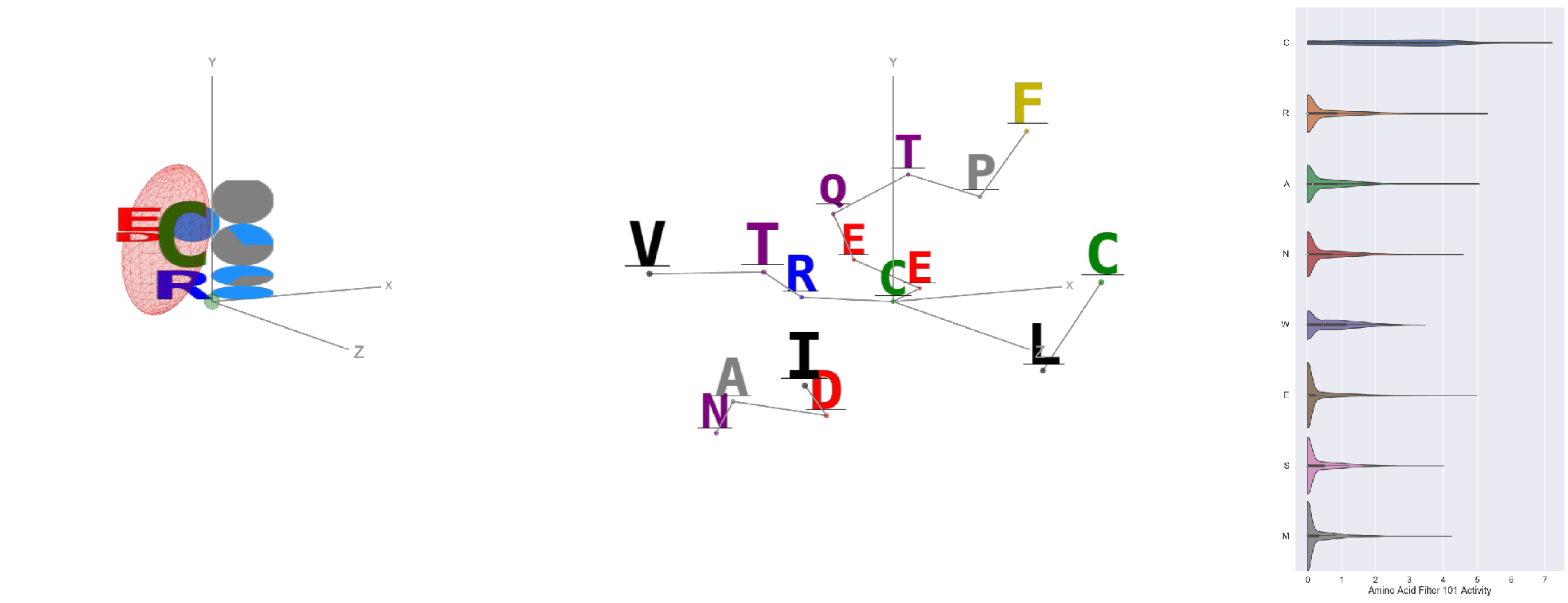


Filter 101
$R_{\text{pred}} = 0.19$
Percent Active = 30
Pooling attention 0.07
Pooling feature 0.49
Top activators
2iwh:B:489 C
4dag:A:182 C
4n4n:A:266 C
4n4n:A:106 C
3tsy:A:433 C
3kov:A:39 C
1b9e:B:7 C
1b9e:B:19 C
3mk7:B:68 C
3w5k:B:159 C

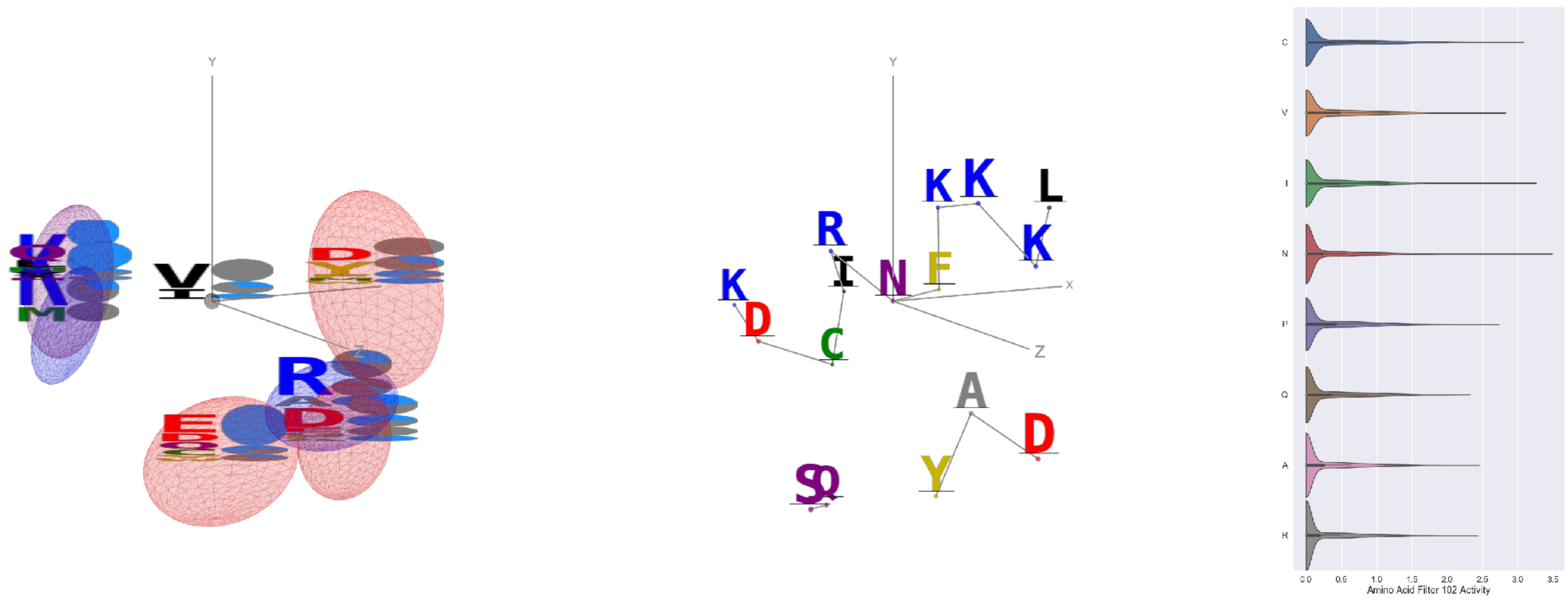


Filter 102
$R_{\text{pred}} = 0.05$
Percent Active = 31
Pooling attention 0.58
Pooling feature 2.56
Top activators
3tsy:A:644 N
4jds:A:213 S
4e0s:B:703 I
1eg1:A:175 C
4pfp:A:474 N
4e0s:B:137 C
4e0s:B:643 V
1j5w:B:21 I
4a7z:A:179 S
1h2i:A:172 L

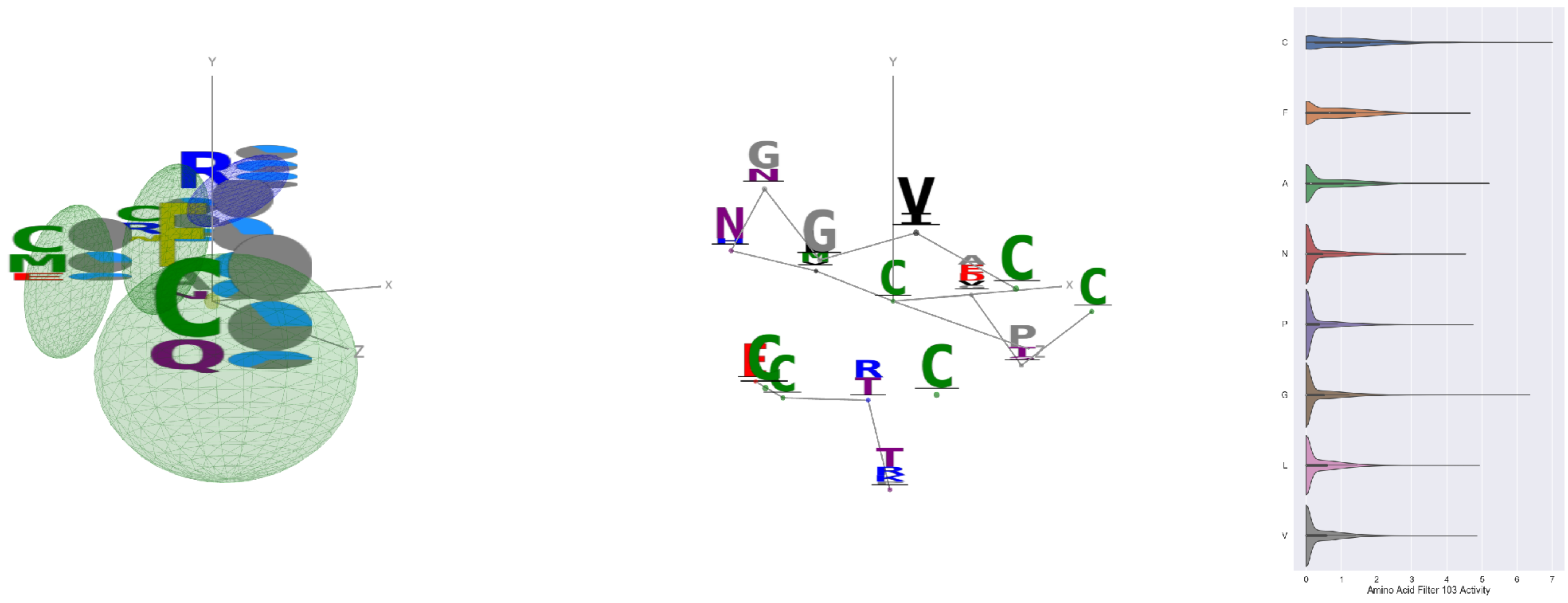


Filter 103
$R_{\text{pred}} = -0.10$
Percent Active = 29
Pooling attention 0.40
Pooling feature 0.74
Top activators
2gy7:B:227 C
1h8p:A:69 C
1h8p:A:24 C
2gy7:B:231 G
4g1m:B:495 C
4e0s:B:505 C
4g1m:B:536 C
2pu9:A:55 C
4g1m:B:575 C
4g1m:B:506 C

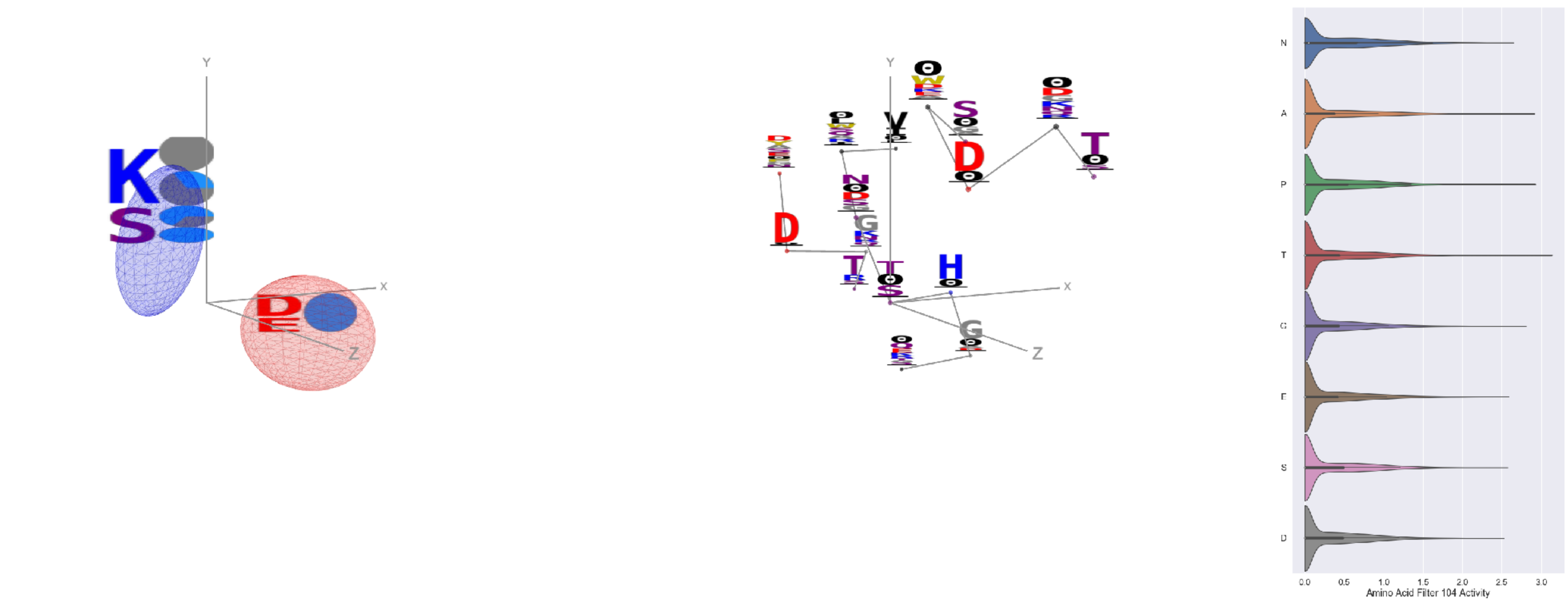


Filter 104
$R_{\text{pred}} = 0.07$
Percent Active = 34
Pooling attention 0.10
Pooling feature 0.27
Top activators
1pgu:B:372 T
3bf0:C:375 P
2iwh:B:704 A
2wk4:B:425 Y
2iwh:B:99 G
1r6m:A:34 G
2iut:A:468 T
2np9:A:398 A
3rzz:A:383 G
1iri:A:361 G

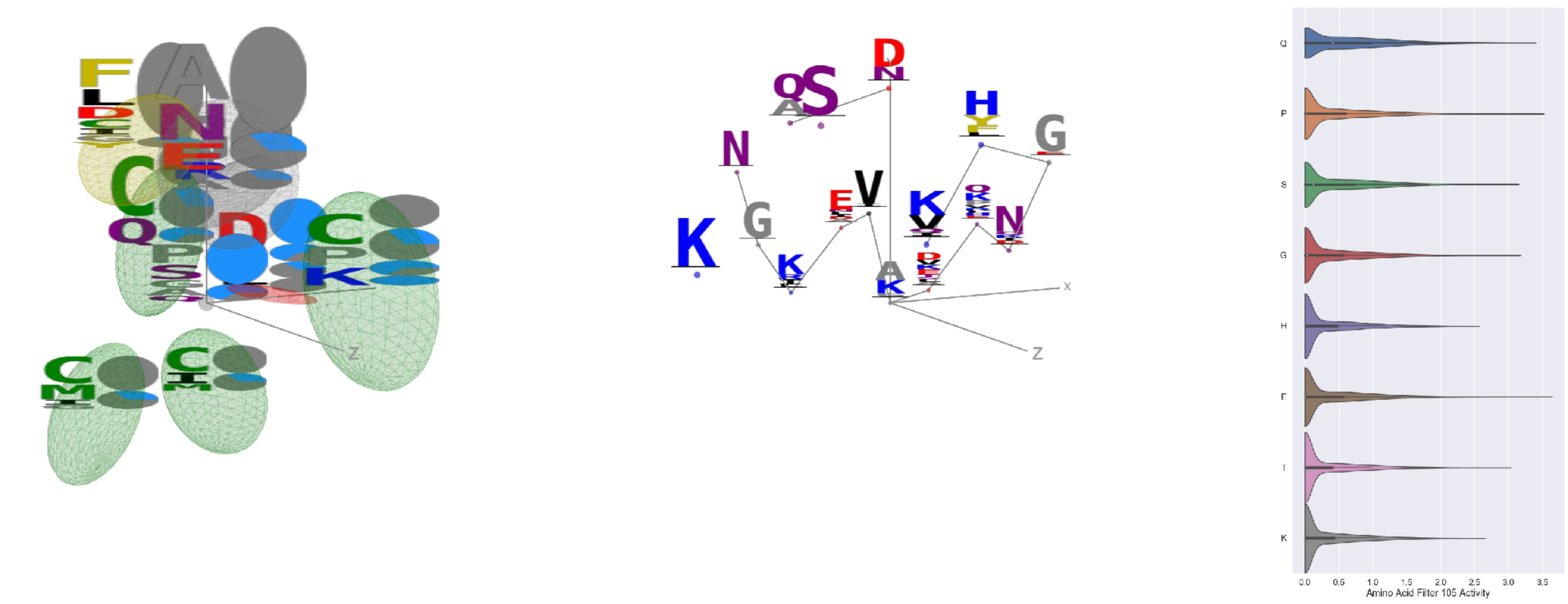


Filter 105
$R_{\text{pred}} = -0.10$
Percent Active = 38
Pooling attention 0.13
Pooling feature 0.44
Top activators
1k90:C:379 A
4v8p:BC:171 E
1bam:A:119 P
5ui2:A:36 Q
2q6b:A:585 A
2gy7:B:279 F
4pfp:A:470 Q
3tsy:A:176 P
1bgv:A:110 Q
4ny2:A:310 Q

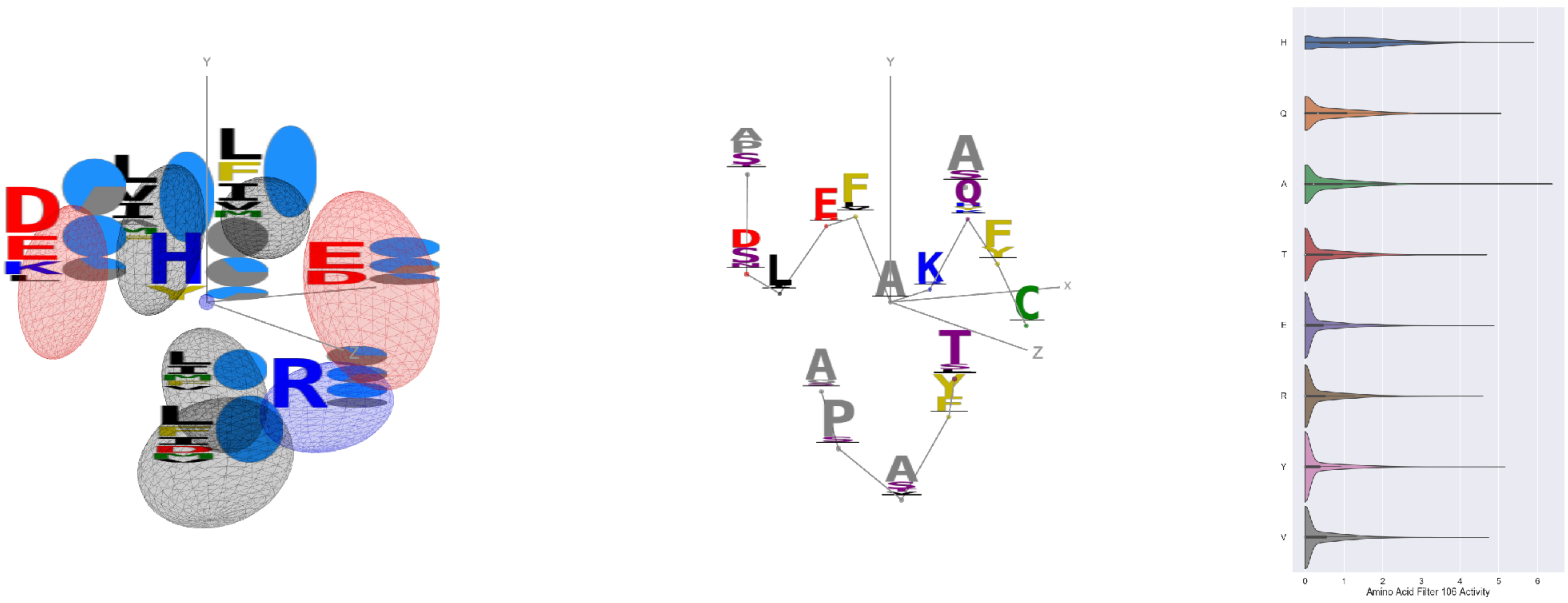


Filter 106
$R_{\text{pred}} = -0.03$
Percent Active = 37
Pooling attention 0.43
Pooling feature 0.34
Top activators
4buj:B:654 A
1ezv:C:197 H
1mj2:A:63 H
1ezv:C:183 H
1dci:A:312 A
3gku:A:179 H
1r7r:A:698 A
1izl:D:331 Y
1w99:A:102 Q
1r7r:A:634 L

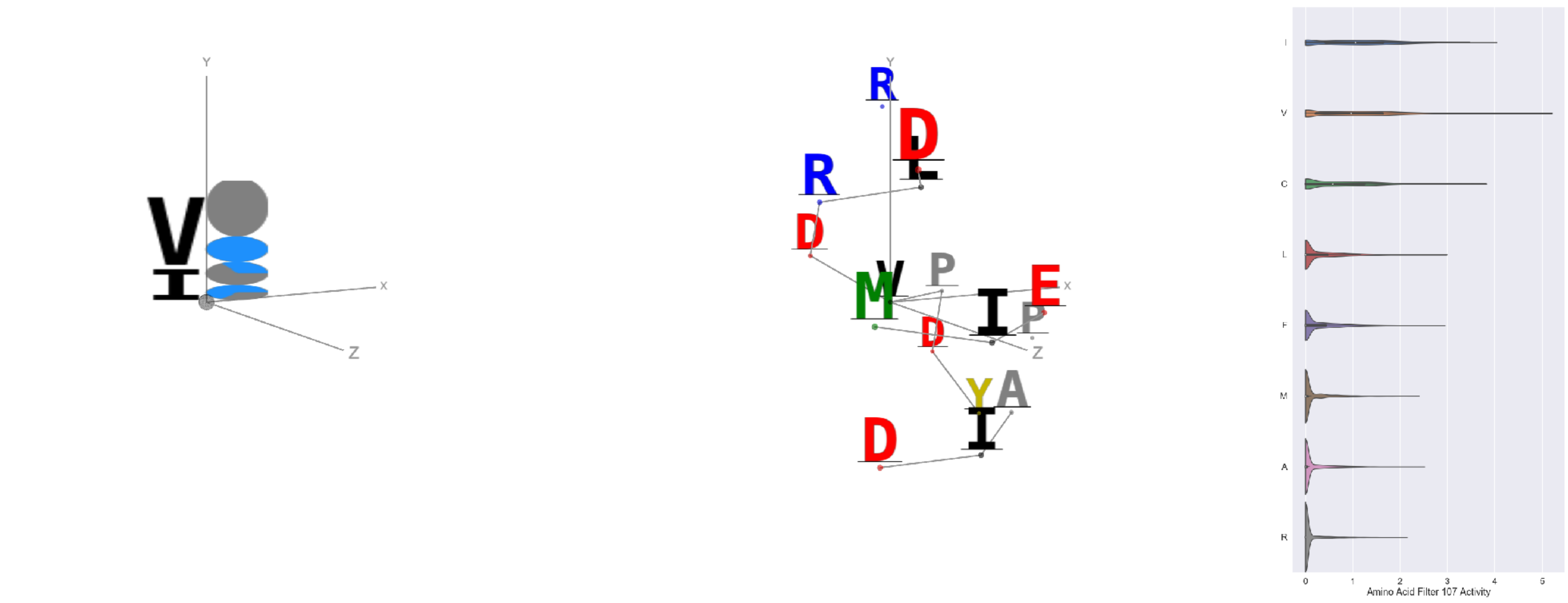


Filter 107
$R_{\text{pred}} = -0.09$
Percent Active = 25
Pooling attention 0.03
Pooling feature 0.54
Top activators
3g8q:A:117 V
3tsy:A:757 V
5ui2:A:200 V
3tsy:A:288 I
3tsy:A:110 V
2iwh:B:91 V
3tsy:A:185 I
2idb:A:381 V
3tsy:A:174 I
4u9c:B:128 V

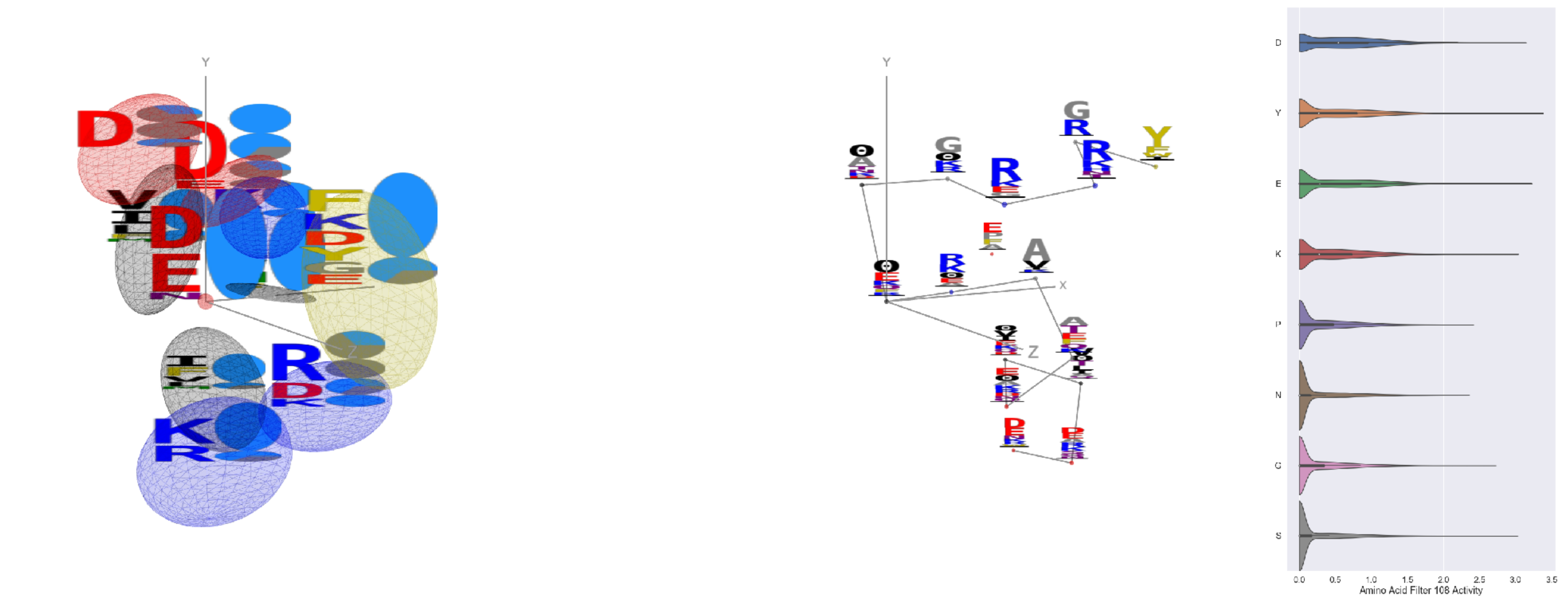


Filter 108
$R_{\text{pred}} = 0.00$
Percent Active = 39
Pooling attention 0.24
Pooling feature 1.17
Top activators
1sl0:A:575 Y
3bow:C:628 Y
4ani:A:72 E
4c0b:D:469 Y
1rq0:A:265 E
3din:D:48 D
1rq0:A:268 D
1yqg:A:263 Y
1r7r:A:735 Y
4asi:C:1783 K

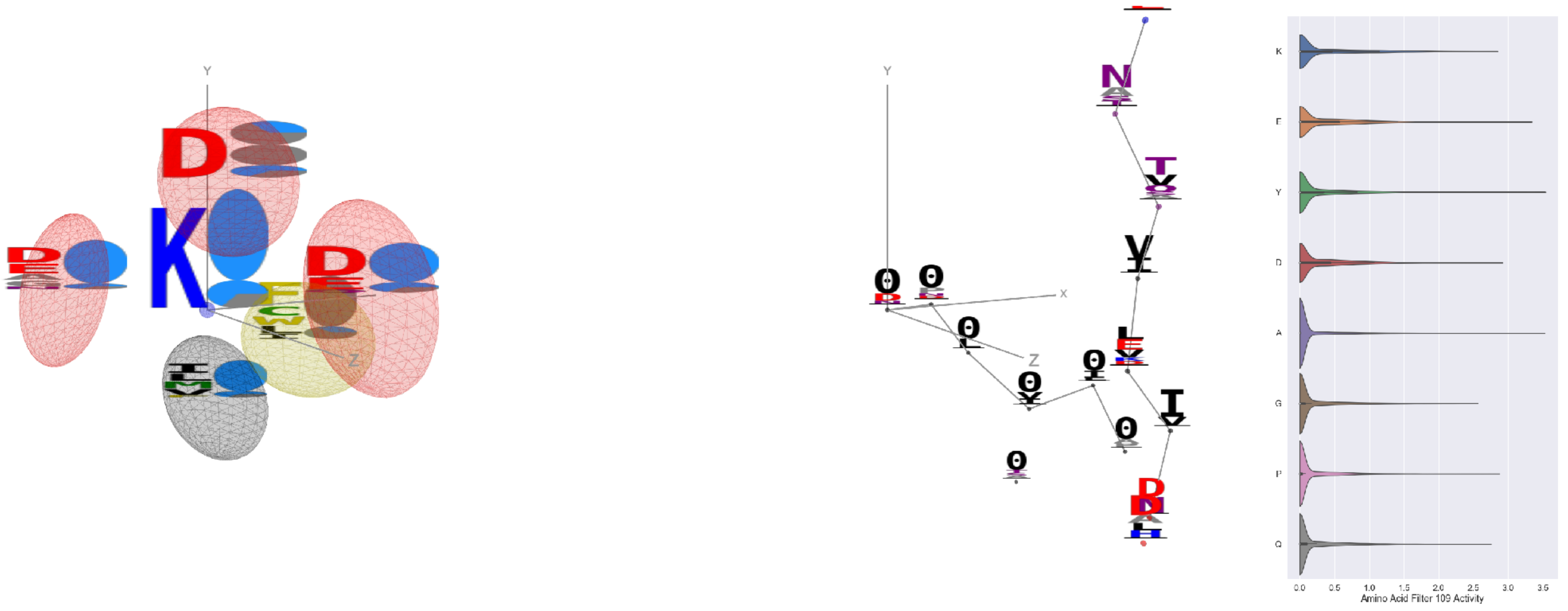


Filter 109
$R_{\text{pred}} = -0.05$
Percent Active = 29
Pooling attention 0.09
Pooling feature 0.41
Top activators
2v6e:A:535 Y
2wgq:B:669 A
4u7u:M:266 Y
2x3l:B:169 E
1o7g:A:405 Y
3ax7:A:165 Y
3b8e:A:487 E
4s20:L:668 A
3drw:A:213 Y
3hgi:A:240 A

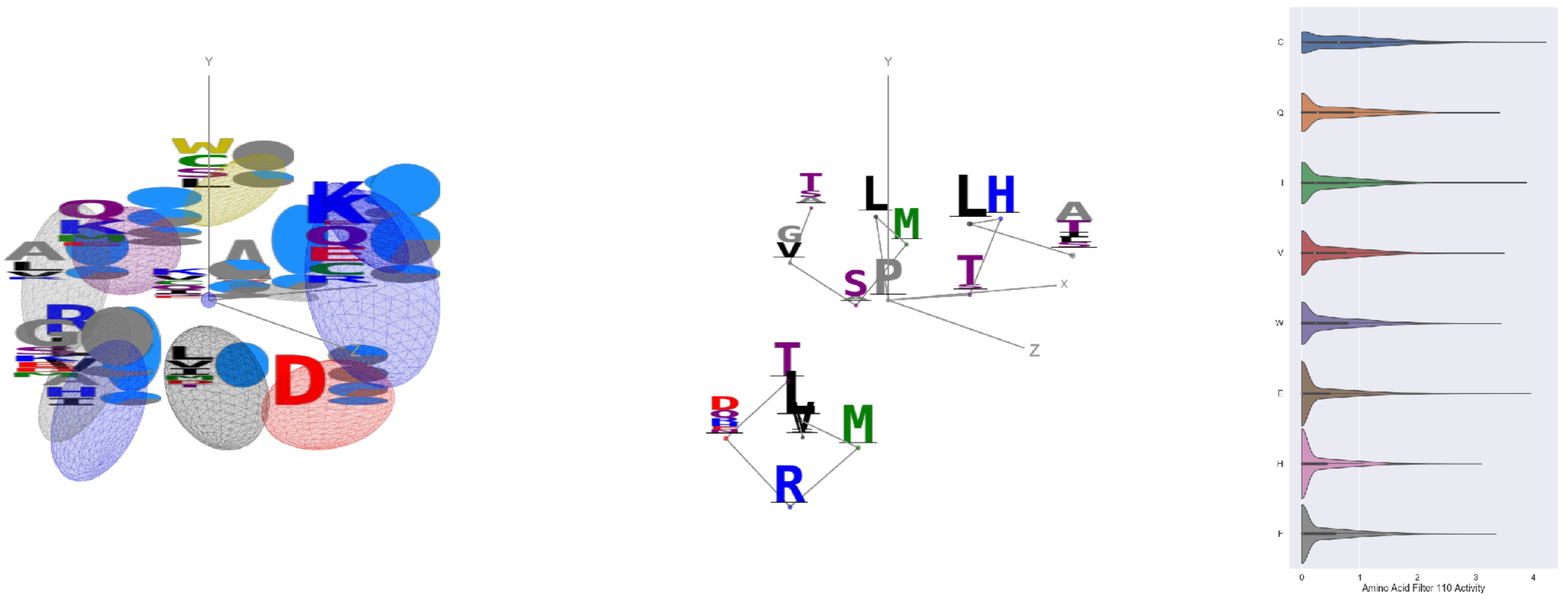


Filter 110
$R_{\text{pred}} = -0.03$
Percent Active = 36
Pooling attention 0.05
Pooling feature 0.09
Top activators
4o93:A:33 P
2gy7:B:227 C
1r7r:A:572 C
2iwh:B:859 E
1eg1:A:219 C
4v81:H:122 I
2iwh:B:690 C
1f93:E:11 E
2qn5:B:98 C
1a92:A:31 E

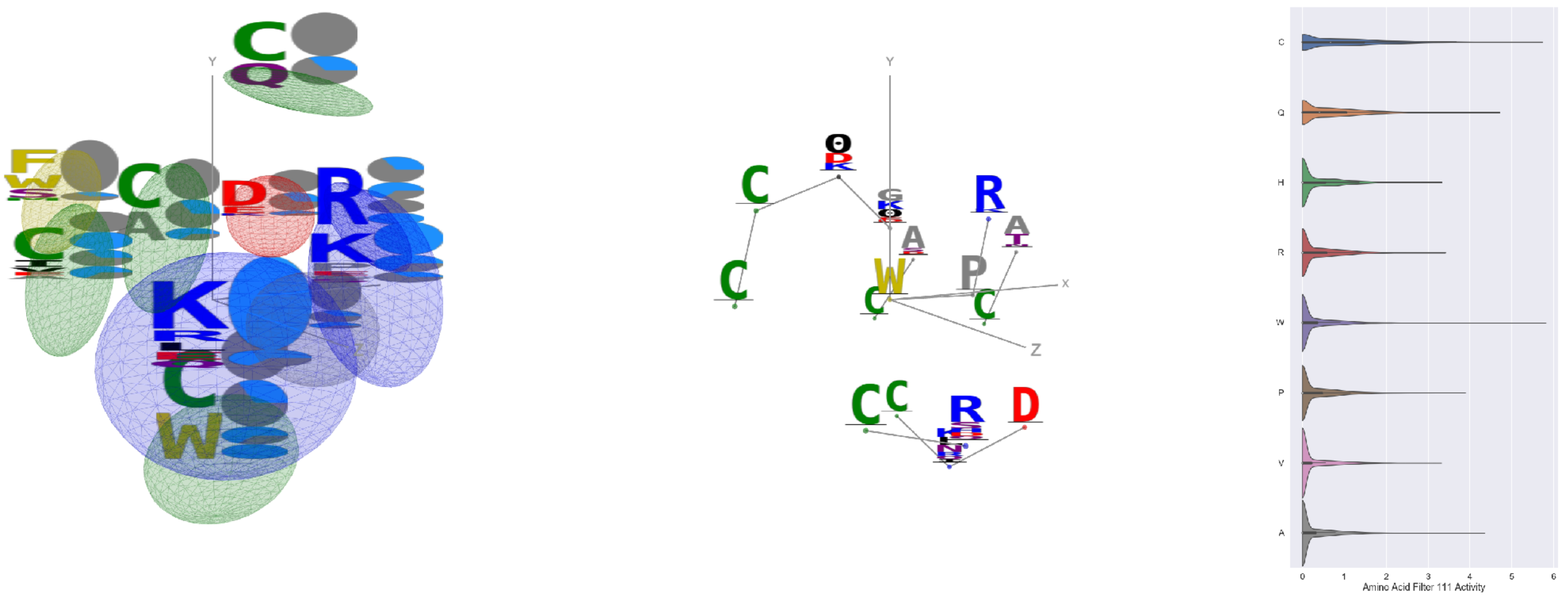


Filter 111
$R_{\text{pred}} = 0.15$
Percent Active = 32
Pooling attention 0.57
Pooling feature 0.18
Top activators
2qn5:B:72 W
2qn5:B:105 C
1izl:D:221 T
4e0s:B:75 C
2qn5:B:41 C
4c8q:H:720 Q
1eg1:A:175 C
2qn5:B:79 A
3irm:A:408 Q
4pht:A:429 C

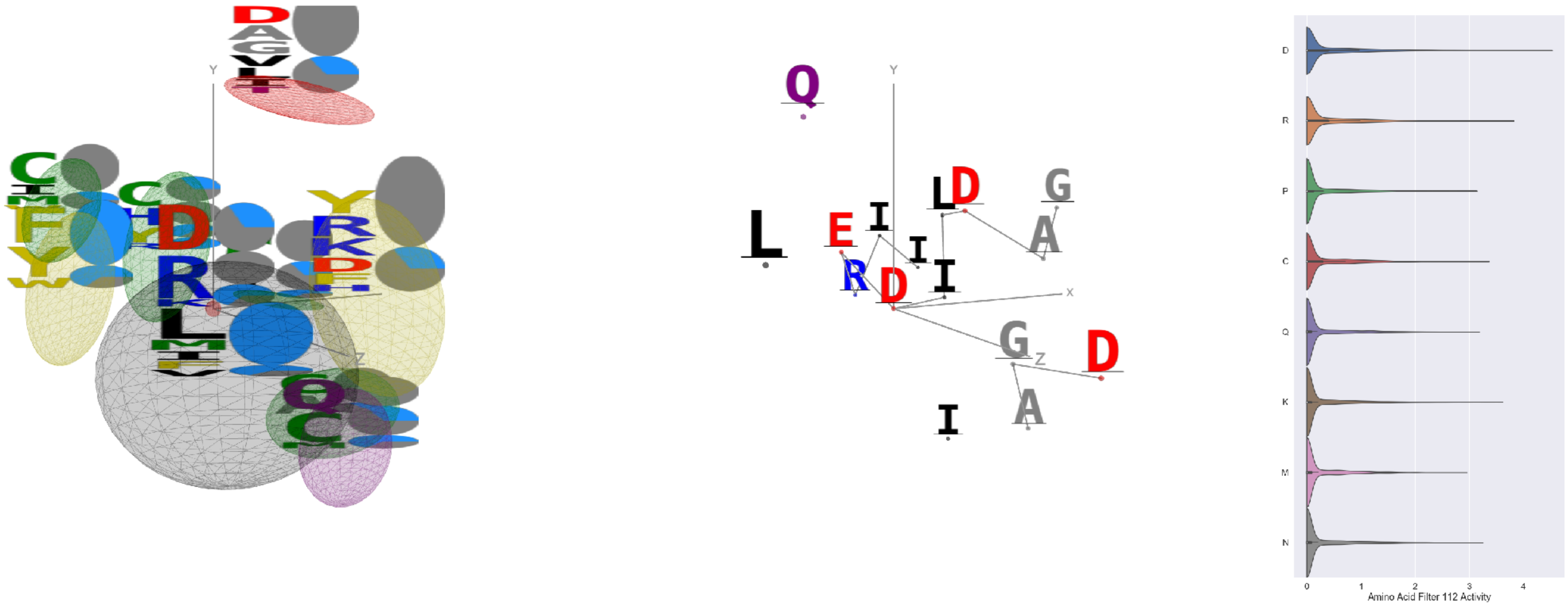


Filter 112
$R_{\text{pred}} = -0.01$
Percent Active = 23
Pooling attention 0.30
Pooling feature 2.12
Top activators
2iwh:B:373 D
4cvw:A:648 D
1bgv:A:165 D
2iwh:B:748 D
1f4a:A:204 R
1b8a:A:368 R
3do6:A:304 R
2x98:A:114 D
1mj2:A:49 R
3ax7:A:89 E

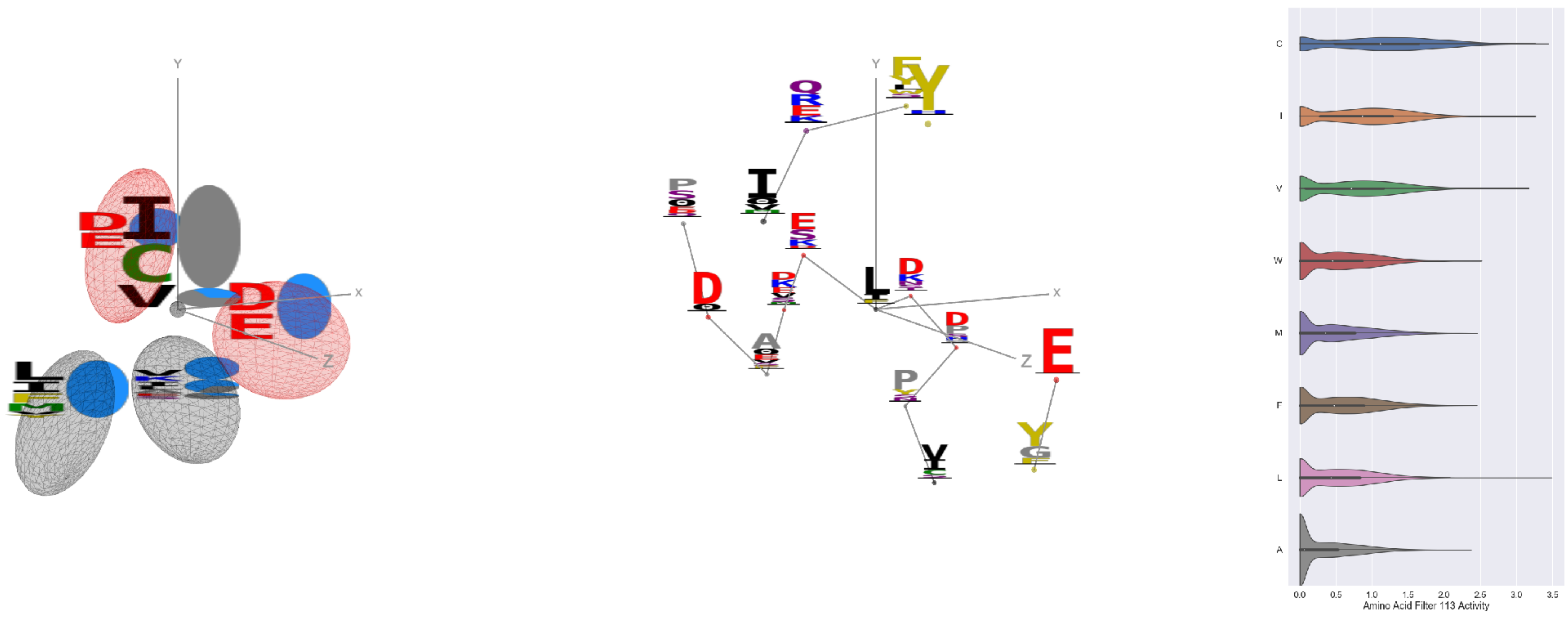


Filter 113
$R_{\text{pred}} = -0.18$
Percent Active = 42
Pooling attention 0.04
Pooling feature 0.50
Top activators
4f5w:A:212 L
1ghq:C:113 C
3l0o:A:86 I
4e0s:B:665 C
2idb:A:256 V
4g1m:B:536 C
5hu6:C:309 C
1a71:A:111 C
2qn5:B:76 C
1mu2:A:254 V

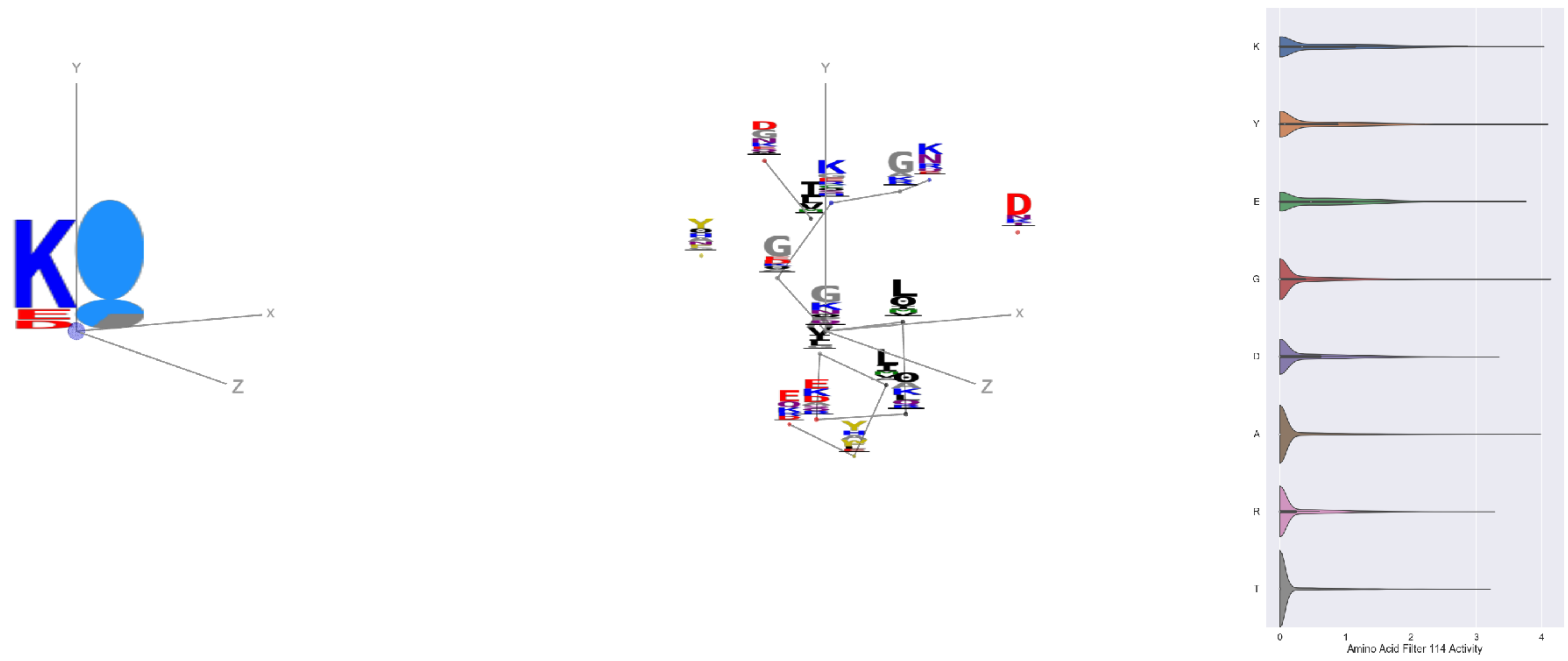


Filter 114
$R_{\text{pred}} = -0.01$
Percent Active = 35
Pooling attention 0.45
Pooling feature 1.33
Top activators
2x7x:A:152 G
1gyt:A:37 Y
1f75:A:133 L
1fpq:A:55 K
2v6e:A:535 Y
4ehi:B:156 A
4pfp:A:217 K
2i8d:A:9 A
4s20:L:449 Y
4s20:L:668 A

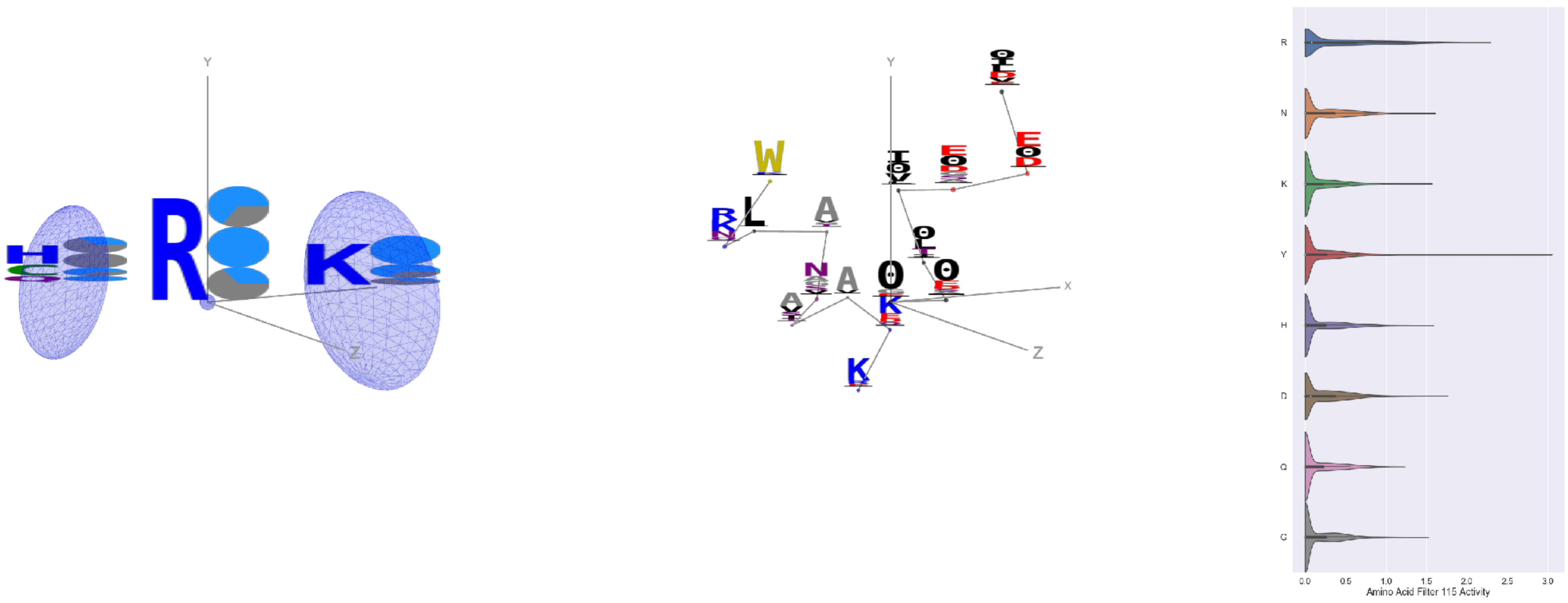


Filter 115
$R_{\text{pred}} = 0.23$
Percent Active = 31
Pooling attention 0.21
Pooling feature 0.37
Top activators
2iut:A:723 Y
2iwh:B:977 W
1hfe:L:398 Y
2v6e:A:535 Y
1o7g:A:448 Y
1qhd:A:145 R
2iwh:B:456 R
4mex:C:394 R
3g8q:A:52 R
1gvn:A:90 Y

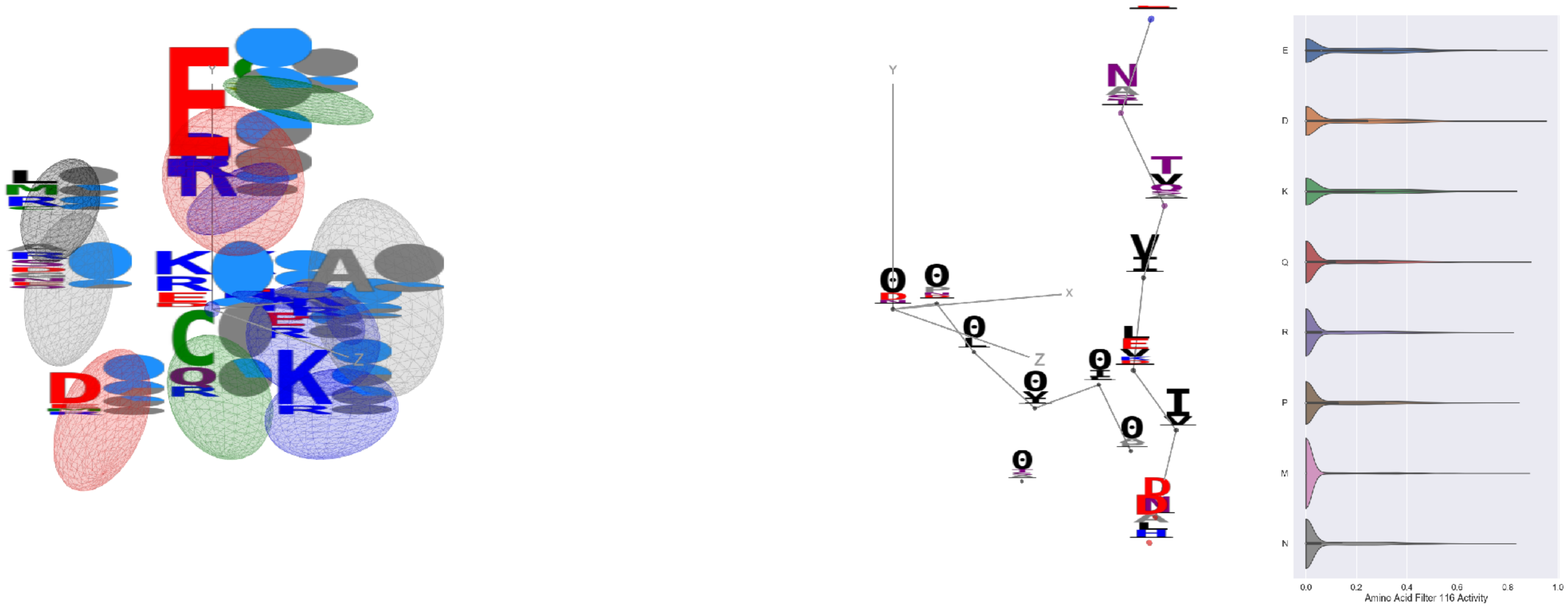


Filter 116
$R_{\mathrm{pred}} = 0.13$
Percent Active = 26
Pooling attention 0.72
Pooling feature 3.63
Top activators
2v6e:A:535 Y
3o47:A:235 E
3g8q:A:41 D
2iwh:B:977 W
4buj:B:13 E
1r7r:A:697 L
3tu3:B:652 D
2f8x:C:278 D
3tsy:A:763 D
1r7r:A:693 E

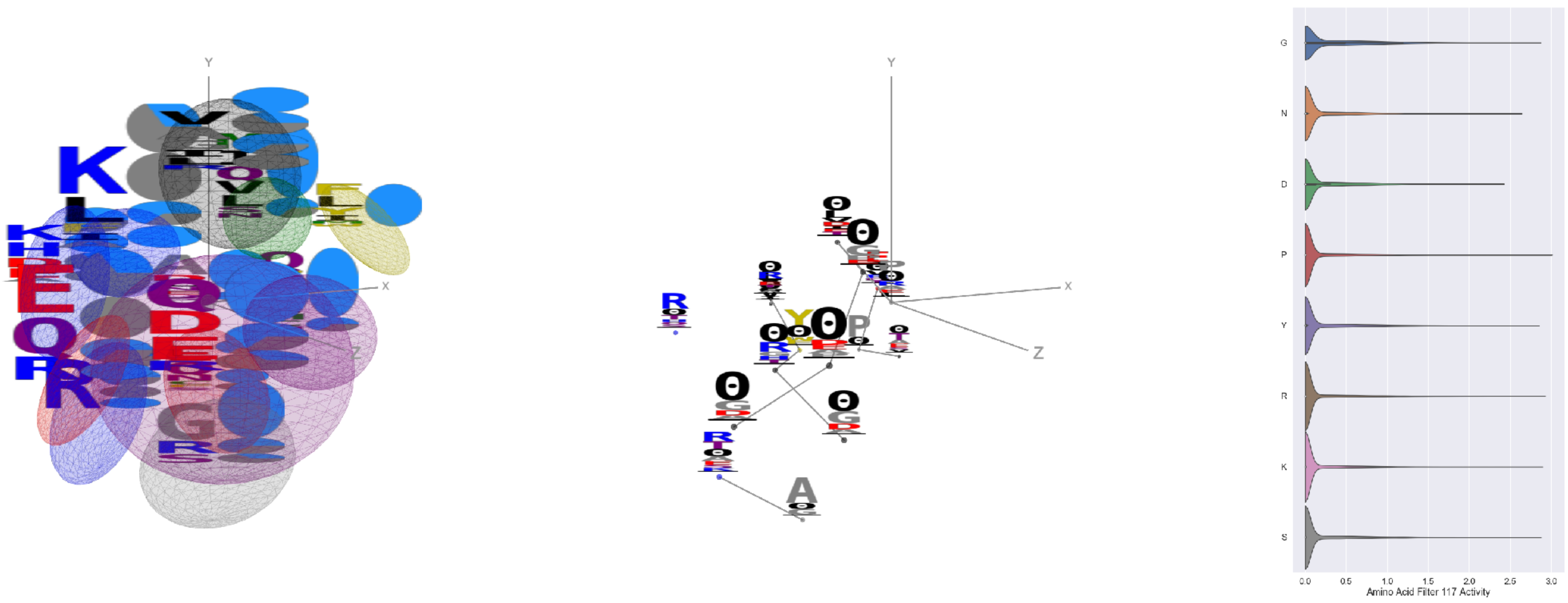


Filter 117
$R_{\mathrm{pred}} = 0.10$
Percent Active = 21
Pooling attention 0.70
Pooling feature 4.04
Top activators
2x0q:A:46 P
1srq:A:118 R
3pio:O:65 K
4wj3:B:263 S
1bh3:A:161 G
3igh:X:263 Y
4pht:A:485 G
1ivu:A:472 T
4r3z:C:353 G
4fzq:A:447 V

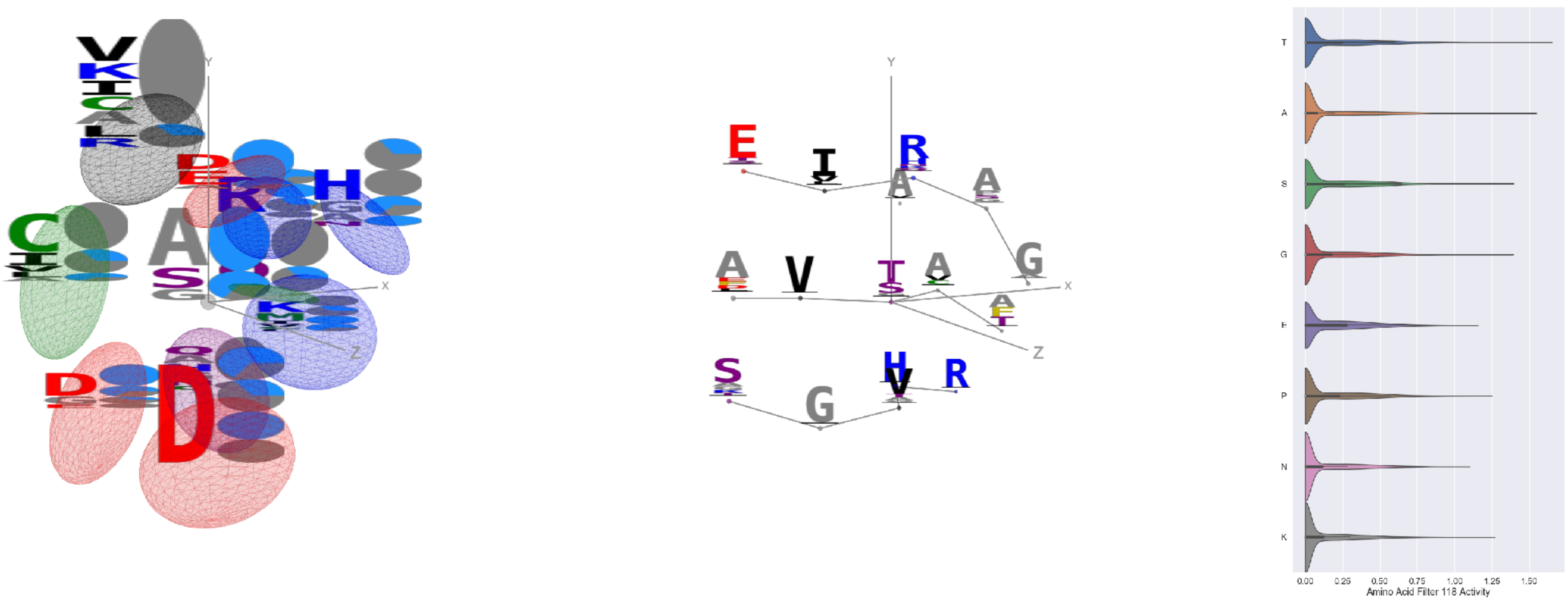


Filter 118
$R_{\mathrm{pred}} = 0.08$
Percent Active = 26
Pooling attention 0.72
Pooling feature 2.63
Top activators
1rq0:A:179 T
4nm9:B:417 A
3tsy:A:451 A
1izl:D:143 A
1izl:D:202 A
1vqv:B:129 S
4d0l:A:551 G
1f75:A:25 A
4bxf:A:351 S
4c0b:B:96 Y

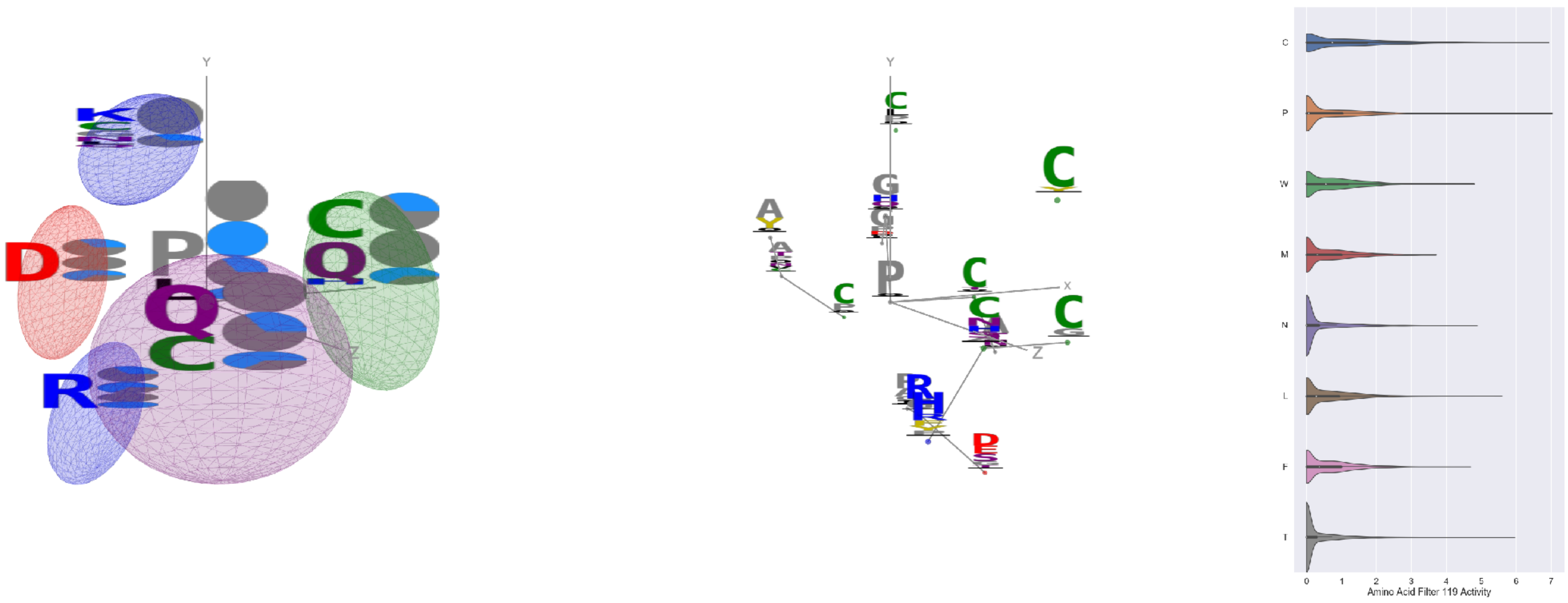


Filter 119
$R_{\mathrm{pred}} = -0.09$
Percent Active = 33
Pooling attention 0.16
Pooling feature 0.48
Top activators
3di4:A:238 P
1eg1:A:219 C
2gy7:B:243 P
3oru:A:208 P
2gy7:B:227 C
4e0s:B:531 C
1cm5:A:335 T
1h8p:A:109 C
2qn5:B:91 C
2qn5:B:27 D

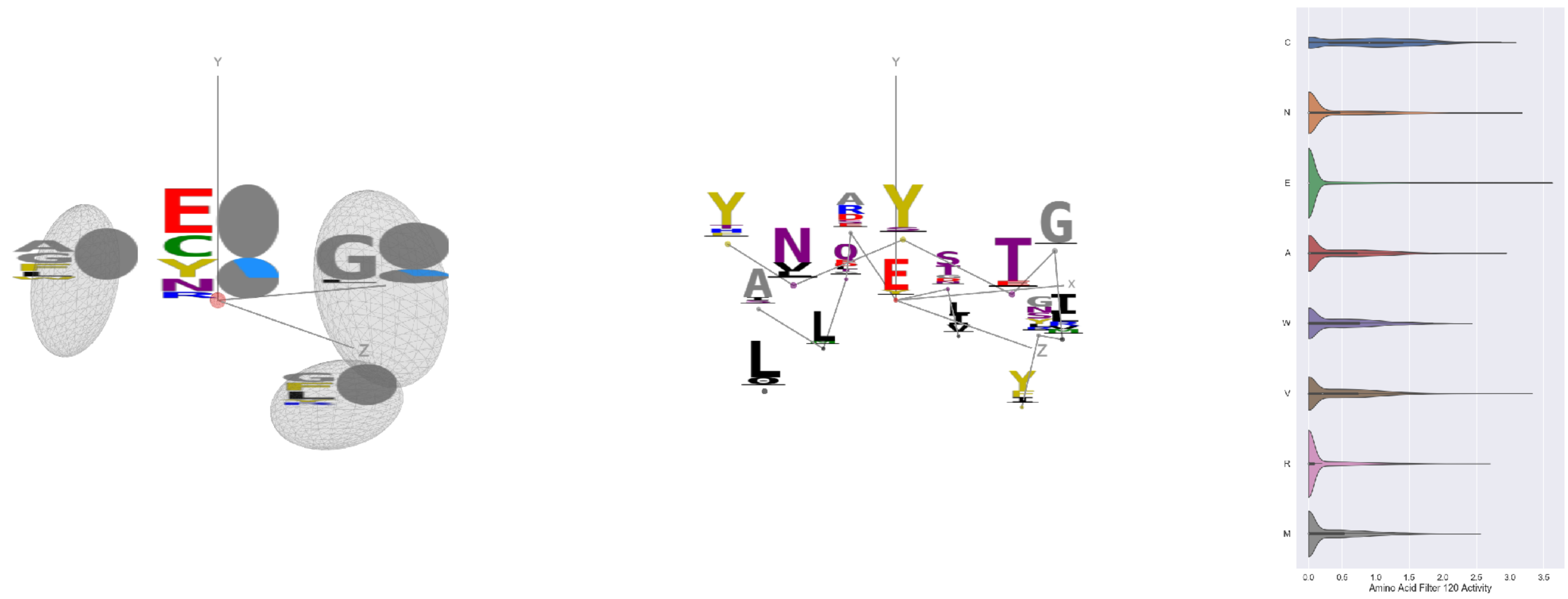

Filter 120
$R_{\text{pred}} = -0.27$
Percent Active = 33
Pooling attention 0.18
Pooling feature 0.82
Top activators
4l8n:A:126 E
1iwa:A:313 V
1lk5:A:107 E
2iwh:B:84 E
3n99:G:86 N
4u4c:A:277 E
4nm9:B:940 N
3do6:A:471 C
3tsy:A:286 N
1qha:A:294 E

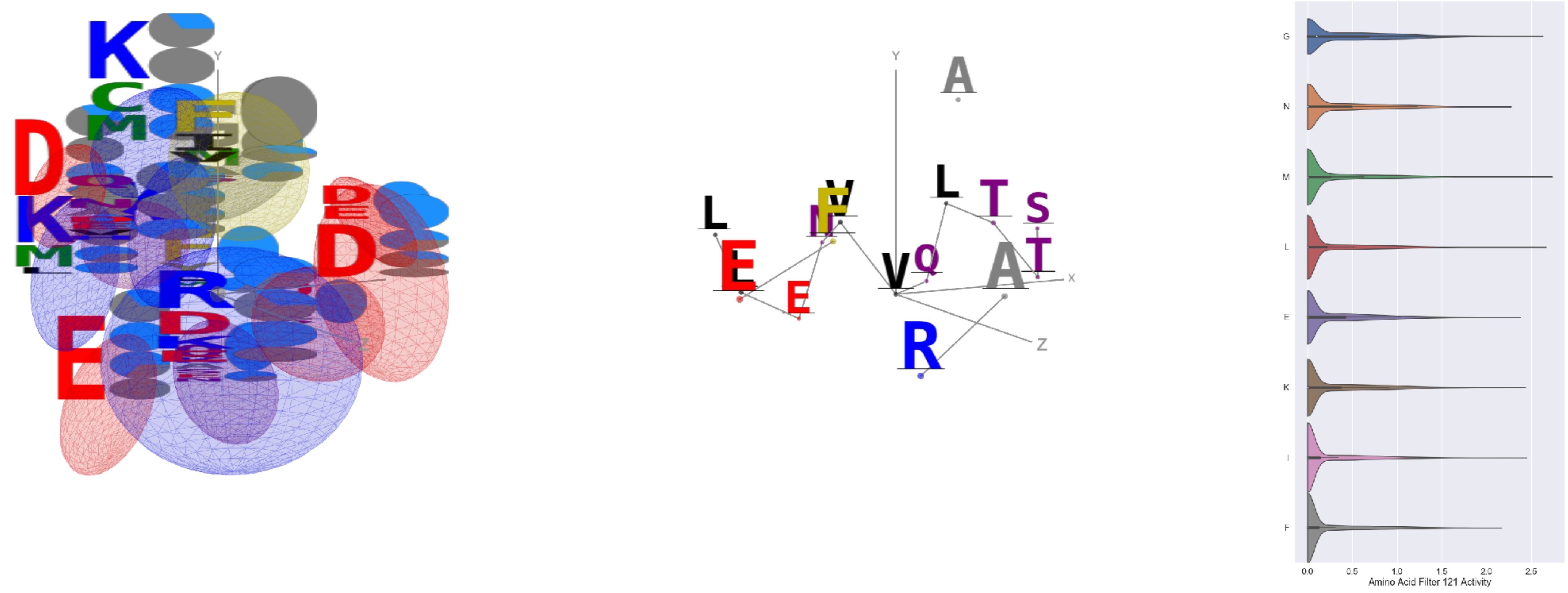

Filter 121
$R_{\text{pred}} = 0.06$
Percent Active = 35
Pooling attention 0.37
Pooling feature 3.07
Top activators
2iwh:B:343 V
1efu:B:7 M
3tsy:A:424 L
5kqv:E:299 R
4itr:A:3662 L
1sl0:A:414 G
1zt2:A:238 V
3ax7:A:758 G
1hfe:L:44 V
3tu3:B:221 Q

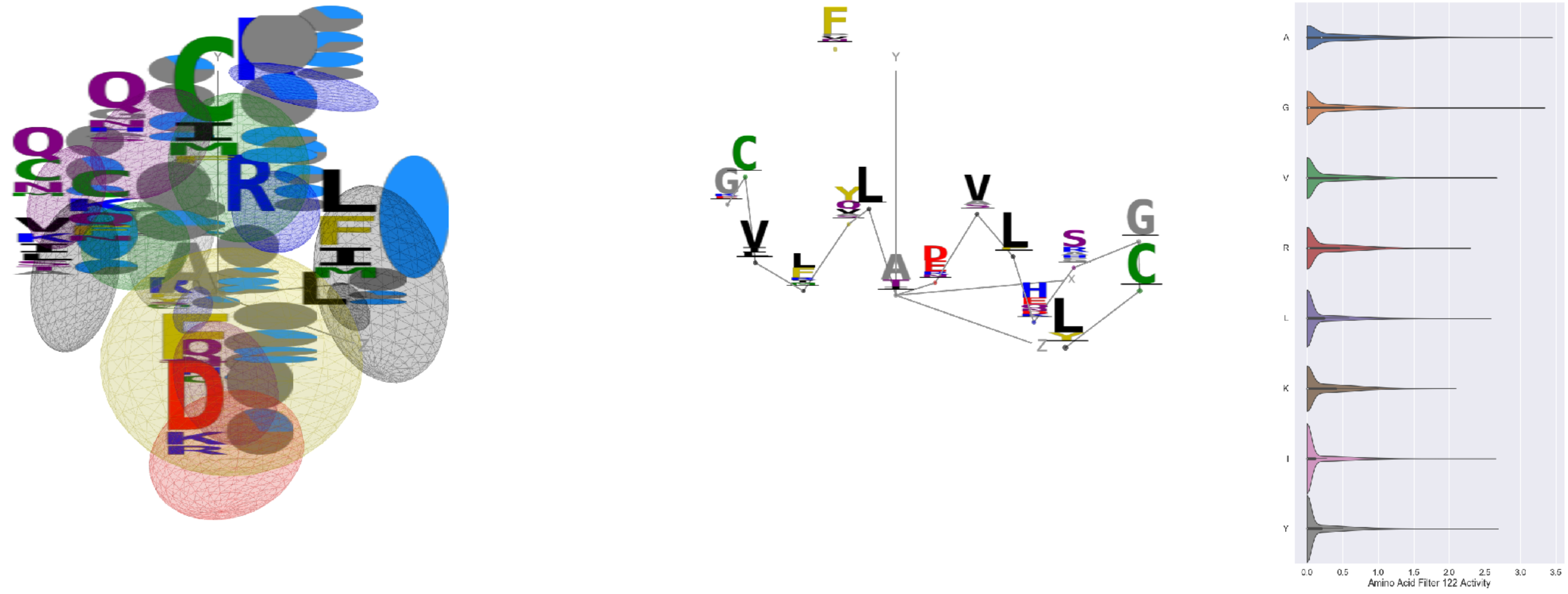

Filter 122
$R_{\text{pred}} = -0.04$
Percent Active = 35
Pooling attention 0.20
Pooling feature 0.32
Top activators
1b9e:B:14 A
1bxr:A:314 A
2wsc:B:184 G
1w36:B:602 A
3kov:A:41 A
2qn5:B:64 G
3tsy:A:800 A
3tsy:A:213 A
1mj2:A:64 A
4e0s:B:509 G

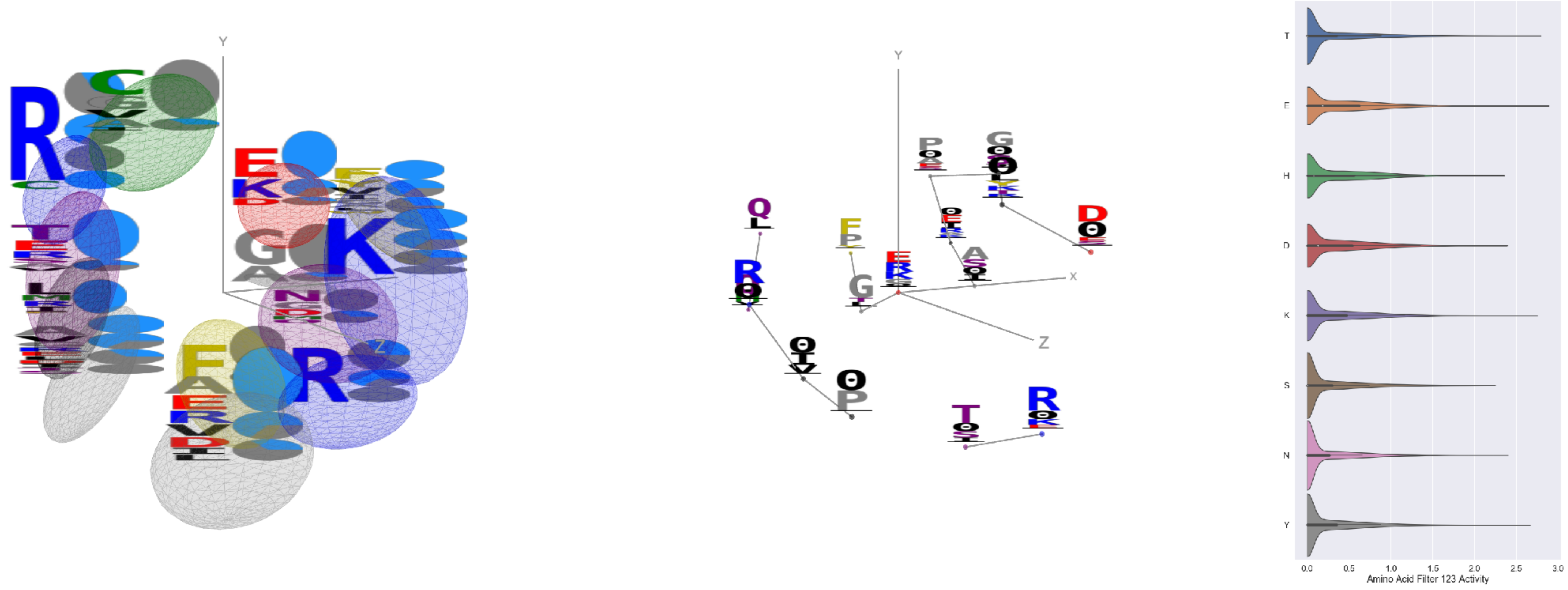

Filter 123
$R_{\text{pred}} = -0.13$
Percent Active = 35
Pooling attention 0.34
Pooling feature 1.21
Top activators
4qiw:A:801 E
2hye:B:48 T
1d0n:A:653 K
2cge:A:446 A
3zxu:A:263 E
2hye:B:147 K
3dlb:A:10 Y
1u6z:B:155 E
3oru:A:109 T
1eg1:A:184 E

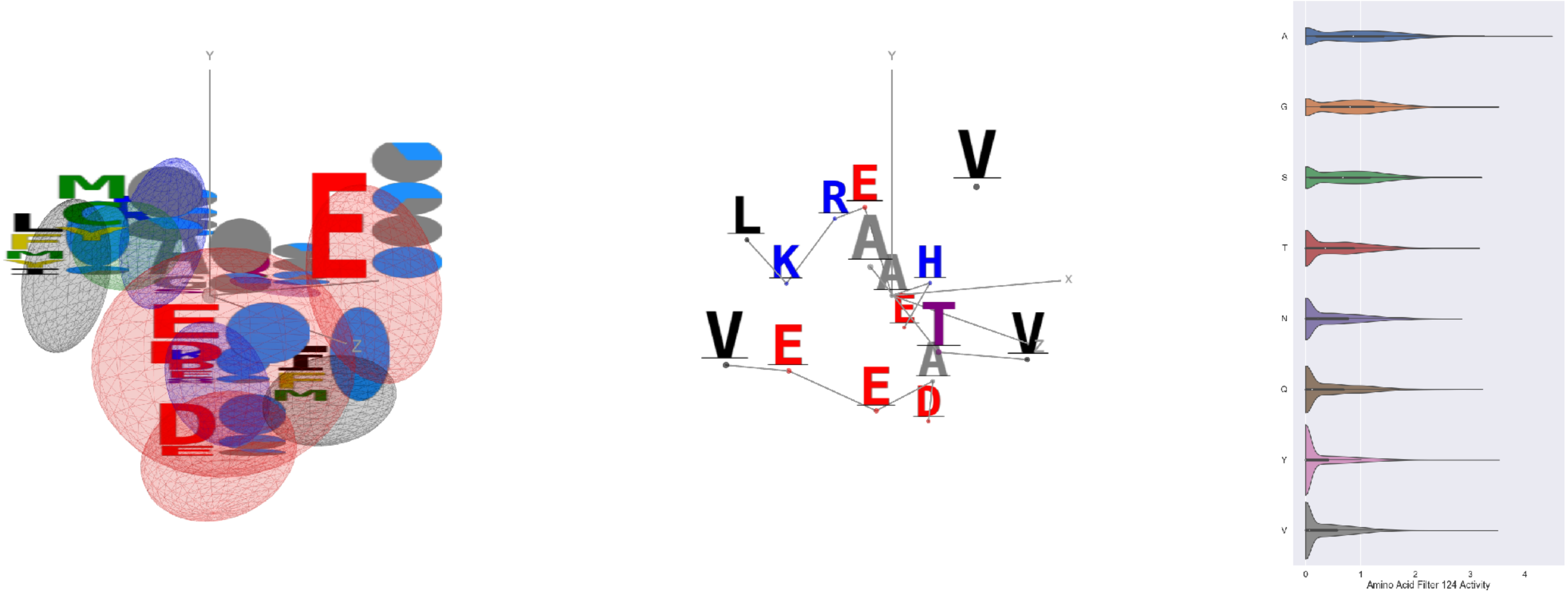


Filter 124
$R_{\text{pred}} = -0.14$
Percent Active = 44
Pooling attention 0.15
Pooling feature 0.20
Top activators
3g8q:A:45 A
3g8q:A:138 A
3g8q:A:224 A
2np9:A:417 A
4v8p:BC:96 A
3o65:A:175 Y
4r3z:C:521 G
3n6r:G:360 V
2iwh:B:697 A
3do6:A:321 A

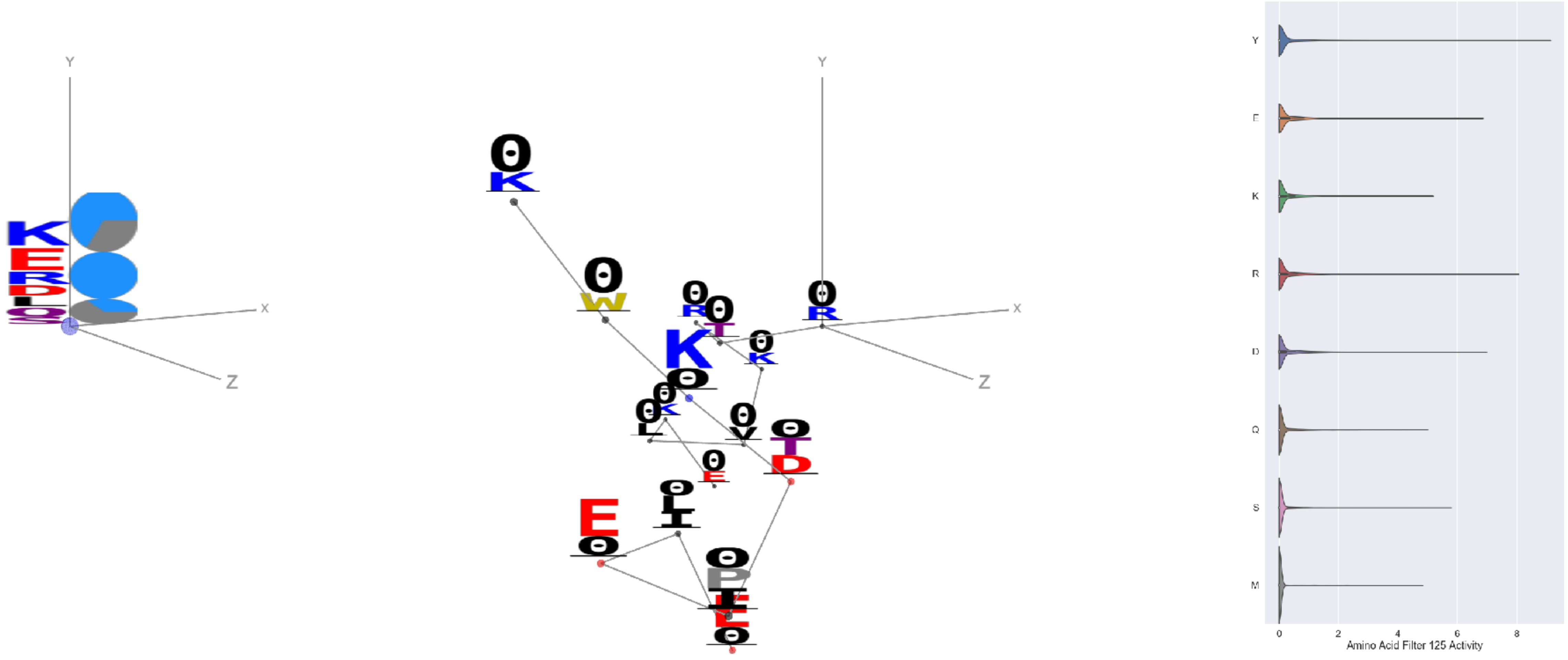


Filter 125
$R_{\text{pred}} = -0.01$
Percent Active = 16
Pooling attention 0.04
Pooling feature 0.18
Top activators
3o47:A:5 Y
1jb0:K:20 I
4c8q:H:656 A
2v6e:A:535 Y
3o47:A:205 R
2yma:A:265 Y
1e7n:A:-2 Y
1r7r:A:669 D
1n1c:A:105 Y
2b5d:X:475 E

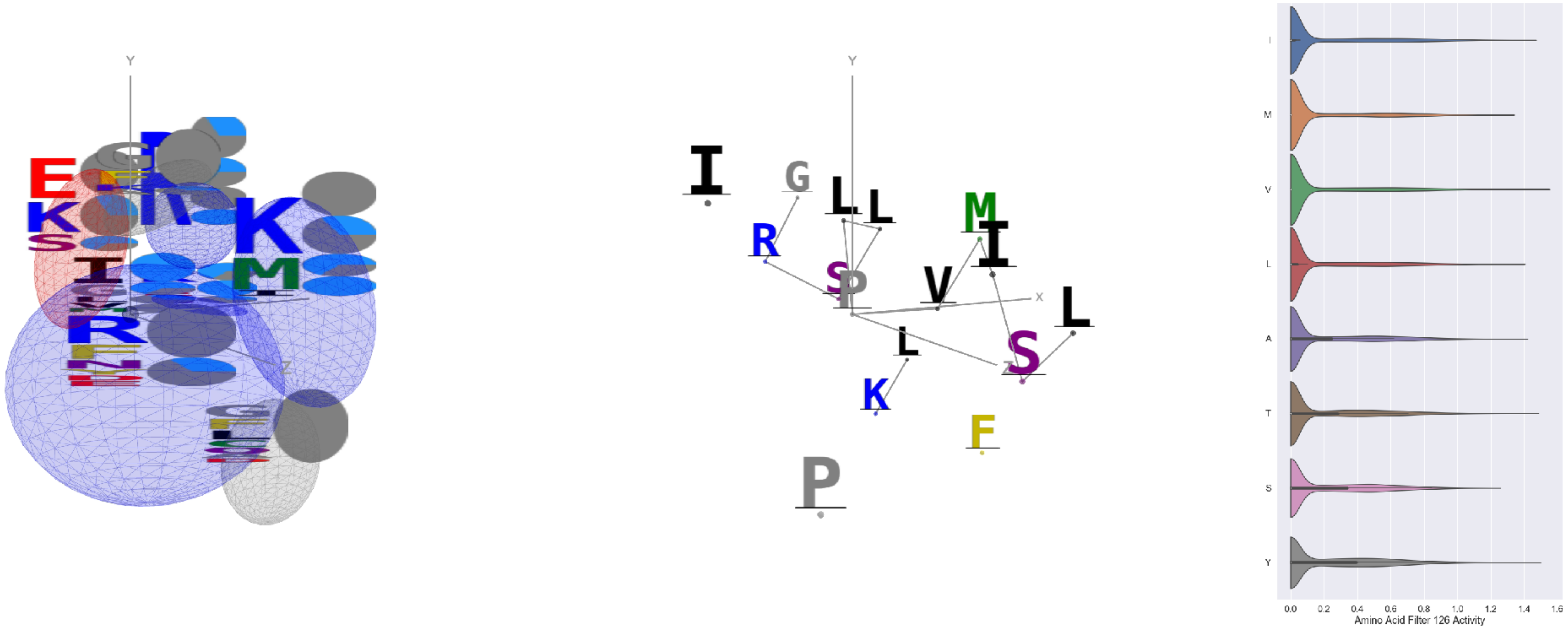


Filter 126
$R_{\text{pred}} = 0.19$
Percent Active = 34
Pooling attention 0.18
Pooling feature 1.73
Top activators
2iwh:B:248 P
2iwh:B:291 P
1mj2:A:45 V
1fx0:B:263 E
4nm9:B:27 Y
1iwa:A:63 T
3g8q:A:35 V
4uzi:A:305 I
1e1c:A:555 V
4oph:A:197 Y

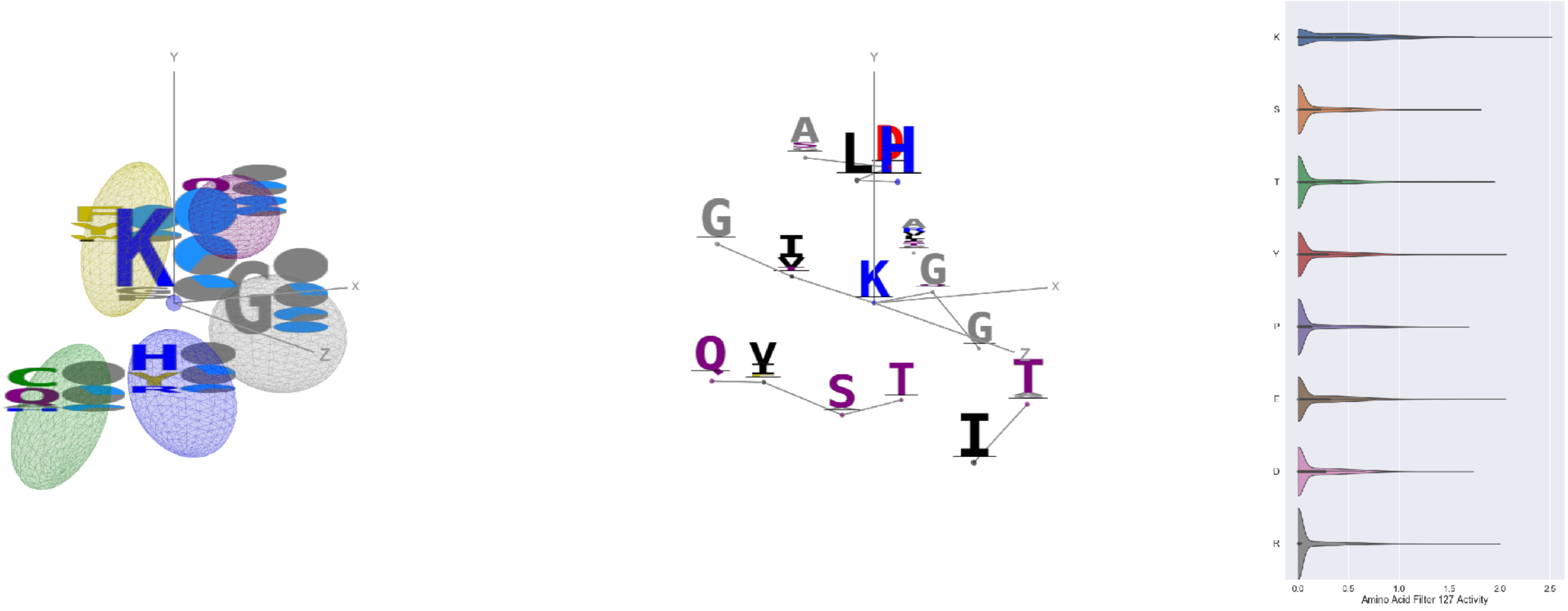


Filter 127
$R_{\text{pred}} = -0.01$
Percent Active = 25
Pooling attention 0.37
Pooling feature 1.68
Top activators
1fx0:B:167 K
1umg:A:86 K
4rcn:B:404 K
2d32:A:386 K
4oph:A:62 K
1cm5:A:103 A
4wj3:B:79 K
4gt2:A:433 Y
4r3z:B:655 E
1ass:A:116 R
