## Supplementary material for "ScanNet: An interpretable geometric deep learning model for structure-based protein binding site prediction": 3D visualisation and activation statistics of all the amino acid filters of the B-cell epitope network

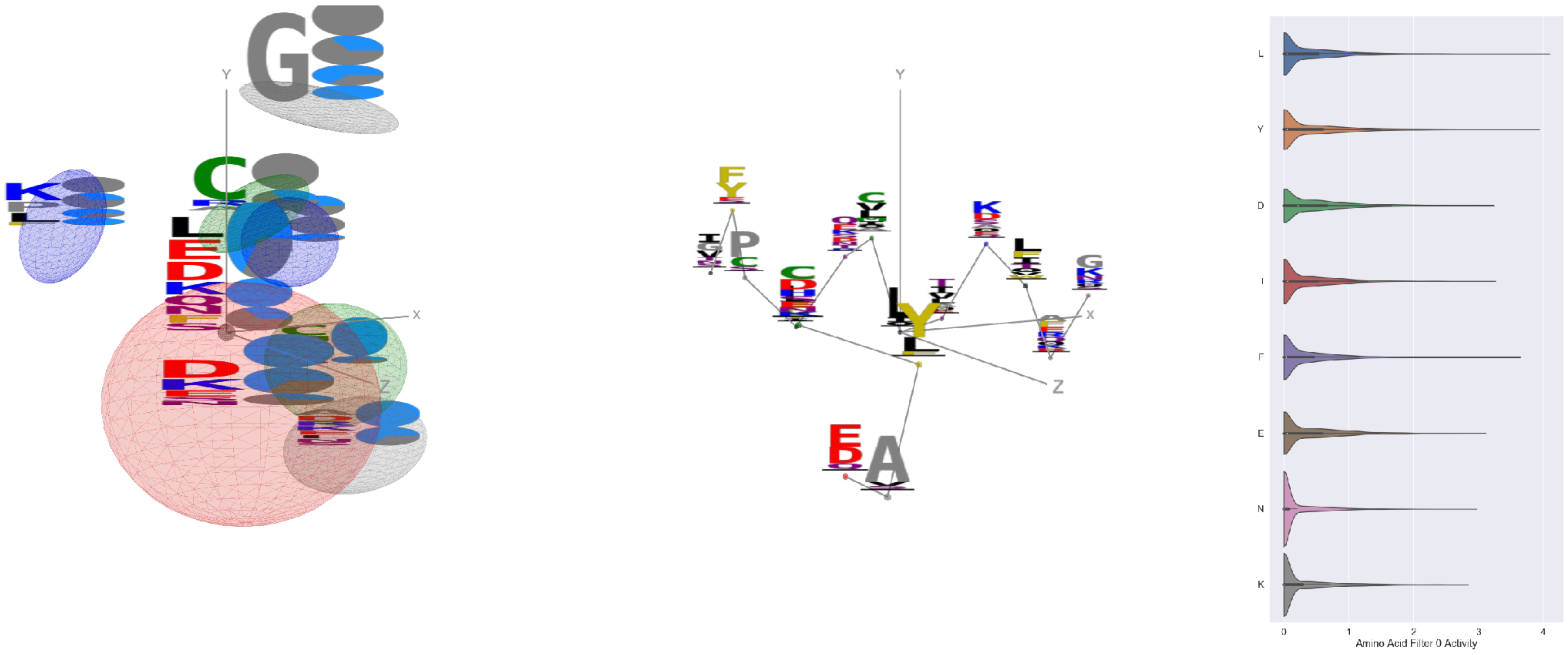


Filter 0
$R_{\text{pred}} = -0.09$
Percent Active = 42
Pooling attention 0.46
Pooling feature 1.18
Top activators
4xmn:E:922 L
6ln2:A:40 Y
4xmn:E:889 L
4xmn:E:981 M
6ln2:A:257 V
4xmn:E:832 F
4xmn:E:908 L
4xmn:E:1030 Y
4xmn:E:1007 L
4xmn:E:876 Y

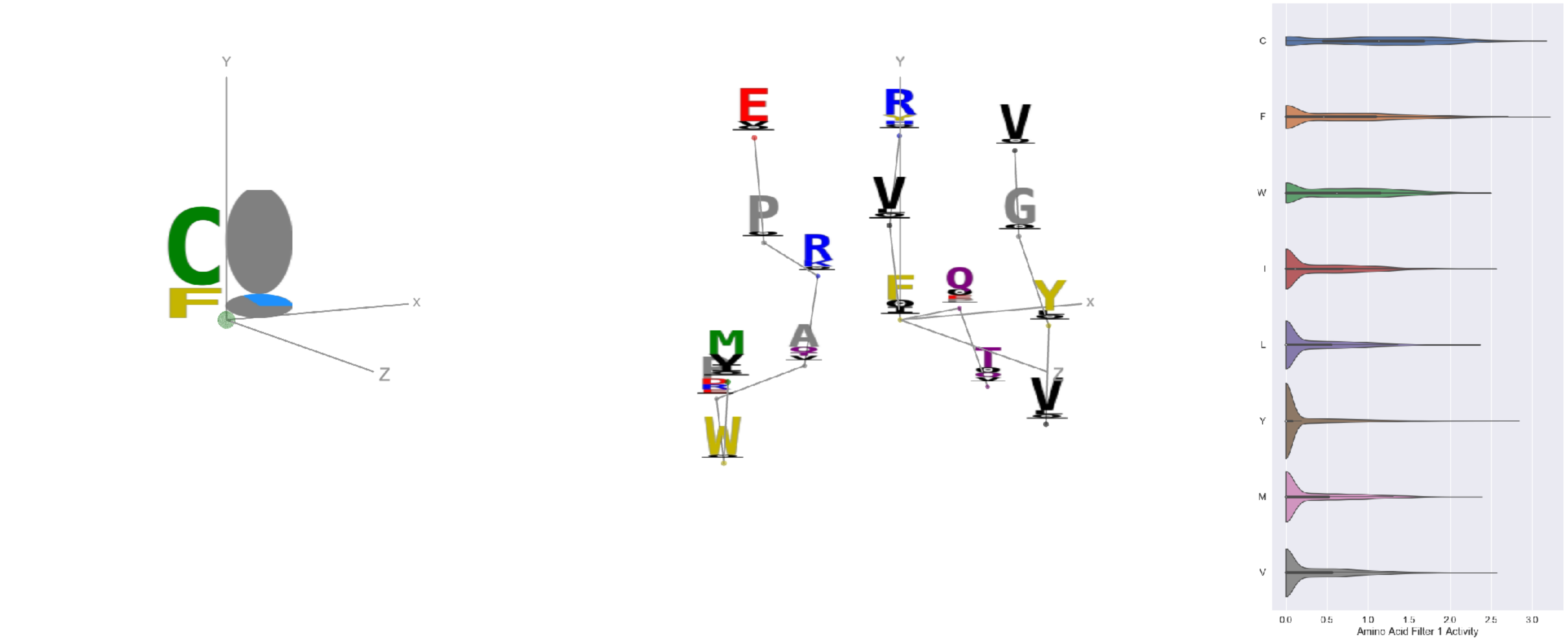


Filter 1
$R_{\text{pred}} = -0.23$
Percent Active = 32
Pooling attention 0.36
Pooling feature 2.55
Top activators
3cvh:A:33 F
4zxb:E:225 C
3iyw:A:190 C
5x0t:E:41 C
7l2e:A:391 C
5kqv:E:237 C
4zxb:E:237 C
7nd9:B:133 Y
6p95:C:86 C
7lcn:A:525 C

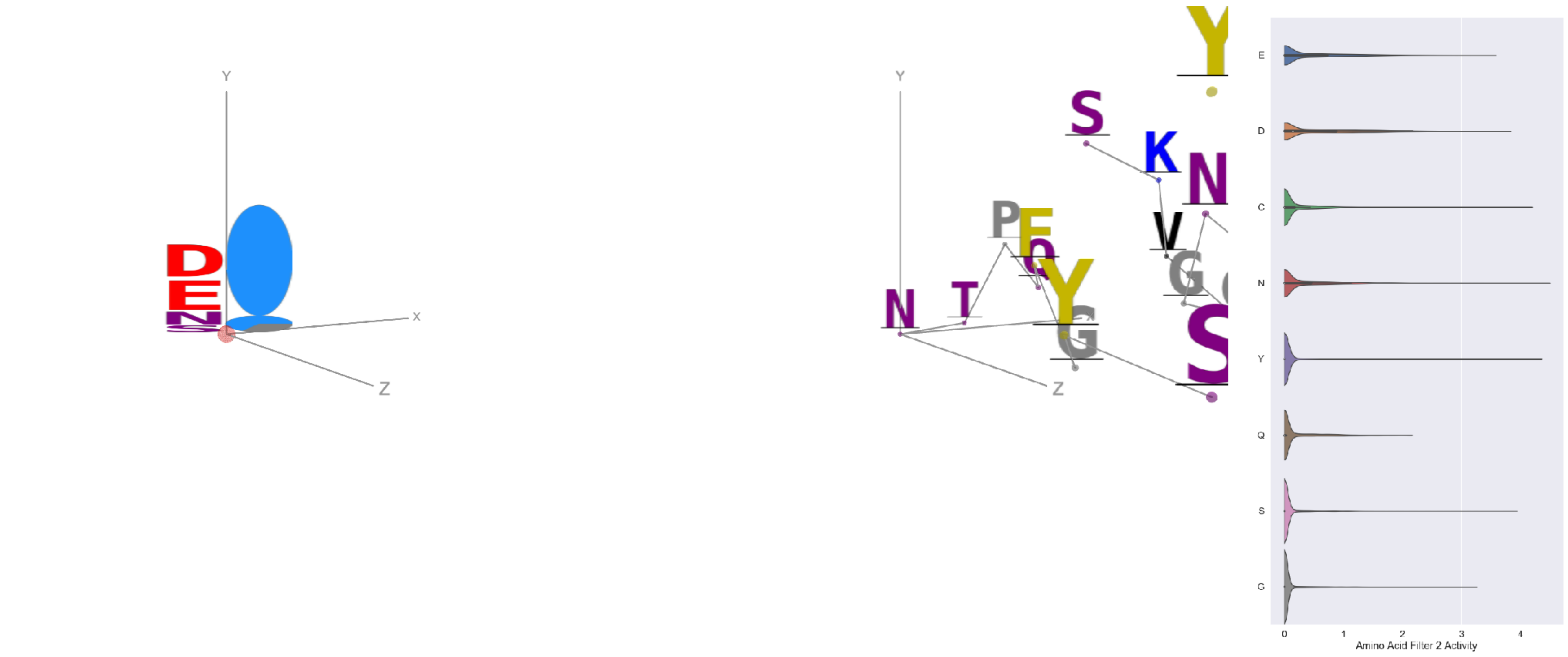


Filter 2
$R_{\text{pred}} = 0.14$
Percent Active = 15
Pooling attention 0.43
Pooling feature 1.53
Top activators
7jv2:A:501 N
7jw0:E:826 V
4cmh:A:287 Y
4g80:I:244 Y
5tlj:X:84 C
3tt3:A:511 Y
7nda:C:1147 Y
7l2d:A:1147 Y
6i07:D:260 S
6nb6:C:808 V

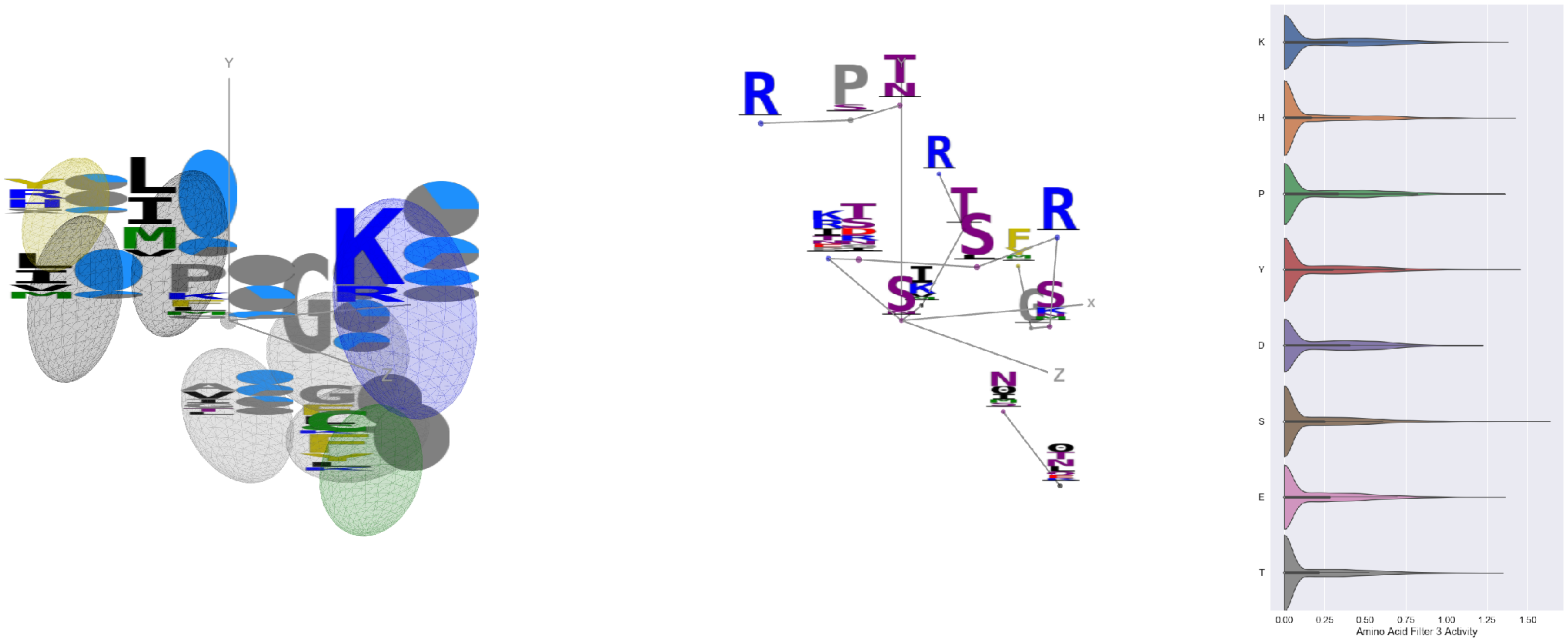


Filter 3
$R_{\text{pred}} = 0.22$
Percent Active = 32
Pooling attention 0.01
Pooling feature 2.35
Top activators
1nca:N:367 S
6q23:A:367 S
6pzw:C:367 S
5i5k:B:1588 Y
6q23:A:263 I
7ce2:A:1271 H
4uao:A:340 V
4f37:A:101 K
4fqv:D:54 S
6c9u:A:577 E

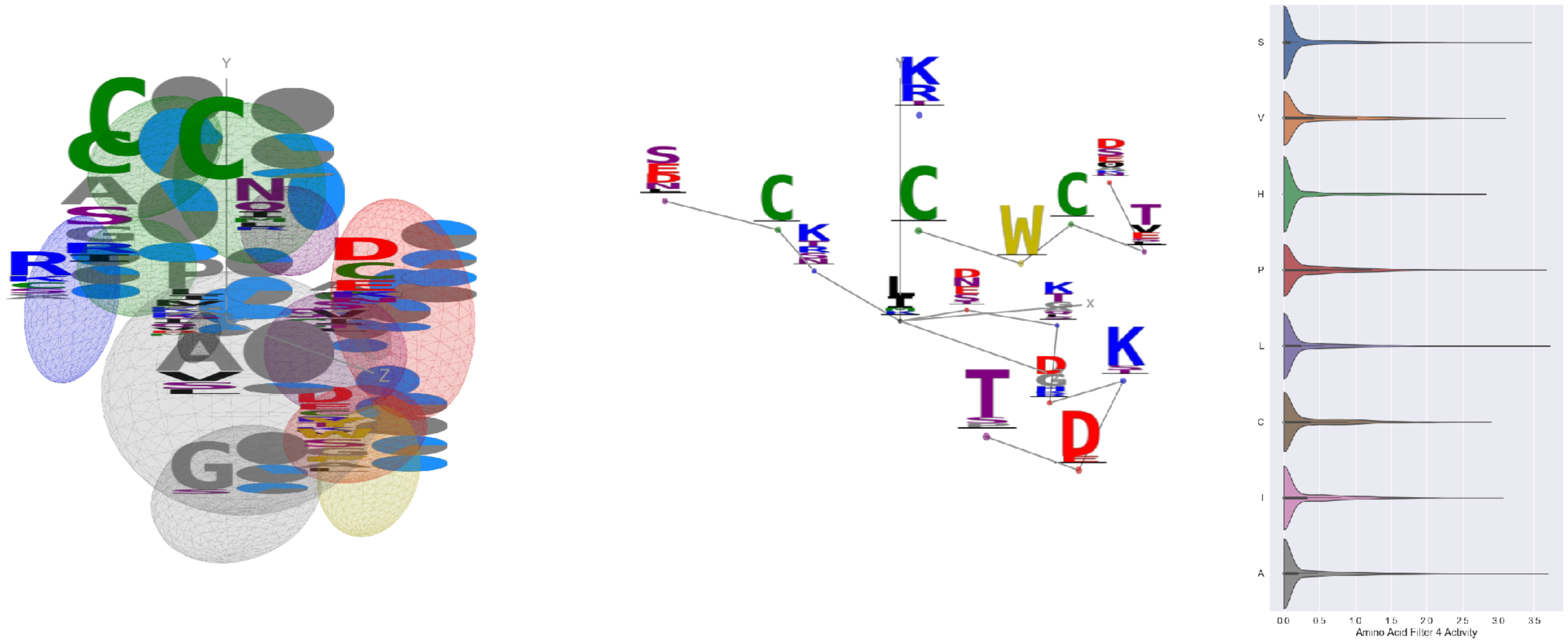


Filter 4
$R_{\mathrm{pred}} = -0.05$
Percent Active = 28
Pooling attention 0.35
Pooling feature 3.12
Top activators
6i07:D:133 L
6c9u:A:891 A
5vod:C:40 P
7l57:B:1051 S
7nda:A:785 L
6ii9:C:255 S
7jw0:E:461 Y
6pt0:A:317 K
6mej:C:419 S
3cvh:A:228 M

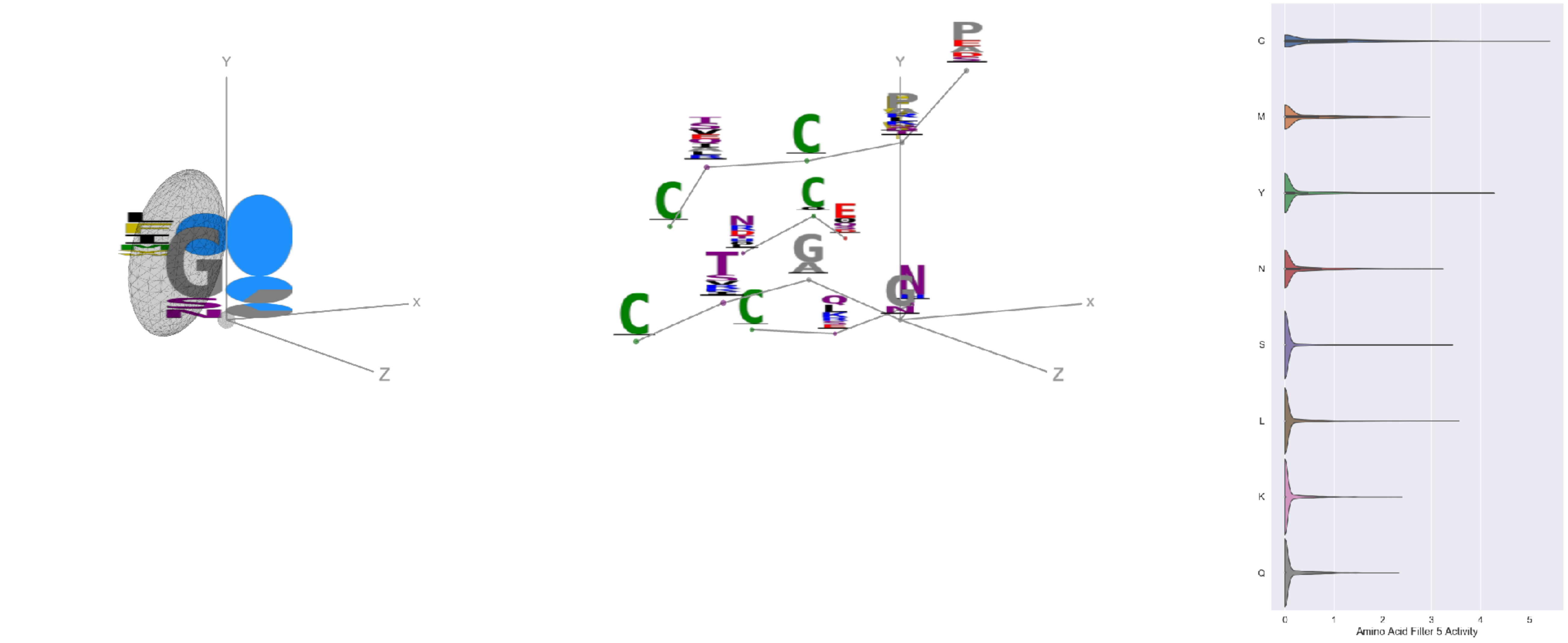


Filter 5
$R_{\mathrm{pred}} = 0.03$
Percent Active = 22
Pooling attention 0.08
Pooling feature 0.63
Top activators
5bo1:A:309 G
7c2l:C:971 G
7l2d:A:971 G
7l57:B:283 G
7jv2:A:446 G
7l2e:A:283 G
6mei:C:523 G
5vkd:A:683 G
7l57:B:971 G
6ulf:K:66 G

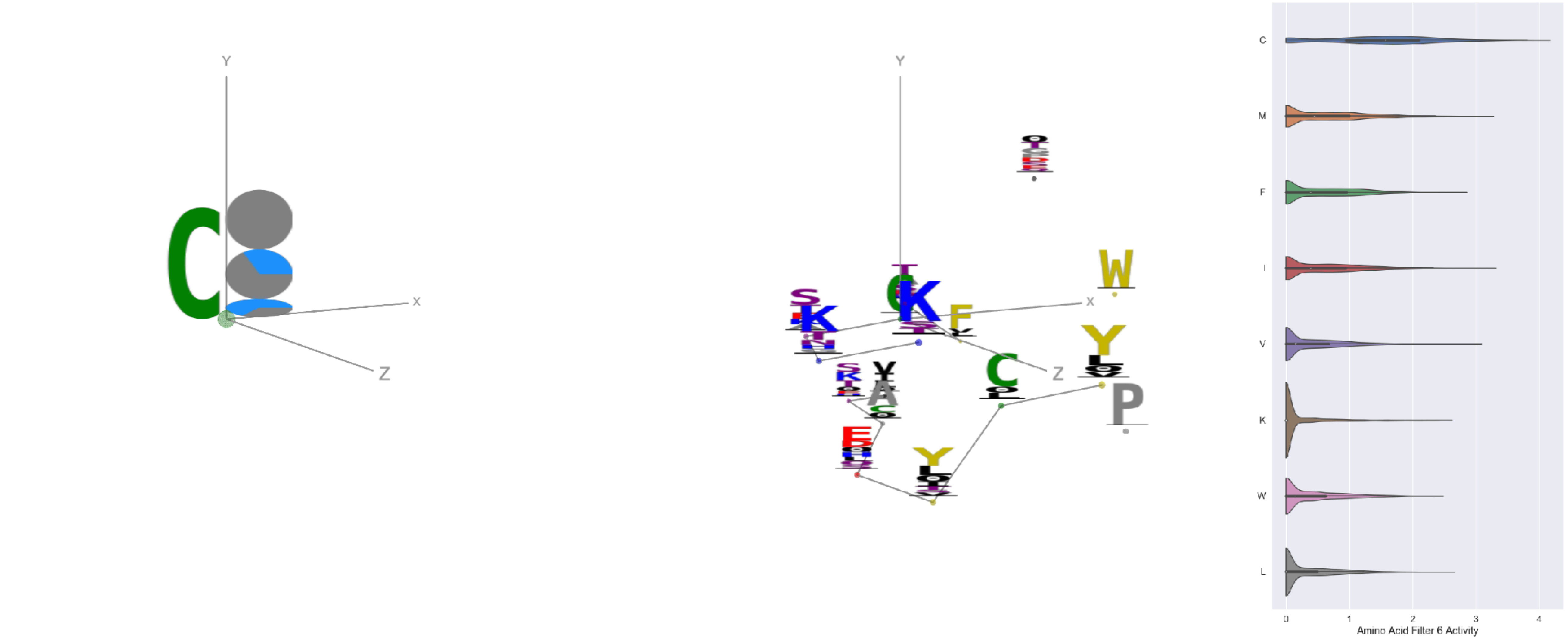


Filter 6
$R_{\mathrm{pred}} = -0.17$
Percent Active = 26
Pooling attention 0.06
Pooling feature 0.38
Top activators
4uta:A:121 C
5n0a:A:121 C
6weq:B:316 C
6p95:c:385 C
7c2l:C:1043 C
6p91:a:385 C
7ly2:A:1043 C
3iyw:A:121 C
7l2e:A:1043 C
4uao:A:265 C

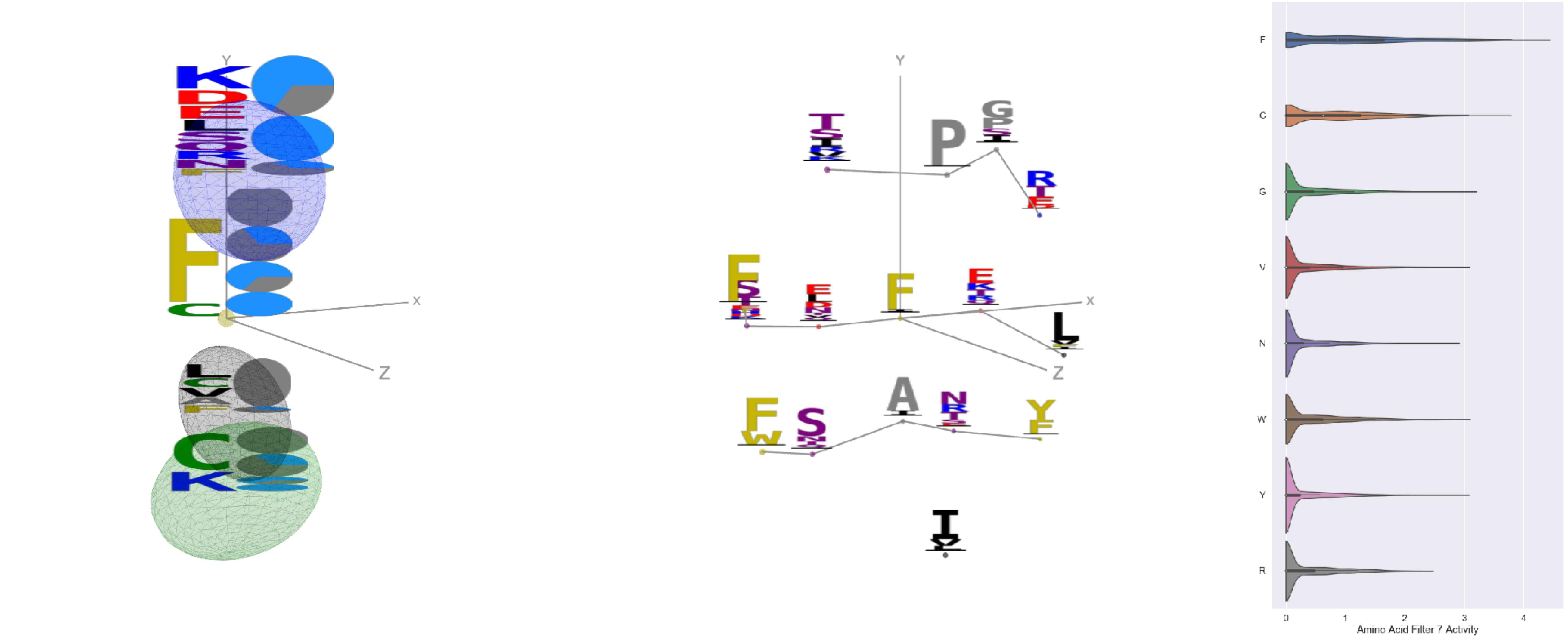


Filter 7
$R_{\mathrm{pred}} = -0.13$
Percent Active = 34
Pooling attention 0.05
Pooling feature 0.58
Top activators
2qql:A:191 F
2kh2:A:101 F
7lxx:A:157 F
7jw0:E:379 C
2kh2:A:121 F
6n5b:B:143 F
6ii9:C:136 F
6lfo:B:278 F
3tt3:A:324 F
6lfo:B:151 F

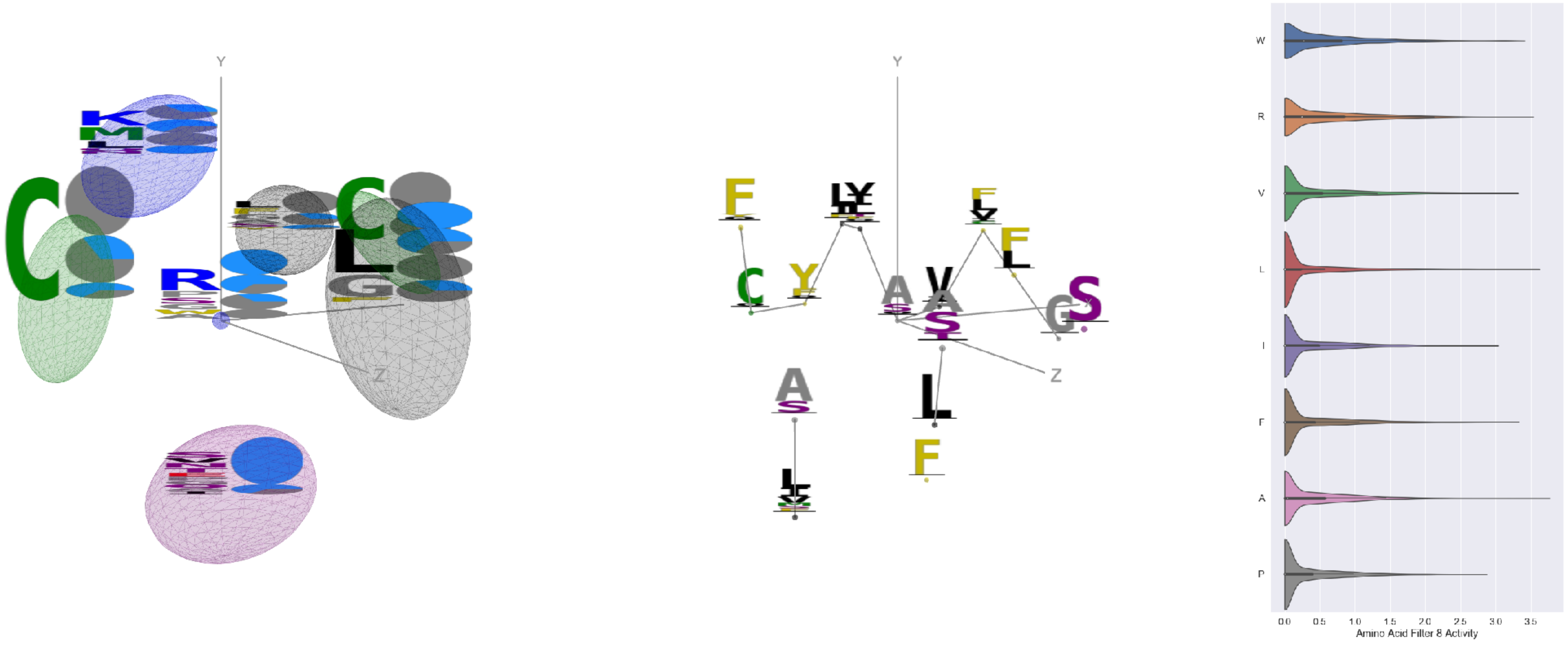


Filter 8
$R_{pred} = -0.16$
Percent Active = 34
Pooling attention 0.42
Pooling feature 0.23
Top activators
6ln2:A:399 A
6ln2:A:271 L
7lxw:A:102 R
6mej:C:606 R
6wha:B:96 W
6ii9:C:171 W
5y11:C:256 F
5tr1:B:290 V
3tt3:A:11 R
6c9u:A:385 V

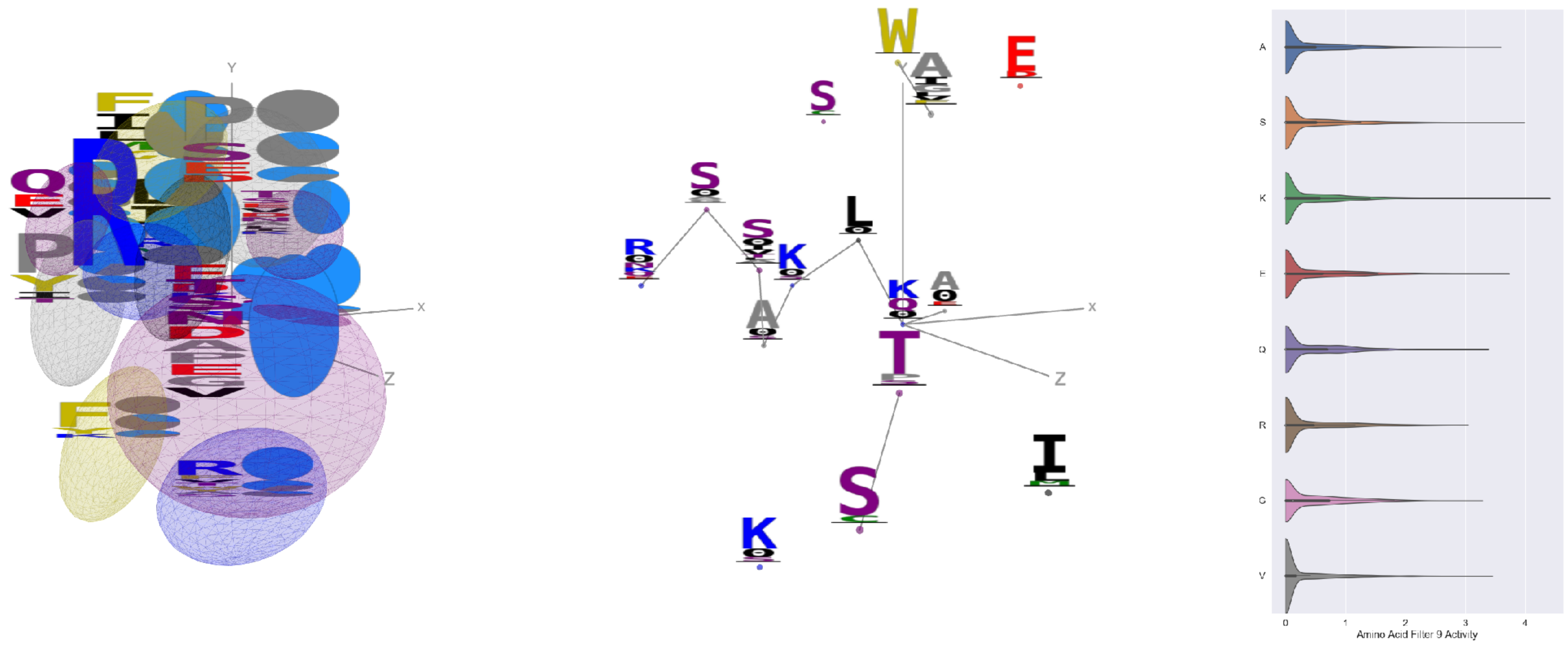


Filter 9
$R_{pred} = -0.00$
Percent Active = 35
Pooling attention 0.01
Pooling feature 1.38
Top activators
7kr5:F:157 K
6nb7:B:795 S
6xob:K:4 E
6r0x:E:44 W
4xmn:E:944 E
7lxx:A:116 S
5tr1:B:37 Y
7cwm:A:989 A
6mjz:B:373 S
1oaz:A:3 Y

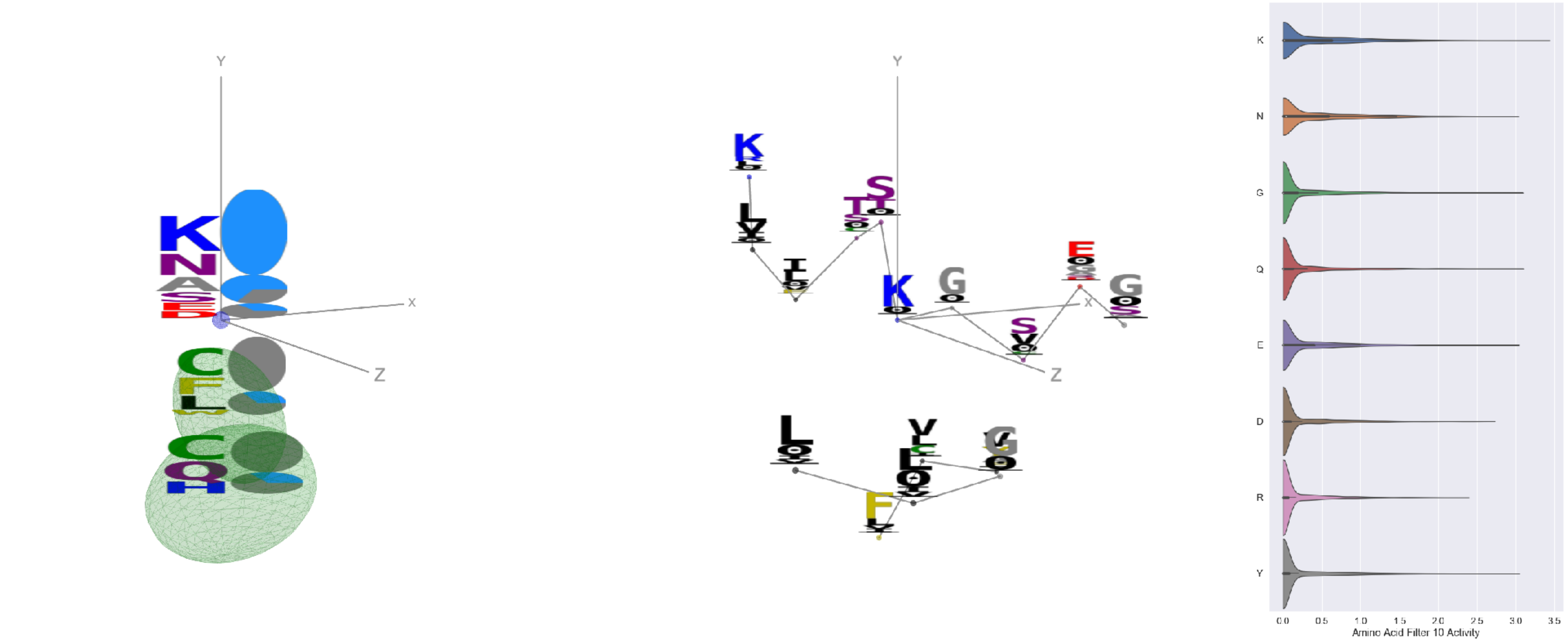


Filter 10
$R_{pred} = 0.09$
Percent Active = 25
Pooling attention 0.54
Pooling feature 0.51
Top activators
6oik:A:46 K
7ce2:A:1055 Q
7lcn:A:536 G
4uta:A:395 Y
3hmx:A:270 E
7cah:A:439 N
6m58:A:298 K
5tq2:A:380 G
6lfo:A:46 K
5tlj:X:37 Y

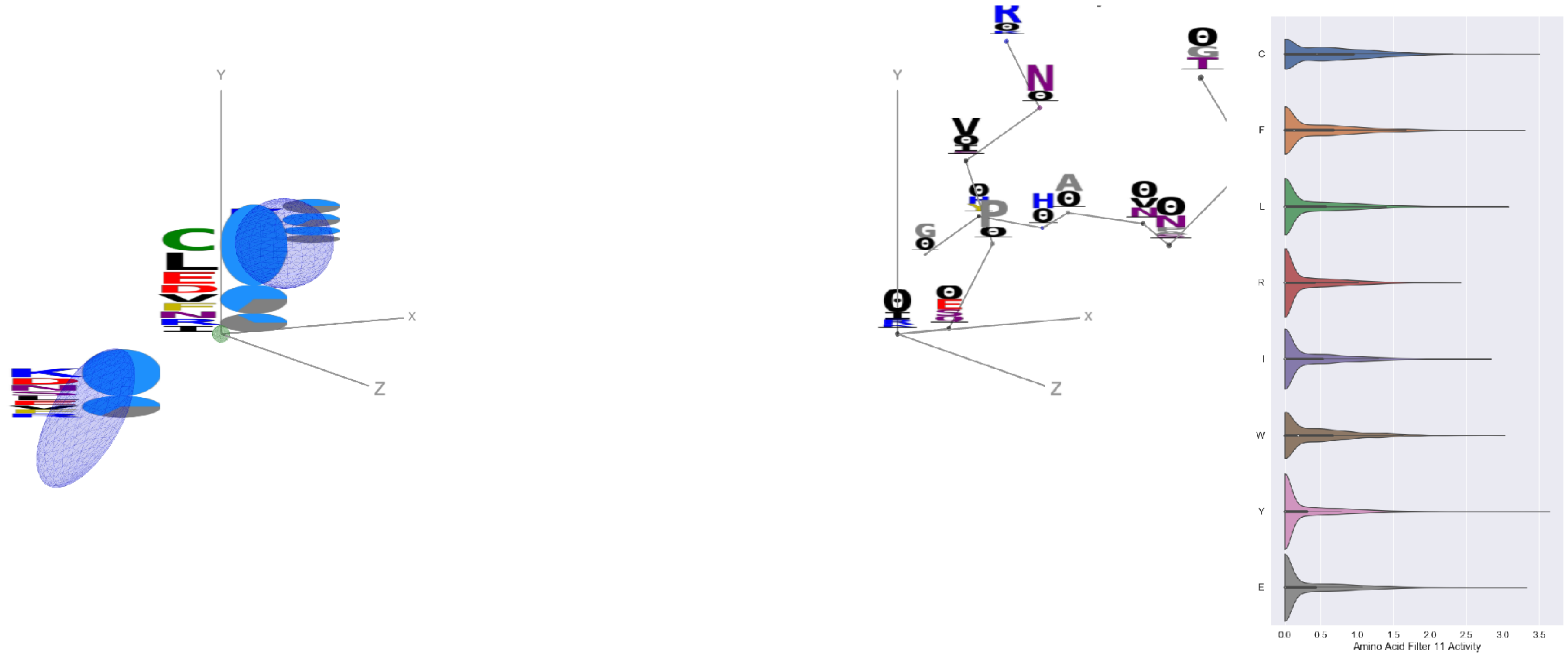


Filter 11
$R_{pred} = -0.00$
Percent Active = 36
Pooling attention 0.30
Pooling feature 0.86
Top activators
6ii9:C:317 Y
1xct:Q:45 G
4ala:C:374 G
4ala:C:382 V
1ynt:G:3113 C
7lm9:A:504 A
3wih:A:30 E
1xct:Q:24 F
4zfg:A:497 Y
6xob:K:412 L

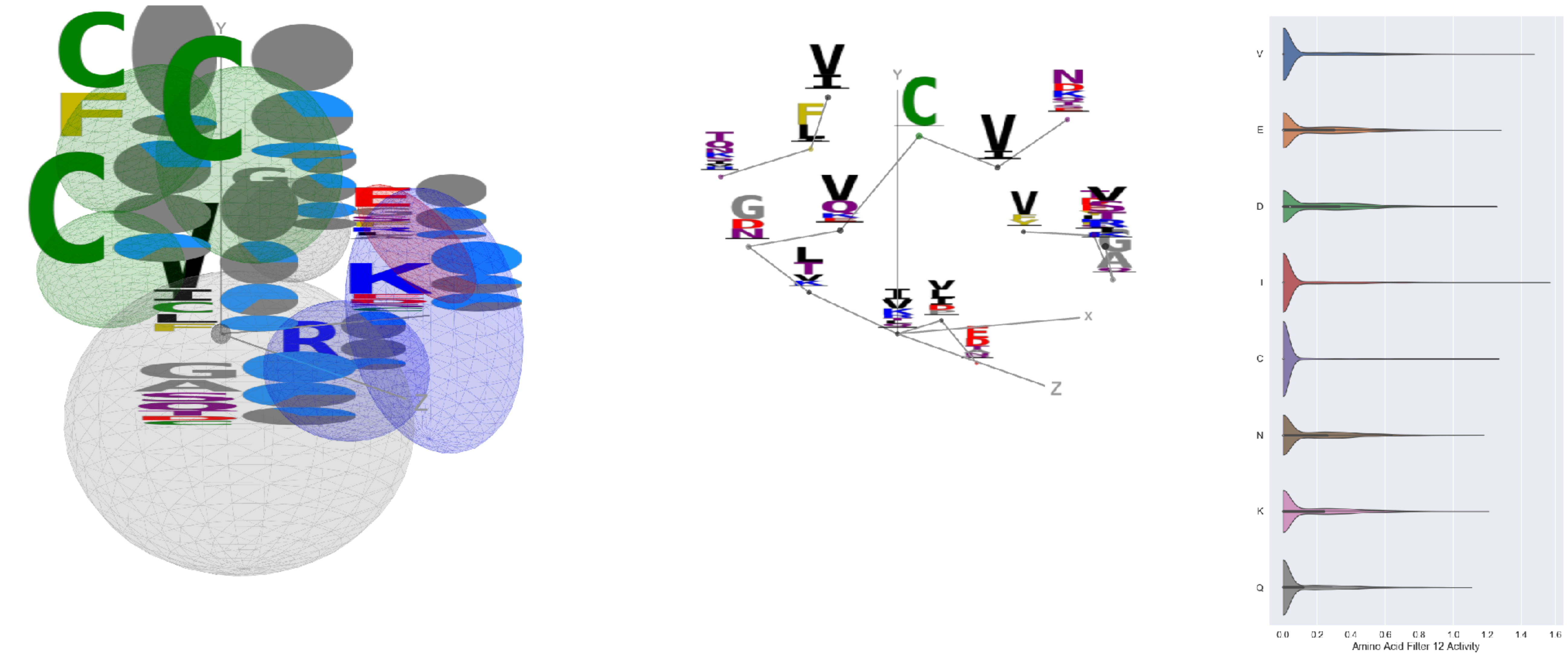


Filter 12
$R_{pred} = 0.15$
Percent Active = 33
Pooling attention 0.32
Pooling feature 3.51
Top activators
6nb7:B:520 I
7l57:B:537 V
7l57:B:1042 F
7l57:B:334 Y
7jw0:E:534 I
7l57:B:660 Y
6xob:K:214 I
6n4b:A:207 E
6nb6:C:321 Y
5f3j:A:429 C

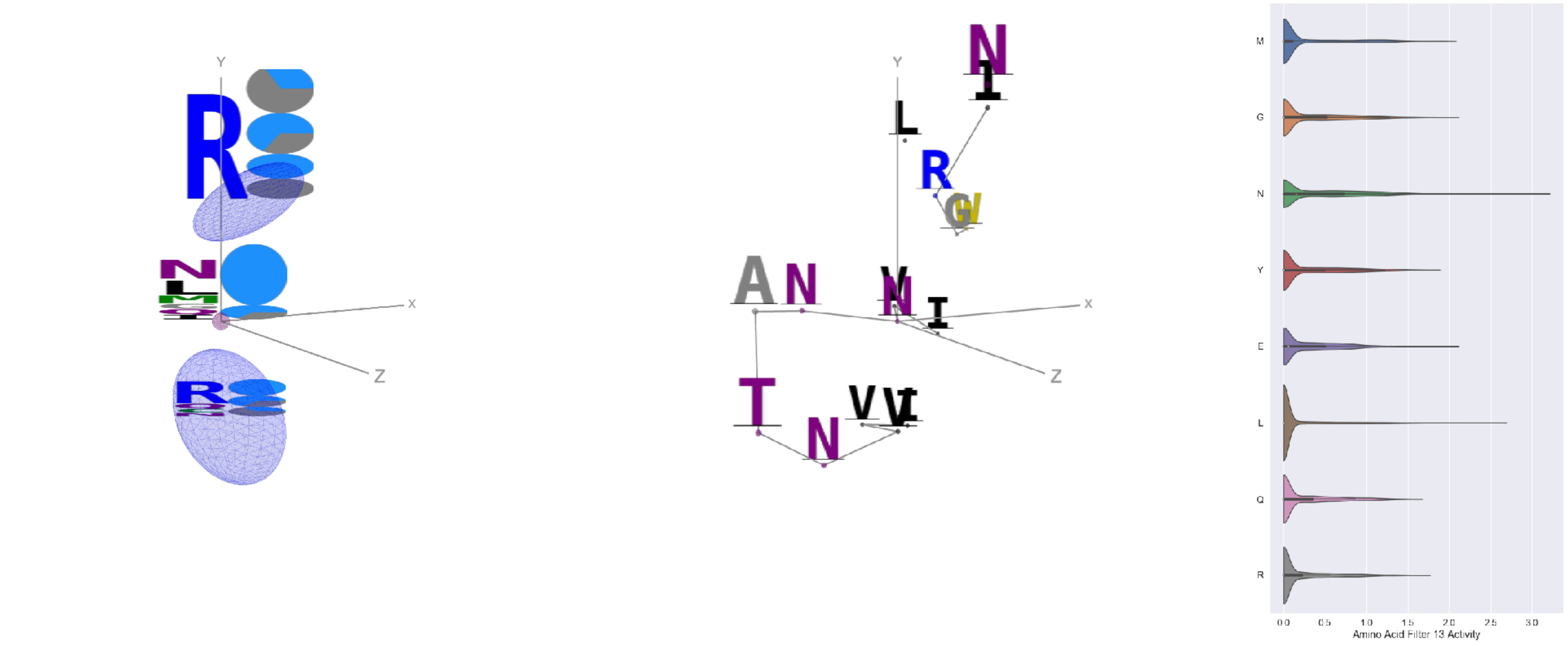


Filter 13
$R_{pred} = 0.31$
Percent Active = 34
Pooling attention 0.11
Pooling feature 0.20
Top activators
7lxx:A:121 N
6p91:a:415 L
6mei:C:645 N
7lxw:A:121 N
6wha:B:87 G
6ulf:K:46 E
6p91:a:359 M
7l2d:A:1012 L
1w72:D:120 G
6wo5:E:605 P

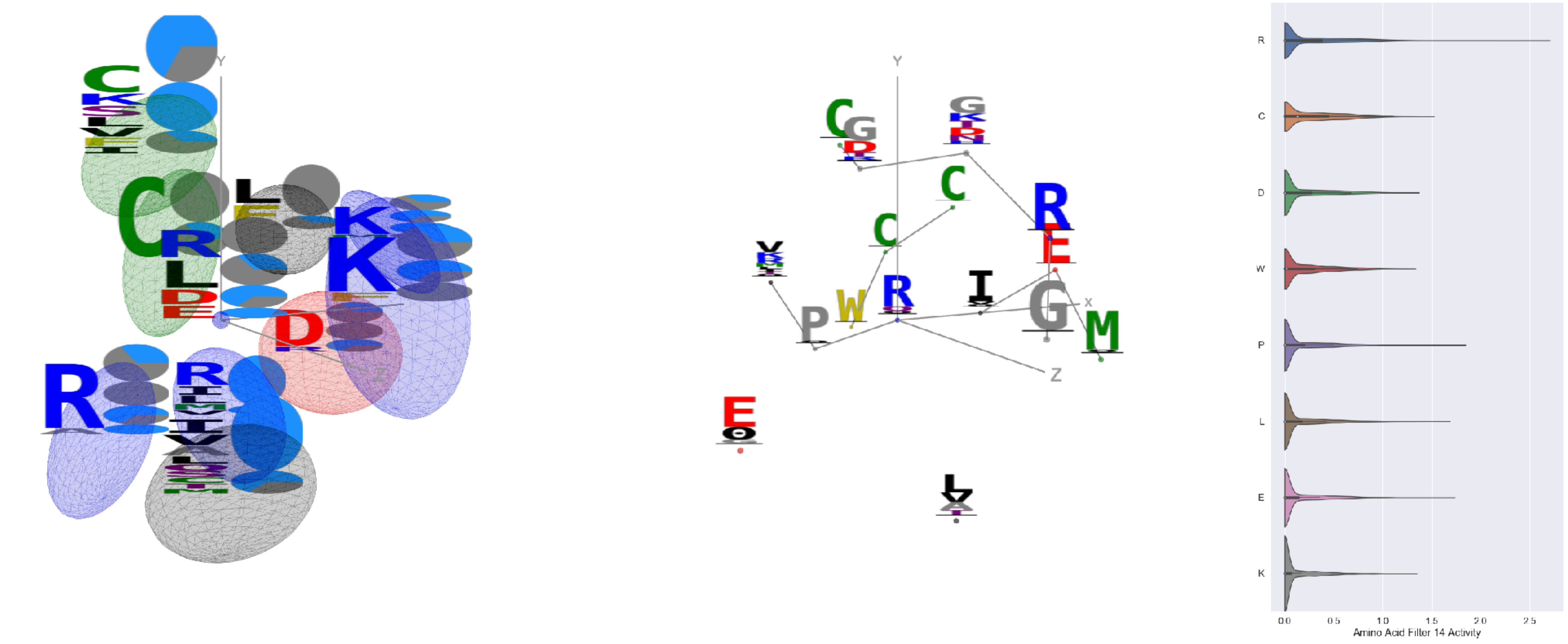


Filter 14
$R_{pred} = -0.12$
Percent Active = 25
Pooling attention 0.12
Pooling feature 0.03
Top activators
7k93:C:336 R
6weq:B:336 R
6gg0:3:54 R
6nha:A:205 P
4uta:A:62 E
6fgb:A:98 L
4zfg:A:401 L
5n0a:A:99 R
6gg0:4:54 R
7lxx:A:118 L

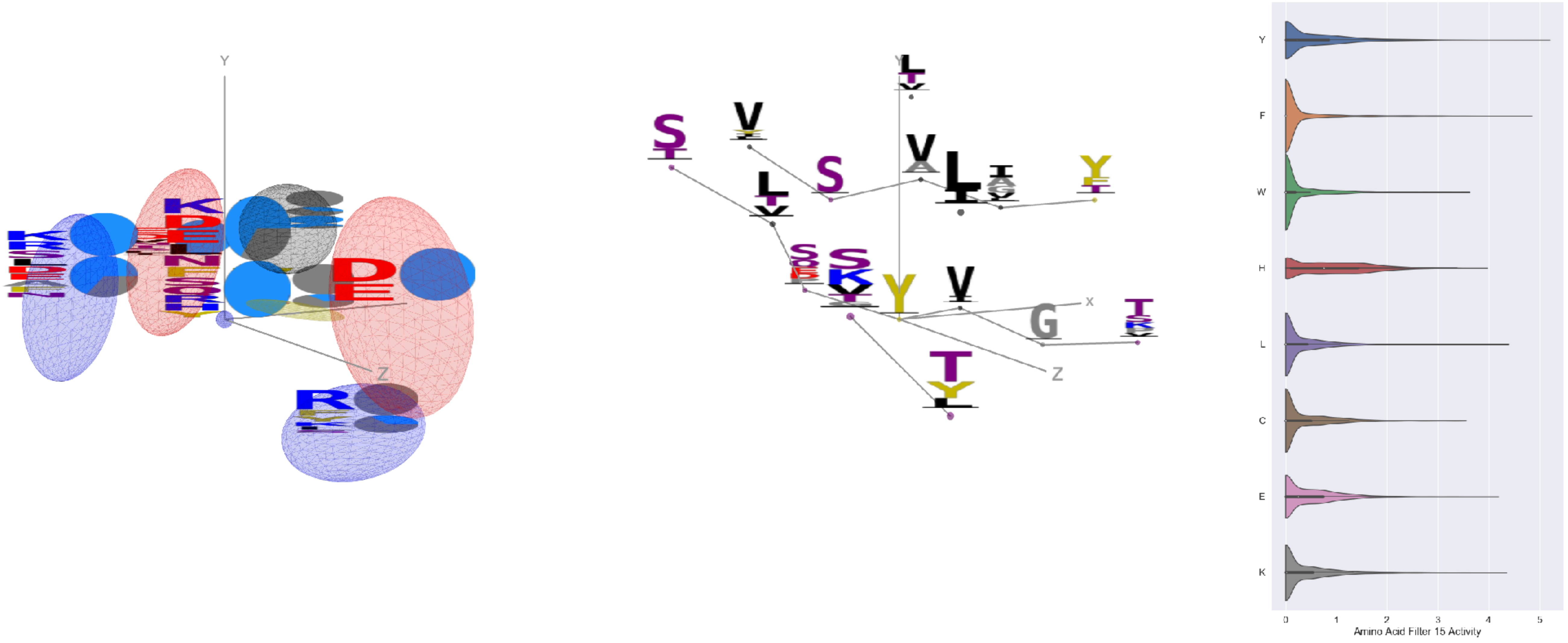


Filter 15
$R_{pred} = -0.10$
Percent Active = 48
Pooling attention 0.44
Pooling feature 0.96
Top activators
7l57:B:313 Y
4xmn:E:886 Y
4xmn:E:927 F
7l57:B:318 Y
4xmn:E:982 Y
7l57:B:38 Y
7jw0:E:154 Y
7l57:B:1067 Y
6nb6:C:325 F
6wpt:C:110 Y

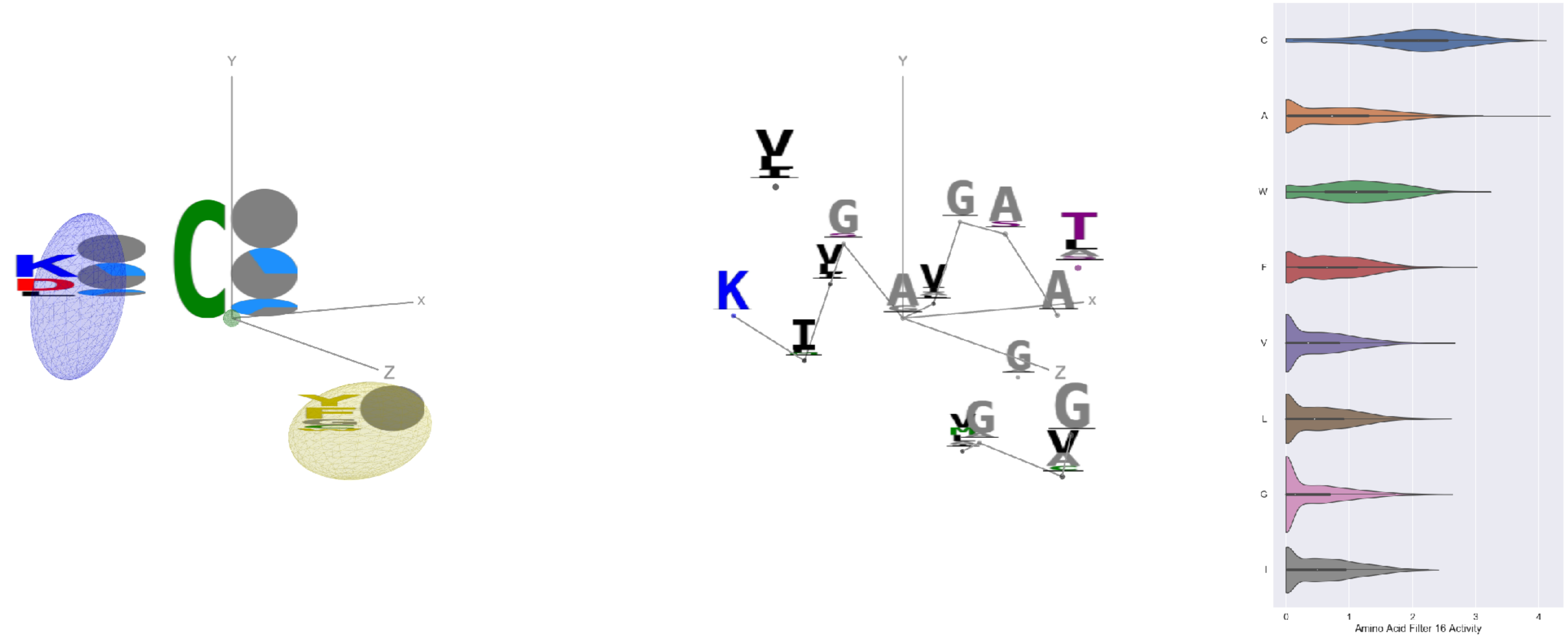


Filter 16
$R_{pred} = -0.28$
Percent Active = 45
Pooling attention 0.13
Pooling feature 0.38
Top activators
6c9u:A:386 A
7lcn:A:525 C
6nb6:A:511 C
6i07:D:66 C
4kxz:E:48 C
7ly2:A:432 C
7jv6:E:525 C
6mei:C:508 C
4kxz:E:44 C
7lm9:A:511 C

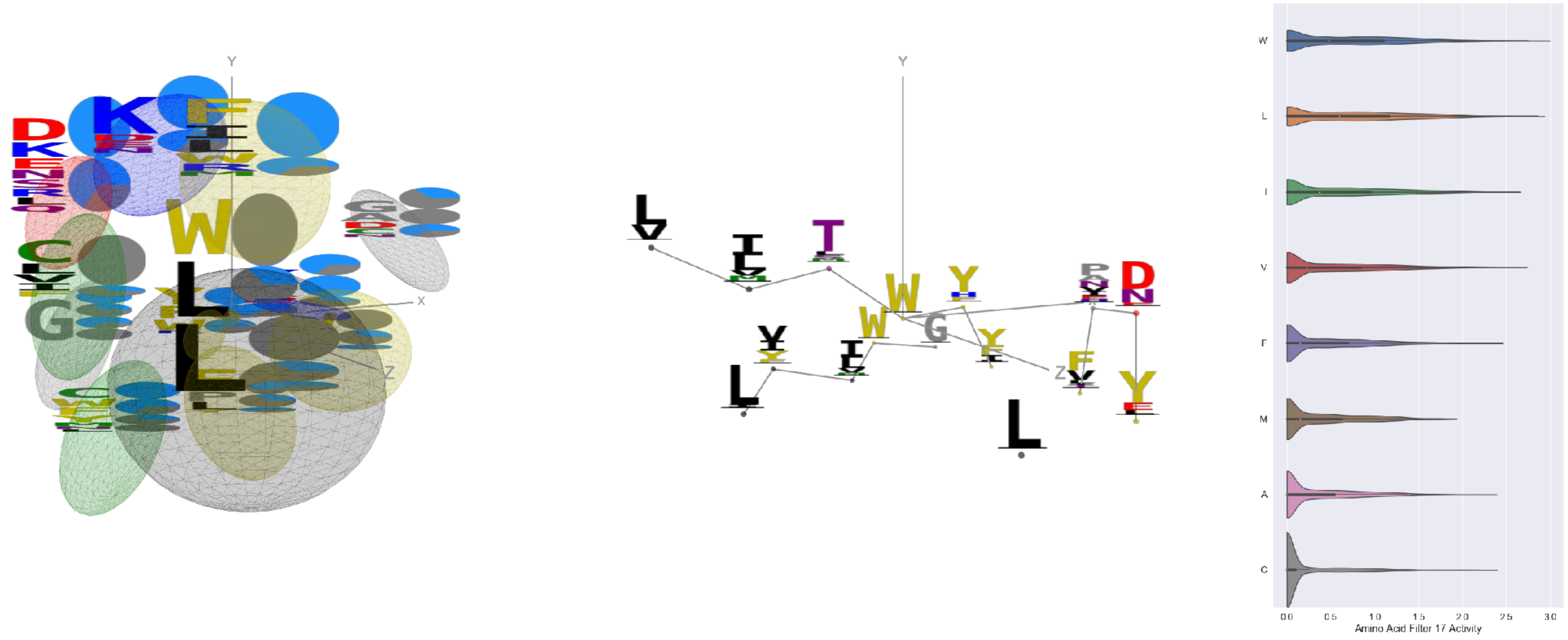


Filter 17
$R_{pred} = -0.08$
Percent Active = 47
Pooling attention 0.15
Pooling feature 1.17
Top activators
6mlm:A:250 W
7ebz:C:209 L
5kqv:E:62 L
6ii9:C:225 W
4fqv:C:234 W
7ebz:B:209 L
6aj7:B:209 L
6wdt:B:209 L
6ii9:C:171 W
6usf:D:237 L

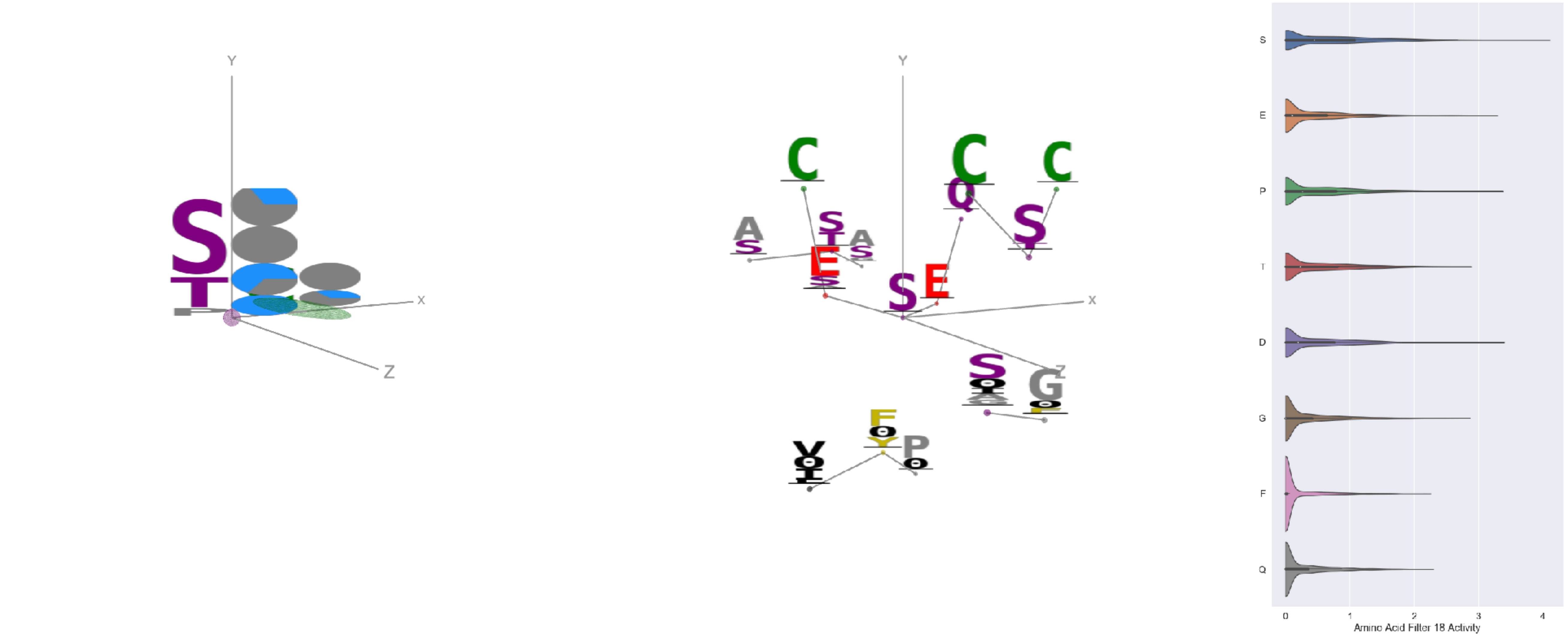


Filter 18
$R_{pred} = -0.04$
Percent Active = 27
Pooling attention 0.05
Pooling feature 0.66
Top activators
6pzw:C:228 S
5i6x:A:438 S
6c9u:A:206 S
5i6x:A:437 D
6q23:A:228 S
3tt3:A:57 P
7nda:A:1031 E
5y11:C:331 S
1nca:N:228 S
6c9u:A:371 S

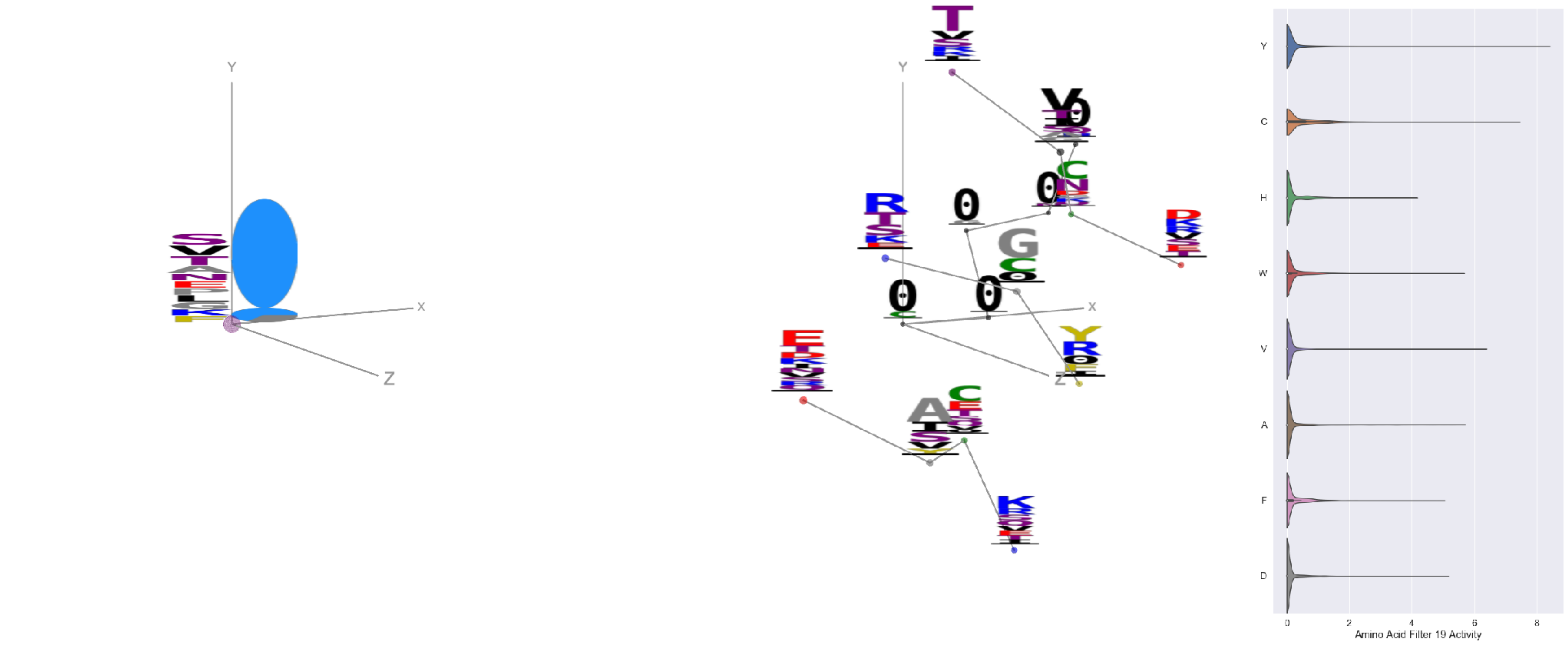


Filter 19
$R_{pred} = -0.06$
Percent Active = 16
Pooling attention 0.08
Pooling feature 1.89
Top activators
3bsz:E:174 Y
7kr5:F:305 Y
5tlj:X:84 C
4mwf:D:585 C
1jrh:I:105 Y
4xmn:E:726 R
5tq2:B:390 Y
6ln2:A:421 Y
4g80:I:244 Y
5otj:C:71 Y

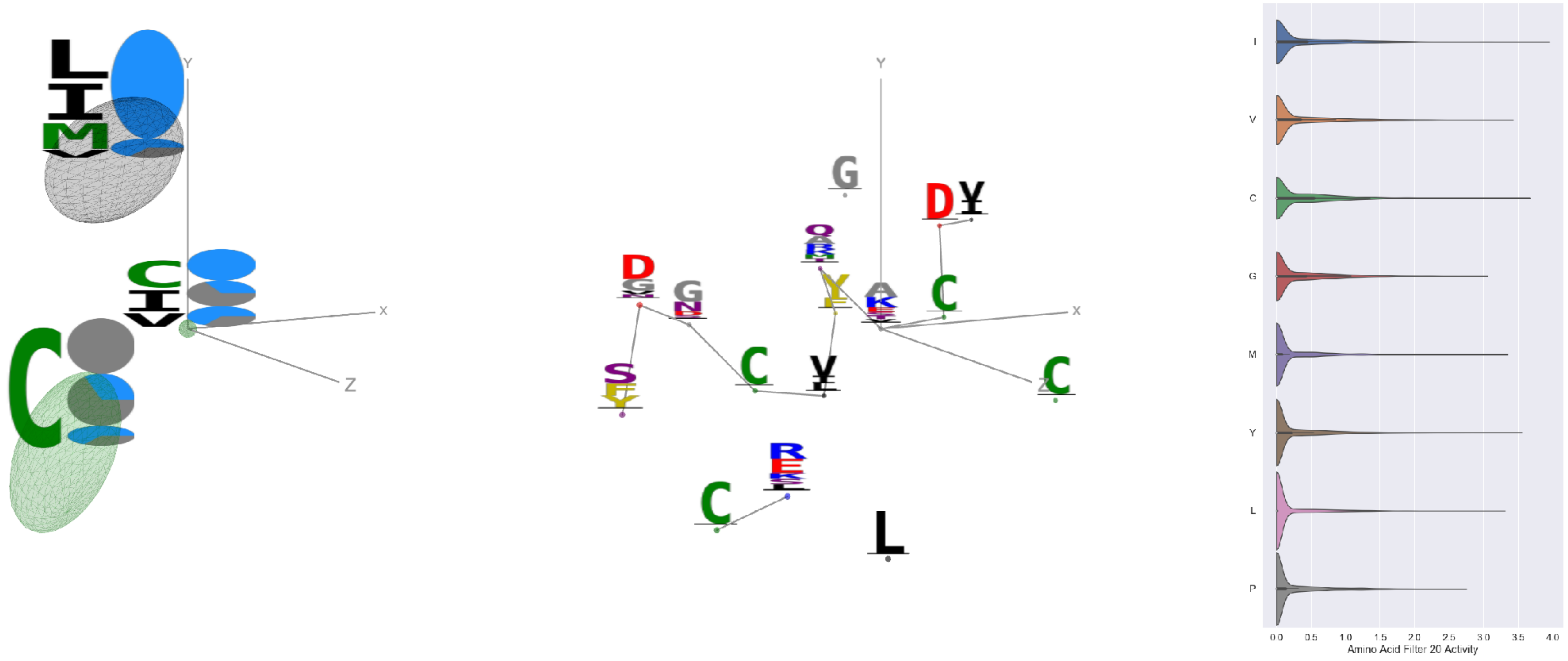


Filter 20
$R_{\mathrm{pred}} = 0.07$
Percent Active = 28
Pooling attention 0.17
Pooling feature 0.18
Top activators
6nb7:B:721 A
6mjz:C:323 I
5y11:C:292 C
4okv:F:205 I
7c2l:C:642 Y
6mjz:B:400 I
5ggv:Y:108 Y
6usf:D:222 V
5esz:C:115 I
7bud:B:301 M

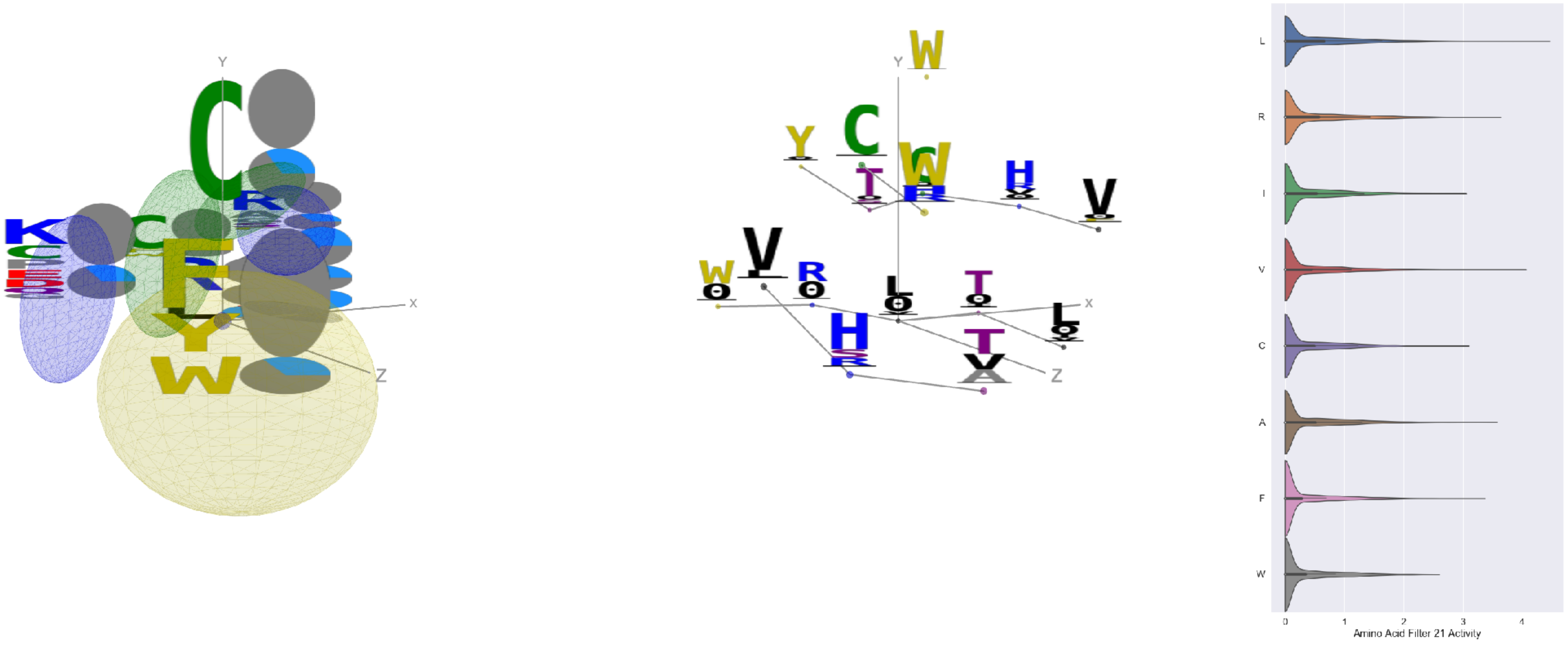


Filter 21
$R_{\mathrm{pred}} = -0.14$
Percent Active = 29
Pooling attention 0.15
Pooling feature 0.20
Top activators
1w72:D:272 L
7nda:A:289 V
7nd9:B:289 V
7l2e:A:289 V
3hae:A:272 L
7nda:C:289 V
6nha:A:265 Y
7ly2:A:289 V
5f3j:A:391 R
7l2d:A:289 V

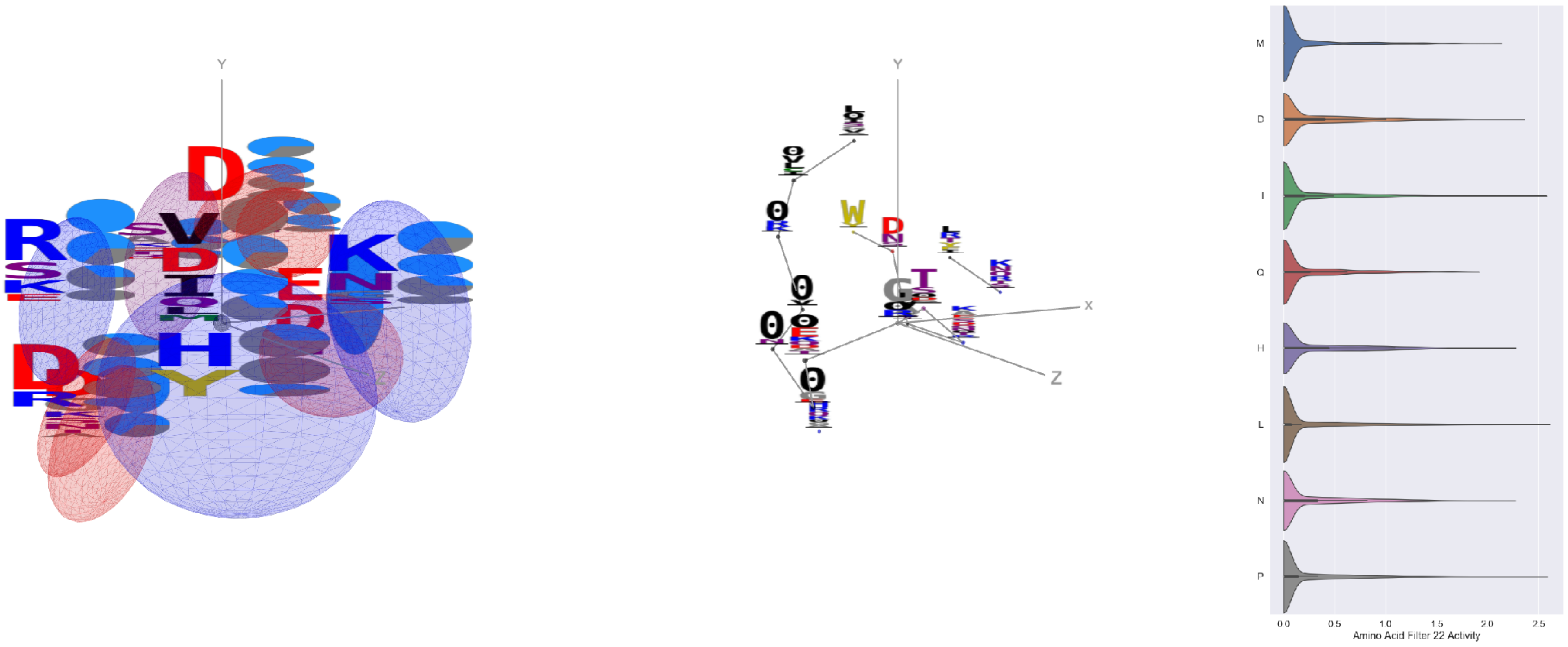


Filter 22
$R_{\mathrm{pred}} = 0.20$
Percent Active = 29
Pooling attention 0.62
Pooling feature 1.41
Top activators
6lfo:B:129 G
6mjz:B:372 L
6p91:a:275 P
1xct:M:28 I
6oik:A:28 Y
6gg0:3:331 V
6gv4:A:62 Y
4okv:F:239 C
6gg0:4:331 V
7l57:B:715 P

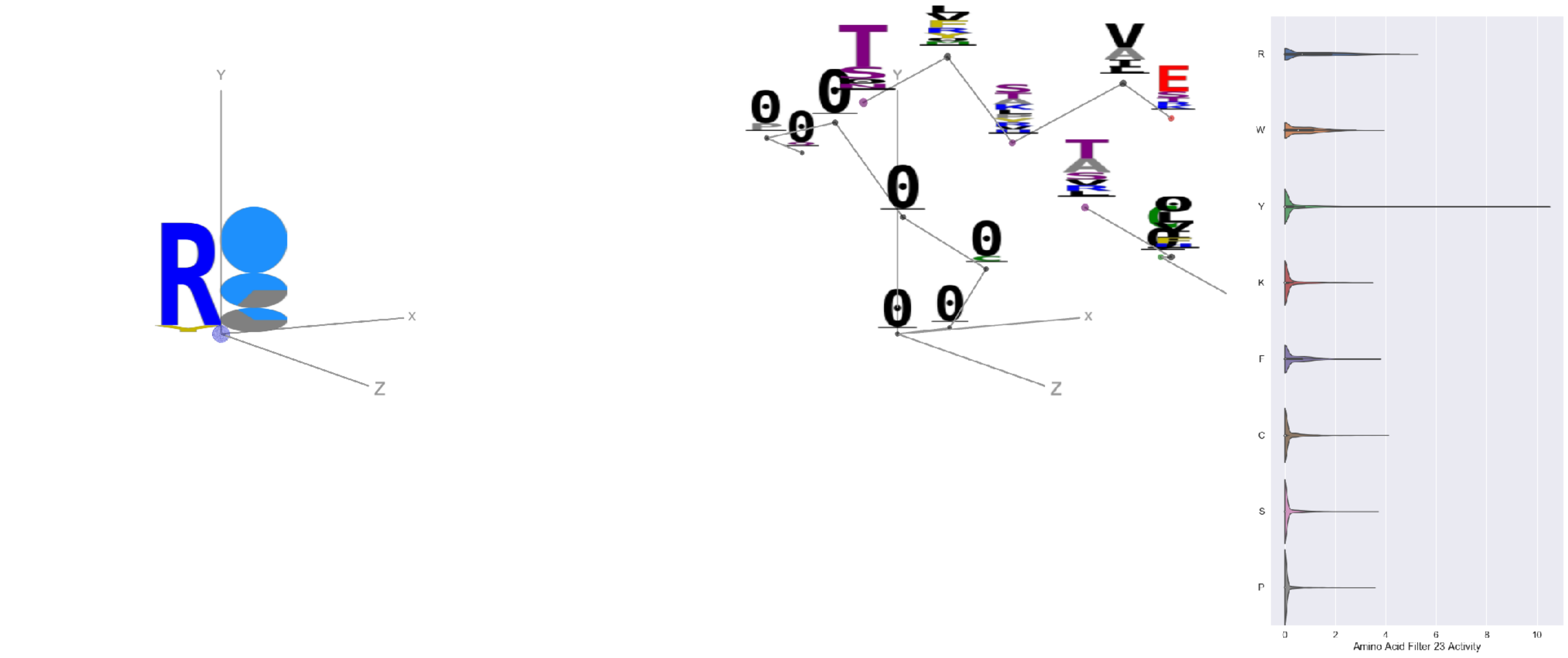


Filter 23
$R_{\mathrm{pred}} = 0.06$
Percent Active = 25
Pooling attention 0.18
Pooling feature 0.18
Top activators
1qfw:B:112 Y
6ii9:C:317 Y
6otc:A:294 R
6p95:C:256 R
7bud:B:471 R
4kuc:A:197 G
7jvr:A:242 R
4mwf:D:543 R
4okv:F:266 R
5vod:C:86 R

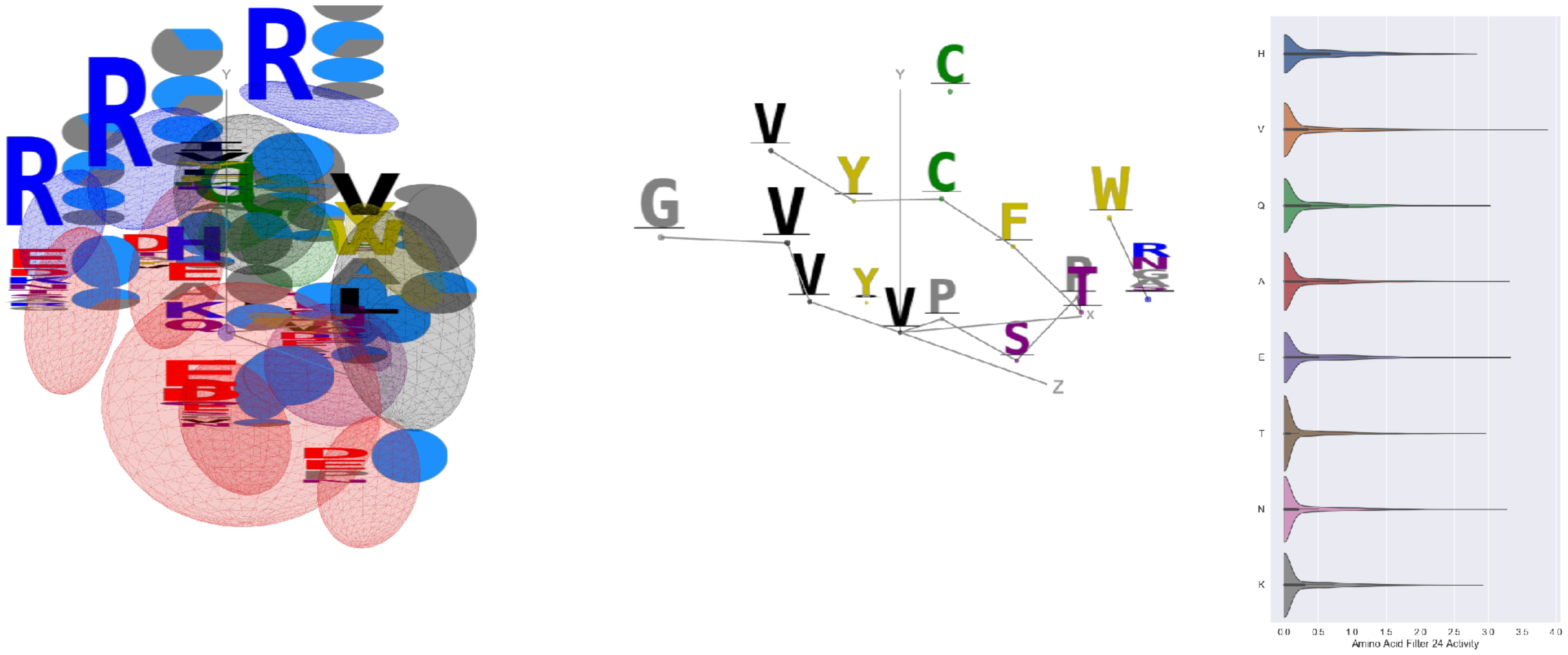


Filter 24
$R_{\text{pred}} = -0.00$
Percent Active = 31
Pooling attention 0.39
Pooling feature 1.76
Top activators
4mwf:D:514 V
6mei:C:514 V
6o25:G:109 V
6ulf:K:48 V
6o25:G:48 V
6mej:C:514 V
6o2b:M:48 V
6o25:G:86 D
6i07:D:258 E
4i3r:G:266 A

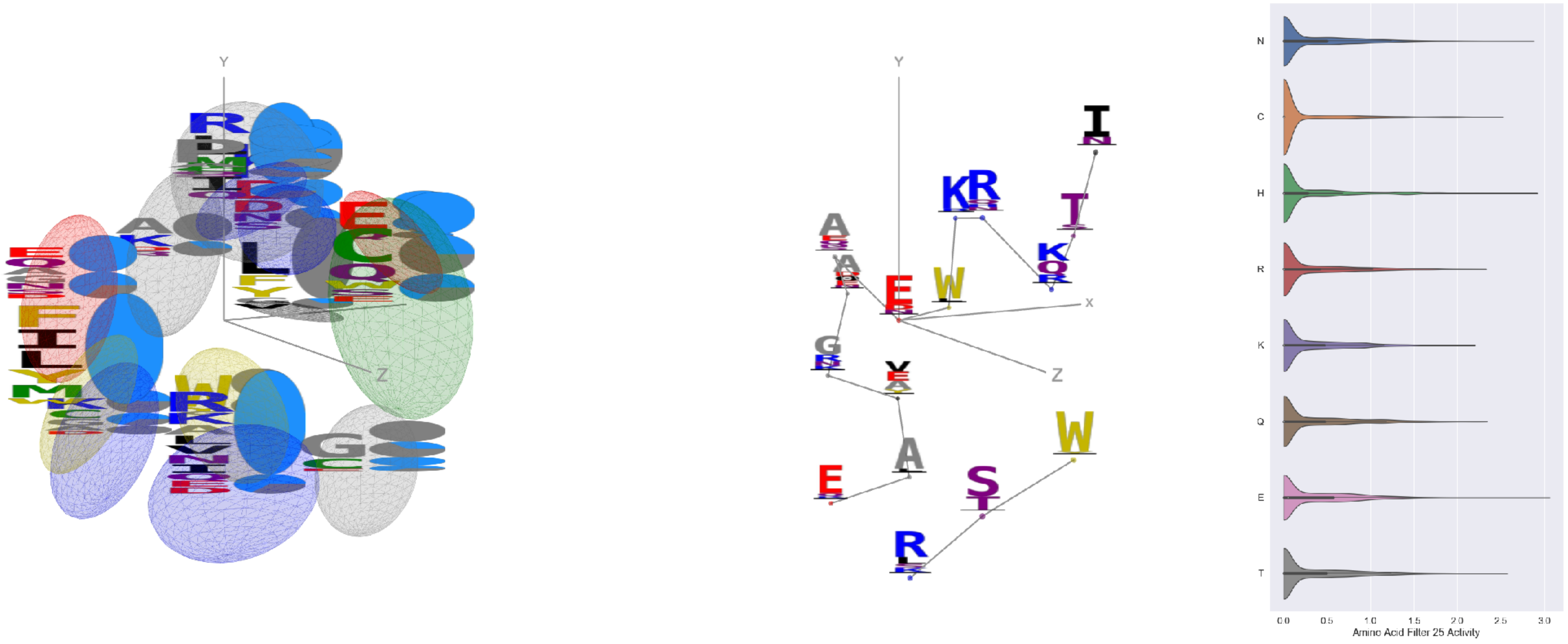


Filter 25
$R_{\text{pred}} = 0.09$
Percent Active = 36
Pooling attention 0.47
Pooling feature 3.35
Top activators
3hae:A:148 E
4zxb:E:429 H
6nb7:B:746 N
6nha:A:142 E
7jw0:E:952 V
6nb6:C:380 S
6fgb:A:142 E
6m58:A:319 F
5f3j:A:458 E
7l57:B:952 V

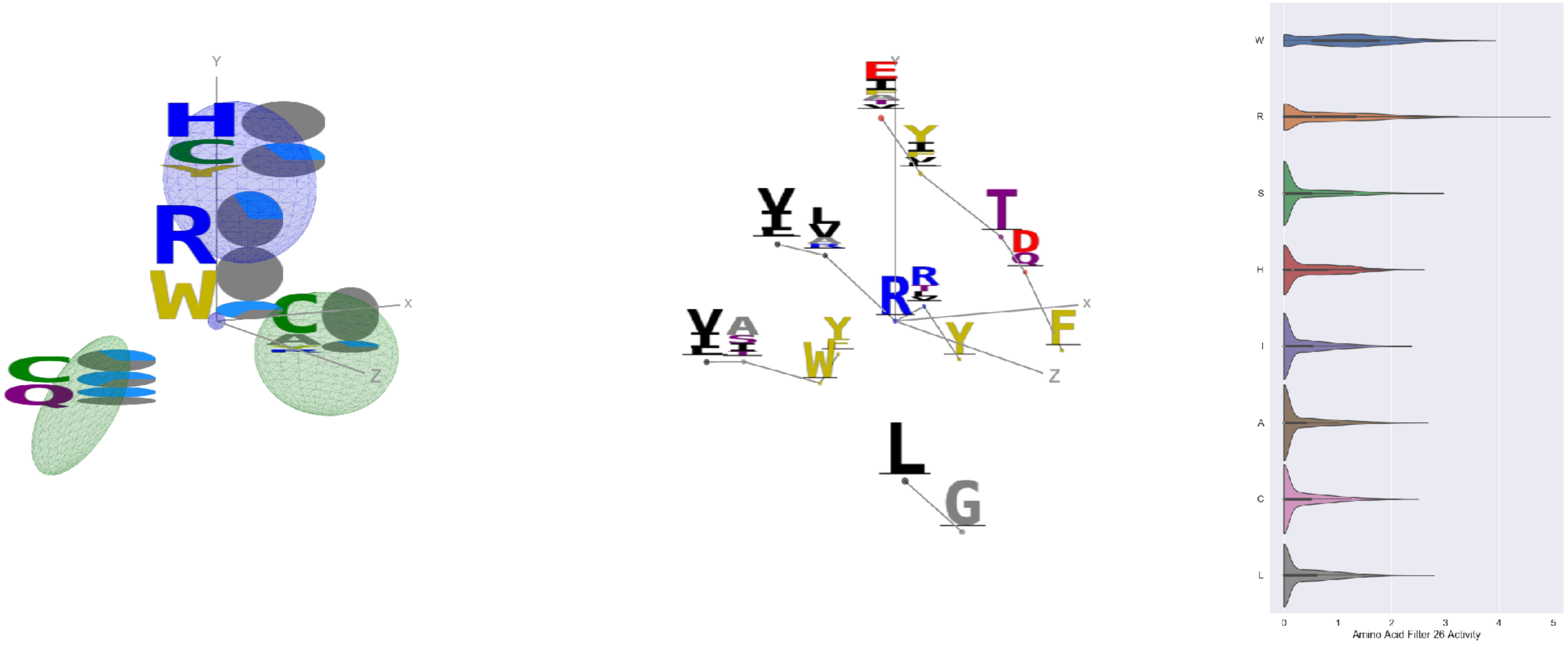


Filter 26
$R_{\text{pred}} = -0.20$
Percent Active = 35
Pooling attention 0.26
Pooling feature 0.34
Top activators
3w9e:C:166 R
4fqv:C:234 W
5y11:C:229 W
6wdt:A:223 R
6mlm:A:250 W
1nca:N:364 R
6pzw:C:300 R
6ur5:C:234 W
6pzw:C:364 R
7ce2:A:991 W

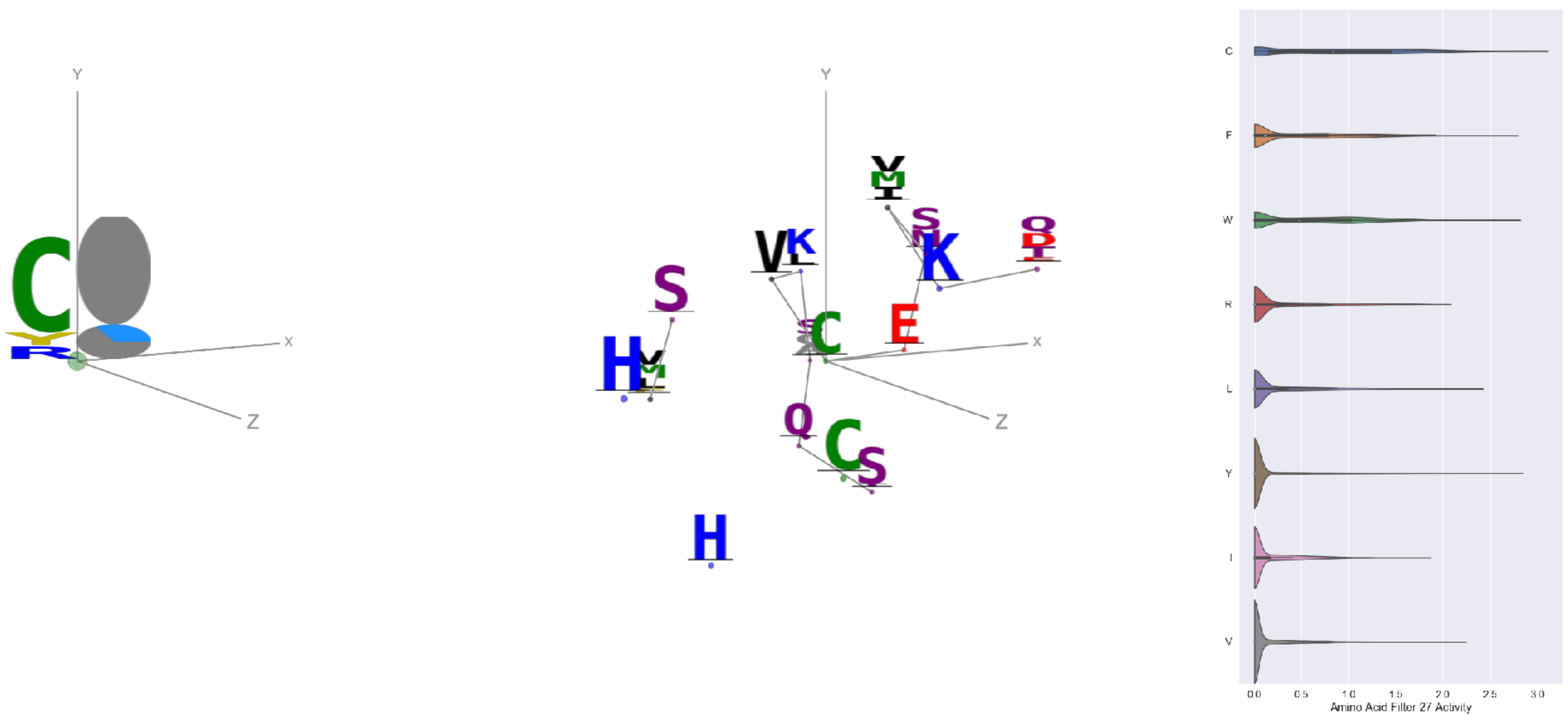


Filter 27
$R_{\text{pred}} = -0.18$
Percent Active = 18
Pooling attention 0.03
Pooling feature 0.78
Top activators
7l2d:A:1032 C
7ly2:A:1032 C
4i3r:G:296 C
7nd9:B:612 Y
6c9u:A:638 W
6nb7:B:1014 C
7l2e:A:612 F
6wdt:A:112 Y
6mei:C:581 C
6mej:C:607 C

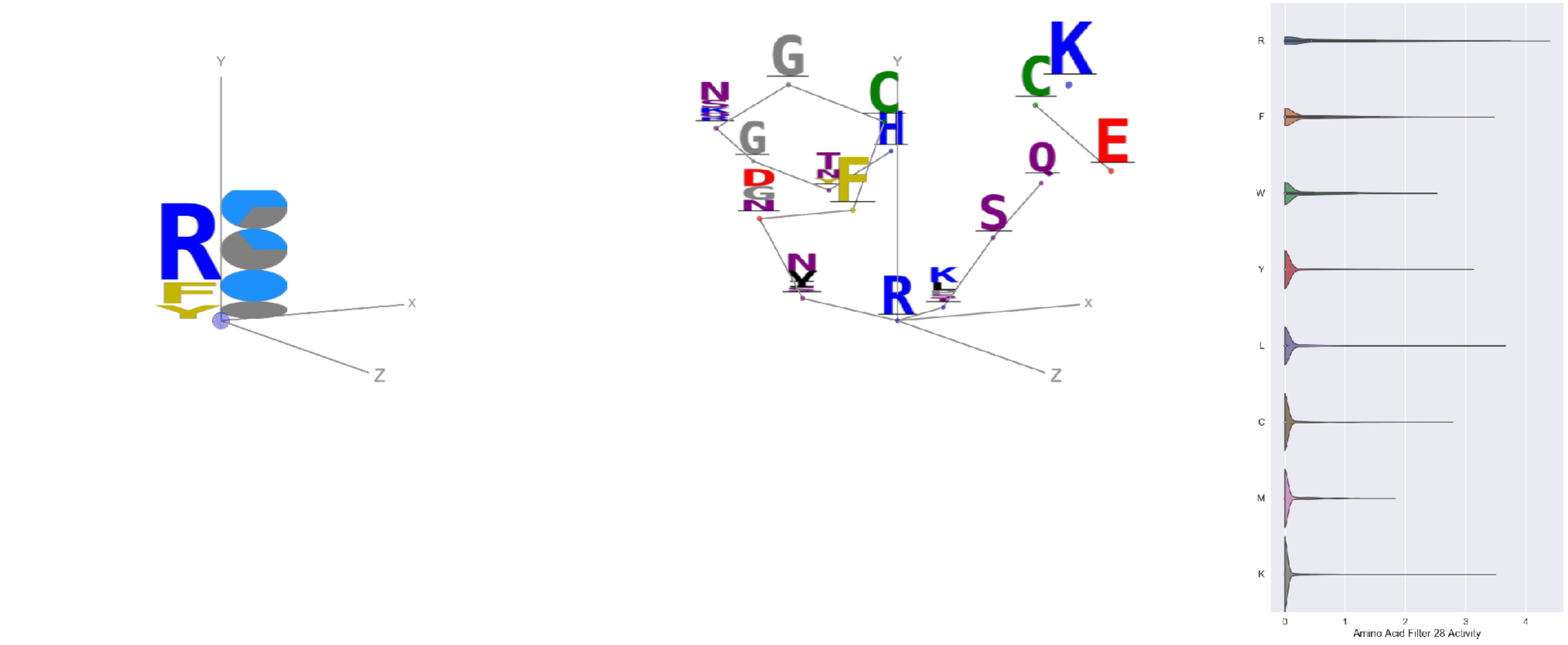

Filter 28
$R_{\text{pred}} = 0.04$
Percent Active = 15
Pooling attention 0.28
Pooling feature 0.23
Top activators
7ly2:A:1039 R
7bud:B:99 R
7jv6:E:1039 R
6fgb:A:176 R
6o2b:M:66 R
6ulf:K:67 R
7c2l:C:1039 R
6o25:G:66 R
7nd9:B:1039 R
6wpt:C:1039 R

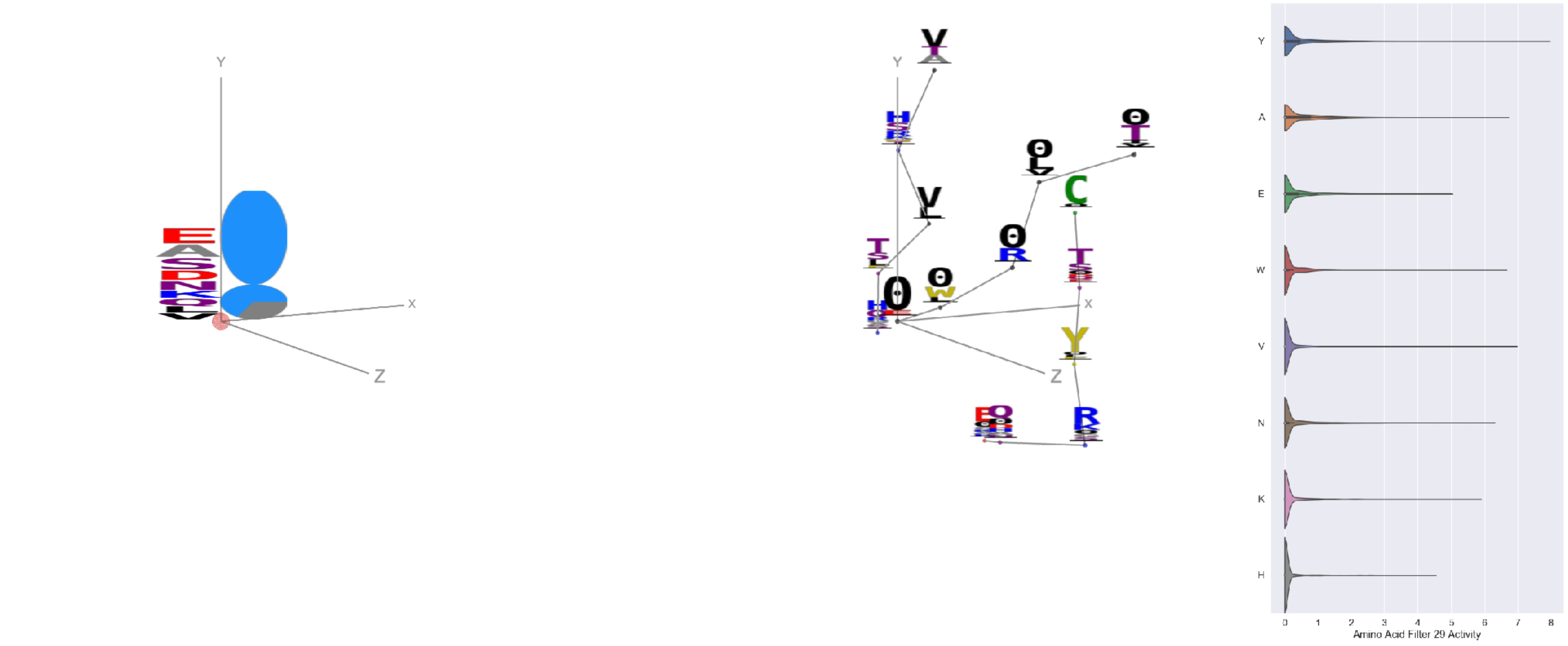

Filter 29
$R_{\text{pred}} = -0.07$
Percent Active = 25
Pooling attention 0.10
Pooling feature 0.74
Top activators
6fgb:A:268 Y
6nb6:C:22 Y
4xmn:E:726 R
7ly2:A:20 Y
6m58:A:582 Y
6ln2:A:421 Y
4uta:A:395 Y
5otj:C:71 Y
7kr5:F:305 Y
6ln2:A:257 V

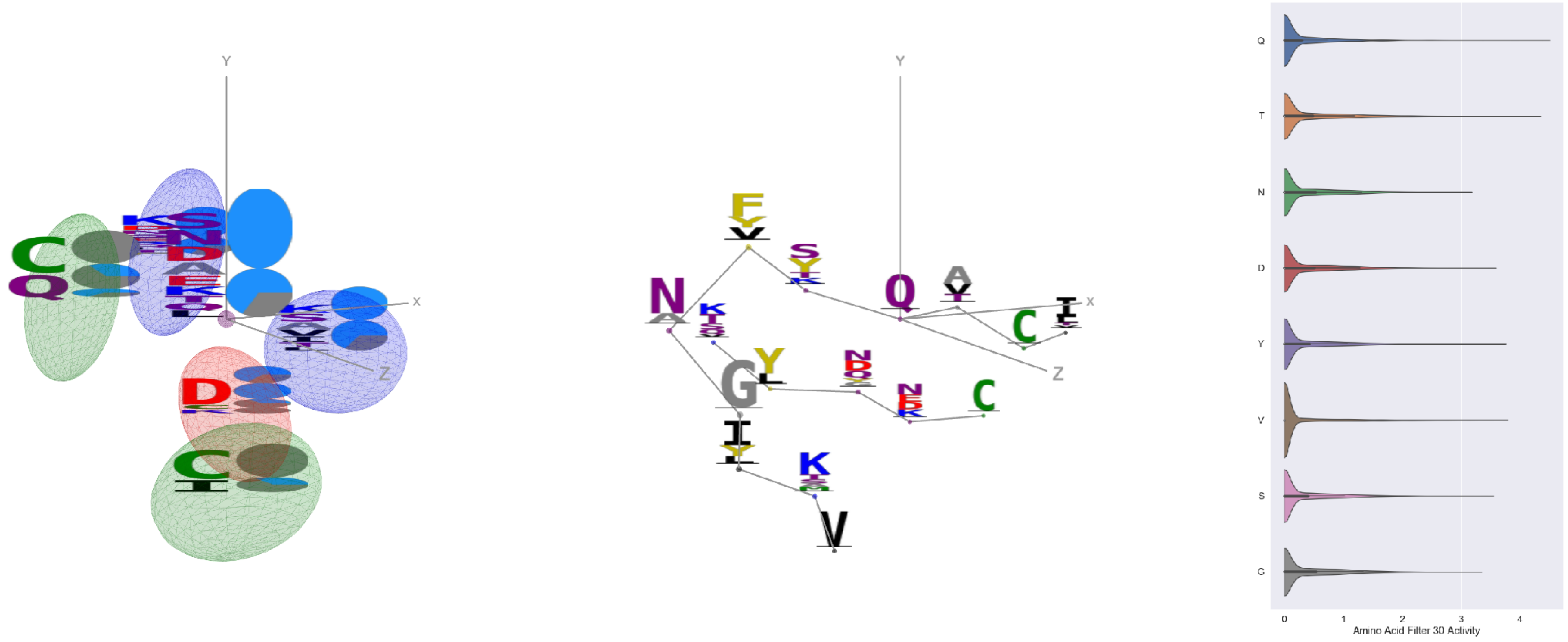

Filter 30
$R_{\text{pred}} = 0.11$
Percent Active = 32
Pooling attention 0.43
Pooling feature 0.82
Top activators
6nb6:A:835 Q
7jw0:E:558 T
6nb7:B:835 Q
7l57:B:558 T
7nd9:B:558 T
6nb7:B:334 V
6wpt:C:134 Y
6nb7:B:364 F
6nb6:C:364 F
6nb6:A:322 D

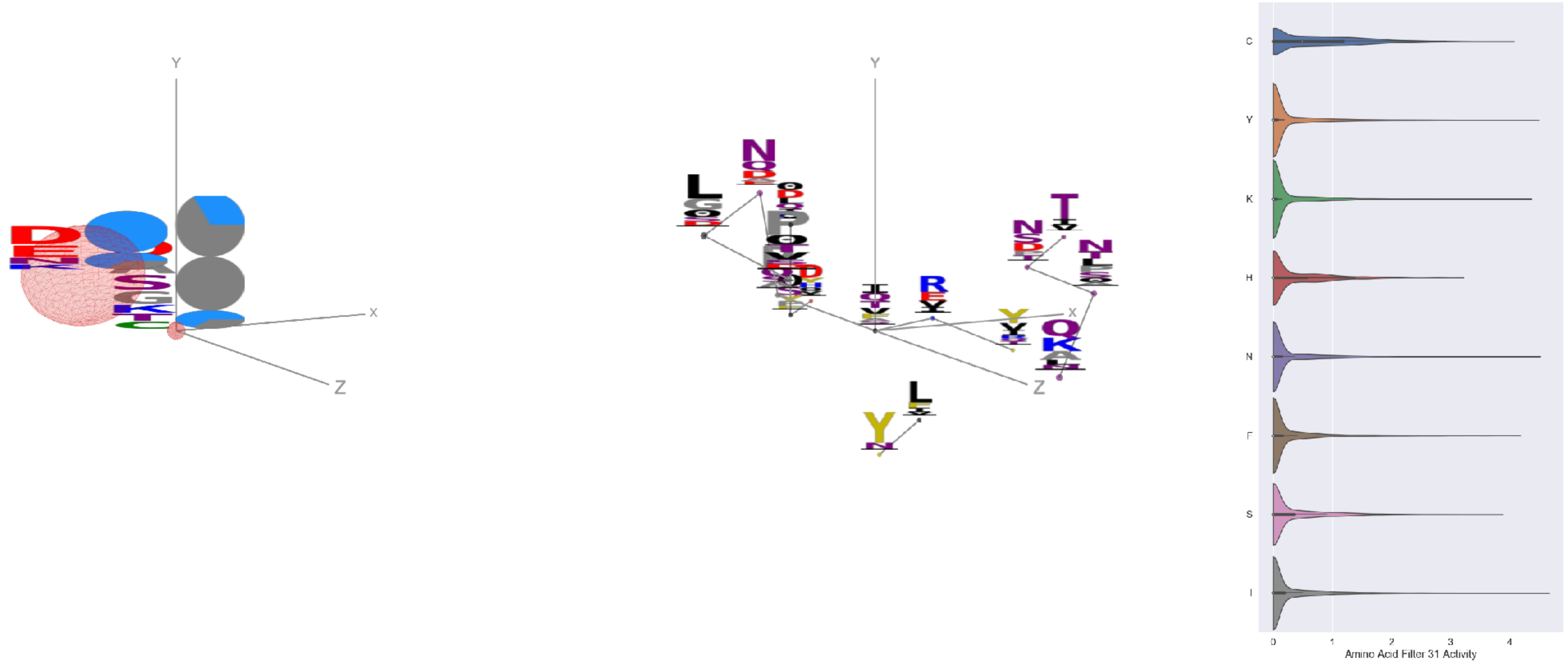

Filter 31
$R_{\text{pred}} = -0.21$
Percent Active = 31
Pooling attention 0.33
Pooling feature 1.22
Top activators
6nb7:B:316 I
7l57:B:819 E
6nb7:B:1111 N
7jw0:E:473 Y
7l57:B:300 K
7jw0:E:133 Y
7l57:B:279 F
7jw0:E:454 Y
7l57:B:733 K
6nb7:B:361 V

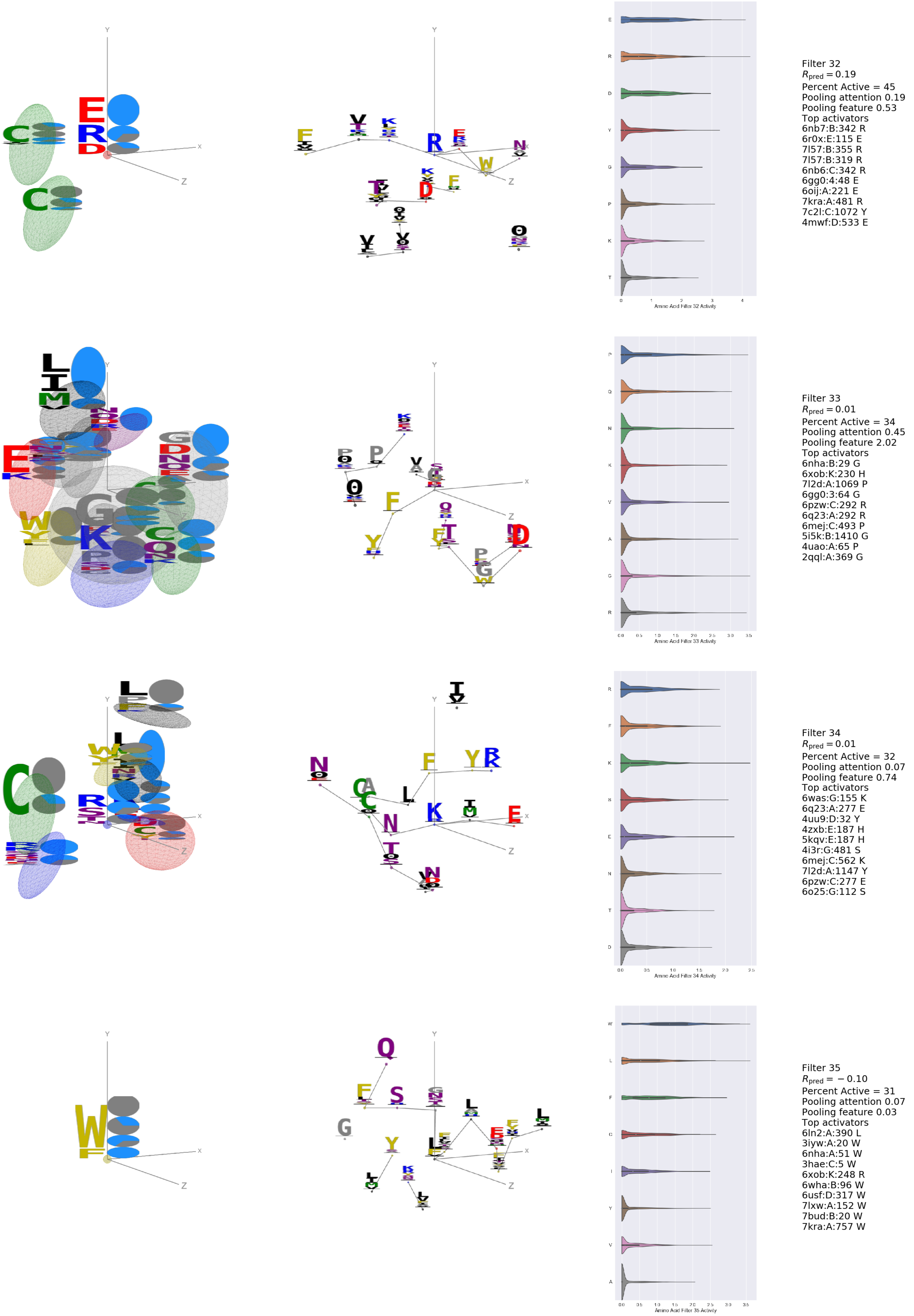

Filter 32
$R_{pred} = 0.19$
Percent Active = 45
Pooling attention 0.19
Pooling feature 0.53
Top activators
6nb7:B:342 R
6r0x:E:115 E
7l57:B:355 R
7l57:B:319 R
6nb6:C:342 R
6gg0:4:48 E
6oij:A:221 E
7kra:A:481 R
7c2l:C:1072 Y
4mwf:D:533 E
Amino Acid Filter 32 Activity
Filter 33
$R_{pred} = 0.01$
Percent Active = 34
Pooling attention 0.45
Pooling feature 2.02
Top activators
6nha:B:29 G
6xob:K:230 H
7l2d:A:1069 P
6gg0:3:64 G
6pzw:C:292 R
6q23:A:292 R
6mej:C:493 P
5i5k:B:1410 G
4uao:A:65 P
2qql:A:369 G
Amino Acid Filter 33 Activity
Filter 34
$R_{pred} = 0.01$
Percent Active = 32
Pooling attention 0.07
Pooling feature 0.74
Top activators
6was:G:155 K
6q23:A:277 E
4uu9:D:32 Y
4zxb:E:187 H
5kqv:E:187 H
4i3r:G:481 S
6mej:C:562 K
7l2d:A:1147 Y
6pzw:C:277 E
6o25:G:112 S
Amino Acid Filter 34 Activity
Filter 35
$R_{pred} = -0.10$
Percent Active = 31
Pooling attention 0.07
Pooling feature 0.03
Top activators
6ln2:A:390 L
3iyw:A:20 W
6nha:A:51 W
3hae:C:5 W
6xob:K:248 R
6wha:B:96 W
6usf:D:317 W
7lxw:A:152 W
7bud:B:20 W
7kra:A:757 W
Amino Acid Filter 35 Activity

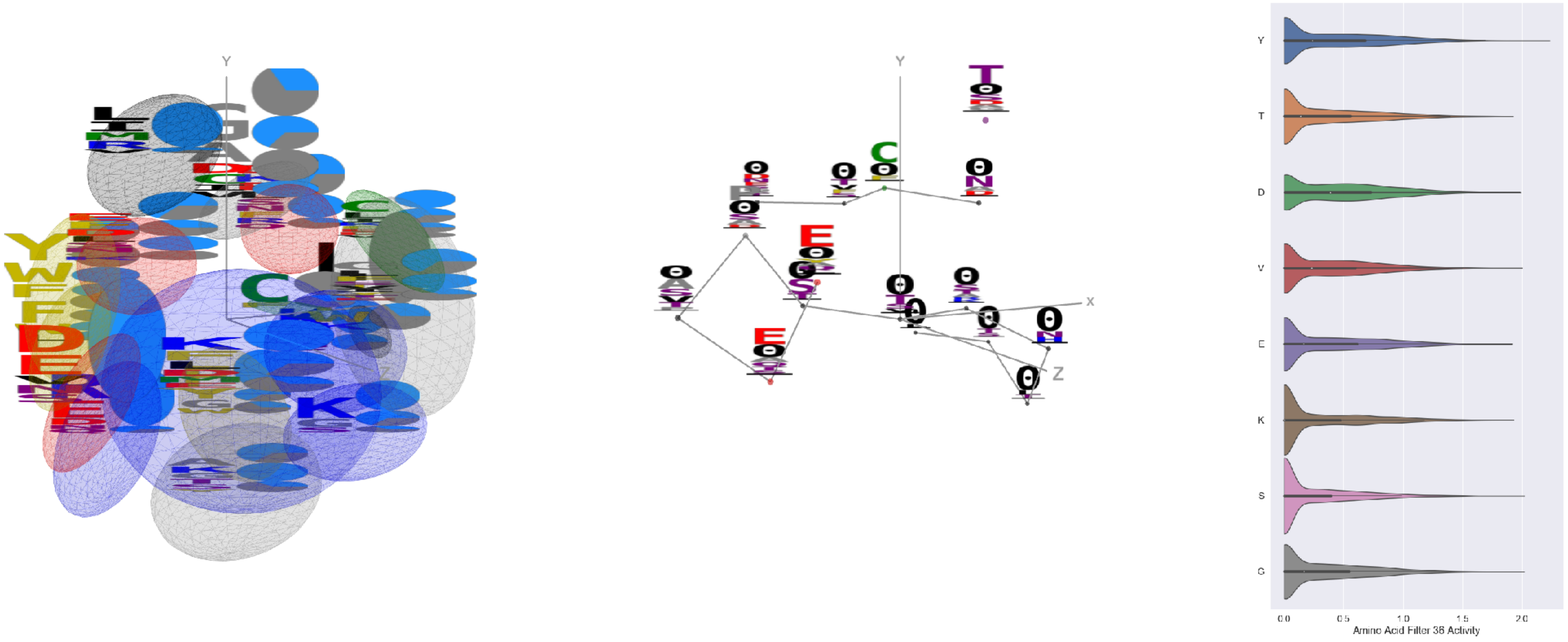


Filter 36
$R_{pred} = 0.01$
Percent Active = 50
Pooling attention 0.20
Pooling feature 0.17
Top activators
5i6x:A:225 Y
4zxb:E:224 A
7l57:B:1060 L
5n0a:A:142 S
6x3x:B:44 G
6rps:B:60 V
7kra:A:67 Y
5tq2:A:373 D
1kb5:B:63 G
4uta:A:229 Y

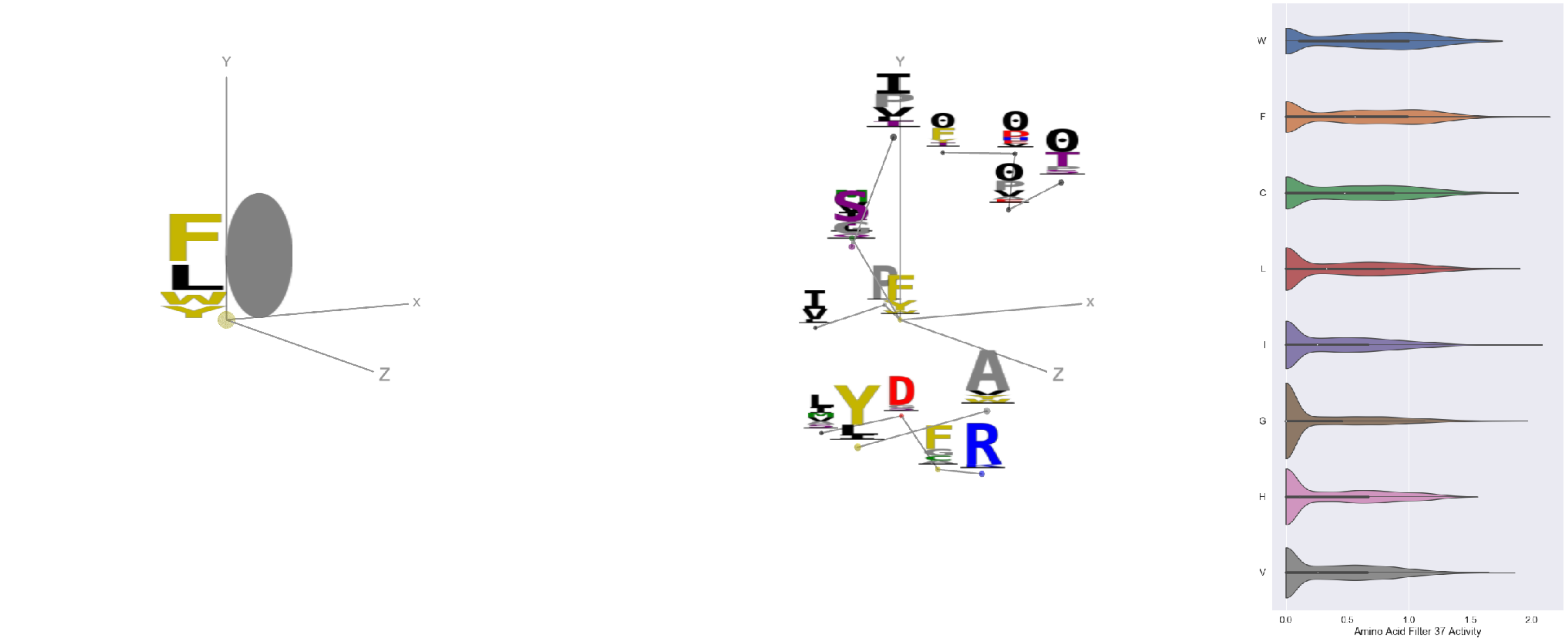


Filter 37
$R_{pred} = -0.31$
Percent Active = 40
Pooling attention 0.12
Pooling feature 0.83
Top activators
7ebz:A:186 F
7l2d:A:1049 I
4dw2:U:103 I
4f3f:C:20 P
6gg0:3:281 G
3w9e:C:179 F
7cws:R:1067 Y
7cwm:A:1067 Y
7ly2:A:1067 Y
7d77:A:216 F

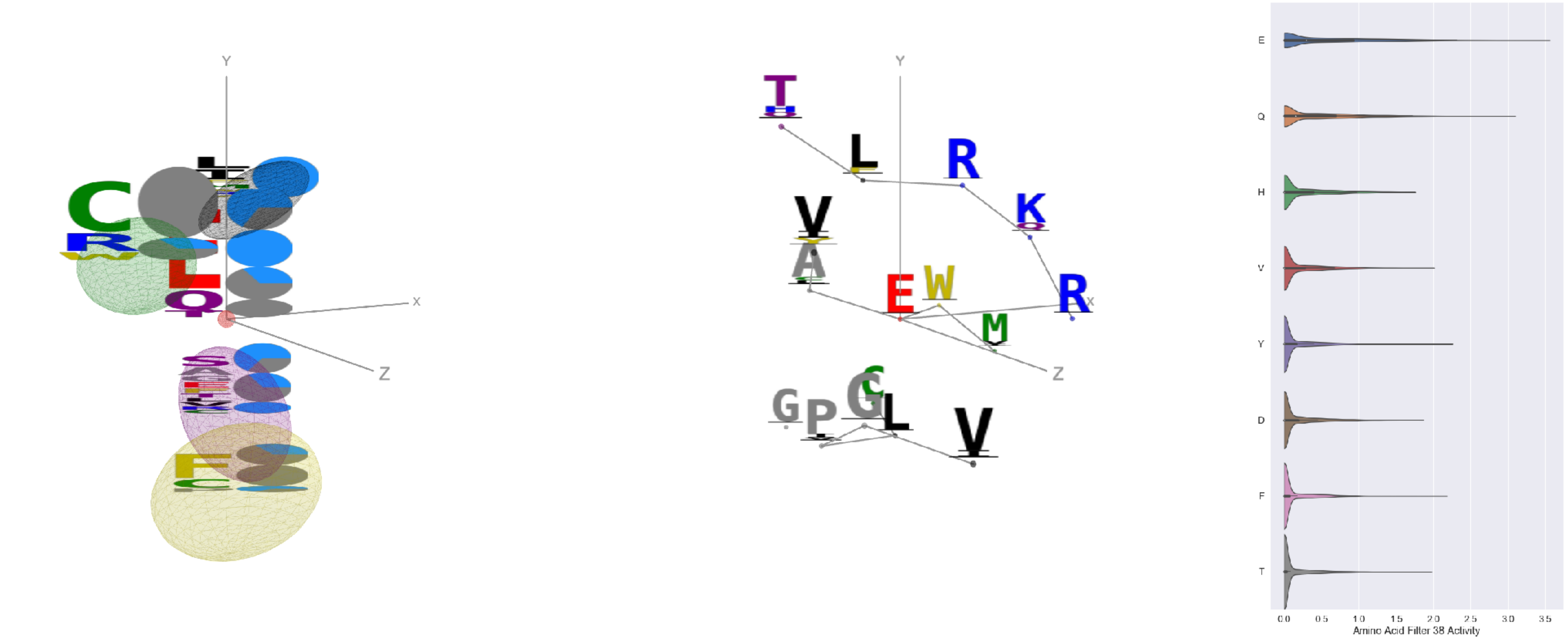


Filter 38
$R_{pred} = -0.01$
Percent Active = 25
Pooling attention 0.05
Pooling feature 0.64
Top activators
6gg0:4:111 E
6gg0:3:111 E
6q23:A:425 E
7kra:A:695 E
6pzw:C:425 E
6bfq:G:93 E
7lxx:A:239 Q
7l2e:A:298 E
6c9u:A:828 E
6rps:B:117 E

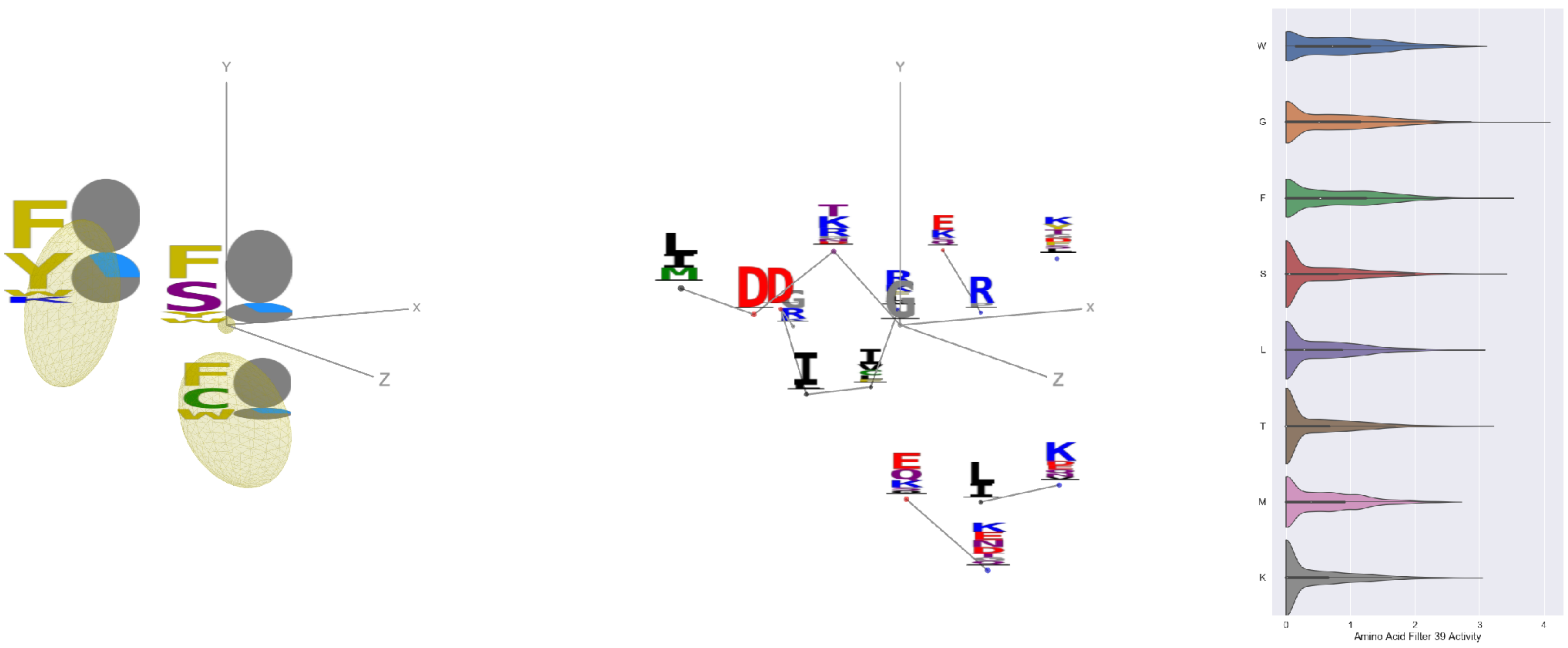


Filter 39
$R_{pred} = -0.06$
Percent Active = 53
Pooling attention 0.27
Pooling feature 0.32
Top activators
5f3j:A:316 G
6yio:B:23 G
4zfg:A:465 Y
5tq0:A:272 D
4uao:A:126 F
6rps:B:49 F
5i5k:B:1223 Y
6wdt:B:117 F
7ebz:C:95 S
7c80:A:259 Y

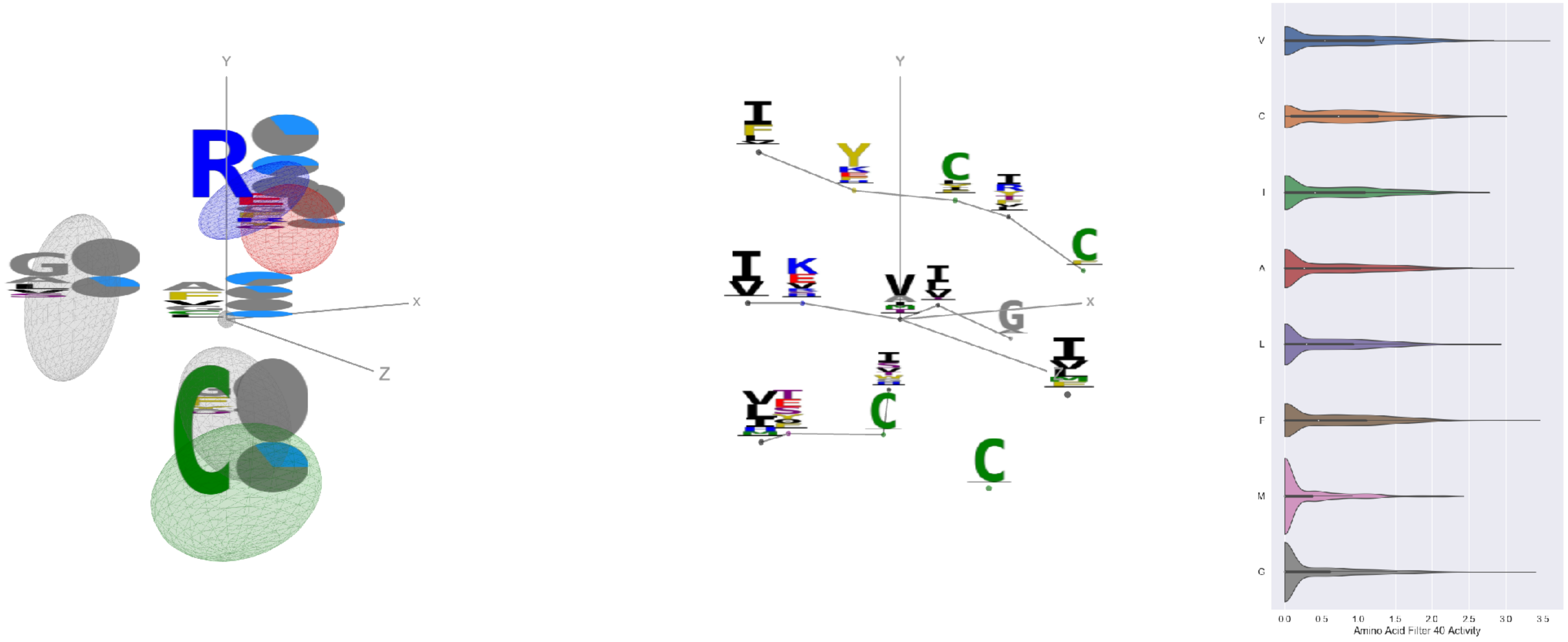


Filter 40
$R_{\mathrm{pred}} = -0.32$
Percent Active = 41
Pooling attention 0.08
Pooling feature 0.23
Top activators
7jum:A:723 V
6c9u:A:270 V
6p91:A:195 F
6mej:C:600 G
6c9u:A:227 A
6p95:C:195 F
7lm9:A:498 V
7ly2:A:512 V
6mej:C:551 G
6q23:A:183 C

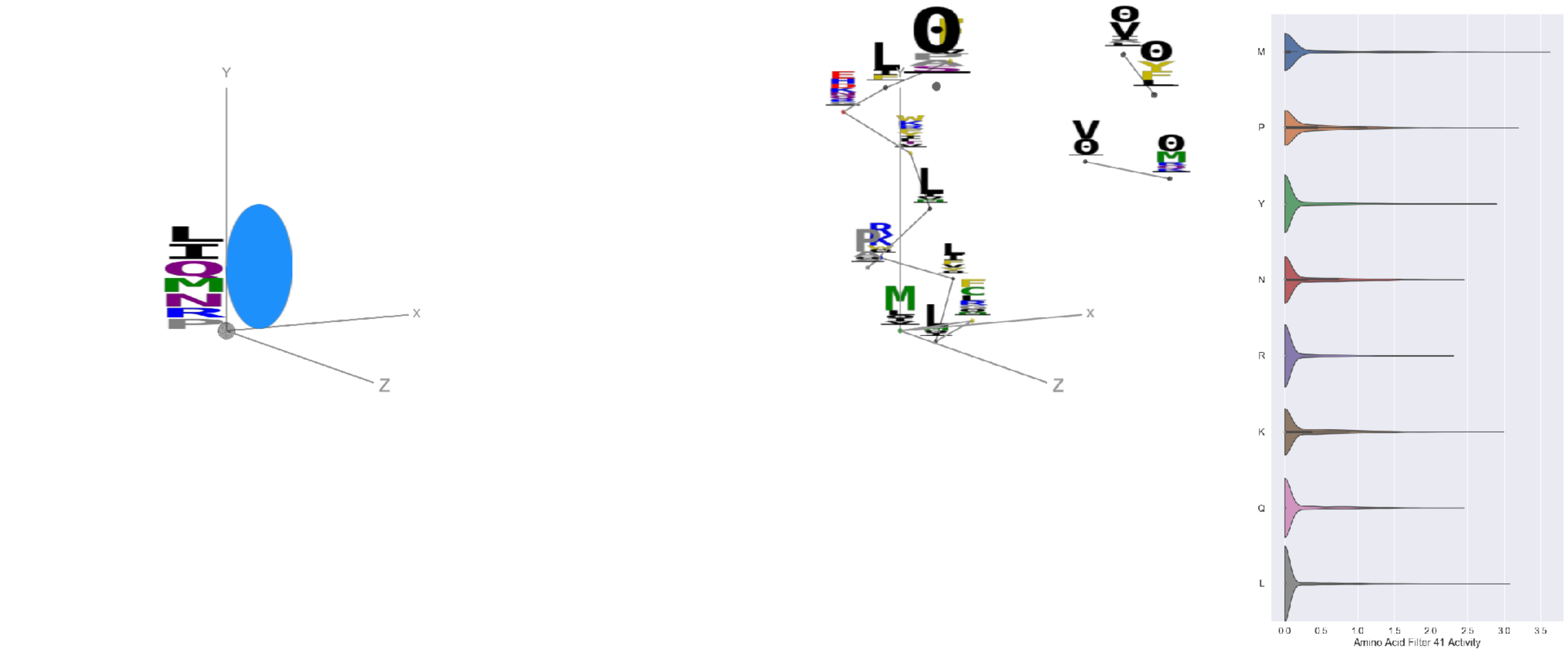


Filter 41
$R_{\mathrm{pred}} = 0.24$
Percent Active = 21
Pooling attention 0.03
Pooling feature 0.25
Top activators
6usf:D:332 M
6p91:a:420 E
6p95:c:420 E
6bfq:G:124 P
7lxx:A:153 M
7lxx:A:24 L
5esv:G:185 P
6wha:B:51 K
5esz:G:185 P
6wo5:E:453 P

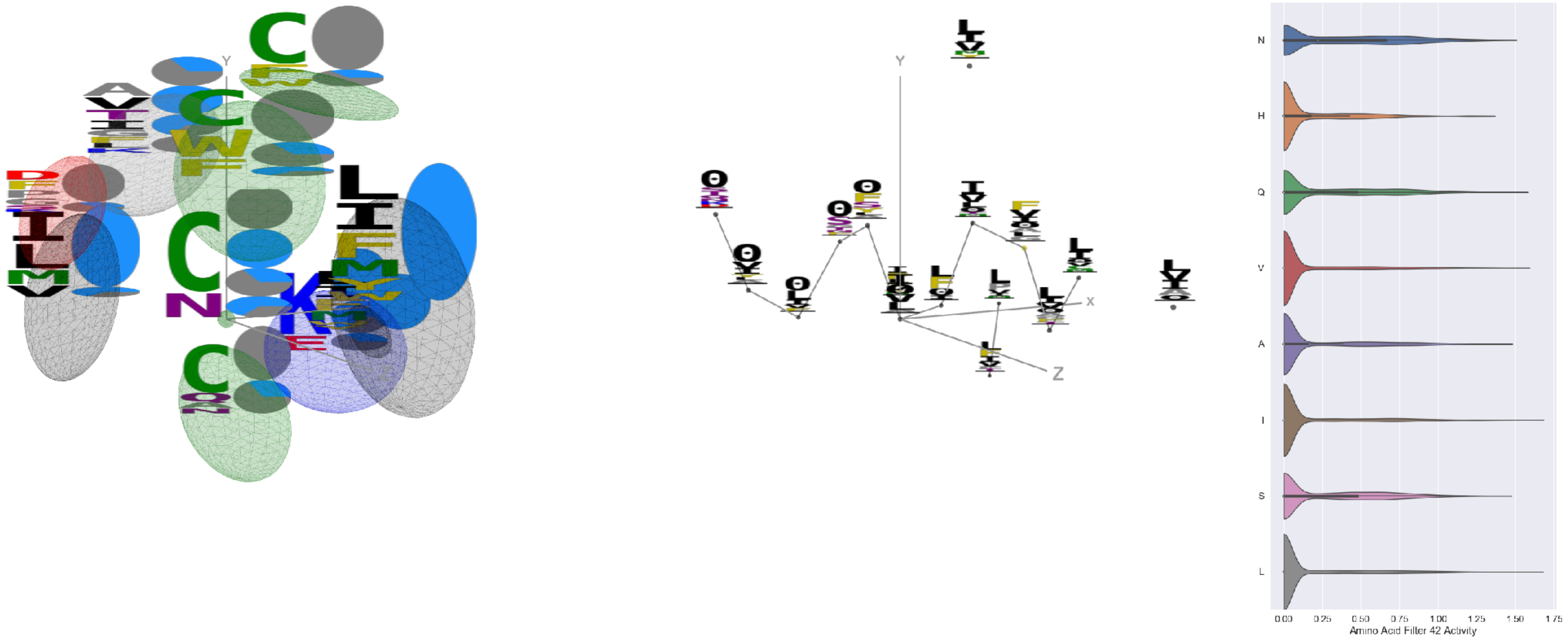


Filter 42
$R_{\mathrm{pred}} = 0.26$
Percent Active = 37
Pooling attention 0.52
Pooling feature 2.41
Top activators
5i6x:A:554 I
3tt3:A:458 L
6m58:A:101 C
5tr1:B:287 L
6x3x:B:328 Y
7bud:B:462 L
6gg0:4:120 V
6oik:A:351 Y
7c80:A:164 Q
7l57:B:321 Q

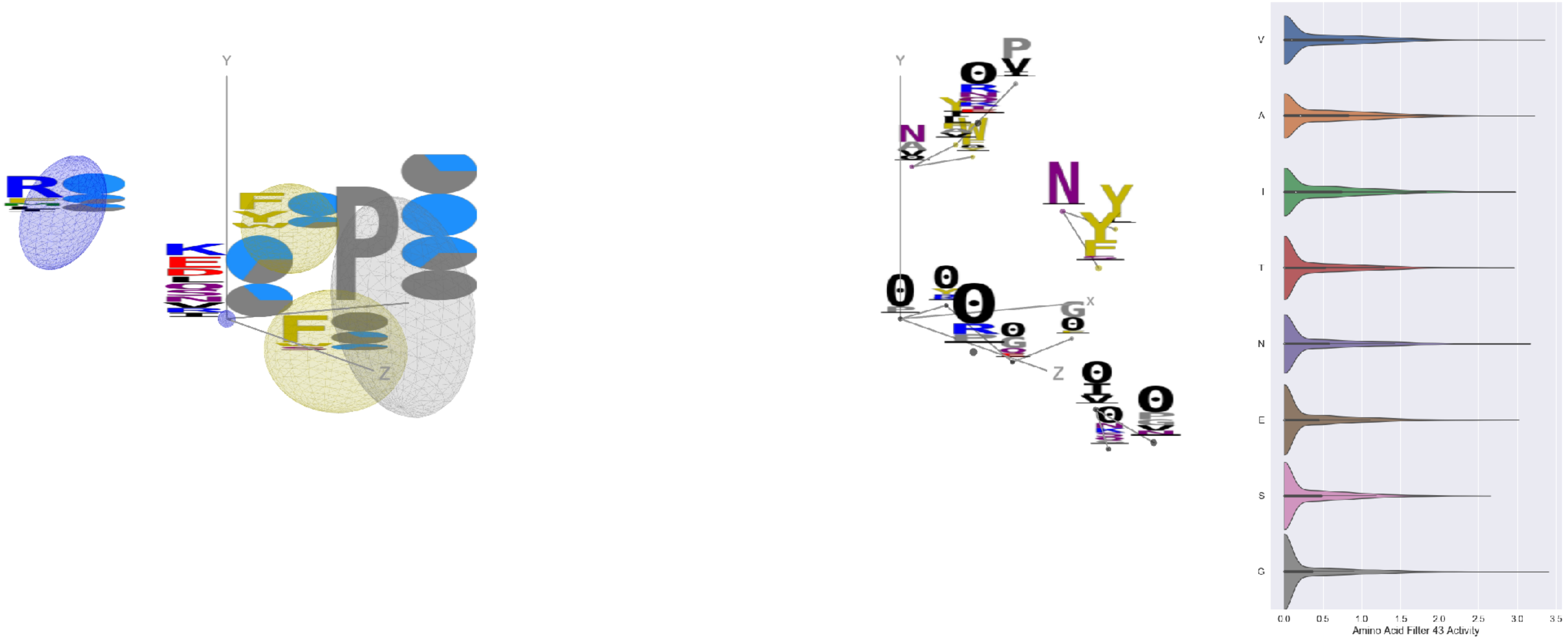


Filter 43
$R_{\mathrm{pred}} = -0.07$
Percent Active = 40
Pooling attention 0.22
Pooling feature 0.82
Top activators
7jv6:E:467 Y
3uze:D:393 K
6ii9:C:317 Y
6ln2:A:283 P
3iyw:A:399 K
7lxx:A:103 G
6o2b:M:12 V
6usf:D:227 P
4i3r:G:266 A
3uze:C:393 K

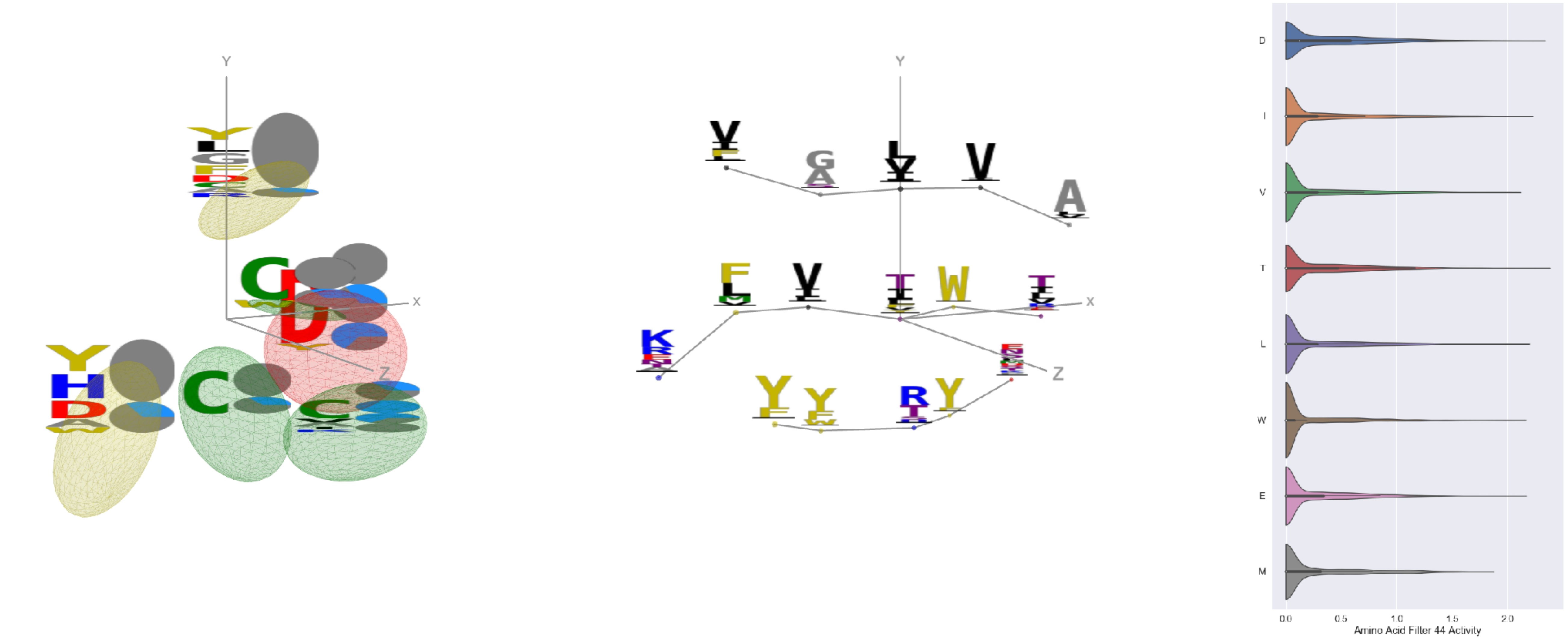


Filter 44
$R_{pred} = 0.08$
Percent Active = 33
Pooling attention 0.30
Pooling feature 0.25
Top activators
6rps:B:210 T
6mei:C:584 D
6n5b:B:151 T
7c2l:C:651 I
6pt0:A:268 L
2qql:A:420 L
5n0a:A:44 E
4fqv:C:69 W
6xob:K:56 D
7lxx:A:141 L

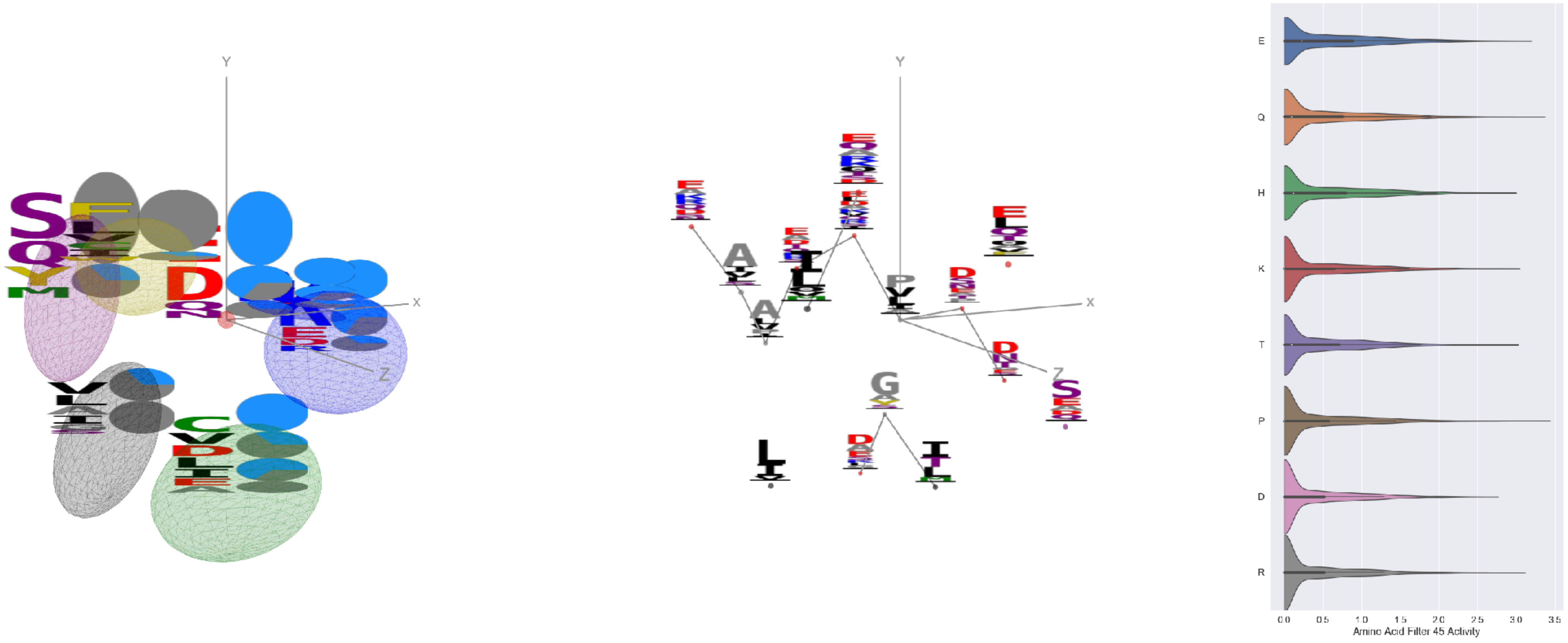


Filter 45
$R_{pred} = 0.06$
Percent Active = 37
Pooling attention 0.47
Pooling feature 1.18
Top activators
2adf:A:1045 P
6wdt:A:273 Y
7lxw:A:115 Q
5tq2:A:298 Y
6fgb:B:77 V
6c9u:A:556 Y
6rps:B:195 E
6weq:B:99 R
5vkd:A:685 E
7l2e:A:100 I

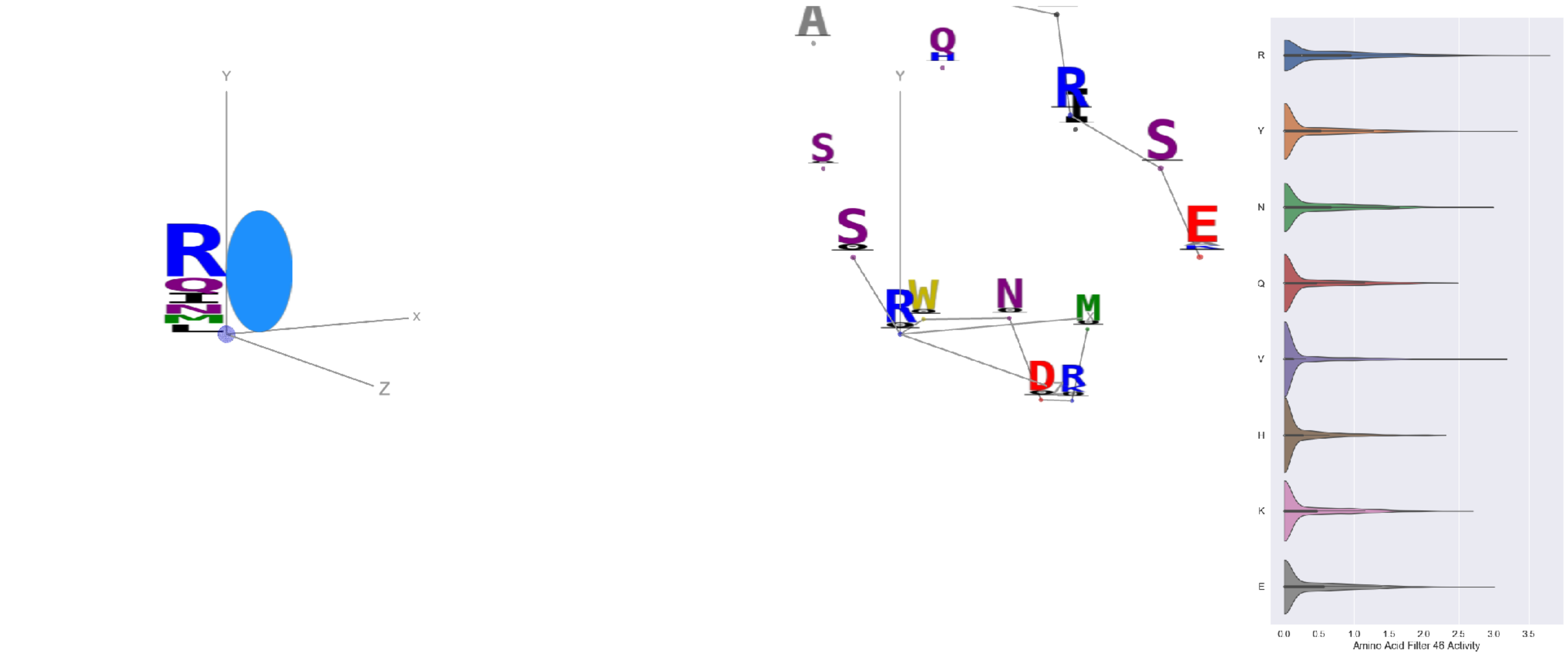


Filter 46
$R_{pred} = 0.26$
Percent Active = 33
Pooling attention 0.25
Pooling feature 0.45
Top activators
4i3r:G:480 R
7bud:B:471 R
4i3r:G:476 R
6ii9:C:317 Y
6r0x:E:83 R
7ebr:C:181 R
6p95:C:256 R
6wo5:E:630 R
7l57:B:382 V
6gg0:4:332 R

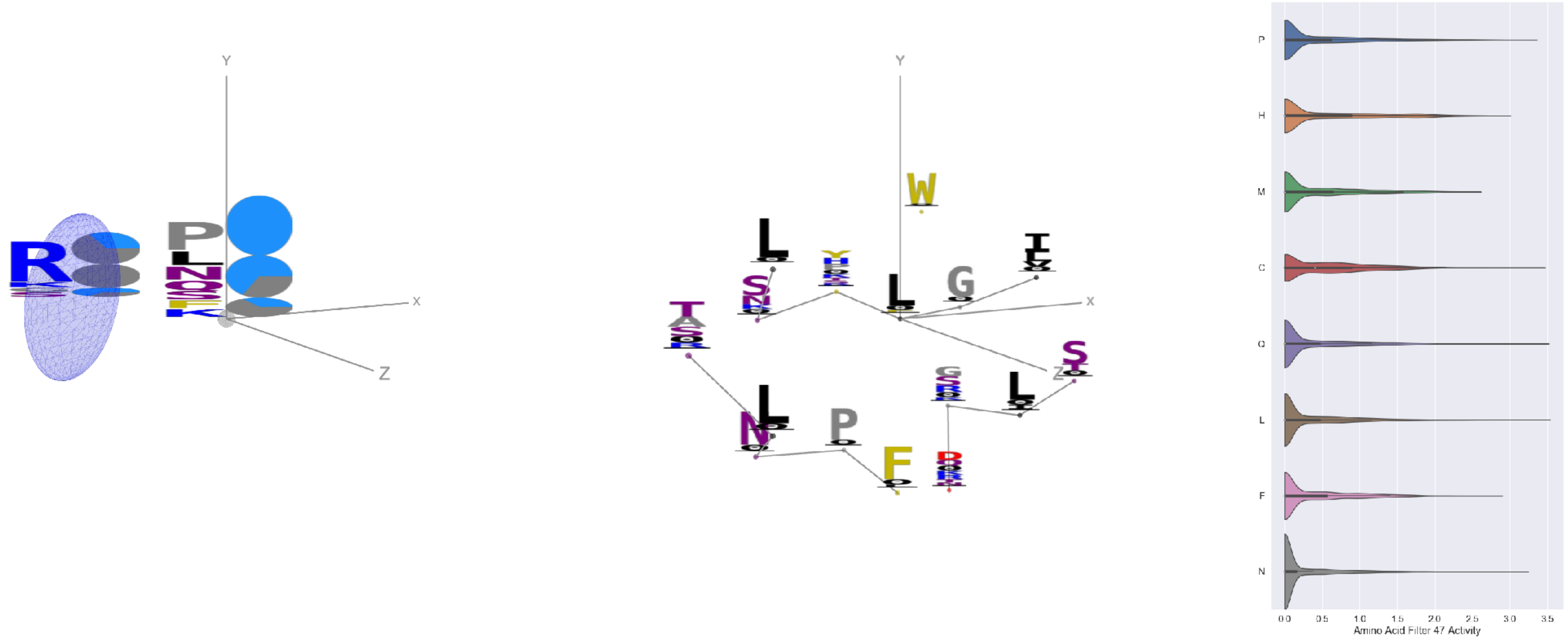


Filter 47
$R_{pred} = -0.17$
Percent Active = 24
Pooling attention 0.08
Pooling feature 0.81
Top activators
4zxb:E:106 L
5f3j:A:388 Q
5kqv:E:106 L
7cah:A:391 C
5f3j:A:240 P
5bk2:B:107 P
7ce2:A:1132 N
4dw2:U:42 C
7k93:C:255 N
6weq:B:255 N

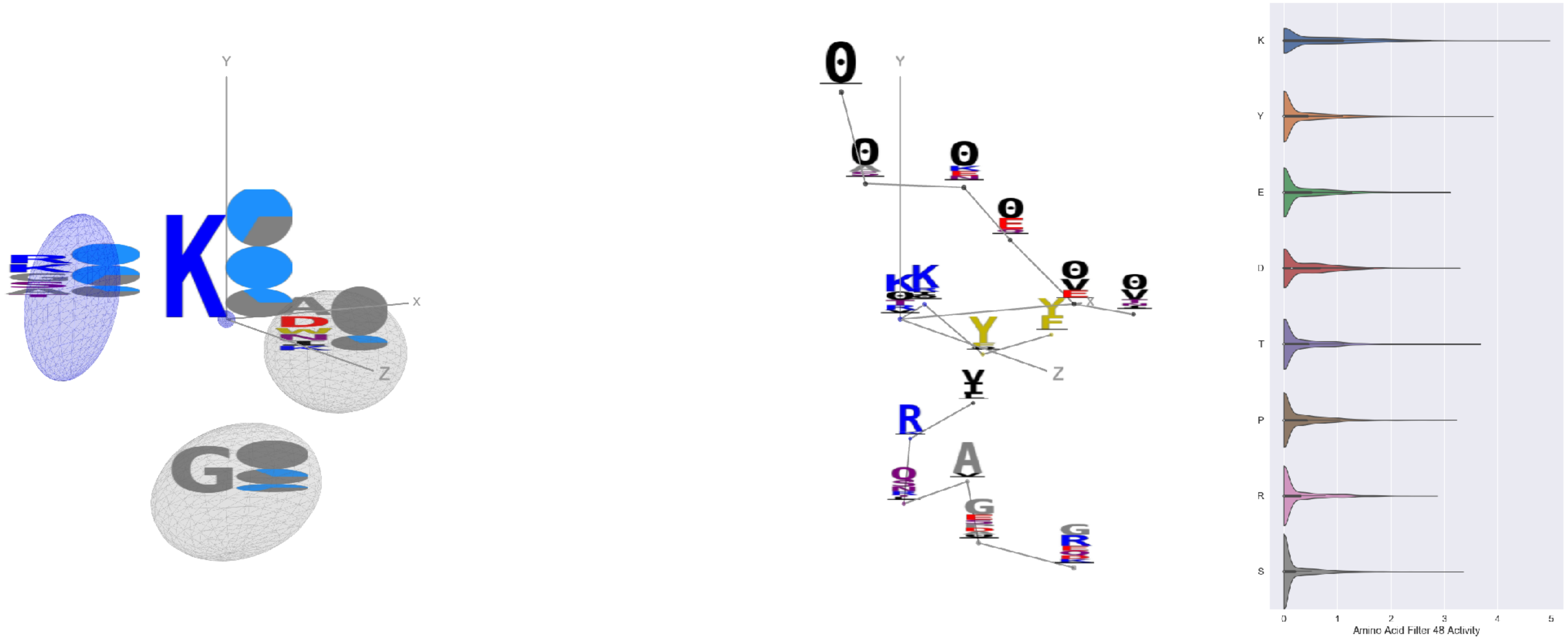

Filter 48
$R_{pred} = 0.09$
Percent Active = 35
Pooling attention 0.27
Pooling feature 0.51
Top activators
3skj:E:176 K
6wdt:A:268 Y
2kh2:A:138 Y
7ebr:A:268 Y
3ks0:A:10 Y
7ebz:A:268 Y
6wdt:A:174 T
7ce2:A:1213 K
1dee:H:2856 Y
1w72:D:176 K

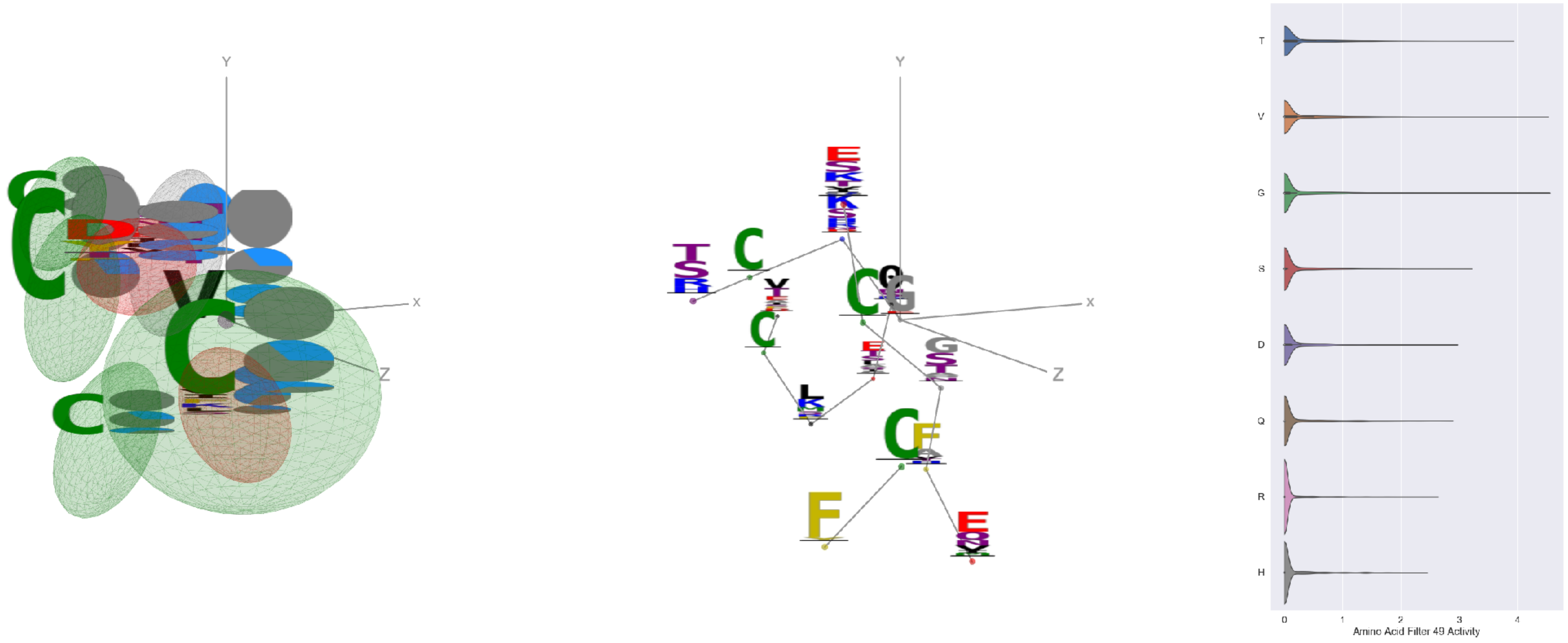

Filter 49
$R_{pred} = -0.03$
Percent Active = 17
Pooling attention 0.28
Pooling feature 0.45
Top activators
5y11:C:278 G
7lxw:A:159 V
7lxx:A:159 V
6pzw:C:235 G
6ii9:C:270 T
6m58:A:164 A
1nca:N:235 G
6mej:C:563 T
6c9u:A:340 T
6m58:A:473 V

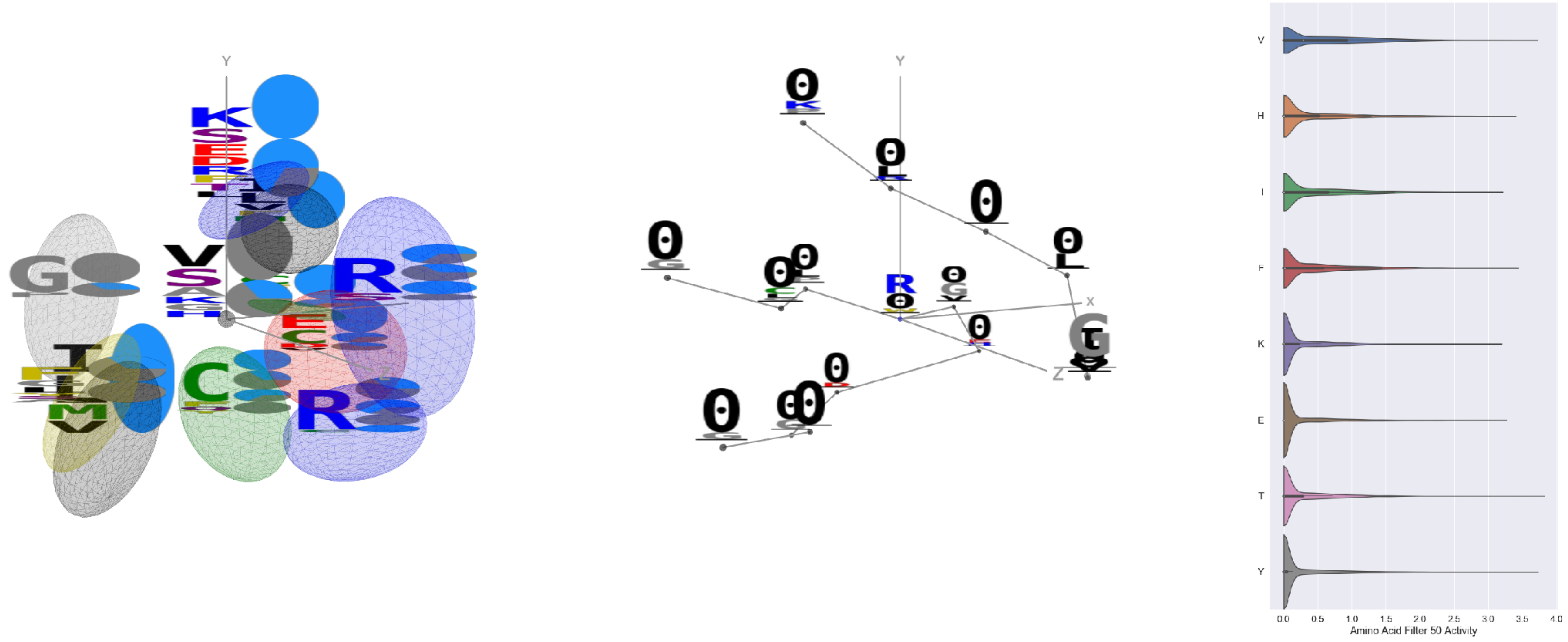

Filter 50
$R_{pred} = -0.24$
Percent Active = 32
Pooling attention 0.13
Pooling feature 0.96
Top activators
5esz:G:155 R
3iyw:A:32 T
6wpt:C:128 Y
7l57:B:433 V
6xkq:A:433 V
6nb7:B:314 Y
7jv6:E:433 V
7l57:B:327 Y
6nb7:B:131 Y
7lm9:A:420 V

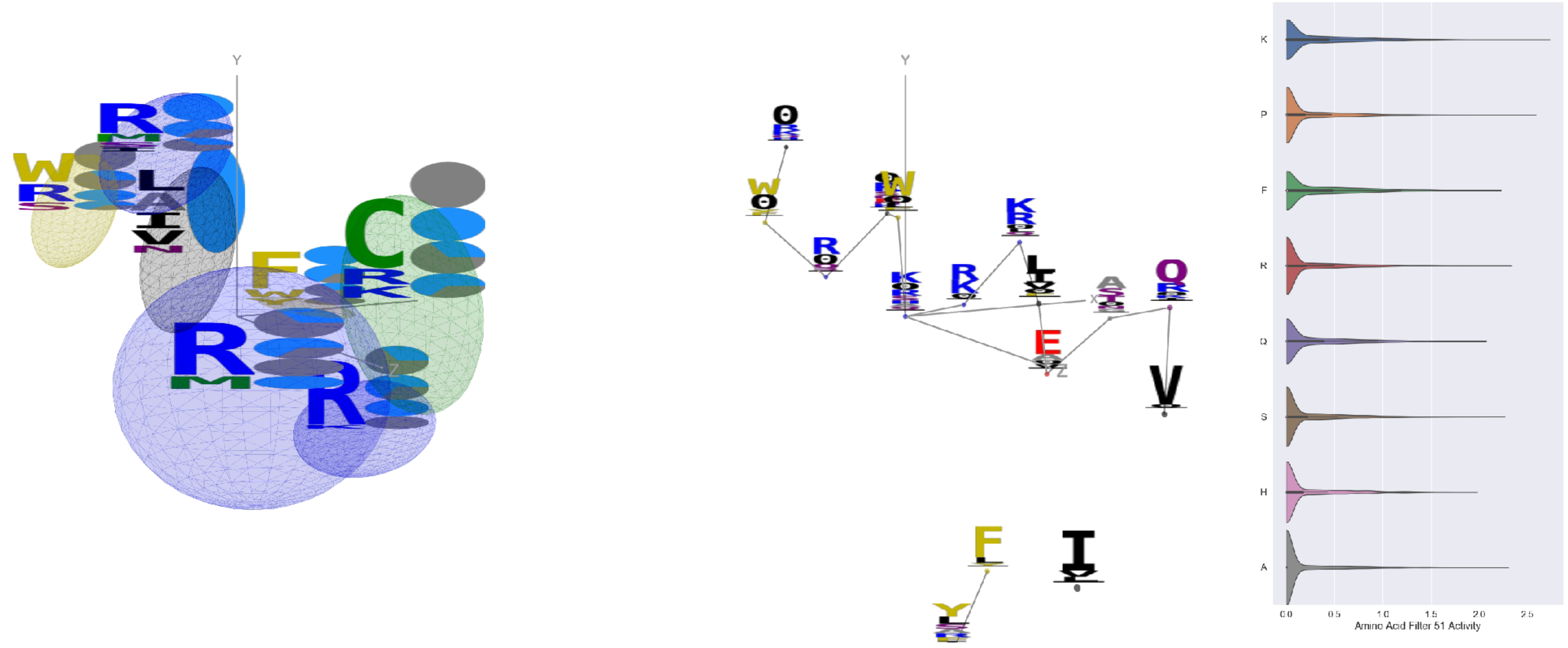

Filter 51
$R_{pred} = -0.05$
Percent Active = 27
Pooling attention 0.50
Pooling feature 0.34
Top activators
6ln2:A:416 K
6mjz:C:185 P
4okv:F:265 K
6ln2:A:312 P
5bo1:A:305 P
5i6x:A:555 Y
7kra:A:746 P
6nb7:B:260 R
4xmn:E:984 Y
7kra:A:755 A

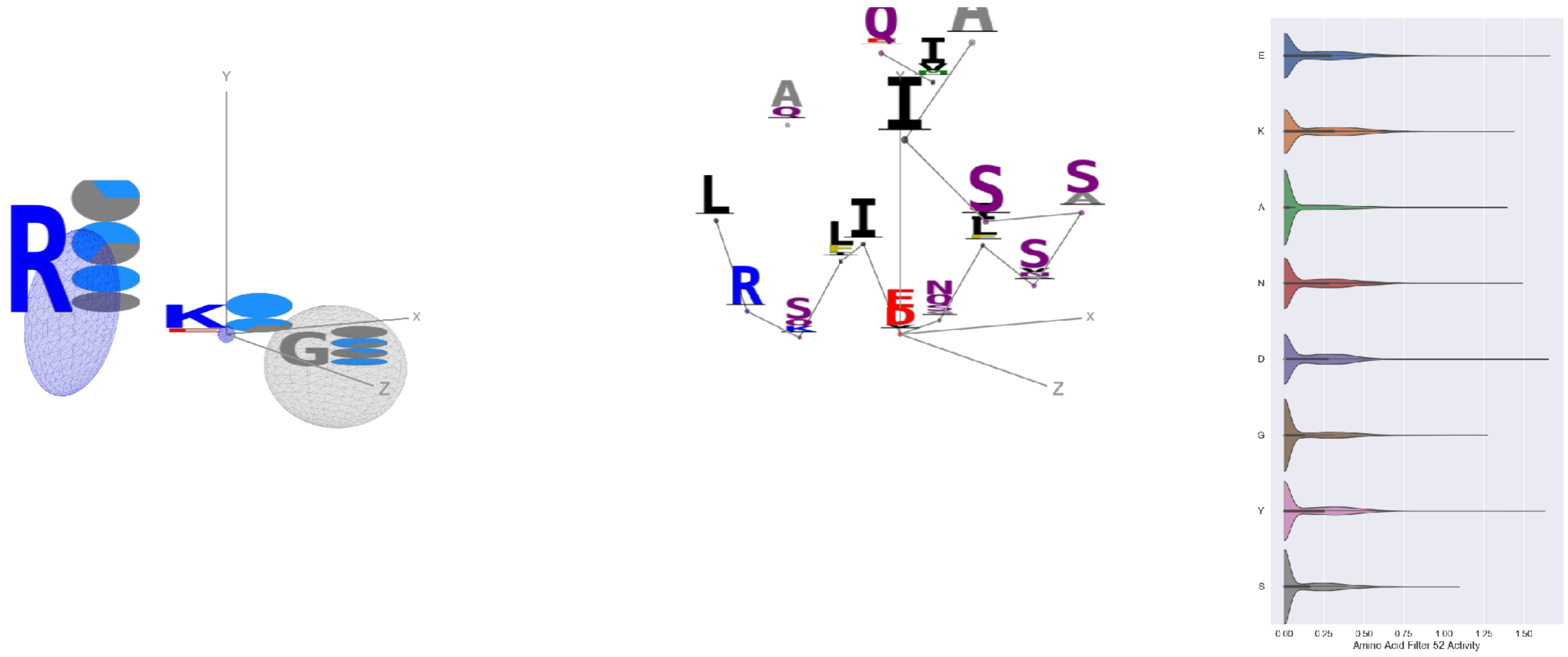


Filter 52
$R_{pred} = 0.21$
Percent Active = 31
Pooling attention 0.13
Pooling feature 1.59
Top activators
7jw0:E:979 E
7l57:B:979 D
7l57:B:811 Y
7lxx:A:99 N
6wo5:E:628 K
6mej:C:628 K
7kra:A:251 A
7l57:B:991 A
6nb7:B:973 A
7lcn:A:991 A

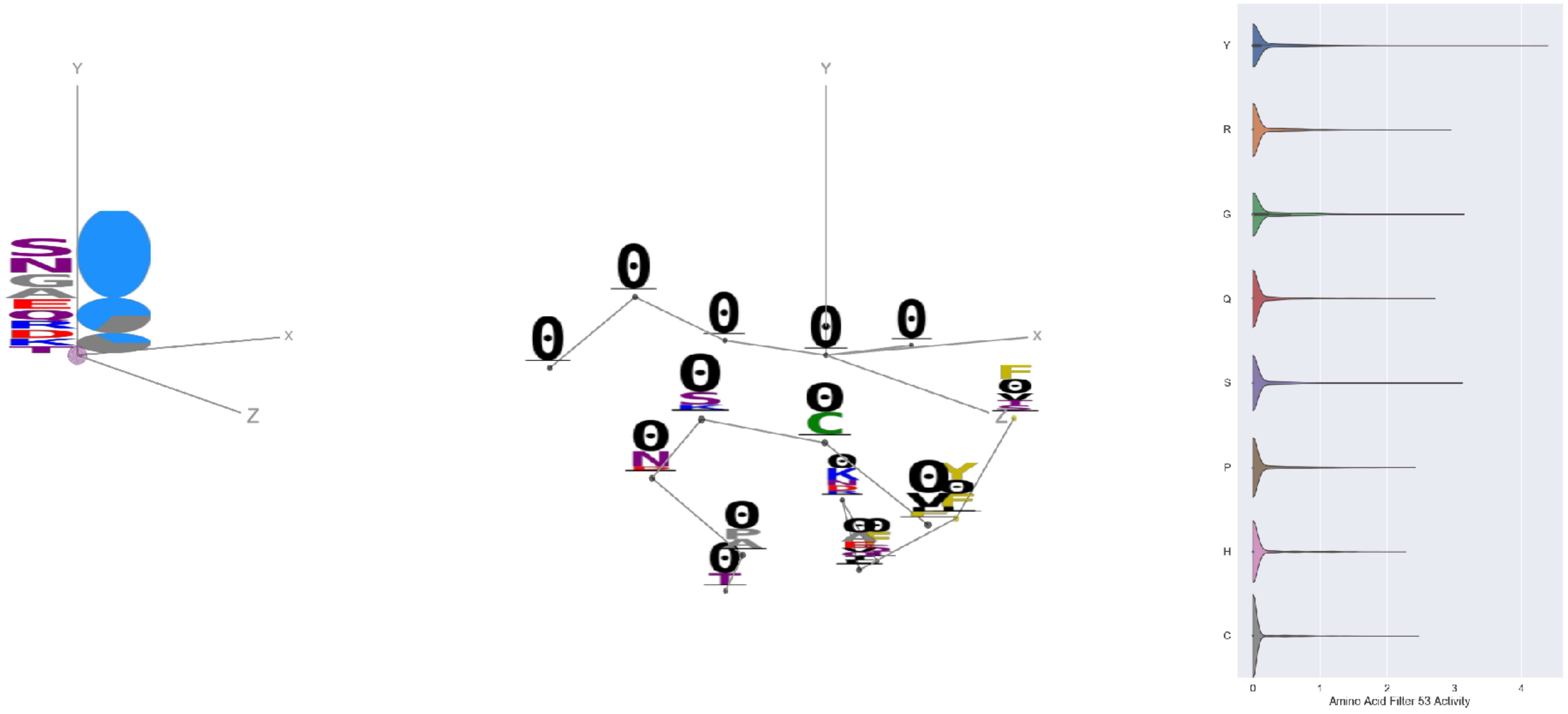


Filter 53
$R_{pred} = 0.07$
Percent Active = 19
Pooling attention 0.58
Pooling feature 0.11
Top activators
7ly2:A:15 Y
7jw0:E:15 Y
7ly0:A:15 Y
7l2e:A:15 Y
4xmn:E:1036 Y
6nb6:A:19 Y
6m58:A:106 K
6m58:A:582 Y
7lcn:A:153 Y
7l57:B:210 Y

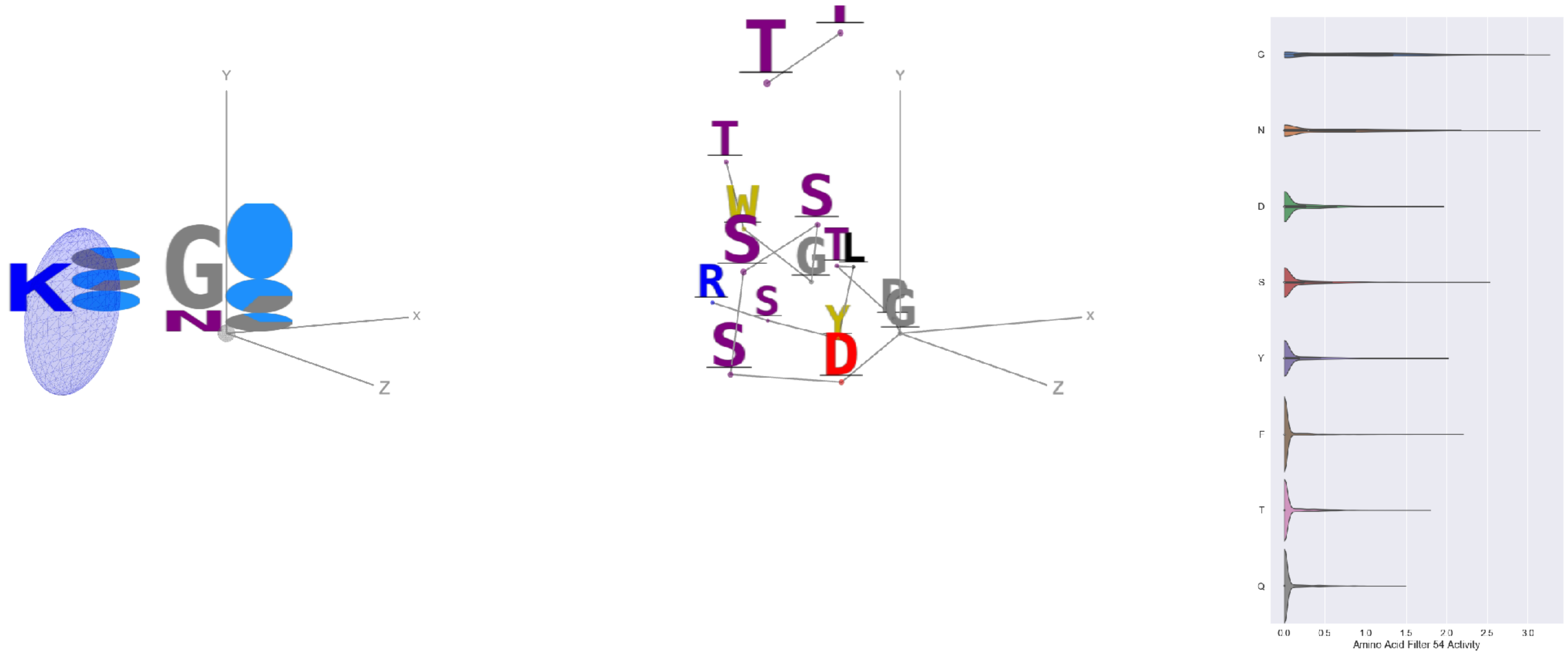


Filter 54
$R_{pred} = 0.10$
Percent Active = 25
Pooling attention 0.04
Pooling feature 0.38
Top activators
7lxx:A:252 G
7lxw:A:148 N
6otc:A:259 G
4i3r:G:380 G
7bud:B:395 G
6pt0:A:40 G
6o2b:M:106 G
7ly2:A:1046 G
7jv2:A:446 G
7cwm:A:1046 G

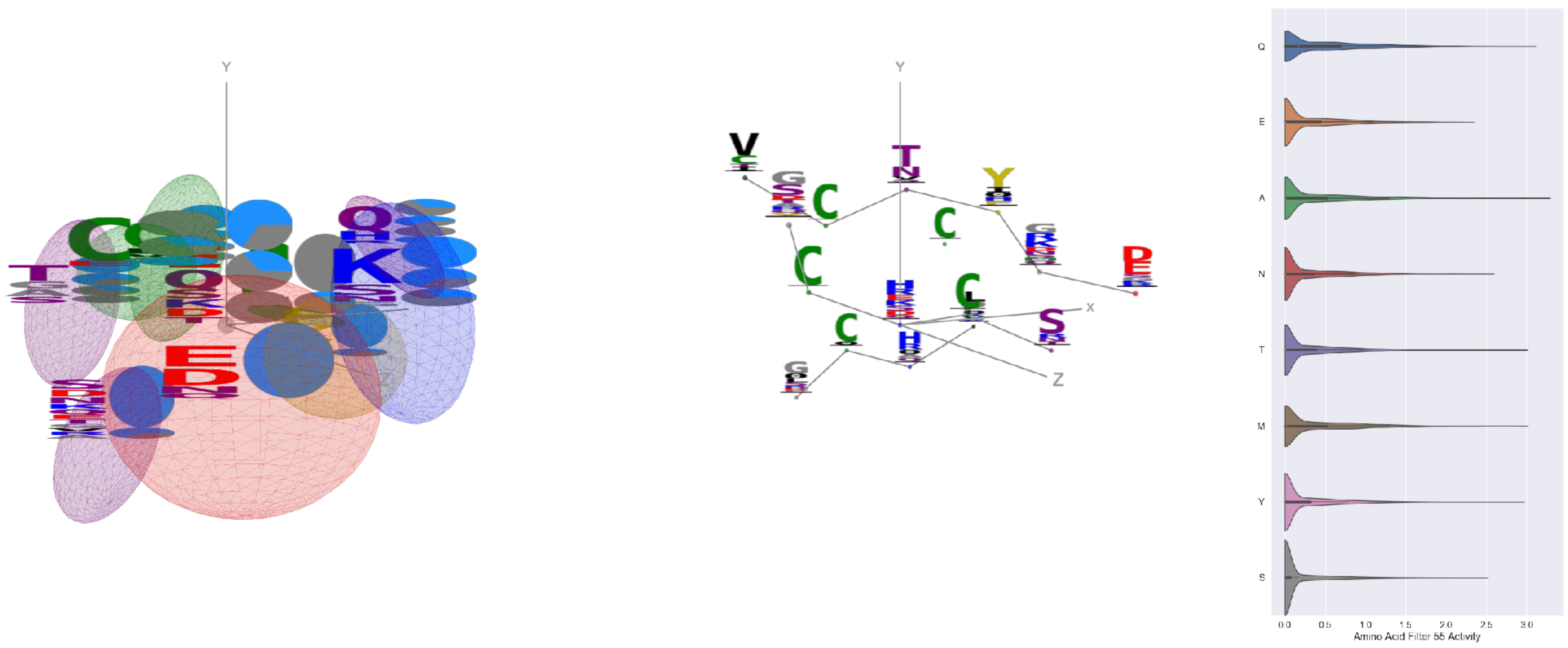


Filter 55
$R_{pred} = -0.04$
Percent Active = 29
Pooling attention 0.11
Pooling feature 0.38
Top activators
1qfw:B:89 H
1xct:M:30 A
7l57:B:761 D
7l57:B:949 Q
4xmn:E:981 M
7jw0:E:1076 T
7jw0:E:887 T
6nb6:C:1126 Y
7jw0:E:761 D
5esv:G:266 Q

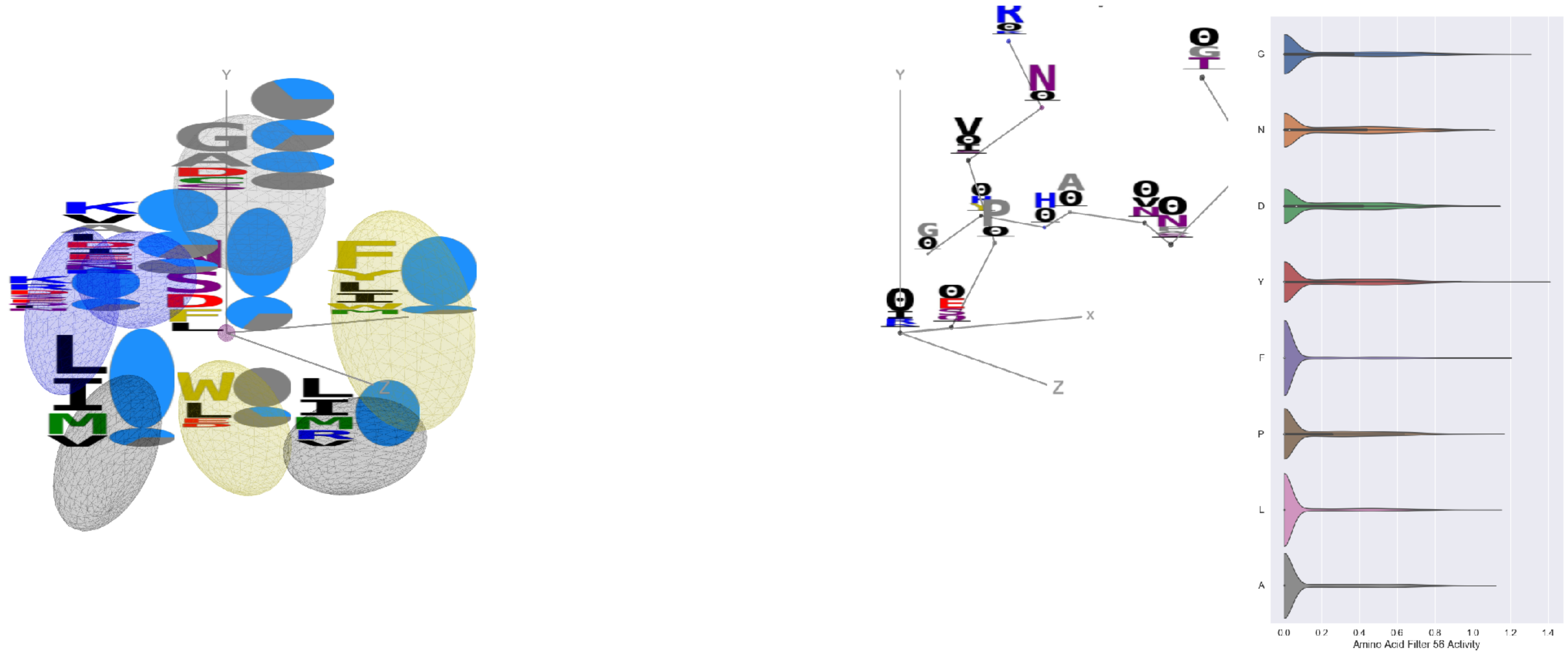

Filter 56
$R_{pred} = 0.24$
Percent Active = 34
Pooling attention 0.03
Pooling feature 1.89
Top activators
6ii9:C:317 Y
6usf:D:467 Y
6nb6:C:168 Y
6oij:A:357 Y
6o25:G:42 G
5d8j:A:78 G
6ulf:K:42 G
7lcn:A:757 G
6nb6:C:1127 Y
6nb7:B:85 Y

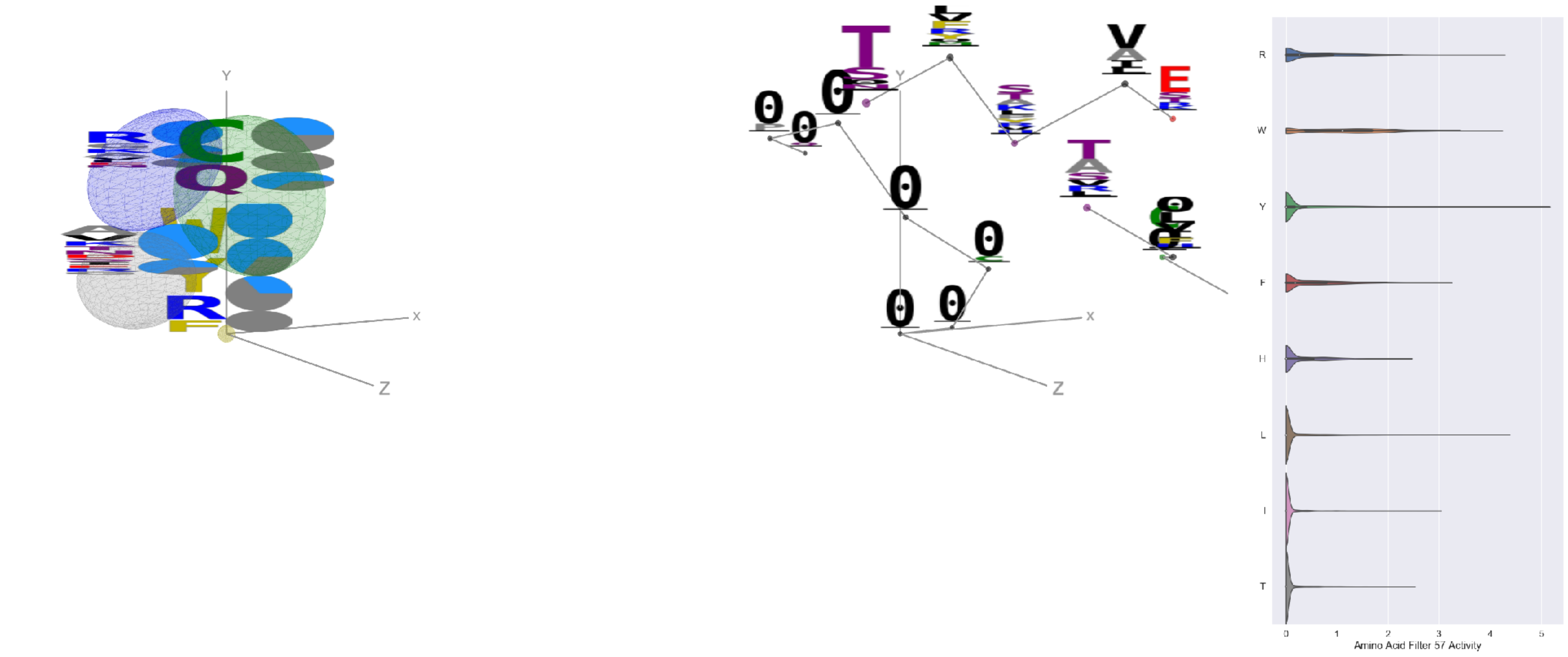

Filter 57
$R_{pred} = 0.05$
Percent Active = 20
Pooling attention 0.11
Pooling feature 1.24
Top activators
1qfw:B:112 Y
4kxz:E:52 L
7bud:B:99 R
3iyw:A:20 W
7cws:R:995 R
7cwm:A:995 R
7lxw:A:152 W
4uta:A:99 R
7c2l:C:995 R
6wpt:C:995 R

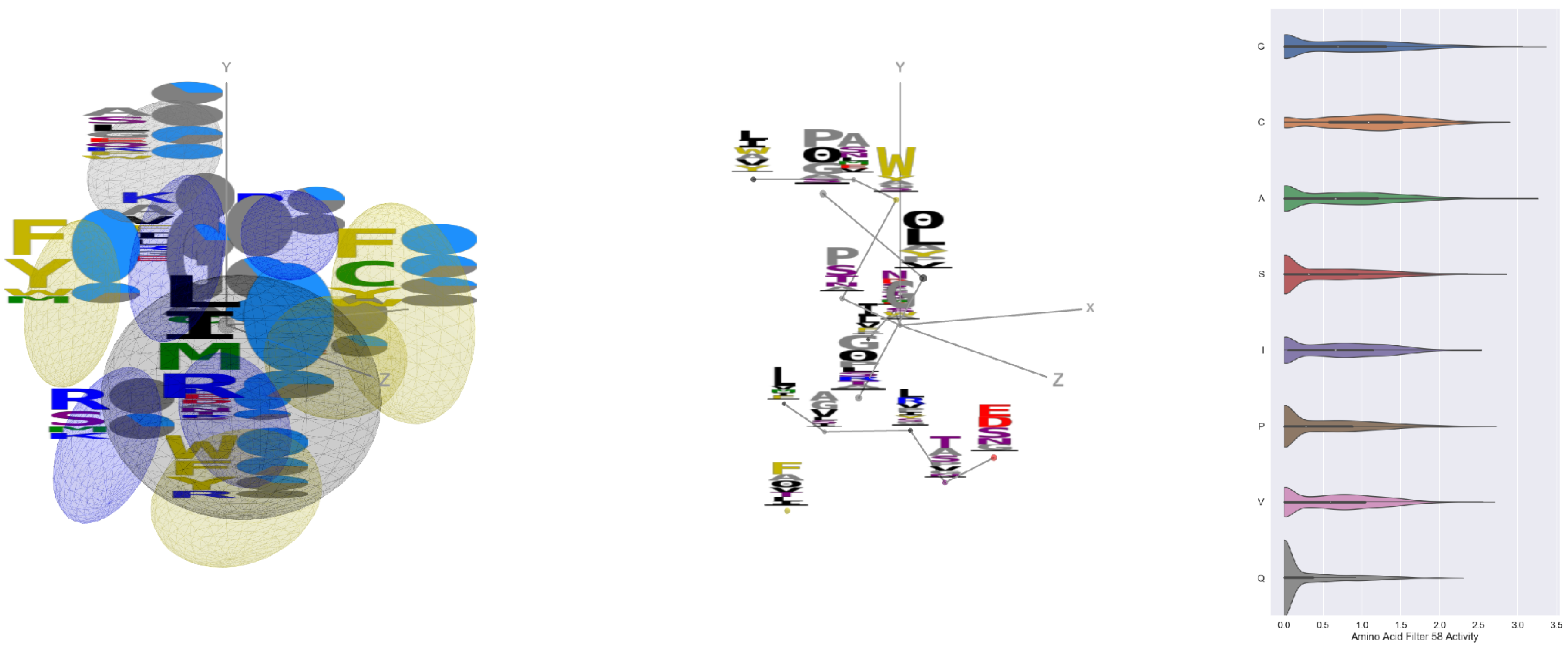

Filter 58
$R_{pred} = -0.25$
Percent Active = 51
Pooling attention 0.14
Pooling feature 0.43
Top activators
5bk2:B:228 G
7kq7:B:184 R
6i8s:C:239 L
7ly2:A:880 A
7nd9:B:880 A
7cwm:A:880 A
7jw0:E:880 A
7cws:R:880 A
6wpt:C:880 A
6q23:A:408 G

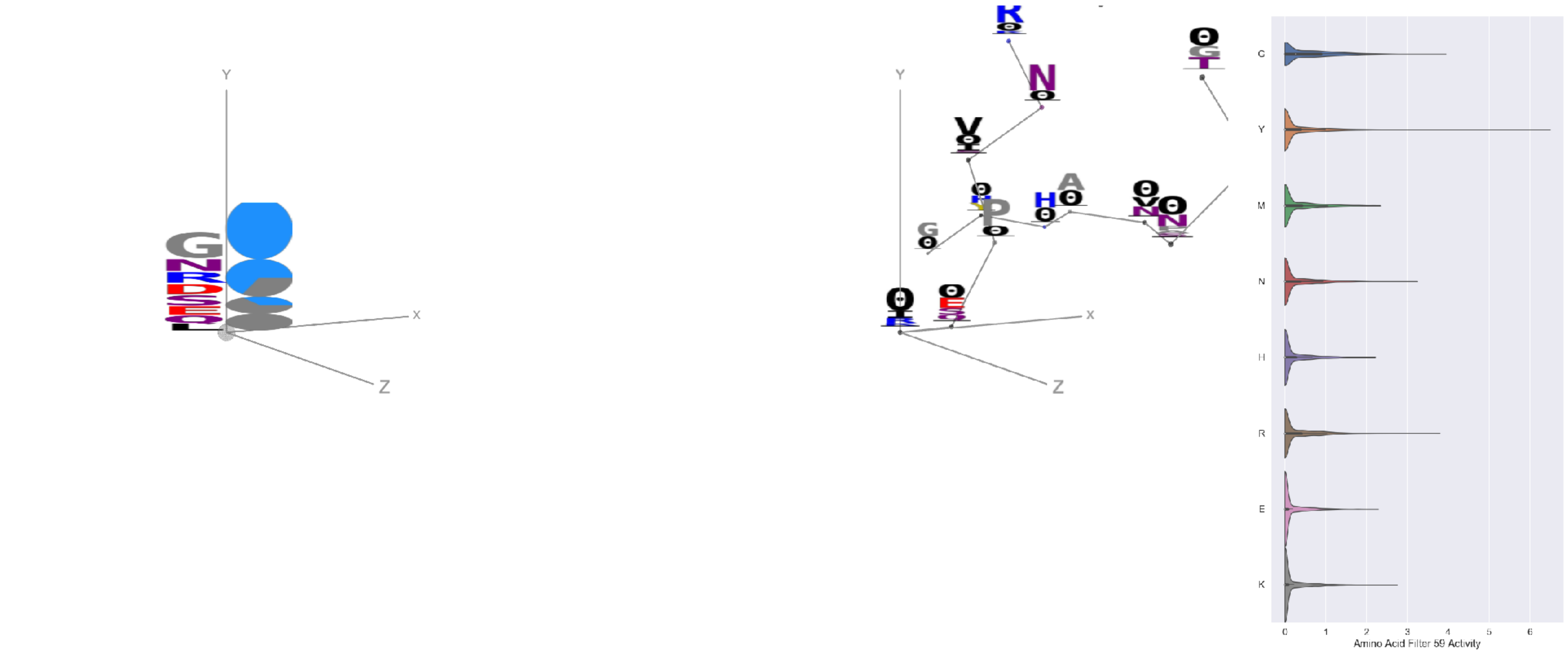

Filter 59
$R_{pred} = -0.04$
Percent Active = 26
Pooling attention 0.02
Pooling feature 0.50
Top activators
6ii9:C:317 Y
6wtv:D:470 Y
4uta:A:395 Y
6fgb:A:268 Y
6nb6:A:21 Y
5tq0:A:53 G
6nb7:B:21 Y
2nr6:B:327 Y
5tq0:A:414 Y
7ly0:A:21 Y

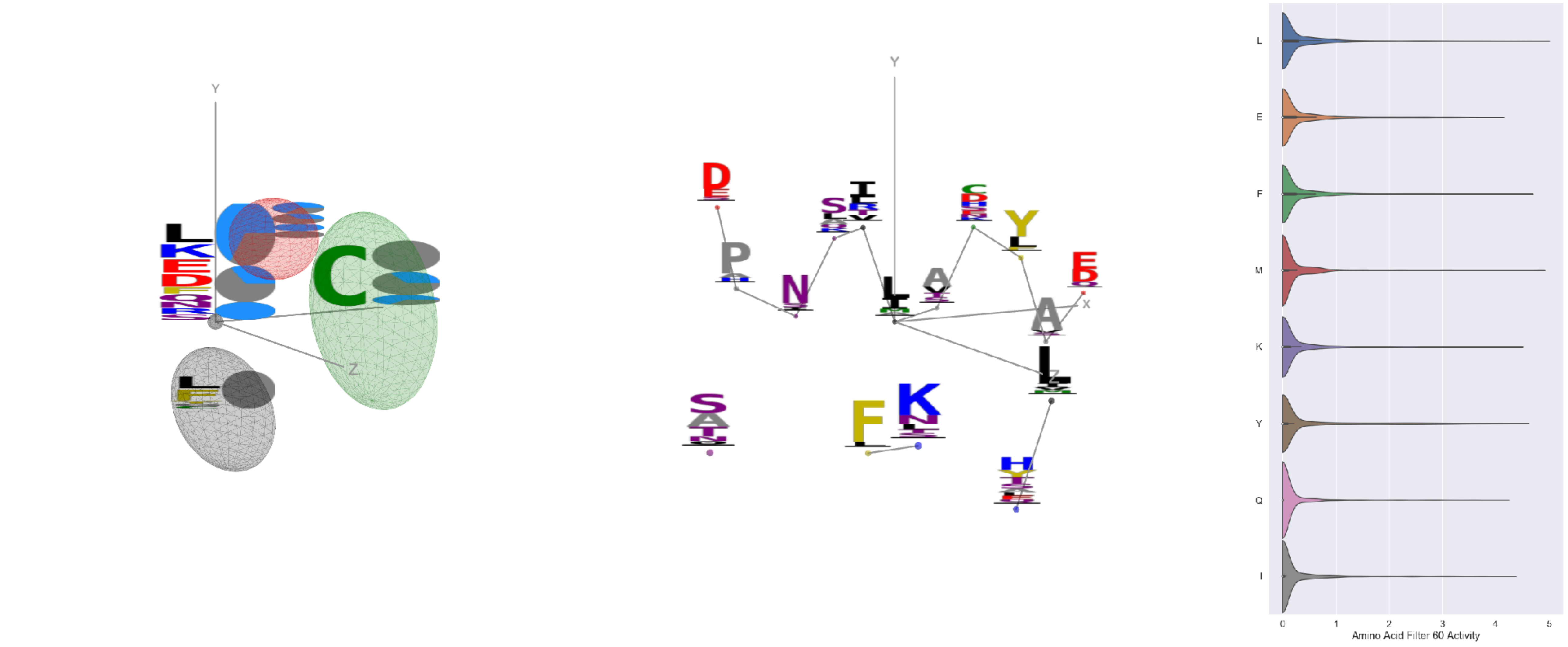

Filter 60
$R_{\text{pred}} = -0.20$
Percent Active = 30
Pooling attention 0.11
Pooling feature 0.14
Top activators
4xmn:E:889 L
4xmn:E:981 M
6nb7:B:263 L
6nb7:B:377 M
4xmn:E:777 F
6nb6:A:263 L
4xmn:E:969 Y
4xmn:E:726 R
4xmn:E:867 N
6nb6:C:343 K

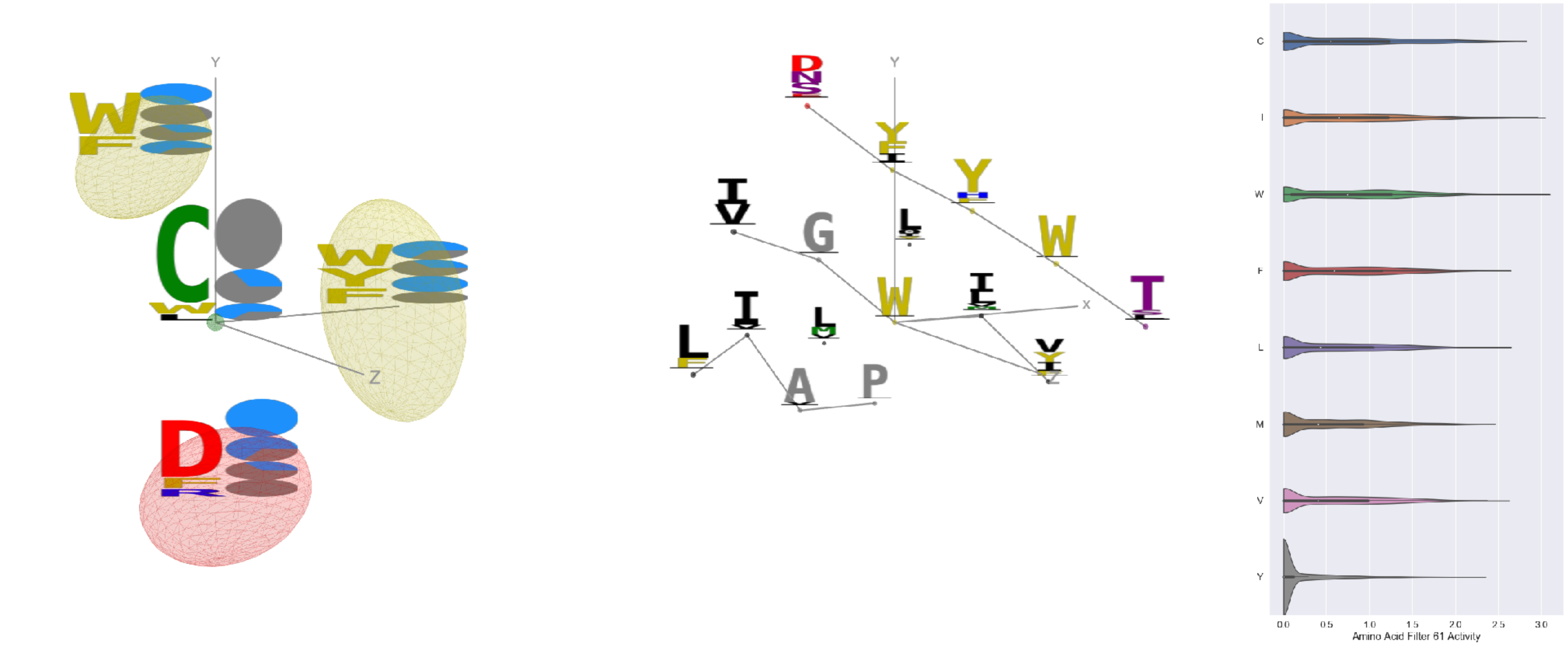

Filter 61
$R_{\text{pred}} = -0.20$
Percent Active = 38
Pooling attention 0.47
Pooling feature 0.95
Top activators
6ii9:C:171 W
6ii9:C:243 I
6ur5:C:180 W
4fqv:C:252 I
6n5b:B:176 W
4i3r:G:385 C
4fqv:C:180 W
7lxw:A:235 I
7bud:B:285 C
6mlm:A:196 W

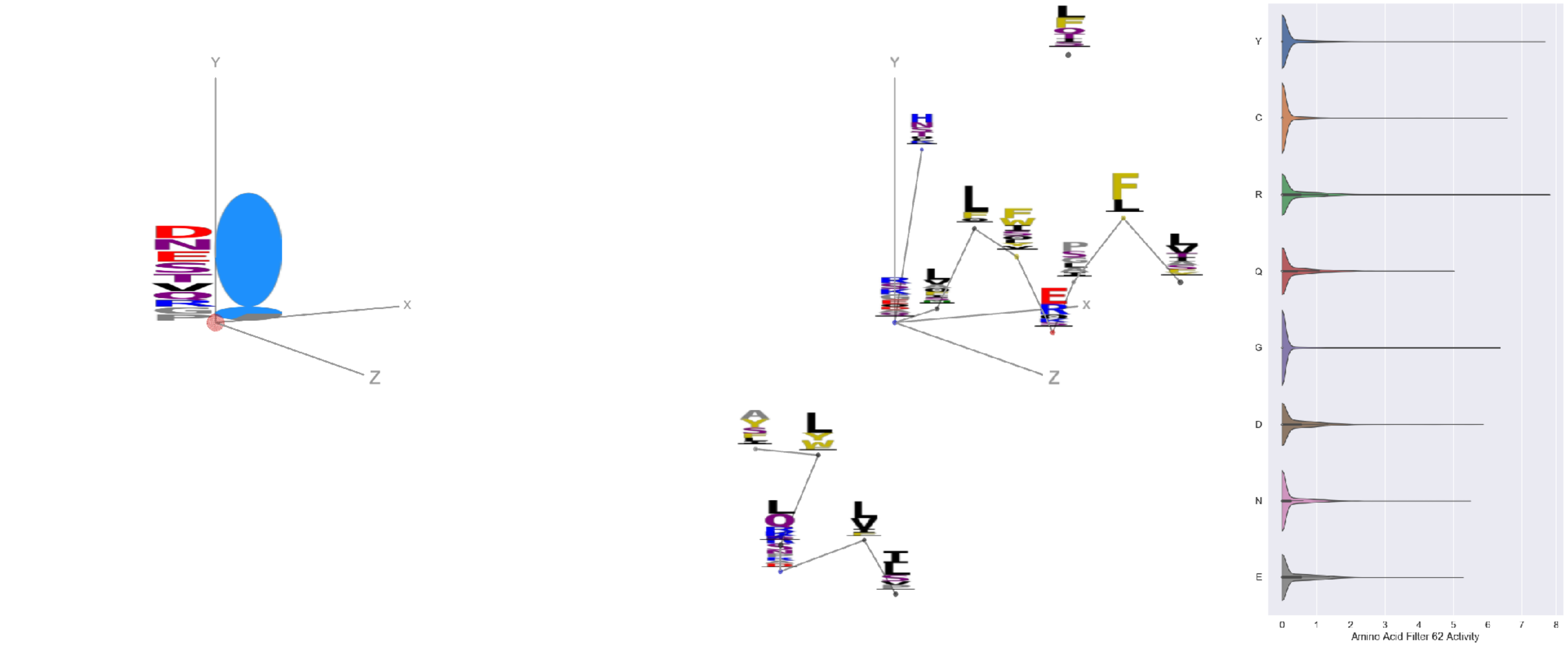

Filter 62
$R_{\text{pred}} = 0.10$
Percent Active = 17
Pooling attention 0.20
Pooling feature 0.87
Top activators
4xmn:E:726 R
6ii9:C:317 Y
6m58:A:582 Y
7jw0:E:826 V
7ly2:A:20 Y
7jw0:E:1147 Y
5otj:C:71 Y
4cmh:A:287 Y
7ly2:A:1145 Y
7l2d:A:1147 Y

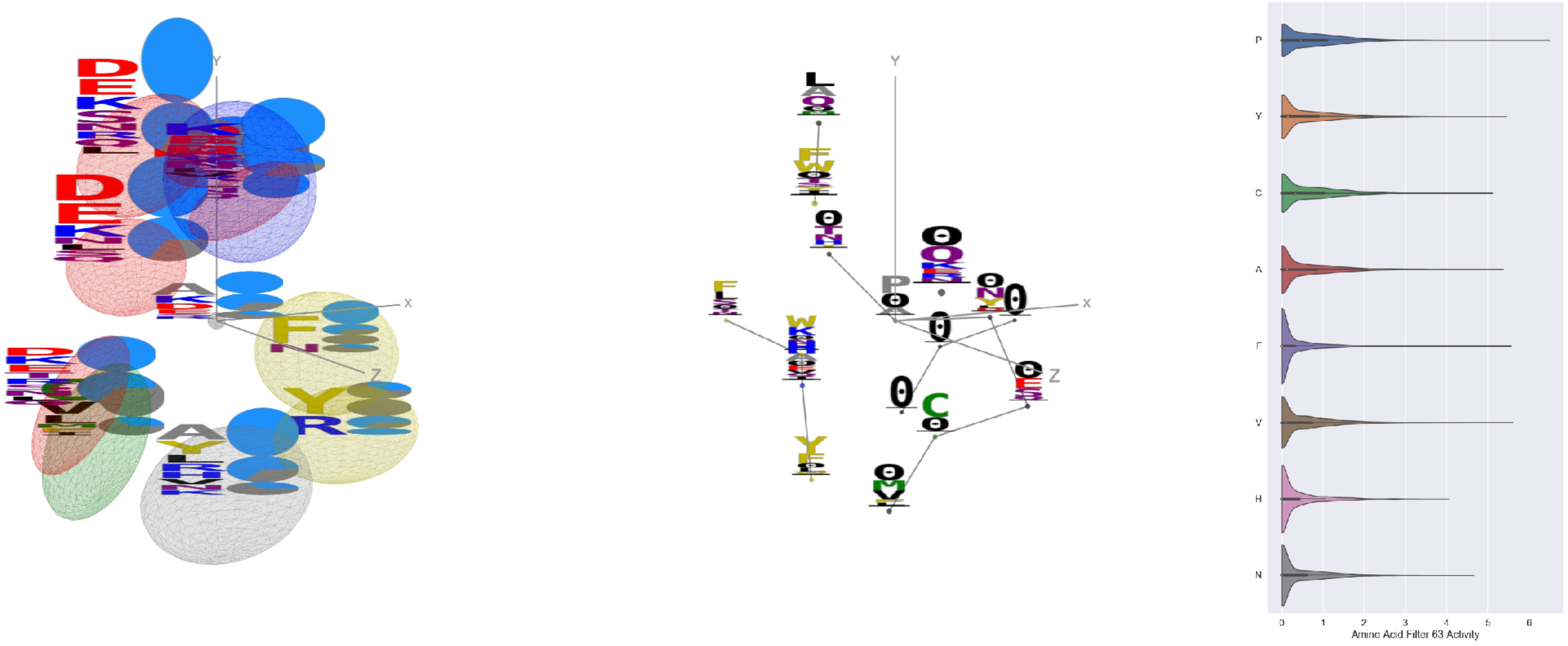

Filter 63
$R_{\text{pred}} = -0.13$
Percent Active = 52
Pooling attention 0.78
Pooling feature 1.56
Top activators
7jw0:E:139 P
6nb7:B:136 P
6nb6:A:136 P
6nb7:B:519 V
6ii9:C:315 P
7jw0:E:497 F
6nb7:B:19 Y
7l57:B:972 A
7jw0:E:972 A
4xmn:E:745 C

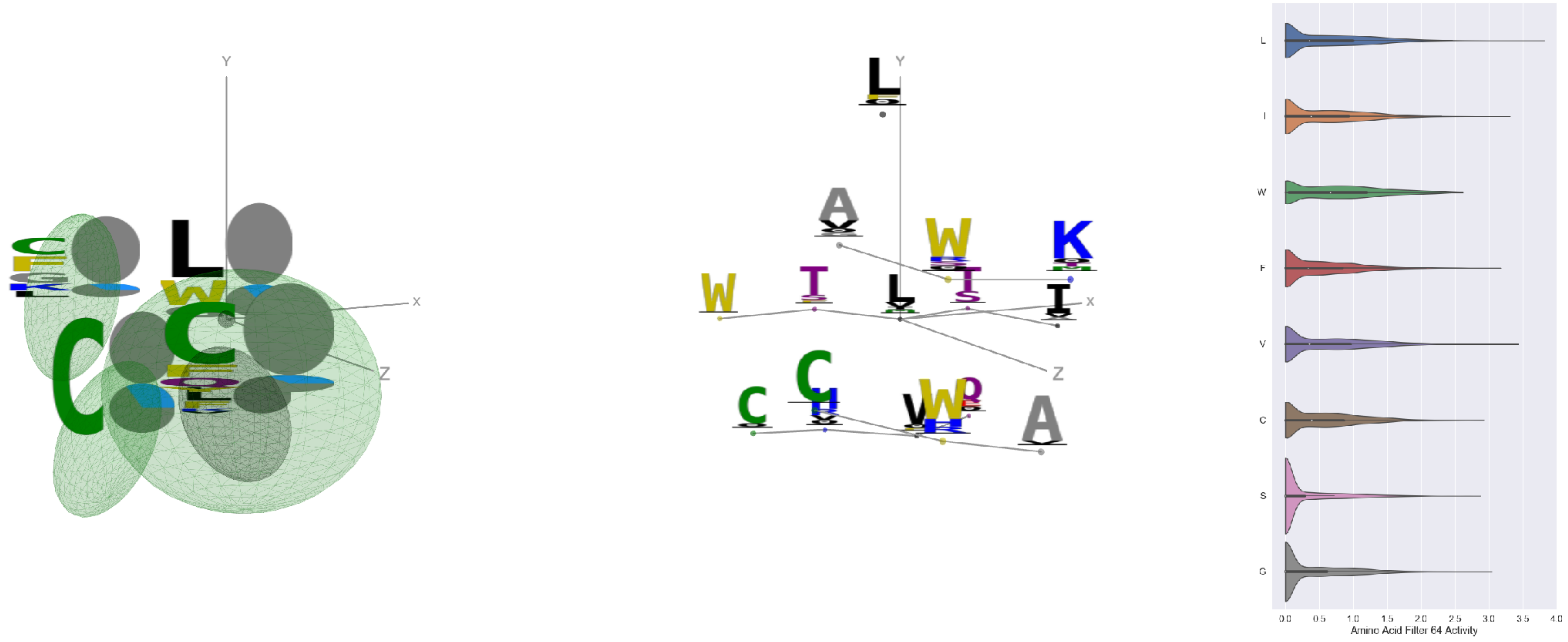

Filter 64
$R_{\text{pred}} = -0.30$
Percent Active = 35
Pooling attention 0.16
Pooling feature 0.51
Top activators
1w72:D:215 L
6nha:B:7 V
3cvh:A:215 L
7nda:A:977 I
7lcn:A:977 I
3hae:A:215 L
6wpt:C:977 L
7jv6:E:977 I
3hae:A:160 L
7ly2:A:977 I

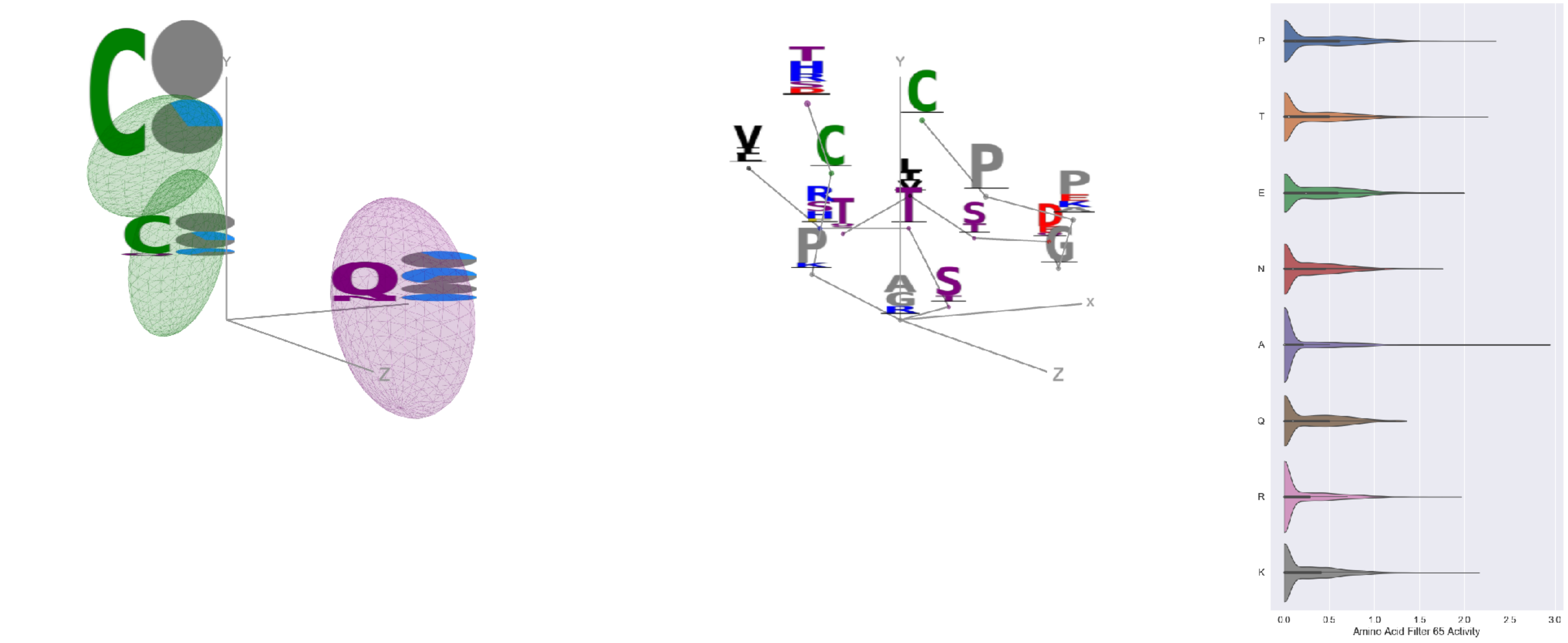

Filter 65
$R_{\text{pred}} = 0.18$
Percent Active = 33
Pooling attention 0.14
Pooling feature 0.46
Top activators
6xob:K:87 A
6mei:C:567 P
4i3r:G:417 P
6nb7:B:509 T
7cwm:A:378 K
6weq:B:284 T
4aei:A:18 P
6m58:A:277 E
4zxb:E:221 Y
7lcn:A:588 R

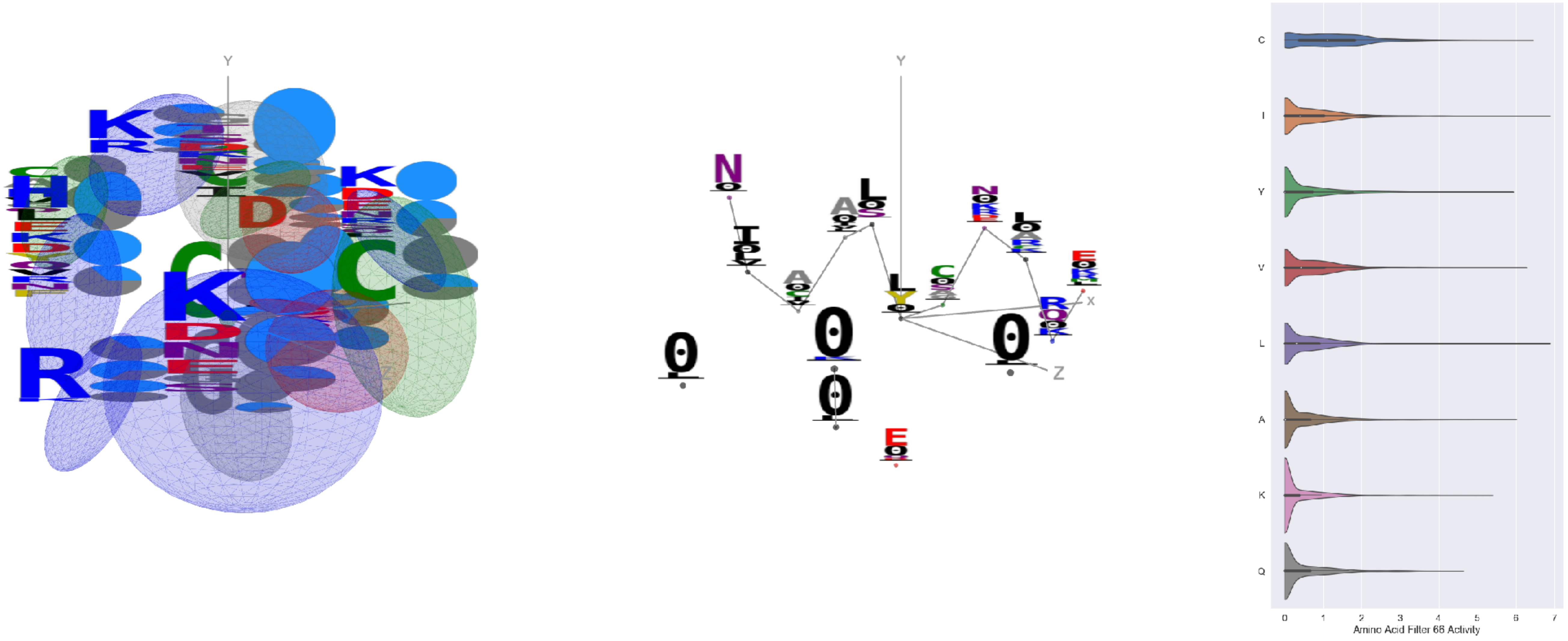

Filter 66
$R_{\text{pred}} = -0.23$
Percent Active = 46
Pooling attention 0.62
Pooling feature 0.48
Top activators
4xmn:E:1000 L
4xmn:E:948 I
4xmn:E:776 C
6nb7:B:570 T
4xmn:E:999 C
4xmn:E:922 L
4xmn:E:868 V
4xmn:E:885 A
4xmn:E:840 G
4xmn:E:875 L

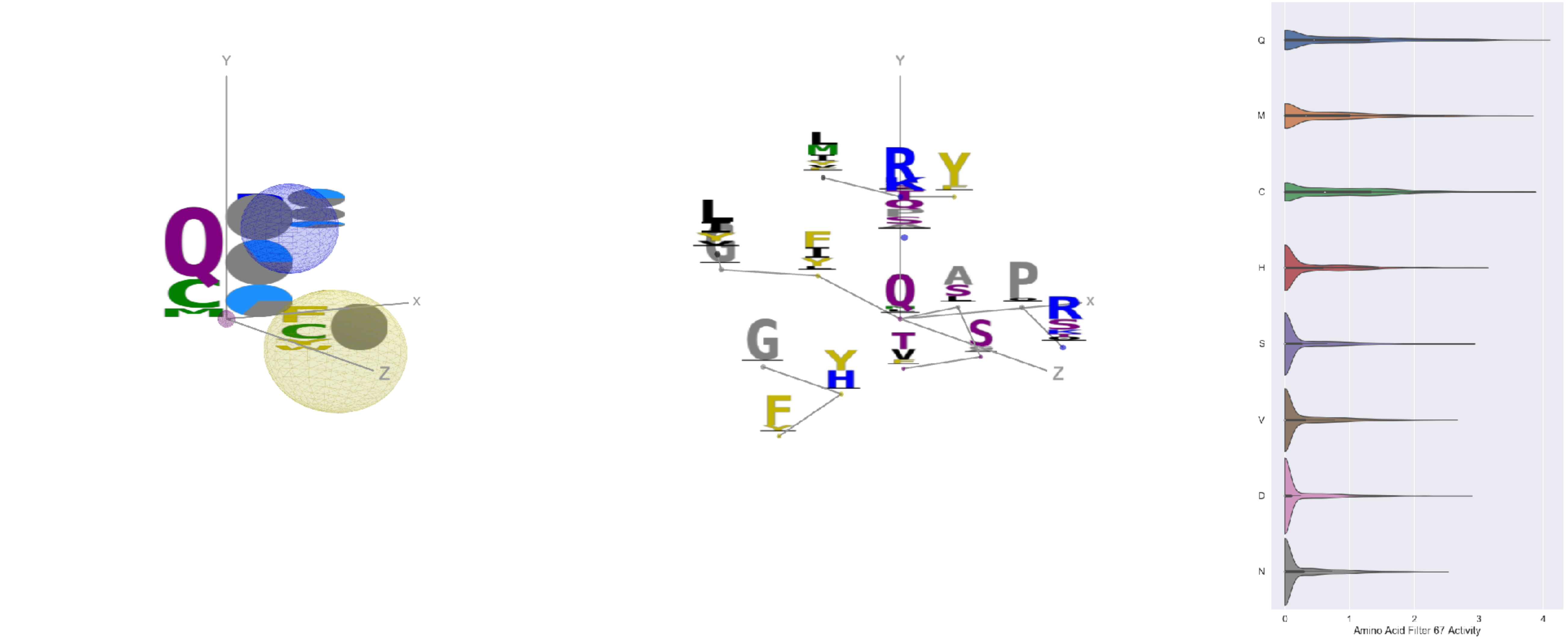

Filter 67
$R_{\text{pred}} = -0.10$
Percent Active = 25
Pooling attention 0.20
Pooling feature 0.26
Top activators
6xob:K:52 Q
6c9u:A:567 Q
6o39:C:109 C
6fgb:A:93 M
3skj:E:159 Q
7jum:A:626 C
3tt3:A:250 Q
4fqv:C:191 Q
3skj:E:164 C
6nb6:C:835 Q

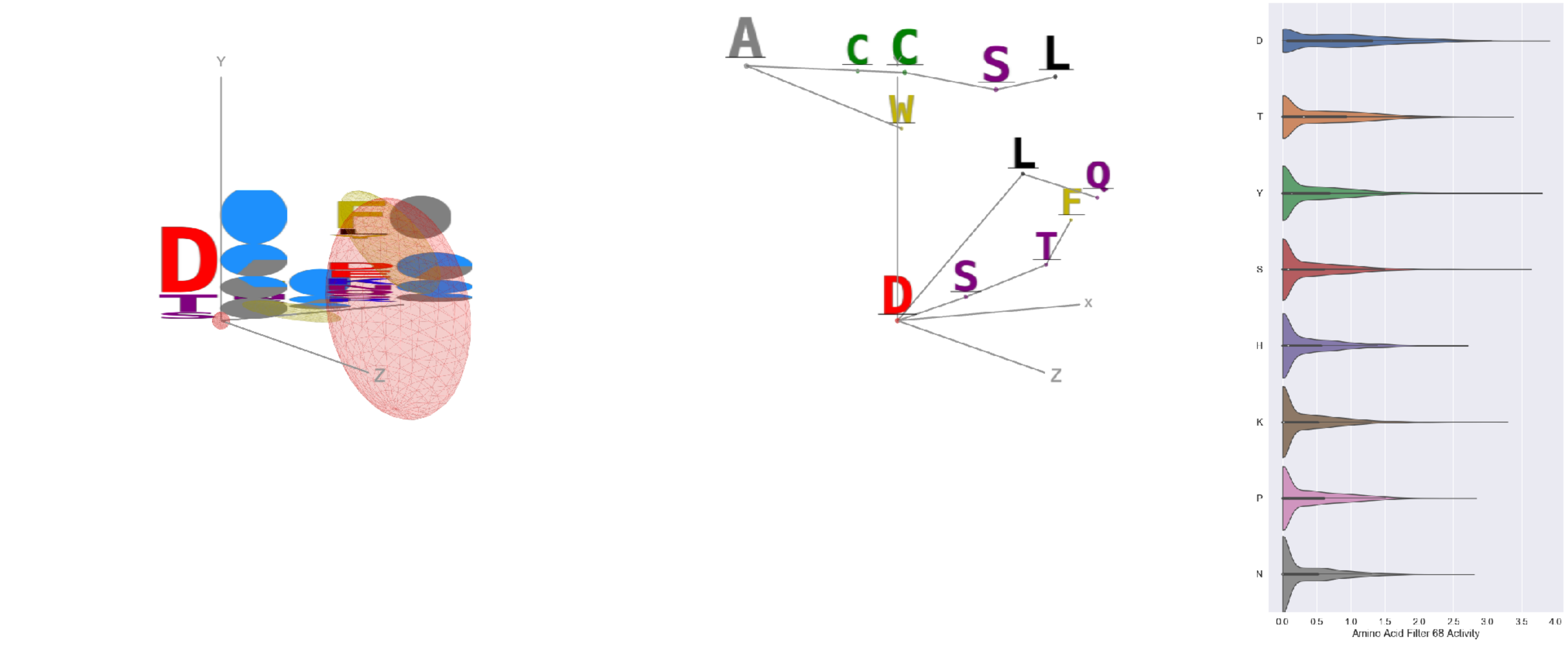

Filter 68
$R_{\mathrm{pred}} = 0.16$
Percent Active = 39
Pooling attention 0.37
Pooling feature 0.41
Top activators
6o2b:M:72 D
6ulf:K:73 D
3uze:C:397 Y
6gg0:4:334 Y
6o2b:M:113 S
6o25:G:72 D
5esv:G:326 Y
6nb6:A:824 D
7a5r:B:279 Y
7l57:B:172 Y

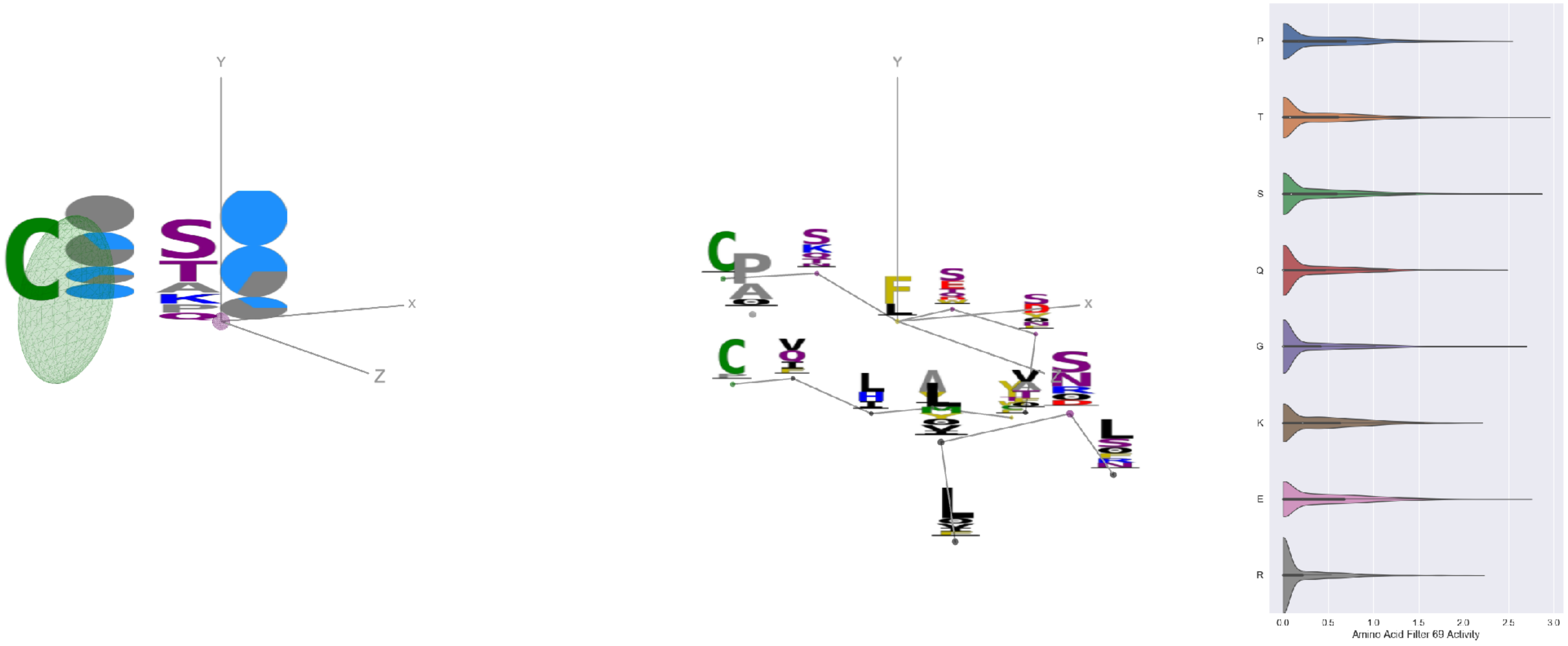

Filter 69
$R_{\mathrm{pred}} = 0.22$
Percent Active = 33
Pooling attention 0.19
Pooling feature 0.36
Top activators
6nb7:B:364 F
7l57:B:549 T
7l57:B:359 S
7a5r:B:530 D
6ii9:C:2 Y
7l57:B:377 F
7jv2:A:443 S
3cvh:A:271 T
6r0x:E:115 E
6nb6:C:364 F

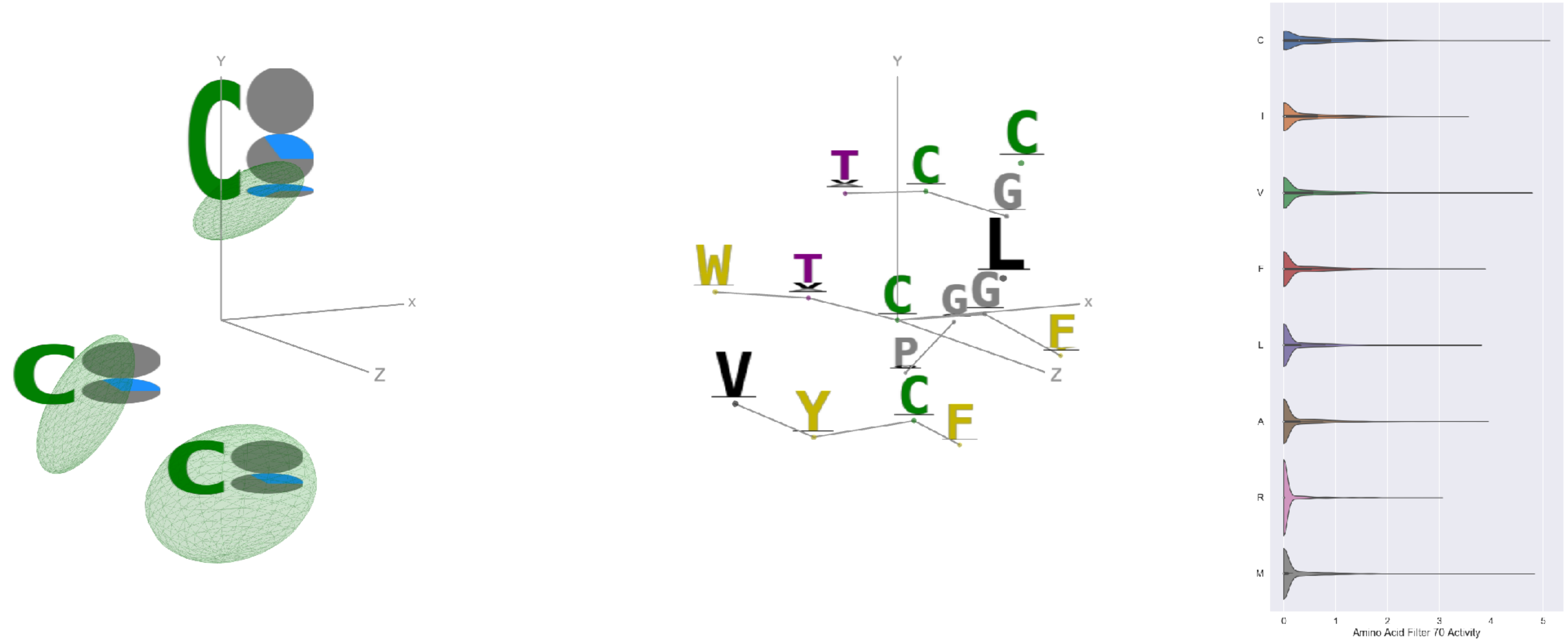

Filter 70
$R_{\mathrm{pred}} = -0.24$
Percent Active = 23
Pooling attention 0.19
Pooling feature 1.01
Top activators
6mej:C:552 C
6e63:A:421 M
7jum:A:723 V
6e62:P:419 G
4mwf:D:552 C
6mei:C:569 C
6e62:P:421 M
3uze:C:358 V
4ala:C:358 V
4uta:A:23 V

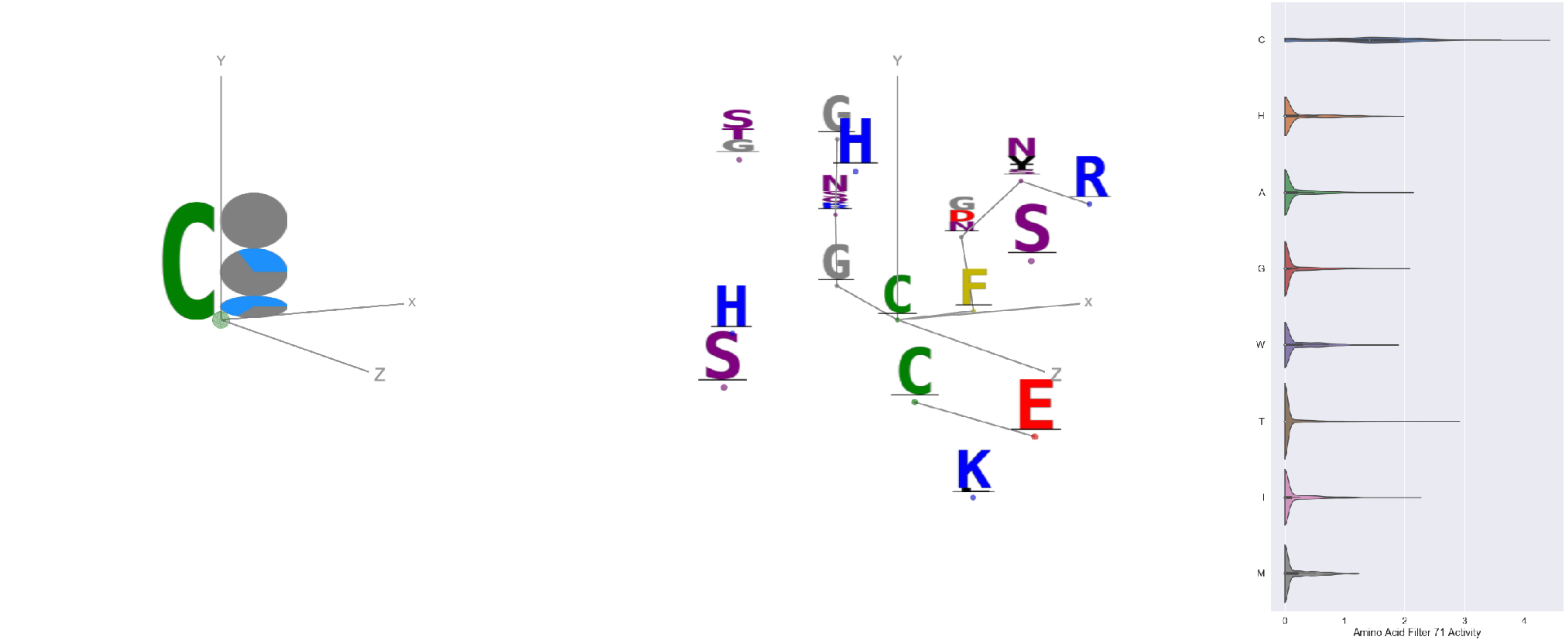

Filter 71
$R_{\mathrm{pred}} = -0.12$
Percent Active = 18
Pooling attention 0.25
Pooling feature 1.01
Top activators
6nb7:B:1025 C
7l2d:A:1043 C
7lxw:A:15 C
7ly2:A:1043 C
7c2l:C:1043 C
4i3r:G:385 C
6nb6:A:1025 C
6nb6:C:1025 C
7l2e:A:1043 C
6bfq:G:96 C

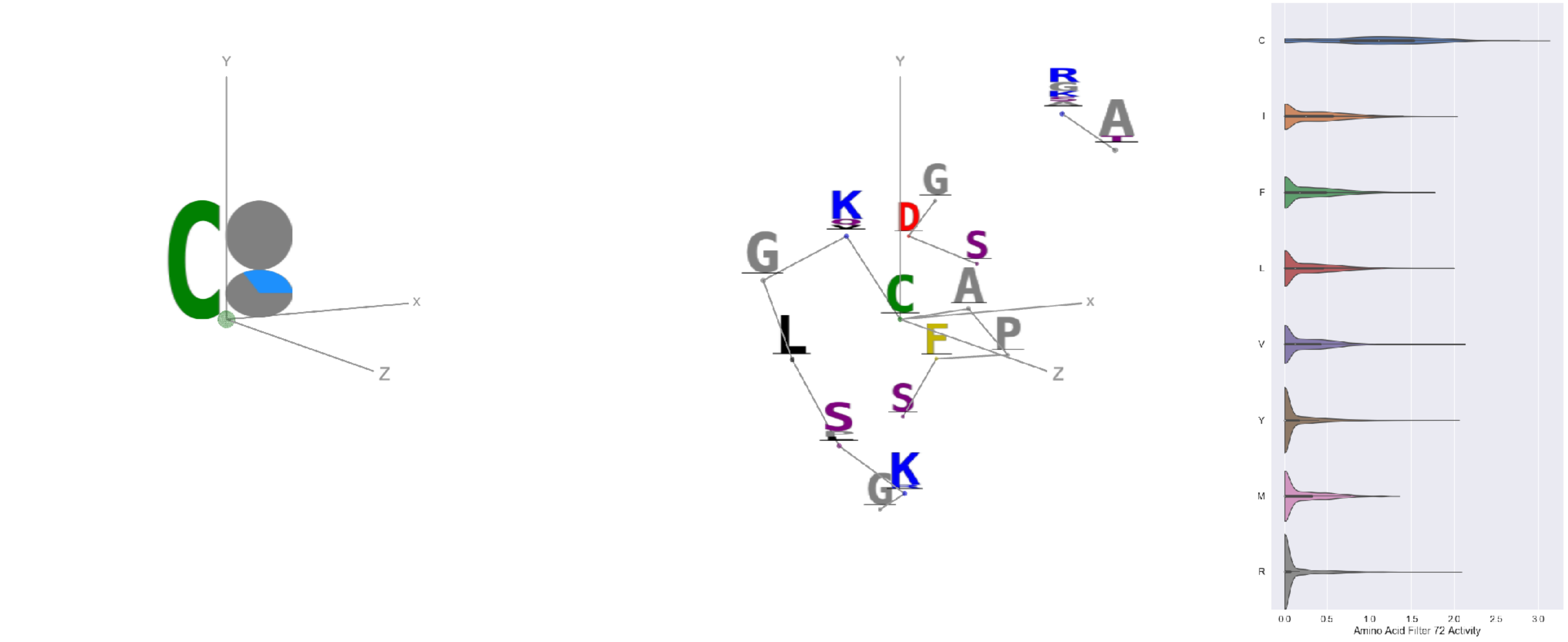

Filter 72
$R_{pred} = -0.14$
Percent Active = 31
Pooling attention 0.04
Pooling feature 0.14
Top activators
6r0x:E:53 C
7l57:B:738 C
7l57:B:749 C
7l57:B:301 C
6nb7:B:348 C
4okv:F:230 C
7jw0:E:749 C
4okv:F:261 C
7lxx:A:131 C
4uao:A:208 C

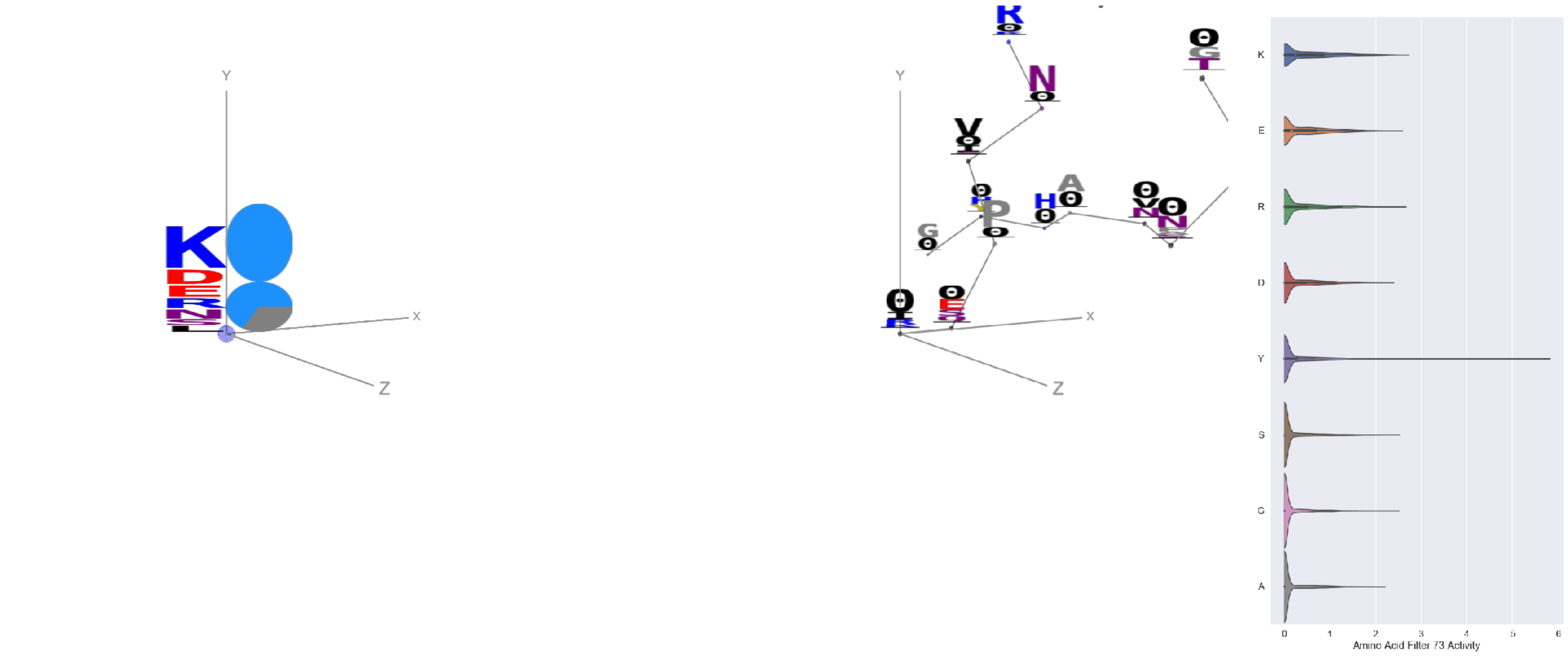

Filter 73
$R_{pred} = 0.15$
Percent Active = 28
Pooling attention 0.13
Pooling feature 0.19
Top activators
6ii9:C:317 Y
5i6x:A:201 Y
3hmx:A:161 Y
3ncy:A:348 Y
6ln2:A:1036 Y
6nb6:A:903 K
2qql:A:209 R
3ncy:B:349 Y
6m58:A:519 Y
7l57:B:702 R

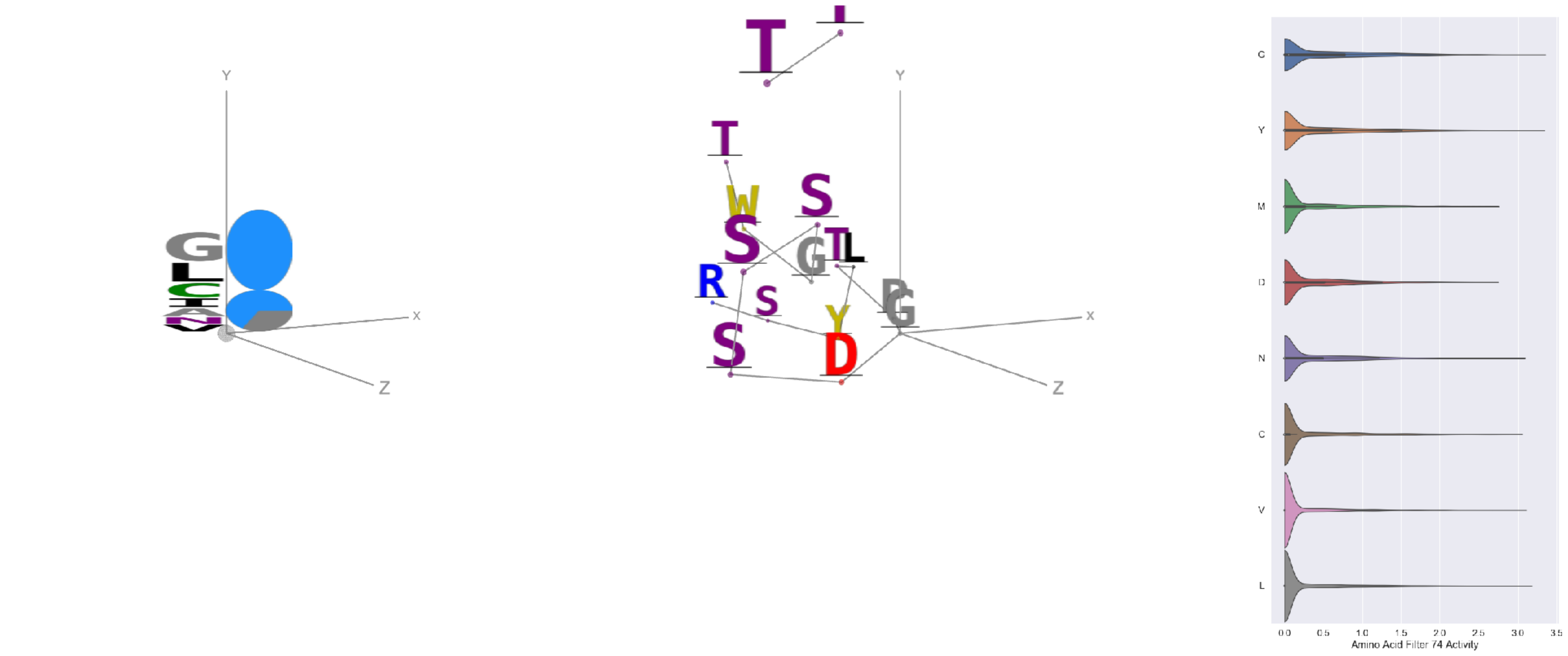

Filter 74
$R_{pred} = 0.20$
Percent Active = 32
Pooling attention 0.44
Pooling feature 0.95
Top activators
7lxx:A:252 G
6oij:A:3 Y
6ii9:C:3 Y
7jv2:A:452 L
6ii9:C:6 G
6weq:B:19 V
7jv2:A:460 N
6nb6:A:833 C
7a5r:B:576 Y
6c9u:A:1 Y

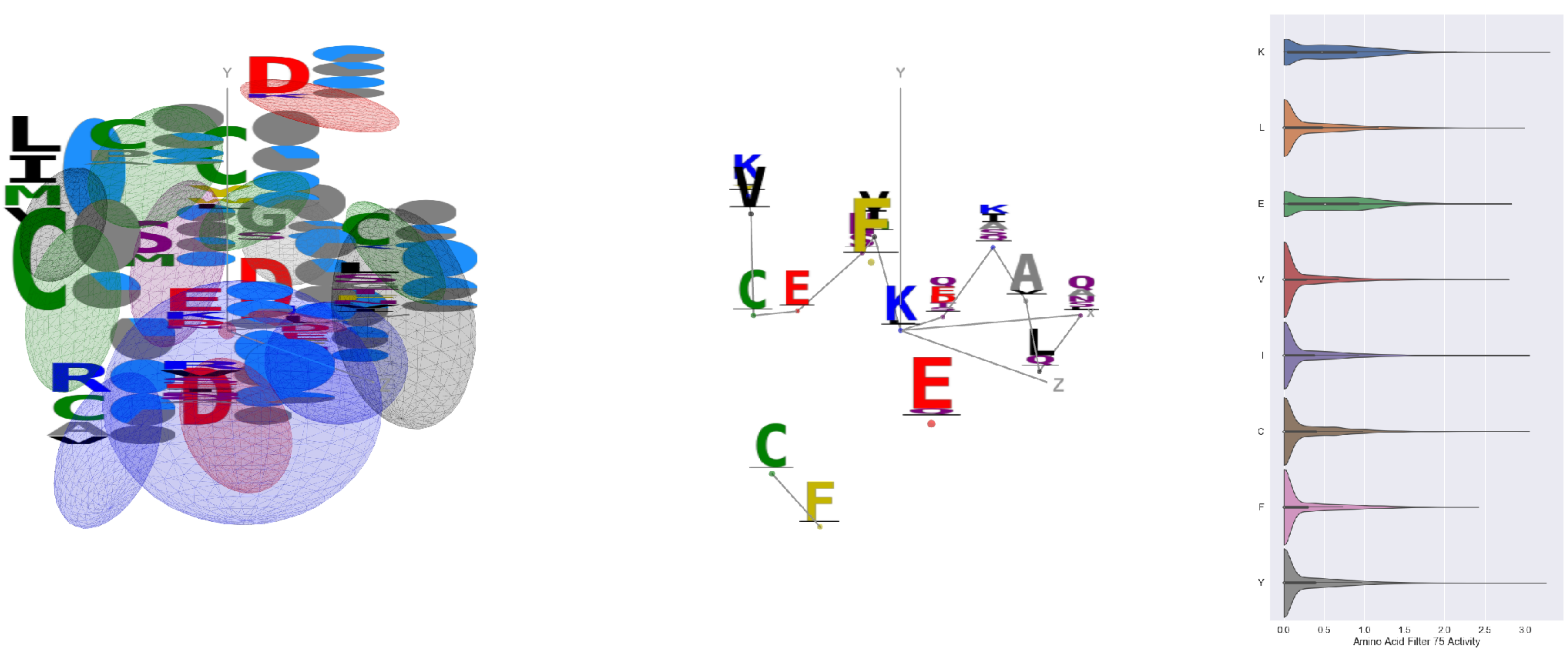

Filter 75
$R_{pred} = -0.06$
Percent Active = 41
Pooling attention 0.15
Pooling feature 1.06
Top activators
7l57:B:1028 K
6nb7:B:90 Y
4xmn:E:827 Y
7jw0:E:513 I
4xmn:E:923 C
4xmn:E:913 R
4xmn:E:765 L
4xmn:E:919 L
6nb6:C:499 I
6m58:A:570 E

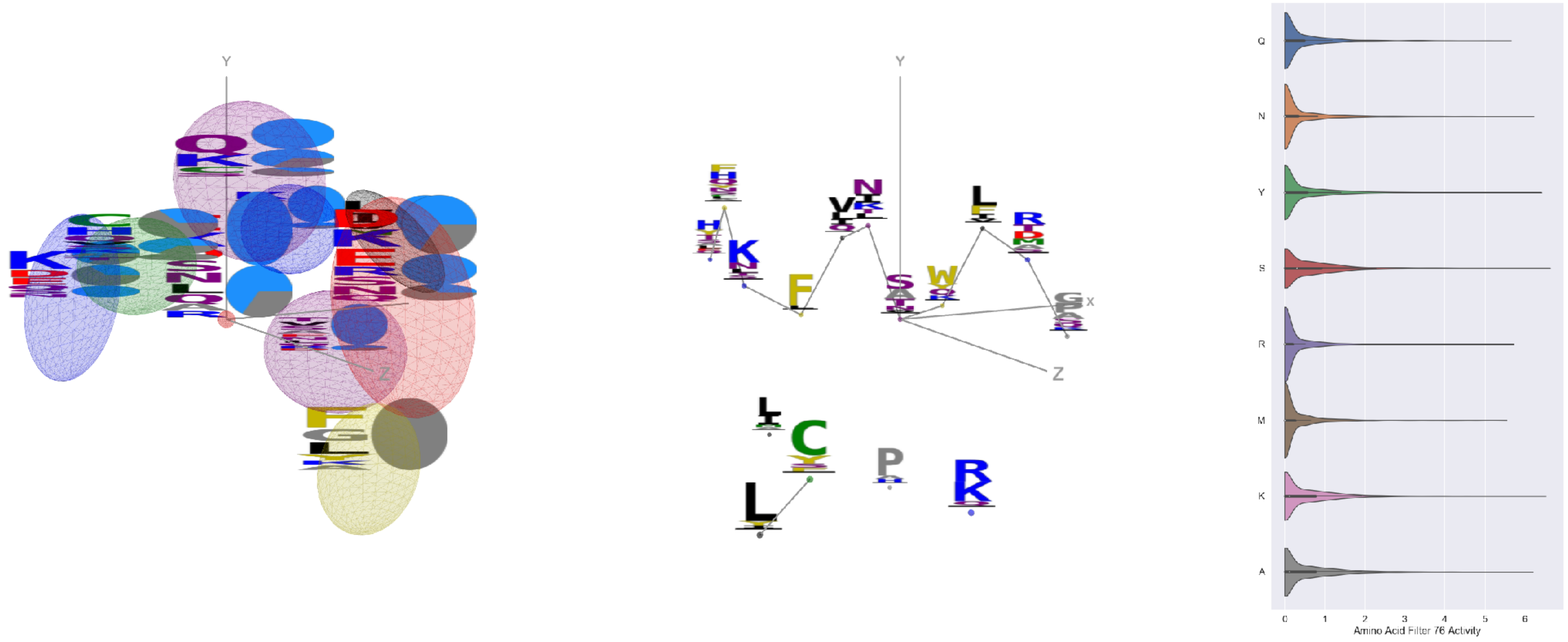


Filter 76
$R_{pred} = -0.19$
Percent Active = 41
Pooling attention 0.36
Pooling feature 0.99
Top activators
4xmn:E:866 S
4xmn:E:952 K
7l57:B:448 Y
6nb6:C:236 Y
6nb7:B:429 N
4xmn:E:966 S
4xmn:E:780 A
6nb7:B:386 K
7jw0:E:188 D
6nb6:C:19 Y

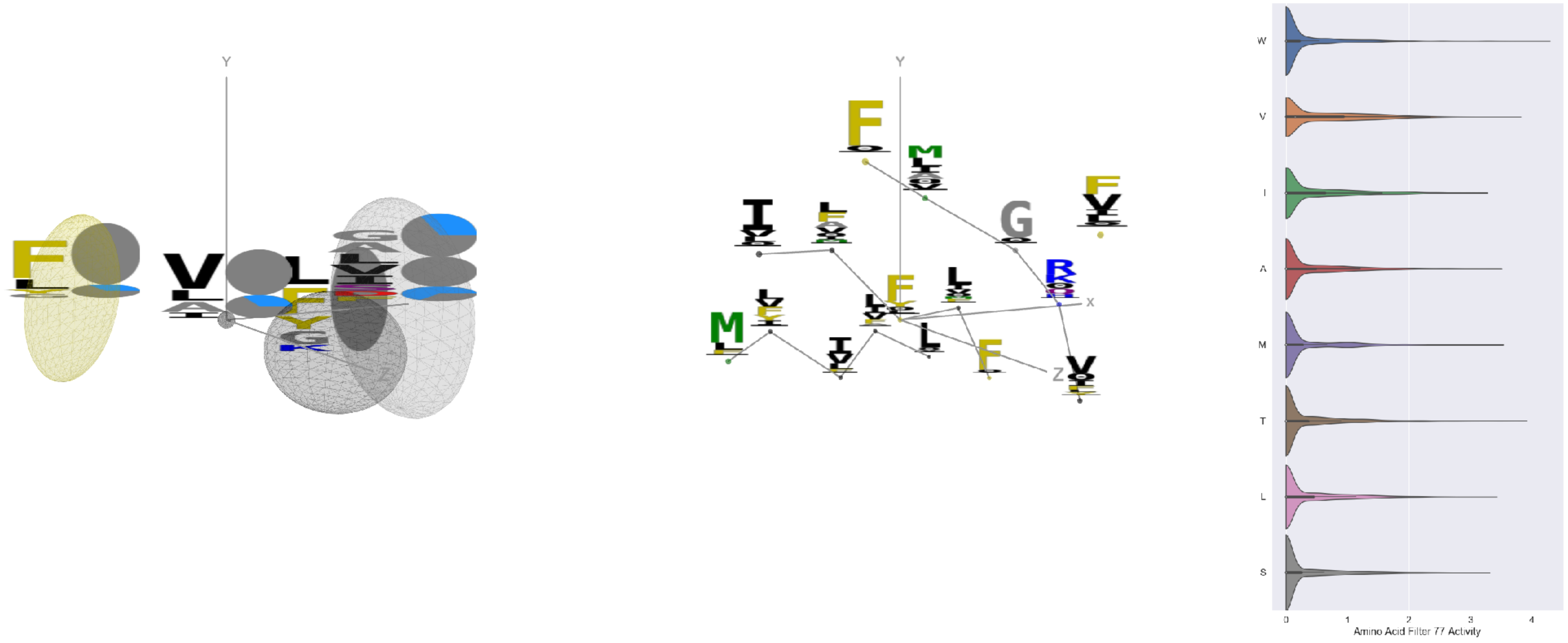


Filter 77
$R_{pred} = -0.15$
Percent Active = 32
Pooling attention 0.30
Pooling feature 1.32
Top activators
6i8s:C:360 F
7c80:B:79 W
3skj:E:59 T
7l2d:A:105 Y
6aod:C:593 V
4dw2:U:231 V
7cws:R:105 Y
7cwm:A:105 Y
7l2e:A:265 Y
7l2e:A:105 Y

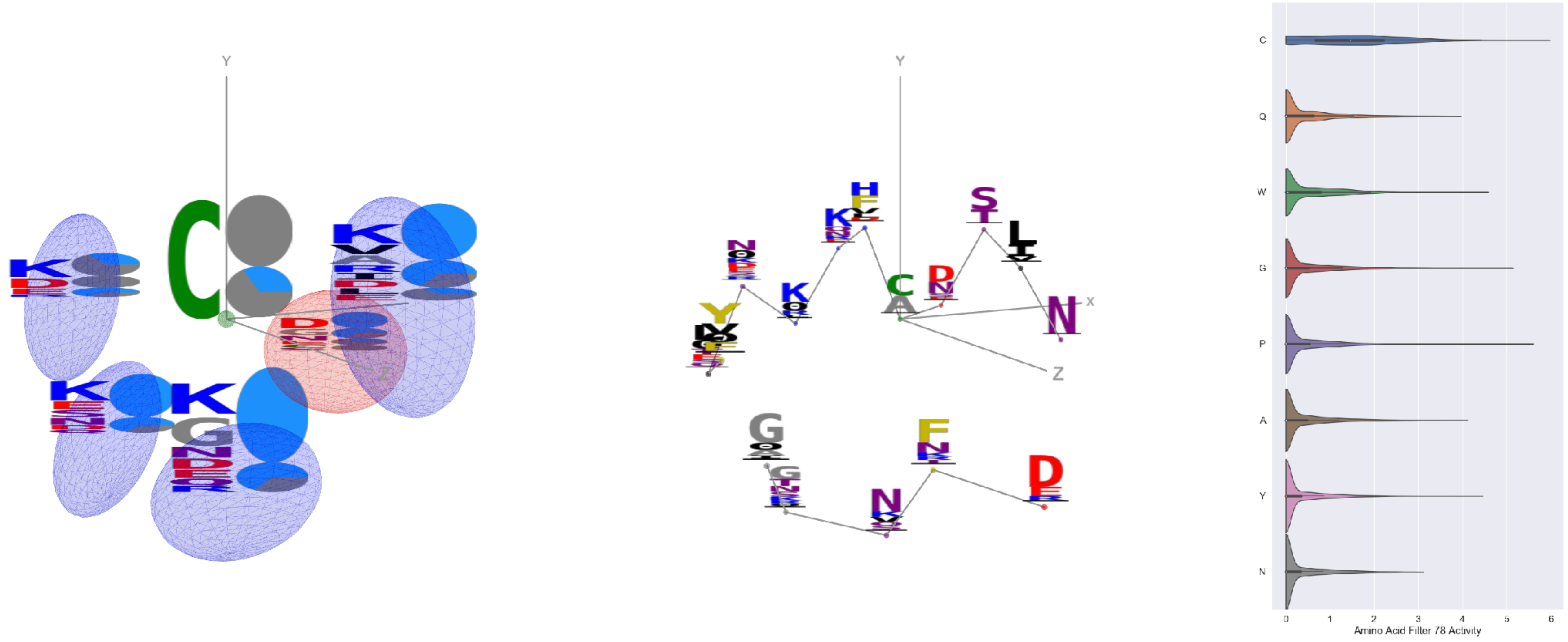


Filter 78
$R_{pred} = -0.06$
Percent Active = 35
Pooling attention 0.33
Pooling feature 0.33
Top activators
6p91:A:123 C
5jym:C:65 P
3iyw:A:305 C
6nb6:C:511 C
3ncy:A:192 G
6aod:C:553 C
3ncy:B:192 G
6p95:C:123 C
5vod:C:43 C
6wtv:D:316 C

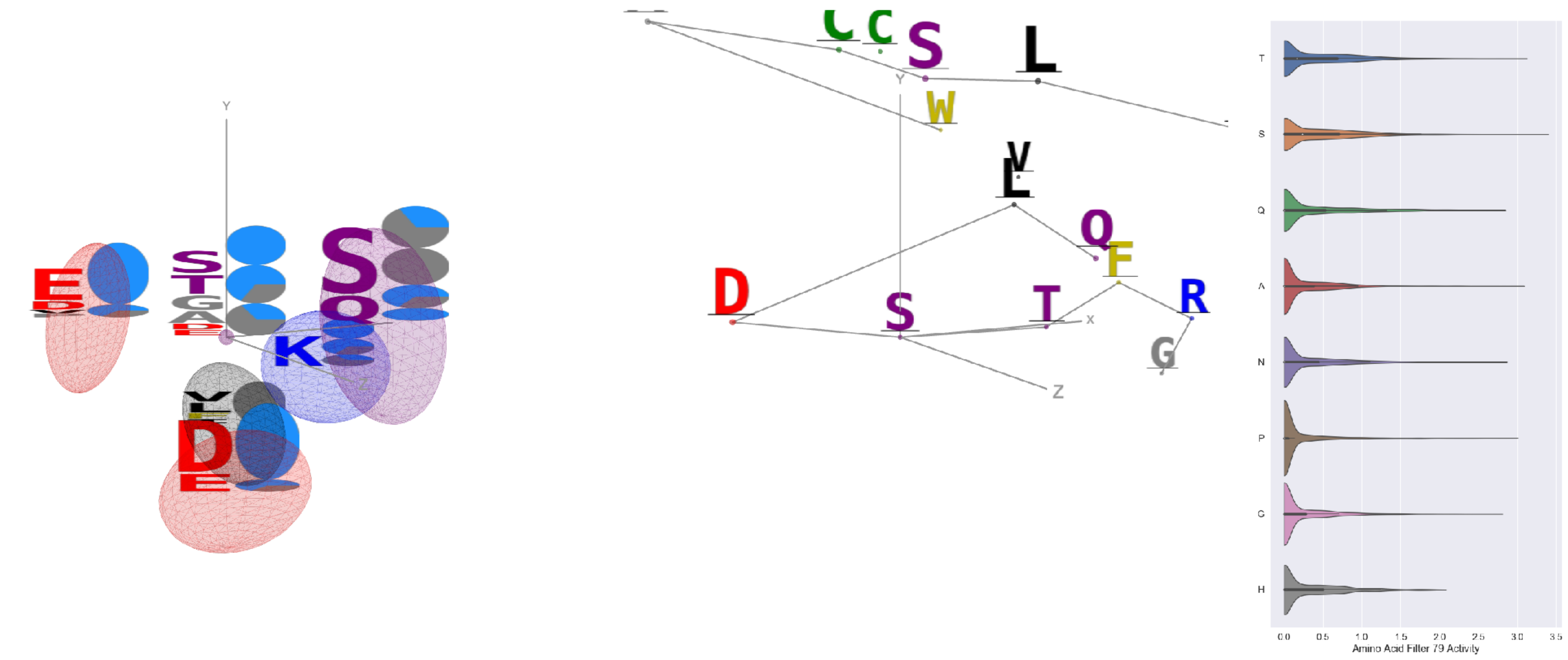


Filter 79
$R_{pred} = -0.01$
Percent Active = 36
Pooling attention 0.05
Pooling feature 0.44
Top activators
6ulf:K:71 S
6o2b:M:70 S
1dee:H:2854 Y
6i07:D:260 S
6umx:A:346 S
7lxw:A:250 T
6o25:G:70 S
3skj:E:122 A
7kr5:E:216 I
7jvr:C:52 P

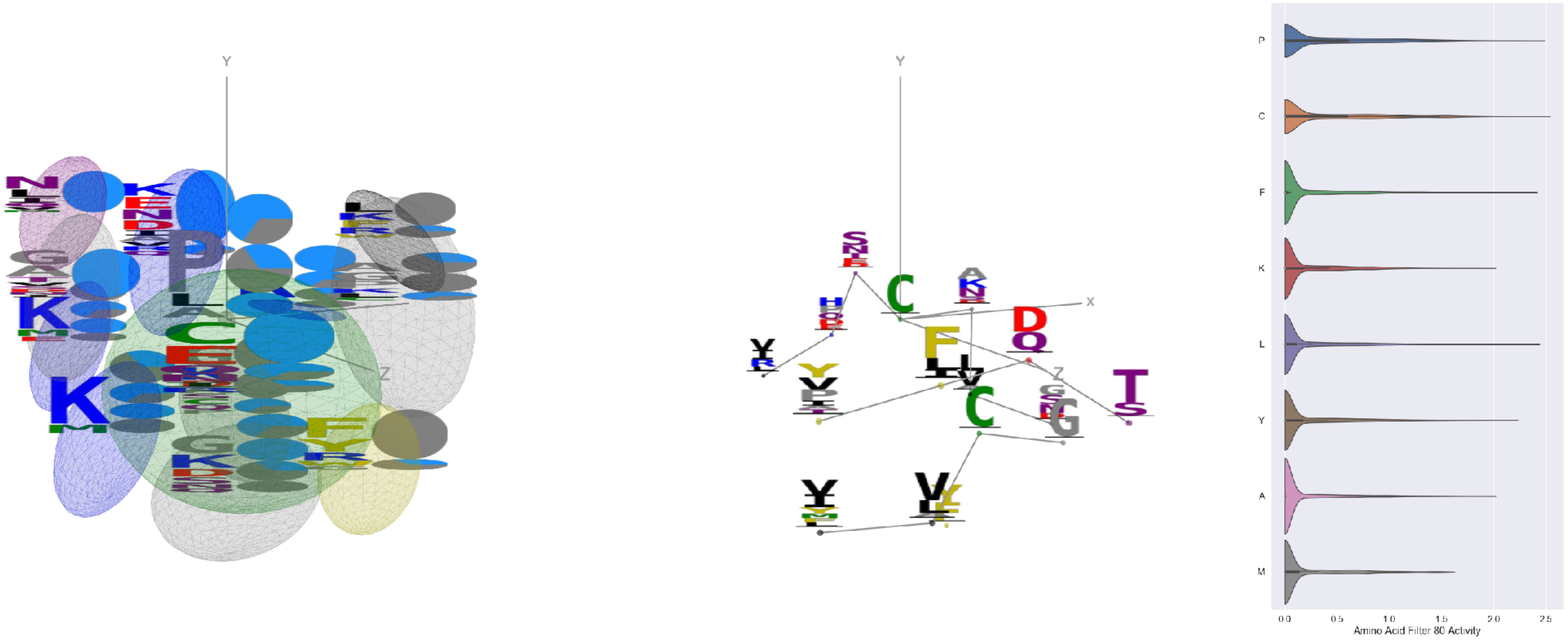


Filter 80
$R_{\mathrm{pred}} = 0.10$
Percent Active = 28
Pooling attention 0.27
Pooling feature 2.94
Top activators
7l57:B:617 C
5n0a:A:75 P
6r0x:E:27 P
6icf:A:195 L
6nb7:B:805 F
6gg0:3:273 Q
6m58:A:279 C
6ii9:C:64 P
6m58:A:361 C
4uta:A:75 P

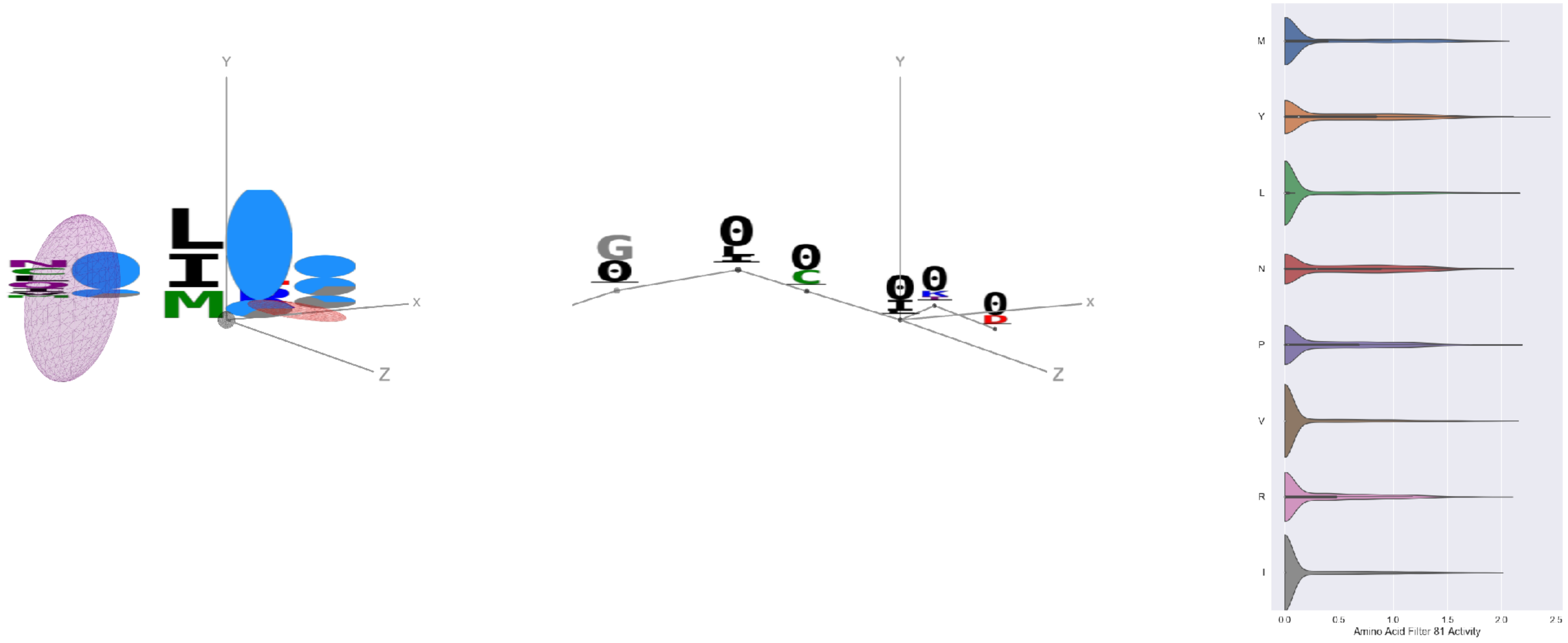


Filter 81
$R_{\mathrm{pred}} = 0.28$
Percent Active = 42
Pooling attention 0.33
Pooling feature 0.67
Top activators
6ii9:C:3 Y
6mlm:A:29 Y
7kra:A:740 G
5vod:C:140 Y
6mhg:E:8 P
7c80:C:5 Y
7kra:A:737 L
6gg0:4:333 V
6aj7:C:25 Y
7ebz:A:4 Y

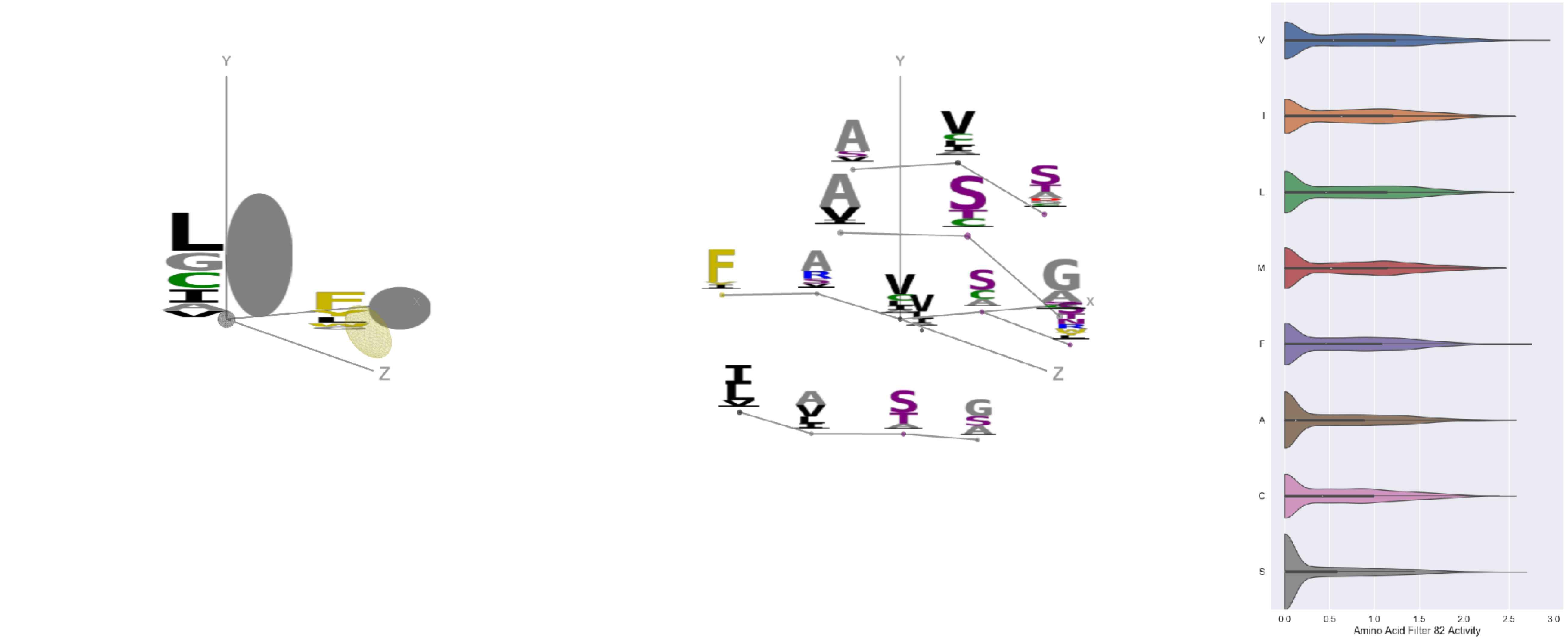


Filter 82
$R_{\mathrm{pred}} = -0.31$
Percent Active = 43
Pooling attention 0.40
Pooling feature 0.92
Top activators
6oik:B:149 V
6pt0:B:318 V
7jvr:B:149 V
2nr6:B:262 F
6pt0:B:149 V
4jr9:A:144 G
6wdt:B:110 V
6wdt:A:235 S
6gv4:B:118 V
5kqv:E:505 V

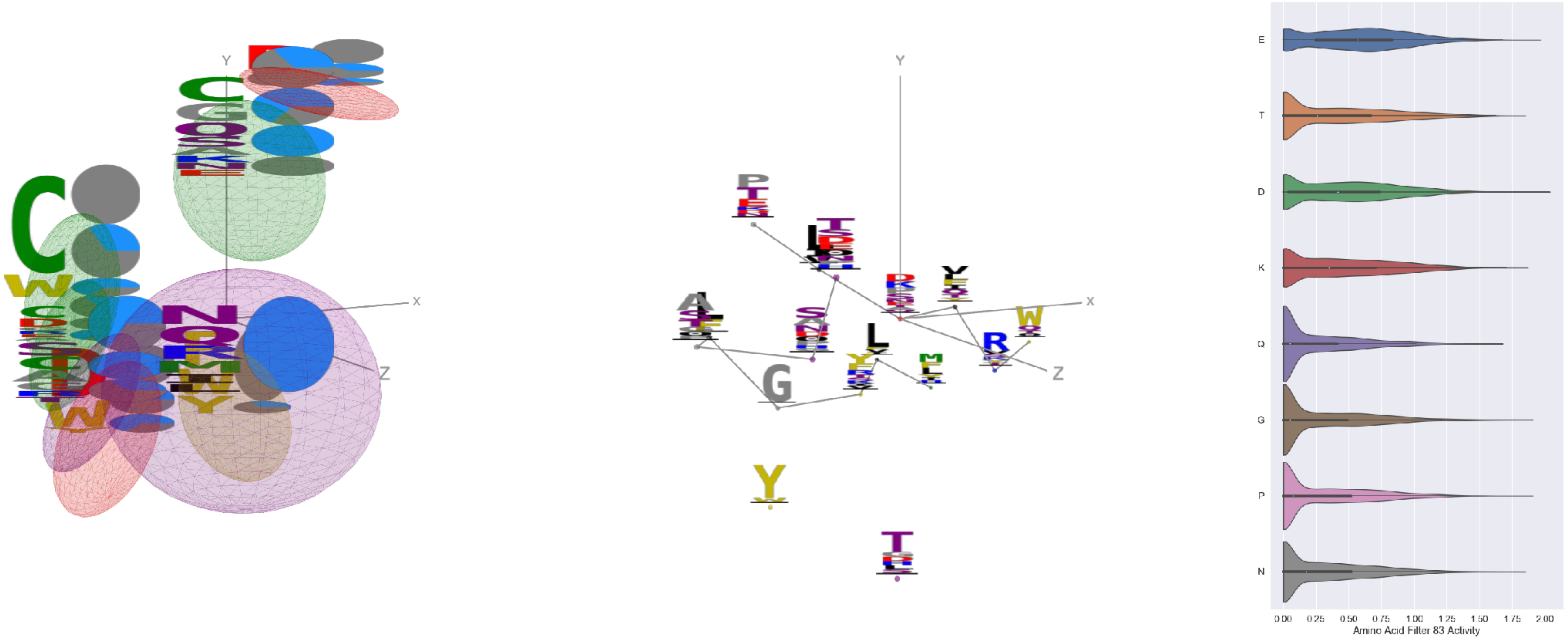


Filter 83
$R_{\mathrm{pred}} = 0.24$
Percent Active = 47
Pooling attention 0.01
Pooling feature 1.04
Top activators
6gv4:B:92 D
6weq:B:310 E
7k93:C:310 E
4zxb:E:250 G
7lm9:A:459 P
5kqv:E:287 E
5tl5:A:18 L
4uao:A:190 K
5i5k:B:1325 T
6wha:C:313 N

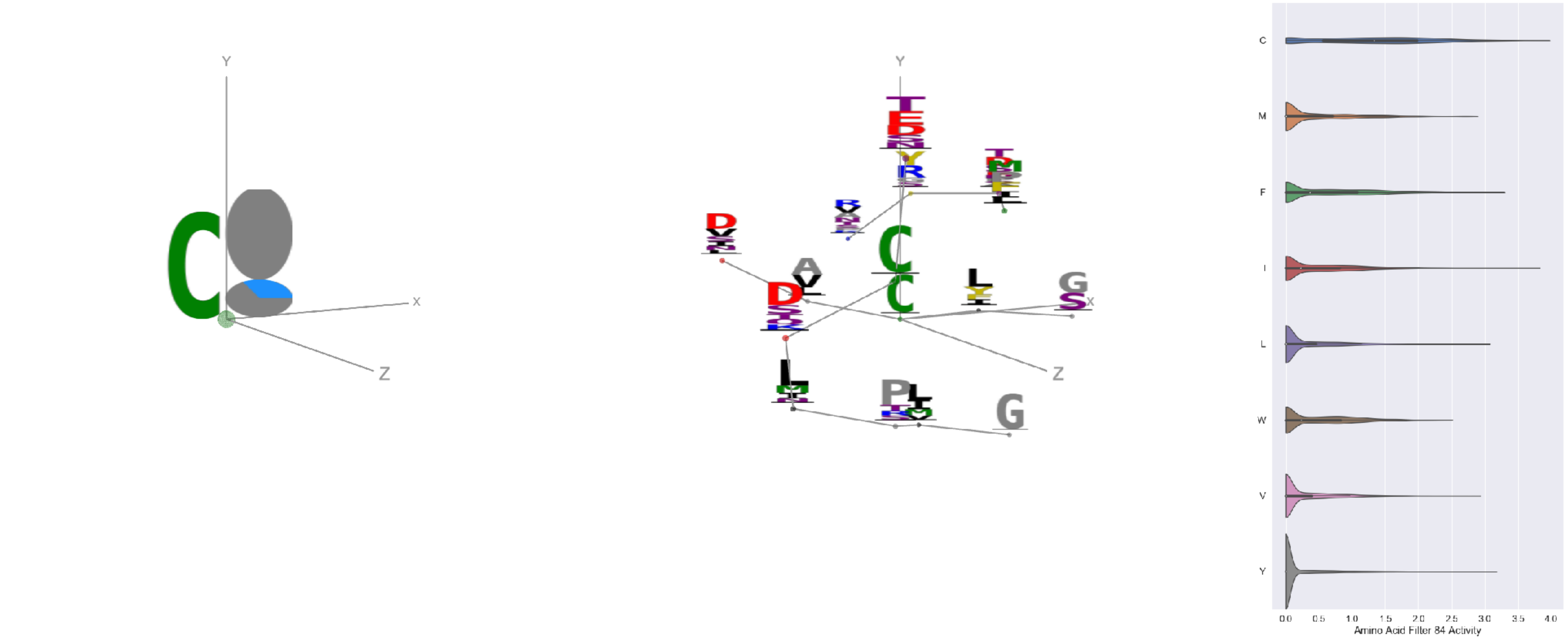


Filter 84
$R_{pred} = -0.17$
Percent Active = 23
Pooling attention 0.09
Pooling feature 0.43
Top activators
6wpt:C:671 C
7jw0:E:671 C
4zxb:E:104 I
6wpt:C:738 C
7jv6:E:738 C
7cws:R:432 C
7nda:A:738 C
7l2d:A:1082 C
7lcn:A:671 C
7l57:B:1082 C

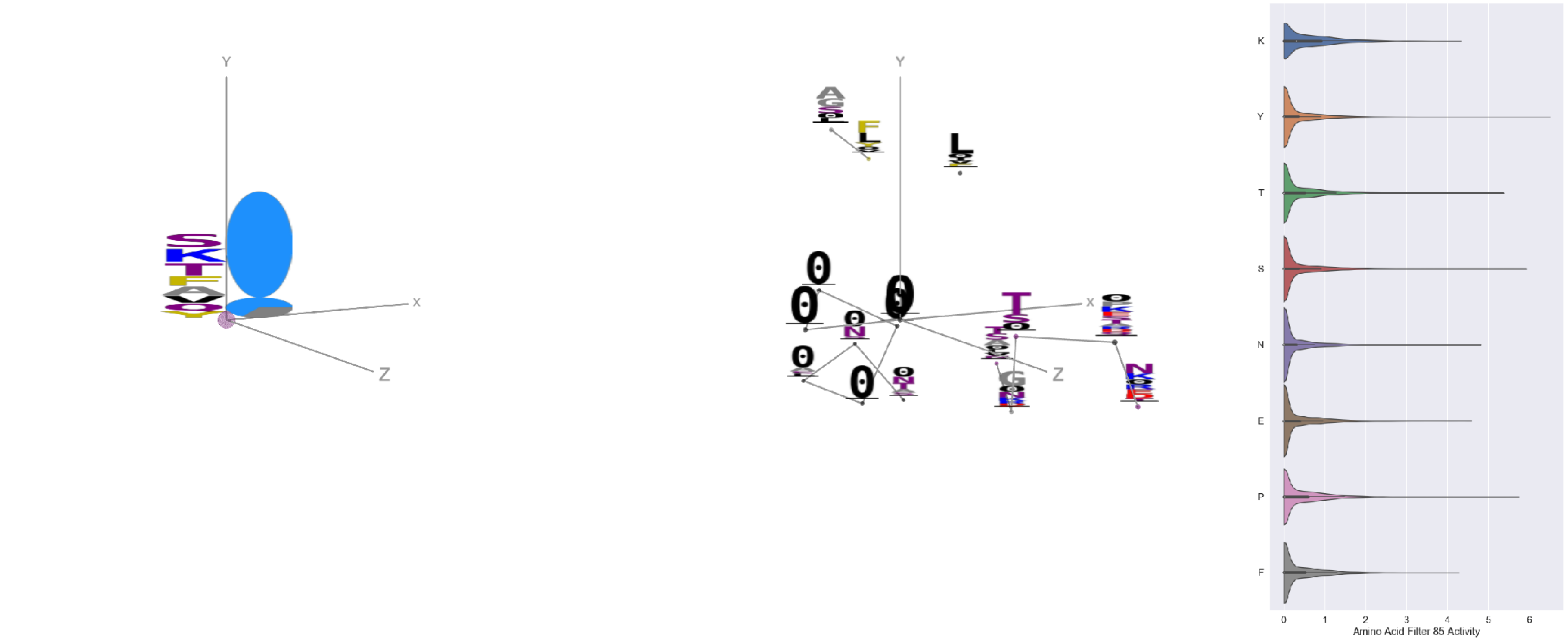


Filter 85
$R_{pred} = 0.05$
Percent Active = 35
Pooling attention 0.01
Pooling feature 0.24
Top activators
6i8s:C:6 Y
3tt3:A:11 R
6p95:c:333 S
6wtv:D:168 Y
5f3j:A:219 Y
4xmn:E:770 P
4cmh:A:48 Y
5tr1:B:36 Y
3jbq:F:532 Y
4uao:A:303 T

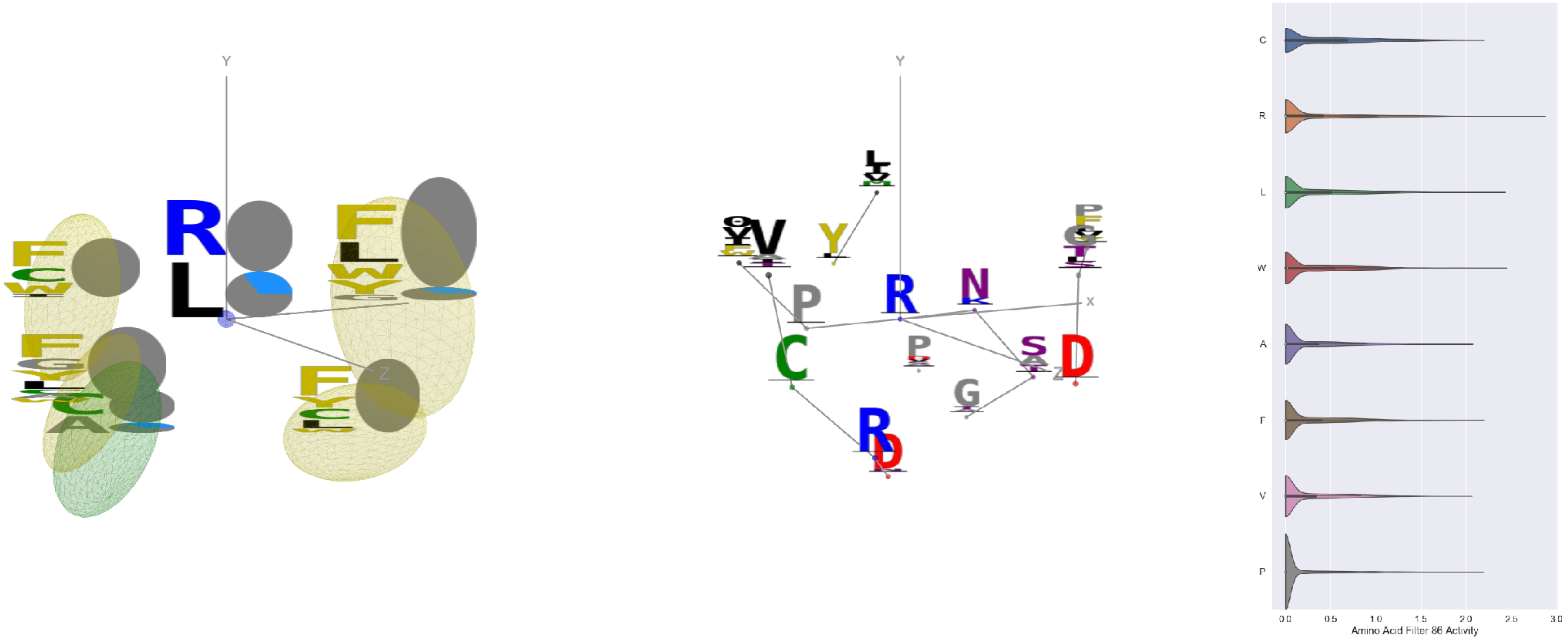


Filter 86
$R_{pred} = -0.27$
Percent Active = 25
Pooling attention 0.48
Pooling feature 1.51
Top activators
6pzw:C:300 R
6q23:A:300 R
1nca:N:300 R
7ce2:A:1303 W
6w7s:A:1693 L
7cws:R:1004 L
7cwm:A:1004 L
7k93:C:336 R
7lcn:A:1004 L
6p95:c:315 L

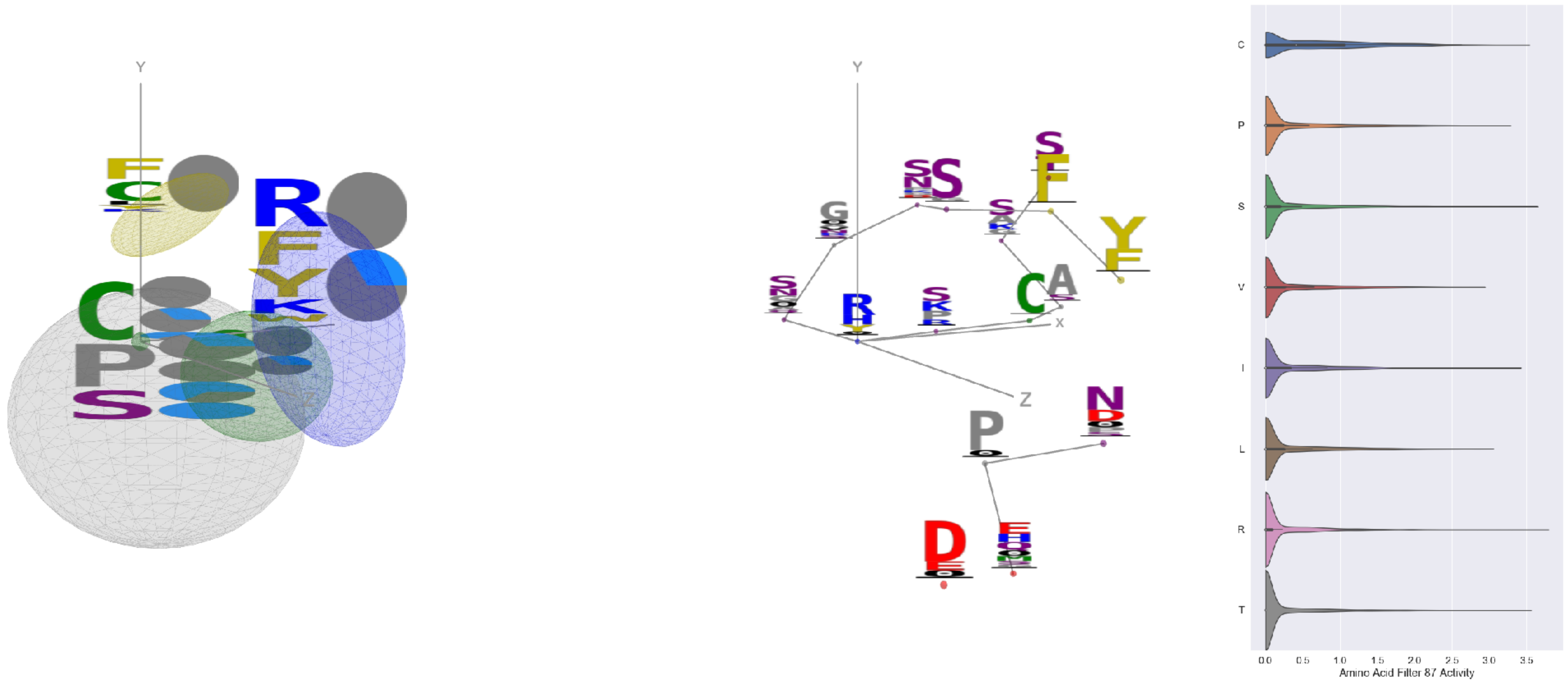


Filter 87
$R_{pred} = -0.01$
Percent Active = 28
Pooling attention 0.28
Pooling feature 0.99
Top activators
6ii9:C:131 R
7lm9:A:362 S
6xkq:A:438 T
6mlm:A:113 C
2qql:A:149 C
6nha:B:57 N
6was:G:161 I
7kra:A:617 T
5y11:C:216 C
4zxb:E:160 P

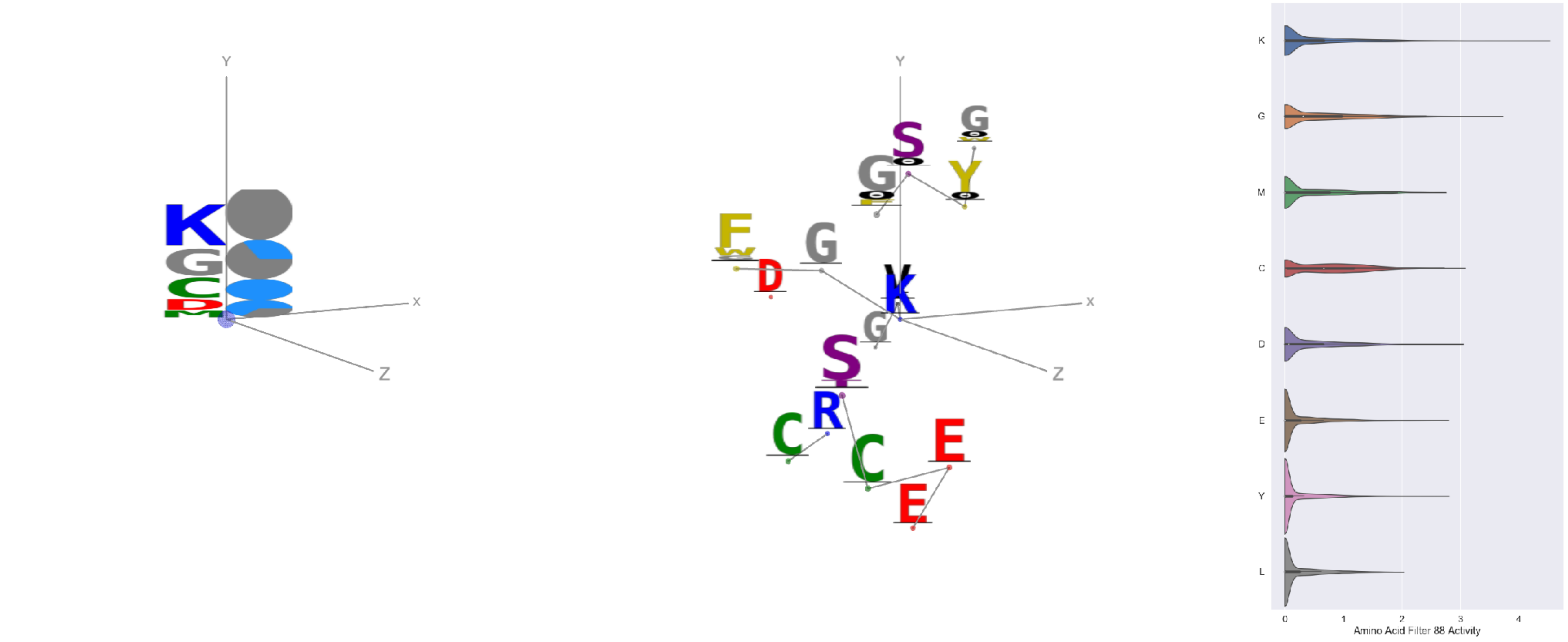

Filter 88
$R_{\text{pred}} = -0.10$
Percent Active = 32
Pooling attention 0.36
Pooling feature 0.69
Top activators
6pzw:C:350 K
1nca:N:350 K
6q23:A:350 K
7ly2:A:1028 K
7lxx:A:77 K
7l2d:A:1028 K
2vh5:R:16 K
7c2l:C:1028 K
6mei:C:517 G
7cwm:A:1028 K

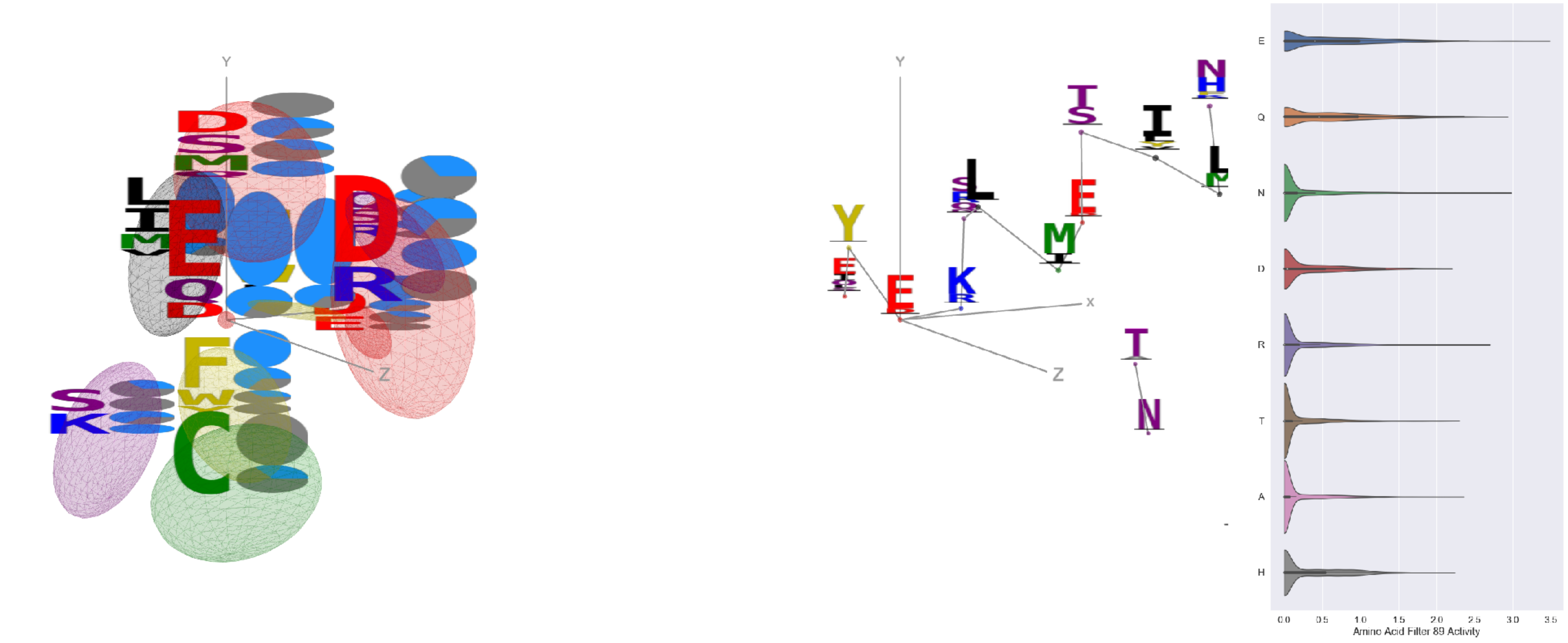

Filter 89
$R_{\text{pred}} = 0.11$
Percent Active = 23
Pooling attention 0.04
Pooling feature 0.10
Top activators
6p95:c:418 E
5d72:B:21 E
7nda:C:819 E
7nda:A:819 E
7l2d:A:819 E
7jv6:E:819 E
6gg0:4:33 E
7nd9:B:819 E
6p95:c:413 E
6gg0:3:33 E

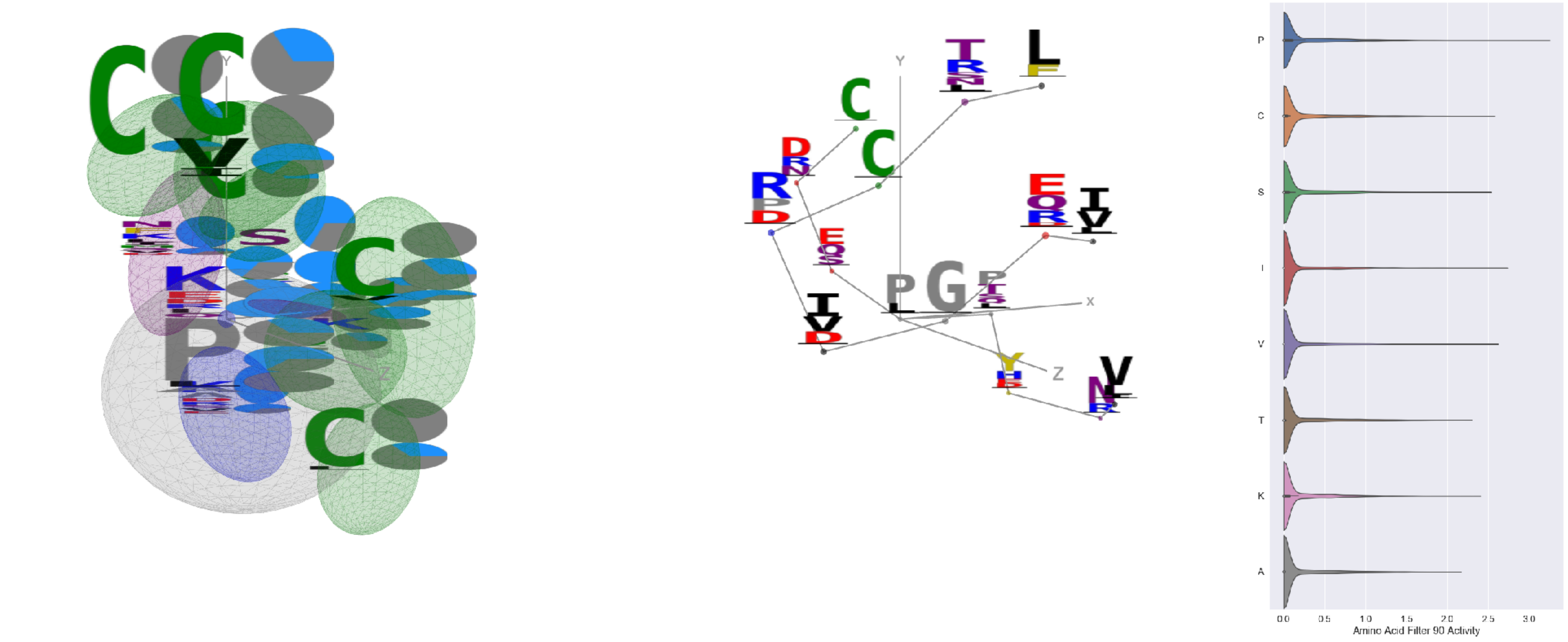

Filter 90
$R_{\text{pred}} = 0.03$
Percent Active = 23
Pooling attention 0.90
Pooling feature 5.02
Top activators
5vod:C:40 P
5bo1:A:305 P
6xob:K:214 I
6xob:K:216 G
7l57:B:495 Y
7l57:B:320 V
3iyw:A:6 V
3uze:D:336 P
6weq:B:291 C
4ala:C:336 P

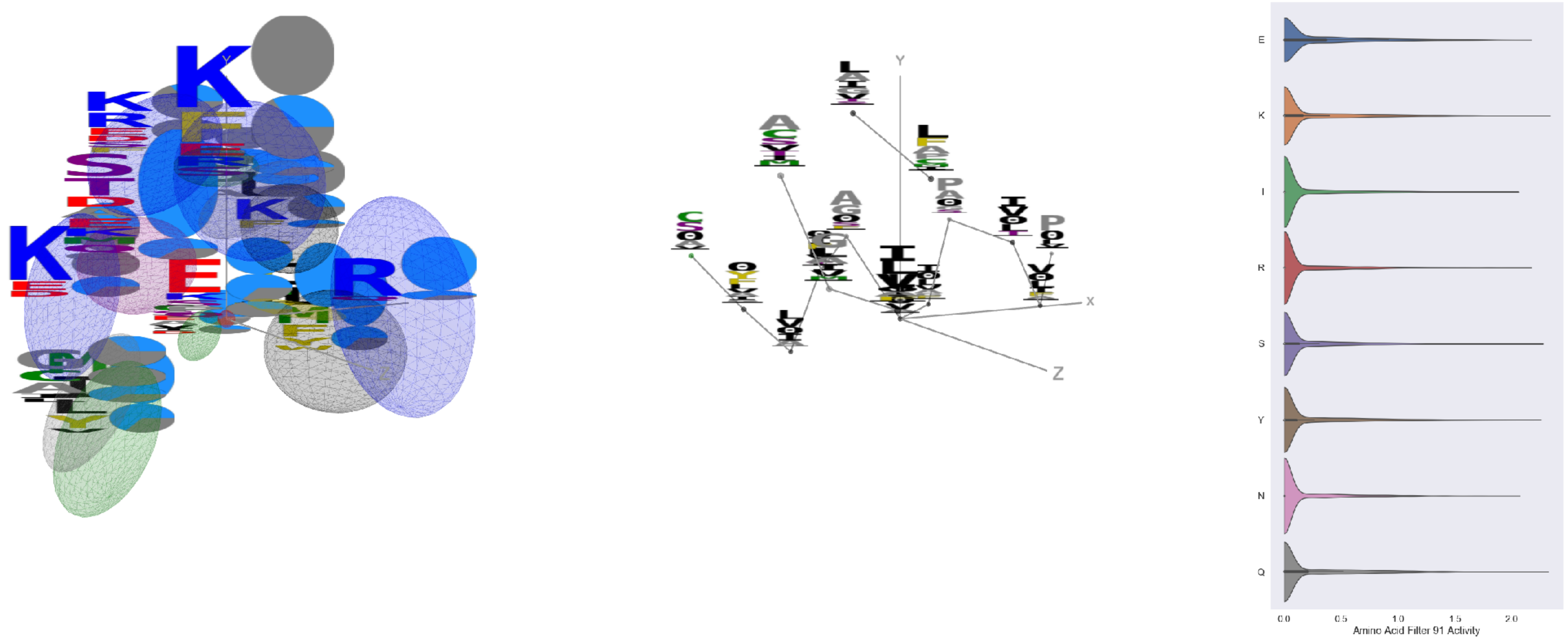

Filter 91
$R_{\text{pred}} = 0.10$
Percent Active = 25
Pooling attention 0.67
Pooling feature 3.48
Top activators
3ncy:B:393 L
3ncy:A:393 L
4fqv:D:3 F
1ynt:G:3091 K
4uao:A:119 Q
6w7s:A:1810 Q
7l2e:A:414 S
7d77:B:56 G
6ln2:A:1044 Y
6oij:A:19 Y

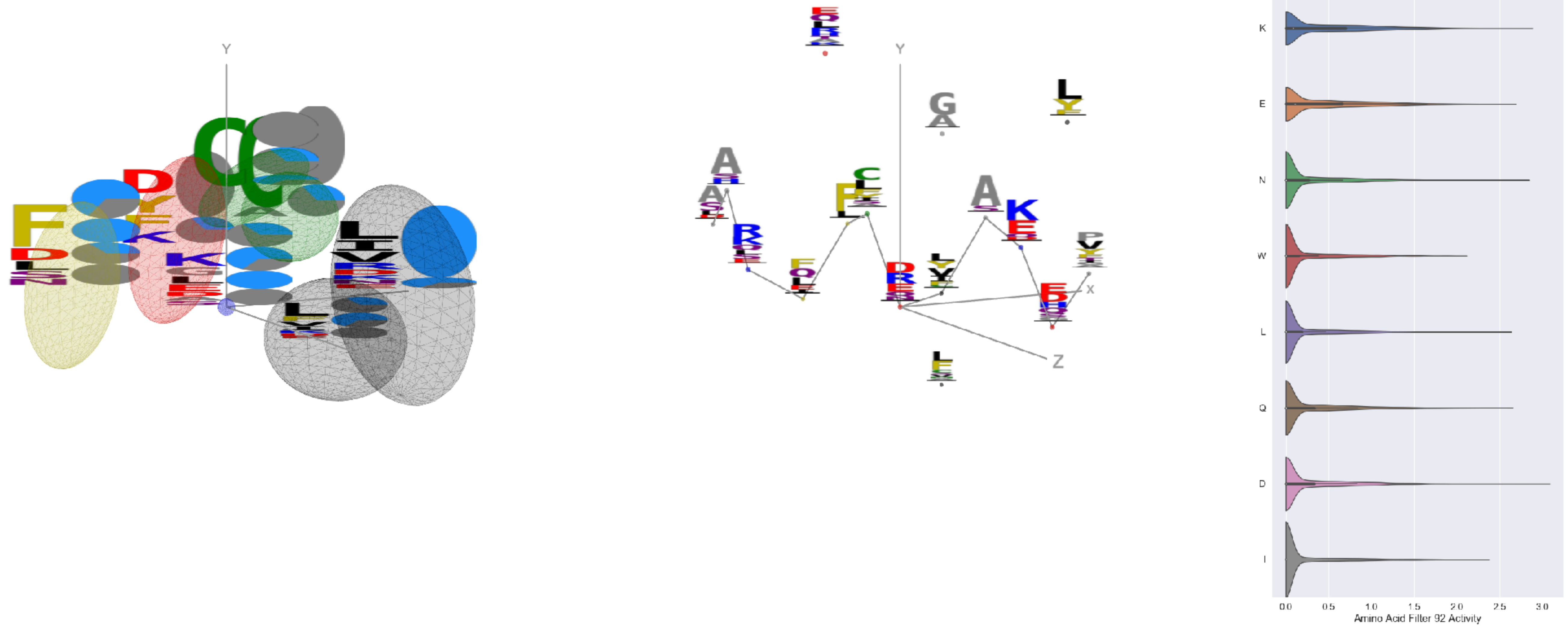


Filter 92
$R_{\mathrm{pred}} = 0.03$
Percent Active = 29
Pooling attention 0.28
Pooling feature 1.11
Top activators
4xmn:E:775 D
4uu9:D:49 K
4jr9:A:353 A
7jw0:E:751 N
6n4b:A:270 K
7c2l:C:1016 E
5i6x:A:609 Y
3iyw:A:93 K
4xmn:E:843 Q
6ln2:A:390 L

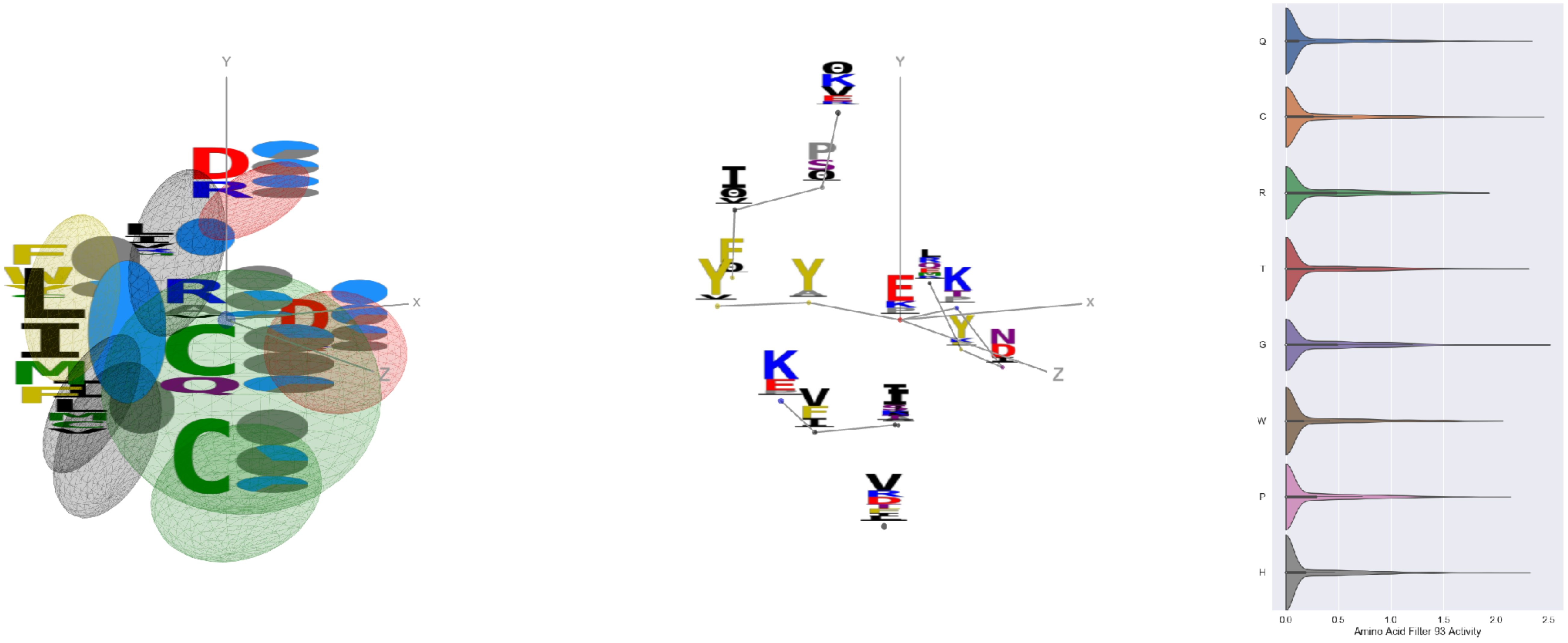


Filter 93
$R_{\mathrm{pred}} = 0.04$
Percent Active = 30
Pooling attention 0.42
Pooling feature 1.71
Top activators
7ce2:A:1127 E
5y11:C:278 G
6p95:C:70 S
6p91:A:70 S
6wpt:C:738 C
7lm9:A:378 C
6nb7:B:54 L
7cws:R:913 Q
7cwm:A:913 Q
5vod:C:90 H

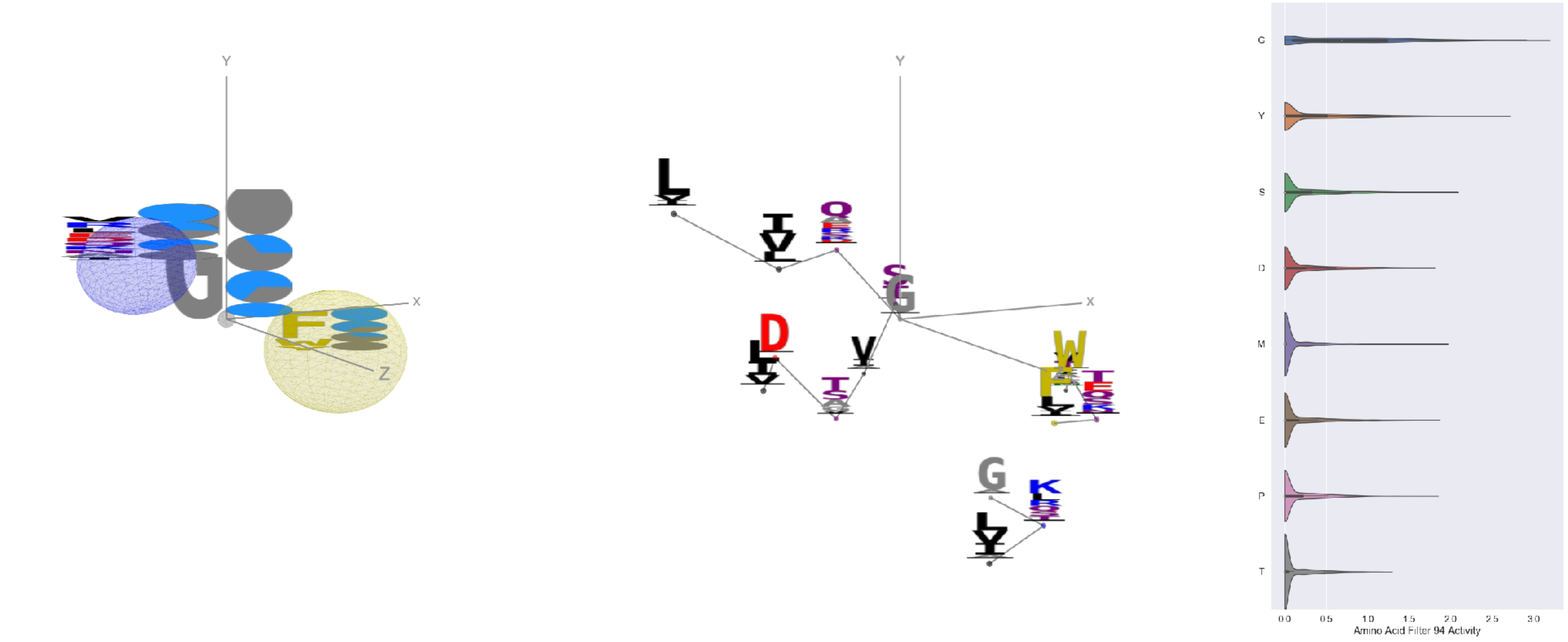


Filter 94
$R_{\mathrm{pred}} = 0.05$
Percent Active = 30
Pooling attention 0.11
Pooling feature 0.58
Top activators
7kra:A:515 G
6c9u:A:566 G
7jv2:A:496 G
6c9u:A:447 G
7l57:B:283 G
6icf:A:277 Y
6pzw:C:196 G
5jym:C:58 G
6ln2:A:1039 Y
5i5k:B:197 G

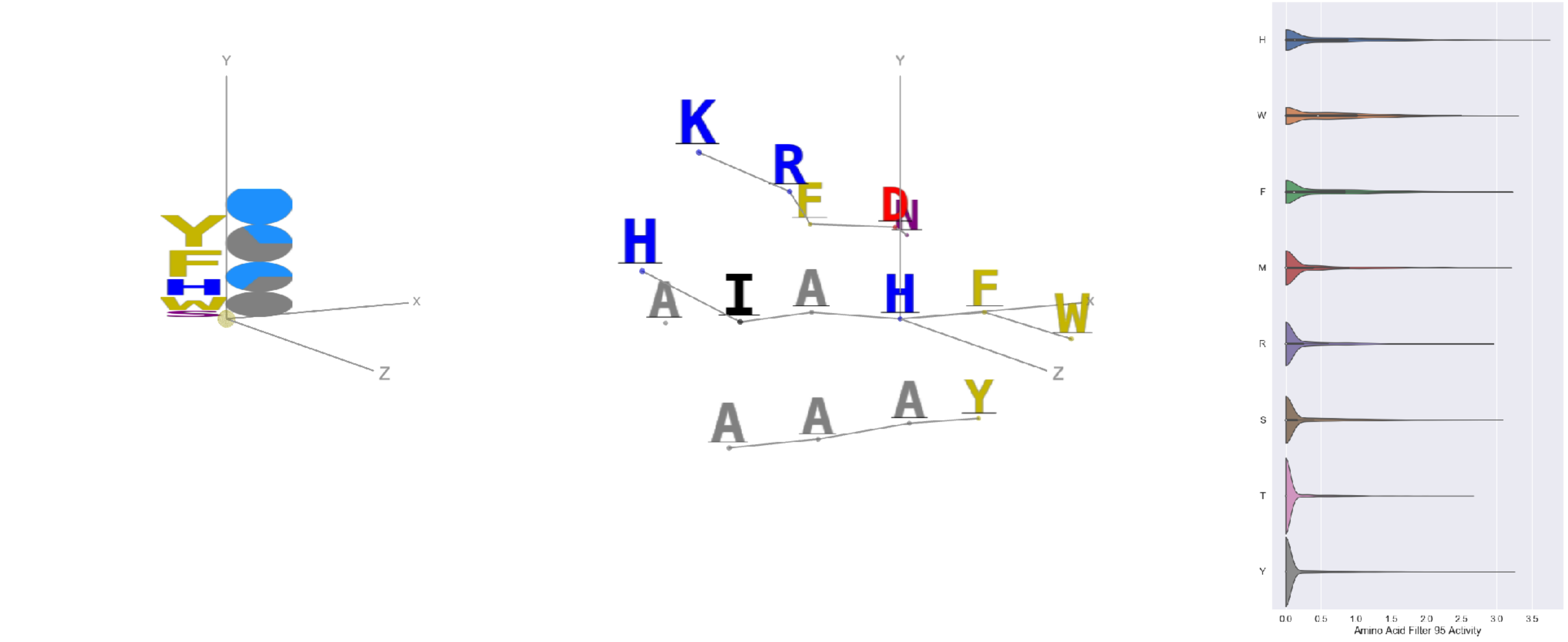


Filter 95
$R_{\mathrm{pred}} = -0.08$
Percent Active = 16
Pooling attention 0.23
Pooling feature 0.90
Top activators
7lxx:A:66 H
4cmh:A:125 W
6fgb:A:29 Y
5i5k:B:1284 F
1nca:N:241 M
7cr5:A:109 Y
6weq:B:200 Y
7ly2:A:1064 H
6o2b:M:36 W
6xkq:A:514 S

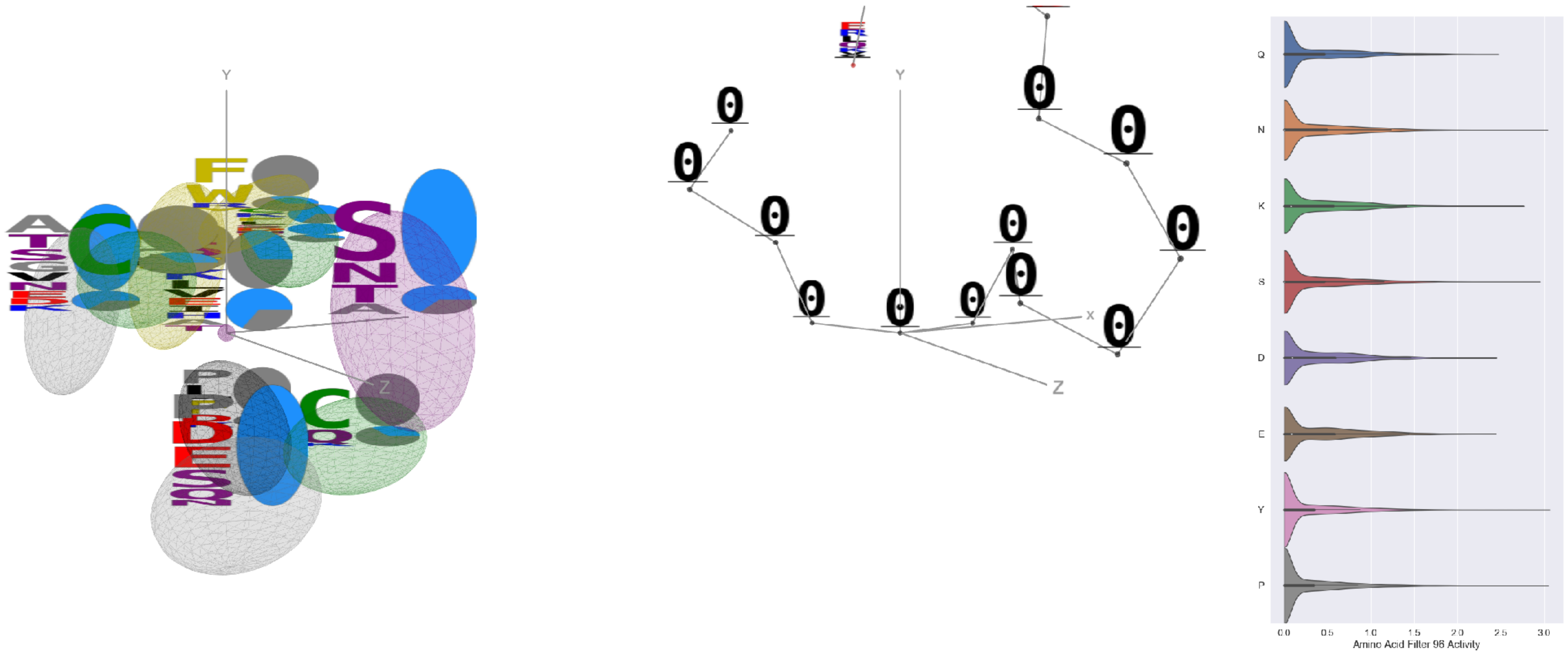

Filter 96
$R_{\text{pred}} = 0.17$
Percent Active = 36
Pooling attention 0.05
Pooling feature 0.31
Top activators
6icf:A:263 Y
6n5b:B:136 P
6ii9:C:287 N
7kq7:B:20 P
4fqv:C:296 N
6was:G:133 N
3hae:A:251 S
6mlm:A:312 N
1w72:D:251 S
6mej:C:493 P

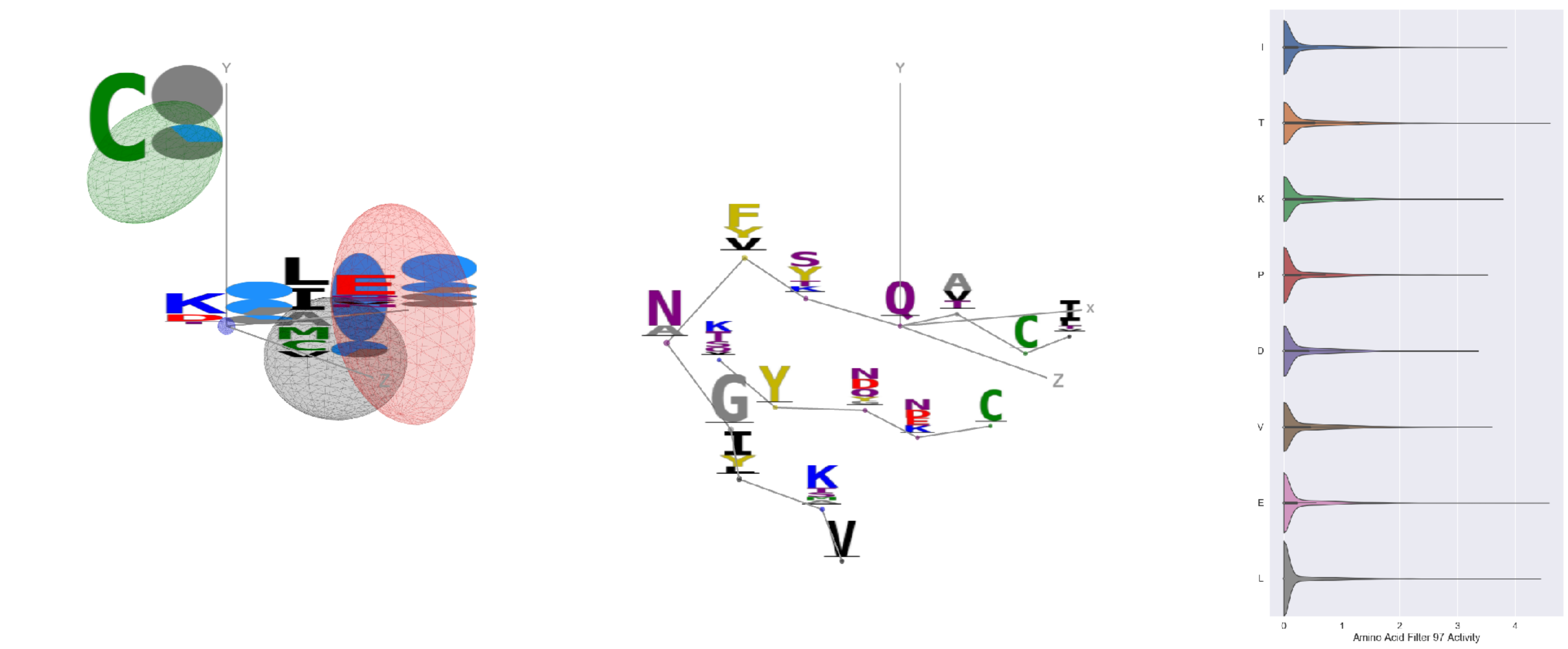

Filter 97
$R_{\text{pred}} = -0.00$
Percent Active = 30
Pooling attention 0.24
Pooling feature 0.32
Top activators
6nb7:B:835 Q
6nb6:A:835 Q
6mej:C:641 T
6usf:D:268 E
7jw0:E:434 L
7c2l:C:573 L
7l57:B:434 I
6c9u:A:390 K
6wo5:E:641 T
1qfw:B:2 Y

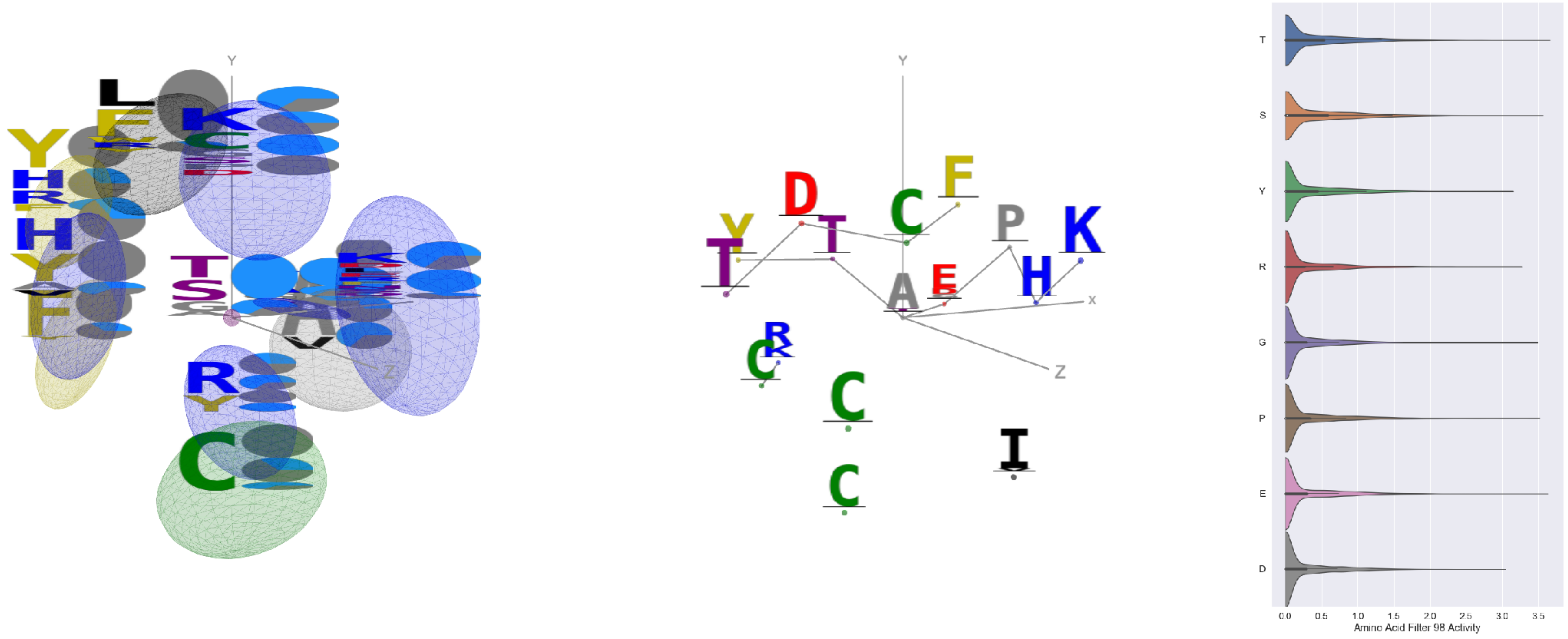

Filter 98
$R_{\text{pred}} = -0.05$
Percent Active = 35
Pooling attention 0.20
Pooling feature 1.85
Top activators
6mei:C:592 A
6wo5:E:563 T
7ce2:A:1127 E
6r0x:E:34 S
6c9u:A:808 P
6mei:C:505 P
7l57:B:1046 G
6mej:C:583 T
6mej:C:592 A
6c9u:A:435 P

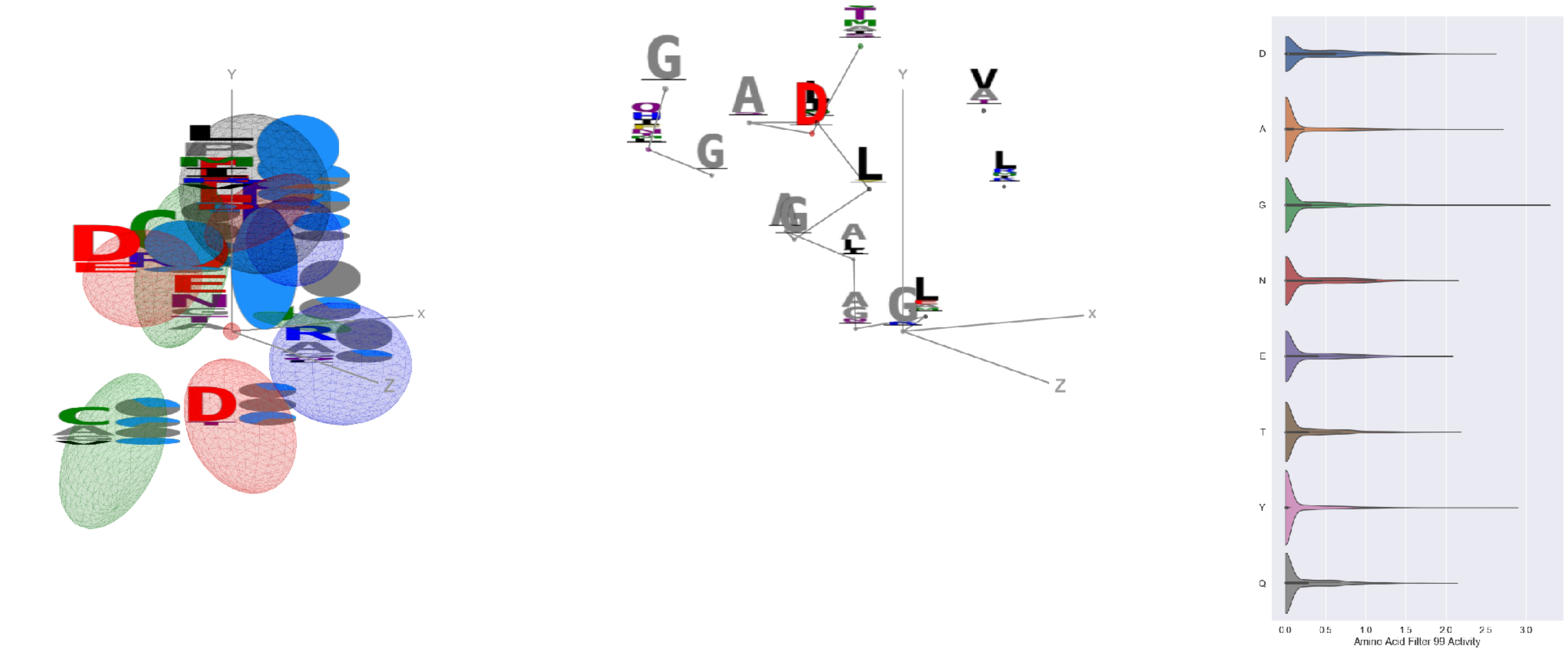

Filter 99
$R_{\text{pred}} = 0.23$
Percent Active = 24
Pooling attention 0.23
Pooling feature 0.54
Top activators
5esz:G:220 G
6wha:B:87 G
1xct:Q:45 G
4xmn:E:1036 Y
3wih:A:82 A
6vmj:Y:243 Y
3wih:A:99 Y
7cwm:A:1127 D
7ly2:A:1127 D
7cws:R:1127 D

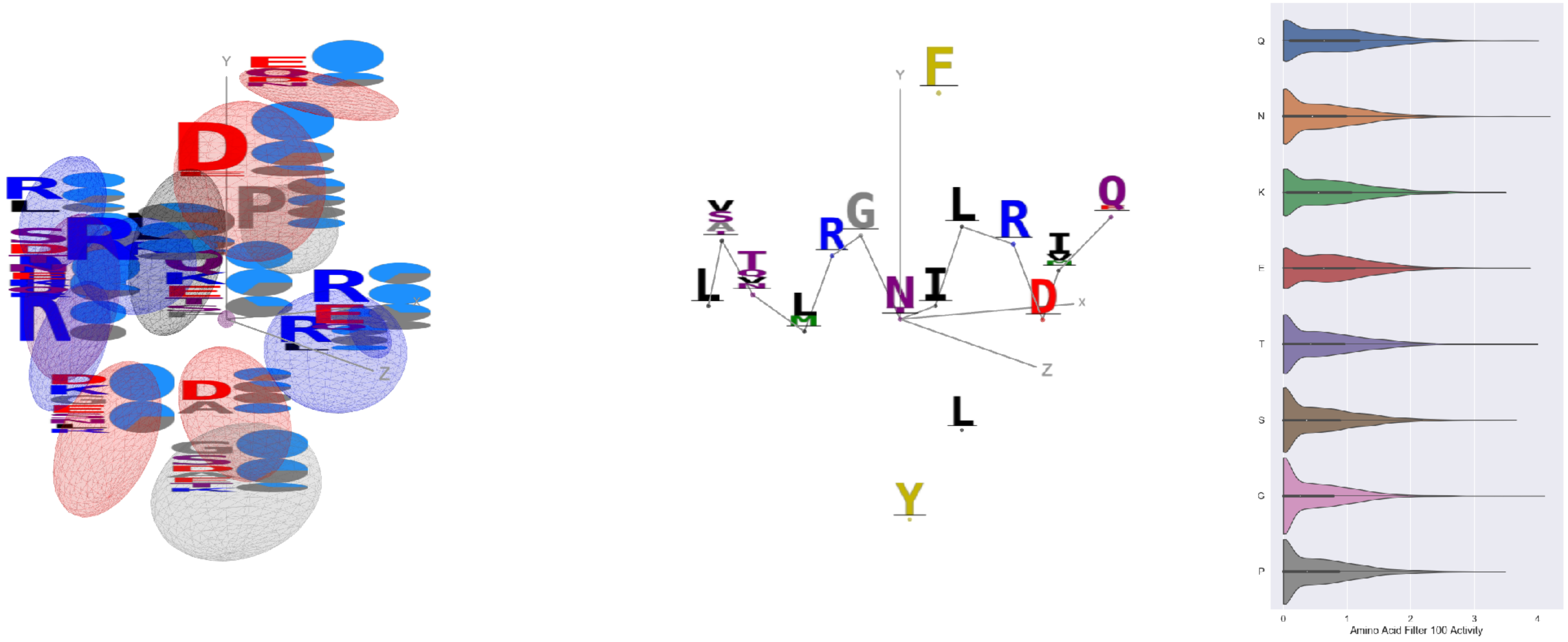


Filter 100
$R_{\text{pred}} = 0.04$
Percent Active = 49
Pooling attention 0.23
Pooling feature 1.13
Top activators
7l57:B:998 N
7l57:B:910 G
7l57:B:805 L
7l57:B:949 Q
6nb7:B:229 T
6ulf:K:69 T
7l57:B:990 E
7jw0:E:994 D
7jw0:E:990 E
7l57:B:903 Q

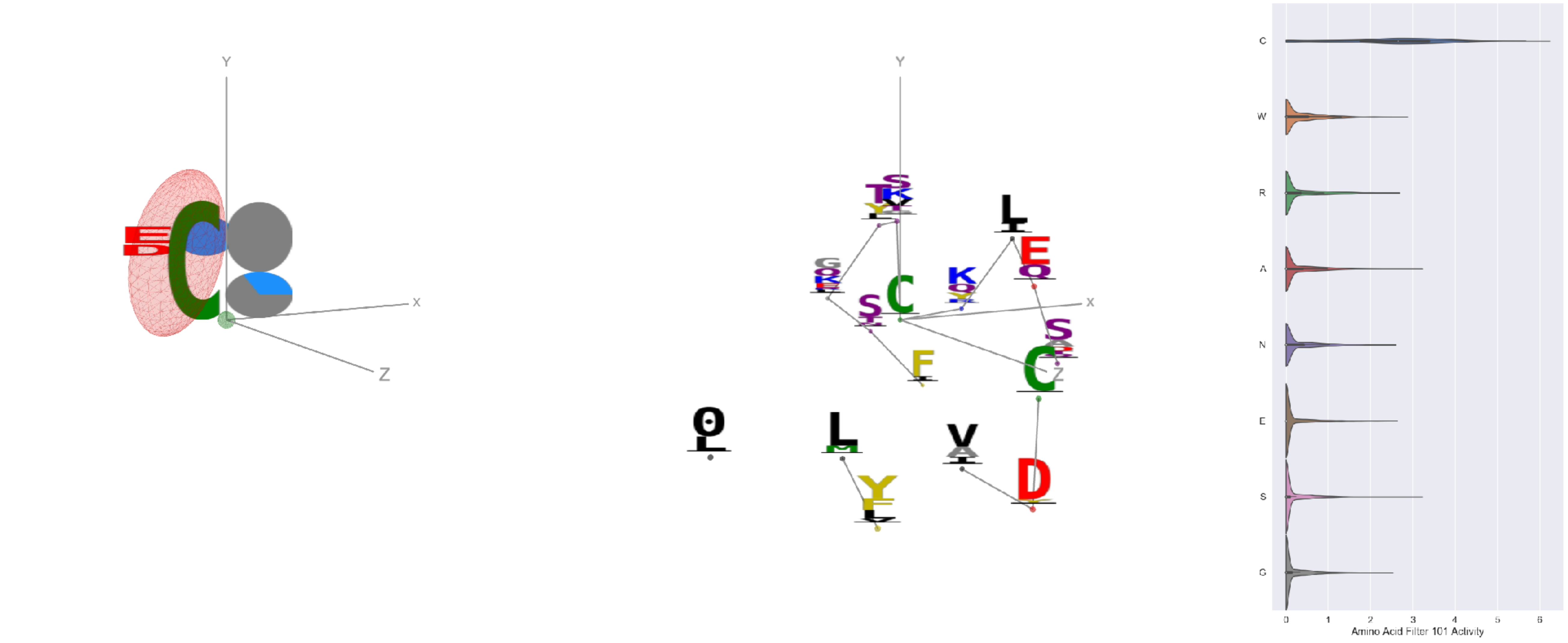


Filter 101
$R_{\text{pred}} = -0.04$
Percent Active = 23
Pooling attention 0.04
Pooling feature 0.33
Top activators
7l57:B:301 C
7l57:B:617 C
6p91:a:301 C
7l57:B:291 C
6p95:c:301 C
6mei:C:452 C
7jw0:E:760 C
7l57:B:649 C
4i3r:G:296 C
7jw0:E:749 C

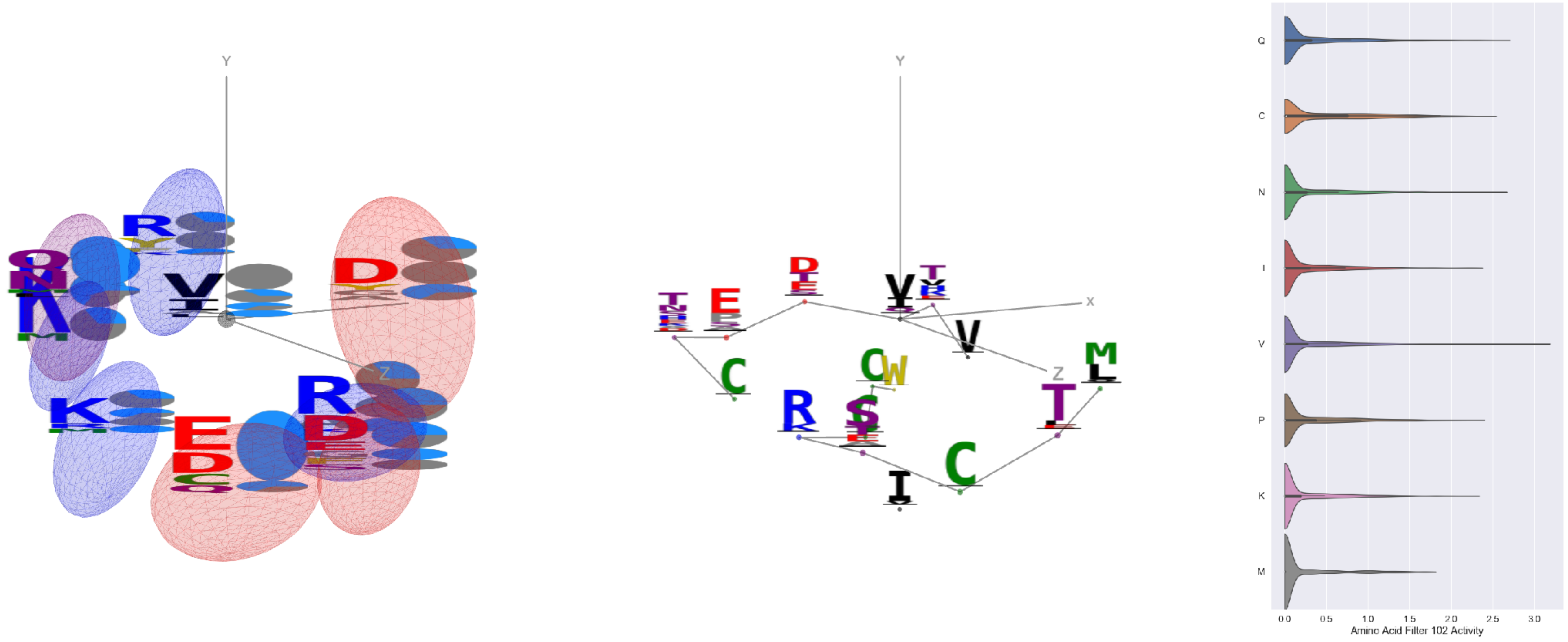


Filter 102
$R_{\text{pred}} = 0.04$
Percent Active = 28
Pooling attention 0.56
Pooling feature 2.57
Top activators
6weq:B:287 V
7k93:C:287 V
7nd9:B:820 D
7ly2:A:1023 Q
7nda:A:764 N
6mei:C:557 S
7l57:B:989 A
7jw0:E:15 Y
7nda:A:1023 Q
7jw0:E:525 C

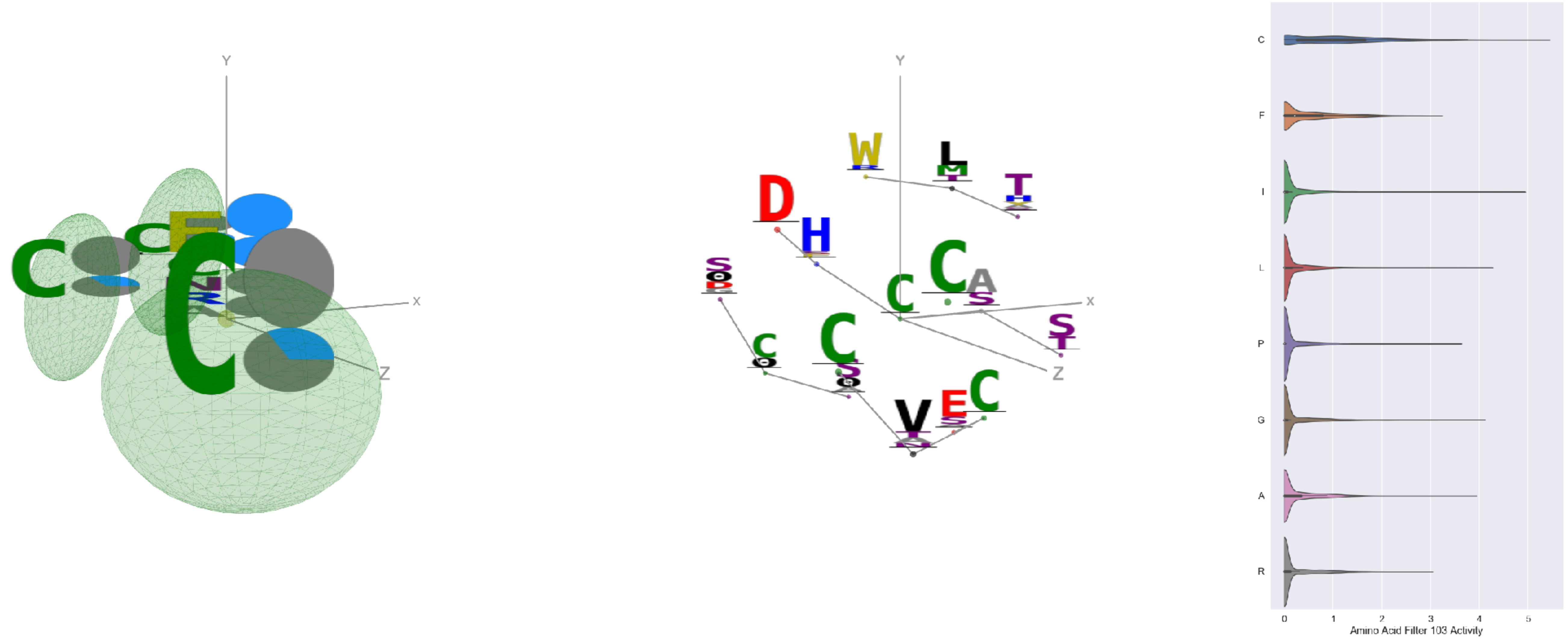


Filter 103
$R_{\text{pred}} = -0.13$
Percent Active = 19
Pooling attention 0.36
Pooling feature 0.71
Top activators
6q23:A:183 C
6pzw:C:183 C
6umx:A:281 C
7jw0:E:977 I
1nca:N:183 C
4aei:A:22 C
1qfw:B:9 C
6nb7:B:959 I
5bo1:A:251 C
7cws:R:977 I

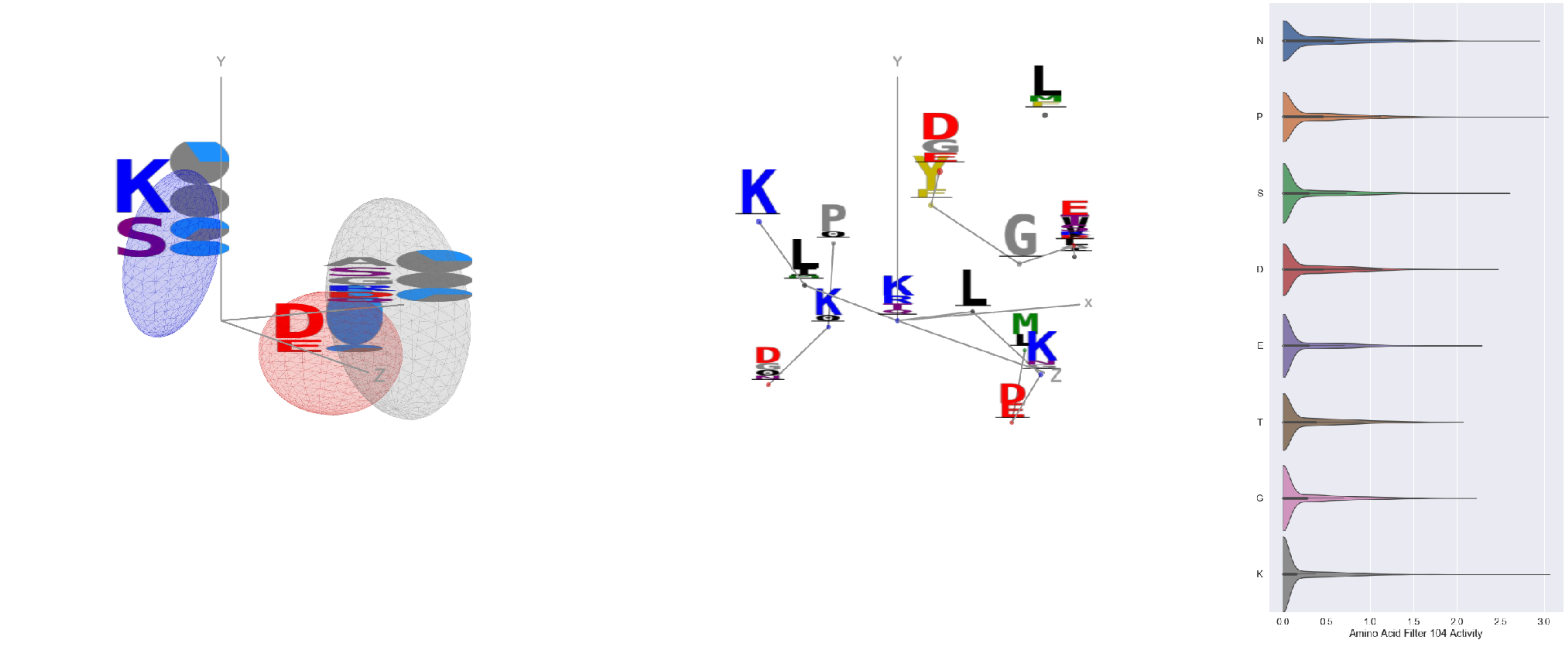


Filter 104
$R_{\text{pred}} = 0.23$
Percent Active = 29
Pooling attention 0.09
Pooling feature 0.40
Top activators
3iyw:A:296 K
7d77:B:65 P
6rps:B:62 N
7jvr:B:65 P
6qno:B:65 P
5d72:B:27 N
6lfo:B:65 P
5f3j:A:334 K
6mej:C:466 R
6wpt:C:420 S

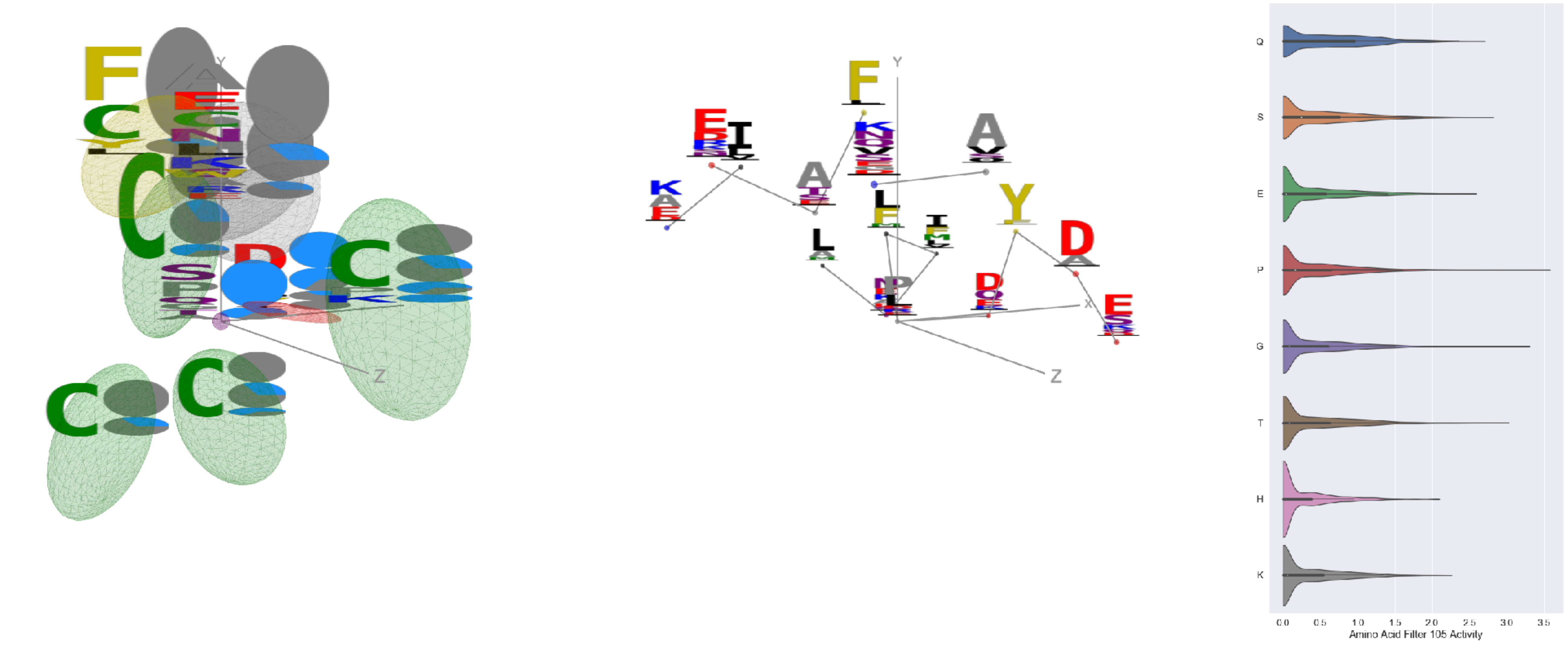


Filter 105
$R_{\text{pred}} = 0.09$
Percent Active = 42
Pooling attention 0.16
Pooling feature 0.26
Top activators
4cmh:A:108 P
7jw0:E:381 G
7jv6:E:381 G
4uao:A:112 T
7nda:A:744 G
6c9u:A:446 S
6mjz:B:185 P
1nca:N:385 D
5vkd:A:659 Q
6mlm:A:108 N

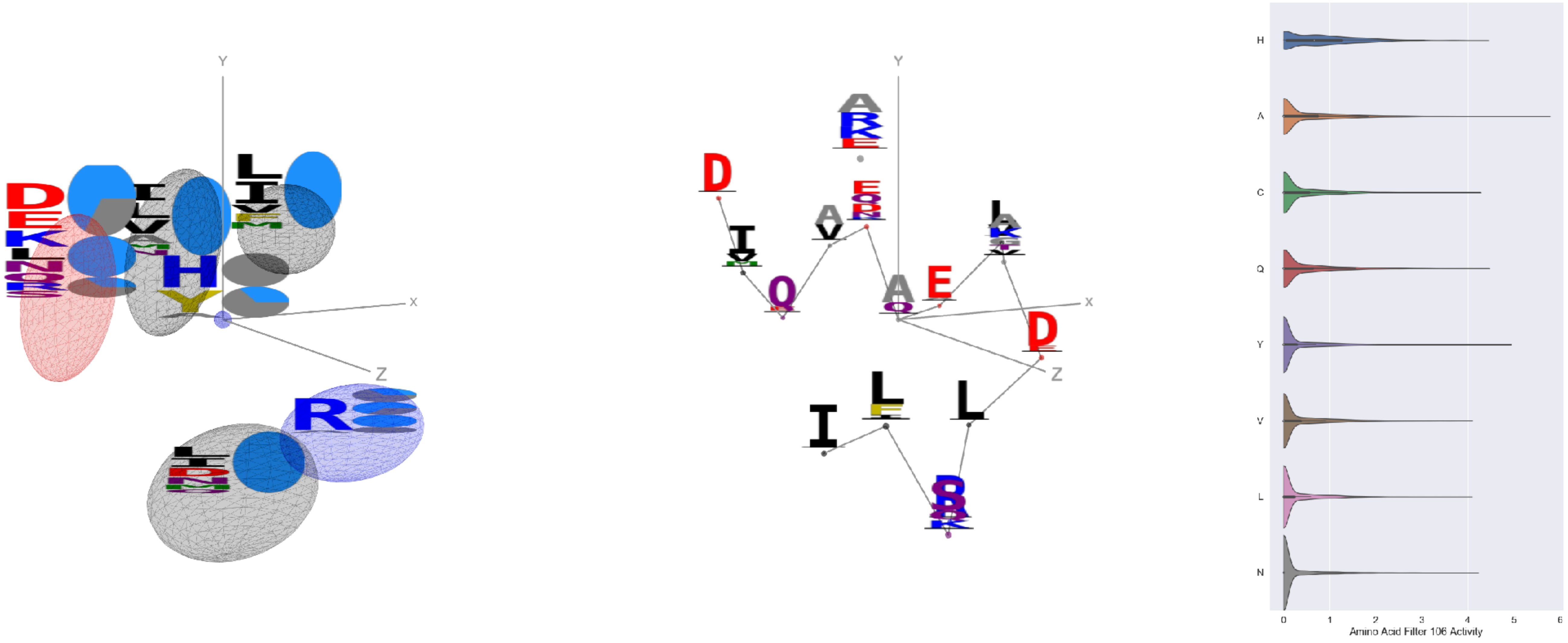


Filter 106
$R_{\text{pred}} = -0.16$
Percent Active = 30
Pooling attention 0.44
Pooling feature 0.29
Top activators
7jw0:E:989 A
7l57:B:989 A
6nb7:B:971 A
7l57:B:924 A
6wha:C:16 Y
7jw0:E:750 R
5h35:D:191 I
3tt3:A:504 Y
6nb7:B:732 R
7jw0:E:819 E

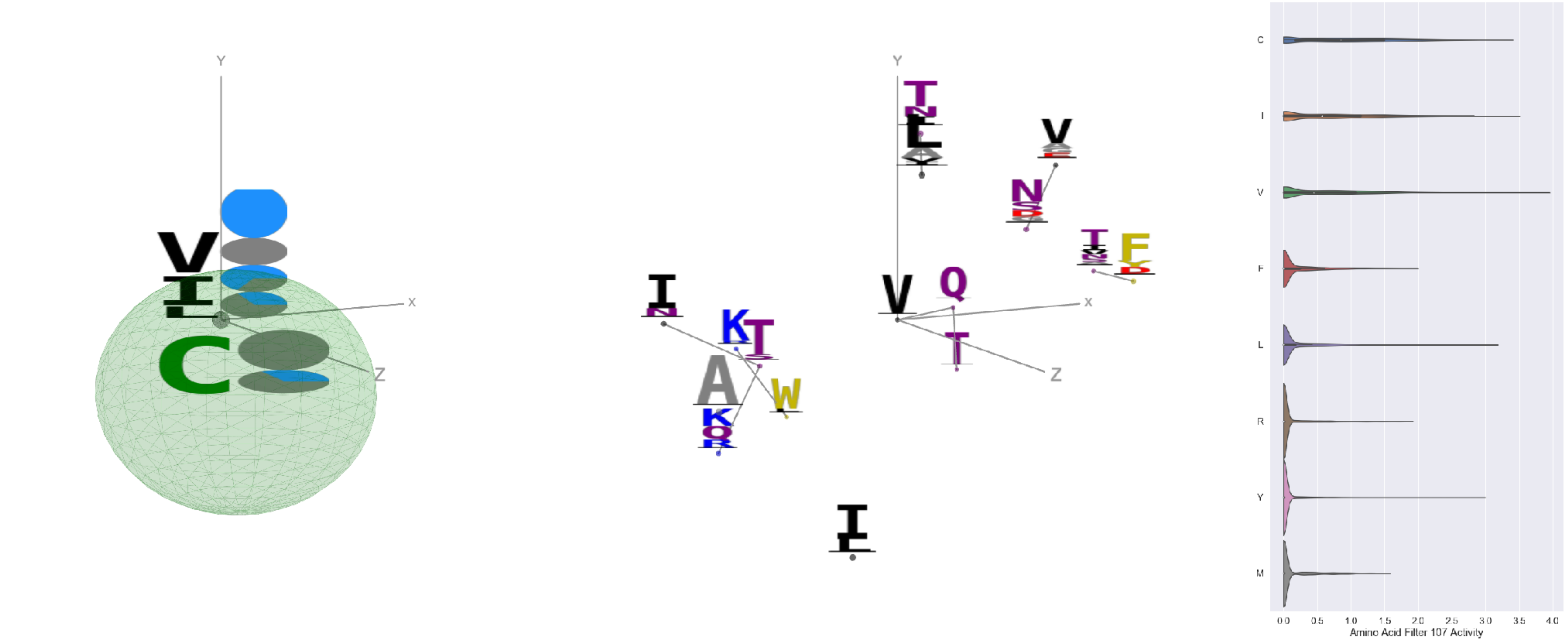


Filter 107
$R_{\text{pred}} = -0.17$
Percent Active = 20
Pooling attention 0.01
Pooling feature 0.58
Top activators
3hae:C:9 V
6nb6:A:808 V
7jw0:E:826 V
7jw0:E:620 V
6aod:C:536 I
7ly2:A:826 V
4aei:A:22 C
6nha:B:37 V
7jv6:E:977 I
6nb6:C:808 V

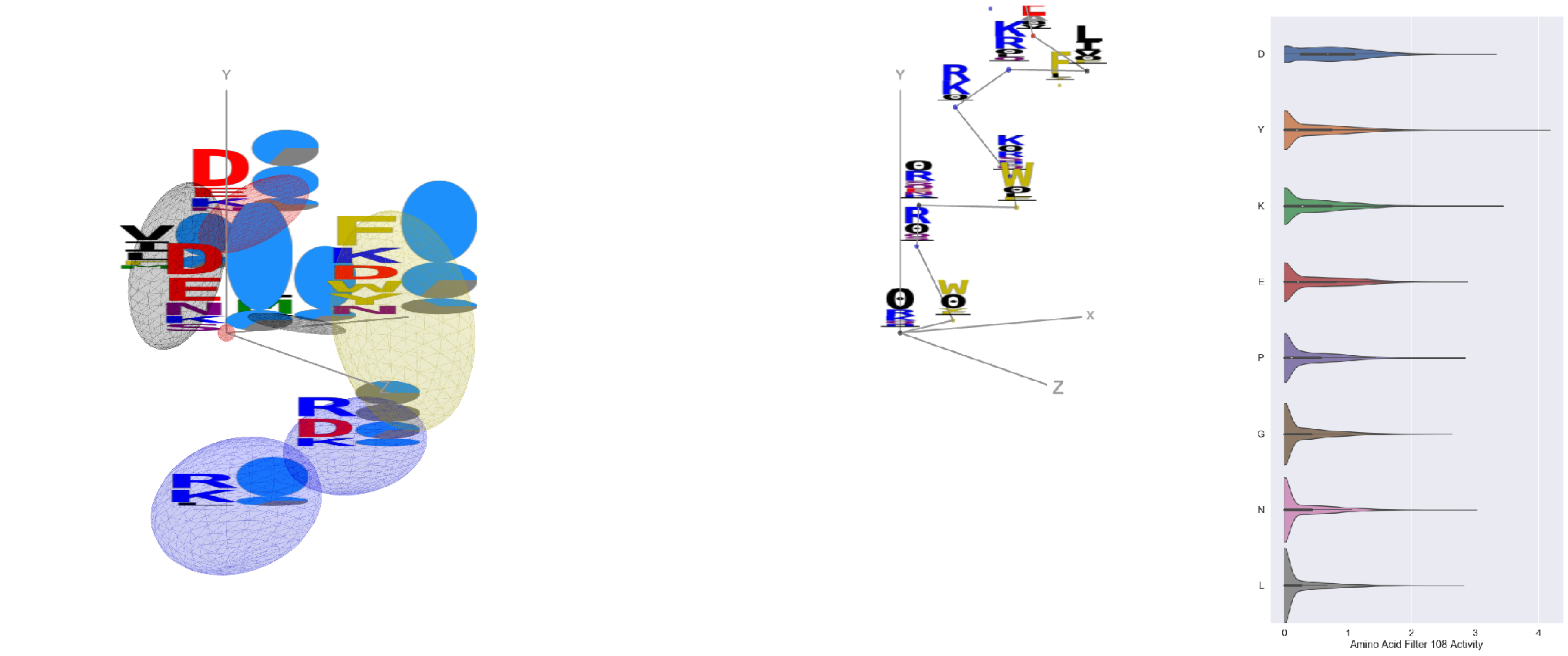


Filter 108
$R_{pred} = 0.11$
Percent Active = 45
Pooling attention 0.20
Pooling feature 0.95
Top activators
6ln2:A:421 Y
4xmn:E:879 F
5otj:C:71 Y
7ly2:A:1145 Y
4xmn:E:726 R
5h35:D:195 W
5esz:C:316 Y
6wha:B:51 K
7l57:B:580 Y
7kr5:E:216 I

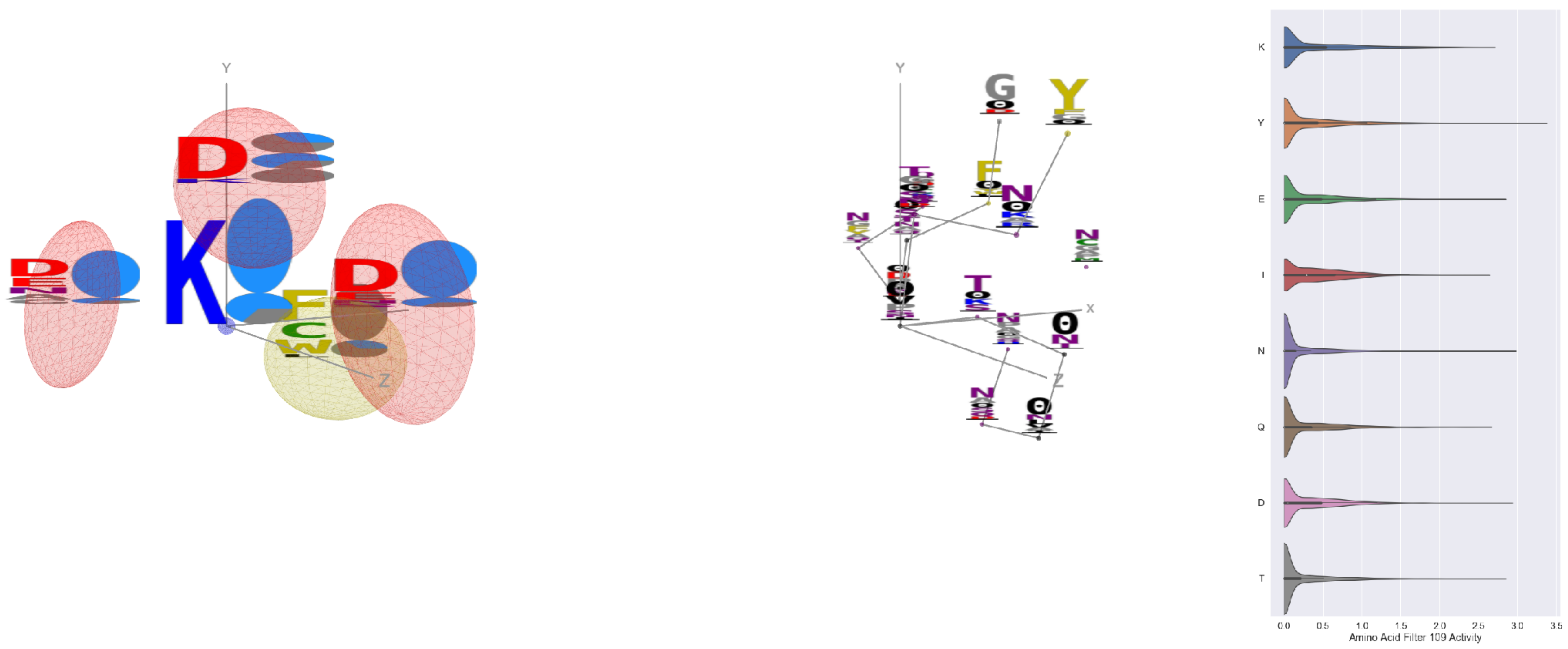


Filter 109
$R_{pred} = 0.06$
Percent Active = 40
Pooling attention 0.13
Pooling feature 0.30
Top activators
7lcn:A:444 Y
7jv6:E:444 Y
6c9u:A:460 A
4dw2:U:217 N
4zfg:A:289 Y
5bk2:B:251 D
4dw2:U:243 Y
5i5k:B:1409 T
4i3r:G:290 E
6yio:B:148 L

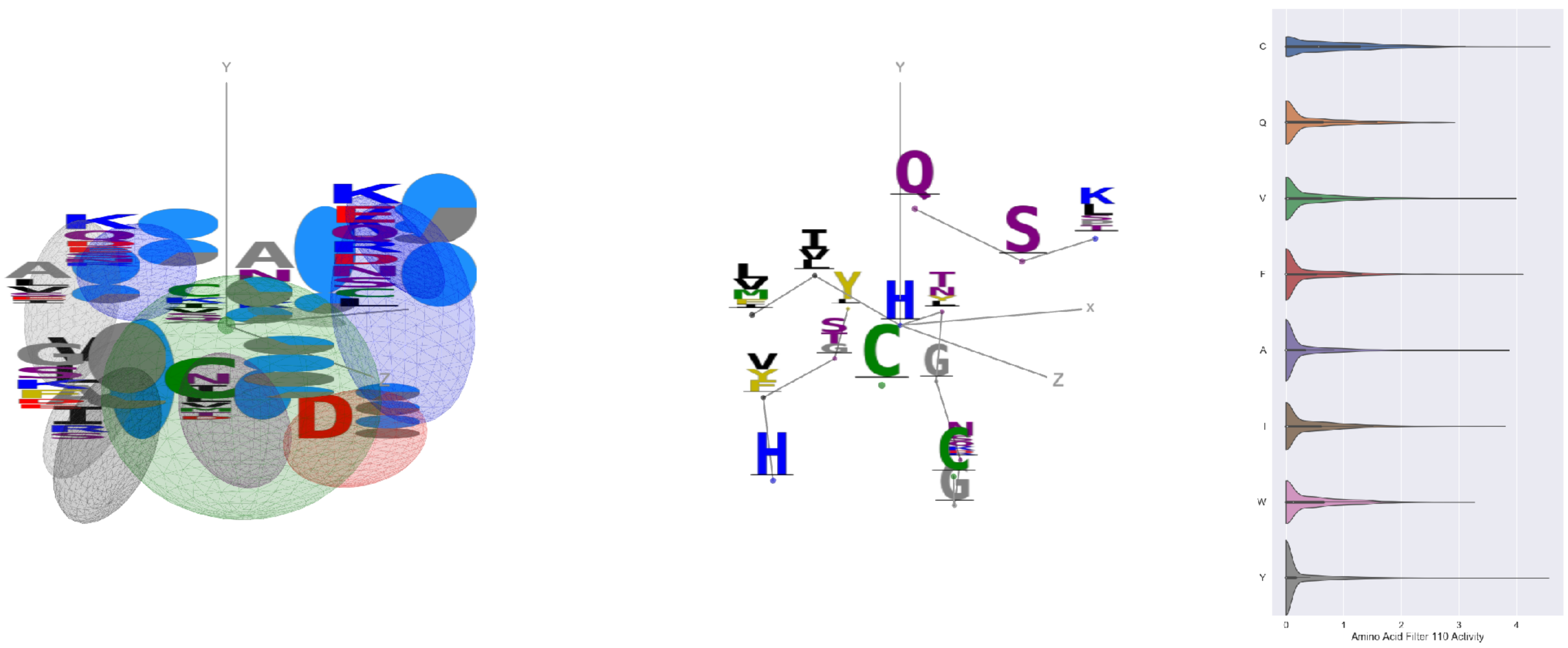


Filter 110
$R_{pred} = -0.12$
Percent Active = 33
Pooling attention 0.05
Pooling feature 0.06
Top activators
7l57:B:1048 H
4xmn:E:919 L
6nb7:B:348 C
7l57:B:240 Y
7l57:B:121 Y
5esz:G:155 R
7lxx:A:133 F
6umx:A:281 C
6nb6:C:236 Y
6nb6:C:361 V

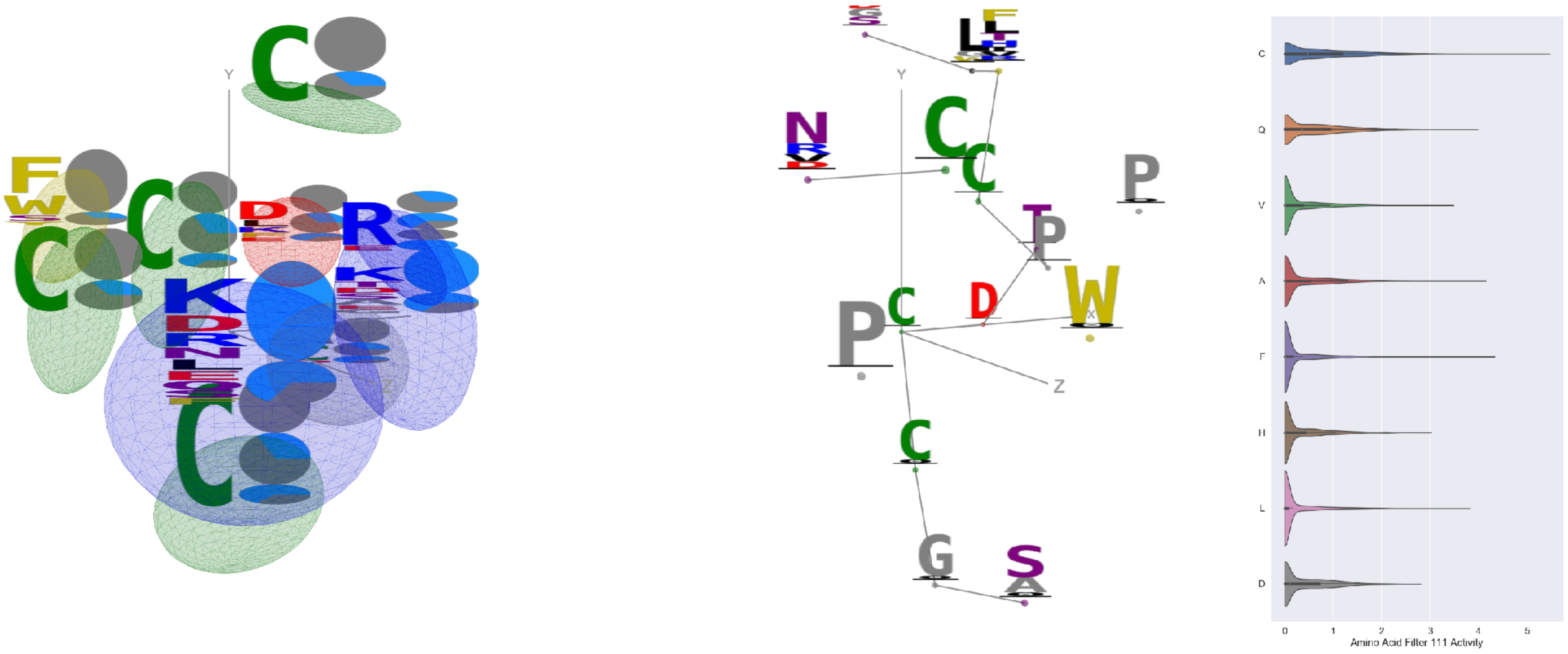


Filter 111
$R_{pred} = 0.01$
Percent Active = 36
Pooling attention 0.55
Pooling feature 0.11
Top activators
4mwf:D:585 C
1oaz:A:49 C
4xmn:E:999 C
5tlj:X:67 C
7l57:B:617 C
4xmn:E:927 F
7l57:B:930 A
4xmn:E:887 C
7l57:B:997 I
4xmn:E:841 A

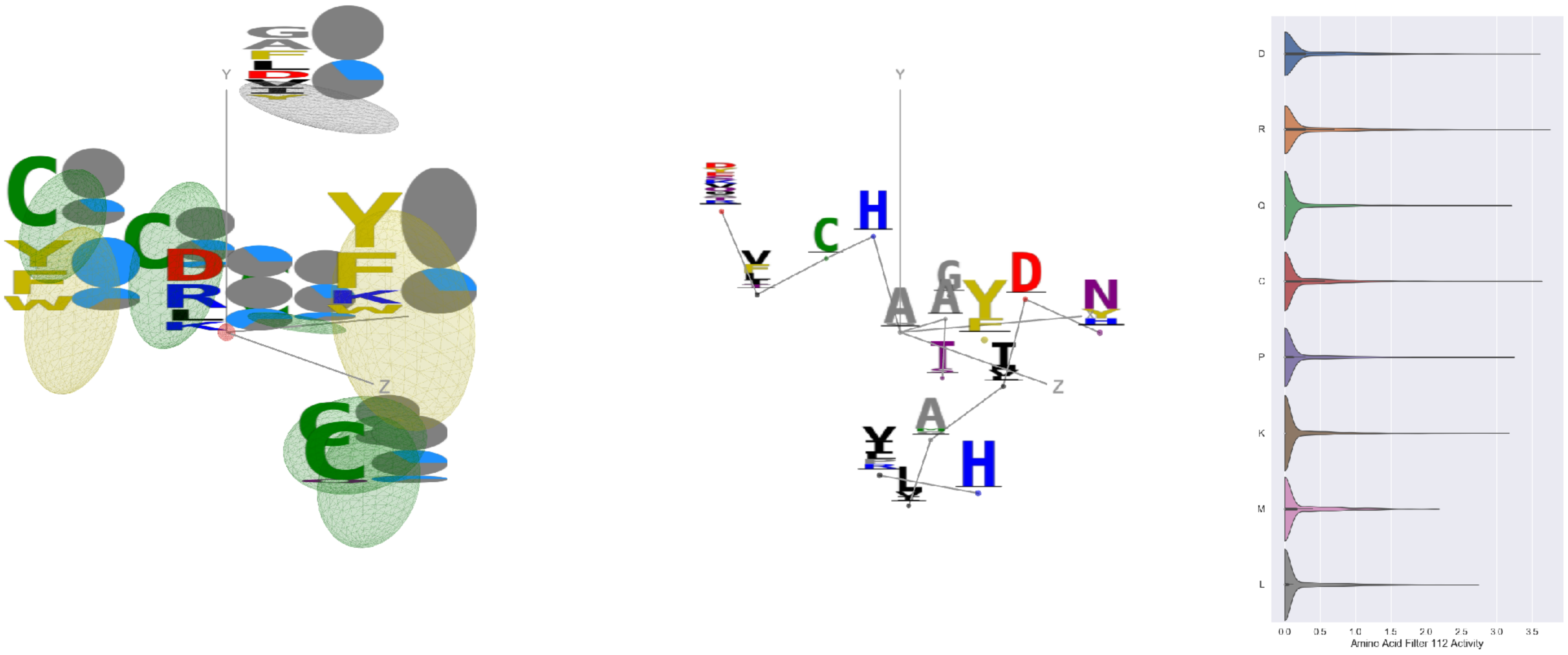


Filter 112
$R_{pred} = -0.02$
Percent Active = 21
Pooling attention 0.29
Pooling feature 2.08
Top activators
6aod:C:412 A
6weq:B:336 R
6aod:C:414 C
6pt0:B:205 D
6lfo:B:333 D
7jvr:B:205 D
1kb5:A:33 S
6ln2:A:246 V
5jym:C:103 D
7c80:A:213 P

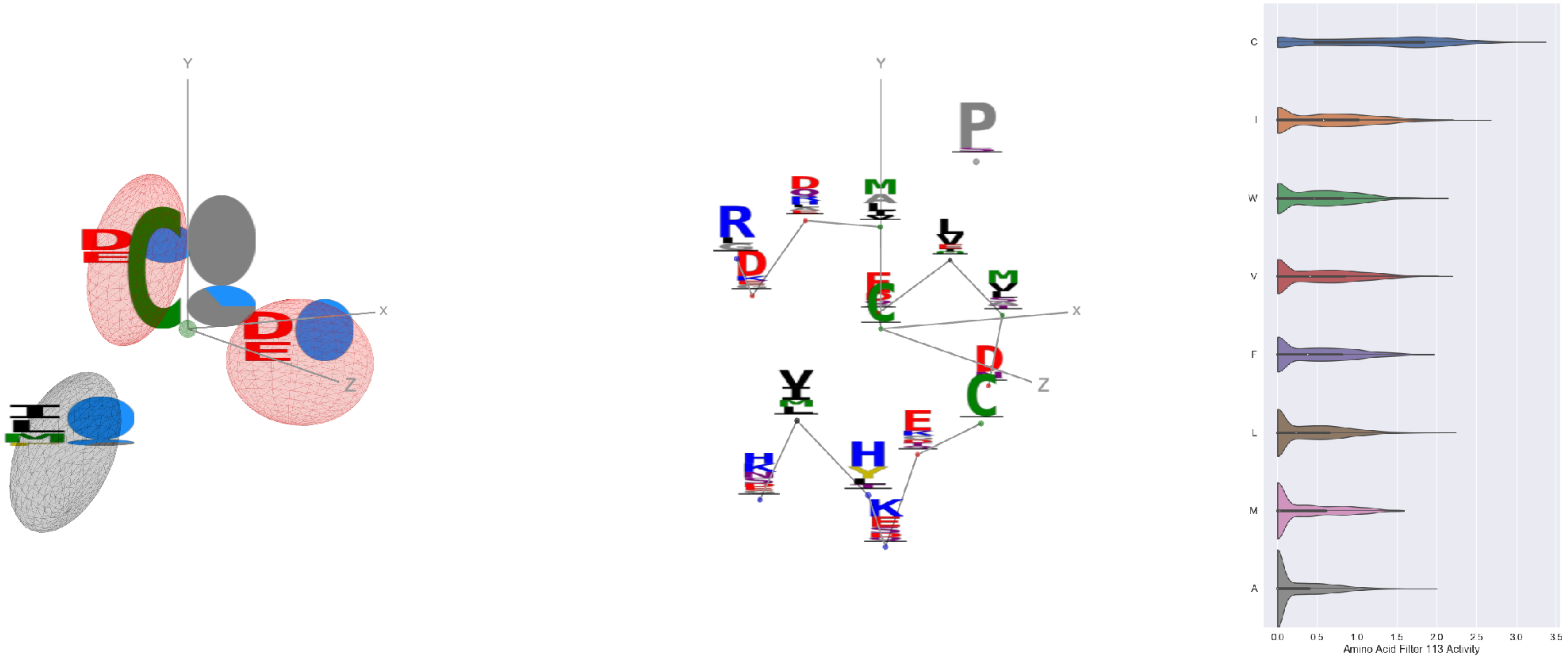


Filter 113
$R_{pred} = -0.27$
Percent Active = 36
Pooling attention 0.03
Pooling feature 0.47
Top activators
6m58:A:253 C
6weq:B:312 C
6mei:C:569 C
6weq:B:280 C
4cmh:A:173 C
7k93:C:312 C
7k93:C:280 C
6m58:A:124 C
6m58:A:278 C
7nda:C:1032 C

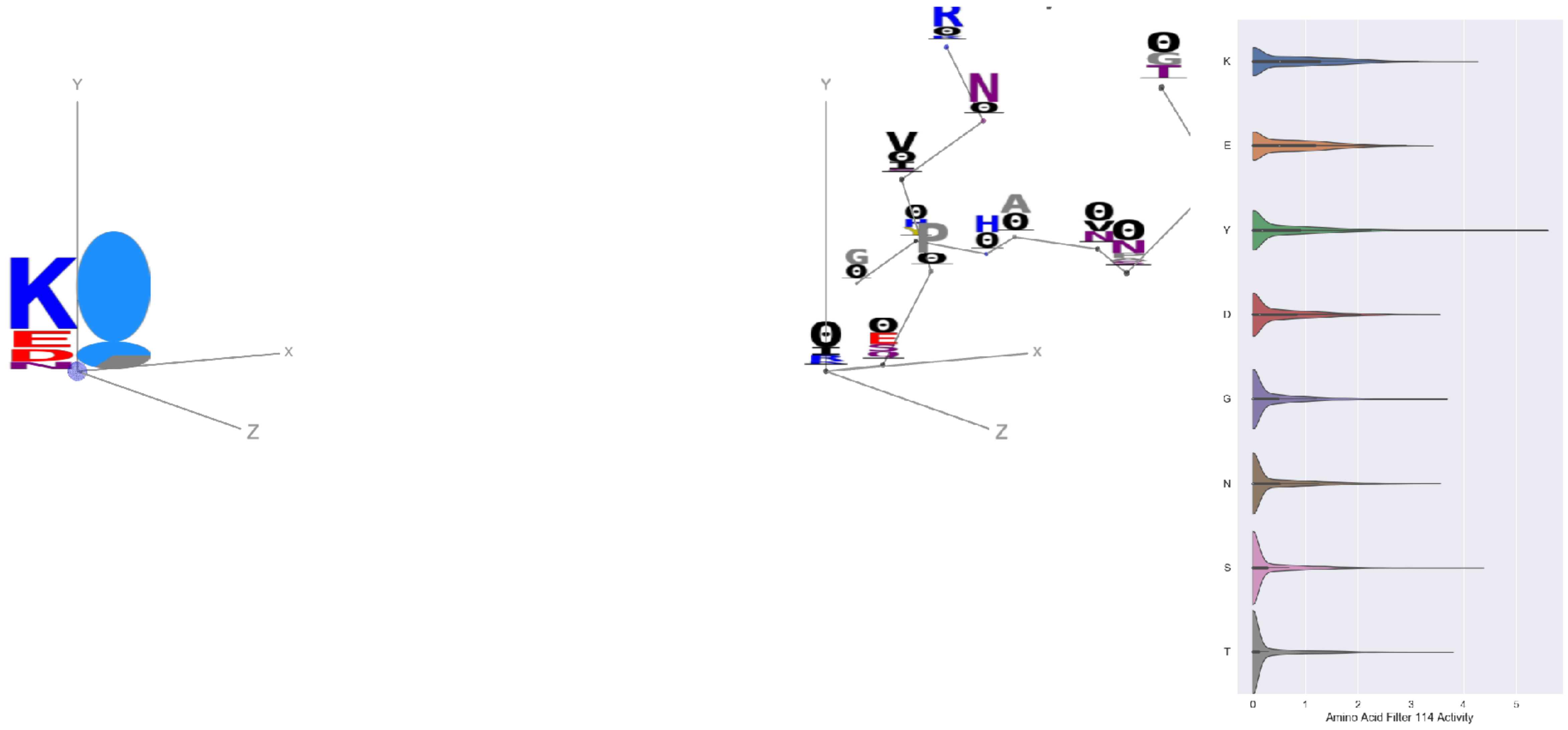


Filter 114
$R_{pred} = 0.14$
Percent Active = 43
Pooling attention 0.40
Pooling feature 1.14
Top activators
6ii9:C:317 Y
5bk2:B:219 S
6wtv:D:445 K
5tq0:A:296 S
5f3j:A:428 K
3uze:D:327 T
5bk2:B:313 Y
7kra:A:331 Y
5n0a:A:88 T
4jr9:A:386 Y

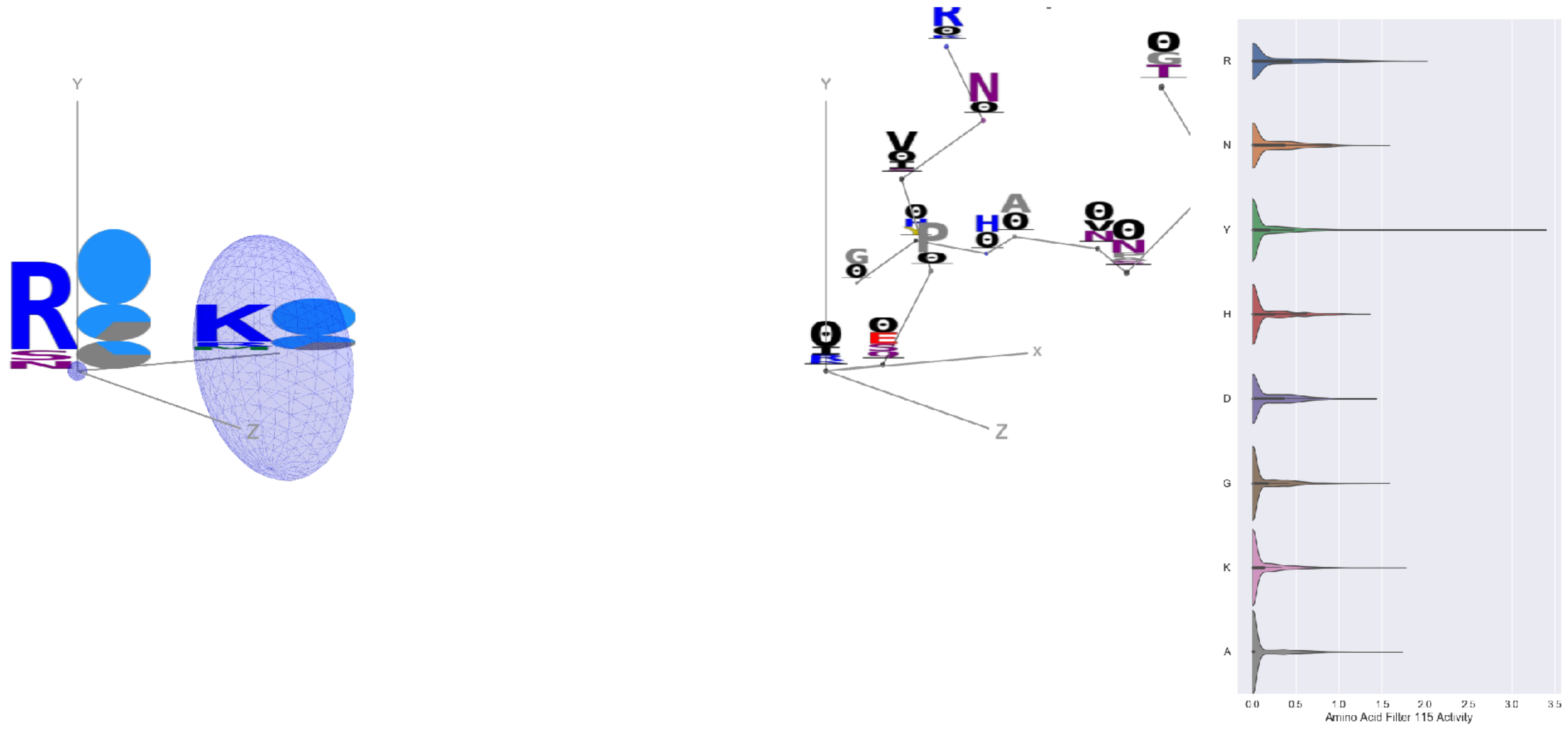


Filter 115
$R_{pred} = 0.22$
Percent Active = 29
Pooling attention 0.22
Pooling feature 0.35
Top activators
6ii9:C:317 Y
1qfw:B:112 Y
6p95:C:256 R
7l2d:A:1147 Y
7ly2:A:1145 Y
5esv:E:139 Y
6nb7:B:1127 Y
3tt3:A:511 Y
7nd9:B:1147 Y
7jw0:E:1147 Y

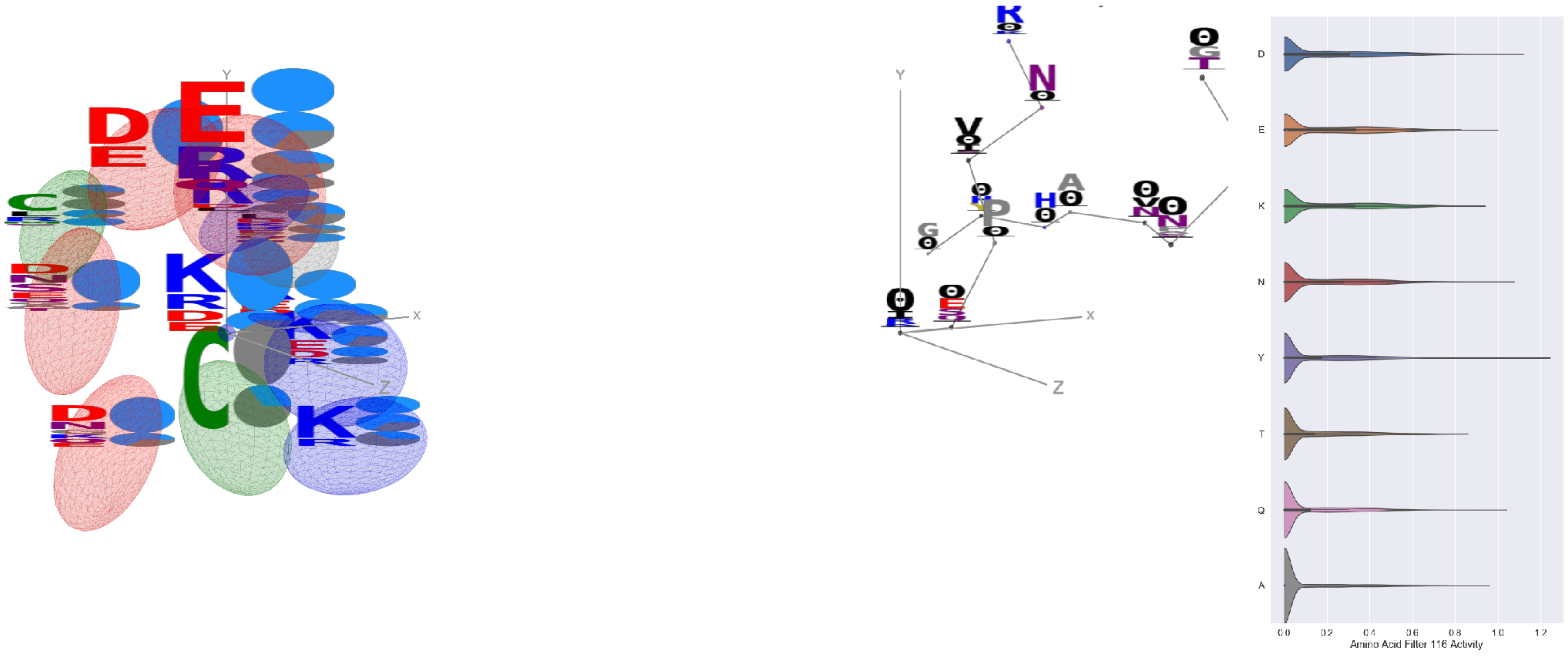

Filter 116
$R_{\text{pred}} = 0.23$
Percent Active = 27
Pooling attention 0.70
Pooling feature 3.70
Top activators
6ii9:C:317 Y
4xmn:E:874 D
6wtv:D:220 N
4xmn:E:947 Q
4xmn:E:863 R
4xmn:E:850 E
4xmn:E:775 D
4xmn:E:771 E
4xmn:E:813 S
5t5n:C:202 D

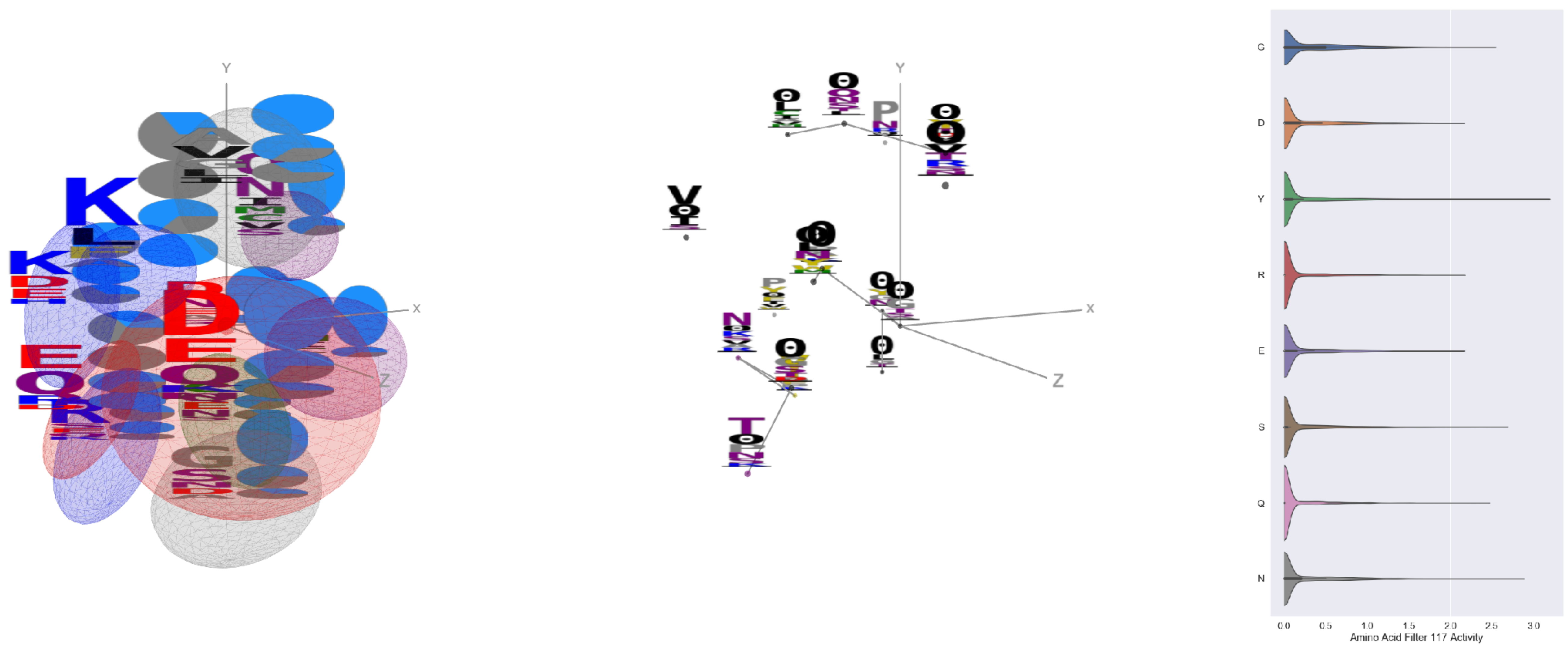

Filter 117
$R_{\text{pred}} = 0.16$
Percent Active = 23
Pooling attention 0.66
Pooling feature 3.89
Top activators
6nb6:C:482 Y
7ce2:A:1234 Y
7ce2:A:988 N
7l57:B:482 Y
3nfp:I:19 S
6n4b:B:97 S
7lcn:A:138 Y
3w9e:C:43 Y
5tq2:B:94 K
7jw0:E:496 G

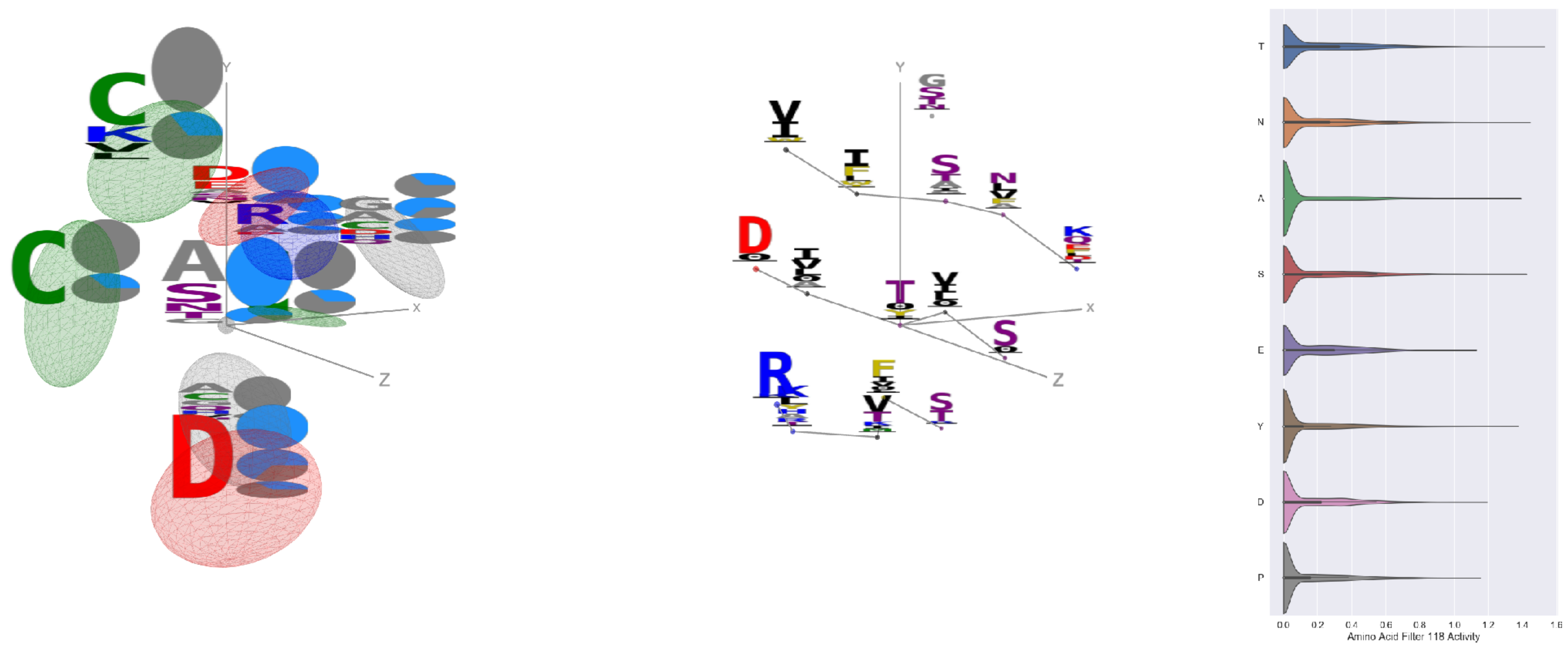

Filter 118
$R_{\text{pred}} = 0.14$
Percent Active = 27
Pooling attention 0.69
Pooling feature 2.59
Top activators
6nb6:C:383 T
6nb7:B:732 R
7jw0:E:540 N
7l57:B:278 S
6nb7:B:383 T
7l57:B:739 A
7l57:B:274 T
7l57:B:202 Y
7jw0:E:542 N
4xmn:E:772 K

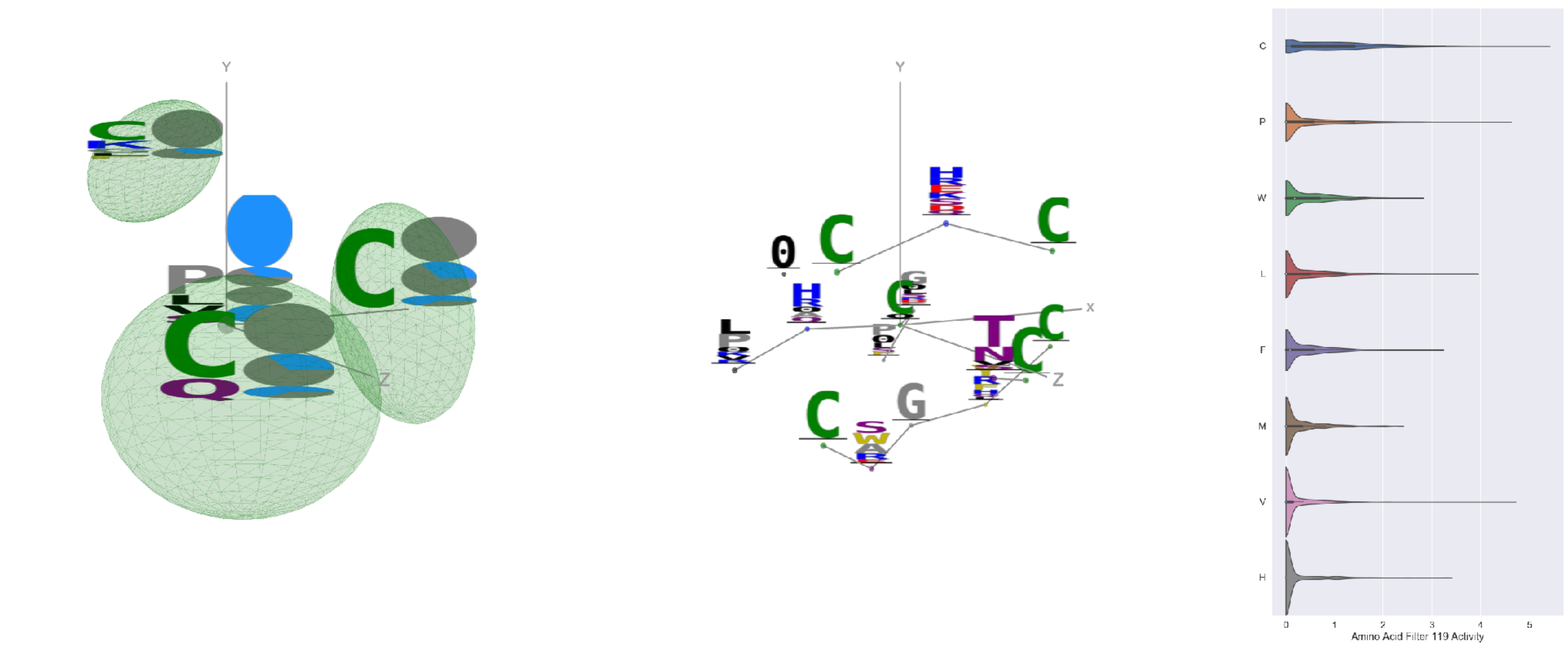

Filter 119
$R_{\text{pred}} = -0.15$
Percent Active = 24
Pooling attention 0.19
Pooling feature 0.48
Top activators
1qfw:B:9 C
7jw0:E:620 V
4kxz:E:48 C
6o25:D:12 P
6wpt:C:620 V
6umx:A:281 C
4zxb:E:595 P
7lcn:A:620 V
3eo1:I:48 C
6pzw:C:183 C

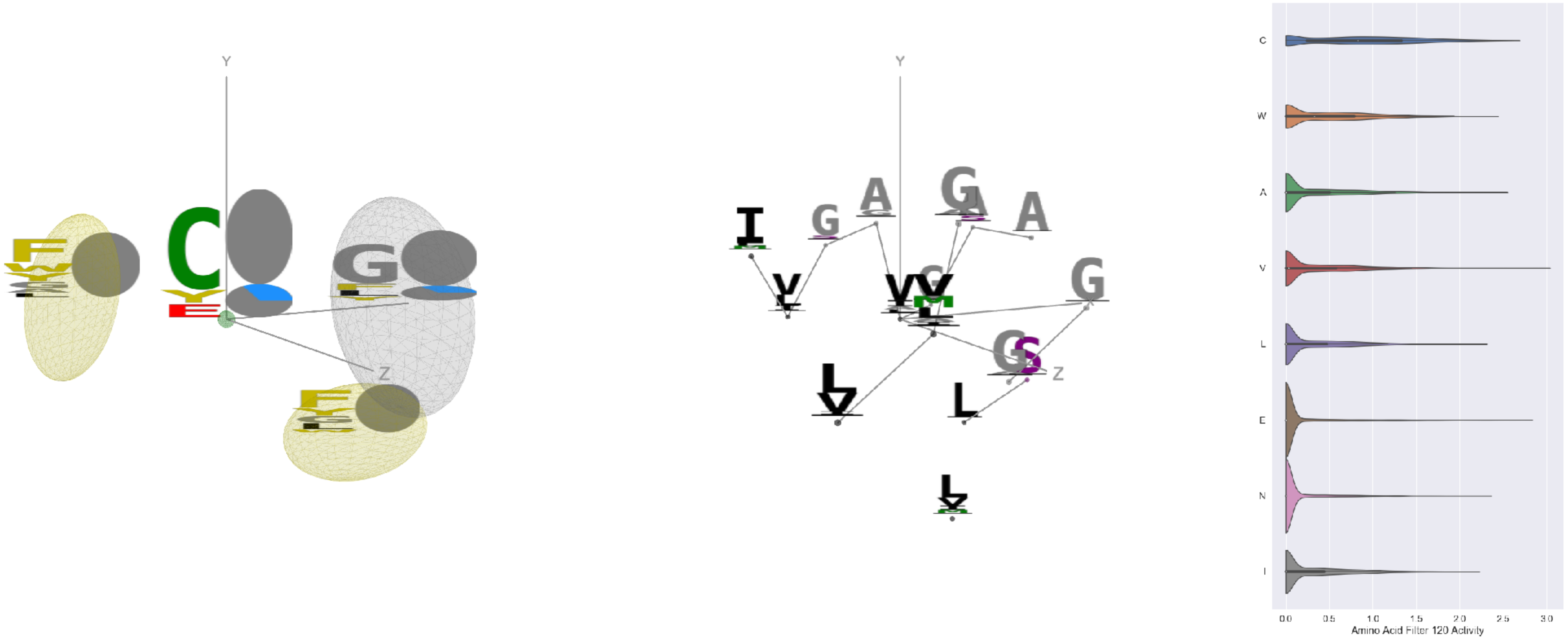


Filter 120
$R_{\mathrm{pred}} = -0.28$
Percent Active = 30
Pooling attention 0.18
Pooling feature 0.75
Top activators
6c9u:A:385 V
5tr1:B:166 E
6nb6:A:576 C
7lcn:A:525 C
6mej:C:494 C
6n5b:B:93 C
6mei:C:494 C
5i6x:A:136 E
7c2l:C:749 C
3w9e:C:118 C

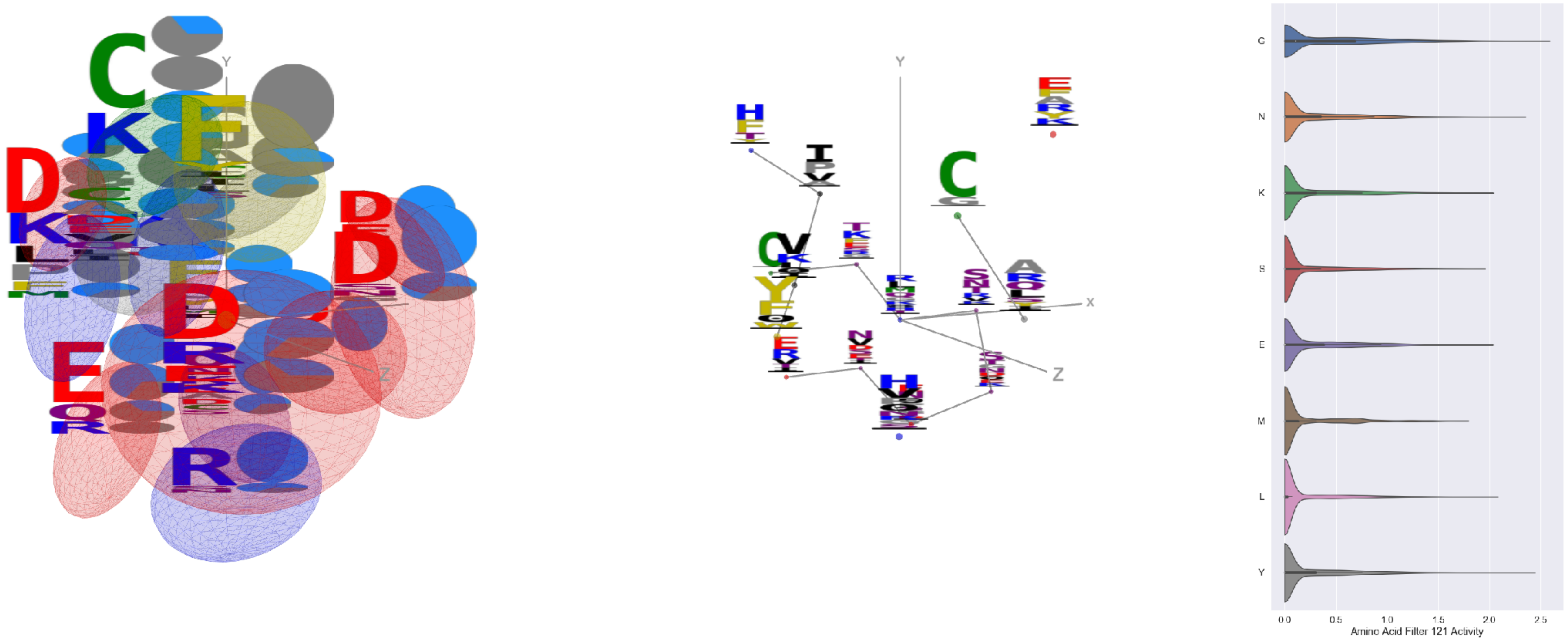


Filter 121
$R_{\mathrm{pred}} = 0.12$
Percent Active = 32
Pooling attention 0.35
Pooling feature 3.04
Top activators
5kqv:E:299 R
7l57:B:648 G
6pt0:A:240 V
6r0x:E:114 D
3skj:E:90 T
7ce2:A:1234 Y
7nd9:B:1119 N
7nda:C:1119 N
7nda:A:1119 N
5y11:C:291 G

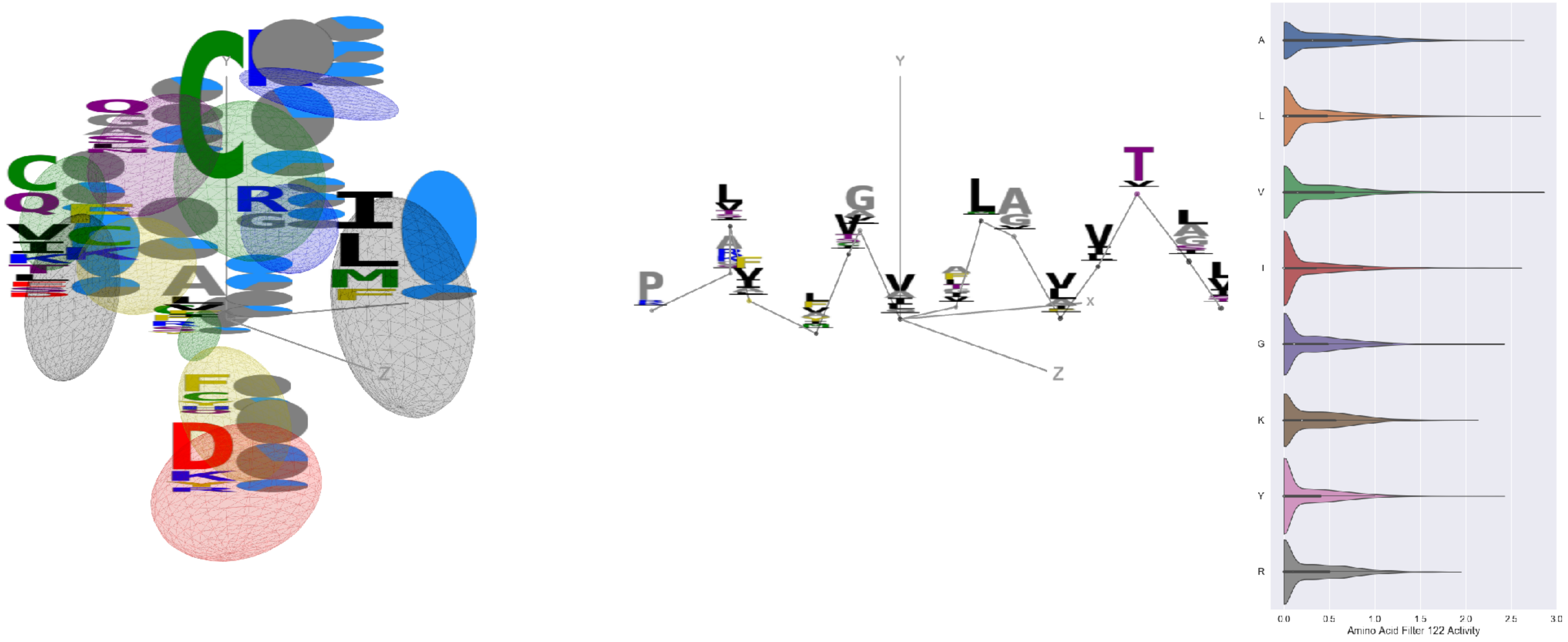


Filter 122
$R_{\mathrm{pred}} = -0.01$
Percent Active = 44
Pooling attention 0.19
Pooling feature 0.30
Top activators
7kra:A:739 V
4jr9:A:447 L
7l2d:A:1056 A
7nda:A:1056 A
5i6x:A:554 I
7lcn:A:1056 A
7l2d:A:766 A
7kra:A:736 A
7l2e:A:1056 A
3ncy:A:150 V

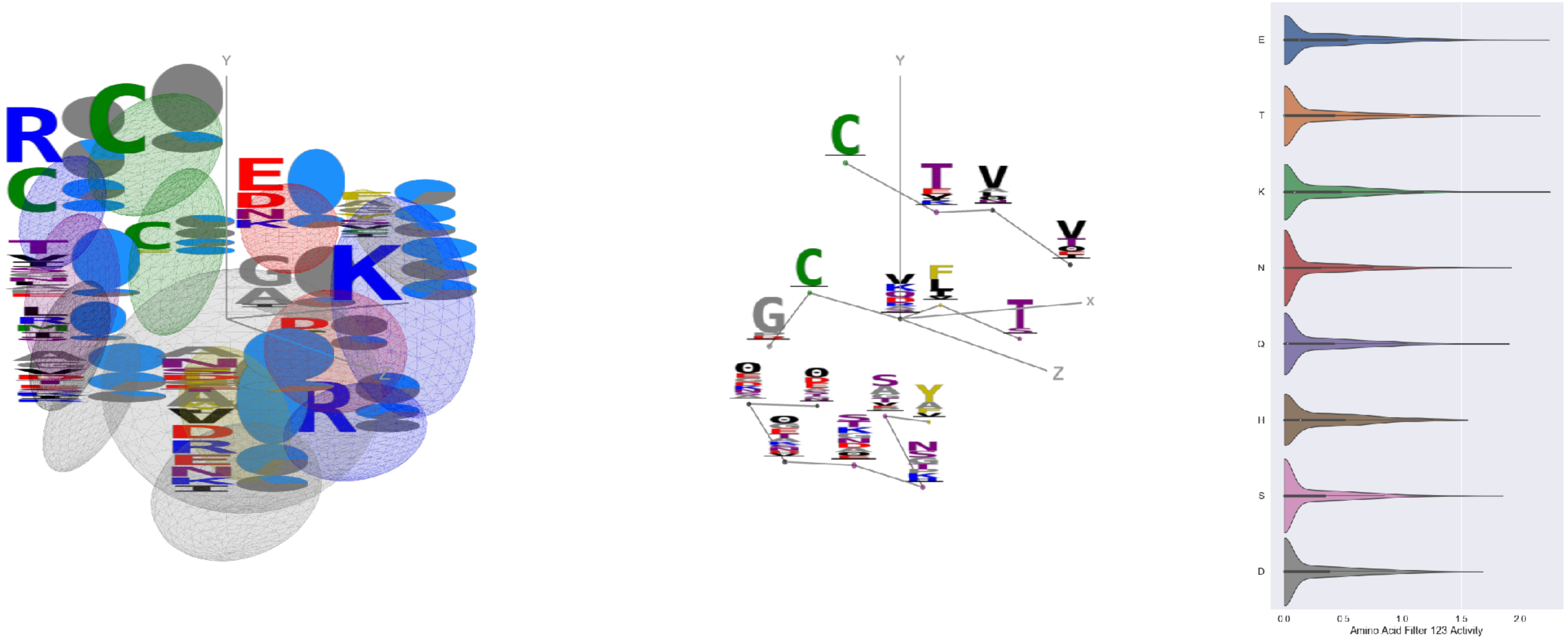


Filter 123
$R_{\mathrm{pred}} = 0.03$
Percent Active = 36
Pooling attention 0.37
Pooling feature 1.35
Top activators
1ynt:G:3166 V
4xmn:E:356 K
7cah:A:516 E
1ynt:G:3033 K
7bud:B:62 E
1nca:N:104 Y
7lcn:A:696 T
6yio:B:54 T
4zxb:E:232 Y
1jrh:I:43 T

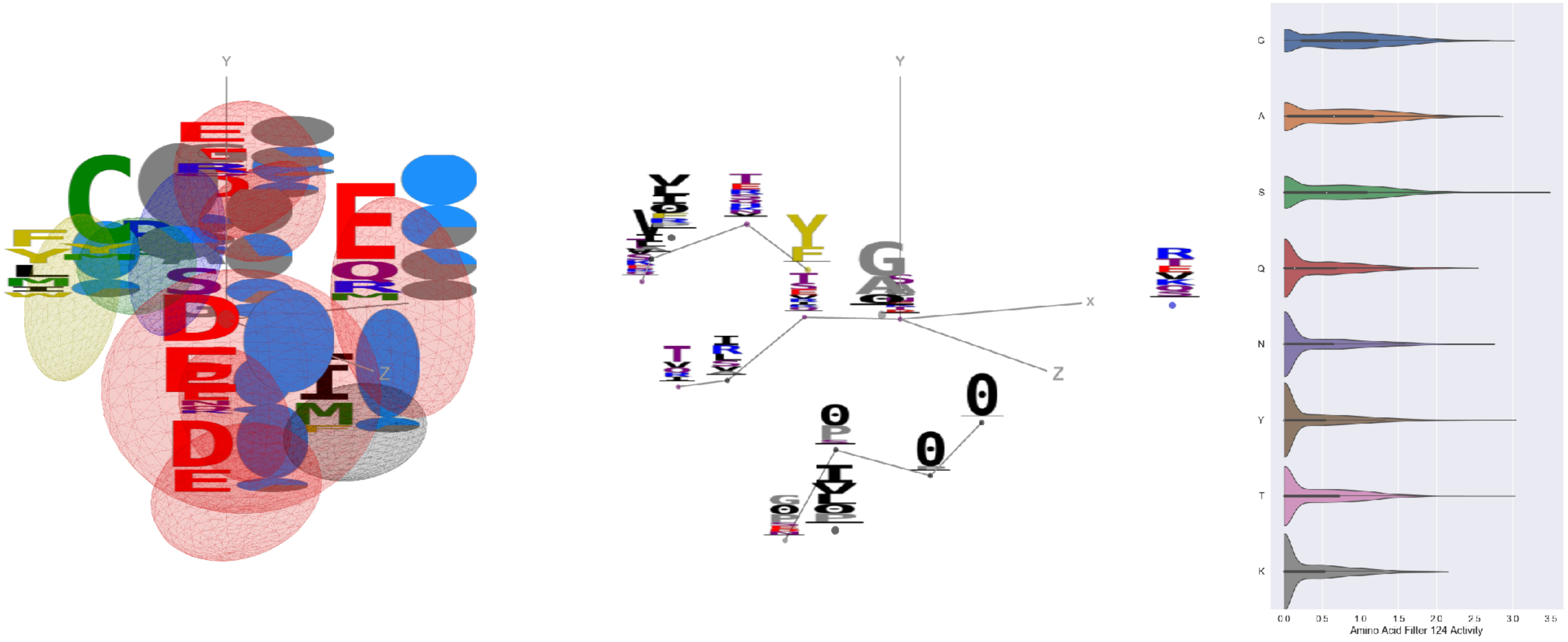

Filter 124
$R_{\text{pred}} = -0.13$
Percent Active = 48
Pooling attention 0.17
Pooling feature 0.26
Top activators
3nfp:K:143 S
3nfp:I:143 S
5t5n:C:2 Y
6mej:C:526 T
6oik:B:144 G
7lxw:A:116 S
6i07:D:107 A
4fqv:D:166 A
5i5k:B:463 S
6wpt:C:813 S

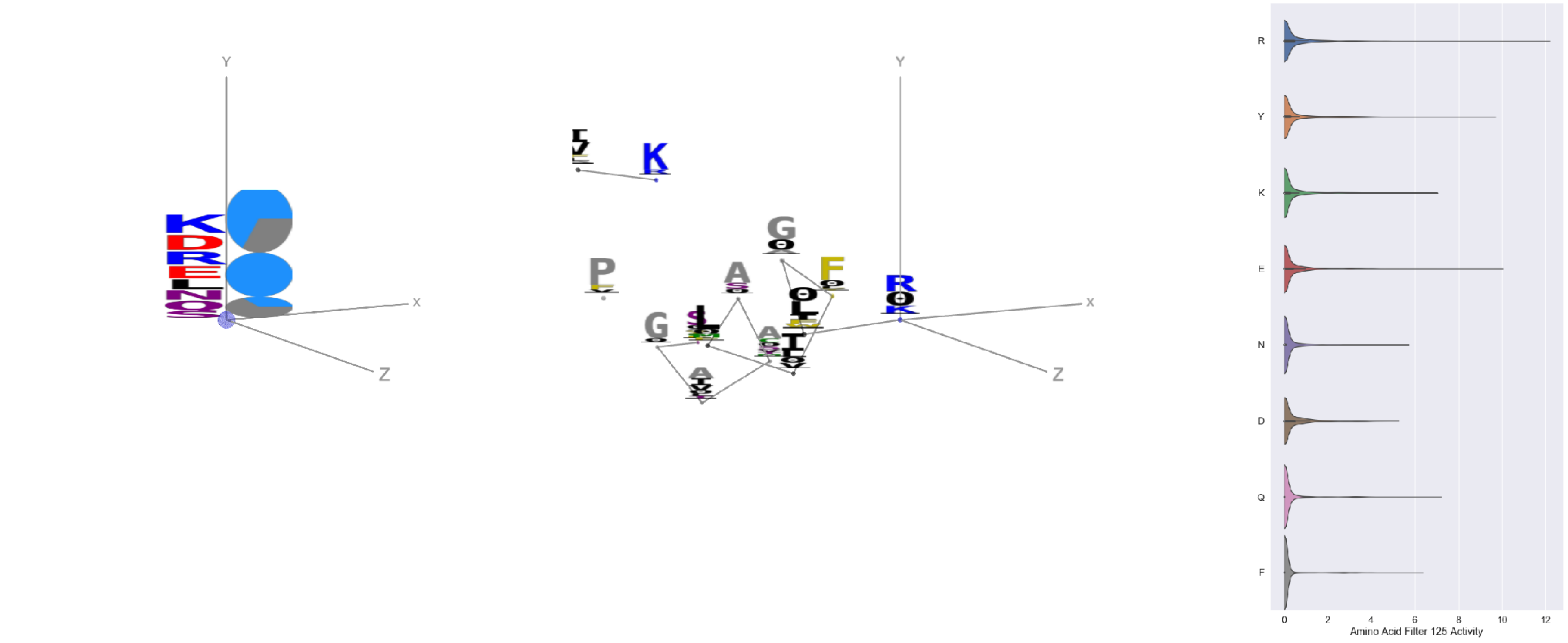

Filter 125
$R_{\text{pred}} = -0.10$
Percent Active = 23
Pooling attention 0.04
Pooling feature 0.16
Top activators
3tt3:A:11 R
4xmn:E:943 E
5otj:C:3 Y
6wtv:D:168 Y
6ii9:C:317 Y
4xmn:E:770 P
7jv6:E:854 S
4xmn:E:821 Y
4xmn:E:838 S
4xmn:E:862 G

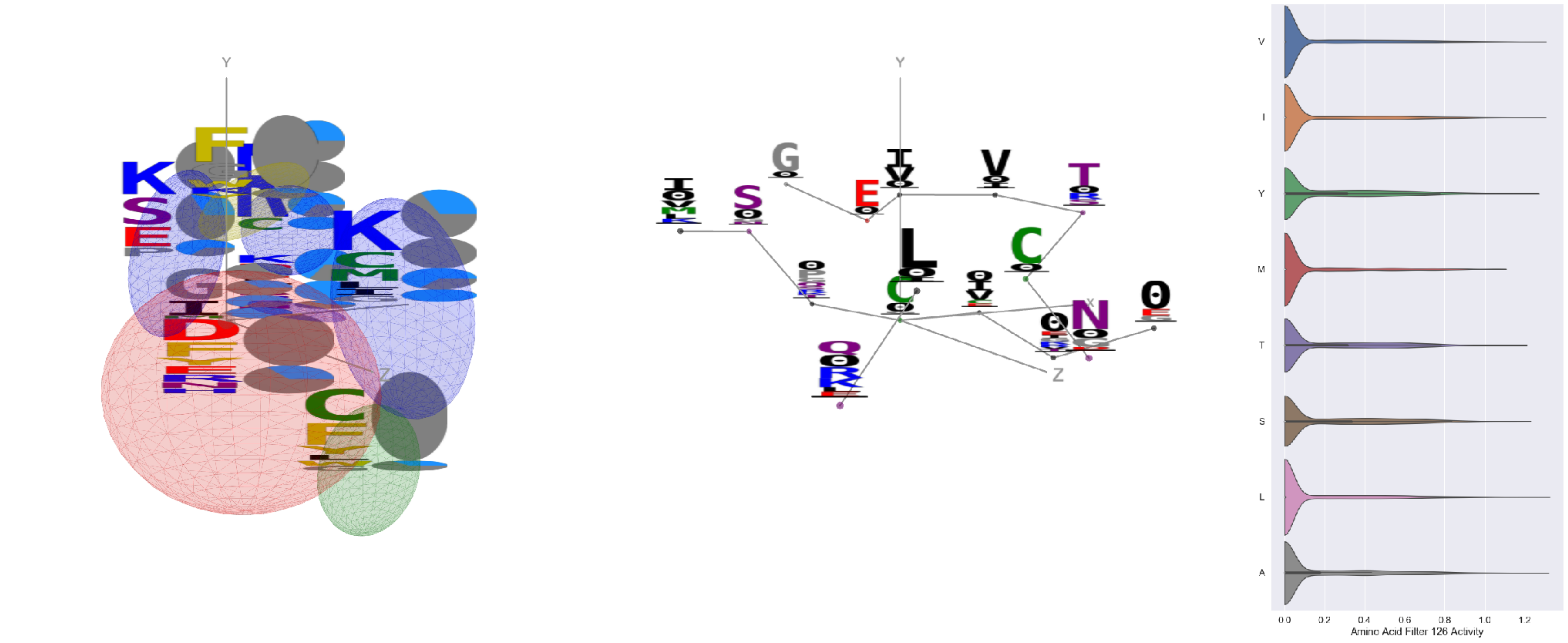

Filter 126
$R_{\text{pred}} = 0.22$
Percent Active = 32
Pooling attention 0.17
Pooling feature 1.65
Top activators
4zxb:E:8 C
5kqv:E:8 C
2qql:A:101 L
6icc:A:223 A
5kqv:E:312 C
6r0x:E:31 V
6pzw:C:263 I
4zxb:E:312 C
1ynt:G:3213 Y
5f3j:A:459 Y

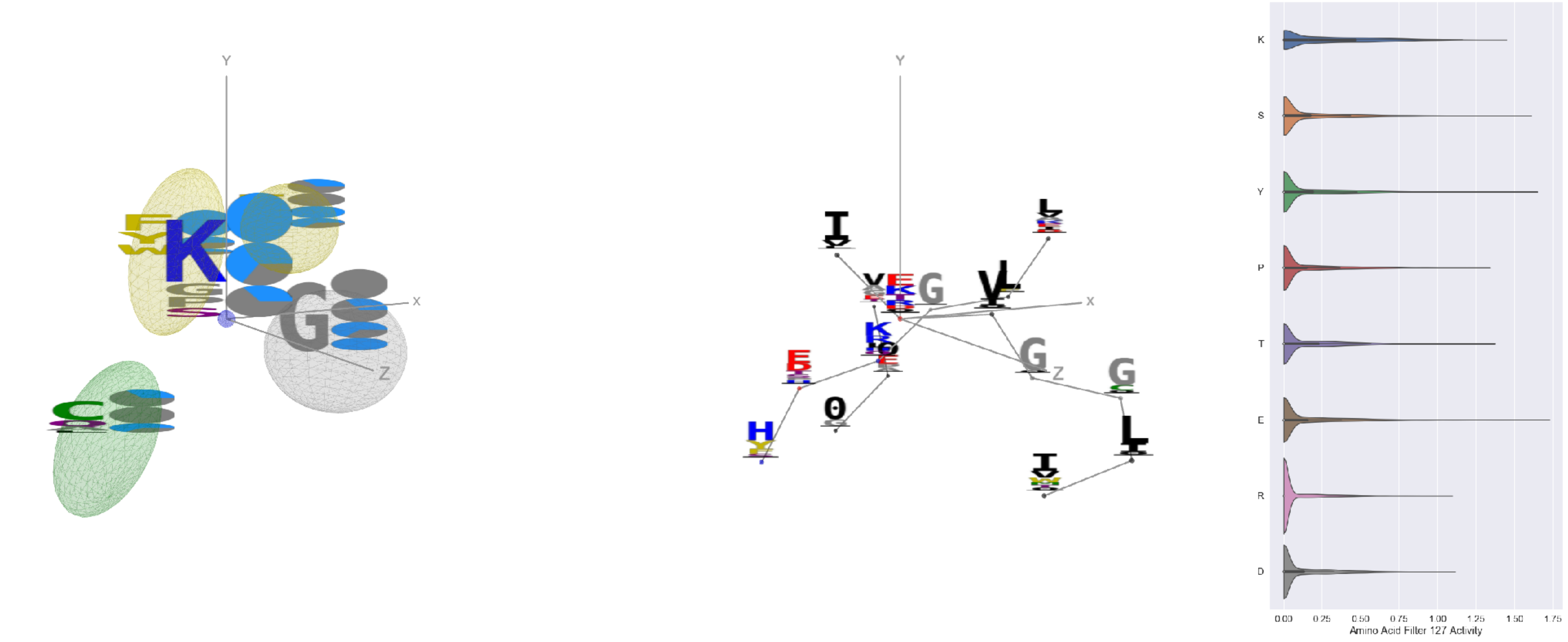

Filter 127
$R_{\text{pred}} = 0.12$
Percent Active = 24
Pooling attention 0.39
Pooling feature 1.82
Top activators
5esz:C:183 E
7nd9:B:224 Y
7cah:A:383 S
2qql:A:136 S
6o2b:M:17 S
7l57:B:224 Y
6xob:K:331 E
6ulf:K:17 S
7ce2:A:1174 K
6icf:A:277 Y
